# Enhancing Wnt signaling lowers fracture incidence in a severe mouse model of Osteogenesis Imperfecta

**DOI:** 10.1101/2025.05.27.656429

**Authors:** Giulia Montagna, Stephanie Lee, Austin Baacke, Matthew Warman, Christina Jacobsen

## Abstract

Increased risk of fracture is a hallmark feature in the skeletal disorder Osteogenesis Imperfecta (OI). One therapy in clinical trials for OI is an antibody that neutralizes sclerostin (a Wnt signaling inhibitor) that is already FDA approved for patients with osteoporosis. In addition to increasing bone mass and lowering incident fractures in patients with osteoporosis, anti-sclerostin antibody increased bone mass and skeletal strength (measured *ex vivo*) in several preclinical models of OI.

Because *in vivo* fracture incidence is the primary outcome measure in human OI trials, we examined the effect of enhancing Wnt signaling in a preclinical mouse model of moderate-severe, autosomal dominant OI (the *Col1a1*^Aga2/+^ mouse). We genetically enhanced Wnt signaling using the *Lrp5*^A214V^ allele, which makes the Wnt co-receptor LRP5 resistant to inhibition by sclerostin. By crossing *Col1a1*^Aga2/+^ sires with *Lrp5*^A214V/+^ dams, we generated pups with OI alone and OI with enhanced Wnt signaling. We radiographically examined these animals for fracture at 5, 9, and 13 weeks of age. We observed that enhanced Wnt signaling significantly reduced the number of fractures in this model by 30% at all 3 ages (e.g., 5.4 ± 1.7 fractures compared to 7.8 ± 1.8 fractures at 5-weeks-old, *p* < 0.001).

These data indicate enhancing Wnt signaling lowers incident fractures in a pre-clinical model of OI and suggest lower fracture incidence should achieved by giving anti-sclerostin antibody to patients with OI. These data also suggest that bone mass could be used as a surrogate marker for lowering fracture risk in patients with OI.

## 1. Introduction

Osteogenesis imperfecta (OI) comprises a group of Mendelian genetic disorders for which skeletal fragility is a principal feature. Most patients affected with OI (∼ 85%) have autosomal dominant mutations in 1 of the 2 genes encoding type 1 collagen (*Col1a1* or *Col1a2*) (Marini *et al*., 2017). Type 1 collagen mutations that cause haploinsufficiency, such as nonsense mutations, typically cause less severe disease than dominant negative mutations, such as missense mutations that alter glycine residues in the protein’s triple-helical domain and in-frame exon skipping mutations, both of which interfere with the assembly and folding of the heterotrimeric type 1 collagen protein (Forlino and Marini, 2016). Therapies, which in other clinical scenarios (e.g., osteoporosis), have been shown to improve bone strength and reduce fracture risk are not FDA-approved for OI; instead, these therapies are prescribed off-label, with differences of opinion within the medical community concerning their efficacy (Dwan *et al*., 2016; Vasanwala *et al*., 2016; Feehan *et al*., 2018; Garganta *et al*., 2018; Ying *et al*., 2020).

In contrast to common skeletal fragility conditions such as age-related osteoporosis, where thousands of participants have been enrolled in randomized controlled trials to demonstrate a drug’s efficacy, intervention studies for patients with OI are limited by having much smaller cohort sizes. Moreover, prospective intervention studies in OI are confounded by the spectra of disease-causing mutations, disease severity, and by the inability for small studies to adequately control for sex, age, genetic background, and environmental effects. For example, the numbers of fractures occurring in family members with the same OI-causing mutation vary widely, likely due to genetic and environmental modifiers (Daley *et al*., 2010).

Several therapies are in Phase III clinical trials for children and adults with Osteogenesis Imperfecta (https://clinicaltrials.gov - NCT05768854, NCT05972551, NCT02352753, NCT00106028, NCT03638128). In each study, fracture incidence is the FDA-mandated primary outcome measure. In preclinical models of OI, these therapies have been shown to increase bone mass and bone strength, the latter measured *ex vivo* (Uveges *et al*., 2009; Grafe *et al*., 2016; Perosky *et al*., 2016; Lee *et al*., 2022). However, the effect of these agents on incident fracture in preclinical models has not been reported. There are several reasons why fracture incidence has not been an outcome in most mouse studies. First, long-bone and vertebral fractures are uncommon in most autosomal dominant mouse OI models (e.g., Jrt, G610C, mov13); therefore, while significant improvements in bone mass and *ex vivo* bone strength have been observed using as few as 10 animals/sex/treatment group, a much larger number of animals would be needed to observe a significant reduction in incident fractures. Second, while human patients can report symptoms of nociceptive pain when a fracture occurs, mice cannot. Furthermore, signs of pain are difficult to observe in mice perhaps, because being prey animals, mice hide pain (Burkholder *et al*., 2012); consequently, incident fractures in mice may not be clinically detected in real time. Third, there is no standard radiographic method for identifying and quantifying fractures in mice.

We and others previously showed that enhancing Wnt signaling in mouse models of OI significantly improves the animals’ bone mass and bone strength (Cui *et al*., 2011; Jacobsen *et al*., 2014; Yorgan *et al*., 2015; Jacobsen *et al*., 2016). Wnt signaling enhancement is being tested in several Phase III clinical trials by using monoclonal antibodies against sclerostin, a Wnt antagonist (https://clinicaltrials.gov - NCT05768854, NCT05972551). Here we report the effect of enhancing Wnt signaling on the incidence of fracture and deformity in a mouse model of autosomal dominant OI (*Col1a1*^Aga2/+^) for which fracture and deformity are common. We also indirectly evaluate the effect of enhancing Wnt signaling on mouse nociceptive and/or nociplastic pain by quantifying animals’ movements and overall activity using a Holeboard assay.

## 2. Materials and methods

### 2.1 Animal husbandry

Two strains of mice were used to generate animals that we prospectively studied. *Col1a1*^Aga2/+^ mice, maintained on a C57BL/6J background, are heterozygous for a point mutation in the final intron of *Col1a1* (Lisse et al., 2008). This mutation creates a cryptic splice acceptor site, and aberrant splicing produces a mutant alpha1 collagen polypeptide that interferes with collagen trafficking in osteoblasts. *Lrp5*^A214V/+^ mice (JAX stock # 012669), maintained in our lab on a 129S1/SvlmJ background, are heterozygous for a missense mutation in the Wnt signaling co-receptor *Lrp5*. In humans and in mice, this mutation increases bone mass and bone strength by making LRP5 resistant to inhibition by Sclerostin (Boyden *et al*., 2002; Little *et al*., 2002; Cui *et al*., 2011). *Col1a1*^Aga2/+^ sires were mated with *Lrp5*^A214V/+^ dams to produce male and female offspring with 4 different genotypes (*Col1a1*^+/+^;*Lrp*5^+/+^, *Col1a1*^Aga2/+^;*Lrp*5^+/+^, *Col1a1*^Aga2/+^;*Lrp*5^+/A214V^, and *Col1a1*^+/+^;*Lrp*5^+/A214V^, henceforth simplified as being WT, Aga2, Aga2/A214V, and A214V, respectively. Offspring were genotyped by tail snip and given ear tags for identification at ∼10 days of age, weaned by 28 days and age, and upon weaning group housed with same sex littermates using standard housing, animal chow, and 12-hour day/night cycles. Cage cards alerted animal care, who performed bi-weekly cage changes to handle the animals gently due to the risk of skeletal injury.

### 2.2 Animal genotyping

Tail snips were collected from ∼10 days old mice and DNA was extracted using the “HotSHOT protocol” (Truett *et al*., 2000) previously developed in-house. Briefly, we digested the tissues in 50 µl of 25 mM NaOH at 95°C for 30 min and inactivated the digestion by adding 50 µl of 40 mM Tris-HCl. We used 2 µl of this concoction as PCR template for the following reactions. The genotyping of *Lrp5*^A214V/+^ mice was performed by PCR using forward 5’ – AGTACTGGCTGGCACAGA – 3’ and reverse primer 5’ – CAGGCTGCCCTTGCAGAT – 3’. The genotyping of *Col1a1*^Aga2/+^ mice was performed by digital droplet PCR (ddPCR) using forward 5’ – GGCAACAGTCGCGCTTCACCTA – 3’ and reverse primer 5’ – GGAGGTCTTGGTGGTTTTGT – 3’ in combination with two different probes: WT-Probe: 5’ –/5HEX/ACCCTCTCCCGCTGTCTTCATTC – 3’ and Aga2-Probe: 5’ – /56-FAM/ACCCTCTCCCGCAGTCTTCATT – 3’. All primers and probes were ordered through Integrated DNA Technologies (IDT).

### 2.3 Animal phenotyping

Other than genotyping, weaning, and cage changing, animals were left undisturbed except at 5 weeks of age, 9 weeks of age, and 13 weeks of age. At these 3 ages, mice were transported to the Animal Behavior and Physiology (AB&P) Core (Riccio *et al*., 2014) at Boston Children’s Hospital to allow for initial acclimation to a new environment. Then individual mice were tested in a Holeboard apparatus to measure their total movement, nose poking, and hindleg rearing, using a camera, infrared sensor, and automated software. The animals were then anesthetized using isoflurane, placed in a prone position, and radiographically imaged using a Faxitron UltraFocus100 scanner (Faxitron Bioptics, LLC, Tucson, AZ, USA). Study animals were euthanized at 13 weeks, prior to the last radiograph.

### 2.4 Radiographic scoring

Investigators reviewed radiographs while blinded to the animals’ genotypes. Radiographs were examined at 7 bilateral skeletal sites (humerus, olecranon, forearm, ischium, femur, tibia, calcaneus) for fracture and deformity. One point was scored for every fracture or deformity at each skeletal element. The presence of scoliosis was scored as Yes or No. The ratio between the distance between the medial surfaces of an animal’s femoral heads and the distance between the animal’s ischial tuberosities was also determined. All radiographs obtained from animals of the same age were scored without looking at radiographs of the same animal taken at the other 2 ages.

### 2.5 Statistical analyses

Primary outcome measures were predetermined before data were unblinded for animal genotype. The primary outcome measure was total number of fractures and deformities by radiograph of Aga2 and Aga2/A214V animals at each time point. Differences in the incidence of scoliosis and in the interfemoral/interischial ratios (IIR) were secondary outcomes. Before unblinding the data, we confirmed that the interobserver correlation coefficients for each outcome measure was > 0.7. We had no *a priori* knowledge regarding expected means and standard deviations for total movement, nose poking, and hindleg rearing in the Holeboard assay. Therefore, differences between Aga2 and Aga2/A214V animals for these the activity measures were considered to be secondary outcomes. Male and female data were initially considered separately. When no sex-dependent differences were observed, the data were pooled. Student’s t-tests and Mann-Whitney U tests were employed to compare the total number of fractures and deformities between Aga and Aga2/A214V mice. IIR differences were assessed with the Mann-Whitney U test, while Student’s t-tests were used to evaluate total movement, nose poking, and hindleg rearing between Aga2 and WT mice and between Aga2 and Aga2/A214V mice. Statistical assessment of scoliosis incidence between Aga2 and Aga2/A214V mice using the two-proportion z-test.

### 2.6 RNA extraction from femurs

WT and Aga2 males were euthanized by CO_2_ inhalation and femurs were dissected. Only fracture-free femurs were processed. Femur epiphyses were discarded and bone marrow was removed by centrifugation. Each diaphysis was stored in -80 in a screw-cap tube with 1 stainless steel metal bead (Qiagen, #69989) per tube. For RNA extraction, 500 µl of TRIzol (Invitrogen, #15596026) and 100 µl of nucleases-free PBS were added to each tube. Homogenization was performed using a Qiagen Tissue Lyser III (Qiagen, #9003240) for 2 rounds of 2 minutes, at 30 Hz of frequency. Samples were allowed to stand for 5 minutes at RT to promote complete dissociation of the nucleoprotein complex. 200 µl of chloroform per tube were added before shaking vigorously and centrifuging the vials for 15 minutes at 12,000 x g and 4°C. The top aqueous phase of the density gradient was collected and mixed with 2 volumes of binding RLT buffer provided by the RNeasy Mini kit (Qiagen, #74104). Following addition of 1 volume of 70% ethanol, the samples were loaded onto the RNA binding columns provided by the kit. In-column genomic DNA digestion was performed using DNase I and the buffer provided in the PureLink kit (PureLink, #12185-010). 80 µl of mix were added per tube and left to incubate for 15 minutes at RT. From this point on we followed the manufacturer guidelines provided in the RNeasy Mini kit. Each sample was eluted in 30 µl of nuclease-free water and quantified using a nanodrop spectrophotometer (DeNovix, DS-11 FX+). RNA quality check was performed using Agilent 2100 Bioanalyzer and RNA 6000 Pico kit (Agilent, #5067-1513).

### 2.7 Aga2 mRNA quantification via ddPCR

RNA was extracted as previously described from the femurs of 5-week-old mice (3 females and 3 males). RNA was then reverse transcribed into cDNA using SuperScript III First-Strand Synthesis System (Invitrogen 18080051) following the manufacturer’s instructions. For reverse transcription, oligo dT was used to achieve better coverage of transcripts’ terminal 3’ ends, where the Aga2 mutation is localized. cDNA was amplified via digital droplet PCR, either using primers specific for the Aga2 mutation (P1_For: 5’-CGATGGATTCCCGTTCGAGT-3’; Aga2_Rev: 5’-TCTGGTGTGAATGAAGACGT-3’) or using primers to amplify all *Col1a1* transcripts (WT_For: 5’-GATGCTAACGTGGTTCGTGA-3’; WT_Rev: 5’-TCCGCTCTTCCAGTCAGAGT-3’). ddPCR master mix was prepared using 12.5 µl EvaGreen Supermix 2x (BioRad, # 1864034), 0.25 µl of each primer (working concentration of 10 µM), 1 ng of cDNA template and water up to 25 µl. 20 µl from each reaction were manually mixed with 70 µl of QX200 Droplet Generation Oil for EvaGreen (BioRad, #1864005) in a DG8 Cartridge (BioRad, #) and droplets were generated using a QX200 Droplet Generator. The reaction droplets were transferred into a dedicated PCR plate and the following PCR protocol was run. After the PCR, the plate was loaded in the QX200 Droplet Reader (BioRad, # 1864003) and the fluorescence of each droplet was measured. The QuantaSoft Software automatically calculate a threshold value and counts the droplets with fluorescence higher than threshold as positive events. Positive droplets for were plotted as average and standard deviation and average difference was inferred using a non-parametric Wilcoxon test.

### 2.8 RNA-sequencing

Individual femurs dissected from 3 Aga2 and 3 WT 10-month-old male mice were used for RNA-sequencing. Genomic DNA contamination was checked using PCR and primers pairing to exon 22-24 of *Col1a1*. We shipped 500 ng of RNA to Azenta laboratories for library preparation and sequencing using 30,000,000 reads (paired-end) per sample. Trimming, deduping, and mapping sequencing reads were performed by Azenta. Reads were visualized using Integrative Genomic Viewer (IGV). RNA sequencing data have been deposited in the publicly available Gene Expression Omnibus data repository (reference number).

## 3. Results

### 3.1 Aga2/A214V mice have fewer fractures and deformities and less scoliosis than Aga2 mice

None of the WT or A214V mice had scoliosis, fractures, or deformities. Among mice that were either Aga2 or Aga2/A214V, the interobserver correlation coefficient (**Supplementary Table 2, tab 1**) for the total number of fractures and deformities was 0.8. **Table 1** provides means and standard deviations for the total number of fractures and deformities observed in Aga2 and Aga2/A214V males and females at 5, 9, and 13 weeks of age, and scoring for all animals is available in **Supplementary Table 1**. **Figure 1** shows representative radiographs from a 5-week-old Aga2 mouse and a 5-week-old WT mouse, and radiographs from every animal and age are available upon request. Fracture and deformity number were not influenced by the animals’ sex; hence male and female data were pooled when determining the effect of the A214V allele. Aga2 mice averaged 7.8 (SD 1.7) fractures and deformities at 5 weeks of age, and this number remained stable to 13 weeks of age. Aga2/A214V mice averaged 5.4 (SD 1.7) fractures and deformities at 5 weeks of age, which also remained stable until 13 weeks of age. At each age the presence of the A214V allele significantly reduced the total number of fractures and deformities in Aga2 mice as determined by T-test and Mann-Whitney test (*p* < 0.001 at all time points).

**Figure 1:**
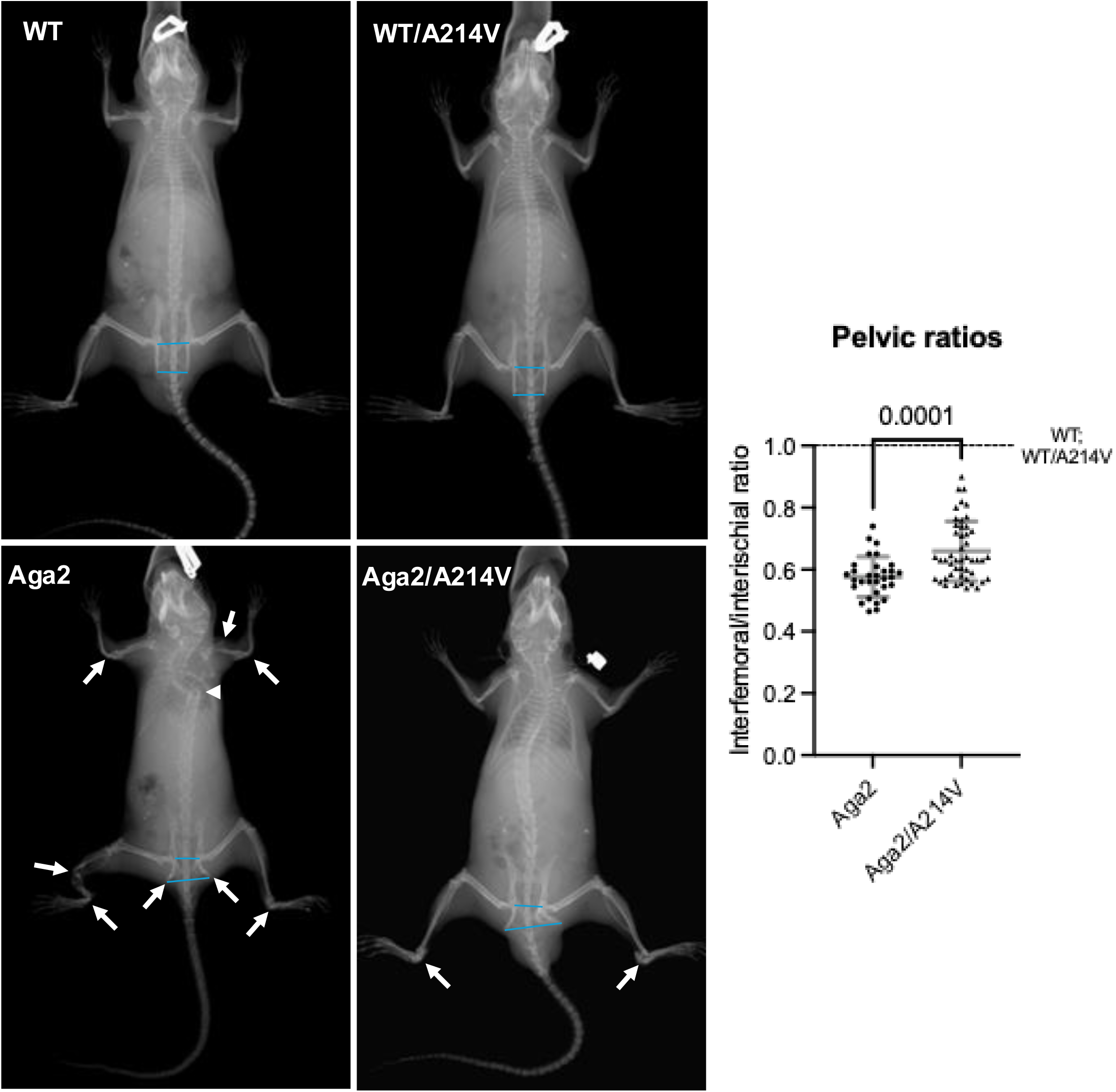
Representative radiographs, scoring method, and pelvic ratios. Representative average radiographs of WT, WT/A214V, Aga2, and Aga2/A214V mice. White arrows indicate deformities counted during scoring. Arrowhead indicates thoracic scoliosis. Blue lines are used to calculate the interfemoral/interischial ratio that is assumed to be 1 for WT and averages around 0.63 for Aga2 and Aga2/A214V. On the right, the interfemoral/interischial ratio (pelvic ratio) data were plotted for Aga2, and Aga2/A214V mice. Dotted line indicates the constant value of 1, which corresponds to interfemoral/interischial ratio average for WT, WT/A214V.

**Table 1:** Average and standard deviations of total number of fractures and deformities observed in Aga2 and Aga2/A214V males and females at 5, 9, and 13 weeks of age. Asterisks indicate a reduction in the number of individuals considered, due to animal death. Specifically, in the Aga2 female cohort we had 8 individuals at 9 weeks and 7 at 13 weeks, while in the Aga2 male cohort 10 individuals were considered at 9 and 13 weeks. No number reduction was recorded in Aga2/A214V cohorts.

|  | Aga2 |  | Aga2/A214V |  | Aga2 | Aga2/<br>A214V | T-test | Mann-Whitney<br>test |
| --- | --- | --- | --- | --- | --- | --- | --- | --- |
|  | Females<br>(n=9) | Males<br>(n=11) | Females<br>(n=16) | Males<br>(n=10) |  |  |  |  |
| 5 weeks | 7.8±2.0 | 7.9±1.4 | 5.9±1.9 | 4.8±1.2 | 7.8±1.7 | 5.4±1.7 | 0.00002 | <0.0001 |
| 9 weeks | 7.7±2.0* | 8.1±1.6* | 6.0±2.1 | 5.1±1.4 | 7.9±1.8 | 5.6±1.9 | 0.00017 | 0.0001 |
| 13 weeks | 7.8±2.2* | 8.0±1.6* | 6.1±2.0 | 5.1±1.7 | 7.9±1.8 | 5.7±1.9 | 0.00054 | 0.0004 |

None of the WT or A214V mice had scoliosis. Among mice that were either Aga2 or Aga2/A214V, the interobserver agreement (https://www.statology.org/fleiss-kappa-excel/) among radiographic scorers for the presence of scoliosis using Fleiss’ Kappa statistics (K) was 0.9, indicating very good agreement between the scorers (**Supplementary Table 2, tab 2**). In mice with the Aga2 allele, the A214V allele significantly reduced the incidence of scoliosis at 5 weeks of age, with 6 of 26 Aga2/A214V mice having scoliosis compared to 12 of 20 Aga2 mice. No difference in scoliosis incidence was seen between females and males with the same genotype.

The IIR was ∼1 for WT and A214V mice. Among mice that were either Aga2 or Aga2/A214V, the interobserver correlation coefficient for IIR was 0.83 (**Supplementary Table 2, tab 3**). The data were not normally distributed and, according to the non-parametric Mann-Whitney test, the IIR did not change significantly with sex or age among the two genotypes. Therefore, we disregarded the variables sex and age and pooled all Aga2 or Aga2/A214V data together. The IIR for Aga2 mice was 0.58 (SD 0.09), while that for Aga2/A214V was 0.66 (SD 0.12). The difference between Aga2 or Aga2/A214V was statistically significant according to the non-parametric Mann-Whitney test (*p* = 0.0001; **Figure 1**).

The number of Aga2 but not the Aga2/AV animals decreased during the 13-week-long experiment. Two Aga2 animals experienced acute long bone fractures during the Holeboard assay and had to be euthanized.

### 3.2 Holeboard assays revealed significant differences in total movement, nose poking, and hindleg rearing between WT and Aga2 mice, but not between Aga2 and Aga2/A214V mice

Each Holeboard assay was performed for one hour and the operators were blinded to mouse genotype. Data were compared across the 4 genotypes (i.e., WT, A214V, Aga2, and Aga2/A214V) at 5, 9, and 13 weeks of age. Means and standard deviations for total movement, hindlimb rearing, and nose poking are plotted in **Figure 2**. Total movement is the total distance the mouse traveled (**Figure 2A**); hindlimb rearing is the total number of times the mouse (**Figure 2B**); nose poking is the total number of times a mouse explored the 9 holes that are the board (**Figure 2C**). Holeboard data from every animal and age are available in **Supplementary Table 3**. Total movement, hindleg rearing, and nose poking were not influenced by animal sex, therefore the two sexes were pooled together for statistical analysis and plotting. Additionally, rearing did not vary with mouse age, so data from the 3 time points were pooled when determining if Aga2 mice differed from WT and whether Aga2 mice differed from Aga2/A214V (**Figure 2B**). Total movement and rearing significantly differed between WT and either Aga2 or Aga2/A214V animals (**Figure 2A and 2B**). However, differences between Aga2 and Aga2/A214V mice did not reach statistical significance, although Aga2 mice with the A214V allele trended toward higher for total movement, rearing, and nose poking.

**Figure 2:**
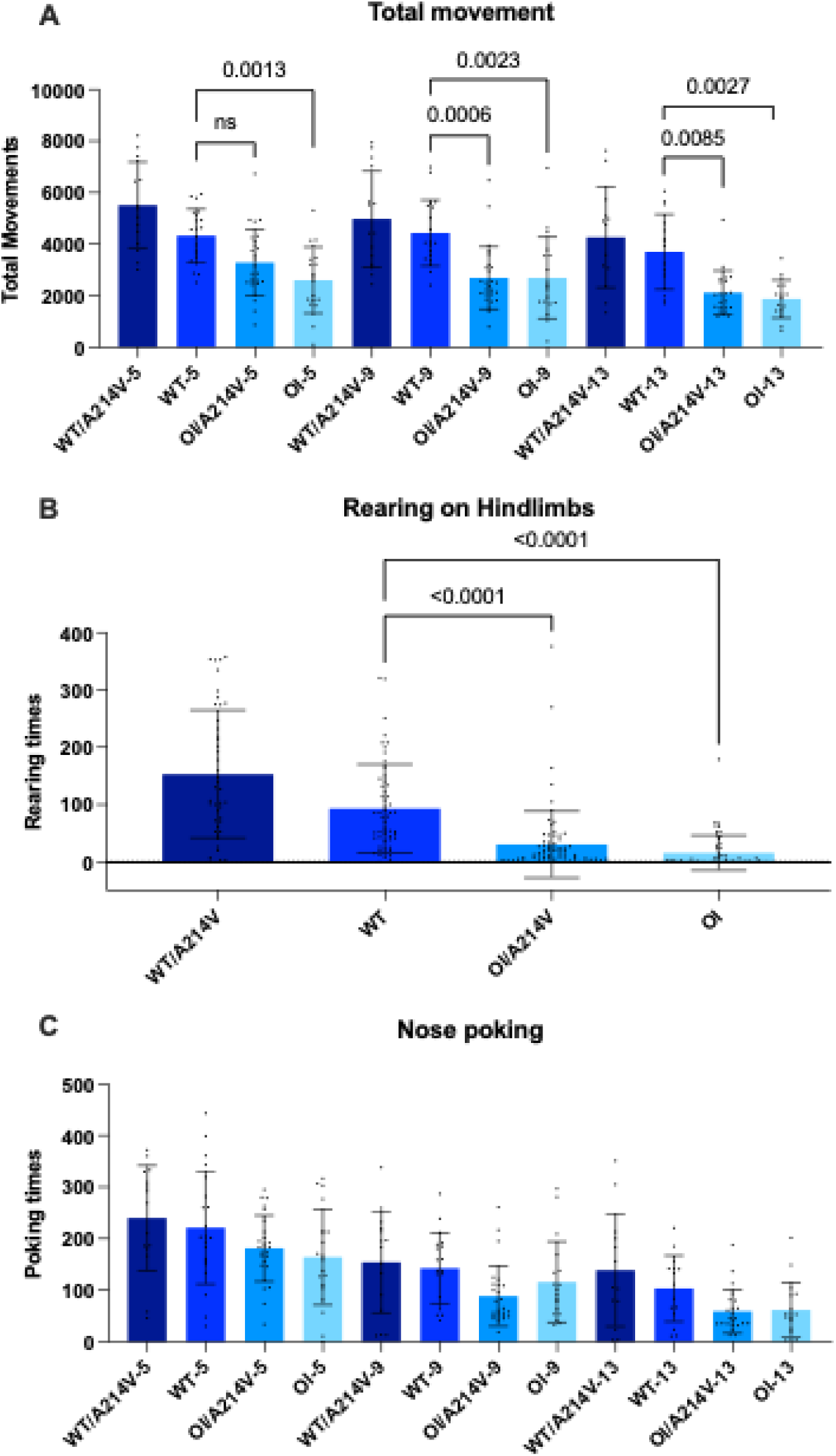
Basic movement, rearing, and poking of Aga2, Aga2/A214V, WT, and WT/A214V were measured using a Holeboard assay. We plotted average and standard deviation per measurement. *p*-values are indicated in the graphs. A. Basic movement was measured in mice with 4 different genotypes (WT, WT/A214V, OI, OI/A214V) at 5, 9, and 13 weeks of age. Labels on the X-axis indicate the genotype and the age. On the Y axis is the distance covered in average by the mice expressed in centimeters. B. Number of times the mice reared on their hindlimbs during 60 min. Data from different times points were pooled together because we did not see significant variation among them. C. Number of times the mice poked their nose into the 9 wholes of the board (considering all 9 wholes together).

### 3.3 Twenty-two percent of bone *Col1a1* transcripts use the cryptic splice site created by the Aga2 mutation

Droplet digital PCR estimated that 22 percent of *Col1a1* transcripts in bone utilize the cryptic splice (Figure S1). A similar percentage of abnormal splicing was observed using RNA sequencing (Figure S2).

## 4. Discussion

The data presented herein indicate that genetic enhancement of Wnt signaling reduces the incidence of fracture and deformity in the Aga2 mouse model of moderate-severe autosomal dominant OI (type 3 OI in humans). The *Lrp5*^A214V^ allele increases bone mass by making LRP5 resistant to inhibition by Sclerostin (Yorgan *et al*., 2015). Humans with equivalent LRP5 missense mutations have high bone mass, as do humans and mice that genetically lack Sclerostin (Brunkow *et al*., 2001; Van Wesenbeeck *et al*., 2003). These observations led to the development of monoclonal antibodies that neutralize Sclerostin as a therapy for disorders of low bone mass, including osteoporosis (Roschger *et al*., 2014; Cardinal *et al*., 2019). For osteoporosis, bone mineral density (BMD) at the spine and at the femoral head correlates with fracture susceptibility. Hence, increased BMD has become a surrogate outcome measure for therapeutic efficacy. However, mutations in type 1 collagen that cause OI affect bone material properties in addition to bone mass. Thus, in the context of abnormal material properties, it was unknown if BMD remains a clinically relevant outcome measure. Therefore, clinical trials in patients with OI have reduction in fracture incidence as the primary outcome. Our fracture data inversely correlate with BMD data for Aga2 and Aga2/A214 mice. Jacobsen et al. (in preparation) found that Aga2/A214V mice have significantly higher trabecular BV/TV than Aga2 mice at 12 weeks of age (0.0.44 vs 0.190, p < 0.0001). Therefore the fracture reductions we observed in Aga2/A214V mice compared to Aga2 mice support the use BMD as a surrogate outcome measure for OI too.

At the study’s onset, our primary outcome measure was the summed number of fractures and deformities at 7 skeletal locations. This measure revealed significant improvements associated with the A214V allele in Aga2 mice. Post-hoc analysis suggests an outcome measure using 4 skeletal locations (humerus, forearm, femur, foreleg) may be better. This is because nearly every animal with an Aga2 allele, independent of the A214V allele, had calcaneal and ischial fracture or deformity. Thus, enhancing Wnt signaling did not protect all skeletal sites in Aga2 mice; however, the incidence of long bone fractures and deformities decreased by 50% in Aga2/A214V mice. Additional studies using Aga2 and similar OI models are needed to confirm if 4 sites are better at detecting significant improvements than the 7 sites we used in this study.

In addition to observing a significant reduction in fracture and deformity, the Aga2/A214V mice also had significantly less scoliosis. We do not know the reason for this improvement. Potential explanations include beneficial LRP5-mediated effects on vertebral bodies, muscle tone and/or strength, and improved type 1 collagen production in ligament, tendon, and/or intervertebral disc. Future work is needed to delineate the responsible mechanism. We also do not know the mechanism by which BMD is increased and fracture incidence is reduced. Possible mechanisms that need to be examined in future studies include changes in the percentage of abnormally spliced transcripts, changes in the sensitivity of osteoblasts to ER stress and apoptosis, or changes in the numbers of osteoblasts that are synthesizing or osteoclasts that are degrading bone.

We speculated that reducing the number of fractures and deformities in mice would produce benefits outside of their skeletal system, such as improved mobility, reduced pain, and increased overall well-being. We attempted to address this hypothesis utilizing the Holeboard assay.

We assumed total movement and hindleg rearing are surrogate measures for acute and chronic pain, and nose poking (a measure of curiosity and exploratory behavior) is a surrogate indicator of well-being (Roughan *et al*., 2009). Aga2 mice performed significantly worse for all three outcome measures compared to WT mice. The A214V allele did not improve performance in Aga2 mice for total mobility at any time point. Given the high prevalence of calcaneal fractures in Aga2 and Aga2/A214V mice, it is not surprising that hindleg rearing was also unchanged. Nose poking had a large SD relative to its mean, so our study was underpowered to detect even a moderate improvement. Because Aga2/A214V animals had an average of 5.4 fractures when the first Holeboard assay was performed, we are unable to determine if decreased mobility is associated with pain rather than impaired mechanics. Many patients with OI experience chronic pain in addition to acute pain associated with fracture (Nghiem *et al*., 2017; Munoz Cortes *et al*., 2022). Studies to determine if bone pain reduces mobility before fractures appear, or if pain originating in other collagen rich tissues (e.g., tendon and ligament) independently affect mobility, will require animals with conditional OI-causing alleles that be activated in a temporal and/or tissue-specific manner.

Other studies have used fracture and deformity as outcome measures in mouse models of OI. Most studies were performed using the autosomal recessive Oim mouse which is homozygous for a mutation in *Col1a2* (Cardinal *et al*., 2019; Cardinal *et al*., 2020; Sun *et al*., 2023). Fracture scoring varied among investigative teams but tended to focus on long bones. For example, Cardinal et al. (2019), defined fractures as “evidence of solution of continuity, callus formation, and patent bone deformity” and counted them in femurs, tibias, humeri, and forearms to assess anti-sclerostin antibody (Scl-Ab) potential to decrease fractures incidence. Sun and colleagues (2023), counted fractures defined as “any evidence of interruption of continuity, bone deformity, or callus formation” in humeri, radii, femurs, tibias, and tail bones to measure Irisin efficacy in fracture incidence reduction. Only one other study used Aga2 mice (Duran *et al*., 2022). This study examined the effect of oral 4-phenylbutyrate, an ER stress reducer, on bone properties including fracture. The investigators reported that 4-phenylbutyrate, added to animals’ drinking water at weaning, reduced fracture and deformity compared to controls. Their scoring system differed from ours, in that they counted fractures of long bone diaphyses (humeri, radii, ulnas, femurs, tibias, and fibulas) but not pelvic, olecranon, or calcaneal fractures.

Our research goal is to benefit patients. Results from this study in Aga2 mice contribute to this goal by showing enhancing Wnt signaling reduces fracture and deformity. Thus, similar effects should be expected in patients with OI. The mouse data also point to challenges for human clinical trials. The mice in our study were all the same age, on a fixed genetic background, and in controlled environments. Aga2 animals averaged 7.8 fractures/deformities by 5 weeks of age, with the SD being ∼1/5 of the mean. This made it possible to detect significant therapy-associated improvement using few animals. In clinical trials involving patients with OI, the SDs for incident fractures and the annualized rates at which new fractures appear are likely to be larger fractions of the means. Furthermore, fractures in patients with milder forms of OI occur infrequently; an individual who has two fractures within 1 calendar year can have no fractures for the next several years. Therefore, individual participants cannot serve as their own controls. Clinical trials whose inclusion criteria require recent fracture and trials that cannot be well-controlled for age, phenotypic severity, genetic background, and environment may have significant difficulties showing therapeutic efficacy unless they can enroll large numbers of participants or the beneficial effect is profound. Indeed, these very factors have played a role in the current pediatric clinical trials for new therapeutics in OI.

Our study has limitations. First, we report fracture data for only 1 mouse model of OI. Second, this autosomal dominant model has an atypical mechanism of mutational effect. The mutation creates a cryptic splice site that affects ∼ 20% of *Col1a1* transcripts in bone. Translation of these abnormal transcripts produces a polypeptide that interferes with collagen trimer assembly, leading to ER stress and to osteoblast apoptosis rather than to the secretion of mutant collagen into the extracellular matrix. Thus, our study does not address whether mice with mutant extracellular matrix collagen will exhibit a similar reduction in fracture incidence. However, we and others have examined the effect of enhanced Wnt signaling in mouse OI models (G610C and Jrt) where mutant collagen is secreted, and we observed significant improvement in their bone mass and *ex vivo* bone strength (Masci *et al*., 2016; Marulanda *et al*., 2023). Unfortunately, G610C and Jrt mice fractured infrequently and the studies were underpowered to detect reductions in fracture incidence. Third, we used a genetic approach to enhance Wnt signaling instead administering a therapeutic agent, such as Sclerostin neutralizing antibody. The genetic approach minimizes animal handling and reduces the chance of inducing iatrogenic fractures. Therefore, the genetic approach provides proof of principle results. Fourth, our genetic approach exposed Aga2/A214V mice to enhanced Wnt signaling *in utero*, whereas neutralizing antibodies would be administered postnatally. Although the former could have been more effective than the latter, it may not be. Neutralizing antibodies enhance Wnt signaling through LRP5 and LRP6 (Li *et al*., 2005), whereas the A214V allele only enhances Wnt signaling via LRP5.

To conclude, using the Aga2 mouse model of OI we show that enhancing Wnt signaling significantly reduces fracture and deformity. This finding is relevant to ongoing clinical trials that are testing the effectiveness of enhancing Wnt signaling in patients with OI.

## Supporting information

SupTable1

SupTable2-tab1

SupTable2-tab2

SupTable2-tab3

SupTable3

## Author contributions and acknowledgements

CMJ and MLW designed the experiments, SL, AB, and GM performed the experiments, GM and MLW wrote the initial draft of the paper, which was reviewed, revised, and approved by all authors. The authors thank the staff in the Animal Behavior and Physiology at BCH and dedicate this paper to staff member Mr. Kamil Moroz, who passed away while this study was being performed. We also thank research assistants Isabelle Hartshorn and Jasmine Nutakki, and summer students Iman Boulouah, Kefan Cui, Beverly Duclas, Islam Elouadi, Darryl Nguyen, Lisa Nguyen, and Rebecca Sannon for helping develop and validate the fracture and deformity scoring system. This work was supported by NIH grants R21AR077292, R21AR082609, P30AR075042, a Michael Geisman Fellowship from the Osteogenesis Imperfecta Foundation. Summer students were supported by Project Success at Harvard Medical School and COACH at Boston Children’s Hospital.

## Funding

NIH R21 to MW and CMJ, OIF Michael Geisman fellowship to GM, P30 for RNA-seq data, P30, COACH, and Project Success for summer students.

## Supplementary material captions

**Figure S1:**
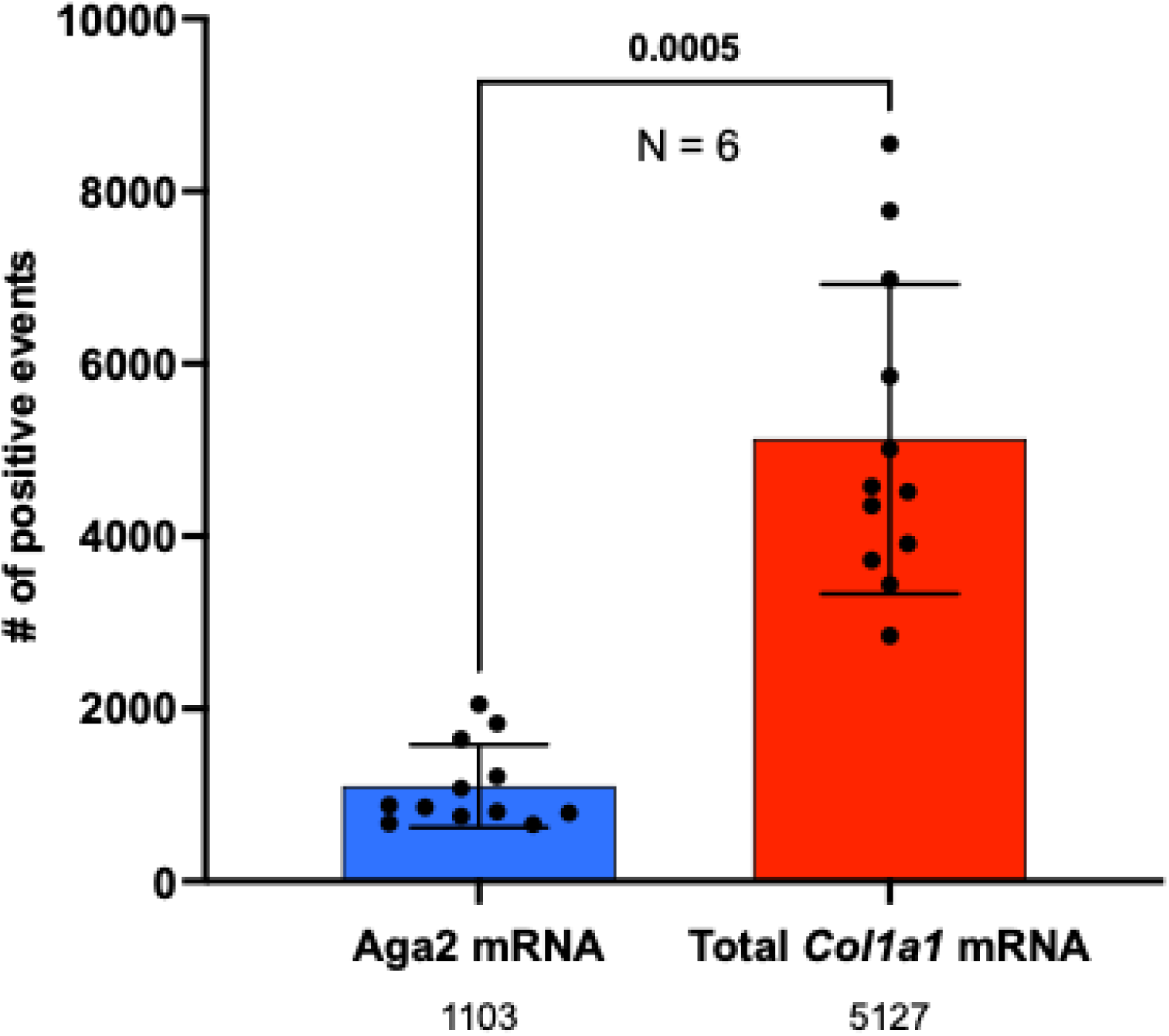
ddPCR quantification of the Aga2 and the WT *Col1a1* mRNAs. The red bar indicates the number of oil droplets in which a molecule of cDNA containing the Aga2 mutation has been amplified. The blue bar shows the number of oil droplets in which a molecule of cDNA containing WT *Col1a1* sequence has been amplified. Bars indicate average numbers and error bars standard deviation. Statistical inference has been calculated using the non-parametric Wilcoxon test that results significant (*p =* 0.0005). The average numbers of events were made explicit below the bars, showing that the Aga2 transcript represents 21.5% of the total number of transcripts detected.

**Figure S2:**
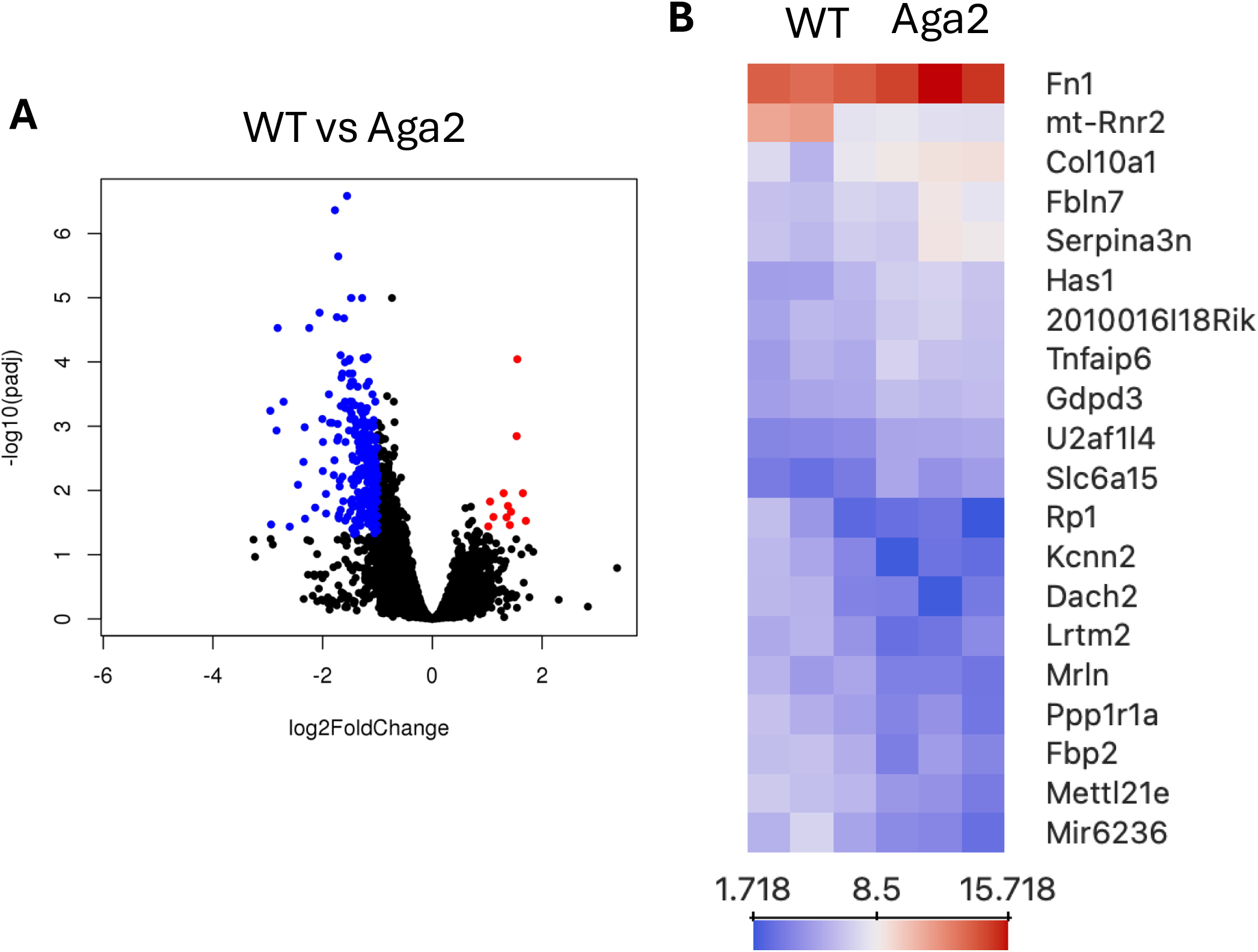
Volcano plot and heatmap representing the top differentially expressed genes between Aga2 and WT femurs. A. Volcano plot showing significantly downregulated, in blue, and upregulated, in red, transcripts in Aga2 femurs with respect to WT. B. Heatmap showing log2 of expression values for the top 20 differentially expressed genes, excluding predicted genes (Gm#####).

**Figure S3:**
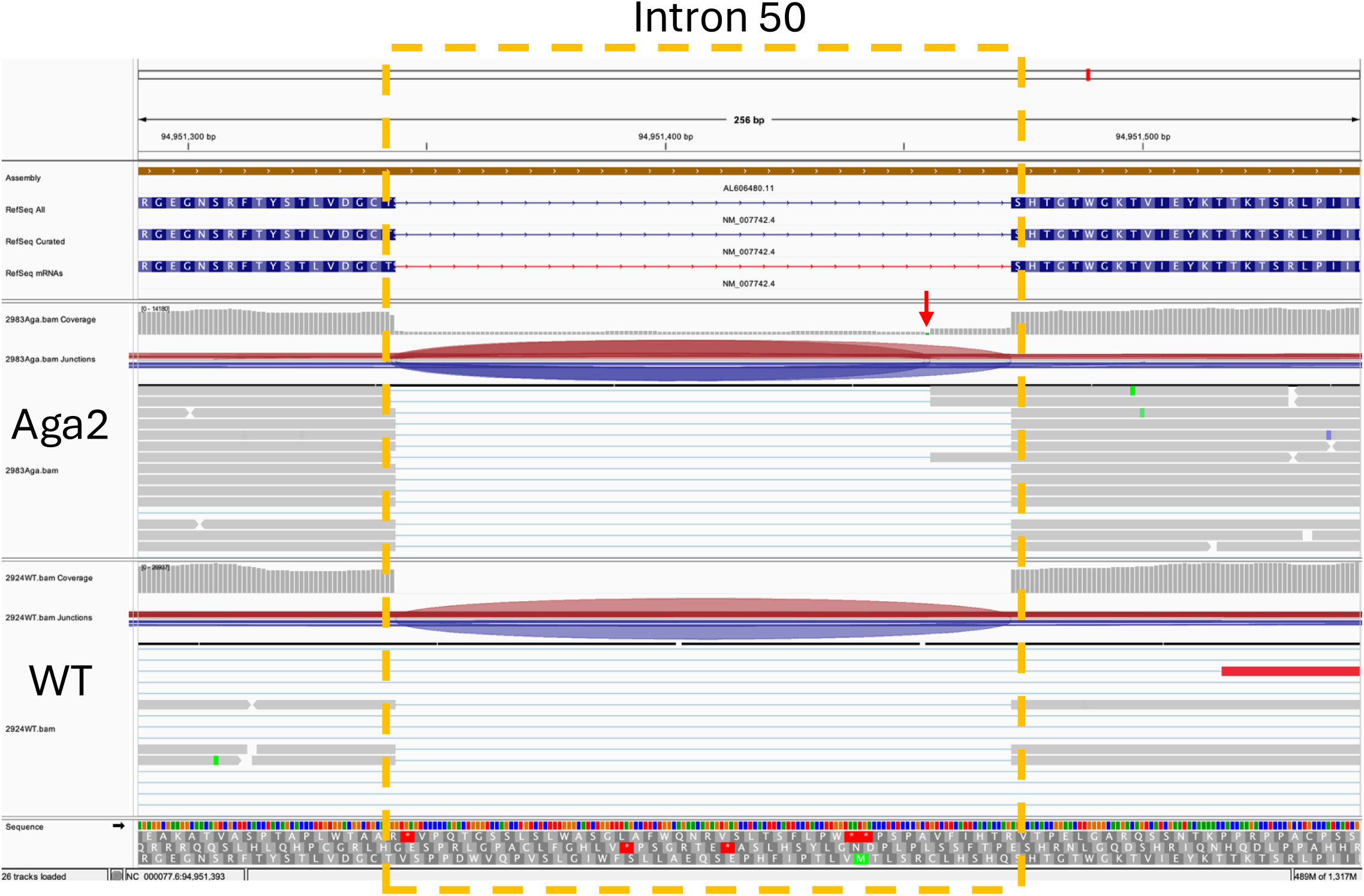
Aga2 and WT transcripts from RNA-seq mapped to the last intron of *Col1a1* gene using Integrative Genomics Viewer (IGV) software. Transcripts derived from RNA-seq of Aga2 (top tracks) and WT (bottom tracks) femurs were mapped to *Col1a1* (reference genome: Mus musculus GFC_000001635.26). In the Aga2 samples, transcripts are mapping to the entire intron, with higher representation of intron 50 3’ end. The red arrow points to the green bar that indicates variation with respect to the reference sequence. That is the position of the T➔A transition that generates the phenotype. No transcripts coming from WT bones mapped to intron 50.

**Figure S4:**
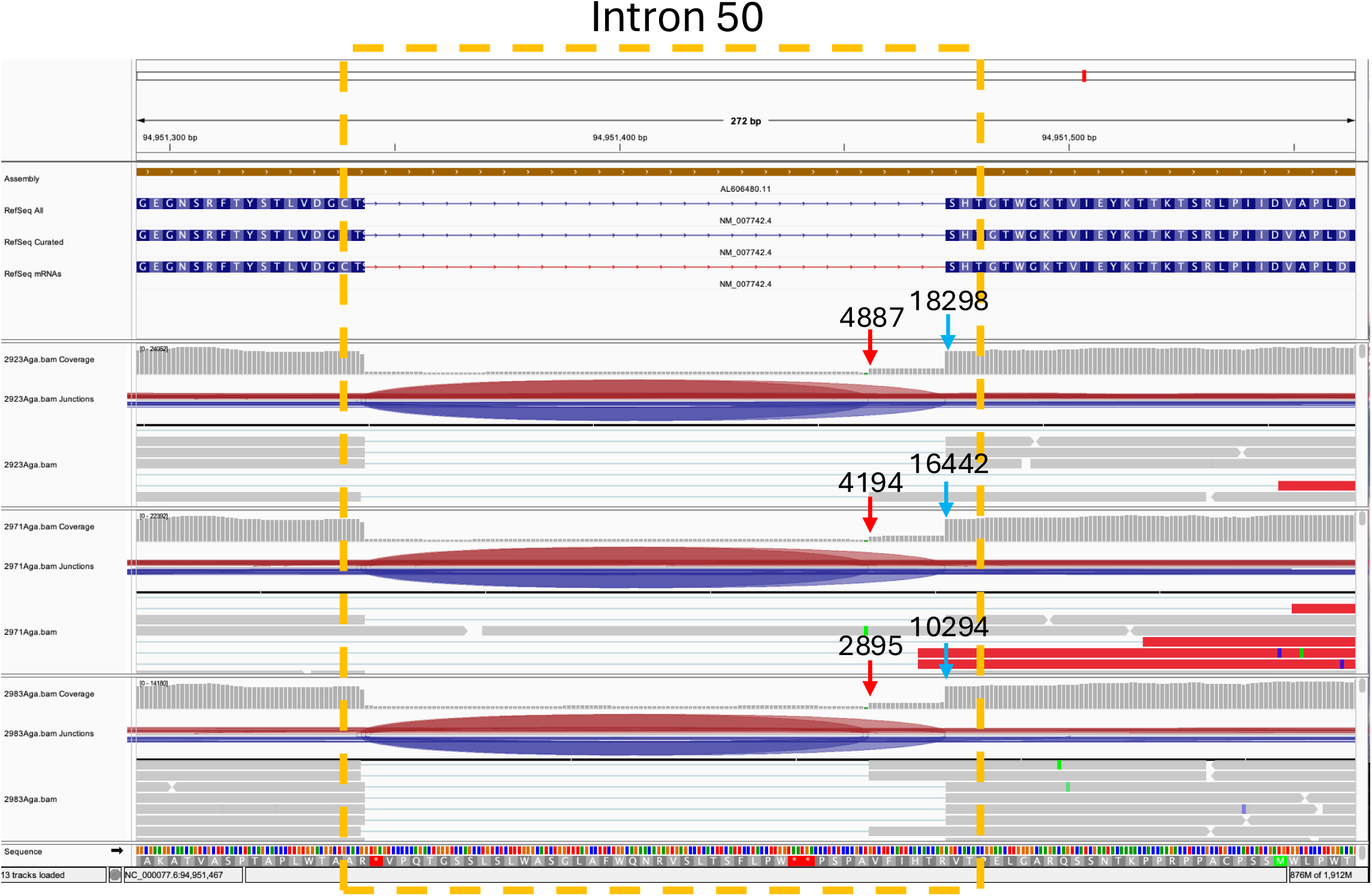
Aga2 transcripts from RNA-seq mapped to the last intron of *Col1a1* gene using Integrative Genomics Viewer (IGV) software. **Femurs of 3 individuals were used to extract RNA for sequencing (mouse IDs: 2923, 2972, and 2983).** The sequenced transcripts mapped to the last intron of *Col1a1* (reference genome: Mus musculus GFC_000001635.26) and showed the presence of the Aga2 aberrant mRNA. Reads counted for Aga2 mRNA (red arrows) are ∼25% of the reads counted for the total amount of *Col1a1* mRNA (blue arrows).

**Supplementary table 1: Scoring data from 8 observers.**

List of animals that have been analyzed using X-rays and Holeboard assay with corresponding numbers of fractures and deformities counted by 8 different observers. Column A contains the ID number present on the ear tag; column B indicates mouse genotype; column C indicates the sex; column D indicates the age. All other columns have been named with the initials of the observer’s name followed by the 7 skeletal sites that have been considered: Humerus, Forearm, Olecranon, Ischium, Femur, Tibia, and Calcaneus. At each skeletal site, fractures (F.) and deformities (D.) were counted separately. For each observer, the measurement of the IIR is reported as “pelvis ratio” along with scoliosis presence (binary variable: Yes or No). For each observer the sum of the total fractures, total deformities, total score (fractures + deformities) was also calculated. In columns ES, ET, and EU, we reported the average total fractures, total deformities, and total score calculated over the 8 different observers. Columns EV and EW indicate how many observers reported scoliosis and how many did not per each mouse.

**Supplementary table 2: Interobserver Correlation Coefficient for total score, scoliosis, and IIR.**

**Tab 1:** Column A contains the ID number present on the ear tag; column B indicates mouse genotype; column C indicates the sex; column D indicates the age. Columns E-L report the total scores measured by the 8 observers. Columns O-T report the Anova Two-Factor Without Replication used to calculate the Interobserver Correlation Coefficient, which indicates the degree of agreement between the different observers. These tests have been run using Excel.

**Tab 2:** Column A contains the ID number present on the ear tag; column B indicates mouse genotype; column C indicates the sex; column D indicates the age. Columns E-L report the “pelvis ratios” (or IIR) measured by the 8 observers. Columns O-T report the Anova Two-Factor Without Replication used to calculate the Interobserver Correlation Coefficient, which indicates the degree of agreement between the different observers. These tests have been run using Excel.

**Tab 3:** Column A contains the ID number present on the ear tag; column B indicates mouse genotype; column C indicates the sex; column D indicates the age. Columns E-L report the presence or absence of scoliosis according to the 8 different observers. Columns N-R report the Fleiss’ Kappa statistic (K), which indicates the degree of agreement between the different observers. These tests have been run using Excel.

**Supplementary table 3: Holeboard data at 30 and 60 minutes.**

Column A contains the ID number present on the ear tag; column B indicates mouse genotype; column C indicates the sex; column D indicates the age. Column E indicates the session interval, while column F the date of recording. Column H indicates the movements in cm that the mice performed throughout the 60 min of the assay. Poking is measured per each of the 9 holes present in the board (columns I-Q) and rearing on the hindlimbs is also reported (column R).

## Notes

### Competing Interest Statement

The authors have declared no competing interest.

