## Supplementary material for "Enhancing Wnt signaling lowers fracture incidence in a severe mouse model of Osteogenesis Imperfecta": SupTable1

| ID | Genotype | Sex | Age (Weeks) | MW-Humerus | MW-Humerus | MW-Forearm (Ulna & Radius) | MW-Forearm (Ulna & [illegible] | MW-Olecranon [illegible] |
|---|---|---|---|---|---|---|---|---|
| 167 | WT/AV | F | 5 | 0 | 0 | 0 | 0 | 0 |
| 168 | WT/AV | M | 5 | 0 | 0 | 0 | 0 | 0 |
| 301 | WT/AV | F | 5 | 0 | 0 | 0 | 0 | 0 |
| 304 | WT/AV | M | 5 | 0 | 0 | 0 | 0 | 0 |
| 332 | WT/AV | F | 5 | 0 | 0 | 0 | 0 | 0 |
| 4689 | WT/AV | M | 5 | 0 | 0 | 0 | 0 | 0 |
| 994 | WT/AV | F | 5 | 0 | 0 | 0 | 0 | 0 |
| 167 | WT/AV | F | 9 | 0 | 0 | 0 | 0 | 0 |
| 168 | WT/AV | M | 9 | 0 | 0 | 0 | 0 | 0 |
| 301 | WT/AV | F | 9 | 0 | 0 | 0 | 0 | 0 |
| 304 | WT/AV | M | 9 | 0 | 0 | 0 | 0 | 0 |
| 332 | WT/AV | F | 9 | 0 | 0 | 0 | 0 | 0 |
| 4689 | WT/AV | M | 9 | 0 | 0 | 0 | 0 | 0 |
| 994 | WT/AV | F | 9 | 0 | 0 | 0 | 0 | 0 |
| 167 | WT/AV | F | 13 | 0 | 0 | 0 | 0 | 0 |
| 168 | WT/AV | M | 13 | 0 | 0 | 0 | 0 | 0 |
| 301 | WT/AV | F | 13 | 0 | 0 | 0 | 0 | 0 |
| 304 | WT/AV | M | 13 | 0 | 0 | 0 | 0 | 0 |
| 332 | WT/AV | F | 13 | 0 | 0 | 0 | 0 | 0 |
| 4689 | WT/AV | M | 13 | 0 | 0 | 0 | 0 | 0 |
| 994 | WT/AV | F | 13 | 0 | 0 | 0 | 0 | 0 |
| 1652 | WT | F | 5 | 0 | 0 | 0 | 0 | 0 |
| 1654 | WT | F | 5 | 0 | 0 | 0 | 0 | 0 |
| 166 | WT | M | 5 | 0 | 0 | 0 | 0 | 0 |
| 169 | WT | M | 5 | 0 | 0 | 0 | 0 | 0 |
| 1931 | WT | F | 5 | 0 | 0 | 0 | 0 | 0 |
| 1932 | WT | F | 5 | 0 | 0 | 0 | 0 | 0 |
| 195 | WT | F | 5 | 0 | 0 | 0 | 0 | 0 |
| 302 | WT | F | 5 | 0 | 0 | 0 | 0 | 0 |

| | | | | | | | | |
|---|---|---|---|---|---|---|---|---|
| 303 | WT | M | 5 | 0 | 0 | 0 | 0 | 0 |
| 317 | WT | M | 5 | 0 | 0 | 0 | 0 | 0 |
| 322 | WT | M | 5 | 0 | 0 | 0 | 0 | 0 |
| 3514 | WT | F | 5 | 0 | 0 | 0 | 0 | 0 |
| 3521 | WT | F | 5 | 0 | 0 | 0 | 0 | 0 |
| 4114 | WT | F | 5 | 0 | 0 | 0 | 0 | 0 |
| 4599 | WT | M | 5 | 0 | 0 | 0 | 0 | 0 |
| 4685 | WT | F | 5 | 0 | 0 | 0 | 0 | 0 |
| 1652 | WT | F | 9 | 0 | 0 | 0 | 0 | 0 |
| 1654 | WT | F | 9 | 0 | 0 | 0 | 0 | 0 |
| 166 | WT | M | 9 | 0 | 0 | 0 | 0 | 0 |
| 169 | WT | M | 9 | 0 | 0 | 0 | 0 | 0 |
| 1931 | WT | F | 9 | 0 | 0 | 0 | 0 | 0 |
| 1932 | WT | F | 9 | 0 | 0 | 0 | 0 | 0 |
| 195 | WT | F | 9 | 0 | 0 | 0 | 0 | 0 |
| 302 | WT | F | 9 | 0 | 0 | 0 | 0 | 0 |
| 303 | WT | M | 9 | 0 | 0 | 0 | 0 | 0 |
| 317 | WT | M | 9 | 0 | 0 | 0 | 0 | 0 |
| 322 | WT | M | 9 | 0 | 0 | 0 | 0 | 0 |
| 3514 | WT | F | 9 | 0 | 0 | 0 | 0 | 0 |
| 3521 | WT | F | 9 | 0 | 0 | 0 | 0 | 0 |
| 4114 | WT | F | 9 | 0 | 0 | 0 | 0 | 0 |
| 4599 | WT | M | 9 | 0 | 0 | 0 | 0 | 0 |
| 4685 | WT | F | 9 | 0 | 0 | 0 | 0 | 0 |
| 1652 | WT | F | 13 | 0 | 0 | 0 | 0 | 0 |
| 1654 | WT | F | 13 | 0 | 0 | 0 | 0 | 0 |
| 0 | WT | M | 13 | 0 | 0 | 0 | 0 | 0 |
| 169 | WT | M | 13 | 0 | 0 | 0 | 0 | 0 |
| 1931 | WT | F | 13 | 0 | 0 | 0 | 0 | 0 |
| 1932 | WT | F | 13 | 0 | 0 | 0 | 0 | 0 |

| | | | | | | | | |
|---|---|---|---|---|---|---|---|---|
| 195 | WT | F | 13 | 0 | 0 | 0 | 0 | 0 |
| 302 | WT | F | 13 | 0 | 0 | 0 | 0 | 0 |
| 303 | WT | M | 13 | 0 | 0 | 0 | 0 | 0 |
| 317 | WT | M | 13 | 0 | 0 | 0 | 0 | 0 |
| 322 | WT | M | 13 | 0 | 0 | 0 | 0 | 0 |
| 3514 | WT | F | 13 | 0 | 0 | 0 | 0 | 0 |
| 3521 | WT | F | 13 | 0 | 0 | 0 | 0 | 0 |
| 4114 | WT | F | 13 | 0 | 0 | 0 | 0 | 0 |
| 4599 | WT | M | 13 | 0 | 0 | 0 | 0 | 0 |
| 4685 | WT | F | 13 | 0 | 0 | 0 | 0 | 0 |
| 1910 | OI | F | 9 | 0 | 0 | 0 | 0 | 0 |
| 351 | OI | M | 9 | 1 | 0 | 0 | 1 | 0 |
| 4115 | OI | F | 9 | 0 | 0 | 0 | 2 | 0 |
| 1214 | OI | M | 9 | 0 | 0 | 0 | 0 | 0 |
| 1910 | OI | F | 5 | 0 | 0 | 0 | 0 | 0 |
| 345 | OI | F | 9 | 0 | 0 | 0 | 2 | 0 |
| 4690 | OI | M | 9 | 0 | 1 | 0 | 2 | 0 |
| 3520 | OI | F | 9 | 1 | 0 | 0 | 2 | 0 |
| 349 | OI | M | 9 | 1 | 1 | 0 | 2 | 0 |
| 989 | OI | F | 9 | 0 | 2 | 0 | 2 | 0 |
| 1210 | OI | M | 9 | 1 | 1 | 0 | 1 | 0 |
| 873 | OI | M | 9 | 1 | 1 | 0 | 2 | 0 |
| 344 | OI | M | 9 | 1 | 1 | 0 | 2 | 0 |
| 319 | OI | M | 9 | 2 | 0 | 0 | 1 | 0 |
| 1910 | OI | F | 13 | 0 | 0 | 0 | 0 | 0 |
| 351 | OI | M | 13 | 0 | 1 | 0 | 1 | 0 |
| 1163 | OI | F | 9 | 1 | 0 | 0 | 2 | 0 |
| 1211 | OI | M | 9 | 0 | 2 | 1 | 0 | 0 |
| 4115 | OI | F | 13 | 0 | 0 | 0 | 2 | 0 |
| 351 | OI | M | 5 | 2 | 0 | 0 | 1 | 0 |

| | | | | | | | | |
|---|---|---|---|---|---|---|---|---|
| 198 | OI | F | 9 | 0 | 1 | 0 | 2 | 0 |
| 987 | OI | F | 9 | 1 | 1 | 0 | 2 | 0 |
| 4596 | OI | M | 9 | 1 | 1 | 0 | 2 | 0 |
| 1214 | OI | M | 13 | 0 | 0 | 0 | 0 | 0 |
| 1214 | OI | M | 5 | 0 | 0 | 0 | 0 | 0 |
| 4115 | OI | F | 5 | 1 | 0 | 0 | 2 | 0 |
| 3520 | OI | F | 5 | 0 | 0 | 0 | 2 | 0 |
| 345 | OI | F | 5 | 1 | 0 | 2 | 0 | 0 |
| 4690 | OI | M | 5 | 1 | 0 | 0 | 2 | 0 |
| 345 | OI | F | 13 | 0 | 0 | 0 | 2 | 0 |
| 319 | OI | M | 5 | 1 | 0 | 0 | 1 | 0 |
| 4690 | OI | M | 13 | 0 | 1 | 0 | 2 | 0 |
| 349 | OI | M | 13 | 0 | 2 | 0 | 2 | 0 |
| 349 | OI | M | 5 | 1 | 0 | 0 | 2 | 0 |
| 1210 | OI | M | 5 | 2 | 0 | 0 | 2 | 0 |
| 989 | OI | F | 5 | 1 | 0 | 0 | 2 | 0 |
| 344 | OI | M | 13 | 0 | 2 | 0 | 2 | 0 |
| 344 | OI | M | 5 | 1 | 0 | 0 | 2 | 0 |
| 1210 | OI | M | 13 | 0 | 2 | 0 | 2 | 0 |
| 873 | OI | M | 5 | 1 | 0 | 0 | 2 | 0 |
| 1211 | OI | M | 5 | 0 | 0 | 0 | 2 | 0 |
| 989 | OI | F | 13 | 1 | 1 | 0 | 2 | 0 |
| 873 | OI | M | 13 | 0 | 2 | 0 | 2 | 0 |
| 943 | OI | M | 5 | 1 | 0 | 0 | 2 | 0 |
| 319 | OI | M | 13 | 0 | 2 | 0 | 1 | 0 |
| 1211 | OI | M | 13 | 1 | 0 | 0 | 1 | 0 |
| 1163 | OI | F | 5 | 1 | 0 | 0 | 2 | 0 |
| 987 | OI | F | 5 | 1 | 1 | 0 | 2 | 0 |
| 198 | OI | F | 5 | 1 | 1 | 0 | 1 | 0 |
| 1163 | OI | F | 13 | 0 | 1 | 0 | 2 | 0 |

| | | | | | | | | |
|---|---|---|---|---|---|---|---|---|
| 941 | OI | F | 5 | 1 | 0 | 0 | 2 | 0 |
| 198 | OI | F | 13 | 0 | 1 | 0 | 2 | 0 |
| 987 | OI | F | 13 | 0 | 1 | 0 | 2 | 0 |
| 4596 | OI | M | 13 | 0 | 2 | 0 | 2 | 0 |
| 4596 | OI | M | 5 | 0 | 1 | 1 | 2 | 0 |
| 1933 | OI/AV | F | 5 | 0 | 1 | 0 | 0 | 0 |
| 1909 | OI/AV | F | 5 | 0 | 1 | 0 | 0 | 0 |
| 165 | OI/AV | F | 13 | 0 | 0 | 0 | 0 | 0 |
| 1933 | OI/AV | F | 13 | 0 | 0 | 0 | 0 | 0 |
| 165 | OI/AV | F | 5 | 0 | 0 | 0 | 0 | 0 |
| 3522 | OI/AV | M | 5 | 0 | 0 | 0 | 0 | 0 |
| 1934 | OI/AV | M | 5 | 0 | 0 | 0 | 0 | 0 |
| 3523 | OI/AV | M | 13 | 0 | 0 | 0 | 0 | 0 |
| 1909 | OI/AV | F | 13 | 0 | 0 | 0 | 0 | 0 |
| 3523 | OI/AV | M | 5 | 0 | 0 | 0 | 0 | 0 |
| 4117 | OI/AV | M | 13 | 0 | 0 | 0 | 0 | 0 |
| 1934 | OI/AV | M | 13 | 0 | 1 | 0 | 0 | 0 |
| 321 | OI/AV | M | 5 | 0 | 0 | 0 | 0 | 0 |
| 340 | OI/AV | F | 5 | 1 | 0 | 0 | 1 | 0 |
| 3522 | OI/AV | M | 13 | 0 | 0 | 0 | 0 | 0 |
| 339 | OI/AV | F | 13 | 0 | 0 | 0 | 0 | 0 |
| 321 | OI/AV | M | 13 | 0 | 0 | 0 | 0 | 0 |
| 340 | OI/AV | F | 13 | 0 | 0 | 0 | 1 | 0 |
| 4117 | OI/AV | M | 5 | 1 | 0 | 0 | 0 | 0 |
| 1935 | OI/AV | M | 5 | 0 | 0 | 0 | 0 | 0 |
| 1935 | OI/AV | M | 13 | 0 | 0 | 0 | 0 | 0 |
| 992 | OI/AV | F | 5 | 0 | 0 | 0 | 0 | 0 |
| 992 | OI/AV | F | 13 | 0 | 0 | 0 | 0 | 0 |
| 339 | OI/AV | F | 5 | 0 | 0 | 0 | 0 | 0 |
| 4686 | OI/AV | F | 5 | 0 | 0 | 0 | 0 | 0 |

| | | | | | | | | |
|---|---|---|---|---|---|---|---|---|
| 4686 | OI/AV | F | 13 | 0 | 1 | 0 | 0 | 0 |
| 350 | OI/AV | M | 5 | 0 | 0 | 0 | 1 | 0 |
| 320 | OI/AV | M | 5 | 0 | 0 | 0 | 2 | 0 |
| 196 | OI/AV | F | 5 | 0 | 0 | 0 | 0 | 0 |
| 3515 | OI/AV | M | 13 | 0 | 0 | 0 | 2 | 0 |
| 4598 | OI/AV | M | 5 | 1 | 0 | 0 | 1 | 0 |
| 1933 | OI/AV | F | 9 | 0 | 0 | 0 | 0 | 0 |
| 350 | OI/AV | M | 13 | 0 | 1 | 0 | 1 | 0 |
| 3523 | OI/AV | M | 9 | 0 | 0 | 0 | 0 | 0 |
| 165 | OI/AV | F | 9 | 0 | 0 | 0 | 0 | 0 |
| 196 | OI/AV | F | 13 | 0 | 0 | 0 | 0 | 0 |
| 3515 | OI/AV | M | 5 | 0 | 0 | 0 | 2 | 0 |
| 1909 | OI/AV | F | 9 | 0 | 0 | 0 | 0 | 0 |
| 4591 | OI/AV | F | 5 | 0 | 0 | 0 | 2 | 0 |
| 992 | OI/AV | F | 9 | 0 | 0 | 0 | 0 | 0 |
| 4591 | OI/AV | F | 13 | 0 | 1 | 0 | 2 | 0 |
| 318 | OI/AV | F | 13 | 0 | 0 | 0 | 2 | 0 |
| 318 | OI/AV | F | 5 | 0 | 0 | 1 | 1 | 0 |
| 339 | OI/AV | F | 9 | 0 | 0 | 0 | 0 | 0 |
| 4113 | OI/AV | F | 13 | 0 | 0 | 0 | 2 | 0 |
| 4113 | OI/AV | F | 5 | 0 | 0 | 0 | 2 | 0 |
| 1934 | OI/AV | M | 9 | 0 | 1 | 0 | 0 | 0 |
| 321 | OI/AV | M | 9 | 0 | 0 | 0 | 0 | 0 |
| 4594 | OI/AV | F | 5 | 0 | 0 | 0 | 2 | 0 |
| 3522 | OI/AV | M | 9 | 0 | 0 | 0 | 0 | 0 |
| 1930 | OI/AV | F | 5 | 0 | 0 | 0 | 2 | 0 |
| 4594 | OI/AV | F | 13 | 0 | 0 | 0 | 2 | 0 |
| 4592 | OI/AV | F | 5 | 0 | 0 | 0 | 2 | 0 |
| 340 | OI/AV | F | 9 | 0 | 0 | 0 | 1 | 0 |
| 4117 | OI/AV | M | 9 | 0 | 0 | 0 | 0 | 0 |

| | | | | | | | | |
|---|---|---|---|---|---|---|---|---|
| 1935 | OI/AV | M | 9 | 0 | 0 | 0 | 0 | 0 |
| 4686 | OI/AV | F | 9 | 0 | 1 | 0 | 0 | 0 |
| 196 | OI/AV | F | 9 | 0 | 0 | 0 | 0 | 0 |
| 4598 | OI/AV | M | 13 | 1 | 1 | 0 | 1 | 0 |
| 320 | OI/AV | M | 13 | 0 | 2 | 0 | 2 | 0 |
| 4592 | OI/AV | F | 13 | 0 | 0 | 0 | 2 | 0 |
| 197 | OI/AV | F | 13 | 0 | 0 | 0 | 2 | 0 |
| 197 | OI/AV | F | 5 | 0 | 0 | 0 | 2 | 0 |
| 164 | OI/AV | F | 5 | 0 | 0 | 0 | 2 | 0 |
| 3515 | OI/AV | M | 9 | 0 | 0 | 0 | 2 | 0 |
| 350 | OI/AV | M | 9 | 0 | 1 | 0 | 1 | 0 |
| 1930 | OI/AV | F | 13 | 0 | 1 | 0 | 2 | 0 |
| 4598 | OI/AV | M | 9 | 1 | 0 | 0 | 1 | 0 |
| 4591 | OI/AV | F | 9 | 0 | 1 | 0 | 2 | 0 |
| 4113 | OI/AV | F | 9 | 0 | 0 | 0 | 2 | 0 |
| 318 | OI/AV | F | 9 | 0 | 0 | 0 | 2 | 0 |
| 4594 | OI/AV | F | 9 | 0 | 0 | 0 | 2 | 0 |
| 320 | OI/AV | M | 9 | 2 | 0 | 0 | 2 | 0 |
| 197 | OI/AV | F | 9 | 0 | 0 | 0 | 2 | 0 |
| 164 | OI/AV | F | 13 | 0 | 2 | 0 | 2 | 0 |
| 1930 | OI/AV | F | 9 | 0 | 1 | 0 | 2 | 0 |
| 4592 | OI/AV | F | 9 | 0 | 0 | 0 | 2 | 0 |
| 164 | OI/AV | F | 9 | 1 | 1 | 0 | 2 | 0 |

| MW-Olecranon | MW-Ischium | MW-Femur | MW-Femur | MW-Tibia F | MW-Tibia D | MW-Calcaneous | MW-Calcaneu | MW-Total Fractures | MW-Total Deformities | MW-Total Score |
|---|---|---|---|---|---|---|---|---|---|---|
| 0 | 0 | 0 | 0 | 0 | 0 | 0 | 0 | 0 | 0 | 0 |
| 0 | 0 | 0 | 0 | 0 | 0 | 0 | 0 | 0 | 0 | 0 |
| 0 | 0 | 0 | 0 | 0 | 0 | 0 | 0 | 0 | 0 | 0 |
| 0 | 0 | 0 | 0 | 0 | 0 | 0 | 0 | 0 | 0 | 0 |
| 0 | 0 | 0 | 0 | 0 | 0 | 0 | 0 | 0 | 0 | 0 |
| 0 | 0 | 0 | 0 | 0 | 0 | 0 | 0 | 0 | 0 | 0 |
| 0 | 0 | 0 | 0 | 0 | 0 | 0 | 0 | 0 | 0 | 0 |
| 0 | 0 | 0 | 0 | 0 | 0 | 0 | 0 | 0 | 0 | 0 |
| 0 | 0 | 0 | 0 | 0 | 0 | 0 | 0 | 0 | 0 | 0 |
| 0 | 0 | 0 | 0 | 0 | 0 | 0 | 0 | 0 | 0 | 0 |
| 0 | 0 | 0 | 0 | 0 | 0 | 0 | 0 | 0 | 0 | 0 |
| 0 | 0 | 0 | 0 | 0 | 0 | 0 | 0 | 0 | 0 | 0 |
| 0 | 0 | 0 | 0 | 0 | 0 | 0 | 0 | 0 | 0 | 0 |
| 0 | 0 | 0 | 0 | 0 | 0 | 0 | 0 | 0 | 0 | 0 |
| 0 | 0 | 0 | 0 | 0 | 0 | 0 | 0 | 0 | 0 | 0 |
| 0 | 0 | 0 | 0 | 0 | 0 | 0 | 0 | 0 | 0 | 0 |
| 0 | 0 | 0 | 0 | 0 | 0 | 0 | 0 | 0 | 0 | 0 |
| 0 | 0 | 0 | 0 | 0 | 0 | 0 | 0 | 0 | 0 | 0 |
| 0 | 0 | 0 | 0 | 0 | 0 | 0 | 0 | 0 | 0 | 0 |
| 0 | 0 | 0 | 0 | 0 | 0 | 0 | 0 | 0 | 0 | 0 |
| 0 | 0 | 0 | 0 | 0 | 0 | 0 | 0 | 0 | 0 | 0 |
| 0 | 0 | 0 | 0 | 0 | 0 | 0 | 0 | 0 | 0 | 0 |
| 0 | 0 | 0 | 0 | 0 | 0 | 0 | 0 | 0 | 0 | 0 |
| 0 | 0 | 0 | 0 | 0 | 0 | 0 | 0 | 0 | 0 | 0 |
| 0 | 0 | 0 | 0 | 0 | 0 | 0 | 0 | 0 | 0 | 0 |
| 0 | 0 | 0 | 0 | 0 | 0 | 0 | 0 | 0 | 0 | 0 |
| 0 | 0 | 0 | 0 | 0 | 0 | 0 | 0 | 0 | 0 | 0 |
| 0 | 0 | 0 | 0 | 0 | 0 | 0 | 0 | 0 | 0 | 0 |
| 0 | 0 | 0 | 0 | 0 | 0 | 0 | 0 | 0 | 0 | 0 |

| | | | | | | | | | | |
|---|---|---|---|---|---|---|---|---|---|---|
| 0 | 0 | 0 | 0 | 0 | 0 | 0 | 0 | 0 | 0 | 0 |
| 0 | 0 | 0 | 0 | 0 | 0 | 0 | 0 | 0 | 0 | 0 |
| 0 | 0 | 0 | 0 | 0 | 0 | 0 | 0 | 0 | 0 | 0 |
| 0 | 0 | 0 | 0 | 0 | 0 | 0 | 0 | 0 | 0 | 0 |
| 0 | 0 | 0 | 0 | 0 | 0 | 0 | 0 | 0 | 0 | 0 |
| 0 | 0 | 0 | 0 | 0 | 0 | 0 | 0 | 0 | 0 | 0 |
| 0 | 0 | 0 | 0 | 0 | 0 | 0 | 0 | 0 | 0 | 0 |
| 0 | 0 | 0 | 0 | 0 | 0 | 0 | 0 | 0 | 0 | 0 |
| 0 | 0 | 0 | 0 | 0 | 0 | 0 | 0 | 0 | 0 | 0 |
| 0 | 0 | 0 | 0 | 0 | 0 | 0 | 0 | 0 | 0 | 0 |
| 0 | 0 | 0 | 0 | 0 | 0 | 0 | 0 | 0 | 0 | 0 |
| 0 | 0 | 0 | 0 | 0 | 0 | 0 | 0 | 0 | 0 | 0 |
| 0 | 0 | 0 | 0 | 0 | 0 | 0 | 0 | 0 | 0 | 0 |
| 0 | 0 | 0 | 0 | 0 | 0 | 0 | 0 | 0 | 0 | 0 |
| 0 | 0 | 0 | 0 | 0 | 0 | 0 | 0 | 0 | 0 | 0 |
| 0 | 0 | 0 | 0 | 0 | 0 | 0 | 0 | 0 | 0 | 0 |
| 0 | 0 | 0 | 0 | 0 | 0 | 0 | 0 | 0 | 0 | 0 |
| 0 | 0 | 0 | 0 | 0 | 0 | 0 | 0 | 0 | 0 | 0 |
| 0 | 0 | 0 | 0 | 0 | 0 | 0 | 0 | 0 | 0 | 0 |
| 0 | 0 | 0 | 0 | 0 | 0 | 0 | 0 | 0 | 0 | 0 |
| 0 | 0 | 0 | 0 | 0 | 0 | 0 | 0 | 0 | 0 | 0 |
| 0 | 0 | 0 | 0 | 0 | 0 | 0 | 0 | 0 | 0 | 0 |
| 0 | 0 | 0 | 0 | 0 | 0 | 0 | 0 | 0 | 0 | 0 |
| 0 | 0 | 0 | 0 | 0 | 0 | 0 | 0 | 0 | 0 | 0 |
| 0 | 0 | 0 | 0 | 0 | 0 | 0 | 0 | 0 | 0 | 0 |
| 0 | 0 | 0 | 0 | 0 | 0 | 0 | 0 | 0 | 0 | 0 |
| 0 | 0 | 0 | 0 | 0 | 0 | 0 | 0 | 0 | 0 | 0 |
| 0 | 0 | 0 | 0 | 0 | 0 | 0 | 0 | 0 | 0 | 0 |
| 0 | 0 | 0 | 0 | 0 | 0 | 0 | 0 | 0 | 0 | 0 |
| 0 | 0 | 0 | 0 | 0 | 0 | 0 | 0 | 0 | 0 | 0 |

| | | | | | | | | | | |
|---|---|---|---|---|---|---|---|---|---|---|
| 0 | 0 | 0 | 0 | 0 | 0 | 0 | 0 | 0 | 0 | 0 |
| 0 | 0 | 0 | 0 | 0 | 0 | 0 | 0 | 0 | 0 | 0 |
| 0 | 0 | 0 | 0 | 0 | 0 | 0 | 0 | 0 | 0 | 0 |
| 0 | 0 | 0 | 0 | 0 | 0 | 0 | 0 | 0 | 0 | 0 |
| 0 | 0 | 0 | 0 | 0 | 0 | 0 | 0 | 0 | 0 | 0 |
| 0 | 0 | 0 | 0 | 0 | 0 | 0 | 0 | 0 | 0 | 0 |
| 0 | 0 | 0 | 0 | 0 | 0 | 0 | 0 | 0 | 0 | 0 |
| 0 | 0 | 0 | 0 | 0 | 0 | 0 | 0 | 0 | 0 | 0 |
| 0 | 0 | 0 | 0 | 0 | 0 | 0 | 0 | 0 | 0 | 0 |
| 0 | 0 | 0 | 0 | 0 | 0 | 0 | 0 | 0 | 0 | 0 |
| 0 | 0 | 0 | 0 | 0 | 2 | 0 | 2 | 0 | 4 | 4 |
| 0 | 2 | 0 | 0 | 0 | 0 | 0 | 2 | 1 | 5 | 6 |
| 0 | 2 | 0 | 1 | 0 | 0 | 0 | 1 | 0 | 6 | 6 |
| 1 | 1 | 0 | 0 | 0 | 1 | 0 | 2 | 0 | 5 | 5 |
| 0 | 0 | 0 | 0 | 1 | 0 | 0 | 2 | 1 | 2 | 3 |
| 0 | 1 | 0 | 0 | 0 | 0 | 0 | 2 | 0 | 5 | 5 |
| 0 | 2 | 1 | 0 | 0 | 0 | 2 | 0 | 3 | 5 | 8 |
| 0 | 2 | 0 | 0 | 1 | 1 | 0 | 1 | 2 | 6 | 8 |
| 0 | 2 | 0 | 0 | 0 | 0 | 0 | 2 | 1 | 7 | 8 |
| 0 | 2 | 0 | 0 | 0 | 0 | 0 | 2 | 0 | 8 | 8 |
| 1 | 2 | 0 | 0 | 0 | 1 | 0 | 2 | 1 | 8 | 9 |
| 0 | 2 | 0 | 0 | 0 | 1 | 0 | 1 | 1 | 7 | 8 |
| 0 | 2 | 0 | 0 | 0 | 1 | 0 | 2 | 1 | 8 | 9 |
| 0 | 2 | 0 | 0 | 1 | 1 | 0 | 1 | 3 | 5 | 8 |
| 0 | 0 | 0 | 0 | 0 | 1 | 0 | 2 | 0 | 3 | 3 |
| 0 | 2 | 0 | 0 | 0 | 0 | 0 | 2 | 0 | 6 | 6 |
| 0 | 2 | 0 | 0 | 0 | 2 | 0 | 2 | 1 | 8 | 9 |
| 2 | 2 | 0 | 0 | 0 | 1 | 0 | 2 | 1 | 9 | 10 |
| 0 | 2 | 0 | 1 | 0 | 0 | 0 | 1 | 0 | 6 | 6 |
| 0 | 2 | 0 | 0 | 0 | 0 | 0 | 2 | 2 | 5 | 7 |

| | | | | | | | | | | |
|---|---|---|---|---|---|---|---|---|---|---|
| 0 | 2 | 0 | 0 | 0 | 2 | 0 | 1 | 0 | 8 | 8 |
| 0 | 2 | 1 | 0 | 0 | 2 | 0 | 2 | 2 | 9 | 11 |
| 0 | 1 | 0 | 1 | 1 | 2 | 0 | 2 | 2 | 9 | 11 |
| 1 | 2 | 0 | 0 | 0 | 0 | 0 | 2 | 0 | 5 | 5 |
| 1 | 1 | 0 | 0 | 0 | 1 | 1 | 1 | 1 | 4 | 5 |
| 0 | 2 | 1 | 0 | 0 | 0 | 0 | 1 | 2 | 5 | 7 |
| 0 | 2 | 0 | 0 | 1 | 0 | 0 | 1 | 1 | 5 | 6 |
| 0 | 2 | 0 | 0 | 0 | 0 | 1 | 1 | 4 | 3 | 7 |
| 0 | 2 | 0 | 0 | 0 | 0 | 2 | 0 | 3 | 4 | 7 |
| 0 | 2 | 0 | 0 | 0 | 0 | 0 | 2 | 0 | 6 | 6 |
| 0 | 2 | 0 | 0 | 1 | 1 | 0 | 0 | 2 | 4 | 6 |
| 0 | 2 | 0 | 1 | 0 | 0 | 0 | 2 | 0 | 8 | 8 |
| 0 | 2 | 0 | 0 | 0 | 0 | 0 | 2 | 0 | 8 | 8 |
| 0 | 2 | 0 | 0 | 0 | 0 | 1 | 1 | 2 | 5 | 7 |
| 0 | 2 | 0 | 0 | 1 | 0 | 0 | 2 | 3 | 6 | 9 |
| 0 | 2 | 0 | 0 | 0 | 0 | 1 | 1 | 2 | 5 | 7 |
| 0 | 2 | 0 | 0 | 0 | 1 | 0 | 2 | 0 | 9 | 9 |
| 0 | 2 | 0 | 0 | 0 | 1 | 0 | 2 | 1 | 7 | 8 |
| 0 | 2 | 0 | 0 | 0 | 1 | 1 | 1 | 1 | 8 | 9 |
| 0 | 2 | 0 | 0 | 1 | 0 | 0 | 2 | 2 | 6 | 8 |
| 0 | 2 | 0 | 0 | 0 | 2 | 0 | 2 | 0 | 8 | 8 |
| 0 | 2 | 0 | 0 | 0 | 0 | 0 | 2 | 1 | 7 | 8 |
| 0 | 2 | 0 | 0 | 0 | 1 | 0 | 1 | 0 | 8 | 8 |
| 0 | 2 | 0 | 0 | 2 | 0 | 0 | 2 | 3 | 6 | 9 |
| 0 | 2 | 0 | 0 | 1 | 1 | 0 | 1 | 1 | 7 | 8 |
| 1 | 2 | 0 | 0 | 0 | 1 | 0 | 2 | 1 | 7 | 8 |
| 0 | 2 | 0 | 0 | 0 | 2 | 0 | 2 | 1 | 8 | 9 |
| 0 | 2 | 0 | 0 | 2 | 0 | 0 | 2 | 3 | 7 | 10 |
| 0 | 2 | 0 | 0 | 1 | 1 | 1 | 1 | 3 | 6 | 9 |
| 0 | 2 | 0 | 0 | 0 | 1 | 0 | 2 | 0 | 8 | 8 |

| | | | | | | | | | | |
|---|---|---|---|---|---|---|---|---|---|---|
| 0 | 2 | 0 | 0 | 0 | 2 | 0 | 2 | 1 | 8 | 9 |
| 1 | 2 | 0 | 0 | 0 | 2 | 0 | 2 | 0 | 10 | 10 |
| 0 | 2 | 0 | 1 | 0 | 2 | 0 | 0 | 0 | 8 | 8 |
| 0 | 2 | 0 | 1 | 0 | 2 | 0 | 2 | 0 | 11 | 11 |
| 0 | 1 | 1 | 0 | 0 | 2 | 1 | 1 | 3 | 7 | 10 |
| 0 | 0 | 0 | 0 | 0 | 0 | 0 | 1 | 0 | 2 | 2 |
| 0 | 0 | 0 | 0 | 0 | 0 | 0 | 2 | 0 | 3 | 3 |
| 0 | 2 | 0 | 0 | 0 | 0 | 0 | 1 | 0 | 3 | 3 |
| 0 | 0 | 0 | 0 | 0 | 0 | 0 | 1 | 0 | 1 | 1 |
| 0 | 2 | 0 | 0 | 0 | 0 | 0 | 1 | 0 | 3 | 3 |
| 0 | 0 | 0 | 0 | 0 | 0 | 2 | 0 | 2 | 0 | 2 |
| 0 | 0 | 0 | 0 | 0 | 0 | 0 | 2 | 0 | 2 | 2 |
| 0 | 0 | 0 | 0 | 0 | 1 | 0 | 2 | 0 | 3 | 3 |
| 0 | 1 | 0 | 0 | 0 | 0 | 0 | 2 | 0 | 3 | 3 |
| 0 | 0 | 0 | 0 | 1 | 0 | 1 | 1 | 2 | 1 | 3 |
| 0 | 1 | 0 | 1 | 0 | 0 | 0 | 1 | 0 | 3 | 3 |
| 0 | 0 | 0 | 0 | 0 | 0 | 0 | 2 | 0 | 3 | 3 |
| 0 | 2 | 0 | 0 | 0 | 0 | 1 | 1 | 1 | 3 | 4 |
| 0 | 2 | 0 | 0 | 0 | 0 | 1 | 0 | 2 | 3 | 5 |
| 0 | 0 | 0 | 0 | 0 | 0 | 0 | 2 | 0 | 2 | 2 |
| 0 | 2 | 0 | 0 | 0 | 1 | 0 | 1 | 0 | 4 | 4 |
| 0 | 2 | 0 | 0 | 0 | 0 | 0 | 2 | 0 | 4 | 4 |
| 0 | 2 | 0 | 0 | 0 | 0 | 0 | 1 | 0 | 4 | 4 |
| 0 | 1 | 1 | 0 | 0 | 0 | 0 | 1 | 2 | 2 | 4 |
| 0 | 2 | 0 | 0 | 0 | 0 | 1 | 1 | 1 | 3 | 4 |
| 0 | 2 | 0 | 0 | 0 | 0 | 0 | 2 | 0 | 4 | 4 |
| 1 | 2 | 0 | 0 | 0 | 0 | 1 | 0 | 1 | 3 | 4 |
| 1 | 2 | 0 | 0 | 0 | 0 | 0 | 1 | 0 | 4 | 4 |
| 0 | 2 | 0 | 0 | 0 | 1 | 1 | 0 | 1 | 3 | 4 |
| 0 | 2 | 0 | 0 | 0 | 0 | 2 | 0 | 2 | 2 | 4 |

| | | | | | | | | | | |
|---|---|---|---|---|---|---|---|---|---|---|
| 0 | 2 | 0 | 0 | 0 | 0 | 0 | 2 | 0 | 5 | 5 |
| 0 | 2 | 0 | 0 | 0 | 0 | 1 | 1 | 1 | 4 | 5 |
| 0 | 2 | 0 | 0 | 0 | 0 | 2 | 0 | 2 | 4 | 6 |
| 0 | 2 | 0 | 0 | 1 | 1 | 0 | 1 | 1 | 4 | 5 |
| 0 | 2 | 0 | 0 | 0 | 0 | 0 | 2 | 0 | 6 | 6 |
| 0 | 2 | 0 | 0 | 0 | 1 | 1 | 1 | 2 | 5 | 7 |
| 0 | 0 | 0 | 0 | 0 | 0 | 0 | 1 | 0 | 1 | 1 |
| 0 | 2 | 0 | 0 | 0 | 0 | 0 | 2 | 0 | 6 | 6 |
| 0 | 0 | 0 | 0 | 0 | 1 | 0 | 2 | 0 | 3 | 3 |
| 0 | 2 | 0 | 0 | 0 | 0 | 0 | 2 | 0 | 4 | 4 |
| 0 | 2 | 0 | 0 | 0 | 2 | 0 | 2 | 0 | 6 | 6 |
| 0 | 2 | 0 | 0 | 1 | 0 | 2 | 0 | 3 | 4 | 7 |
| 0 | 1 | 0 | 0 | 0 | 0 | 0 | 2 | 0 | 3 | 3 |
| 0 | 2 | 0 | 0 | 0 | 0 | 0 | 2 | 0 | 6 | 6 |
| 1 | 2 | 0 | 0 | 0 | 0 | 0 | 1 | 0 | 4 | 4 |
| 0 | 2 | 0 | 0 | 0 | 0 | 0 | 2 | 0 | 7 | 7 |
| 0 | 2 | 0 | 0 | 0 | 2 | 0 | 1 | 0 | 7 | 7 |
| 0 | 2 | 0 | 0 | 1 | 1 | 1 | 0 | 3 | 4 | 7 |
| 0 | 2 | 0 | 0 | 0 | 1 | 0 | 1 | 0 | 4 | 4 |
| 0 | 2 | 0 | 0 | 0 | 2 | 0 | 0 | 0 | 6 | 6 |
| 0 | 2 | 0 | 0 | 2 | 0 | 0 | 0 | 2 | 4 | 6 |
| 0 | 0 | 0 | 0 | 0 | 0 | 0 | 2 | 0 | 3 | 3 |
| 0 | 2 | 0 | 0 | 0 | 0 | 0 | 2 | 0 | 4 | 4 |
| 0 | 2 | 0 | 0 | 0 | 2 | 0 | 1 | 0 | 7 | 7 |
| 0 | 0 | 0 | 0 | 0 | 0 | 0 | 2 | 0 | 2 | 2 |
| 0 | 2 | 0 | 0 | 0 | 2 | 0 | 0 | 0 | 6 | 6 |
| 0 | 2 | 0 | 0 | 0 | 2 | 0 | 1 | 0 | 7 | 7 |
| 0 | 2 | 0 | 0 | 0 | 2 | 1 | 1 | 1 | 7 | 8 |
| 0 | 2 | 0 | 0 | 0 | 0 | 0 | 1 | 0 | 4 | 4 |
| 0 | 1 | 0 | 1 | 0 | 0 | 0 | 1 | 0 | 3 | 3 |

| | | | | | | | | | | |
|---|---|---|---|---|---|---|---|---|---|---|
| 0 | 2 | 0 | 0 | 0 | 0 | 0 | 2 | 0 | 4 | 4 |
| 0 | 2 | 0 | 0 | 0 | 0 | 0 | 2 | 0 | 5 | 5 |
| 0 | 2 | 0 | 0 | 0 | 2 | 0 | 1 | 0 | 5 | 5 |
| 1 | 2 | 0 | 0 | 0 | 1 | 0 | 2 | 1 | 8 | 9 |
| 0 | 2 | 0 | 0 | 0 | 0 | 0 | 2 | 0 | 8 | 8 |
| 0 | 2 | 0 | 0 | 0 | 2 | 0 | 2 | 0 | 8 | 8 |
| 0 | 2 | 0 | 0 | 0 | 2 | 0 | 1 | 0 | 7 | 7 |
| 0 | 2 | 0 | 0 | 1 | 1 | 1 | 0 | 2 | 5 | 7 |
| 0 | 1 | 0 | 0 | 1 | 0 | 2 | 0 | 3 | 3 | 6 |
| 0 | 2 | 0 | 0 | 0 | 0 | 0 | 2 | 0 | 6 | 6 |
| 0 | 2 | 0 | 0 | 0 | 0 | 0 | 2 | 0 | 6 | 6 |
| 0 | 2 | 0 | 0 | 0 | 2 | 0 | 0 | 0 | 7 | 7 |
| 0 | 2 | 0 | 0 | 0 | 1 | 0 | 2 | 1 | 6 | 7 |
| 0 | 2 | 0 | 0 | 0 | 0 | 0 | 2 | 0 | 7 | 7 |
| 0 | 2 | 0 | 0 | 0 | 2 | 1 | 0 | 1 | 6 | 7 |
| 0 | 2 | 0 | 0 | 0 | 2 | 0 | 1 | 0 | 7 | 7 |
| 0 | 2 | 0 | 0 | 0 | 2 | 0 | 1 | 0 | 7 | 7 |
| 0 | 2 | 0 | 0 | 0 | 0 | 0 | 2 | 2 | 6 | 8 |
| 0 | 2 | 0 | 0 | 0 | 2 | 0 | 1 | 0 | 7 | 7 |
| 0 | 1 | 0 | 0 | 0 | 1 | 0 | 2 | 0 | 8 | 8 |
| 0 | 2 | 0 | 0 | 0 | 2 | 0 | 0 | 0 | 7 | 7 |
| 0 | 2 | 0 | 0 | 0 | 2 | 0 | 2 | 0 | 8 | 8 |
| 0 | 1 | 0 | 0 | 0 | 1 | 0 | 2 | 1 | 7 | 8 |

| MW-Pelvis Ratio | MW-Scoliosis | GM-Humerus F | GM-Humerus D | GM-Forearm (Ulna & | GM-Forearm (Ulna & | GM-Olecranon F | GM-Olecranon D | GM-Ischium | GM-Femur F. |
|---|---|---|---|---|---|---|---|---|---|
| 1.00 | N | 0 | 0 | 0 | 0 | 0 | 0 | 0 | 0 |
| 1.00 | N | 0 | 0 | 0 | 0 | 0 | 0 | 0 | 0 |
| 1.00 | N | 0 | 0 | 0 | 0 | 0 | 0 | 0 | 0 |
| 1.00 | N | 0 | 0 | 0 | 0 | 0 | 0 | 0 | 0 |
| 1.00 | N | 0 | 0 | 0 | 0 | 0 | 0 | 0 | 0 |
| 1.00 | N | 0 | 0 | 0 | 0 | 0 | 0 | 0 | 0 |
| 1.00 | N | 0 | 0 | 0 | 0 | 0 | 0 | 0 | 0 |
| 1.00 | N | 0 | 0 | 0 | 0 | 0 | 0 | 0 | 0 |
| 1.00 | N | 0 | 0 | 0 | 0 | 0 | 0 | 0 | 0 |
| 1.00 | N | 0 | 0 | 0 | 0 | 0 | 0 | 0 | 0 |
| 1.00 | N | 0 | 0 | 0 | 0 | 0 | 0 | 0 | 0 |
| 1.00 | N | 0 | 0 | 0 | 0 | 0 | 0 | 0 | 0 |
| 1.00 | N | 0 | 0 | 0 | 0 | 0 | 0 | 0 | 0 |
| 1.00 | N | 0 | 0 | 0 | 0 | 0 | 0 | 0 | 0 |
| 1.00 | N | 0 | 0 | 0 | 0 | 0 | 0 | 0 | 0 |
| 1.00 | N | 0 | 0 | 0 | 0 | 0 | 0 | 0 | 0 |
| 1.00 | N | 0 | 0 | 0 | 0 | 0 | 0 | 0 | 0 |
| 1.00 | N | 0 | 0 | 0 | 0 | 0 | 0 | 0 | 0 |
| 1.00 | N | 0 | 0 | 0 | 0 | 0 | 0 | 0 | 0 |
| 1.00 | N | 0 | 0 | 0 | 0 | 0 | 0 | 0 | 0 |
| 1.00 | N | 0 | 0 | 0 | 0 | 0 | 0 | 0 | 0 |
| 1.00 | N | 0 | 0 | 0 | 0 | 0 | 0 | 0 | 0 |
| 1.00 | N | 0 | 0 | 0 | 0 | 0 | 0 | 0 | 0 |
| 1.00 | N | 0 | 0 | 0 | 0 | 0 | 0 | 0 | 0 |
| 1.00 | N | 0 | 0 | 0 | 0 | 0 | 0 | 0 | 0 |
| 1.00 | N | 0 | 0 | 0 | 0 | 0 | 0 | 0 | 0 |
| 1.00 | N | 0 | 0 | 0 | 0 | 0 | 0 | 0 | 0 |
| 1.00 | N | 0 | 0 | 0 | 0 | 0 | 0 | 0 | 0 |
| 1.00 | N | 0 | 0 | 0 | 0 | 0 | 0 | 0 | 0 |

| | | | | | | | | | |
|---|---|---|---|---|---|---|---|---|---|
| 1.00 | N | 0 | 0 | 0 | 0 | 0 | 0 | 0 | 0 |
| 1.00 | N | 0 | 0 | 0 | 0 | 0 | 0 | 0 | 0 |
| 1.00 | N | 0 | 0 | 0 | 0 | 0 | 0 | 0 | 0 |
| 1.00 | N | 0 | 0 | 0 | 0 | 0 | 0 | 0 | 0 |
| 1.00 | N | 0 | 0 | 0 | 0 | 0 | 0 | 0 | 0 |
| 1.00 | N | 0 | 0 | 0 | 0 | 0 | 0 | 0 | 0 |
| 1.00 | N | 0 | 0 | 0 | 0 | 0 | 0 | 0 | 0 |
| 1.00 | N | 0 | 0 | 0 | 0 | 0 | 0 | 0 | 0 |
| 1.00 | N | 0 | 0 | 0 | 0 | 0 | 0 | 0 | 0 |
| 1.00 | N | 0 | 0 | 0 | 0 | 0 | 0 | 0 | 0 |
| 1.00 | N | 0 | 0 | 0 | 0 | 0 | 0 | 0 | 0 |
| 1.00 | N | 0 | 0 | 0 | 0 | 0 | 0 | 0 | 0 |
| 1.00 | N | 0 | 0 | 0 | 0 | 0 | 0 | 0 | 0 |
| 1.00 | N | 0 | 0 | 0 | 0 | 0 | 0 | 0 | 0 |
| 1.00 | N | 0 | 0 | 0 | 0 | 0 | 0 | 0 | 0 |
| 1.00 | N | 0 | 0 | 0 | 0 | 0 | 0 | 0 | 0 |
| 1.00 | N | 0 | 0 | 0 | 0 | 0 | 0 | 0 | 0 |
| 1.00 | N | 0 | 0 | 0 | 0 | 0 | 0 | 0 | 0 |
| 1.00 | N | 0 | 0 | 0 | 0 | 0 | 0 | 0 | 0 |
| 1.00 | N | 0 | 0 | 0 | 0 | 0 | 0 | 0 | 0 |
| 1.00 | N | 0 | 0 | 0 | 0 | 0 | 0 | 0 | 0 |
| 1.00 | N | 0 | 0 | 0 | 0 | 0 | 0 | 0 | 0 |
| 1.00 | N | 0 | 0 | 0 | 0 | 0 | 0 | 0 | 0 |
| 1.00 | N | 0 | 0 | 0 | 0 | 0 | 0 | 0 | 0 |
| 1.00 | N | 0 | 0 | 0 | 0 | 0 | 0 | 0 | 0 |
| 1.00 | N | 0 | 0 | 0 | 0 | 0 | 0 | 0 | 0 |
| 1.00 | N | 0 | 0 | 0 | 0 | 0 | 0 | 0 | 0 |
| 1.00 | N | 0 | 0 | 0 | 0 | 0 | 0 | 0 | 0 |
| 1.00 | N | 0 | 0 | 0 | 0 | 0 | 0 | 0 | 0 |
| 1.00 | N | 0 | 0 | 0 | 0 | 0 | 0 | 0 | 0 |

| | | | | | | | | | |
|---|---|---|---|---|---|---|---|---|---|
| 1.00 | N | 0 | 0 | 0 | 0 | 0 | 0 | 0 | 0 |
| 1.00 | N | 0 | 0 | 0 | 0 | 0 | 0 | 0 | 0 |
| 1.00 | N | 0 | 0 | 0 | 0 | 0 | 0 | 0 | 0 |
| 1.00 | N | 0 | 0 | 0 | 0 | 0 | 0 | 0 | 0 |
| 1.00 | N | 0 | 0 | 0 | 0 | 0 | 0 | 0 | 0 |
| 1.00 | N | 0 | 0 | 0 | 0 | 0 | 0 | 0 | 0 |
| 1.00 | N | 0 | 0 | 0 | 0 | 0 | 0 | 0 | 0 |
| 1.00 | N | 0 | 0 | 0 | 0 | 0 | 0 | 0 | 0 |
| 1.00 | N | 0 | 0 | 0 | 0 | 0 | 0 | 0 | 0 |
| 1.00 | N | 0 | 0 | 0 | 0 | 0 | 0 | 0 | 0 |
| 0.90 | N | 0 | 0 | 0 | 1 | 0 | 0 | 0 | 0 |
| 0.00 | N | 1 | 0 | 0 | 1 | 0 | 0 | 2 | 0 |
| 0.46 | N | 0 | 0 | 0 | 2 | 0 | 0 | 2 | 0 |
| 0.58 | N | 0 | 1 | 0 | 1 | 0 | 1 | 2 | 0 |
| 0.85 | N | 0 | 0 | 0 | 0 | 0 | 0 | 0 | 0 |
| 0.53 | Y | 0 | 1 | 0 | 2 | 0 | 0 | 2 | 0 |
| 0.46 | N | 0 | 1 | 0 | 2 | 0 | 1 | 2 | 1 |
| 0.55 | Y | 0 | 1 | 0 | 2 | 0 | 0 | 2 | 0 |
| 0.51 | Y | 0 | 2 | 0 | 3 | 0 | 0 | 2 | 0 |
| 0.49 | N | 0 | 2 | 0 | 2 | 0 | 0 | 2 | 0 |
| 0.42 | Y | 0 | 2 | 0 | 2 | 0 | 0 | 2 | 0 |
| 0.55 | N | 1 | 1 | 0 | 2 | 0 | 0 | 2 | 0 |
| 0.44 | Y | 1 | 1 | 0 | 2 | 0 | 0 | 2 | 0 |
| 0.59 | Y | 1 | 3 | 0 | 2 | 0 | 1 | 2 | 0 |
| 0.87 | N | 0 | 0 | 0 | 2 | 0 | 0 | 1 | 0 |
| 0.57 | N | 0 | 1 | 0 | 1 | 0 | 0 | 2 | 0 |
| 0.53 | N | 0 | 1 | 0 | 0 | 0 | 2 | 2 | 0 |
| 0.47 | Y | 0 | 2 | 1 | 2 | 0 | 0 | 2 | 0 |
| 0.44 | N | 0 | 0 | 0 | 2 | 0 | 0 | 2 | 0 |
| 0.56 | N | 1 | 0 | 0 | 1 | 0 | 0 | 2 | 0 |

| | | | | | | | | | |
|---|---|---|---|---|---|---|---|---|---|
| 0.64 | Y | 0 | 1 | 0 | 2 | 0 | 1 | 2 | 0 |
| 0.56 | Y | 0 | 2 | 0 | 2 | 0 | 0 | 2 | 0 |
| 0.54 | N | 0 | 2 | 0 | 2 | 0 | 1 | 2 | 0 |
| 0.58 | Y | 0 | 1 | 0 | 1 | 0 | 0 | 2 | 0 |
| 0.68 | N | 0 | 1 | 0 | 1 | 1 | 0 | 2 | 0 |
| 0.53 | N | 0 | 0 | 0 | 2 | 0 | 0 | 2 | 0 |
| 0.65 | Y | 0 | 0 | 0 | 2 | 0 | 0 | 2 | 0 |
| 0.49 | Y | 0 | 1 | 0 | 2 | 0 | 0 | 2 | 0 |
| 0.59 | N | 0 | 0 | 0 | 2 | 0 | 2 | 2 | 0 |
| 0.58 | Y | 0 | 0 | 0 | 2 | 0 | 0 | 2 | 0 |
| 0.67 | Y | 0 | 1 | 0 | 1 | 0 | 1 | 2 | 0 |
| 0.50 | N | 0 | 1 | 0 | 2 | 0 | 0 | 2 | 0 |
| 0.55 | Y | 0 | 2 | 0 | 2 | 0 | 0 | 2 | 0 |
| 0.67 | Y | 0 | 1 | 0 | 2 | 0 | 0 | 2 | 0 |
| 0.53 | Y | 0 | 1 | 0 | 2 | 0 | 0 | 2 | 0 |
| 0.42 | Y | 0 | 1 | 0 | 2 | 0 | 0 | 2 | 0 |
| 0.43 | Y | 0 | 2 | 0 | 2 | 0 | 0 | 2 | 0 |
| 0.43 | Y | 0 | 2 | 0 | 2 | 0 | 0 | 2 | 0 |
| 0.46 | Y | 0 | 2 | 0 | 2 | 0 | 0 | 2 | 0 |
| 0.61 | Y | 1 | 1 | 0 | 2 | 0 | 0 | 2 | 0 |
| 0.50 | Y | 1 | 0 | 0 | 1 | 0 | 1 | 2 | 0 |
| 0.48 | N | 1 | 2 | 0 | 2 | 0 | 0 | 2 | 0 |
| 0.52 | N | 1 | 2 | 0 | 3 | 0 | 0 | 2 | 0 |
| 0.57 | Y | 0 | 2 | 0 | 2 | 0 | 0 | 2 | 0 |
| 0.68 | Y | 0 | 2 | 0 | 2 | 0 | 0 | 2 | 0 |
| 0.46 | Y | 2 | 1 | 0 | 2 | 0 | 0 | 2 | 0 |
| 0.53 | Y | 1 | 0 | 0 | 2 | 0 | 0 | 2 | 0 |
| 0.57 | Y | 0 | 1 | 0 | 2 | 0 | 0 | 2 | 0 |
| 0.70 | Y | 0 | 1 | 0 | 0 | 0 | 0 | 2 | 0 |
| 0.48 | Y | 0 | 2 | 0 | 2 | 0 | 0 | 2 | 0 |

| | | | | | | | | | |
|---|---|---|---|---|---|---|---|---|---|
| 0.68 | Y | 1 | 0 | 0 | 2 | 0 | 0 | 2 | 0 |
| 0.65 | Y | 0 | 1 | 0 | 2 | 0 | 0 | 2 | 0 |
| 0.47 | N | 1 | 1 | 0 | 2 | 0 | 0 | 2 | 1 |
| 0.68 | N | 0 | 2 | 0 | 2 | 0 | 0 | 2 | 0 |
| 0.58 | N | 0 | 2 | 0 | 2 | 0 | 1 | 2 | 0 |
| 0.77 | N | 0 | 0 | 0 | 0 | 0 | 0 | 0 | 0 |
| 0.91 | N | 0 | 0 | 0 | 0 | 0 | 0 | 2 | 0 |
| 0.66 | N | 0 | 0 | 0 | 0 | 0 | 0 | 2 | 0 |
| 0.84 | N | 0 | 0 | 0 | 2 | 0 | 0 | 1 | 0 |
| 0.63 | N | 0 | 0 | 0 | 0 | 0 | 0 | 2 | 0 |
| 0.85 | N | 0 | 0 | 0 | 0 | 0 | 0 | 2 | 0 |
| 1.03 | N | 0 | 1 | 0 | 0 | 0 | 2 | 2 | 0 |
| 0.88 | N | 0 | 0 | 0 | 0 | 0 | 0 | 1 | 0 |
| 0.94 | N | 0 | 0 | 0 | 1 | 0 | 0 | 1 | 0 |
| 1.00 | N | 0 | 0 | 0 | 0 | 0 | 0 | 2 | 0 |
| 0.62 | N | 0 | 0 | 0 | 0 | 0 | 0 | 2 | 0 |
| 0.96 | N | 0 | 1 | 0 | 2 | 0 | 0 | 1 | 0 |
| 0.69 | N | 0 | 0 | 0 | 0 | 0 | 0 | 2 | 0 |
| 0.62 | N | 0 | 0 | 0 | 2 | 0 | 0 | 2 | 0 |
| 0.86 | N | 0 | 0 | 0 | 0 | 0 | 0 | 2 | 0 |
| 0.68 | N | 0 | 0 | 0 | 2 | 0 | 0 | 2 | 0 |
| 0.67 | N | 0 | 0 | 0 | 2 | 0 | 0 | 2 | 0 |
| 0.65 | N | 0 | 0 | 0 | 3 | 0 | 0 | 2 | 0 |
| 0.67 | N | 0 | 0 | 0 | 0 | 0 | 0 | 2 | 1 |
| 0.47 | N | 0 | 1 | 0 | 0 | 0 | 0 | 2 | 0 |
| 0.45 | N | 0 | 1 | 0 | 0 | 0 | 2 | 0 | 0 |
| 0.60 | N | 0 | 0 | 0 | 0 | 0 | 1 | 2 | 0 |
| 0.59 | N | 0 | 0 | 0 | 2 | 0 | 1 | 2 | 0 |
| 0.60 | N | 0 | 0 | 0 | 0 | 0 | 2 | 2 | 0 |
| 0.55 | N | 0 | 1 | 0 | 1 | 0 | 0 | 2 | 0 |

| | | | | | | | | | |
|---|---|---|---|---|---|---|---|---|---|
| 0.62 | N | 0 | 1 | 0 | 1 | 0 | 0 | 2 | 0 |
| 0.58 | N | 1 | 0 | 0 | 1 | 0 | 0 | 2 | 0 |
| 0.69 | N | 0 | 0 | 0 | 2 | 0 | 0 | 2 | 0 |
| 0.54 | Y | 0 | 0 | 0 | 2 | 0 | 0 | 2 | 0 |
| 0.61 | Y | 0 | 0 | 0 | 2 | 0 | 0 | 2 | 0 |
| 0.63 | Y | 0 | 0 | 0 | 1 | 0 | 0 | 2 | 0 |
| 0.86 | N | 0 | 0 | 0 | 0 | 0 | 0 | 1 | 0 |
| 0.60 | N | 0 | 1 | 0 | 1 | 0 | 0 | 2 | 0 |
| 0.89 | N | 0 | 0 | 0 | 0 | 0 | 0 | 0 | 0 |
| 0.73 | Y | 0 | 0 | 0 | 0 | 0 | 0 | 2 | 0 |
| 0.50 | Y | 0 | 0 | 0 | 2 | 0 | 0 | 2 | 0 |
| 0.60 | Y | 0 | 0 | 0 | 2 | 0 | 0 | 2 | 0 |
| 0.89 | N | 0 | 0 | 0 | 1 | 0 | 0 | 1 | 0 |
| 0.50 | Y | 1 | 0 | 0 | 2 | 0 | 0 | 2 | 0 |
| 0.59 | N | 0 | 0 | 0 | 0 | 0 | 1 | 2 | 0 |
| 0.51 | Y | 0 | 1 | 0 | 2 | 0 | 0 | 2 | 0 |
| 0.63 | N | 0 | 0 | 0 | 2 | 0 | 0 | 2 | 0 |
| 0.68 | N | 0 | 0 | 0 | 2 | 0 | 0 | 2 | 0 |
| 0.62 | N | 0 | 0 | 0 | 2 | 0 | 0 | 2 | 0 |
| 0.56 | N | 0 | 0 | 0 | 2 | 0 | 0 | 2 | 0 |
| 0.57 | N | 0 | 0 | 0 | 2 | 0 | 0 | 2 | 0 |
| 0.97 | N | 0 | 1 | 0 | 1 | 0 | 0 | 2 | 0 |
| 0.67 | N | 0 | 0 | 0 | 0 | 0 | 0 | 2 | 0 |
| 0.53 | N | 0 | 1 | 0 | 2 | 0 | 0 | 2 | 0 |
| 0.83 | N | 0 | 0 | 0 | 0 | 0 | 0 | 2 | 0 |
| 0.51 | N | 0 | 1 | 0 | 2 | 0 | 0 | 0 | 0 |
| 0.54 | Y | 0 | 0 | 0 | 2 | 0 | 0 | 2 | 0 |
| 0.44 | Y | 0 | 0 | 0 | 2 | 0 | 0 | 2 | 0 |
| 0.71 | N | 0 | 0 | 0 | 2 | 0 | 0 | 2 | 0 |
| 0.70 | N | 0 | 0 | 0 | 1 | 0 | 0 | 2 | 0 |

| | | | | | | | | | |
|---|---|---|---|---|---|---|---|---|---|
| 0.47 | N | 0 | 1 | 0 | 0 | 0 | 0 | 2 | 0 |
| 0.62 | N | 0 | 2 | 0 | 1 | 0 | 0 | 2 | 0 |
| 0.59 | Y | 0 | 0 | 0 | 2 | 0 | 0 | 2 | 0 |
| 0.60 | Y | 0 | 2 | 0 | 1 | 0 | 0 | 2 | 0 |
| 0.58 | N | 0 | 2 | 0 | 2 | 0 | 0 | 2 | 0 |
| 0.54 | Y | 0 | 0 | 0 | 2 | 0 | 0 | 2 | 0 |
| 0.61 | Y | 0 | 0 | 0 | 2 | 0 | 0 | 2 | 0 |
| 0.67 | Y | 0 | 0 | 0 | 2 | 0 | 0 | 2 | 0 |
| 0.88 | N | 0 | 1 | 0 | 2 | 0 | 0 | 2 | 0 |
| 0.60 | Y | 0 | 0 | 0 | 2 | 0 | 0 | 2 | 0 |
| 0.50 | N | 0 | 1 | 0 | 1 | 0 | 0 | 2 | 0 |
| 0.55 | N | 0 | 1 | 0 | 2 | 0 | 0 | 2 | 0 |
| 0.65 | Y | 0 | 1 | 0 | 1 | 0 | 0 | 2 | 0 |
| 0.58 | Y | 0 | 1 | 0 | 2 | 0 | 0 | 2 | 0 |
| 0.55 | N | 0 | 0 | 0 | 2 | 0 | 0 | 2 | 0 |
| 0.63 | N | 0 | 0 | 0 | 2 | 0 | 1 | 2 | 0 |
| 0.49 | Y | 0 | 0 | 0 | 2 | 0 | 0 | 2 | 0 |
| 0.65 | N | 0 | 2 | 0 | 2 | 0 | 0 | 2 | 0 |
| 0.55 | Y | 0 | 0 | 0 | 2 | 0 | 0 | 2 | 0 |
| 0.78 | Y | 0 | 2 | 0 | 2 | 0 | 0 | 2 | 0 |
| 0.60 | N | 0 | 1 | 0 | 2 | 0 | 0 | 2 | 0 |
| 0.58 | Y | 0 | 0 | 0 | 2 | 0 | 0 | 2 | 0 |
| 0.80 | Y | 0 | 2 | 0 | 2 | 0 | 0 | 2 | 0 |

| GM-Femur D. | GM-Tibia F. | GM-Tibia D. | GM-Calcaneous F | GM-Calcaneous D | GM-Total Fractures | GM-Total Deformities | GM-Total Score | GM-Pelvis Ratio | GM-Scoliosis |
|---|---|---|---|---|---|---|---|---|---|
| 0 | 0 | 0 | 0 | 0 | 0 | 0 | 0 | 1.00 | N |
| 0 | 0 | 0 | 0 | 0 | 0 | 0 | 0 | 1.00 | N |
| 0 | 0 | 0 | 0 | 0 | 0 | 0 | 0 | 1.00 | N |
| 0 | 0 | 0 | 0 | 0 | 0 | 0 | 0 | 1.00 | N |
| 0 | 0 | 0 | 0 | 0 | 0 | 0 | 0 | 1.00 | N |
| 0 | 0 | 0 | 0 | 0 | 0 | 0 | 0 | 1.00 | N |
| 0 | 0 | 0 | 0 | 0 | 0 | 0 | 0 | 1.00 | N |
| 0 | 0 | 0 | 0 | 0 | 0 | 0 | 0 | 1.00 | N |
| 0 | 0 | 0 | 0 | 0 | 0 | 0 | 0 | 1.00 | N |
| 0 | 0 | 0 | 0 | 0 | 0 | 0 | 0 | 1.00 | N |
| 0 | 0 | 0 | 0 | 0 | 0 | 0 | 0 | 1.00 | N |
| 0 | 0 | 0 | 0 | 0 | 0 | 0 | 0 | 1.00 | N |
| 0 | 0 | 0 | 0 | 0 | 0 | 0 | 0 | 1.00 | N |
| 0 | 0 | 0 | 0 | 0 | 0 | 0 | 0 | 1.00 | N |
| 0 | 0 | 0 | 0 | 0 | 0 | 0 | 0 | 1.00 | N |
| 0 | 0 | 0 | 0 | 0 | 0 | 0 | 0 | 1.00 | N |
| 0 | 0 | 0 | 0 | 0 | 0 | 0 | 0 | 1.00 | N |
| 0 | 0 | 0 | 0 | 0 | 0 | 0 | 0 | 1.00 | N |
| 0 | 0 | 0 | 0 | 0 | 0 | 0 | 0 | 1.00 | N |
| 0 | 0 | 0 | 0 | 0 | 0 | 0 | 0 | 1.00 | N |
| 0 | 0 | 0 | 0 | 0 | 0 | 0 | 0 | 1.00 | N |
| 0 | 0 | 0 | 0 | 0 | 0 | 0 | 0 | 1.00 | N |
| 0 | 0 | 0 | 0 | 0 | 0 | 0 | 0 | 1.00 | N |
| 0 | 0 | 0 | 0 | 0 | 0 | 0 | 0 | 1.00 | N |
| 0 | 0 | 0 | 0 | 0 | 0 | 0 | 0 | 1.00 | N |
| 0 | 0 | 0 | 0 | 0 | 0 | 0 | 0 | 1.00 | N |
| 0 | 0 | 0 | 0 | 0 | 0 | 0 | 0 | 1.00 | N |
| 0 | 0 | 0 | 0 | 0 | 0 | 0 | 0 | 1.00 | N |
| 0 | 0 | 0 | 0 | 0 | 0 | 0 | 0 | 1.00 | N |

| | | | | | | | | | |
|---|---|---|---|---|---|---|---|---|---|
| 0 | 0 | 0 | 0 | 0 | 0 | 0 | 0 | 1.00 | N |
| 0 | 0 | 0 | 0 | 0 | 0 | 0 | 0 | 1.00 | N |
| 0 | 0 | 0 | 0 | 0 | 0 | 0 | 0 | 1.00 | N |
| 0 | 0 | 0 | 0 | 0 | 0 | 0 | 0 | 1.00 | N |
| 0 | 0 | 0 | 0 | 0 | 0 | 0 | 0 | 1.00 | N |
| 0 | 0 | 0 | 0 | 0 | 0 | 0 | 0 | 1.00 | N |
| 0 | 0 | 0 | 0 | 0 | 0 | 0 | 0 | 1.00 | N |
| 0 | 0 | 0 | 0 | 0 | 0 | 0 | 0 | 1.00 | N |
| 0 | 0 | 0 | 0 | 0 | 0 | 0 | 0 | 1.00 | N |
| 0 | 0 | 0 | 0 | 0 | 0 | 0 | 0 | 1.00 | N |
| 0 | 0 | 0 | 0 | 0 | 0 | 0 | 0 | 1.00 | N |
| 0 | 0 | 0 | 0 | 0 | 0 | 0 | 0 | 1.00 | N |
| 0 | 0 | 0 | 0 | 0 | 0 | 0 | 0 | 1.00 | N |
| 0 | 0 | 0 | 0 | 0 | 0 | 0 | 0 | 1.00 | N |
| 0 | 0 | 0 | 0 | 0 | 0 | 0 | 0 | 1.00 | N |
| 0 | 0 | 0 | 0 | 0 | 0 | 0 | 0 | 1.00 | N |
| 0 | 0 | 0 | 0 | 0 | 0 | 0 | 0 | 1.00 | N |
| 0 | 0 | 0 | 0 | 0 | 0 | 0 | 0 | 1.00 | N |
| 0 | 0 | 0 | 0 | 0 | 0 | 0 | 0 | 1.00 | N |
| 0 | 0 | 0 | 0 | 0 | 0 | 0 | 0 | 1.00 | N |
| 0 | 0 | 0 | 0 | 0 | 0 | 0 | 0 | 1.00 | N |
| 0 | 0 | 0 | 0 | 0 | 0 | 0 | 0 | 1.00 | N |
| 0 | 0 | 0 | 0 | 0 | 0 | 0 | 0 | 1.00 | N |
| 0 | 0 | 0 | 0 | 0 | 0 | 0 | 0 | 1.00 | N |
| 0 | 0 | 0 | 0 | 0 | 0 | 0 | 0 | 1.00 | N |
| 0 | 0 | 0 | 0 | 0 | 0 | 0 | 0 | 1.00 | N |
| 0 | 0 | 0 | 0 | 0 | 0 | 0 | 0 | 1.00 | N |
| 0 | 0 | 0 | 0 | 0 | 0 | 0 | 0 | 1.00 | N |
| 0 | 0 | 0 | 0 | 0 | 0 | 0 | 0 | 1.00 | N |
| 0 | 0 | 0 | 0 | 0 | 0 | 0 | 0 | 1.00 | N |

| | | | | | | | | | |
|---|---|---|---|---|---|---|---|---|---|
| 0 | 0 | 0 | 0 | 0 | 0 | 0 | 0 | 1.00 | N |
| 0 | 0 | 0 | 0 | 0 | 0 | 0 | 0 | 1.00 | N |
| 0 | 0 | 0 | 0 | 0 | 0 | 0 | 0 | 1.00 | N |
| 0 | 0 | 0 | 0 | 0 | 0 | 0 | 0 | 1.00 | N |
| 0 | 0 | 0 | 0 | 0 | 0 | 0 | 0 | 1.00 | N |
| 0 | 0 | 0 | 0 | 0 | 0 | 0 | 0 | 1.00 | N |
| 0 | 0 | 0 | 0 | 0 | 0 | 0 | 0 | 1.00 | N |
| 0 | 0 | 0 | 0 | 0 | 0 | 0 | 0 | 1.00 | N |
| 0 | 0 | 0 | 0 | 0 | 0 | 0 | 0 | 1.00 | N |
| 0 | 0 | 0 | 0 | 0 | 0 | 0 | 0 | 1.00 | N |
| 0 | 0 | 2 | 0 | 2 | 0 | 5 | 5 | 0.90 | N |
| 0 | 0 | 0 | 0 | 2 | 1 | 5 | 6 | 0.63 | N |
| 1 | 0 | 0 | 0 | 1 | 0 | 6 | 6 | 0.48 | Y |
| 0 | 0 | 1 | 0 | 2 | 0 | 8 | 8 | 0.73 | N |
| 0 | 0 | 2 | 0 | 2 | 0 | 4 | 4 | 0.85 | N |
| 0 | 0 | 0 | 0 | 2 | 0 | 7 | 7 | 0.55 | Y |
| 0 | 0 | 0 | 0 | 2 | 1 | 8 | 9 | 0.57 | N |
| 0 | 0 | 2 | 0 | 2 | 0 | 9 | 9 | 0.60 | Y |
| 0 | 0 | 0 | 0 | 2 | 0 | 9 | 9 | 0.54 | Y |
| 0 | 0 | 0 | 0 | 2 | 0 | 8 | 8 | 0.48 | Y |
| 0 | 0 | 1 | 0 | 2 | 0 | 9 | 9 | 0.53 | Y |
| 0 | 0 | 1 | 0 | 2 | 1 | 8 | 9 | 0.53 | Y |
| 0 | 0 | 1 | 0 | 2 | 1 | 8 | 9 | 0.54 | Y |
| 0 | 0 | 2 | 0 | 1 | 1 | 11 | 12 | 0.61 | Y |
| 0 | 0 | 2 | 0 | 2 | 0 | 7 | 7 | 0.84 | N |
| 0 | 0 | 0 | 0 | 2 | 0 | 6 | 6 | 0.60 | N |
| 0 | 0 | 2 | 0 | 2 | 0 | 9 | 9 | 0.53 | Y |
| 0 | 0 | 2 | 0 | 2 | 1 | 10 | 11 | 0.50 | Y |
| 0 | 0 | 0 | 0 | 2 | 0 | 6 | 6 | 0.43 | N |
| 0 | 0 | 0 | 0 | 2 | 1 | 5 | 6 | 0.64 | N |

| | | | | | | | | | |
|---|---|---|---|---|---|---|---|---|---|
| 0 | 0 | 3 | 0 | 2 | 0 | 11 | 11 | 0.61 | Y |
| 1 | 0 | 2 | 0 | 2 | 0 | 11 | 11 | 0.57 | Y |
| 1 | 0 | 3 | 0 | 2 | 0 | 13 | 13 | 0.64 | N |
| 0 | 0 | 1 | 0 | 2 | 0 | 7 | 7 | 0.65 | Y |
| 0 | 0 | 1 | 1 | 1 | 2 | 6 | 8 | 0.77 | N |
| 1 | 0 | 0 | 0 | 2 | 0 | 7 | 7 | 0.45 | Y |
| 0 | 0 | 1 | 0 | 1 | 0 | 6 | 6 | 0.57 | Y |
| 0 | 0 | 0 | 0 | 2 | 0 | 7 | 7 | 0.58 | Y |
| 0 | 0 | 0 | 1 | 1 | 1 | 7 | 8 | 0.61 | N |
| 0 | 0 | 0 | 0 | 2 | 0 | 6 | 6 | 0.56 | Y |
| 0 | 0 | 2 | 0 | 2 | 0 | 9 | 9 | 0.59 | Y |
| 1 | 0 | 0 | 0 | 2 | 0 | 8 | 8 | 0.52 | N |
| 0 | 0 | 0 | 0 | 2 | 0 | 8 | 8 | 0.56 | Y |
| 0 | 0 | 0 | 1 | 2 | 1 | 7 | 8 | 0.55 | Y |
| 0 | 1 | 0 | 0 | 2 | 1 | 7 | 8 | 0.60 | Y |
| 0 | 0 | 0 | 1 | 1 | 1 | 6 | 7 | 0.53 | Y |
| 0 | 0 | 1 | 0 | 1 | 0 | 8 | 8 | 0.43 | Y |
| 0 | 0 | 1 | 0 | 2 | 0 | 9 | 9 | 0.57 | Y |
| 0 | 0 | 2 | 0 | 2 | 0 | 10 | 10 | 0.45 | Y |
| 0 | 0 | 1 | 0 | 2 | 1 | 8 | 9 | 0.56 | Y |
| 0 | 1 | 1 | 0 | 2 | 2 | 7 | 9 | 0.49 | Y |
| 0 | 0 | 0 | 0 | 2 | 1 | 8 | 9 | 0.46 | Y |
| 0 | 0 | 2 | 0 | 2 | 1 | 11 | 12 | 0.49 | Y |
| 0 | 0 | 2 | 1 | 1 | 1 | 9 | 10 | 0.62 | Y |
| 0 | 0 | 2 | 0 | 1 | 0 | 9 | 9 | 0.61 | Y |
| 0 | 0 | 2 | 0 | 2 | 2 | 9 | 11 | 0.46 | Y |
| 0 | 0 | 2 | 0 | 0 | 1 | 6 | 7 | 0.51 | Y |
| 0 | 0 | 2 | 0 | 2 | 0 | 9 | 9 | 0.56 | Y |
| 0 | 0 | 2 | 0 | 2 | 0 | 7 | 7 | 0.58 | Y |
| 0 | 0 | 2 | 0 | 2 | 0 | 10 | 10 | 0.48 | Y |

| | | | | | | | | | |
|---|---|---|---|---|---|---|---|---|---|
| 0 | 0 | 3 | 0 | 2 | 1 | 9 | 10 | 0.62 | Y |
| 0 | 0 | 2 | 0 | 2 | 0 | 9 | 9 | 0.62 | Y |
| 0 | 0 | 2 | 0 | 2 | 2 | 9 | 11 | 0.53 | Y |
| 1 | 0 | 4 | 0 | 2 | 0 | 13 | 13 | 0.63 | N |
| 1 | 0 | 2 | 0 | 2 | 0 | 12 | 12 | 0.61 | N |
| 0 | 0 | 0 | 0 | 1 | 0 | 1 | 1 | 0.90 | N |
| 0 | 0 | 0 | 0 | 2 | 0 | 4 | 4 | 0.88 | N |
| 0 | 0 | 0 | 0 | 1 | 0 | 3 | 3 | 0.57 | N |
| 0 | 0 | 0 | 0 | 1 | 0 | 4 | 4 | 0.82 | N |
| 0 | 0 | 0 | 0 | 2 | 0 | 4 | 4 | 0.68 | N |
| 0 | 0 | 0 | 0 | 2 | 0 | 4 | 4 | 0.92 | N |
| 0 | 0 | 0 | 0 | 2 | 0 | 7 | 7 | 1.00 | N |
| 0 | 0 | 0 | 0 | 2 | 0 | 3 | 3 | 0.81 | N |
| 0 | 0 | 0 | 0 | 2 | 0 | 4 | 4 | 0.93 | N |
| 0 | 1 | 0 | 1 | 1 | 2 | 3 | 5 | 1.00 | N |
| 1 | 0 | 0 | 0 | 1 | 0 | 4 | 4 | 0.59 | N |
| 0 | 0 | 0 | 0 | 2 | 0 | 6 | 6 | 0.93 | N |
| 0 | 0 | 0 | 0 | 2 | 0 | 4 | 4 | 0.68 | N |
| 0 | 0 | 0 | 1 | 0 | 1 | 4 | 5 | 0.68 | N |
| 0 | 0 | 0 | 0 | 2 | 0 | 4 | 4 | 0.88 | N |
| 0 | 0 | 1 | 0 | 1 | 0 | 6 | 6 | 0.61 | N |
| 0 | 0 | 0 | 0 | 2 | 0 | 6 | 6 | 0.66 | N |
| 0 | 0 | 0 | 0 | 1 | 0 | 6 | 6 | 0.61 | Y |
| 0 | 0 | 0 | 0 | 2 | 1 | 4 | 5 | 0.65 | N |
| 0 | 0 | 0 | 0 | 2 | 0 | 5 | 5 | 0.53 | N |
| 0 | 0 | 0 | 0 | 2 | 0 | 5 | 5 | 0.50 | N |
| 0 | 0 | 0 | 1 | 0 | 1 | 3 | 4 | 0.54 | N |
| 0 | 0 | 2 | 0 | 2 | 0 | 9 | 9 | 0.54 | Y |
| 0 | 0 | 1 | 0 | 2 | 0 | 7 | 7 | 0.58 | N |
| 0 | 0 | 0 | 0 | 2 | 0 | 6 | 6 | 0.66 | N |

| | | | | | | | | | |
|---|---|---|---|---|---|---|---|---|---|
| 0 | 0 | 0 | 0 | 2 | 0 | 6 | 6 | 0.62 | N |
| 0 | 0 | 0 | 1 | 1 | 2 | 4 | 6 | 0.61 | N |
| 0 | 0 | 0 | 2 | 0 | 2 | 4 | 6 | 0.60 | N |
| 0 | 0 | 2 | 0 | 2 | 0 | 8 | 8 | 0.59 | Y |
| 0 | 0 | 0 | 0 | 2 | 0 | 6 | 6 | 0.58 | N |
| 0 | 0 | 1 | 0 | 2 | 0 | 6 | 6 | 0.67 | Y |
| 0 | 0 | 0 | 0 | 1 | 0 | 2 | 2 | 0.85 | N |
| 0 | 0 | 0 | 0 | 2 | 0 | 6 | 6 | 0.55 | N |
| 0 | 0 | 0 | 0 | 2 | 0 | 2 | 2 | 0.93 | N |
| 0 | 0 | 0 | 0 | 2 | 0 | 4 | 4 | 0.73 | N |
| 0 | 0 | 2 | 0 | 2 | 0 | 8 | 8 | 0.58 | Y |
| 0 | 0 | 0 | 1 | 1 | 1 | 5 | 6 | 0.56 | Y |
| 0 | 0 | 0 | 0 | 2 | 0 | 4 | 4 | 0.93 | N |
| 0 | 0 | 0 | 0 | 2 | 1 | 6 | 7 | 0.67 | Y |
| 0 | 0 | 0 | 0 | 1 | 0 | 4 | 4 | 0.61 | N |
| 0 | 0 | 0 | 0 | 2 | 0 | 7 | 7 | 0.60 | Y |
| 0 | 0 | 2 | 0 | 1 | 0 | 7 | 7 | 0.59 | N |
| 0 | 0 | 2 | 1 | 1 | 1 | 7 | 8 | 0.54 | N |
| 0 | 0 | 1 | 0 | 1 | 0 | 6 | 6 | 0.60 | N |
| 0 | 0 | 2 | 0 | 2 | 0 | 8 | 8 | 0.50 | N |
| 0 | 0 | 2 | 0 | 2 | 0 | 8 | 8 | 0.54 | N |
| 0 | 0 | 1 | 0 | 2 | 0 | 7 | 7 | 1.10 | N |
| 0 | 0 | 0 | 0 | 2 | 0 | 4 | 4 | 0.69 | Y |
| 0 | 0 | 2 | 0 | 2 | 0 | 9 | 9 | 0.55 | N |
| 0 | 0 | 0 | 0 | 2 | 0 | 4 | 4 | 0.93 | N |
| 0 | 0 | 2 | 0 | 2 | 0 | 7 | 7 | 0.53 | N |
| 0 | 0 | 2 | 0 | 2 | 0 | 8 | 8 | 0.57 | Y |
| 0 | 0 | 2 | 0 | 2 | 0 | 8 | 8 | 0.60 | Y |
| 0 | 0 | 0 | 0 | 1 | 0 | 5 | 5 | 0.68 | N |
| 1 | 0 | 0 | 0 | 1 | 0 | 5 | 5 | 0.60 | N |

| | | | | | | | | | |
|---|---|---|---|---|---|---|---|---|---|
| 0 | 0 | 0 | 0 | 2 | 0 | 5 | 5 | 0.50 | N |
| 0 | 0 | 0 | 0 | 2 | 0 | 7 | 7 | 0.62 | N |
| 0 | 0 | 2 | 0 | 2 | 0 | 8 | 8 | 0.62 | Y |
| 0 | 0 | 1 | 0 | 2 | 0 | 8 | 8 | 0.57 | Y |
| 0 | 0 | 0 | 0 | 2 | 0 | 8 | 8 | 0.59 | N |
| 0 | 0 | 2 | 0 | 2 | 0 | 8 | 8 | 0.56 | Y |
| 0 | 0 | 2 | 0 | 2 | 0 | 8 | 8 | 0.56 | Y |
| 0 | 0 | 2 | 0 | 2 | 0 | 8 | 8 | 0.52 | Y |
| 0 | 0 | 1 | 1 | 1 | 1 | 7 | 8 | 0.92 | N |
| 1 | 0 | 0 | 0 | 2 | 0 | 7 | 7 | 0.57 | Y |
| 0 | 0 | 0 | 0 | 2 | 0 | 6 | 6 | 0.60 | N |
| 0 | 0 | 2 | 0 | 1 | 0 | 8 | 8 | 0.60 | N |
| 0 | 0 | 1 | 0 | 2 | 0 | 7 | 7 | 0.63 | Y |
| 0 | 0 | 0 | 0 | 2 | 0 | 7 | 7 | 0.62 | Y |
| 0 | 0 | 2 | 0 | 2 | 0 | 8 | 8 | 0.57 | N |
| 0 | 0 | 2 | 0 | 1 | 0 | 8 | 8 | 0.63 | N |
| 0 | 0 | 2 | 0 | 2 | 0 | 8 | 8 | 0.60 | Y |
| 0 | 0 | 0 | 0 | 2 | 0 | 8 | 8 | 0.60 | N |
| 0 | 0 | 2 | 0 | 2 | 0 | 8 | 8 | 0.53 | Y |
| 0 | 0 | 1 | 0 | 2 | 0 | 9 | 9 | 0.84 | N |
| 0 | 0 | 2 | 0 | 2 | 0 | 9 | 9 | 0.57 | N |
| 0 | 0 | 2 | 0 | 2 | 0 | 8 | 8 | 0.61 | Y |
| 0 | 0 | 1 | 0 | 2 | 0 | 9 | 9 | 0.84 | N |

| IH-Humerus F | IH-Humerus D | IH-Forearm (Ulna & | IH-Forearm (Ulna & | IH-Olecranon F | IH-Olecranon D | IH-Ischium | IH-Femur F. | IH-Femur D. | IH-Tibia F. |
|---|---|---|---|---|---|---|---|---|---|
| 0 | 0 | 0 | 0 | 0 | 0 | 0 | 0 | 0 | 0 |
| 0 | 0 | 0 | 0 | 0 | 0 | 0 | 0 | 0 | 0 |
| 0 | 0 | 0 | 0 | 0 | 0 | 0 | 0 | 0 | 0 |
| 0 | 0 | 0 | 0 | 0 | 0 | 0 | 0 | 0 | 0 |
| 0 | 0 | 0 | 0 | 0 | 0 | 0 | 0 | 0 | 0 |
| 0 | 0 | 0 | 0 | 0 | 0 | 0 | 0 | 0 | 0 |
| 0 | 0 | 0 | 0 | 0 | 0 | 0 | 0 | 0 | 0 |
| 0 | 0 | 0 | 0 | 0 | 0 | 0 | 0 | 0 | 0 |
| 0 | 0 | 0 | 0 | 0 | 0 | 0 | 0 | 0 | 0 |
| 0 | 0 | 0 | 0 | 0 | 0 | 0 | 0 | 0 | 0 |
| 0 | 0 | 0 | 0 | 0 | 0 | 0 | 0 | 0 | 0 |
| 0 | 0 | 0 | 0 | 0 | 0 | 0 | 0 | 0 | 0 |
| 0 | 0 | 0 | 0 | 0 | 0 | 0 | 0 | 0 | 0 |
| 0 | 0 | 0 | 0 | 0 | 0 | 0 | 0 | 0 | 0 |
| 0 | 0 | 0 | 0 | 0 | 0 | 0 | 0 | 0 | 0 |
| 0 | 0 | 0 | 0 | 0 | 0 | 0 | 0 | 0 | 0 |
| 0 | 0 | 0 | 0 | 0 | 0 | 0 | 0 | 0 | 0 |
| 0 | 0 | 0 | 0 | 0 | 0 | 0 | 0 | 0 | 0 |
| 0 | 0 | 0 | 0 | 0 | 0 | 0 | 0 | 0 | 0 |
| 0 | 0 | 0 | 0 | 0 | 0 | 0 | 0 | 0 | 0 |
| 0 | 0 | 0 | 0 | 0 | 0 | 0 | 0 | 0 | 0 |
| 0 | 0 | 0 | 0 | 0 | 0 | 0 | 0 | 0 | 0 |
| 0 | 0 | 0 | 0 | 0 | 0 | 0 | 0 | 0 | 0 |
| 0 | 0 | 0 | 0 | 0 | 0 | 0 | 0 | 0 | 0 |
| 0 | 0 | 0 | 0 | 0 | 0 | 0 | 0 | 0 | 0 |
| 0 | 0 | 0 | 0 | 0 | 0 | 0 | 0 | 0 | 0 |
| 0 | 0 | 0 | 0 | 0 | 0 | 0 | 0 | 0 | 0 |
| 0 | 0 | 0 | 0 | 0 | 0 | 0 | 0 | 0 | 0 |
| 0 | 0 | 0 | 0 | 0 | 0 | 0 | 0 | 0 | 0 |

| | | | | | | | | | |
|---|---|---|---|---|---|---|---|---|---|
| 0 | 0 | 0 | 0 | 0 | 0 | 0 | 0 | 0 | 0 |
| 0 | 0 | 0 | 0 | 0 | 0 | 0 | 0 | 0 | 0 |
| 0 | 0 | 0 | 0 | 0 | 0 | 0 | 0 | 0 | 0 |
| 0 | 0 | 0 | 0 | 0 | 0 | 0 | 0 | 0 | 0 |
| 0 | 0 | 0 | 0 | 0 | 0 | 0 | 0 | 0 | 0 |
| 0 | 0 | 0 | 0 | 0 | 0 | 0 | 0 | 0 | 0 |
| 0 | 0 | 0 | 0 | 0 | 0 | 0 | 0 | 0 | 0 |
| 0 | 0 | 0 | 0 | 0 | 0 | 0 | 0 | 0 | 0 |
| 0 | 0 | 0 | 0 | 0 | 0 | 0 | 0 | 0 | 0 |
| 0 | 0 | 0 | 0 | 0 | 0 | 0 | 0 | 0 | 0 |
| 0 | 0 | 0 | 0 | 0 | 0 | 0 | 0 | 0 | 0 |
| 0 | 0 | 0 | 0 | 0 | 0 | 0 | 0 | 0 | 0 |
| 0 | 0 | 0 | 0 | 0 | 0 | 0 | 0 | 0 | 0 |
| 0 | 0 | 0 | 0 | 0 | 0 | 0 | 0 | 0 | 0 |
| 0 | 0 | 0 | 0 | 0 | 0 | 0 | 0 | 0 | 0 |
| 0 | 0 | 0 | 0 | 0 | 0 | 0 | 0 | 0 | 0 |
| 0 | 0 | 0 | 0 | 0 | 0 | 0 | 0 | 0 | 0 |
| 0 | 0 | 0 | 0 | 0 | 0 | 0 | 0 | 0 | 0 |
| 0 | 0 | 0 | 0 | 0 | 0 | 0 | 0 | 0 | 0 |
| 0 | 0 | 0 | 0 | 0 | 0 | 0 | 0 | 0 | 0 |
| 0 | 0 | 0 | 0 | 0 | 0 | 0 | 0 | 0 | 0 |
| 0 | 0 | 0 | 0 | 0 | 0 | 0 | 0 | 0 | 0 |
| 0 | 0 | 0 | 0 | 0 | 0 | 0 | 0 | 0 | 0 |
| 0 | 0 | 0 | 0 | 0 | 0 | 0 | 0 | 0 | 0 |
| 0 | 0 | 0 | 0 | 0 | 0 | 0 | 0 | 0 | 0 |
| 0 | 0 | 0 | 0 | 0 | 0 | 0 | 0 | 0 | 0 |
| 0 | 0 | 0 | 0 | 0 | 0 | 0 | 0 | 0 | 0 |
| 0 | 0 | 0 | 0 | 0 | 0 | 0 | 0 | 0 | 0 |
| 0 | 0 | 0 | 0 | 0 | 0 | 0 | 0 | 0 | 0 |
| 0 | 0 | 0 | 0 | 0 | 0 | 0 | 0 | 0 | 0 |

| | | | | | | | | | |
|---|---|---|---|---|---|---|---|---|---|
| 0 | 0 | 0 | 0 | 0 | 0 | 0 | 0 | 0 | 0 |
| 0 | 0 | 0 | 0 | 0 | 0 | 0 | 0 | 0 | 0 |
| 0 | 0 | 0 | 0 | 0 | 0 | 0 | 0 | 0 | 0 |
| 0 | 0 | 0 | 0 | 0 | 0 | 0 | 0 | 0 | 0 |
| 0 | 0 | 0 | 0 | 0 | 0 | 0 | 0 | 0 | 0 |
| 0 | 0 | 0 | 0 | 0 | 0 | 0 | 0 | 0 | 0 |
| 0 | 0 | 0 | 0 | 0 | 0 | 0 | 0 | 0 | 0 |
| 0 | 0 | 0 | 0 | 0 | 0 | 0 | 0 | 0 | 0 |
| 0 | 0 | 0 | 0 | 0 | 0 | 0 | 0 | 0 | 0 |
| 0 | 0 | 0 | 0 | 0 | 0 | 0 | 0 | 0 | 0 |
| 0 | 0 | 0 | 0 | 0 | 0 | 1 | 0 | 0 | 0 |
| 1 | 0 | 0 | 1 | 0 | 0 | 2 | 0 | 0 | 0 |
| 0 | 0 | 0 | 2 | 0 | 0 | 2 | 0 | 0 | 0 |
| 0 | 0 | 0 | 0 | 0 | 1 | 2 | 0 | 0 | 0 |
| 0 | 0 | 0 | 0 | 0 | 0 | 1 | 0 | 0 | 0 |
| 0 | 1 | 0 | 2 | 0 | 1 | 2 | 0 | 0 | 0 |
| 0 | 0 | 0 | 0 | 0 | 0 | 2 | 0 | 0 | 0 |
| 1 | 0 | 0 | 2 | 0 | 0 | 2 | 0 | 0 | 0 |
| 2 | 0 | 0 | 1 | 0 | 1 | 2 | 0 | 0 | 0 |
| 0 | 2 | 0 | 1 | 0 | 0 | 2 | 0 | 0 | 0 |
| 0 | 1 | 0 | 1 | 0 | 0 | 2 | 0 | 0 | 0 |
| 1 | 1 | 0 | 1 | 0 | 0 | 2 | 0 | 0 | 1 |
| 0 | 2 | 0 | 2 | 0 | 0 | 0 | 0 | 0 | 0 |
| 2 | 0 | 0 | 1 | 0 | 1 | 2 | 0 | 0 | 1 |
| 0 | 0 | 0 | 0 | 0 | 0 | 1 | 0 | 0 | 0 |
| 0 | 1 | 0 | 1 | 0 | 0 | 2 | 0 | 0 | 0 |
| 0 | 1 | 0 | 1 | 0 | 1 | 2 | 0 | 0 | 0 |
| 0 | 2 | 1 | 0 | 0 | 0 | 2 | 0 | 0 | 0 |
| 0 | 0 | 0 | 2 | 0 | 0 | 2 | 0 | 0 | 0 |
| 1 | 0 | 0 | 1 | 0 | 0 | 2 | 0 | 0 | 0 |

| | | | | | | | | | |
|---|---|---|---|---|---|---|---|---|---|
| 0 | 1 | 0 | 2 | 0 | 0 | 2 | 0 | 0 | 0 |
| 0 | 2 | 0 | 1 | 0 | 0 | 2 | 0 | 0 | 1 |
| 1 | 1 | 1 | 1 | 0 | 0 | 2 | 0 | 1 | 0 |
| 0 | 0 | 0 | 0 | 0 | 0 | 2 | 0 | 0 | 0 |
| 0 | 0 | 0 | 0 | 0 | 0 | 2 | 0 | 0 | 0 |
| 0 | 0 | 0 | 1 | 0 | 0 | 2 | 0 | 0 | 0 |
| 0 | 0 | 0 | 2 | 0 | 0 | 2 | 0 | 0 | 0 |
| 0 | 1 | 0 | 1 | 0 | 1 | 2 | 0 | 0 | 0 |
| 0 | 0 | 0 | 2 | 0 | 0 | 2 | 0 | 0 | 0 |
| 0 | 0 | 0 | 2 | 0 | 1 | 2 | 0 | 0 | 0 |
| 1 | 0 | 0 | 1 | 1 | 0 | 2 | 0 | 0 | 0 |
| 0 | 0 | 0 | 2 | 0 | 0 | 2 | 0 | 0 | 0 |
| 0 | 1 | 0 | 1 | 0 | 0 | 2 | 0 | 0 | 0 |
| 1 | 0 | 0 | 1 | 0 | 1 | 2 | 0 | 0 | 0 |
| 0 | 0 | 0 | 1 | 0 | 0 | 2 | 0 | 0 | 1 |
| 0 | 1 | 0 | 1 | 0 | 1 | 2 | 0 | 0 | 0 |
| 1 | 1 | 0 | 2 | 0 | 1 | 0 | 0 | 0 | 0 |
| 1 | 1 | 0 | 2 | 0 | 0 | 2 | 0 | 0 | 0 |
| 0 | 1 | 0 | 1 | 0 | 0 | 2 | 0 | 0 | 0 |
| 1 | 1 | 0 | 1 | 0 | 0 | 2 | 0 | 0 | 0 |
| 1 | 0 | 0 | 1 | 0 | 0 | 2 | 0 | 0 | 1 |
| 0 | 2 | 0 | 2 | 0 | 0 | 2 | 0 | 0 | 0 |
| 0 | 2 | 0 | 1 | 0 | 0 | 2 | 0 | 0 | 1 |
| 0 | 2 | 0 | 0 | 0 | 0 | 2 | 0 | 0 | 1 |
| 1 | 1 | 0 | 1 | 0 | 1 | 2 | 0 | 0 | 1 |
| 0 | 2 | 0 | 1 | 0 | 1 | 2 | 0 | 0 | 0 |
| 1 | 0 | 0 | 1 | 0 | 0 | 2 | 0 | 0 | 0 |
| 1 | 1 | 0 | 1 | 0 | 0 | 2 | 0 | 0 | 1 |
| 0 | 0 | 0 | 1 | 0 | 1 | 2 | 0 | 0 | 2 |
| 0 | 1 | 0 | 1 | 0 | 1 | 2 | 0 | 0 | 0 |

| | | | | | | | | | |
|---|---|---|---|---|---|---|---|---|---|
| 1 | 0 | 0 | 1 | 0 | 0 | 2 | 0 | 0 | 2 |
| 0 | 1 | 0 | 2 | 0 | 1 | 2 | 0 | 0 | 0 |
| 1 | 0 | 0 | 1 | 0 | 0 | 2 | 0 | 0 | 0 |
| 0 | 2 | 0 | 2 | 0 | 0 | 2 | 0 | 0 | 1 |
| 1 | 1 | 1 | 1 | 0 | 0 | 2 | 1 | 0 | 0 |
| 0 | 0 | 0 | 0 | 0 | 0 | 2 | 0 | 0 | 0 |
| 0 | 0 | 0 | 0 | 0 | 0 | 1 | 0 | 0 | 0 |
| 0 | 0 | 0 | 0 | 0 | 0 | 2 | 0 | 0 | 0 |
| 0 | 0 | 0 | 2 | 0 | 0 | 2 | 0 | 0 | 0 |
| 0 | 0 | 0 | 0 | 0 | 0 | 2 | 0 | 0 | 0 |
| 0 | 0 | 0 | 0 | 0 | 0 | 1 | 0 | 0 | 0 |
| 0 | 1 | 0 | 0 | 0 | 0 | 1 | 0 | 0 | 0 |
| 0 | 0 | 0 | 0 | 0 | 0 | 1 | 0 | 0 | 0 |
| 0 | 0 | 0 | 0 | 0 | 0 | 1 | 0 | 0 | 0 |
| 0 | 0 | 0 | 0 | 0 | 0 | 1 | 0 | 0 | 0 |
| 0 | 0 | 0 | 0 | 0 | 0 | 2 | 0 | 1 | 0 |
| 0 | 1 | 0 | 0 | 0 | 0 | 2 | 0 | 0 | 0 |
| 0 | 0 | 0 | 0 | 0 | 0 | 2 | 0 | 0 | 0 |
| 0 | 0 | 0 | 0 | 0 | 0 | 2 | 0 | 0 | 0 |
| 0 | 0 | 0 | 0 | 0 | 0 | 1 | 0 | 0 | 0 |
| 0 | 0 | 0 | 0 | 0 | 0 | 2 | 0 | 0 | 0 |
| 0 | 0 | 0 | 0 | 0 | 0 | 2 | 0 | 0 | 0 |
| 0 | 0 | 0 | 1 | 0 | 0 | 2 | 0 | 0 | 0 |
| 0 | 0 | 0 | 0 | 0 | 0 | 1 | 1 | 0 | 0 |
| 0 | 1 | 0 | 1 | 0 | 0 | 2 | 0 | 0 | 0 |
| 0 | 1 | 0 | 1 | 0 | 0 | 2 | 0 | 0 | 0 |
| 0 | 0 | 0 | 0 | 0 | 1 | 2 | 0 | 0 | 0 |
| 0 | 0 | 0 | 0 | 0 | 0 | 2 | 0 | 0 | 0 |
| 0 | 0 | 0 | 0 | 0 | 0 | 2 | 0 | 0 | 0 |
| 0 | 0 | 0 | 0 | 0 | 0 | 2 | 0 | 0 | 0 |

| | | | | | | | | | |
|---|---|---|---|---|---|---|---|---|---|
| 0 | 1 | 0 | 0 | 0 | 0 | 2 | 0 | 0 | 0 |
| 1 | 0 | 0 | 0 | 0 | 1 | 2 | 0 | 0 | 0 |
| 0 | 0 | 0 | 2 | 0 | 0 | 2 | 0 | 0 | 0 |
| 0 | 0 | 0 | 0 | 0 | 0 | 2 | 0 | 0 | 1 |
| 0 | 0 | 0 | 2 | 0 | 0 | 2 | 0 | 0 | 0 |
| 0 | 0 | 0 | 1 | 0 | 0 | 2 | 0 | 0 | 0 |
| 0 | 0 | 0 | 0 | 0 | 0 | 2 | 0 | 0 | 0 |
| 0 | 1 | 0 | 1 | 0 | 0 | 2 | 0 | 0 | 0 |
| 0 | 0 | 0 | 0 | 0 | 0 | 2 | 0 | 0 | 0 |
| 0 | 0 | 0 | 0 | 0 | 0 | 2 | 0 | 0 | 0 |
| 0 | 0 | 0 | 0 | 0 | 1 | 2 | 0 | 0 | 0 |
| 0 | 0 | 0 | 2 | 0 | 1 | 2 | 0 | 0 | 0 |
| 0 | 0 | 0 | 0 | 0 | 0 | 1 | 0 | 0 | 0 |
| 1 | 0 | 0 | 2 | 0 | 0 | 2 | 0 | 0 | 0 |
| 0 | 0 | 0 | 0 | 0 | 0 | 2 | 0 | 0 | 0 |
| 0 | 1 | 0 | 2 | 0 | 0 | 2 | 0 | 0 | 0 |
| 0 | 0 | 0 | 2 | 0 | 0 | 2 | 0 | 0 | 0 |
| 0 | 0 | 0 | 2 | 0 | 0 | 2 | 0 | 0 | 0 |
| 0 | 0 | 0 | 0 | 0 | 0 | 2 | 0 | 0 | 0 |
| 0 | 0 | 0 | 2 | 0 | 0 | 2 | 0 | 0 | 0 |
| 0 | 0 | 0 | 2 | 0 | 0 | 2 | 0 | 0 | 0 |
| 1 | 0 | 0 | 0 | 0 | 0 | 1 | 0 | 0 | 0 |
| 0 | 0 | 0 | 0 | 0 | 0 | 2 | 0 | 0 | 0 |
| 0 | 0 | 0 | 2 | 0 | 0 | 2 | 0 | 0 | 0 |
| 0 | 0 | 0 | 0 | 0 | 0 | 1 | 0 | 0 | 0 |
| 0 | 0 | 0 | 2 | 0 | 0 | 2 | 0 | 0 | 0 |
| 0 | 0 | 0 | 2 | 0 | 0 | 2 | 0 | 0 | 0 |
| 0 | 0 | 0 | 2 | 0 | 0 | 2 | 0 | 0 | 0 |
| 0 | 0 | 1 | 0 | 0 | 0 | 2 | 0 | 0 | 0 |
| 0 | 0 | 0 | 0 | 0 | 0 | 2 | 1 | 0 | 0 |

| | | | | | | | | | |
|---|---|---|---|---|---|---|---|---|---|
| 0 | 1 | 0 | 1 | 0 | 0 | 2 | 0 | 0 | 0 |
| 1 | 0 | 0 | 0 | 0 | 0 | 2 | 0 | 0 | 0 |
| 0 | 0 | 0 | 0 | 0 | 0 | 2 | 0 | 0 | 0 |
| 0 | 1 | 0 | 1 | 0 | 0 | 2 | 0 | 0 | 0 |
| 1 | 1 | 0 | 2 | 0 | 1 | 2 | 0 | 0 | 0 |
| 0 | 0 | 0 | 2 | 0 | 0 | 2 | 0 | 0 | 0 |
| 0 | 0 | 0 | 2 | 0 | 0 | 2 | 0 | 0 | 0 |
| 0 | 0 | 0 | 2 | 0 | 1 | 2 | 0 | 0 | 1 |
| 1 | 1 | 0 | 2 | 0 | 2 | 2 | 0 | 0 | 1 |
| 0 | 0 | 0 | 2 | 0 | 0 | 2 | 0 | 0 | 0 |
| 0 | 1 | 0 | 1 | 0 | 0 | 2 | 0 | 0 | 0 |
| 0 | 1 | 0 | 1 | 0 | 1 | 2 | 0 | 0 | 0 |
| 0 | 1 | 0 | 1 | 0 | 0 | 2 | 0 | 0 | 0 |
| 0 | 1 | 0 | 2 | 0 | 0 | 2 | 0 | 0 | 0 |
| 0 | 0 | 0 | 2 | 0 | 0 | 2 | 0 | 0 | 0 |
| 0 | 0 | 0 | 2 | 0 | 1 | 2 | 0 | 0 | 1 |
| 0 | 0 | 0 | 2 | 0 | 0 | 2 | 0 | 0 | 0 |
| 2 | 0 | 0 | 2 | 0 | 0 | 2 | 0 | 0 | 0 |
| 0 | 0 | 0 | 2 | 0 | 1 | 2 | 0 | 0 | 0 |
| 0 | 2 | 0 | 2 | 0 | 2 | 2 | 0 | 0 | 0 |
| 0 | 1 | 0 | 1 | 0 | 0 | 2 | 0 | 0 | 0 |
| 0 | 0 | 0 | 2 | 0 | 0 | 2 | 0 | 0 | 0 |
| 2 | 0 | 0 | 2 | 0 | 2 | 2 | 0 | 0 | 0 |

| IH-Tibia D. | IH-Calcaneus E | IH-Calcaneus D | IH-Total Fractures | IH-Total Deformities | IH-Total Score | IH-Pelvis Ratio | IH-Scoliosis | IB-Humerus E | IB-Humerus D |
|---|---|---|---|---|---|---|---|---|---|
| 0 | 0 | 0 | 0 | 0 | 0 | 1.00 | N | 0 | 0 |
| 0 | 0 | 0 | 0 | 0 | 0 | 1.00 | N | 0 | 0 |
| 0 | 0 | 0 | 0 | 0 | 0 | 1.00 | N | 0 | 0 |
| 0 | 0 | 0 | 0 | 0 | 0 | 1.00 | N | 0 | 0 |
| 0 | 0 | 0 | 0 | 0 | 0 | 1.00 | N | 0 | 0 |
| 0 | 0 | 0 | 0 | 0 | 0 | 1.00 | N | 0 | 0 |
| 0 | 0 | 0 | 0 | 0 | 0 | 1.00 | N | 0 | 0 |
| 0 | 0 | 0 | 0 | 0 | 0 | 1.00 | N | 0 | 0 |
| 0 | 0 | 0 | 0 | 0 | 0 | 1.00 | N | 0 | 0 |
| 0 | 0 | 0 | 0 | 0 | 0 | 1.00 | N | 0 | 0 |
| 0 | 0 | 0 | 0 | 0 | 0 | 1.00 | N | 0 | 0 |
| 0 | 0 | 0 | 0 | 0 | 0 | 1.00 | N | 0 | 0 |
| 0 | 0 | 0 | 0 | 0 | 0 | 1.00 | N | 0 | 0 |
| 0 | 0 | 0 | 0 | 0 | 0 | 1.00 | N | 0 | 0 |
| 0 | 0 | 0 | 0 | 0 | 0 | 1.00 | N | 0 | 0 |
| 0 | 0 | 0 | 0 | 0 | 0 | 1.00 | N | 0 | 0 |
| 0 | 0 | 0 | 0 | 0 | 0 | 1.00 | N | 0 | 0 |
| 0 | 0 | 0 | 0 | 0 | 0 | 1.00 | N | 0 | 0 |
| 0 | 0 | 0 | 0 | 0 | 0 | 1.00 | N | 0 | 0 |
| 0 | 0 | 0 | 0 | 0 | 0 | 1.00 | N | 0 | 0 |
| 0 | 0 | 0 | 0 | 0 | 0 | 1.00 | N | 0 | 0 |
| 0 | 0 | 0 | 0 | 0 | 0 | 1.00 | N | 0 | 0 |
| 0 | 0 | 0 | 0 | 0 | 0 | 1.00 | N | 0 | 0 |
| 0 | 0 | 0 | 0 | 0 | 0 | 1.00 | N | 0 | 0 |
| 0 | 0 | 0 | 0 | 0 | 0 | 1.00 | N | 0 | 0 |
| 0 | 0 | 0 | 0 | 0 | 0 | 1.00 | N | 0 | 0 |
| 0 | 0 | 0 | 0 | 0 | 0 | 1.00 | N | 0 | 0 |
| 0 | 0 | 0 | 0 | 0 | 0 | 1.00 | N | 0 | 0 |
| 0 | 0 | 0 | 0 | 0 | 0 | 1.00 | N | 0 | 0 |

| | | | | | | | | | |
|---|---|---|---|---|---|---|---|---|---|
| 0 | 0 | 0 | 0 | 0 | 0 | 1.00 | N | 0 | 0 |
| 0 | 0 | 0 | 0 | 0 | 0 | 1.00 | N | 0 | 0 |
| 0 | 0 | 0 | 0 | 0 | 0 | 1.00 | N | 0 | 0 |
| 0 | 0 | 0 | 0 | 0 | 0 | 1.00 | N | 0 | 0 |
| 0 | 0 | 0 | 0 | 0 | 0 | 1.00 | N | 0 | 0 |
| 0 | 0 | 0 | 0 | 0 | 0 | 1.00 | N | 0 | 0 |
| 0 | 0 | 0 | 0 | 0 | 0 | 1.00 | N | 0 | 0 |
| 0 | 0 | 0 | 0 | 0 | 0 | 1.00 | N | 0 | 0 |
| 0 | 0 | 0 | 0 | 0 | 0 | 1.00 | N | 0 | 0 |
| 0 | 0 | 0 | 0 | 0 | 0 | 1.00 | N | 0 | 0 |
| 0 | 0 | 0 | 0 | 0 | 0 | 1.00 | N | 0 | 0 |
| 0 | 0 | 0 | 0 | 0 | 0 | 1.00 | N | 0 | 0 |
| 0 | 0 | 0 | 0 | 0 | 0 | 1.00 | N | 0 | 0 |
| 0 | 0 | 0 | 0 | 0 | 0 | 1.00 | N | 0 | 0 |
| 0 | 0 | 0 | 0 | 0 | 0 | 1.00 | N | 0 | 0 |
| 0 | 0 | 0 | 0 | 0 | 0 | 1.00 | N | 0 | 0 |
| 0 | 0 | 0 | 0 | 0 | 0 | 1.00 | N | 0 | 0 |
| 0 | 0 | 0 | 0 | 0 | 0 | 1.00 | N | 0 | 0 |
| 0 | 0 | 0 | 0 | 0 | 0 | 1.00 | N | 0 | 0 |
| 0 | 0 | 0 | 0 | 0 | 0 | 1.00 | N | 0 | 0 |
| 0 | 0 | 0 | 0 | 0 | 0 | 1.00 | N | 0 | 0 |
| 0 | 0 | 0 | 0 | 0 | 0 | 1.00 | N | 0 | 0 |
| 0 | 0 | 0 | 0 | 0 | 0 | 1.00 | N | 0 | 0 |
| 0 | 0 | 0 | 0 | 0 | 0 | 1.00 | N | 0 | 0 |
| 0 | 0 | 0 | 0 | 0 | 0 | 1.00 | N | 0 | 0 |
| 0 | 0 | 0 | 0 | 0 | 0 | 1.00 | N | 0 | 0 |
| 0 | 0 | 0 | 0 | 0 | 0 | 1.00 | N | 0 | 0 |
| 0 | 0 | 0 | 0 | 0 | 0 | 1.00 | N | 0 | 0 |
| 0 | 0 | 0 | 0 | 0 | 0 | 1.00 | N | 0 | 0 |
| 0 | 0 | 0 | 0 | 0 | 0 | 1.00 | N | 0 | 0 |

| | | | | | | | | | |
|---|---|---|---|---|---|---|---|---|---|
| 0 | 0 | 0 | 0 | 0 | 0 | 1.00 | N | 0 | 0 |
| 0 | 0 | 0 | 0 | 0 | 0 | 1.00 | N | 0 | 0 |
| 0 | 0 | 0 | 0 | 0 | 0 | 1.00 | N | 0 | 0 |
| 0 | 0 | 0 | 0 | 0 | 0 | 1.00 | N | 0 | 0 |
| 0 | 0 | 0 | 0 | 0 | 0 | 1.00 | N | 0 | 0 |
| 0 | 0 | 0 | 0 | 0 | 0 | 1.00 | N | 0 | 0 |
| 0 | 0 | 0 | 0 | 0 | 0 | 1.00 | N | 0 | 0 |
| 0 | 0 | 0 | 0 | 0 | 0 | 1.00 | N | 0 | 0 |
| 0 | 0 | 0 | 0 | 0 | 0 | 1.00 | N | 0 | 0 |
| 0 | 0 | 0 | 0 | 0 | 0 | 1.00 | N | 0 | 0 |
| 1 | 0 | 0 | 0 | 2 | 2 | 0.79 | N | 0 | 0 |
| 0 | 0 | 0 | 1 | 3 | 4 | 0.60 | N | 1 | 0 |
| 0 | 0 | 1 | 0 | 5 | 5 | 0.52 | Y | 0 | 0 |
| 1 | 0 | 1 | 0 | 5 | 5 | 0.82 | N | 0 | 0 |
| 1 | 0 | 0 | 0 | 2 | 2 | 0.81 | N | 0 | 0 |
| 0 | 0 | 1 | 0 | 7 | 7 | 0.63 | Y | 0 | 0 |
| 0 | 0 | 2 | 0 | 4 | 4 | 0.59 | Y | 0 | 0 |
| 1 | 0 | 2 | 1 | 7 | 8 | 0.58 | Y | 1 | 0 |
| 0 | 0 | 2 | 2 | 6 | 8 | 0.57 | Y | 0 | 2 |
| 0 | 0 | 2 | 0 | 7 | 7 | 0.53 | Y | 2 | 0 |
| 1 | 0 | 2 | 0 | 7 | 7 | 0.64 | Y | 1 | 1 |
| 0 | 0 | 2 | 2 | 6 | 8 | 0.63 | Y | 1 | 1 |
| 1 | 0 | 2 | 0 | 7 | 7 | 0.57 | Y | 1 | 1 |
| 1 | 0 | 1 | 3 | 6 | 9 | 0.69 | Y | 2 | 1 |
| 1 | 0 | 0 | 0 | 2 | 2 | 0.79 | N | 0 | 0 |
| 0 | 0 | 0 | 0 | 4 | 4 | 0.65 | N | 1 | 0 |
| 2 | 0 | 2 | 0 | 9 | 9 | 0.64 | Y | 0 | 0 |
| 2 | 0 | 0 | 1 | 6 | 7 | 0.53 | Y | 1 | 1 |
| 0 | 0 | 1 | 0 | 5 | 5 | 0.52 | Y | 0 | 0 |
| 0 | 0 | 0 | 1 | 3 | 4 | 0.68 | N | 1 | 0 |

| | | | | | | | | | |
|---|---|---|---|---|---|---|---|---|---|
| 2 | 0 | 1 | 0 | 8 | 8 | 0.71 | Y | 1 | 1 |
| 1 | 0 | 1 | 1 | 7 | 8 | 0.60 | Y | 1 | 0 |
| 2 | 0 | 1 | 2 | 8 | 10 | 0.64 | N | 1 | 1 |
| 1 | 0 | 1 | 0 | 4 | 4 | 0.75 | N | 0 | 0 |
| 1 | 0 | 1 | 0 | 4 | 4 | 0.81 | N | 0 | 1 |
| 0 | 0 | 1 | 0 | 4 | 4 | 0.57 | Y | 0 | 0 |
| 1 | 0 | 1 | 0 | 6 | 6 | 0.64 | Y | 0 | 0 |
| 0 | 0 | 2 | 0 | 7 | 7 | 0.65 | Y | 0 | 0 |
| 0 | 1 | 0 | 1 | 4 | 5 | 0.62 | Y | 0 | 0 |
| 0 | 0 | 2 | 0 | 7 | 7 | 0.56 | Y | 0 | 0 |
| 2 | 1 | 0 | 3 | 5 | 8 | 0.73 | Y | 1 | 0 |
| 0 | 0 | 2 | 0 | 6 | 6 | 0.64 | Y | 0 | 0 |
| 0 | 0 | 1 | 0 | 5 | 5 | 0.62 | Y | 0 | 1 |
| 0 | 1 | 1 | 2 | 5 | 7 | 0.65 | Y | 1 | 0 |
| 0 | 0 | 1 | 1 | 4 | 5 | 0.69 | Y | 2 | 0 |
| 0 | 0 | 2 | 0 | 7 | 7 | 0.57 | Y | 1 | 1 |
| 1 | 0 | 1 | 1 | 6 | 7 | 0.56 | Y | 1 | 1 |
| 1 | 0 | 2 | 1 | 8 | 9 | 0.64 | Y | 1 | 0 |
| 1 | 0 | 2 | 0 | 7 | 7 | 0.54 | Y | 2 | 0 |
| 1 | 0 | 1 | 1 | 6 | 7 | 0.65 | Y | 1 | 0 |
| 0 | 0 | 0 | 2 | 3 | 5 | 0.55 | Y | 1 | 1 |
| 0 | 0 | 2 | 0 | 8 | 8 | 0.55 | Y | 1 | 1 |
| 0 | 0 | 1 | 1 | 6 | 7 | 0.63 | Y | 0 | 2 |
| 0 | 1 | 0 | 2 | 4 | 6 | 0.69 | Y | 2 | 0 |
| 1 | 0 | 1 | 2 | 7 | 9 | 0.73 | Y | 0 | 2 |
| 2 | 0 | 0 | 0 | 8 | 8 | 0.62 | Y | 1 | 1 |
| 2 | 0 | 2 | 1 | 7 | 8 | 0.60 | Y | 1 | 1 |
| 1 | 0 | 1 | 2 | 6 | 8 | 0.56 | Y | 1 | 0 |
| 0 | 1 | 1 | 3 | 5 | 8 | 0.71 | Y | 0 | 1 |
| 2 | 0 | 2 | 0 | 9 | 9 | 0.56 | Y | 1 | 1 |

| | | | | | | | | | |
|---|---|---|---|---|---|---|---|---|---|
| 0 | 0 | 0 | 3 | 3 | 6 | 0.71 | Y | 1 | 0 |
| 2 | 0 | 1 | 0 | 9 | 9 | 0.73 | Y | 0 | 2 |
| 2 | 0 | 1 | 1 | 6 | 7 | 0.51 | Y | 1 | 0 |
| 1 | 0 | 1 | 1 | 8 | 9 | 0.73 | N | 1 | 1 |
| 2 | 0 | 1 | 3 | 7 | 10 | 0.71 | N | 1 | 1 |
| 0 | 0 | 1 | 0 | 3 | 3 | 0.76 | N | 0 | 0 |
| 0 | 0 | 0 | 0 | 1 | 1 | 0.92 | N | 0 | 0 |
| 0 | 0 | 1 | 0 | 3 | 3 | 0.68 | N | 0 | 0 |
| 0 | 0 | 1 | 0 | 5 | 5 | 0.69 | Y | 0 | 0 |
| 0 | 0 | 1 | 0 | 3 | 3 | 0.65 | N | 0 | 0 |
| 0 | 0 | 2 | 0 | 3 | 3 | 0.93 | N | 0 | 0 |
| 0 | 0 | 2 | 0 | 4 | 4 | 0.81 | N | 0 | 1 |
| 0 | 0 | 2 | 0 | 3 | 3 | 0.89 | Y | 0 | 0 |
| 0 | 0 | 0 | 0 | 1 | 1 | 0.94 | N | 0 | 0 |
| 0 | 0 | 1 | 0 | 2 | 2 | 1.00 | N | 0 | 0 |
| 0 | 0 | 1 | 0 | 4 | 4 | 0.68 | Y | 0 | 0 |
| 1 | 0 | 2 | 0 | 6 | 6 | 0.93 | N | 0 | 0 |
| 0 | 0 | 1 | 0 | 3 | 3 | 0.87 | N | 0 | 0 |
| 0 | 1 | 0 | 1 | 2 | 3 | 0.66 | N | 0 | 0 |
| 0 | 0 | 2 | 0 | 3 | 3 | 0.90 | N | 0 | 0 |
| 1 | 0 | 1 | 0 | 4 | 4 | 0.77 | N | 0 | 0 |
| 0 | 0 | 2 | 0 | 4 | 4 | 0.81 | N | 0 | 0 |
| 0 | 0 | 1 | 0 | 4 | 4 | 0.74 | Y | 0 | 0 |
| 0 | 0 | 1 | 1 | 2 | 3 | 0.70 | N | 0 | 0 |
| 0 | 0 | 2 | 0 | 6 | 6 | 0.56 | N | 0 | 0 |
| 0 | 0 | 2 | 0 | 6 | 6 | 0.58 | Y | 0 | 0 |
| 0 | 0 | 1 | 0 | 4 | 4 | 0.51 | N | 0 | 0 |
| 0 | 0 | 1 | 0 | 3 | 3 | 0.63 | Y | 0 | 0 |
| 1 | 0 | 1 | 0 | 4 | 4 | 0.75 | N | 0 | 0 |
| 0 | 0 | 1 | 0 | 3 | 3 | 0.66 | Y | 1 | 0 |

| | | | | | | | | | |
|---|---|---|---|---|---|---|---|---|---|
| 0 | 0 | 2 | 0 | 5 | 5 | 0.71 | Y | 0 | 1 |
| 0 | 0 | 1 | 1 | 4 | 5 | 0.65 | N | 1 | 0 |
| 0 | 1 | 1 | 1 | 5 | 6 | 0.72 | N | 0 | 0 |
| 1 | 0 | 2 | 1 | 5 | 6 | 0.54 | Y | 0 | 0 |
| 0 | 0 | 2 | 0 | 6 | 6 | 0.63 | Y | 0 | 0 |
| 1 | 0 | 1 | 0 | 5 | 5 | 0.62 | Y | 0 | 0 |
| 0 | 0 | 1 | 0 | 3 | 3 | 0.73 | N | 0 | 0 |
| 0 | 0 | 2 | 0 | 6 | 6 | 0.66 | N | 0 | 1 |
| 0 | 0 | 1 | 0 | 3 | 3 | 0.93 | N | 0 | 0 |
| 0 | 0 | 1 | 0 | 3 | 3 | 0.64 | N | 0 | 0 |
| 2 | 0 | 2 | 0 | 7 | 7 | 0.59 | Y | 0 | 0 |
| 0 | 0 | 2 | 0 | 7 | 7 | 0.72 | Y | 0 | 0 |
| 0 | 0 | 0 | 0 | 1 | 1 | 0.94 | N | 0 | 0 |
| 0 | 0 | 1 | 1 | 5 | 6 | 0.62 | Y | 1 | 0 |
| 0 | 0 | 1 | 0 | 3 | 3 | 0.63 | N | 0 | 0 |
| 0 | 0 | 2 | 0 | 7 | 7 | 0.67 | Y | 0 | 1 |
| 2 | 0 | 1 | 0 | 7 | 7 | 0.74 | N | 0 | 0 |
| 2 | 0 | 1 | 0 | 7 | 7 | 0.68 | N | 0 | 0 |
| 1 | 0 | 1 | 0 | 4 | 4 | 0.67 | N | 0 | 0 |
| 2 | 0 | 0 | 0 | 6 | 6 | 0.62 | Y | 0 | 0 |
| 2 | 0 | 0 | 0 | 6 | 6 | 0.57 | N | 0 | 0 |
| 1 | 0 | 2 | 1 | 4 | 5 | 0.90 | N | 0 | 1 |
| 0 | 0 | 2 | 0 | 4 | 4 | 0.80 | N | 0 | 0 |
| 2 | 0 | 0 | 0 | 6 | 6 | 0.57 | N | 0 | 0 |
| 0 | 0 | 2 | 0 | 3 | 3 | 0.83 | N | 1 | 0 |
| 2 | 0 | 1 | 0 | 7 | 7 | 0.63 | N | 0 | 1 |
| 2 | 0 | 1 | 0 | 7 | 7 | 0.56 | Y | 0 | 0 |
| 2 | 0 | 0 | 0 | 6 | 6 | 0.70 | Y | 0 | 0 |
| 0 | 0 | 1 | 1 | 3 | 4 | 0.70 | N | 0 | 0 |
| 0 | 0 | 1 | 1 | 3 | 4 | 0.58 | N | 0 | 0 |

| | | | | | | | | | |
|---|---|---|---|---|---|---|---|---|---|
| 0 | 0 | 2 | 0 | 6 | 6 | 0.57 | Y | 0 | 0 |
| 0 | 0 | 2 | 1 | 4 | 5 | 0.67 | Y | 1 | 0 |
| 2 | 0 | 2 | 0 | 6 | 6 | 0.63 | Y | 0 | 0 |
| 1 | 0 | 2 | 0 | 7 | 7 | 0.67 | Y | 1 | 1 |
| 0 | 0 | 2 | 1 | 8 | 9 | 0.64 | N | 0 | 2 |
| 2 | 0 | 1 | 0 | 7 | 7 | 0.64 | Y | 0 | 0 |
| 2 | 0 | 2 | 0 | 8 | 8 | 0.59 | Y | 0 | 0 |
| 1 | 0 | 2 | 1 | 8 | 9 | 0.59 | Y | 0 | 0 |
| 0 | 1 | 1 | 3 | 8 | 11 | 0.86 | N | 1 | 0 |
| 0 | 0 | 2 | 0 | 6 | 6 | 0.64 | Y | 0 | 0 |
| 0 | 0 | 1 | 0 | 5 | 5 | 0.68 | N | 0 | 1 |
| 2 | 0 | 1 | 0 | 8 | 8 | 0.61 | N | 0 | 1 |
| 1 | 0 | 2 | 0 | 7 | 7 | 0.66 | Y | 1 | 0 |
| 0 | 0 | 1 | 0 | 6 | 6 | 0.62 | Y | 0 | 1 |
| 2 | 0 | 0 | 0 | 6 | 6 | 0.61 | N | 0 | 0 |
| 2 | 0 | 1 | 1 | 8 | 9 | 0.68 | N | 0 | 0 |
| 2 | 0 | 1 | 0 | 7 | 7 | 0.58 | Y | 0 | 0 |
| 0 | 0 | 2 | 2 | 6 | 8 | 0.70 | N | 0 | 2 |
| 2 | 0 | 2 | 0 | 9 | 9 | 0.60 | Y | 0 | 0 |
| 1 | 0 | 2 | 0 | 11 | 11 | 0.79 | Y | 0 | 2 |
| 2 | 0 | 0 | 0 | 6 | 6 | 0.56 | N | 0 | 1 |
| 2 | 0 | 1 | 0 | 7 | 7 | 0.63 | Y | 0 | 0 |
| 1 | 0 | 2 | 2 | 9 | 11 | 0.81 | N | 0 | 2 |

| IB-Forearm (Ulna & [illegible] | IB-Forearm (Ulna & [illegible] | IB-Olecranon F [illegible] | IB-Olecranon D [illegible] | IB-Ischium | IB-Femur F. | IB-Femur D. | IB-Tibia F. | IB-Tibia D. | IB-Calcaneus F [illegible] |
|---|---|---|---|---|---|---|---|---|---|
| 0 | 0 | 0 | 0 | 0 | 0 | 0 | 0 | 0 | 0 |
| 0 | 0 | 0 | 0 | 0 | 0 | 0 | 0 | 0 | 0 |
| 0 | 0 | 0 | 0 | 0 | 0 | 0 | 0 | 0 | 0 |
| 0 | 0 | 0 | 0 | 0 | 0 | 0 | 0 | 0 | 0 |
| 0 | 0 | 0 | 0 | 0 | 0 | 0 | 0 | 0 | 0 |
| 0 | 0 | 0 | 0 | 0 | 0 | 0 | 0 | 0 | 0 |
| 0 | 0 | 0 | 0 | 0 | 0 | 0 | 0 | 0 | 0 |
| 0 | 0 | 0 | 0 | 0 | 0 | 0 | 0 | 0 | 0 |
| 0 | 0 | 0 | 0 | 0 | 0 | 0 | 0 | 0 | 0 |
| 0 | 0 | 0 | 0 | 0 | 0 | 0 | 0 | 0 | 0 |
| 0 | 0 | 0 | 0 | 0 | 0 | 0 | 0 | 0 | 0 |
| 0 | 0 | 0 | 0 | 0 | 0 | 0 | 0 | 0 | 0 |
| 0 | 0 | 0 | 0 | 0 | 0 | 0 | 0 | 0 | 0 |
| 0 | 0 | 0 | 0 | 0 | 0 | 0 | 0 | 0 | 0 |
| 0 | 0 | 0 | 0 | 0 | 0 | 0 | 0 | 0 | 0 |
| 0 | 0 | 0 | 0 | 0 | 0 | 0 | 0 | 0 | 0 |
| 0 | 0 | 0 | 0 | 0 | 0 | 0 | 0 | 0 | 0 |
| 0 | 0 | 0 | 0 | 0 | 0 | 0 | 0 | 0 | 0 |
| 0 | 0 | 0 | 0 | 0 | 0 | 0 | 0 | 0 | 0 |
| 0 | 0 | 0 | 0 | 0 | 0 | 0 | 0 | 0 | 0 |
| 0 | 0 | 0 | 0 | 0 | 0 | 0 | 0 | 0 | 0 |
| 0 | 0 | 0 | 0 | 0 | 0 | 0 | 0 | 0 | 0 |
| 0 | 0 | 0 | 0 | 0 | 0 | 0 | 0 | 0 | 0 |
| 0 | 0 | 0 | 0 | 0 | 0 | 0 | 0 | 0 | 0 |
| 0 | 0 | 0 | 0 | 0 | 0 | 0 | 0 | 0 | 0 |
| 0 | 0 | 0 | 0 | 0 | 0 | 0 | 0 | 0 | 0 |
| 0 | 0 | 0 | 0 | 0 | 0 | 0 | 0 | 0 | 0 |
| 0 | 0 | 0 | 0 | 0 | 0 | 0 | 0 | 0 | 0 |
| 0 | 0 | 0 | 0 | 0 | 0 | 0 | 0 | 0 | 0 |

| | | | | | | | | | |
|---|---|---|---|---|---|---|---|---|---|
| 0 | 0 | 0 | 0 | 0 | 0 | 0 | 0 | 0 | 0 |
| 0 | 0 | 0 | 0 | 0 | 0 | 0 | 0 | 0 | 0 |
| 0 | 0 | 0 | 0 | 0 | 0 | 0 | 0 | 0 | 0 |
| 0 | 0 | 0 | 0 | 0 | 0 | 0 | 0 | 0 | 0 |
| 0 | 0 | 0 | 0 | 0 | 0 | 0 | 0 | 0 | 0 |
| 0 | 0 | 0 | 0 | 0 | 0 | 0 | 0 | 0 | 0 |
| 0 | 0 | 0 | 0 | 0 | 0 | 0 | 0 | 0 | 0 |
| 0 | 0 | 0 | 0 | 0 | 0 | 0 | 0 | 0 | 0 |
| 0 | 0 | 0 | 0 | 0 | 0 | 0 | 0 | 0 | 0 |
| 0 | 0 | 0 | 0 | 0 | 0 | 0 | 0 | 0 | 0 |
| 0 | 0 | 0 | 0 | 0 | 0 | 0 | 0 | 0 | 0 |
| 0 | 0 | 0 | 0 | 0 | 0 | 0 | 0 | 0 | 0 |
| 0 | 0 | 0 | 0 | 0 | 0 | 0 | 0 | 0 | 0 |
| 0 | 0 | 0 | 0 | 0 | 0 | 0 | 0 | 0 | 0 |
| 0 | 0 | 0 | 0 | 0 | 0 | 0 | 0 | 0 | 0 |
| 0 | 0 | 0 | 0 | 0 | 0 | 0 | 0 | 0 | 0 |
| 0 | 0 | 0 | 0 | 0 | 0 | 0 | 0 | 0 | 0 |
| 0 | 0 | 0 | 0 | 0 | 0 | 0 | 0 | 0 | 0 |
| 0 | 0 | 0 | 0 | 0 | 0 | 0 | 0 | 0 | 0 |
| 0 | 0 | 0 | 0 | 0 | 0 | 0 | 0 | 0 | 0 |
| 0 | 0 | 0 | 0 | 0 | 0 | 0 | 0 | 0 | 0 |
| 0 | 0 | 0 | 0 | 0 | 0 | 0 | 0 | 0 | 0 |
| 0 | 0 | 0 | 0 | 0 | 0 | 0 | 0 | 0 | 0 |
| 0 | 0 | 0 | 0 | 0 | 0 | 0 | 0 | 0 | 0 |
| 0 | 0 | 0 | 0 | 0 | 0 | 0 | 0 | 0 | 0 |
| 0 | 0 | 0 | 0 | 0 | 0 | 0 | 0 | 0 | 0 |
| 0 | 0 | 0 | 0 | 0 | 0 | 0 | 0 | 0 | 0 |
| 0 | 0 | 0 | 0 | 0 | 0 | 0 | 0 | 0 | 0 |
| 0 | 0 | 0 | 0 | 0 | 0 | 0 | 0 | 0 | 0 |
| 0 | 0 | 0 | 0 | 0 | 0 | 0 | 0 | 0 | 0 |

| | | | | | | | | | |
|---|---|---|---|---|---|---|---|---|---|
| 0 | 0 | 0 | 0 | 0 | 0 | 0 | 0 | 0 | 0 |
| 0 | 0 | 0 | 0 | 0 | 0 | 0 | 0 | 0 | 0 |
| 0 | 0 | 0 | 0 | 0 | 0 | 0 | 0 | 0 | 0 |
| 0 | 0 | 0 | 0 | 0 | 0 | 0 | 0 | 0 | 0 |
| 0 | 0 | 0 | 0 | 0 | 0 | 0 | 0 | 0 | 0 |
| 0 | 0 | 0 | 0 | 0 | 0 | 0 | 0 | 0 | 0 |
| 0 | 0 | 0 | 0 | 0 | 0 | 0 | 0 | 0 | 0 |
| 0 | 0 | 0 | 0 | 0 | 0 | 0 | 0 | 0 | 0 |
| 0 | 0 | 0 | 0 | 0 | 0 | 0 | 0 | 0 | 0 |
| 0 | 0 | 0 | 0 | 0 | 0 | 0 | 0 | 0 | 0 |
| 0 | 0 | 0 | 0 | 2 | 0 | 0 | 0 | 1 | 0 |
| 0 | 1 | 0 | 0 | 2 | 0 | 0 | 0 | 0 | 0 |
| 0 | 2 | 0 | 0 | 2 | 0 | 1 | 0 | 0 | 0 |
| 0 | 0 | 0 | 0 | 2 | 0 | 0 | 0 | 0 | 0 |
| 0 | 0 | 0 | 0 | 1 | 0 | 0 | 1 | 0 | 0 |
| 0 | 2 | 0 | 0 | 2 | 0 | 0 | 0 | 0 | 0 |
| 0 | 2 | 0 | 0 | 2 | 1 | 0 | 0 | 0 | 0 |
| 0 | 2 | 0 | 0 | 2 | 0 | 0 | 1 | 0 | 0 |
| 0 | 2 | 0 | 0 | 2 | 0 | 0 | 0 | 0 | 0 |
| 0 | 1 | 0 | 0 | 2 | 0 | 0 | 0 | 0 | 0 |
| 0 | 0 | 0 | 0 | 2 | 0 | 0 | 0 | 0 | 0 |
| 0 | 2 | 0 | 0 | 2 | 0 | 0 | 1 | 0 | 0 |
| 0 | 2 | 0 | 0 | 2 | 0 | 0 | 0 | 1 | 0 |
| 0 | 1 | 0 | 0 | 2 | 0 | 0 | 0 | 2 | 0 |
| 0 | 0 | 0 | 0 | 2 | 0 | 0 | 0 | 1 | 0 |
| 0 | 1 | 0 | 0 | 2 | 0 | 0 | 0 | 0 | 0 |
| 0 | 2 | 0 | 0 | 0 | 0 | 0 | 2 | 0 | 0 |
| 0 | 0 | 0 | 0 | 2 | 0 | 0 | 0 | 0 | 0 |
| 0 | 2 | 0 | 0 | 2 | 0 | 1 | 0 | 0 | 0 |
| 0 | 1 | 0 | 0 | 2 | 0 | 0 | 0 | 0 | 0 |

| | | | | | | | | | |
|---|---|---|---|---|---|---|---|---|---|
| 1 | 2 | 0 | 0 | 2 | 0 | 0 | 2 | 0 | 0 |
| 0 | 2 | 0 | 0 | 2 | 0 | 0 | 2 | 0 | 0 |
| 0 | 2 | 0 | 0 | 0 | 0 | 1 | 1 | 1 | 0 |
| 0 | 0 | 0 | 0 | 2 | 0 | 0 | 0 | 0 | 0 |
| 0 | 0 | 0 | 0 | 2 | 0 | 0 | 0 | 0 | 0 |
| 0 | 2 | 0 | 0 | 2 | 1 | 0 | 0 | 0 | 0 |
| 0 | 2 | 0 | 0 | 2 | 0 | 0 | 1 | 0 | 0 |
| 0 | 2 | 0 | 0 | 2 | 0 | 0 | 0 | 0 | 0 |
| 0 | 2 | 0 | 0 | 2 | 0 | 0 | 0 | 0 | 1 |
| 0 | 2 | 0 | 0 | 2 | 1 | 0 | 0 | 0 | 0 |
| 0 | 1 | 0 | 0 | 2 | 0 | 0 | 2 | 0 | 0 |
| 0 | 2 | 0 | 0 | 2 | 1 | 0 | 0 | 0 | 0 |
| 0 | 2 | 0 | 0 | 2 | 0 | 0 | 0 | 0 | 0 |
| 0 | 2 | 0 | 0 | 2 | 0 | 0 | 0 | 0 | 0 |
| 0 | 0 | 0 | 0 | 2 | 0 | 0 | 1 | 0 | 0 |
| 0 | 2 | 0 | 0 | 2 | 0 | 0 | 0 | 0 | 0 |
| 0 | 2 | 0 | 0 | 2 | 0 | 0 | 0 | 1 | 0 |
| 0 | 2 | 0 | 0 | 2 | 0 | 0 | 1 | 0 | 0 |
| 0 | 0 | 0 | 0 | 2 | 0 | 0 | 0 | 0 | 0 |
| 0 | 2 | 0 | 0 | 2 | 0 | 0 | 1 | 0 | 0 |
| 0 | 0 | 0 | 0 | 2 | 0 | 0 | 0 | 0 | 0 |
| 0 | 2 | 0 | 0 | 2 | 0 | 0 | 0 | 0 | 0 |
| 0 | 1 | 0 | 0 | 2 | 0 | 0 | 1 | 0 | 0 |
| 0 | 2 | 0 | 0 | 2 | 0 | 0 | 2 | 0 | 0 |
| 0 | 1 | 0 | 0 | 2 | 0 | 0 | 2 | 2 | 0 |
| 0 | 1 | 0 | 0 | 2 | 0 | 0 | 0 | 0 | 0 |
| 0 | 2 | 0 | 0 | 2 | 0 | 0 | 1 | 0 | 0 |
| 0 | 2 | 0 | 0 | 2 | 0 | 0 | 2 | 0 | 0 |
| 1 | 2 | 0 | 0 | 2 | 0 | 0 | 2 | 0 | 0 |
| 0 | 2 | 0 | 0 | 2 | 0 | 0 | 1 | 0 | 0 |

| | | | | | | | | | |
|---|---|---|---|---|---|---|---|---|---|
| 0 | 2 | 0 | 0 | 2 | 0 | 0 | 2 | 1 | 0 |
| 1 | 2 | 0 | 0 | 2 | 0 | 0 | 0 | 2 | 0 |
| 0 | 2 | 0 | 0 | 2 | 0 | 0 | 2 | 0 | 0 |
| 0 | 2 | 0 | 0 | 0 | 0 | 1 | 2 | 0 | 0 |
| 0 | 2 | 0 | 0 | 2 | 1 | 0 | 0 | 0 | 0 |
| 0 | 0 | 0 | 0 | 2 | 0 | 0 | 0 | 0 | 0 |
| 0 | 0 | 0 | 0 | 2 | 0 | 0 | 1 | 0 | 0 |
| 0 | 0 | 0 | 0 | 2 | 0 | 0 | 0 | 0 | 0 |
| 0 | 0 | 0 | 0 | 2 | 0 | 0 | 0 | 0 | 0 |
| 0 | 0 | 0 | 0 | 2 | 0 | 0 | 0 | 0 | 0 |
| 0 | 0 | 0 | 0 | 1 | 0 | 0 | 0 | 0 | 1 |
| 0 | 0 | 0 | 0 | 0 | 0 | 0 | 0 | 0 | 0 |
| 0 | 0 | 0 | 0 | 2 | 0 | 0 | 0 | 0 | 0 |
| 0 | 0 | 0 | 0 | 2 | 0 | 0 | 0 | 0 | 0 |
| 0 | 0 | 0 | 0 | 0 | 0 | 0 | 0 | 0 | 0 |
| 0 | 0 | 0 | 0 | 0 | 0 | 0 | 0 | 0 | 0 |
| 0 | 0 | 0 | 0 | 0 | 0 | 0 | 0 | 0 | 0 |
| 0 | 0 | 0 | 0 | 2 | 0 | 0 | 0 | 0 | 0 |
| 0 | 0 | 0 | 0 | 2 | 0 | 0 | 0 | 0 | 0 |
| 0 | 2 | 0 | 0 | 2 | 0 | 0 | 0 | 0 | 0 |
| 0 | 0 | 0 | 0 | 2 | 0 | 0 | 0 | 2 | 0 |
| 0 | 0 | 0 | 0 | 2 | 0 | 0 | 0 | 0 | 0 |
| 0 | 1 | 0 | 0 | 2 | 0 | 0 | 0 | 0 | 0 |
| 0 | 0 | 0 | 0 | 2 | 0 | 0 | 1 | 0 | 0 |
| 0 | 0 | 0 | 0 | 2 | 0 | 0 | 0 | 0 | 0 |
| 0 | 0 | 0 | 0 | 2 | 0 | 0 | 0 | 0 | 0 |
| 0 | 1 | 0 | 0 | 2 | 0 | 0 | 0 | 0 | 0 |
| 0 | 0 | 0 | 1 | 0 | 0 | 0 | 0 | 0 | 0 |
| 0 | 2 | 0 | 0 | 2 | 0 | 0 | 2 | 0 | 0 |
| 0 | 0 | 0 | 0 | 2 | 0 | 0 | 0 | 0 | 0 |

| | | | | | | | | | |
|---|---|---|---|---|---|---|---|---|---|
| 0 | 2 | 0 | 0 | 2 | 0 | 0 | 1 | 0 | 0 |
| 0 | 1 | 0 | 0 | 2 | 0 | 0 | 0 | 0 | 0 |
| 0 | 2 | 0 | 0 | 2 | 0 | 0 | 0 | 0 | 0 |
| 0 | 0 | 0 | 0 | 2 | 0 | 0 | 1 | 1 | 0 |
| 0 | 2 | 0 | 0 | 2 | 0 | 0 | 0 | 0 | 0 |
| 0 | 1 | 0 | 0 | 2 | 0 | 0 | 0 | 0 | 0 |
| 0 | 0 | 0 | 0 | 2 | 0 | 0 | 0 | 0 | 0 |
| 0 | 2 | 0 | 0 | 2 | 0 | 0 | 0 | 0 | 0 |
| 0 | 0 | 0 | 0 | 1 | 0 | 0 | 0 | 0 | 0 |
| 0 | 0 | 0 | 0 | 2 | 0 | 0 | 0 | 0 | 0 |
| 0 | 0 | 0 | 0 | 2 | 0 | 0 | 1 | 1 | 0 |
| 0 | 2 | 0 | 0 | 2 | 0 | 0 | 0 | 0 | 0 |
| 0 | 0 | 0 | 0 | 2 | 0 | 0 | 0 | 0 | 0 |
| 0 | 2 | 0 | 0 | 2 | 0 | 0 | 0 | 0 | 0 |
| 0 | 0 | 0 | 1 | 0 | 0 | 0 | 0 | 0 | 0 |
| 0 | 2 | 0 | 0 | 2 | 0 | 0 | 0 | 0 | 0 |
| 0 | 2 | 0 | 0 | 2 | 0 | 0 | 0 | 2 | 0 |
| 0 | 2 | 0 | 0 | 2 | 0 | 0 | 2 | 0 | 0 |
| 0 | 0 | 0 | 0 | 2 | 0 | 0 | 0 | 2 | 0 |
| 0 | 2 | 0 | 0 | 2 | 0 | 0 | 2 | 0 | 0 |
| 0 | 2 | 0 | 0 | 2 | 0 | 0 | 2 | 0 | 0 |
| 0 | 0 | 0 | 0 | 0 | 0 | 0 | 0 | 0 | 0 |
| 0 | 0 | 0 | 0 | 2 | 0 | 0 | 0 | 0 | 0 |
| 0 | 2 | 0 | 0 | 2 | 0 | 0 | 0 | 0 | 0 |
| 0 | 2 | 0 | 0 | 2 | 0 | 0 | 1 | 0 | 0 |
| 0 | 2 | 0 | 0 | 2 | 0 | 0 | 2 | 0 | 0 |
| 0 | 2 | 0 | 0 | 2 | 0 | 0 | 0 | 2 | 0 |
| 0 | 2 | 0 | 0 | 2 | 0 | 0 | 2 | 0 | 0 |
| 0 | 1 | 0 | 0 | 2 | 0 | 0 | 0 | 0 | 0 |
| 0 | 0 | 0 | 0 | 2 | 0 | 0 | 0 | 1 | 0 |

| | | | | | | | | | |
|---|---|---|---|---|---|---|---|---|---|
| 0 | 0 | 0 | 0 | 2 | 0 | 0 | 0 | 0 | 0 |
| 0 | 1 | 0 | 0 | 2 | 0 | 0 | 1 | 0 | 0 |
| 0 | 0 | 0 | 0 | 2 | 0 | 0 | 2 | 0 | 0 |
| 0 | 2 | 0 | 0 | 2 | 0 | 0 | 1 | 0 | 0 |
| 0 | 2 | 0 | 0 | 2 | 0 | 0 | 0 | 0 | 0 |
| 0 | 2 | 0 | 0 | 2 | 0 | 0 | 0 | 2 | 0 |
| 0 | 2 | 0 | 0 | 2 | 0 | 0 | 0 | 2 | 0 |
| 0 | 2 | 0 | 0 | 2 | 0 | 0 | 2 | 0 | 0 |
| 0 | 2 | 0 | 0 | 2 | 0 | 0 | 1 | 0 | 0 |
| 0 | 2 | 0 | 0 | 2 | 0 | 0 | 0 | 0 | 0 |
| 0 | 1 | 0 | 0 | 2 | 0 | 0 | 0 | 0 | 0 |
| 0 | 2 | 0 | 0 | 2 | 0 | 0 | 2 | 0 | 0 |
| 0 | 1 | 0 | 0 | 2 | 0 | 0 | 1 | 0 | 0 |
| 0 | 2 | 0 | 0 | 2 | 0 | 0 | 0 | 0 | 0 |
| 0 | 2 | 0 | 0 | 2 | 0 | 0 | 2 | 0 | 0 |
| 0 | 2 | 0 | 0 | 2 | 0 | 0 | 0 | 2 | 0 |
| 0 | 2 | 0 | 0 | 2 | 0 | 0 | 0 | 2 | 0 |
| 0 | 2 | 0 | 0 | 2 | 0 | 0 | 0 | 0 | 0 |
| 0 | 2 | 0 | 0 | 2 | 0 | 0 | 1 | 1 | 0 |
| 0 | 2 | 0 | 0 | 2 | 0 | 0 | 1 | 1 | 0 |
| 0 | 2 | 0 | 0 | 2 | 0 | 0 | 2 | 0 | 0 |
| 0 | 2 | 0 | 0 | 2 | 0 | 0 | 2 | 0 | 0 |
| 0 | 2 | 0 | 0 | 2 | 0 | 0 | 0 | 1 | 0 |

| IB-Calcaneus D | IB-Total Fractures | IB-Total Deformities | IB-Total Score | IB-Pelvis Ratio | IB-Scoliosis | IE-Humerus F | IE-Humerus D | IE-Forearm (Ulna & | IE-Forearm (Ulna & |
|---|---|---|---|---|---|---|---|---|---|
| 0 | 0 | 0 | 0 | 1.00 | N | 0 | 0 | 0 | 0 |
| 0 | 0 | 0 | 0 | 1.00 | N | 0 | 0 | 0 | 0 |
| 0 | 0 | 0 | 0 | 1.00 | N | 0 | 0 | 0 | 0 |
| 0 | 0 | 0 | 0 | 1.00 | N | 0 | 0 | 0 | 0 |
| 0 | 0 | 0 | 0 | 1.00 | N | 0 | 0 | 0 | 0 |
| 0 | 0 | 0 | 0 | 1.00 | N | 0 | 0 | 0 | 0 |
| 0 | 0 | 0 | 0 | 1.00 | N | 0 | 0 | 0 | 0 |
| 0 | 0 | 0 | 0 | 1.00 | N | 0 | 0 | 0 | 0 |
| 0 | 0 | 0 | 0 | 1.00 | N | 0 | 0 | 0 | 0 |
| 0 | 0 | 0 | 0 | 1.00 | N | 0 | 0 | 0 | 0 |
| 0 | 0 | 0 | 0 | 1.00 | N | 0 | 0 | 0 | 0 |
| 0 | 0 | 0 | 0 | 1.00 | N | 0 | 0 | 0 | 0 |
| 0 | 0 | 0 | 0 | 1.00 | N | 0 | 0 | 0 | 0 |
| 0 | 0 | 0 | 0 | 1.00 | N | 0 | 0 | 0 | 0 |
| 0 | 0 | 0 | 0 | 1.00 | N | 0 | 0 | 0 | 0 |
| 0 | 0 | 0 | 0 | 1.00 | N | 0 | 0 | 0 | 0 |
| 0 | 0 | 0 | 0 | 1.00 | N | 0 | 0 | 0 | 0 |
| 0 | 0 | 0 | 0 | 1.00 | N | 0 | 0 | 0 | 0 |
| 0 | 0 | 0 | 0 | 1.00 | N | 0 | 0 | 0 | 0 |
| 0 | 0 | 0 | 0 | 1.00 | N | 0 | 0 | 0 | 0 |
| 0 | 0 | 0 | 0 | 1.00 | N | 0 | 0 | 0 | 0 |
| 0 | 0 | 0 | 0 | 1.00 | N | 0 | 0 | 0 | 0 |
| 0 | 0 | 0 | 0 | 1.00 | N | 0 | 0 | 0 | 0 |
| 0 | 0 | 0 | 0 | 1.00 | N | 0 | 0 | 0 | 0 |
| 0 | 0 | 0 | 0 | 1.00 | N | 0 | 0 | 0 | 0 |
| 0 | 0 | 0 | 0 | 1.00 | N | 0 | 0 | 0 | 0 |
| 0 | 0 | 0 | 0 | 1.00 | N | 0 | 0 | 0 | 0 |
| 0 | 0 | 0 | 0 | 1.00 | N | 0 | 0 | 0 | 0 |
| 0 | 0 | 0 | 0 | 1.00 | N | 0 | 0 | 0 | 0 |

| | | | | | | | | | |
|---|---|---|---|---|---|---|---|---|---|
| 0 | 0 | 0 | 0 | 1.00 | N | 0 | 0 | 0 | 0 |
| 0 | 0 | 0 | 0 | 1.00 | N | 0 | 0 | 0 | 0 |
| 0 | 0 | 0 | 0 | 1.00 | N | 0 | 0 | 0 | 0 |
| 0 | 0 | 0 | 0 | 1.00 | N | 0 | 0 | 0 | 0 |
| 0 | 0 | 0 | 0 | 1.00 | N | 0 | 0 | 0 | 0 |
| 0 | 0 | 0 | 0 | 1.00 | N | 0 | 0 | 0 | 0 |
| 0 | 0 | 0 | 0 | 1.00 | N | 0 | 0 | 0 | 0 |
| 0 | 0 | 0 | 0 | 1.00 | N | 0 | 0 | 0 | 0 |
| 0 | 0 | 0 | 0 | 1.00 | N | 0 | 0 | 0 | 0 |
| 0 | 0 | 0 | 0 | 1.00 | N | 0 | 0 | 0 | 0 |
| 0 | 0 | 0 | 0 | 1.00 | N | 0 | 0 | 0 | 0 |
| 0 | 0 | 0 | 0 | 1.00 | N | 0 | 0 | 0 | 0 |
| 0 | 0 | 0 | 0 | 1.00 | N | 0 | 0 | 0 | 0 |
| 0 | 0 | 0 | 0 | 1.00 | N | 0 | 0 | 0 | 0 |
| 0 | 0 | 0 | 0 | 1.00 | N | 0 | 0 | 0 | 0 |
| 0 | 0 | 0 | 0 | 1.00 | N | 0 | 0 | 0 | 0 |
| 0 | 0 | 0 | 0 | 1.00 | N | 0 | 0 | 0 | 0 |
| 0 | 0 | 0 | 0 | 1.00 | N | 0 | 0 | 0 | 0 |
| 0 | 0 | 0 | 0 | 1.00 | N | 0 | 0 | 0 | 0 |
| 0 | 0 | 0 | 0 | 1.00 | N | 0 | 0 | 0 | 0 |
| 0 | 0 | 0 | 0 | 1.00 | N | 0 | 0 | 0 | 0 |
| 0 | 0 | 0 | 0 | 1.00 | N | 0 | 0 | 0 | 0 |
| 0 | 0 | 0 | 0 | 1.00 | N | 0 | 0 | 0 | 0 |
| 0 | 0 | 0 | 0 | 1.00 | N | 0 | 0 | 0 | 0 |
| 0 | 0 | 0 | 0 | 1.00 | N | 0 | 0 | 0 | 0 |
| 0 | 0 | 0 | 0 | 1.00 | N | 0 | 0 | 0 | 0 |
| 0 | 0 | 0 | 0 | 1.00 | N | 0 | 0 | 0 | 0 |
| 0 | 0 | 0 | 0 | 1.00 | N | 0 | 0 | 0 | 0 |
| 0 | 0 | 0 | 0 | 1.00 | N | 0 | 0 | 0 | 0 |
| 0 | 0 | 0 | 0 | 1.00 | N | 0 | 0 | 0 | 0 |

| | | | | | | | | | |
|---|---|---|---|---|---|---|---|---|---|
| 0 | 0 | 0 | 0 | 1.00 | N | 0 | 0 | 0 | 0 |
| 0 | 0 | 0 | 0 | 1.00 | N | 0 | 0 | 0 | 0 |
| 0 | 0 | 0 | 0 | 1.00 | N | 0 | 0 | 0 | 0 |
| 0 | 0 | 0 | 0 | 1.00 | N | 0 | 0 | 0 | 0 |
| 0 | 0 | 0 | 0 | 1.00 | N | 0 | 0 | 0 | 0 |
| 0 | 0 | 0 | 0 | 1.00 | N | 0 | 0 | 0 | 0 |
| 0 | 0 | 0 | 0 | 1.00 | N | 0 | 0 | 0 | 0 |
| 0 | 0 | 0 | 0 | 1.00 | N | 0 | 0 | 0 | 0 |
| 0 | 0 | 0 | 0 | 1.00 | N | 0 | 0 | 0 | 0 |
| 0 | 0 | 0 | 0 | 1.00 | N | 0 | 0 | 0 | 0 |
| 2 | 0 | 5 | 5 | 0.84 | N | 0 | 0 | 0 | 0 |
| 2 | 1 | 5 | 6 | 0.50 | N | 1 | 0 | 0 | 1 |
| 1 | 0 | 6 | 6 | 0.50 | N | 0 | 0 | 0 | 1 |
| 2 | 0 | 4 | 4 | 0.69 | N | 0 | 1 | 0 | 1 |
| 2 | 1 | 3 | 4 | 0.89 | N | 0 | 0 | 0 | 0 |
| 2 | 0 | 6 | 6 | 0.63 | Y | 0 | 0 | 0 | 2 |
| 2 | 1 | 6 | 7 | 0.60 | N | 0 | 2 | 0 | 2 |
| 2 | 2 | 6 | 8 | 0.64 | N | 1 | 0 | 0 | 2 |
| 2 | 0 | 8 | 8 | 0.53 | N | 0 | 2 | 0 | 2 |
| 2 | 2 | 5 | 7 | 0.52 | Y | 1 | 1 | 0 | 2 |
| 2 | 1 | 5 | 6 | 0.51 | Y | 1 | 1 | 0 | 2 |
| 2 | 2 | 7 | 9 | 0.48 | Y | 2 | 0 | 0 | 2 |
| 2 | 1 | 8 | 9 | 0.49 | Y | 1 | 1 | 0 | 2 |
| 1 | 2 | 7 | 9 | 0.59 | Y | 1 | 1 | 1 | 1 |
| 2 | 0 | 5 | 5 | 0.79 | N | 0 | 0 | 0 | 0 |
| 2 | 1 | 5 | 6 | 0.60 | N | 1 | 0 | 0 | 1 |
| 2 | 2 | 4 | 6 | 0.53 | N | 0 | 2 | 0 | 2 |
| 2 | 1 | 5 | 6 | 0.47 | Y | 2 | 0 | 1 | 1 |
| 1 | 0 | 6 | 6 | 0.52 | N | 0 | 0 | 0 | 2 |
| 2 | 1 | 5 | 6 | 0.57 | N | 1 | 0 | 0 | 1 |

| | | | | | | | | | |
|---|---|---|---|---|---|---|---|---|---|
| 2 | 4 | 7 | 11 | 0.64 | Y | 0 | 2 | 0 | 2 |
| 2 | 3 | 6 | 9 | 0.54 | Y | 2 | 0 | 0 | 2 |
| 2 | 2 | 7 | 9 | 0.61 | N | 0 | 2 | 0 | 2 |
| 2 | 0 | 4 | 4 | 0.55 | N | 0 | 1 | 0 | 1 |
| 2 | 0 | 5 | 5 | 0.61 | N | 0 | 1 | 0 | 1 |
| 1 | 1 | 5 | 6 | 0.49 | N | 0 | 1 | 0 | 2 |
| 1 | 1 | 5 | 6 | 0.60 | N | 1 | 0 | 0 | 2 |
| 2 | 0 | 6 | 6 | 0.67 | Y | 0 | 0 | 0 | 2 |
| 1 | 1 | 5 | 6 | 0.64 | N | 0 | 2 | 0 | 2 |
| 2 | 1 | 6 | 7 | 0.65 | Y | 0 | 0 | 0 | 2 |
| 2 | 3 | 5 | 8 | 0.58 | N | 1 | 0 | 0 | 1 |
| 2 | 1 | 6 | 7 | 0.62 | N | 0 | 2 | 0 | 2 |
| 2 | 0 | 7 | 7 | 0.53 | N | 0 | 2 | 0 | 2 |
| 2 | 1 | 6 | 7 | 0.52 | N | 1 | 1 | 0 | 2 |
| 2 | 3 | 4 | 7 | 0.66 | Y | 1 | 1 | 0 | 2 |
| 2 | 1 | 7 | 8 | 0.51 | N | 1 | 1 | 0 | 2 |
| 2 | 1 | 8 | 9 | 0.53 | Y | 1 | 1 | 0 | 2 |
| 2 | 2 | 6 | 8 | 0.59 | Y | 1 | 1 | 0 | 2 |
| 2 | 2 | 4 | 6 | 0.47 | Y | 1 | 1 | 0 | 2 |
| 2 | 2 | 6 | 8 | 0.48 | Y | 1 | 1 | 0 | 2 |
| 2 | 1 | 5 | 6 | 0.51 | Y | 2 | 0 | 0 | 2 |
| 2 | 1 | 7 | 8 | 0.52 | Y | 1 | 1 | 0 | 2 |
| 2 | 1 | 7 | 8 | 0.44 | Y | 0 | 2 | 0 | 2 |
| 2 | 4 | 6 | 10 | 0.67 | Y | 0 | 2 | 0 | 2 |
| 2 | 2 | 9 | 11 | 0.56 | Y | 1 | 1 | 1 | 1 |
| 2 | 1 | 6 | 7 | 0.49 | Y | 0 | 2 | 0 | 2 |
| 2 | 2 | 7 | 9 | 0.49 | N | 1 | 1 | 0 | 2 |
| 2 | 3 | 6 | 9 | 0.53 | N | 1 | 1 | 0 | 2 |
| 2 | 3 | 7 | 10 | 0.61 | Y | 0 | 2 | 0 | 2 |
| 2 | 2 | 7 | 9 | 0.50 | N | 0 | 2 | 0 | 2 |

| | | | | | | | | | |
|---|---|---|---|---|---|---|---|---|---|
| 2 | 3 | 7 | 10 | 0.60 | N | 1 | 0 | 0 | 2 |
| 2 | 1 | 10 | 11 | 0.60 | Y | 0 | 2 | 0 | 2 |
| 2 | 3 | 6 | 9 | 0.50 | Y | 1 | 1 | 0 | 2 |
| 0 | 3 | 4 | 7 | 0.70 | N | 0 | 2 | 0 | 2 |
| 2 | 2 | 7 | 9 | 0.44 | N | 1 | 1 | 0 | 2 |
| 1 | 0 | 3 | 3 | 0.82 | N | 0 | 0 | 0 | 0 |
| 0 | 1 | 2 | 3 | 0.92 | N | 0 | 0 | 0 | 0 |
| 2 | 0 | 4 | 4 | 0.59 | N | 0 | 0 | 0 | 0 |
| 1 | 0 | 3 | 3 | 0.94 | N | 0 | 0 | 0 | 0 |
| 1 | 0 | 3 | 3 | 0.69 | N | 0 | 0 | 0 | 0 |
| 1 | 1 | 2 | 3 | 0.88 | N | 0 | 0 | 0 | 0 |
| 2 | 0 | 3 | 3 | 1.00 | N | 0 | 1 | 0 | 0 |
| 2 | 0 | 4 | 4 | 0.90 | N | 0 | 0 | 0 | 0 |
| 2 | 0 | 4 | 4 | 0.88 | N | 0 | 0 | 0 | 0 |
| 2 | 0 | 2 | 2 | 0.88 | N | 0 | 0 | 0 | 0 |
| 0 | 0 | 0 | 0 | 0.49 | N | 0 | 0 | 0 | 0 |
| 2 | 0 | 2 | 2 | 1.00 | N | 0 | 1 | 0 | 0 |
| 2 | 0 | 4 | 4 | 0.66 | N | 0 | 0 | 0 | 0 |
| 1 | 0 | 3 | 3 | 0.68 | N | 0 | 0 | 0 | 0 |
| 2 | 0 | 6 | 6 | 0.90 | Y | 0 | 0 | 0 | 0 |
| 2 | 0 | 6 | 6 | 0.63 | N | 0 | 0 | 0 | 0 |
| 2 | 0 | 4 | 4 | 0.69 | N | 0 | 0 | 0 | 0 |
| 1 | 0 | 4 | 4 | 0.63 | N | 0 | 0 | 0 | 2 |
| 1 | 1 | 3 | 4 | 0.73 | N | 0 | 0 | 0 | 0 |
| 2 | 0 | 4 | 4 | 0.53 | N | 0 | 0 | 0 | 0 |
| 2 | 0 | 4 | 4 | 0.51 | N | 0 | 0 | 0 | 0 |
| 2 | 0 | 5 | 5 | 0.53 | N | 0 | 0 | 0 | 0 |
| 1 | 0 | 2 | 2 | 0.51 | N | 0 | 0 | 0 | 0 |
| 2 | 2 | 6 | 8 | 0.58 | N | 0 | 0 | 0 | 0 |
| 2 | 1 | 4 | 5 | 0.63 | N | 1 | 0 | 0 | 0 |

| | | | | | | | | | |
|---|---|---|---|---|---|---|---|---|---|
| 2 | 1 | 7 | 8 | 0.68 | N | 0 | 1 | 0 | 0 |
| 2 | 1 | 5 | 6 | 0.61 | N | 1 | 0 | 1 | 0 |
| 2 | 0 | 6 | 6 | 0.60 | N | 0 | 0 | 0 | 2 |
| 1 | 1 | 4 | 5 | 0.59 | Y | 0 | 0 | 0 | 0 |
| 2 | 0 | 6 | 6 | 0.63 | Y | 0 | 0 | 0 | 2 |
| 2 | 0 | 5 | 5 | 0.70 | Y | 0 | 0 | 0 | 1 |
| 1 | 0 | 3 | 3 | 0.81 | N | 0 | 0 | 0 | 0 |
| 2 | 0 | 7 | 7 | 0.59 | N | 0 | 1 | 0 | 1 |
| 2 | 0 | 3 | 3 | 0.90 | N | 0 | 0 | 0 | 0 |
| 2 | 0 | 4 | 4 | 0.66 | N | 0 | 0 | 0 | 0 |
| 1 | 1 | 4 | 5 | 0.53 | Y | 0 | 0 | 0 | 0 |
| 2 | 0 | 6 | 6 | 0.59 | Y | 0 | 0 | 0 | 2 |
| 2 | 0 | 4 | 4 | 0.90 | N | 0 | 0 | 0 | 0 |
| 0 | 1 | 4 | 5 | 0.71 | Y | 1 | 0 | 0 | 2 |
| 1 | 0 | 2 | 2 | 0.59 | N | 0 | 0 | 0 | 0 |
| 2 | 0 | 7 | 7 | 0.61 | Y | 0 | 1 | 0 | 2 |
| 2 | 0 | 8 | 8 | 0.65 | N | 0 | 2 | 0 | 0 |
| 1 | 2 | 5 | 7 | 0.59 | N | 0 | 2 | 0 | 0 |
| 2 | 0 | 6 | 6 | 0.73 | N | 0 | 0 | 0 | 0 |
| 2 | 2 | 6 | 8 | 0.58 | N | 0 | 0 | 0 | 2 |
| 2 | 2 | 6 | 8 | 0.59 | N | 0 | 0 | 0 | 2 |
| 2 | 0 | 3 | 3 | 1.00 | N | 0 | 1 | 0 | 0 |
| 2 | 0 | 4 | 4 | 0.62 | N | 0 | 0 | 0 | 0 |
| 2 | 0 | 6 | 6 | 0.58 | N | 0 | 0 | 0 | 2 |
| 2 | 2 | 6 | 8 | 0.90 | N | 0 | 0 | 0 | 0 |
| 1 | 2 | 6 | 8 | 0.53 | N | 0 | 1 | 0 | 2 |
| 2 | 0 | 8 | 8 | 0.58 | N | 0 | 0 | 0 | 2 |
| 2 | 2 | 6 | 8 | 0.55 | N | 0 | 0 | 0 | 2 |
| 1 | 0 | 4 | 4 | 0.84 | N | 0 | 0 | 0 | 2 |
| 1 | 0 | 4 | 4 | 0.62 | N | 0 | 0 | 0 | 0 |

| | | | | | | | | | |
|---|---|---|---|---|---|---|---|---|---|
| 2 | 0 | 4 | 4 | 0.44 | N | 0 | 0 | 0 | 0 |
| 2 | 2 | 5 | 7 | 0.64 | N | 1 | 0 | 0 | 0 |
| 1 | 2 | 3 | 5 | 0.63 | Y | 0 | 0 | 0 | 0 |
| 2 | 2 | 7 | 9 | 0.63 | Y | 2 | 0 | 0 | 1 |
| 2 | 0 | 8 | 8 | 0.58 | N | 2 | 0 | 0 | 2 |
| 2 | 0 | 8 | 8 | 0.60 | N | 0 | 0 | 0 | 2 |
| 2 | 0 | 8 | 8 | 0.58 | Y | 0 | 0 | 0 | 2 |
| 2 | 2 | 6 | 8 | 0.51 | Y | 0 | 0 | 0 | 2 |
| 2 | 2 | 6 | 8 | 0.92 | N | 1 | 0 | 0 | 2 |
| 2 | 0 | 6 | 6 | 0.57 | Y | 0 | 0 | 0 | 2 |
| 2 | 0 | 6 | 6 | 0.60 | N | 0 | 1 | 0 | 1 |
| 1 | 2 | 6 | 8 | 0.55 | N | 0 | 1 | 0 | 2 |
| 2 | 2 | 5 | 7 | 0.64 | Y | 1 | 0 | 0 | 1 |
| 2 | 0 | 7 | 7 | 0.63 | Y | 0 | 1 | 0 | 2 |
| 2 | 2 | 6 | 8 | 0.58 | N | 0 | 0 | 0 | 2 |
| 2 | 0 | 8 | 8 | 0.63 | N | 0 | 0 | 0 | 2 |
| 2 | 0 | 8 | 8 | 0.70 | N | 0 | 0 | 0 | 2 |
| 2 | 0 | 8 | 8 | 0.57 | N | 2 | 0 | 0 | 2 |
| 2 | 1 | 7 | 8 | 0.52 | Y | 0 | 0 | 0 | 2 |
| 2 | 1 | 9 | 10 | 0.78 | Y | 1 | 1 | 0 | 2 |
| 1 | 2 | 6 | 8 | 0.56 | N | 0 | 1 | 0 | 2 |
| 2 | 2 | 6 | 8 | 0.61 | N | 0 | 0 | 0 | 2 |
| 2 | 0 | 9 | 9 | 0.81 | Y | 1 | 1 | 0 | 2 |

| IE-Olecranon F. | IE-Olecranon D. | IE-Ischium | IE-Femur F. | IE-Femur D. | IE-Tibia F. | IE-Tibia D. | IE-Calcaneus F. | IE-Calcaneus D. | IE-Total Fractures |
|---|---|---|---|---|---|---|---|---|---|
| 0 | 0 | 0 | 0 | 0 | 0 | 0 | 0 | 0 | 0 |
| 0 | 0 | 0 | 0 | 0 | 0 | 0 | 0 | 0 | 0 |
| 0 | 0 | 0 | 0 | 0 | 0 | 0 | 0 | 0 | 0 |
| 0 | 0 | 0 | 0 | 0 | 0 | 0 | 0 | 0 | 0 |
| 0 | 0 | 0 | 0 | 0 | 0 | 0 | 0 | 0 | 0 |
| 0 | 0 | 0 | 0 | 0 | 0 | 0 | 0 | 0 | 0 |
| 0 | 0 | 0 | 0 | 0 | 0 | 0 | 0 | 0 | 0 |
| 0 | 0 | 0 | 0 | 0 | 0 | 0 | 0 | 0 | 0 |
| 0 | 0 | 0 | 0 | 0 | 0 | 0 | 0 | 0 | 0 |
| 0 | 0 | 0 | 0 | 0 | 0 | 0 | 0 | 0 | 0 |
| 0 | 0 | 0 | 0 | 0 | 0 | 0 | 0 | 0 | 0 |
| 0 | 0 | 0 | 0 | 0 | 0 | 0 | 0 | 0 | 0 |
| 0 | 0 | 0 | 0 | 0 | 0 | 0 | 0 | 0 | 0 |
| 0 | 0 | 0 | 0 | 0 | 0 | 0 | 0 | 0 | 0 |
| 0 | 0 | 0 | 0 | 0 | 0 | 0 | 0 | 0 | 0 |
| 0 | 0 | 0 | 0 | 0 | 0 | 0 | 0 | 0 | 0 |
| 0 | 0 | 0 | 0 | 0 | 0 | 0 | 0 | 0 | 0 |
| 0 | 0 | 0 | 0 | 0 | 0 | 0 | 0 | 0 | 0 |
| 0 | 0 | 0 | 0 | 0 | 0 | 0 | 0 | 0 | 0 |
| 0 | 0 | 0 | 0 | 0 | 0 | 0 | 0 | 0 | 0 |
| 0 | 0 | 0 | 0 | 0 | 0 | 0 | 0 | 0 | 0 |
| 0 | 0 | 0 | 0 | 0 | 0 | 0 | 0 | 0 | 0 |
| 0 | 0 | 0 | 0 | 0 | 0 | 0 | 0 | 0 | 0 |
| 0 | 0 | 0 | 0 | 0 | 0 | 0 | 0 | 0 | 0 |
| 0 | 0 | 0 | 0 | 0 | 0 | 0 | 0 | 0 | 0 |
| 0 | 0 | 0 | 0 | 0 | 0 | 0 | 0 | 0 | 0 |
| 0 | 0 | 0 | 0 | 0 | 0 | 0 | 0 | 0 | 0 |
| 0 | 0 | 0 | 0 | 0 | 0 | 0 | 0 | 0 | 0 |
| 0 | 0 | 0 | 0 | 0 | 0 | 0 | 0 | 0 | 0 |

| | | | | | | | | | |
|---|---|---|---|---|---|---|---|---|---|
| 0 | 0 | 0 | 0 | 0 | 0 | 0 | 0 | 0 | 0 |
| 0 | 0 | 0 | 0 | 0 | 0 | 0 | 0 | 0 | 0 |
| 0 | 0 | 0 | 0 | 0 | 0 | 0 | 0 | 0 | 0 |
| 0 | 0 | 0 | 0 | 0 | 0 | 0 | 0 | 0 | 0 |
| 0 | 0 | 0 | 0 | 0 | 0 | 0 | 0 | 0 | 0 |
| 0 | 0 | 0 | 0 | 0 | 0 | 0 | 0 | 0 | 0 |
| 0 | 0 | 0 | 0 | 0 | 0 | 0 | 0 | 0 | 0 |
| 0 | 0 | 0 | 0 | 0 | 0 | 0 | 0 | 0 | 0 |
| 0 | 0 | 0 | 0 | 0 | 0 | 0 | 0 | 0 | 0 |
| 0 | 0 | 0 | 0 | 0 | 0 | 0 | 0 | 0 | 0 |
| 0 | 0 | 0 | 0 | 0 | 0 | 0 | 0 | 0 | 0 |
| 0 | 0 | 0 | 0 | 0 | 0 | 0 | 0 | 0 | 0 |
| 0 | 0 | 0 | 0 | 0 | 0 | 0 | 0 | 0 | 0 |
| 0 | 0 | 0 | 0 | 0 | 0 | 0 | 0 | 0 | 0 |
| 0 | 0 | 0 | 0 | 0 | 0 | 0 | 0 | 0 | 0 |
| 0 | 0 | 0 | 0 | 0 | 0 | 0 | 0 | 0 | 0 |
| 0 | 0 | 0 | 0 | 0 | 0 | 0 | 0 | 0 | 0 |
| 0 | 0 | 0 | 0 | 0 | 0 | 0 | 0 | 0 | 0 |
| 0 | 0 | 0 | 0 | 0 | 0 | 0 | 0 | 0 | 0 |
| 0 | 0 | 0 | 0 | 0 | 0 | 0 | 0 | 0 | 0 |
| 0 | 0 | 0 | 0 | 0 | 0 | 0 | 0 | 0 | 0 |
| 0 | 0 | 0 | 0 | 0 | 0 | 0 | 0 | 0 | 0 |
| 0 | 0 | 0 | 0 | 0 | 0 | 0 | 0 | 0 | 0 |
| 0 | 0 | 0 | 0 | 0 | 0 | 0 | 0 | 0 | 0 |
| 0 | 0 | 0 | 0 | 0 | 0 | 0 | 0 | 0 | 0 |
| 0 | 0 | 0 | 0 | 0 | 0 | 0 | 0 | 0 | 0 |
| 0 | 0 | 0 | 0 | 0 | 0 | 0 | 0 | 0 | 0 |
| 0 | 0 | 0 | 0 | 0 | 0 | 0 | 0 | 0 | 0 |
| 0 | 0 | 0 | 0 | 0 | 0 | 0 | 0 | 0 | 0 |
| 0 | 0 | 0 | 0 | 0 | 0 | 0 | 0 | 0 | 0 |

| | | | | | | | | | |
|---|---|---|---|---|---|---|---|---|---|
| 0 | 0 | 0 | 0 | 0 | 0 | 0 | 0 | 0 | 0 |
| 0 | 0 | 0 | 0 | 0 | 0 | 0 | 0 | 0 | 0 |
| 0 | 0 | 0 | 0 | 0 | 0 | 0 | 0 | 0 | 0 |
| 0 | 0 | 0 | 0 | 0 | 0 | 0 | 0 | 0 | 0 |
| 0 | 0 | 0 | 0 | 0 | 0 | 0 | 0 | 0 | 0 |
| 0 | 0 | 0 | 0 | 0 | 0 | 0 | 0 | 0 | 0 |
| 0 | 0 | 0 | 0 | 0 | 0 | 0 | 0 | 0 | 0 |
| 0 | 0 | 0 | 0 | 0 | 0 | 0 | 0 | 0 | 0 |
| 0 | 0 | 0 | 0 | 0 | 0 | 0 | 0 | 0 | 0 |
| 0 | 0 | 0 | 0 | 0 | 0 | 0 | 0 | 0 | 0 |
| 0 | 0 | 1 | 0 | 0 | 0 | 1 | 0 | 2 | 0 |
| 0 | 0 | 2 | 0 | 0 | 0 | 0 | 0 | 2 | 1 |
| 0 | 2 | 2 | 0 | 0 | 0 | 0 | 0 | 1 | 0 |
| 0 | 0 | 2 | 0 | 0 | 0 | 1 | 0 | 2 | 0 |
| 0 | 0 | 2 | 0 | 0 | 1 | 0 | 0 | 2 | 1 |
| 0 | 0 | 2 | 0 | 0 | 0 | 0 | 0 | 2 | 0 |
| 0 | 0 | 2 | 0 | 0 | 0 | 0 | 0 | 2 | 0 |
| 0 | 0 | 2 | 0 | 0 | 0 | 1 | 0 | 1 | 1 |
| 0 | 0 | 2 | 0 | 0 | 0 | 0 | 0 | 2 | 0 |
| 0 | 1 | 2 | 0 | 0 | 0 | 0 | 0 | 2 | 1 |
| 0 | 0 | 2 | 0 | 0 | 1 | 0 | 0 | 2 | 2 |
| 0 | 0 | 2 | 0 | 0 | 1 | 0 | 0 | 2 | 3 |
| 0 | 0 | 2 | 0 | 0 | 1 | 0 | 0 | 2 | 2 |
| 0 | 0 | 2 | 0 | 0 | 0 | 2 | 0 | 1 | 2 |
| 0 | 0 | 1 | 0 | 0 | 0 | 1 | 0 | 2 | 0 |
| 0 | 0 | 2 | 0 | 0 | 0 | 0 | 0 | 2 | 1 |
| 0 | 0 | 2 | 0 | 0 | 0 | 2 | 0 | 2 | 0 |
| 0 | 0 | 2 | 0 | 0 | 1 | 0 | 0 | 2 | 4 |
| 0 | 0 | 2 | 0 | 0 | 0 | 0 | 0 | 1 | 0 |
| 0 | 0 | 2 | 0 | 0 | 0 | 0 | 0 | 2 | 1 |

| | | | | | | | | | |
|---|---|---|---|---|---|---|---|---|---|
| 0 | 1 | 2 | 0 | 0 | 0 | 2 | 0 | 2 | 0 |
| 0 | 0 | 1 | 1 | 0 | 2 | 0 | 0 | 2 | 5 |
| 0 | 0 | 2 | 0 | 1 | 0 | 2 | 0 | 2 | 0 |
| 0 | 0 | 2 | 0 | 0 | 0 | 1 | 0 | 2 | 0 |
| 0 | 0 | 2 | 0 | 0 | 0 | 1 | 0 | 2 | 0 |
| 0 | 0 | 2 | 1 | 0 | 0 | 0 | 0 | 1 | 1 |
| 0 | 0 | 2 | 0 | 0 | 0 | 1 | 0 | 1 | 1 |
| 0 | 0 | 2 | 0 | 0 | 0 | 0 | 2 | 0 | 0 |
| 0 | 0 | 0 | 0 | 0 | 0 | 2 | 1 | 1 | 1 |
| 0 | 1 | 2 | 0 | 0 | 0 | 0 | 0 | 2 | 0 |
| 0 | 0 | 2 | 0 | 0 | 1 | 1 | 0 | 2 | 2 |
| 0 | 0 | 2 | 1 | 0 | 0 | 0 | 0 | 2 | 1 |
| 0 | 0 | 2 | 0 | 0 | 0 | 0 | 0 | 2 | 0 |
| 0 | 0 | 2 | 0 | 0 | 0 | 0 | 0 | 2 | 1 |
| 0 | 0 | 2 | 0 | 0 | 1 | 0 | 0 | 2 | 2 |
| 0 | 1 | 2 | 0 | 0 | 0 | 0 | 0 | 2 | 1 |
| 0 | 0 | 2 | 0 | 0 | 1 | 0 | 0 | 2 | 2 |
| 0 | 0 | 2 | 0 | 0 | 1 | 0 | 0 | 2 | 2 |
| 0 | 0 | 2 | 0 | 0 | 1 | 0 | 0 | 2 | 2 |
| 0 | 0 | 2 | 0 | 0 | 1 | 0 | 0 | 2 | 2 |
| 0 | 0 | 2 | 0 | 0 | 1 | 0 | 0 | 2 | 3 |
| 0 | 1 | 2 | 0 | 0 | 0 | 0 | 0 | 2 | 1 |
| 0 | 0 | 2 | 0 | 0 | 0 | 1 | 0 | 2 | 0 |
| 0 | 0 | 2 | 0 | 0 | 2 | 0 | 0 | 2 | 2 |
| 0 | 0 | 2 | 0 | 0 | 1 | 2 | 0 | 1 | 3 |
| 0 | 0 | 2 | 0 | 0 | 1 | 1 | 0 | 2 | 1 |
| 0 | 0 | 2 | 0 | 0 | 0 | 2 | 0 | 2 | 1 |
| 0 | 1 | 2 | 0 | 0 | 2 | 0 | 0 | 2 | 3 |
| 0 | 1 | 2 | 0 | 0 | 1 | 1 | 0 | 2 | 1 |
| 0 | 0 | 2 | 0 | 0 | 2 | 0 | 0 | 2 | 2 |

| | | | | | | | | | |
|---|---|---|---|---|---|---|---|---|---|
| 0 | 1 | 2 | 0 | 0 | 0 | 2 | 0 | 2 | 1 |
| 0 | 1 | 2 | 0 | 0 | 0 | 2 | 0 | 2 | 0 |
| 0 | 0 | 2 | 1 | 0 | 1 | 1 | 0 | 2 | 2 |
| 0 | 0 | 2 | 0 | 1 | 0 | 2 | 0 | 2 | 0 |
| 0 | 0 | 2 | 1 | 0 | 0 | 2 | 0 | 2 | 2 |
| 0 | 0 | 2 | 0 | 0 | 0 | 0 | 0 | 1 | 0 |
| 0 | 0 | 2 | 0 | 0 | 0 | 0 | 0 | 2 | 0 |
| 0 | 0 | 2 | 0 | 0 | 0 | 0 | 0 | 1 | 0 |
| 0 | 0 | 2 | 0 | 0 | 0 | 0 | 0 | 1 | 0 |
| 0 | 0 | 2 | 0 | 0 | 0 | 0 | 0 | 2 | 0 |
| 0 | 0 | 2 | 0 | 0 | 0 | 0 | 0 | 2 | 0 |
| 0 | 0 | 0 | 0 | 0 | 0 | 0 | 0 | 2 | 0 |
| 0 | 0 | 2 | 0 | 0 | 0 | 0 | 0 | 2 | 0 |
| 0 | 0 | 2 | 0 | 0 | 0 | 0 | 0 | 2 | 0 |
| 0 | 0 | 2 | 0 | 0 | 0 | 0 | 0 | 2 | 0 |
| 0 | 0 | 2 | 0 | 1 | 0 | 0 | 0 | 1 | 0 |
| 0 | 0 | 0 | 0 | 0 | 0 | 0 | 0 | 2 | 0 |
| 0 | 0 | 2 | 0 | 0 | 0 | 0 | 0 | 2 | 0 |
| 0 | 2 | 0 | 0 | 0 | 0 | 0 | 1 | 1 | 1 |
| 0 | 0 | 2 | 0 | 0 | 0 | 0 | 0 | 2 | 0 |
| 0 | 0 | 2 | 0 | 0 | 0 | 0 | 0 | 1 | 0 |
| 0 | 0 | 2 | 0 | 0 | 0 | 0 | 0 | 2 | 0 |
| 0 | 0 | 2 | 0 | 0 | 0 | 0 | 0 | 1 | 0 |
| 0 | 0 | 2 | 1 | 0 | 0 | 0 | 0 | 1 | 1 |
| 0 | 0 | 2 | 0 | 0 | 0 | 0 | 0 | 2 | 0 |
| 0 | 0 | 2 | 0 | 0 | 0 | 0 | 0 | 2 | 0 |
| 0 | 1 | 2 | 0 | 0 | 0 | 0 | 1 | 0 | 1 |
| 0 | 1 | 2 | 0 | 0 | 0 | 0 | 0 | 2 | 0 |
| 0 | 0 | 2 | 0 | 0 | 1 | 0 | 0 | 1 | 1 |
| 0 | 0 | 2 | 0 | 0 | 0 | 0 | 0 | 2 | 1 |

| | | | | | | | | | |
|---|---|---|---|---|---|---|---|---|---|
| 0 | 0 | 2 | 0 | 0 | 0 | 0 | 0 | 2 | 0 |
| 0 | 2 | 0 | 0 | 0 | 0 | 0 | 0 | 2 | 1 |
| 0 | 0 | 2 | 0 | 0 | 0 | 0 | 2 | 0 | 2 |
| 0 | 0 | 2 | 0 | 0 | 0 | 2 | 0 | 2 | 0 |
| 0 | 0 | 2 | 0 | 0 | 0 | 0 | 0 | 2 | 0 |
| 0 | 0 | 2 | 0 | 0 | 0 | 1 | 0 | 2 | 0 |
| 0 | 0 | 2 | 0 | 0 | 0 | 0 | 0 | 1 | 0 |
| 0 | 0 | 2 | 0 | 0 | 0 | 0 | 0 | 2 | 0 |
| 0 | 0 | 2 | 0 | 0 | 0 | 0 | 0 | 2 | 0 |
| 0 | 0 | 2 | 0 | 0 | 0 | 0 | 0 | 1 | 0 |
| 0 | 0 | 2 | 0 | 0 | 1 | 1 | 0 | 1 | 2 |
| 0 | 0 | 2 | 0 | 0 | 0 | 0 | 0 | 2 | 0 |
| 0 | 0 | 2 | 0 | 0 | 0 | 0 | 0 | 2 | 0 |
| 0 | 0 | 2 | 0 | 0 | 0 | 0 | 0 | 2 | 1 |
| 0 | 1 | 2 | 0 | 0 | 0 | 0 | 0 | 1 | 0 |
| 0 | 0 | 2 | 0 | 0 | 0 | 0 | 0 | 2 | 0 |
| 0 | 0 | 2 | 0 | 0 | 1 | 1 | 0 | 1 | 1 |
| 0 | 0 | 2 | 0 | 0 | 1 | 1 | 1 | 0 | 2 |
| 0 | 0 | 2 | 0 | 0 | 0 | 0 | 0 | 1 | 0 |
| 0 | 1 | 2 | 0 | 0 | 0 | 0 | 0 | 2 | 2 |
| 0 | 0 | 2 | 0 | 0 | 2 | 0 | 0 | 2 | 2 |
| 0 | 0 | 0 | 0 | 0 | 0 | 0 | 0 | 2 | 0 |
| 0 | 0 | 2 | 0 | 0 | 0 | 0 | 0 | 2 | 0 |
| 0 | 0 | 2 | 0 | 0 | 0 | 2 | 0 | 2 | 0 |
| 0 | 0 | 2 | 0 | 0 | 0 | 0 | 0 | 2 | 0 |
| 0 | 0 | 2 | 0 | 0 | 0 | 2 | 0 | 1 | 0 |
| 0 | 0 | 2 | 0 | 0 | 0 | 2 | 0 | 2 | 0 |
| 0 | 0 | 2 | 0 | 0 | 0 | 2 | 0 | 2 | 0 |
| 0 | 0 | 2 | 0 | 0 | 0 | 0 | 0 | 2 | 0 |
| 0 | 0 | 2 | 0 | 1 | 0 | 0 | 0 | 1 | 0 |

| | | | | | | | | | |
|---|---|---|---|---|---|---|---|---|---|
| 0 | 0 | 2 | 0 | 0 | 0 | 0 | 0 | 2 | 0 |
| 0 | 0 | 2 | 0 | 0 | 0 | 0 | 0 | 2 | 1 |
| 0 | 0 | 2 | 0 | 0 | 1 | 1 | 0 | 1 | 1 |
| 0 | 0 | 2 | 0 | 0 | 0 | 1 | 0 | 2 | 2 |
| 0 | 0 | 2 | 0 | 0 | 0 | 0 | 0 | 2 | 2 |
| 0 | 0 | 2 | 0 | 0 | 0 | 2 | 0 | 2 | 0 |
| 0 | 0 | 2 | 0 | 0 | 0 | 2 | 0 | 2 | 0 |
| 0 | 0 | 2 | 0 | 0 | 2 | 0 | 0 | 1 | 2 |
| 0 | 1 | 1 | 0 | 0 | 1 | 0 | 0 | 2 | 2 |
| 0 | 0 | 2 | 0 | 0 | 0 | 0 | 0 | 2 | 0 |
| 0 | 0 | 2 | 0 | 0 | 0 | 0 | 0 | 2 | 0 |
| 0 | 0 | 2 | 0 | 0 | 0 | 2 | 0 | 1 | 0 |
| 0 | 0 | 2 | 0 | 0 | 0 | 1 | 0 | 2 | 1 |
| 0 | 0 | 2 | 0 | 0 | 0 | 0 | 0 | 2 | 0 |
| 0 | 1 | 2 | 0 | 0 | 2 | 0 | 0 | 2 | 2 |
| 0 | 0 | 2 | 0 | 0 | 0 | 2 | 0 | 1 | 0 |
| 0 | 0 | 2 | 0 | 0 | 0 | 2 | 0 | 2 | 0 |
| 0 | 0 | 2 | 0 | 0 | 0 | 0 | 0 | 2 | 2 |
| 0 | 0 | 2 | 0 | 0 | 0 | 2 | 0 | 2 | 0 |
| 0 | 1 | 1 | 0 | 0 | 1 | 1 | 0 | 2 | 2 |
| 0 | 0 | 2 | 0 | 0 | 0 | 2 | 0 | 1 | 0 |
| 0 | 0 | 2 | 0 | 0 | 0 | 2 | 0 | 2 | 0 |
| 0 | 1 | 1 | 0 | 0 | 1 | 0 | 1 | 1 | 3 |

| IE-Total Deformities | IE-Total Score | IE-Pelvis Ratio | IE-Scoliosis | KC-Humerus F | KC-Humerus D | KC-Forearm (Ulna & | KC-Forearm (Ulna & | KC-Olecranon F | KC-Olecranon D |
|---|---|---|---|---|---|---|---|---|---|
| 0 | 0 | 1 | N | 0 | 0 | 0 | 0 | 0 | 0 |
| 0 | 0 | 1 | N | 0 | 0 | 0 | 0 | 0 | 0 |
| 0 | 0 | 1 | N | 0 | 0 | 0 | 0 | 0 | 0 |
| 0 | 0 | 1 | N | 0 | 0 | 0 | 0 | 0 | 0 |
| 0 | 0 | 1 | N | 0 | 0 | 0 | 0 | 0 | 0 |
| 0 | 0 | 1 | N | 0 | 0 | 0 | 0 | 0 | 0 |
| 0 | 0 | 1 | N | 0 | 0 | 0 | 0 | 0 | 0 |
| 0 | 0 | 1 | N | 0 | 0 | 0 | 0 | 0 | 0 |
| 0 | 0 | 1 | N | 0 | 0 | 0 | 0 | 0 | 0 |
| 0 | 0 | 1 | N | 0 | 0 | 0 | 0 | 0 | 0 |
| 0 | 0 | 1 | N | 0 | 0 | 0 | 0 | 0 | 0 |
| 0 | 0 | 1 | N | 0 | 0 | 0 | 0 | 0 | 0 |
| 0 | 0 | 1 | N | 0 | 0 | 0 | 0 | 0 | 0 |
| 0 | 0 | 1 | N | 0 | 0 | 0 | 0 | 0 | 0 |
| 0 | 0 | 1 | N | 0 | 0 | 0 | 0 | 0 | 0 |
| 0 | 0 | 1 | N | 0 | 0 | 0 | 0 | 0 | 0 |
| 0 | 0 | 1 | N | 0 | 0 | 0 | 0 | 0 | 0 |
| 0 | 0 | 1 | N | 0 | 0 | 0 | 0 | 0 | 0 |
| 0 | 0 | 1 | N | 0 | 0 | 0 | 0 | 0 | 0 |
| 0 | 0 | 1 | N | 0 | 0 | 0 | 0 | 0 | 0 |
| 0 | 0 | 1 | N | 0 | 0 | 0 | 0 | 0 | 0 |
| 0 | 0 | 1 | N | 0 | 0 | 0 | 0 | 0 | 0 |
| 0 | 0 | 1 | N | 0 | 0 | 0 | 0 | 0 | 0 |
| 0 | 0 | 1 | N | 0 | 0 | 0 | 0 | 0 | 0 |
| 0 | 0 | 1 | N | 0 | 0 | 0 | 0 | 0 | 0 |
| 0 | 0 | 1 | N | 0 | 0 | 0 | 0 | 0 | 0 |
| 0 | 0 | 1 | N | 0 | 0 | 0 | 0 | 0 | 0 |
| 0 | 0 | 1 | N | 0 | 0 | 0 | 0 | 0 | 0 |
| 0 | 0 | 1 | N | 0 | 0 | 0 | 0 | 0 | 0 |

| | | | | | | | | | |
|---|---|---|---|---|---|---|---|---|---|
| 0 | 0 | 1 | N | 0 | 0 | 0 | 0 | 0 | 0 |
| 0 | 0 | 1 | N | 0 | 0 | 0 | 0 | 0 | 0 |
| 0 | 0 | 1 | N | 0 | 0 | 0 | 0 | 0 | 0 |
| 0 | 0 | 1 | N | 0 | 0 | 0 | 0 | 0 | 0 |
| 0 | 0 | 1 | N | 0 | 0 | 0 | 0 | 0 | 0 |
| 0 | 0 | 1 | N | 0 | 0 | 0 | 0 | 0 | 0 |
| 0 | 0 | 1 | N | 0 | 0 | 0 | 0 | 0 | 0 |
| 0 | 0 | 1 | N | 0 | 0 | 0 | 0 | 0 | 0 |
| 0 | 0 | 1 | N | 0 | 0 | 0 | 0 | 0 | 0 |
| 0 | 0 | 1 | N | 0 | 0 | 0 | 0 | 0 | 0 |
| 0 | 0 | 1 | N | 0 | 0 | 0 | 0 | 0 | 0 |
| 0 | 0 | 1 | N | 0 | 0 | 0 | 0 | 0 | 0 |
| 0 | 0 | 1 | N | 0 | 0 | 0 | 0 | 0 | 0 |
| 0 | 0 | 1 | N | 0 | 0 | 0 | 0 | 0 | 0 |
| 0 | 0 | 1 | N | 0 | 0 | 0 | 0 | 0 | 0 |
| 0 | 0 | 1 | N | 0 | 0 | 0 | 0 | 0 | 0 |
| 0 | 0 | 1 | N | 0 | 0 | 0 | 0 | 0 | 0 |
| 0 | 0 | 1 | N | 0 | 0 | 0 | 0 | 0 | 0 |
| 0 | 0 | 1 | N | 0 | 0 | 0 | 0 | 0 | 0 |
| 0 | 0 | 1 | N | 0 | 0 | 0 | 0 | 0 | 0 |
| 0 | 0 | 1 | N | 0 | 0 | 0 | 0 | 0 | 0 |
| 0 | 0 | 1 | N | 0 | 0 | 0 | 0 | 0 | 0 |
| 0 | 0 | 1 | N | 0 | 0 | 0 | 0 | 0 | 0 |
| 0 | 0 | 1 | N | 0 | 0 | 0 | 0 | 0 | 0 |
| 0 | 0 | 1 | N | 0 | 0 | 0 | 0 | 0 | 0 |
| 0 | 0 | 1 | N | 0 | 0 | 0 | 0 | 0 | 0 |
| 0 | 0 | 1 | N | 0 | 0 | 0 | 0 | 0 | 0 |
| 0 | 0 | 1 | N | 0 | 0 | 0 | 0 | 0 | 0 |
| 0 | 0 | 1 | N | 0 | 0 | 0 | 0 | 0 | 0 |
| 0 | 0 | 1 | N | 0 | 0 | 0 | 0 | 0 | 0 |

| | | | | | | | | | |
|---|---|---|---|---|---|---|---|---|---|
| 0 | 0 | 1 | N | 0 | 0 | 0 | 0 | 0 | 0 |
| 0 | 0 | 1 | N | 0 | 0 | 0 | 0 | 0 | 0 |
| 0 | 0 | 1 | N | 0 | 0 | 0 | 0 | 0 | 0 |
| 0 | 0 | 1 | N | 0 | 0 | 0 | 0 | 0 | 0 |
| 0 | 0 | 1 | N | 0 | 0 | 0 | 0 | 0 | 0 |
| 0 | 0 | 1 | N | 0 | 0 | 0 | 0 | 0 | 0 |
| 0 | 0 | 1 | N | 0 | 0 | 0 | 0 | 0 | 0 |
| 0 | 0 | 1 | N | 0 | 0 | 0 | 0 | 0 | 0 |
| 0 | 0 | 1 | N | 0 | 0 | 0 | 0 | 0 | 0 |
| 0 | 0 | 1 | N | 0 | 0 | 0 | 0 | 0 | 0 |
| 5 | 5 | 0.84 | N | 0 | 0 | 0 | 0 | 0 | 0 |
| 5 | 6 | 0.61 | N | 0 | 1 | 1 | 0 | 0 | 0 |
| 6 | 6 | 0.5 | N | 0 | 0 | 0 | 2 | 0 | 0 |
| 7 | 7 | 0.67 | N | 0 | 1 | 0 | 1 | 0 | 0 |
| 4 | 5 | 0.82 | N | 0 | 0 | 0 | 0 | 0 | 0 |
| 6 | 6 | 0.53 | Y | 0 | 0 | 0 | 2 | 0 | 0 |
| 8 | 8 | 0.46 | N | 0 | 0 | 0 | 2 | 0 | 0 |
| 6 | 7 | 0.51 | N | 1 | 0 | 0 | 2 | 0 | 0 |
| 8 | 8 | 0.63 | N | 0 | 2 | 0 | 2 | 0 | 0 |
| 8 | 9 | 0.63 | Y | 1 | 1 | 0 | 2 | 0 | 0 |
| 7 | 9 | 0.52 | N | 0 | 2 | 0 | 2 | 0 | 0 |
| 6 | 9 | 0.5 | Y | 1 | 1 | 0 | 2 | 0 | 0 |
| 7 | 9 | 0.53 | N | 2 | 0 | 0 | 2 | 0 | 0 |
| 7 | 9 | 0.61 | N | 2 | 0 | 1 | 1 | 0 | 0 |
| 5 | 5 | 0.8 | N | 0 | 0 | 0 | 0 | 0 | 0 |
| 5 | 6 | 0.6 | N | 0 | 1 | 0 | 1 | 0 | 0 |
| 10 | 10 | 0.55 | N | 0 | 2 | 0 | 2 | 0 | 0 |
| 5 | 9 | 0.52 | N | 0 | 2 | 1 | 1 | 0 | 0 |
| 5 | 5 | 0.41 | N | 0 | 0 | 0 | 2 | 0 | 0 |
| 5 | 6 | 0.61 | N | 0 | 1 | 0 | 1 | 0 | 0 |

| | | | | | | | | | |
|---|---|---|---|---|---|---|---|---|---|
| 11 | 11 | 0.68 | Y | 0 | 2 | 0 | 2 | 0 | 0 |
| 6 | 11 | 0.49 | Y | 1 | 1 | 0 | 2 | 0 | 0 |
| 11 | 11 | 0.63 | N | 1 | 1 | 1 | 1 | 0 | 0 |
| 7 | 7 | 0.66 | N | 0 | 1 | 0 | 1 | 0 | 0 |
| 7 | 7 | 0.73 | N | 0 | 1 | 0 | 1 | 0 | 0 |
| 6 | 7 | 0.49 | N | 0 | 0 | 0 | 2 | 0 | 0 |
| 6 | 7 | 0.54 | N | 0 | 0 | 0 | 2 | 0 | 0 |
| 6 | 6 | 0.54 | Y | 0 | 0 | 0 | 2 | 0 | 0 |
| 7 | 8 | 0.59 | N | 0 | 0 | 0 | 2 | 0 | 0 |
| 7 | 7 | 0.59 | Y | 0 | 0 | 0 | 2 | 0 | 0 |
| 6 | 8 | 0.68 | N | 1 | 0 | 0 | 1 | 0 | 0 |
| 8 | 9 | 0.5 | N | 0 | 0 | 0 | 2 | 0 | 0 |
| 8 | 8 | 0.57 | N | 0 | 2 | 0 | 2 | 0 | 0 |
| 7 | 8 | 0.55 | N | 1 | 1 | | 2 | 0 | 0 |
| 7 | 9 | 0.63 | N | 0 | 1 | 0 | 2 | 0 | 0 |
| 8 | 9 | 0.55 | Y | 1 | 1 | 0 | 2 | 0 | 1 |
| 7 | 9 | 0.51 | N | 1 | 1 | 0 | 2 | 0 | 0 |
| 7 | 9 | 0.58 | N | 1 | 1 | 0 | 2 | 0 | 0 |
| 7 | 9 | 0.5 | N | 0 | 2 | 0 | 2 | 0 | 0 |
| 7 | 9 | 0.56 | Y | 1 | 1 | 0 | 2 | 0 | 0 |
| 6 | 9 | 0.53 | N | 1 | 1 | 0 | 2 | 0 | 0 |
| 8 | 9 | 0.5 | Y | 1 | 1 | 0 | 2 | 0 | 0 |
| 9 | 9 | 0.47 | Y | 0 | 2 | 0 | 2 | 0 | 0 |
| 8 | 10 | 0.62 | N | 0 | 2 | 0 | 2 | 0 | 0 |
| 7 | 10 | 0.66 | N | 1 | 1 | 1 | 1 | 0 | 0 |
| 9 | 10 | 0.5 | N | 0 | 2 | 0 | 2 | 0 | 0 |
| 9 | 10 | 0.54 | N | 1 | 1 | 0 | 2 | 0 | 0 |
| 8 | 11 | 0.65 | Y | 1 | 0 | 0 | 2 | 0 | 0 |
| 10 | 11 | 0.68 | Y | 1 | 1 | 0 | 2 | 0 | 1 |
| 8 | 10 | 0.57 | N | 0 | 2 | 0 | 2 | 0 | 0 |

| | | | | | | | | | |
|---|---|---|---|---|---|---|---|---|---|
| 9 | 10 | 0.71 | Y | 1 | 0 | 0 | 2 | 0 | 1 |
| 11 | 11 | 0.68 | Y | 0 | 1 | 0 | 2 | 0 | 0 |
| 8 | 10 | 0.48 | y | 1 | 1 | 1 | 1 | 0 | 0 |
| 11 | 11 | 0.62 | N | 1 | 1 | 1 | 1 | 0 | 0 |
| 9 | 11 | 0.63 | N | 1 | 1 | 1 | 1 | 0 | 0 |
| 3 | 3 | 0.68 | N | 0 | 0 | 0 | 0 | 0 | 0 |
| 4 | 4 | 0.78 | N | 0 | 0 | 0 | 0 | 0 | 0 |
| 3 | 3 | 0.68 | N | 0 | 0 | 0 | 0 | 0 | 0 |
| 3 | 3 | 0.81 | N | 0 | 0 | 0 | 0 | 0 | 0 |
| 4 | 4 | 0.62 | N | 0 | 0 | 0 | 0 | 0 | 0 |
| 4 | 4 | 0.88 | N | 0 | 0 | 0 | 0 | 0 | 0 |
| 3 | 3 | 0.76 | N | 0 | 0 | 0 | 0 | 0 | 0 |
| 4 | 4 | 0.88 | N | 0 | 0 | 0 | 0 | 0 | 0 |
| 4 | 4 | 0.81 | N | 0 | 0 | 0 | 0 | 0 | 0 |
| 4 | 4 | 1 | N | | 0 | 0 | 0 | 0 | 0 |
| 4 | 4 | 0.56 | N | 0 | 0 | 0 | 0 | 0 | 0 |
| 3 | 3 | 0.68 | N | 0 | 0 | 0 | 0 | 0 | 0 |
| 4 | 4 | 0.68 | N | 0 | 0 | 0 | 0 | 0 | 0 |
| 3 | 4 | 0.66 | N | 0 | 0 | 0 | 0 | 0 | 0 |
| 4 | 4 | 0.86 | N | 0 | 0 | 0 | 0 | 0 | 0 |
| 3 | 3 | 0.62 | N | 0 | 0 | 0 | 0 | 0 | 0 |
| 4 | 4 | 0.67 | N | 0 | 0 | 0 | 0 | 0 | 0 |
| 5 | 5 | 0.61 | N | 0 | 0 | 0 | 0 | 0 | 0 |
| 3 | 4 | 0.56 | N | 0 | 0 | 0 | 0 | 0 | 0 |
| 4 | 4 | 0.59 | N | 0 | 0 | 0 | 0 | 0 | 0 |
| 4 | 4 | 0.58 | N | 0 | 0 | 0 | 0 | 0 | 0 |
| 3 | 4 | 0.6 | N | 0 | 0 | 0 | 0 | 0 | 1 |
| 5 | 5 | 0.46 | N | 0 | 0 | 0 | 0 | 1 | 0 |
| 3 | 4 | 0.61 | N | 0 | 0 | 0 | 0 | 0 | 0 |
| 4 | 5 | 0.58 | Y | 1 | 0 | 0 | 0 | 0 | 0 |

| | | | | | | | | | |
|---|---|---|---|---|---|---|---|---|---|
| 5 | 5 | 0.63 | Y | 0 | 1 | 0 | 0 | 0 | 0 |
| 5 | 6 | 0.57 | N | 1 | 0 | 0 | 1 | 0 | 0 |
| 4 | 6 | 0.66 | N | 0 | 0 | 0 | 2 | 0 | 0 |
| 6 | 6 | 0.55 | N | 0 | 0 | 0 | 0 | 0 | 0 |
| 6 | 6 | 0.62 | N | 0 | 0 | 0 | 2 | 0 | 0 |
| 6 | 6 | 0.64 | Y | 0 | 0 | 0 | 1 | 0 | 0 |
| 3 | 3 | 0.83 | N | 0 | 0 | 0 | 0 | 0 | 0 |
| 6 | 6 | 0.49 | N | 0 | 1 | 0 | 1 | 0 | 0 |
| 4 | 4 | 0.89 | N | 0 | 0 | 0 | 0 | 0 | 0 |
| 3 | 3 | 0.73 | N | 0 | 0 | 0 | 0 | 0 | 0 |
| 4 | 6 | 0.58 | N | 0 | 0 | 0 | 0 | 0 | 1 |
| 6 | 6 | 0.59 | N | 0 | 0 | 0 | 2 | 0 | 0 |
| 4 | 4 | 0.84 | N | 0 | 0 | 0 | 0 | 0 | 0 |
| 6 | 7 | 0.68 | N | 1 | 0 | 0 | 2 | 0 | 0 |
| 4 | 4 | 0.58 | N | 0 | 0 | 0 | 0 | 1 | 0 |
| 7 | 7 | 0.61 | N | 0 | 1 | 0 | 2 | 0 | 0 |
| 6 | 7 | 0.65 | N | 0 | 0 | 0 | 2 | 0 | 0 |
| 5 | 7 | 0.59 | N | 0 | 0 | 0 | 2 | 0 | 0 |
| 3 | 3 | 0.58 | N | 0 | 0 | 0 | 0 | 0 | 0 |
| 7 | 7 | 0.56 | N | 0 | 0 | 0 | 2 | 0 | 0 |
| 6 | 8 | 0.59 | N | 0 | 0 | 0 | 2 | 0 | 0 |
| 3 | 3 | 0.77 | N | 0 | 0 | 0 | 0 | 0 | 0 |
| 4 | 4 | 0.63 | N | 0 | 0 | 0 | 0 | 0 | 0 |
| 8 | 8 | 0.58 | N | 0 | 0 | 0 | 2 | 0 | 0 |
| 4 | 4 | 0.86 | N | 0 | 0 | 0 | 0 | 0 | 0 |
| 8 | 8 | 0.56 | N | 0 | 0 | 0 | 2 | 0 | 0 |
| 8 | 8 | 0.57 | N | 0 | 0 | 0 | 2 | 0 | 0 |
| 8 | 8 | 0.53 | N | 0 | 0 | 0 | 2 | 0 | 0 |
| 6 | 6 | 0.57 | N | 0 | 0 | 0 | 0 | 0 | 0 |
| 4 | 4 | 0.56 | N | 0 | 0 | 0 | 0 | 0 | 0 |

| | | | | | | | | | |
|---|---|---|---|---|---|---|---|---|---|
| 4 | 4 | 0.58 | N | 0 | 0 | 0 | 0 | 0 | 0 |
| 4 | 5 | 0.63 | Y | 1 | 0 | 0 | 0 | 0 | 0 |
| 4 | 5 | 0.61 | N | 0 | 0 | 0 | 0 | 0 | 0 |
| 6 | 8 | 0.65 | Y | 1 | 1 | 0 | 1 | 0 | 0 |
| 6 | 8 | 0.62 | N | 2 | 0 | 0 | 2 | 0 | 0 |
| 8 | 8 | 0.6 | Y | 0 | 0 | 0 | 2 | 0 | 0 |
| 8 | 8 | 0.64 | N | 0 | 0 | 0 | 2 | 0 | 0 |
| 5 | 7 | 0.67 | N | 0 | 0 | 0 | 2 | 0 | 0 |
| 6 | 8 | 0.85 | N | 1 | 0 | 0 | 2 | 0 | 0 |
| 6 | 6 | 0.54 | N | 0 | 0 | 0 | 2 | 0 | 0 |
| 6 | 6 | 0.55 | N | 0 | 1 | 0 | 1 | 0 | 0 |
| 8 | 8 | 0.5 | N | 0 | 1 | 0 | 2 | 0 | 0 |
| 6 | 7 | 0.64 | Y | 1 | 0 | 0 | 1 | 0 | 0 |
| 7 | 7 | 0.63 | N | 1 | 0 | 0 | 2 | 0 | 0 |
| 7 | 9 | 0.55 | N | 0 | 0 | 0 | 2 | 0 | 0 |
| 7 | 7 | 0.63 | N | 0 | 0 | 0 | 2 | 0 | 0 |
| 8 | 8 | 0.57 | N | 0 | 0 | 0 | 2 | 0 | 0 |
| 6 | 8 | 0.64 | N | 2 | 0 | 0 | 2 | 0 | 0 |
| 8 | 8 | 0.52 | N | 0 | 0 | 0 | 2 | 0 | 0 |
| 8 | 10 | 0.78 | N | 1 | 1 | 0 | 2 | 0 | 1 |
| 8 | 8 | 0.61 | N | 1 | 0 | 0 | 2 | 0 | 0 |
| 8 | 8 | 0.64 | N | 0 | 0 | 0 | 2 | 0 | 0 |
| 6 | 9 | 0.83 | N | 1 | 1 | 0 | 2 | 0 | 1 |

| KC-Ischium | KC-Femur F. | KC-Femur D. | KC-Tibia F. | KC-Tibia D. | KC-Calcaneous F | KC-Calcaneous D | KC-Total Fractures | KC-Total Deformities | KC-Total Score |
|---|---|---|---|---|---|---|---|---|---|
| 0 | 0 | 0 | 0 | 0 | 0 | 0 | 0 | 0 | 0 |
| 0 | 0 | 0 | 0 | 0 | 0 | 0 | 0 | 0 | 0 |
| 0 | 0 | 0 | 0 | 0 | 0 | 0 | 0 | 0 | 0 |
| 0 | 0 | 0 | 0 | 0 | 0 | 0 | 0 | 0 | 0 |
| 0 | 0 | 0 | 0 | 0 | 0 | 0 | 0 | 0 | 0 |
| 0 | 0 | 0 | 0 | 0 | 0 | 0 | 0 | 0 | 0 |
| 0 | 0 | 0 | 0 | 0 | 0 | 0 | 0 | 0 | 0 |
| 0 | 0 | 0 | 0 | 0 | 0 | 0 | 0 | 0 | 0 |
| 0 | 0 | 0 | 0 | 0 | 0 | 0 | 0 | 0 | 0 |
| 0 | 0 | 0 | 0 | 0 | 0 | 0 | 0 | 0 | 0 |
| 0 | 0 | 0 | 0 | 0 | 0 | 0 | 0 | 0 | 0 |
| 0 | 0 | 0 | 0 | 0 | 0 | 0 | 0 | 0 | 0 |
| 0 | 0 | 0 | 0 | 0 | 0 | 0 | 0 | 0 | 0 |
| 0 | 0 | 0 | 0 | 0 | 0 | 0 | 0 | 0 | 0 |
| 0 | 0 | 0 | 0 | 0 | 0 | 0 | 0 | 0 | 0 |
| 0 | 0 | 0 | 0 | 0 | 0 | 0 | 0 | 0 | 0 |
| 0 | 0 | 0 | 0 | 0 | 0 | 0 | 0 | 0 | 0 |
| 0 | 0 | 0 | 0 | 0 | 0 | 0 | 0 | 0 | 0 |
| 0 | 0 | 0 | 0 | 0 | 0 | 0 | 0 | 0 | 0 |
| 0 | 0 | 0 | 0 | 0 | 0 | 0 | 0 | 0 | 0 |
| 0 | 0 | 0 | 0 | 0 | 0 | 0 | 0 | 0 | 0 |
| 0 | 0 | 0 | 0 | 0 | 0 | 0 | 0 | 0 | 0 |
| 0 | 0 | 0 | 0 | 0 | 0 | 0 | 0 | 0 | 0 |
| 0 | 0 | 0 | 0 | 0 | 0 | 0 | 0 | 0 | 0 |
| 0 | 0 | 0 | 0 | 0 | 0 | 0 | 0 | 0 | 0 |
| 0 | 0 | 0 | 0 | 0 | 0 | 0 | 0 | 0 | 0 |
| 0 | 0 | 0 | 0 | 0 | 0 | 0 | 0 | 0 | 0 |
| 0 | 0 | 0 | 0 | 0 | 0 | 0 | 0 | 0 | 0 |
| 0 | 0 | 0 | 0 | 0 | 0 | 0 | 0 | 0 | 0 |

| | | | | | | | | | |
|---|---|---|---|---|---|---|---|---|---|
| 0 | 0 | 0 | 0 | 0 | 0 | 0 | 0 | 0 | 0 |
| 0 | 0 | 0 | 0 | 0 | 0 | 0 | 0 | 0 | 0 |
| 0 | 0 | 0 | 0 | 0 | 0 | 0 | 0 | 0 | 0 |
| 0 | 0 | 0 | 0 | 0 | 0 | 0 | 0 | 0 | 0 |
| 0 | 0 | 0 | 0 | 0 | 0 | 0 | 0 | 0 | 0 |
| 0 | 0 | 0 | 0 | 0 | 0 | 0 | 0 | 0 | 0 |
| 0 | 0 | 0 | 0 | 0 | 0 | 0 | 0 | 0 | 0 |
| 0 | 0 | 0 | 0 | 0 | 0 | 0 | 0 | 0 | 0 |
| 0 | 0 | 0 | 0 | 0 | 0 | 0 | 0 | 0 | 0 |
| 0 | 0 | 0 | 0 | 0 | 0 | 0 | 0 | 0 | 0 |
| 0 | 0 | 0 | 0 | 0 | 0 | 0 | 0 | 0 | 0 |
| 0 | 0 | 0 | 0 | 0 | 0 | 0 | 0 | 0 | 0 |
| 0 | 0 | 0 | 0 | 0 | 0 | 0 | 0 | 0 | 0 |
| 0 | 0 | 0 | 0 | 0 | 0 | 0 | 0 | 0 | 0 |
| 0 | 0 | 0 | 0 | 0 | 0 | 0 | 0 | 0 | 0 |
| 0 | 0 | 0 | 0 | 0 | 0 | 0 | 0 | 0 | 0 |
| 0 | 0 | 0 | 0 | 0 | 0 | 0 | 0 | 0 | 0 |
| 0 | 0 | 0 | 0 | 0 | 0 | 0 | 0 | 0 | 0 |
| 0 | 0 | 0 | 0 | 0 | 0 | 0 | 0 | 0 | 0 |
| 0 | 0 | 0 | 0 | 0 | 0 | 0 | 0 | 0 | 0 |
| 0 | 0 | 0 | 0 | 0 | 0 | 0 | 0 | 0 | 0 |
| 0 | 0 | 0 | 0 | 0 | 0 | 0 | 0 | 0 | 0 |
| 0 | 0 | 0 | 0 | 0 | 0 | 0 | 0 | 0 | 0 |
| 0 | 0 | 0 | 0 | 0 | 0 | 0 | 0 | 0 | 0 |
| 0 | 0 | 0 | 0 | 0 | 0 | 0 | 0 | 0 | 0 |
| 0 | 0 | 0 | 0 | 0 | 0 | 0 | 0 | 0 | 0 |
| 0 | 0 | 0 | 0 | 0 | 0 | 0 | 0 | 0 | 0 |
| 0 | 0 | 0 | 0 | 0 | 0 | 0 | 0 | 0 | 0 |
| 0 | 0 | 0 | 0 | 0 | 0 | 0 | 0 | 0 | 0 |
| 0 | 0 | 0 | 0 | 0 | 0 | 0 | 0 | 0 | 0 |

| | | | | | | | | | |
|---|---|---|---|---|---|---|---|---|---|
| 0 | 0 | 0 | 0 | 0 | 0 | 0 | 0 | 0 | 0 |
| 0 | 0 | 0 | 0 | 0 | 0 | 0 | 0 | 0 | 0 |
| 0 | 0 | 0 | 0 | 0 | 0 | 0 | 0 | 0 | 0 |
| 0 | 0 | 0 | 0 | 0 | 0 | 0 | 0 | 0 | 0 |
| 0 | 0 | 0 | 0 | 0 | 0 | 0 | 0 | 0 | 0 |
| 0 | 0 | 0 | 0 | 0 | 0 | 0 | 0 | 0 | 0 |
| 0 | 0 | 0 | 0 | 0 | 0 | 0 | 0 | 0 | 0 |
| 0 | 0 | 0 | 0 | 0 | 0 | 0 | 0 | 0 | 0 |
| 0 | 0 | 0 | 0 | 0 | 0 | 0 | 0 | 0 | 0 |
| 0 | 0 | 0 | 0 | 0 | 0 | 0 | 0 | 0 | 0 |
| 2 | 0 | 0 | 0 | 0 | 0 | 2 | 0 | 4 | 4 |
| 2 | 0 | 0 | 0 | 0 | 0 | 2 | 1 | 5 | 6 |
| 2 | 0 | 0 | 0 | 0 | 0 | 1 | 0 | 5 | 5 |
| 2 | 0 | 0 | 0 | 1 | 0 | 2 | 0 | 7 | 7 |
| 2 | 0 | 0 | 0 | 0 | 0 | 2 | 0 | 4 | 4 |
| 2 | 0 | 0 | 0 | 0 | 0 | 2 | 0 | 6 | 6 |
| 2 | 1 | 0 | 0 | 0 | 0 | 2 | 1 | 6 | 7 |
| 2 | 0 | 0 | 1 | 0 | 0 | 1 | 2 | 5 | 7 |
| 2 | 0 | 0 | 0 | 0 | 0 | 2 | 0 | 8 | 8 |
| 2 | 0 | 0 | 0 | 0 | 0 | 2 | 1 | 7 | 8 |
| 2 | 0 | 0 | 0 | 1 | 0 | 2 | 0 | 9 | 9 |
| 2 | 0 | 0 | 0 | 1 | 0 | 1 | 1 | 7 | 8 |
| 2 | 0 | 0 | 0 | 1 | 0 | 2 | 2 | 7 | 9 |
| 2 | 0 | 0 | 1 | 1 | 0 | 1 | 4 | 5 | 9 |
| 2 | 0 | 0 | 0 | 0 | 0 | 2 | 0 | 4 | 4 |
| 2 | 0 | 0 | 0 | 0 | 0 | 0 | 0 | 4 | 4 |
| 2 | 0 | 0 | 0 | 2 | 0 | 2 | 0 | 10 | 10 |
| 2 | 0 | 0 | 0 | 2 | 0 | 2 | 1 | 9 | 10 |
| 2 | 0 | 0 | 0 | 0 | 0 | 1 | 0 | 5 | 5 |
| 2 | 0 | 0 | 0 | 0 | 0 | 2 | 0 | 6 | 6 |

| | | | | | | | | | |
|---|---|---|---|---|---|---|---|---|---|
| 2 | 0 | 0 | 1 | 1 | 0 | 2 | 1 | 9 | 10 |
| 2 | 0 | 1 | 1 | 0 | 0 | 2 | 2 | 8 | 10 |
| 2 | 0 | 1 | 0 | 2 | 0 | 2 | 2 | 9 | 11 |
| 2 | 0 | 0 | 0 | 1 | 0 | 2 | 0 | 7 | 7 |
| 2 | 0 | 0 | 0 | 1 | 0 | 2 | 0 | 7 | 7 |
| 2 | 1 | 0 | 0 | 0 | 0 | 1 | 1 | 5 | 6 |
| 2 | 0 | 0 | 0 | 1 | 0 | 1 | 0 | 6 | 6 |
| 2 | 0 | 0 | 0 | 0 | 0 | 2 | 0 | 6 | 6 |
| 2 | 0 | 0 | 0 | 0 | 0 | 2 | 0 | 6 | 6 |
| 2 | 0 | 0 | 1 | 0 | 0 | 2 | 1 | 6 | 7 |
| 2 | 0 | 0 | 0 | 2 | 0 | 0 | 1 | 5 | 6 |
| 2 | 1 | 0 | 0 | 0 | 0 | 2 | 1 | 6 | 7 |
| 2 | 0 | 0 | 0 | 0 | 0 | 2 | 0 | 8 | 8 |
| 2 | 0 | 0 | 0 | 0 | 0 | 2 | 1 | 7 | 8 |
| 2 | 0 | 0 | 1 | 0 | 0 | 2 | 1 | 7 | 8 |
| 2 | 0 | 0 | 0 | 0 | 0 | 2 | 1 | 8 | 9 |
| 2 | 0 | 0 | 0 | 0 | 0 | 2 | 1 | 7 | 8 |
| 2 | 0 | 0 | 0 | 1 | 0 | 2 | 1 | 8 | 9 |
| 2 | 0 | 0 | 0 | 1 | 0 | 2 | 0 | 9 | 9 |
| 2 | 0 | 0 | 0 | 1 | 0 | 1 | 1 | 7 | 8 |
| 2 | 0 | 0 | 0 | 2 | 0 | 2 | 1 | 9 | 10 |
| 2 | 0 | 0 | 0 | 0 | 0 | 2 | 1 | 7 | 8 |
| 2 | 0 | 0 | 0 | 1 | 0 | 1 | 0 | 8 | 8 |
| 2 | 0 | 0 | 1 | 1 | 0 | 1 | 1 | 8 | 9 |
| 2 | 0 | 0 | 2 | 0 | 0 | 1 | 4 | 5 | 9 |
| 2 | 0 | 0 | 0 | 2 | 0 | 2 | 0 | 10 | 10 |
| 2 | 0 | 0 | 0 | 2 | 0 | 2 | 1 | 9 | 10 |
| 2 | 0 | 0 | 1 | 1 | 0 | 2 | 2 | 7 | 9 |
| 2 | 0 | 0 | 1 | 1 | 0 | 2 | 2 | 9 | 11 |
| 2 | 0 | 0 | 0 | 2 | 0 | 2 | 0 | 10 | 10 |

| | | | | | | | | | |
|---|---|---|---|---|---|---|---|---|---|
| 2 | 0 | 0 | 0 | 2 | 0 | 2 | 1 | 9 | 10 |
| 2 | 0 | 0 | 0 | 2 | 0 | 2 | 0 | 9 | 9 |
| 2 | 0 | 1 | 0 | 2 | 0 | 2 | 2 | 9 | 11 |
| 2 | 0 | 1 | 1 | 1 | 0 | 2 | 3 | 8 | 11 |
| 2 | 0 | 1 | 0 | 2 | 0 | 2 | 2 | 9 | 11 |
| 2 | 0 | 0 | 0 | 0 | 0 | 1 | 0 | 3 | 3 |
| 2 | 0 | 0 | 0 | 0 | 0 | 2 | 0 | 4 | 4 |
| 2 | 0 | 0 | 0 | 0 | 0 | 1 | 0 | 3 | 3 |
| 2 | 0 | 0 | 0 | 0 | 0 | 1 | 0 | 3 | 3 |
| 2 | 0 | 0 | 0 | 0 | 0 | 2 | 0 | 4 | 4 |
| 2 | 0 | 0 | 0 | 0 | 0 | 2 | 0 | 4 | 4 |
| 0 | 0 | 0 | 0 | 0 | 0 | 2 | 0 | 2 | 2 |
| 2 | 0 | 0 | 0 | 0 | 0 | 2 | 0 | 4 | 4 |
| 2 | 0 | 0 | 0 | 0 | 0 | 2 | 0 | 4 | 4 |
| 2 | 0 | 0 | 0 | 0 | 0 | 2 | 0 | 4 | 4 |
| 2 | 0 | 1 | 0 | 0 | 0 | 1 | 0 | 4 | 4 |
| 0 | 0 | 0 | 0 | 0 | 0 | 2 | 0 | 2 | 2 |
| 2 | 0 | 0 | 0 | 0 | 0 | 2 | 0 | 4 | 4 |
| 2 | 0 | 0 | 0 | 0 | 0 | 1 | 0 | 3 | 3 |
| 2 | 0 | 0 | 0 | 0 | 0 | 2 | 0 | 4 | 4 |
| 2 | 0 | 0 | 0 | 0 | 0 | 1 | 0 | 3 | 3 |
| 2 | 0 | 0 | 0 | 0 | 0 | 2 | 0 | 4 | 4 |
| 2 | 0 | 0 | 0 | 0 | 0 | 1 | 0 | 3 | 3 |
| 2 | 1 | 0 | 0 | 0 | 0 | 1 | 1 | 3 | 4 |
| 2 | 0 | 0 | 0 | 0 | 0 | 2 | 0 | 4 | 4 |
| 2 | 0 | 0 | 0 | 0 | 0 | 2 | 0 | 4 | 4 |
| 2 | 0 | 0 | 0 | 0 | 0 | 1 | 0 | 4 | 4 |
| 2 | 0 | 0 | 0 | 0 | 0 | 1 | 1 | 3 | 4 |
| 2 | 0 | 0 | 0 | 0 | 0 | 1 | 0 | 3 | 3 |
| 2 | 0 | 0 | 0 | 0 | 0 | 2 | 1 | 4 | 5 |

| | | | | | | | | | |
|---|---|---|---|---|---|---|---|---|---|
| 2 | 0 | 0 | 0 | 0 | 0 | 2 | 0 | 5 | 5 |
| 2 | 0 | 0 | 0 | 0 | 0 | 2 | 1 | 5 | 6 |
| 2 | 0 | 0 | 0 | 0 | 0 | 2 | 0 | 6 | 6 |
| 2 | 0 | 0 | 1 | 1 | 0 | 2 | 1 | 5 | 6 |
| 2 | 0 | 0 | 0 | 0 | 0 | 2 | 0 | 6 | 6 |
| 2 | 0 | 0 | 0 | 1 | 0 | 2 | 0 | 6 | 6 |
| 2 | 0 | 0 | 0 | 0 | 0 | 1 | 0 | 3 | 3 |
| 2 | 0 | 0 | 0 | 0 | 0 | 2 | 0 | 6 | 6 |
| 2 | 0 | 0 | 0 | 0 | 0 | 2 | 0 | 4 | 4 |
| 2 | 0 | 0 | 0 | 0 | 0 | 1 | 0 | 3 | 3 |
| 2 | 0 | 0 | 0 | 2 | 0 | 2 | 0 | 7 | 7 |
| 2 | 0 | 0 | 0 | 0 | 0 | 2 | 0 | 6 | 6 |
| 2 | 0 | 0 | 0 | 0 | 0 | 2 | 0 | 4 | 4 |
| 2 | 0 | 0 | 0 | 0 | 0 | 2 | 1 | 6 | 7 |
| 2 | 0 | 0 | 0 | 0 | 0 | 1 | 1 | 3 | 4 |
| 2 | 0 | 0 | 0 | 0 | 0 | 2 | 0 | 7 | 7 |
| 2 | 0 | 0 | 0 | 2 | 0 | 1 | 0 | 7 | 7 |
| 2 | 0 | 0 | 0 | 2 | 0 | 1 | 0 | 7 | 7 |
| 2 | 0 | 0 | 0 | 0 | 0 | 1 | 0 | 3 | 3 |
| 2 | 0 | 0 | 0 | 2 | 0 | 0 | 0 | 6 | 6 |
| 2 | 0 | 0 | 0 | 2 | 0 | 0 | 0 | 6 | 6 |
| 0 | 0 | 0 | 0 | 0 | 0 | 2 | 0 | 2 | 2 |
| 2 | 0 | 0 | 0 | 0 | 0 | 2 | 0 | 4 | 4 |
| 2 | 0 | 0 | 0 | 2 | 0 | 1 | 0 | 7 | 7 |
| 2 | 0 | 0 | 0 | 0 | 0 | 2 | 0 | 4 | 4 |
| 2 | 0 | 0 | 0 | 2 | 0 | 0 | 0 | 6 | 6 |
| 2 | 0 | 0 | 0 | 2 | 0 | 1 | 0 | 7 | 7 |
| 2 | 0 | 0 | 0 | 2 | 0 | 2 | 0 | 8 | 8 |
| 2 | 0 | 0 | 0 | 0 | 0 | 1 | 0 | 3 | 3 |
| 2 | 0 | 1 | 0 | 0 | 0 | 1 | 0 | 4 | 4 |

| | | | | | | | | | |
|---|---|---|---|---|---|---|---|---|---|
| 2 | 0 | 0 | 0 | 0 | 0 | 2 | 0 | 4 | 4 |
| 2 | 0 | 0 | 0 | 0 | 0 | 2 | 1 | 4 | 5 |
| 2 | 0 | 0 | 1 | 1 | 0 | 2 | 1 | 5 | 6 |
| 2 | 0 | 0 | 0 | 1 | 0 | 2 | 1 | 7 | 8 |
| 2 | 0 | 0 | 0 | 0 | 0 | 2 | 2 | 6 | 8 |
| 2 | 0 | 0 | 0 | 2 | 0 | 2 | 0 | 8 | 8 |
| 2 | 0 | 0 | 0 | 2 | 0 | 2 | 0 | 8 | 8 |
| 2 | 0 | 0 | 0 | 2 | 0 | 2 | 0 | 8 | 8 |
| 2 | 0 | 0 | 1 | 0 | 0 | 2 | 2 | 6 | 8 |
| 2 | 0 | 0 | 0 | 0 | 0 | 2 | 0 | 6 | 6 |
| 2 | 0 | 0 | 0 | 0 | 0 | 2 | 0 | 6 | 6 |
| 2 | 0 | 0 | 0 | 2 | 0 | 0 | 0 | 7 | 7 |
| 2 | 0 | 0 | 0 | 1 | 0 | 2 | 1 | 6 | 7 |
| 2 | 0 | 0 | 0 | 0 | 0 | 2 | 1 | 6 | 7 |
| 2 | 0 | 0 | 0 | 2 | 0 | 0 | 0 | 6 | 6 |
| 2 | 0 | 0 | 0 | 2 | 0 | 1 | 0 | 7 | 7 |
| 2 | 0 | 0 | 0 | 2 | 0 | 1 | 0 | 7 | 7 |
| 2 | 0 | 0 | 0 | 0 | 0 | 2 | 2 | 6 | 8 |
| 2 | 0 | 0 | 0 | 2 | 0 | 2 | 0 | 8 | 8 |
| 2 | 0 | 0 | 1 | 0 | 0 | 2 | 2 | 8 | 10 |
| 2 | 0 | 0 | 0 | 2 | 0 | 0 | 1 | 6 | 7 |
| 2 | 0 | 0 | 0 | 2 | 0 | 2 | 0 | 8 | 8 |
| 2 | 0 | 0 | 1 | 0 | 0 | 2 | 2 | 8 | 10 |

| KC-Pelvis Ratio | KC-Scoliosis | DN-Humerus [illegible] | DN-Humerus D | DN-Forearm (Ulna & [illegible] | DN-Forearm (Ulna & [illegible] | DN-Olecranon F | DN-Olecranon D | DN-Ischium | DN-Femur F. |
|---|---|---|---|---|---|---|---|---|---|
| 1 | N | 0 | 0 | 0 | 0 | 0 | 0 | 0 | 0 |
| 1 | N | 0 | 0 | 0 | 0 | 0 | 0 | 0 | 0 |
| 1 | N | 0 | 0 | 0 | 0 | 0 | 0 | 0 | 0 |
| 1 | N | 0 | 0 | 0 | 0 | 0 | 0 | 0 | 0 |
| 1 | N | 0 | 0 | 0 | 0 | 0 | 0 | 0 | 0 |
| 1 | N | 0 | 0 | 0 | 0 | 0 | 0 | 0 | 0 |
| 1 | N | 0 | 0 | 0 | 0 | 0 | 0 | 0 | 0 |
| 1 | N | 0 | 0 | 0 | 0 | 0 | 0 | 0 | 0 |
| 1 | N | 0 | 0 | 0 | 0 | 0 | 0 | 0 | 0 |
| 1 | N | 0 | 0 | 0 | 0 | 0 | 0 | 0 | 0 |
| 1 | N | 0 | 0 | 0 | 0 | 0 | 0 | 0 | 0 |
| 1 | N | 0 | 0 | 0 | 0 | 0 | 0 | 0 | 0 |
| 1 | N | 0 | 0 | 0 | 0 | 0 | 0 | 0 | 0 |
| 1 | N | 0 | 0 | 0 | 0 | 0 | 0 | 0 | 0 |
| 1 | N | 0 | 0 | 0 | 0 | 0 | 0 | 0 | 0 |
| 1 | N | 0 | 0 | 0 | 0 | 0 | 0 | 0 | 0 |
| 1 | N | 0 | 0 | 0 | 0 | 0 | 0 | 0 | 0 |
| 1 | N | 0 | 0 | 0 | 0 | 0 | 0 | 0 | 0 |
| 1 | N | 0 | 0 | 0 | 0 | 0 | 0 | 0 | 0 |
| 1 | N | 0 | 0 | 0 | 0 | 0 | 0 | 0 | 0 |
| 1 | N | 0 | 0 | 0 | 0 | 0 | 0 | 0 | 0 |
| 1 | N | 0 | 0 | 0 | 0 | 0 | 0 | 0 | 0 |
| 1 | N | 0 | 0 | 0 | 0 | 0 | 0 | 0 | 0 |
| 1 | N | 0 | 0 | 0 | 0 | 0 | 0 | 0 | 0 |
| 1 | N | 0 | 0 | 0 | 0 | 0 | 0 | 0 | 0 |
| 1 | N | 0 | 0 | 0 | 0 | 0 | 0 | 0 | 0 |
| 1 | N | 0 | 0 | 0 | 0 | 0 | 0 | 0 | 0 |
| 1 | N | 0 | 0 | 0 | 0 | 0 | 0 | 0 | 0 |
| 1 | N | 0 | 0 | 0 | 0 | 0 | 0 | 0 | 0 |

| | | | | | | | | | |
|---|---|---|---|---|---|---|---|---|---|
| 1 | N | 0 | 0 | 0 | 0 | 0 | 0 | 0 | 0 |
| 1 | N | 0 | 0 | 0 | 0 | 0 | 0 | 0 | 0 |
| 1 | N | 0 | 0 | 0 | 0 | 0 | 0 | 0 | 0 |
| 1 | N | 0 | 0 | 0 | 0 | 0 | 0 | 0 | 0 |
| 1 | N | 0 | 0 | 0 | 0 | 0 | 0 | 0 | 0 |
| 1 | N | 0 | 0 | 0 | 0 | 0 | 0 | 0 | 0 |
| 1 | N | 0 | 0 | 0 | 0 | 0 | 0 | 0 | 0 |
| 1 | N | 0 | 0 | 0 | 0 | 0 | 0 | 0 | 0 |
| 1 | N | 0 | 0 | 0 | 0 | 0 | 0 | 0 | 0 |
| 1 | N | 0 | 0 | 0 | 0 | 0 | 0 | 0 | 0 |
| 1 | N | 0 | 0 | 0 | 0 | 0 | 0 | 0 | 0 |
| 1 | N | 0 | 0 | 0 | 0 | 0 | 0 | 0 | 0 |
| 1 | N | 0 | 0 | 0 | 0 | 0 | 0 | 0 | 0 |
| 1 | N | 0 | 0 | 0 | 0 | 0 | 0 | 0 | 0 |
| 1 | N | 0 | 0 | 0 | 0 | 0 | 0 | 0 | 0 |
| 1 | N | 0 | 0 | 0 | 0 | 0 | 0 | 0 | 0 |
| 1 | N | 0 | 0 | 0 | 0 | 0 | 0 | 0 | 0 |
| 1 | N | 0 | 0 | 0 | 0 | 0 | 0 | 0 | 0 |
| 1 | N | 0 | 0 | 0 | 0 | 0 | 0 | 0 | 0 |
| 1 | N | 0 | 0 | 0 | 0 | 0 | 0 | 0 | 0 |
| 1 | N | 0 | 0 | 0 | 0 | 0 | 0 | 0 | 0 |
| 1 | N | 0 | 0 | 0 | 0 | 0 | 0 | 0 | 0 |
| 1 | N | 0 | 0 | 0 | 0 | 0 | 0 | 0 | 0 |
| 1 | N | 0 | 0 | 0 | 0 | 0 | 0 | 0 | 0 |
| 1 | N | 0 | 0 | 0 | 0 | 0 | 0 | 0 | 0 |
| 1 | N | 0 | 0 | 0 | 0 | 0 | 0 | 0 | 0 |
| 1 | N | 0 | 0 | 0 | 0 | 0 | 0 | 0 | 0 |
| 1 | N | 0 | 0 | 0 | 0 | 0 | 0 | 0 | 0 |
| 1 | N | 0 | 0 | 0 | 0 | 0 | 0 | 0 | 0 |
| 1 | N | 0 | 0 | 0 | 0 | 0 | 0 | 0 | 0 |

| | | | | | | | | | |
|---|---|---|---|---|---|---|---|---|---|
| 1 | N | 0 | 0 | 0 | 0 | 0 | 0 | 0 | 0 |
| 1 | N | 0 | 0 | 0 | 0 | 0 | 0 | 0 | 0 |
| 1 | N | 0 | 0 | 0 | 0 | 0 | 0 | 0 | 0 |
| 1 | N | 0 | 0 | 0 | 0 | 0 | 0 | 0 | 0 |
| 1 | N | 0 | 0 | 0 | 0 | 0 | 0 | 0 | 0 |
| 1 | N | 0 | 0 | 0 | 0 | 0 | 0 | 0 | 0 |
| 1 | N | 0 | 0 | 0 | 0 | 0 | 0 | 0 | 0 |
| 1 | N | 0 | 0 | 0 | 0 | 0 | 0 | 0 | 0 |
| 1 | N | 0 | 0 | 0 | 0 | 0 | 0 | 0 | 0 |
| 1 | N | 0 | 0 | 0 | 0 | 0 | 0 | 0 | 0 |
| 0.84 | Y | 0 | 0 | 0 | 1 | 0 | 0 | 1 | 0 |
| 0.58 | N | 1 | 0 | 0 | 1 | 0 | 0 | 2 | 0 |
| 0.53 | Y | 0 | 1 | 0 | 2 | 0 | 0 | 2 | 0 |
| 0.7 | N | 0 | 0 | 0 | 1 | 0 | 1 | 2 | 0 |
| 0.82 | Y | 0 | 0 | 0 | 0 | 0 | 0 | 1 | 0 |
| 0.58 | Y | 0 | 1 | 0 | 2 | 0 | 0 | 2 | 0 |
| 0.58 | Y | 0 | 1 | 0 | 2 | 0 | 0 | 2 | 0 |
| 0.53 | Y | 0 | 1 | 0 | 2 | 0 | 0 | 2 | 0 |
| 0.52 | N | 0 | 2 | 0 | 2 | 0 | 1 | 2 | 0 |
| 0.48 | Y | 0 | 2 | 0 | 2 | 0 | 1 | 2 | 0 |
| 0.49 | Y | 0 | 2 | 0 | 2 | 0 | 0 | 2 | 0 |
| 0.49 | Y | 1 | 1 | 0 | 2 | 0 | 0 | 2 | 0 |
| 0.5 | N | 2 | 0 | 0 | 2 | 0 | 0 | 2 | 0 |
| 0.59 | Y | 1 | 1 | 0 | 1 | 0 | 0 | 2 | 0 |
| 0.81 | Y | 0 | 0 | 0 | 1 | 0 | 0 | 1 | 0 |
| 0.6 | N | 0 | 1 | 0 | 1 | 0 | 0 | 2 | 0 |
| 0.56 | N | 0 | 2 | 0 | 2 | 0 | 1 | 2 | 0 |
| 0.48 | Y | 0 | 2 | 0 | 1 | 0 | 2 | 2 | 0 |
| 0.39 | Y | 0 | 1 | 0 | 2 | 0 | 0 | 2 | 0 |
| 0.54 | N | 1 | 0 | 0 | 1 | 0 | 0 | 2 | 0 |

| | | | | | | | | | |
|---|---|---|---|---|---|---|---|---|---|
| 0.62 | Y | 1 | 1 | 0 | 2 | 0 | 0 | 2 | 0 |
| 0.5 | Y | 1 | 1 | 0 | 2 | 0 | 1 | 2 | 0 |
| 0.61 | N | 1 | 1 | 0 | 2 | 0 | 0 | 2 | 0 |
| 0.71 | N | 0 | 0 | 0 | 2 | 0 | 1 | 2 | 0 |
| 0.73 | N | 0 | 0 | 0 | 0 | 0 | 1 | 2 | 0 |
| 0.51 | Y | 0 | 1 | 0 | 2 | 0 | 0 | 2 | 1 |
| 0.5 | Y | 0 | 1 | 0 | 2 | 0 | 0 | 2 | 0 |
| 0.56 | Y | 0 | 1 | 0 | 2 | 0 | 0 | 2 | 0 |
| 1.82 | Y | 0 | 1 | 0 | 2 | 0 | 0 | 2 | 0 |
| 0.57 | Y | 0 | 0 | 0 | 2 | 0 | 1 | 2 | 0 |
| 0.6 | Y | 1 | 1 | 0 | 1 | 0 | 0 | 2 | 0 |
| 0.59 | Y | 0 | 1 | 0 | 2 | 0 | 0 | 2 | 0 |
| 0.5 | N | 0 | 2 | 0 | 2 | 0 | 1 | 2 | 0 |
| 0.54 | N | 1 | 1 | 0 | 2 | 0 | 0 | 2 | 0 |
| 0.6 | Y | 0 | 2 | 0 | 2 | 0 | 0 | 2 | 0 |
| 0.58 | Y | 1 | 1 | 0 | 2 | 0 | 1 | 2 | 0 |
| 0.48 | N | 1 | 1 | 0 | 2 | 0 | 0 | 2 | 0 |
| 0.49 | N | 1 | 0 | 0 | 2 | 0 | 0 | 2 | 0 |
| 0.52 | Y | 0 | 2 | 0 | 2 | 0 | 0 | 2 | 0 |
| 0.56 | Y | 1 | 1 | 0 | 2 | 0 | 0 | 2 | 0 |
| 0.49 | Y | 2 | 0 | 0 | 1 | 0 | 2 | 2 | 0 |
| 0.44 | Y | 0 | 2 | 0 | 2 | 0 | 1 | 2 | 0 |
| 0.47 | Y | 0 | 2 | 0 | 2 | 0 | 0 | 2 | 0 |
| 0.62 | N | 1 | 1 | 0 | 2 | 0 | 0 | 2 | 0 |
| 0.53 | Y | 0 | 2 | 0 | 1 | 0 | 0 | 2 | 0 |
| 0.47 | Y | 0 | 2 | 0 | 1 | 0 | 2 | 2 | 0 |
| 0.49 | N | 1 | 1 | 0 | 2 | 0 | 1 | 2 | 0 |
| 0.53 | Y | 1 | 1 | 0 | 2 | 0 | 1 | 2 | 0 |
| 0.63 | Y | 0 | 2 | 0 | 2 | 0 | 1 | 2 | 0 |
| 0.56 | N | 0 | 2 | 0 | 2 | 0 | 1 | 2 | 0 |

| | | | | | | | | | |
|---|---|---|---|---|---|---|---|---|---|
| 0.63 | N | 1 | 0 | 0 | 2 | 0 | 1 | 2 | 0 |
| 0.61 | Y | 0 | 2 | 0 | 2 | 0 | 1 | 2 | 0 |
| 0.46 | Y | 1 | 1 | 0 | 2 | 0 | 1 | 2 | 0 |
| 0.62 | N | 1 | 1 | 1 | 2 | 0 | 0 | 2 | 0 |
| 0.58 | N | 1 | 1 | 2 | 1 | 0 | 0 | 2 | 0 |
| 0.76 | N | 0 | 0 | 0 | 0 | 0 | 0 | 2 | 0 |
| 0.95 | N | 0 | 0 | 0 | 0 | 0 | 0 | 2 | 0 |
| 0.53 | N | 0 | 0 | 0 | 0 | 0 | 0 | 2 | 0 |
| 0.78 | N | 0 | 0 | 0 | 0 | 0 | 0 | 2 | 0 |
| 0.73 | N | 0 | 0 | 0 | 0 | 0 | 0 | 2 | 0 |
| 0.86 | N | 0 | 0 | 0 | 0 | 0 | 0 | 2 | 0 |
| 1 | N | 0 | 0 | 0 | 1 | 0 | 0 | 1 | 0 |
| 0.86 | N | 0 | 0 | 0 | 0 | 0 | 0 | 2 | 0 |
| 0.83 | N | 0 | 0 | 0 | 0 | 0 | 0 | 2 | 0 |
| 0.9 | N | 0 | 0 | 0 | 0 | 0 | 0 | 2 | 0 |
| 0.5 | Y | 0 | 0 | 0 | 0 | 0 | 0 | 2 | 0 |
| 0.91 | N | 0 | 0 | 0 | 1 | 0 | 0 | 1 | 0 |
| 0.67 | N | 0 | 0 | 0 | 0 | 0 | 0 | 2 | 0 |
| 0.71 | N | 0 | 0 | 0 | 1 | 0 | 0 | 2 | 0 |
| 0.88 | N | 0 | 0 | 0 | 0 | 0 | 0 | 2 | 0 |
| 0.6 | N | 0 | 0 | 0 | 0 | 0 | 0 | 2 | 0 |
| 0.62 | N | 0 | 0 | 0 | 0 | 0 | 0 | 2 | 0 |
| 1.47 | N | 0 | 0 | 0 | 2 | 0 | 0 | 2 | 0 |
| 0.65 | Y | 0 | 0 | 0 | 0 | 0 | 0 | 2 | 1 |
| 0.5 | N | 0 | 0 | 0 | 0 | 0 | 0 | 2 | 0 |
| 0.5 | N | 0 | 0 | 0 | 0 | 0 | 0 | 2 | 0 |
| 0.65 | N | 0 | 0 | 0 | 1 | 0 | 2 | 2 | 0 |
| 0.55 | N | 0 | 0 | 0 | 1 | 0 | 0 | 2 | 0 |
| 0.64 | N | 0 | 0 | 0 | 0 | 0 | 0 | 2 | 0 |
| 1.58 | N | 0 | 0 | 0 | 1 | 0 | 0 | 2 | 0 |

| | | | | | | | | | |
|---|---|---|---|---|---|---|---|---|---|
| 0.59 | N | 2 | 0 | 0 | 1 | 0 | 0 | 2 | 0 |
| 0.59 | N | 1 | 0 | 0 | 1 | 0 | 1 | 2 | 0 |
| 0.6 | N | 0 | 0 | 0 | 2 | 0 | 0 | 2 | 0 |
| 0.59 | Y | 0 | 0 | 0 | 0 | 0 | 0 | 2 | 0 |
| 0.53 | N | 0 | 0 | 0 | 2 | 0 | 0 | 2 | 0 |
| 0.64 | Y | 0 | 0 | 0 | 1 | 0 | 0 | 2 | 0 |
| 0.78 | N | 0 | 0 | 0 | 0 | 0 | 0 | 2 | 0 |
| 0.57 | N | 0 | 1 | 0 | 1 | 0 | 1 | 2 | 0 |
| 0.86 | N | 0 | 0 | 0 | 0 | 0 | 0 | 2 | 0 |
| 0.75 | N | 0 | 0 | 0 | 0 | 0 | 0 | 2 | 0 |
| 0.53 | Y | 0 | 0 | 0 | 0 | 0 | 0 | 2 | 0 |
| 0.52 | N | 0 | 0 | 0 | 2 | 0 | 0 | 2 | 0 |
| 0.91 | N | 0 | 0 | 0 | 0 | 0 | 0 | 2 | 0 |
| 0.54 | Y | 1 | 0 | 0 | 2 | 0 | 1 | 2 | 0 |
| 0.58 | N | 0 | 0 | 0 | 0 | 0 | 1 | 2 | 0 |
| 0.55 | Y | 0 | 1 | 0 | 2 | 0 | 1 | 2 | 0 |
| 0.6 | N | 0 | 0 | 0 | 2 | 0 | 0 | 2 | 0 |
| 0.6 | N | 0 | 0 | 0 | 2 | 0 | 0 | 2 | 0 |
| 0.6 | N | 0 | 0 | 0 | 0 | 0 | 0 | 2 | 0 |
| 0.54 | N | 0 | 0 | 0 | 2 | 0 | 1 | 2 | 0 |
| 0.52 | N | 0 | 0 | 0 | 2 | 0 | 1 | 2 | 0 |
| 1 | N | 0 | 0 | 0 | 1 | 0 | 0 | 1 | 0 |
| 0.68 | N | 0 | 0 | 0 | 0 | 0 | 0 | 2 | 0 |
| 0.54 | N | 0 | 0 | 0 | 2 | 0 | 0 | 2 | 0 |
| 0.79 | N | 0 | 0 | 0 | 0 | 0 | 0 | 2 | 0 |
| 0.54 | N | 0 | 1 | 0 | 2 | 0 | 0 | 2 | 0 |
| 0.53 | N | 0 | 0 | 0 | 2 | 0 | 0 | 2 | 0 |
| 0.61 | Y | 0 | 0 | 0 | 2 | 0 | 0 | 2 | 0 |
| 1.4 | N | 0 | 0 | 0 | 2 | 0 | 0 | 2 | 0 |
| 0.6 | Y | 0 | 0 | 0 | 0 | 0 | 0 | 2 | 0 |

| | | | | | | | | | |
|---|---|---|---|---|---|---|---|---|---|
| 0.45 | N | 0 | 0 | 0 | 0 | 0 | 0 | 2 | 0 |
| 0.63 | N | 0 | 0 | 0 | 1 | 0 | 0 | 2 | 0 |
| 0.57 | Y | 0 | 0 | 0 | 0 | 0 | 0 | 2 | 0 |
| 0.65 | Y | 2 | 0 | 0 | 1 | 0 | 0 | 2 | 0 |
| 1.74 | N | 0 | 2 | 0 | 0 | 0 | 2 | 2 | 0 |
| 0.57 | Y | 0 | 0 | 0 | 2 | 0 | 0 | 2 | 0 |
| 0.53 | Y | 0 | 0 | 0 | 2 | 0 | 0 | 2 | 0 |
| 0.53 | Y | 0 | 0 | 0 | 2 | 0 | 0 | 2 | 0 |
| 0.94 | N | 1 | 0 | 0 | 2 | 0 | 1 | 1 | 0 |
| 0.55 | N | 0 | 0 | 0 | 2 | 0 | 0 | 2 | 0 |
| 0.59 | N | 0 | 1 | 0 | 1 | 0 | 1 | 2 | 0 |
| 0.5 | N | 0 | 1 | 0 | 0 | 0 | 2 | 2 | 0 |
| 0.64 | Y | 0 | 0 | 0 | 1 | 0 | 0 | 2 | 0 |
| 0.55 | Y | 0 | 1 | 0 | 2 | 0 | 1 | 2 | 0 |
| 0.45 | N | 0 | 0 | 0 | 2 | 0 | 1 | 2 | 0 |
| 0.6 | N | 0 | 0 | 0 | 2 | 0 | 0 | 2 | 0 |
| 0.55 | N | 0 | 0 | 0 | 2 | 0 | 0 | 2 | 0 |
| 0.61 | N | 2 | 0 | 0 | 2 | 0 | 0 | 2 | 0 |
| 0.53 | Y | 0 | 0 | 0 | 2 | 0 | 0 | 2 | 0 |
| 0.78 | N | 2 | 0 | 0 | 2 | 0 | 1 | 1 | 0 |
| 0.56 | N | 0 | 1 | 0 | 2 | 0 | 0 | 2 | 0 |
| 0.57 | Y | 0 | 0 | 0 | 2 | 0 | 0 | 2 | 0 |
| 0.78 | N | 2 | 0 | 0 | 2 | 0 | 1 | 1 | 0 |

| DN-Femur D. | DN-Tibia F. | DN-Tibia D. | DN-Calcaneus F. | DN-Calcaneus D. | DN-Total Fractures | DN-Total Deformities | DN-Total Score | DN-Pelvis Ratio | DN-Scoliosis |
|---|---|---|---|---|---|---|---|---|---|
| 0 | 0 | 0 | 0 | 0 | 0 | 0 | 0 | 1.00 | N |
| 0 | 0 | 0 | 0 | 0 | 0 | 0 | 0 | 1.00 | N |
| 0 | 0 | 0 | 0 | 0 | 0 | 0 | 0 | 1.00 | N |
| 0 | 0 | 0 | 0 | 0 | 0 | 0 | 0 | 1.00 | N |
| 0 | 0 | 0 | 0 | 0 | 0 | 0 | 0 | 1.00 | N |
| 0 | 0 | 0 | 0 | 0 | 0 | 0 | 0 | 1.00 | N |
| 0 | 0 | 0 | 0 | 0 | 0 | 0 | 0 | 1.00 | N |
| 0 | 0 | 0 | 0 | 0 | 0 | 0 | 0 | 1.00 | N |
| 0 | 0 | 0 | 0 | 0 | 0 | 0 | 0 | 1.00 | N |
| 0 | 0 | 0 | 0 | 0 | 0 | 0 | 0 | 1.00 | N |
| 0 | 0 | 0 | 0 | 0 | 0 | 0 | 0 | 1.00 | N |
| 0 | 0 | 0 | 0 | 0 | 0 | 0 | 0 | 1.00 | N |
| 0 | 0 | 0 | 0 | 0 | 0 | 0 | 0 | 1.00 | N |
| 0 | 0 | 0 | 0 | 0 | 0 | 0 | 0 | 1.00 | N |
| 0 | 0 | 0 | 0 | 0 | 0 | 0 | 0 | 1.00 | N |
| 0 | 0 | 0 | 0 | 0 | 0 | 0 | 0 | 1.00 | N |
| 0 | 0 | 0 | 0 | 0 | 0 | 0 | 0 | 1.00 | N |
| 0 | 0 | 0 | 0 | 0 | 0 | 0 | 0 | 1.00 | N |
| 0 | 0 | 0 | 0 | 0 | 0 | 0 | 0 | 1.00 | N |
| 0 | 0 | 0 | 0 | 0 | 0 | 0 | 0 | 1.00 | N |
| 0 | 0 | 0 | 0 | 0 | 0 | 0 | 0 | 1.00 | N |
| 0 | 0 | 0 | 0 | 0 | 0 | 0 | 0 | 1.00 | N |
| 0 | 0 | 0 | 0 | 0 | 0 | 0 | 0 | 1.00 | N |
| 0 | 0 | 0 | 0 | 0 | 0 | 0 | 0 | 1.00 | N |
| 0 | 0 | 0 | 0 | 0 | 0 | 0 | 0 | 1.00 | N |
| 0 | 0 | 0 | 0 | 0 | 0 | 0 | 0 | 1.00 | N |
| 0 | 0 | 0 | 0 | 0 | 0 | 0 | 0 | 1.00 | N |
| 0 | 0 | 0 | 0 | 0 | 0 | 0 | 0 | 1.00 | N |
| 0 | 0 | 0 | 0 | 0 | 0 | 0 | 0 | 1.00 | N |

| | | | | | | | | | |
|---|---|---|---|---|---|---|---|---|---|
| 0 | 0 | 0 | 0 | 0 | 0 | 0 | 0 | 1.00 | N |
| 0 | 0 | 0 | 0 | 0 | 0 | 0 | 0 | 1.00 | N |
| 0 | 0 | 0 | 0 | 0 | 0 | 0 | 0 | 1.00 | N |
| 0 | 0 | 0 | 0 | 0 | 0 | 0 | 0 | 1.00 | N |
| 0 | 0 | 0 | 0 | 0 | 0 | 0 | 0 | 1.00 | N |
| 0 | 0 | 0 | 0 | 0 | 0 | 0 | 0 | 1.00 | N |
| 0 | 0 | 0 | 0 | 0 | 0 | 0 | 0 | 1.00 | N |
| 0 | 0 | 0 | 0 | 0 | 0 | 0 | 0 | 1.00 | N |
| 0 | 0 | 0 | 0 | 0 | 0 | 0 | 0 | 1.00 | N |
| 0 | 0 | 0 | 0 | 0 | 0 | 0 | 0 | 1.00 | N |
| 0 | 0 | 0 | 0 | 0 | 0 | 0 | 0 | 1.00 | N |
| 0 | 0 | 0 | 0 | 0 | 0 | 0 | 0 | 1.00 | N |
| 0 | 0 | 0 | 0 | 0 | 0 | 0 | 0 | 1.00 | N |
| 0 | 0 | 0 | 0 | 0 | 0 | 0 | 0 | 1.00 | N |
| 0 | 0 | 0 | 0 | 0 | 0 | 0 | 0 | 1.00 | N |
| 0 | 0 | 0 | 0 | 0 | 0 | 0 | 0 | 1.00 | N |
| 0 | 0 | 0 | 0 | 0 | 0 | 0 | 0 | 1.00 | N |
| 0 | 0 | 0 | 0 | 0 | 0 | 0 | 0 | 1.00 | N |
| 0 | 0 | 0 | 0 | 0 | 0 | 0 | 0 | 1.00 | N |
| 0 | 0 | 0 | 0 | 0 | 0 | 0 | 0 | 1.00 | N |
| 0 | 0 | 0 | 0 | 0 | 0 | 0 | 0 | 1.00 | N |
| 0 | 0 | 0 | 0 | 0 | 0 | 0 | 0 | 1.00 | N |
| 0 | 0 | 0 | 0 | 0 | 0 | 0 | 0 | 1.00 | N |
| 0 | 0 | 0 | 0 | 0 | 0 | 0 | 0 | 1.00 | N |
| 0 | 0 | 0 | 0 | 0 | 0 | 0 | 0 | 1.00 | N |
| 0 | 0 | 0 | 0 | 0 | 0 | 0 | 0 | 1.00 | N |
| 0 | 0 | 0 | 0 | 0 | 0 | 0 | 0 | 1.00 | N |
| 0 | 0 | 0 | 0 | 0 | 0 | 0 | 0 | 1.00 | N |
| 0 | 0 | 0 | 0 | 0 | 0 | 0 | 0 | 1.00 | N |
| 0 | 0 | 0 | 0 | 0 | 0 | 0 | 0 | 1.00 | N |

| | | | | | | | | | |
|---|---|---|---|---|---|---|---|---|---|
| 0 | 0 | 0 | 0 | 0 | 0 | 0 | 0 | 1.00 | N |
| 0 | 0 | 0 | 0 | 0 | 0 | 0 | 0 | 1.00 | N |
| 0 | 0 | 0 | 0 | 0 | 0 | 0 | 0 | 1.00 | N |
| 0 | 0 | 0 | 0 | 0 | 0 | 0 | 0 | 1.00 | N |
| 0 | 0 | 0 | 0 | 0 | 0 | 0 | 0 | 1.00 | N |
| 0 | 0 | 0 | 0 | 0 | 0 | 0 | 0 | 1.00 | N |
| 0 | 0 | 0 | 0 | 0 | 0 | 0 | 0 | 1.00 | N |
| 0 | 0 | 0 | 0 | 0 | 0 | 0 | 0 | 1.00 | N |
| 0 | 0 | 0 | 0 | 0 | 0 | 0 | 0 | 1.00 | N |
| 0 | 0 | 0 | 0 | 0 | 0 | 0 | 0 | 1.00 | N |
| 0 | 1 | 0 | 0 | 2 | 1 | 4 | 5 | 0.85 | N |
| 0 | 0 | 0 | 0 | 2 | 1 | 5 | 6 | 0.56 | N |
| 0 | 0 | 0 | 0 | 1 | 0 | 6 | 6 | 0.40 | Y |
| 0 | 0 | 1 | 0 | 2 | 0 | 7 | 7 | 0.66 | N |
| 0 | 1 | 0 | 1 | 1 | 2 | 2 | 4 | 0.81 | N |
| 0 | 0 | 0 | 0 | 2 | 0 | 7 | 7 | 0.61 | Y |
| 0 | 0 | 0 | 0 | 2 | 0 | 7 | 7 | 0.58 | N |
| 0 | 1 | 0 | 0 | 2 | 1 | 7 | 8 | 0.67 | Y |
| 0 | 0 | 0 | 0 | 2 | 0 | 9 | 9 | 0.55 | N |
| 0 | 0 | 0 | 0 | 2 | 0 | 9 | 9 | 0.45 | Y |
| 0 | 0 | 1 | 0 | 2 | 0 | 9 | 9 | 0.49 | N |
| 0 | 1 | 0 | 0 | 2 | 2 | 7 | 9 | 0.51 | Y |
| 0 | 1 | 0 | 0 | 2 | 3 | 6 | 9 | 0.49 | Y |
| 0 | 1 | 1 | 0 | 1 | 2 | 6 | 8 | 0.59 | Y |
| 0 | 1 | 0 | 0 | 2 | 1 | 4 | 5 | 0.84 | N |
| 0 | 0 | 0 | 0 | 2 | 0 | 6 | 6 | 0.55 | N |
| 0 | 1 | 1 | 0 | 2 | 1 | 10 | 11 | 0.50 | N |
| 0 | 0 | 2 | 0 | 2 | 0 | 11 | 11 | 0.45 | N |
| 0 | 0 | 0 | 0 | 1 | 0 | 6 | 6 | 0.40 | Y |
| 0 | 0 | 0 | 0 | 2 | 1 | 5 | 6 | 0.59 | N |

| | | | | | | | | | |
|---|---|---|---|---|---|---|---|---|---|
| 0 | 2 | 0 | 0 | 2 | 3 | 7 | 10 | 0.58 | Y |
| 0 | 2 | 0 | 0 | 2 | 3 | 8 | 11 | 0.57 | Y |
| 1 | 2 | 0 | 0 | 2 | 3 | 8 | 11 | 0.61 | N |
| 0 | 0 | 1 | 0 | 2 | 0 | 8 | 8 | 0.66 | N |
| 0 | 0 | 1 | 1 | 1 | 1 | 5 | 6 | 0.69 | N |
| 0 | 0 | 0 | 0 | 1 | 1 | 6 | 7 | 0.52 | Y |
| 0 | 0 | 1 | 0 | 1 | 0 | 7 | 7 | 0.50 | Y |
| 0 | 0 | 0 | 0 | 2 | 0 | 7 | 7 | 0.58 | Y |
| 0 | 0 | 0 | 0 | 2 | 0 | 7 | 7 | 0.61 | N |
| 0 | 0 | 0 | 0 | 2 | 0 | 7 | 7 | 0.56 | Y |
| 0 | 2 | 0 | 0 | 1 | 3 | 5 | 8 | 0.61 | Y |
| 0 | 0 | 0 | 0 | 2 | 0 | 7 | 7 | 0.53 | N |
| 0 | 0 | 0 | 1 | 1 | 1 | 8 | 9 | 0.53 | N |
| 0 | 0 | 0 | 0 | 2 | 1 | 7 | 8 | 0.54 | N |
| 0 | 0 | 1 | 0 | 2 | 0 | 9 | 9 | 0.49 | N |
| 0 | 0 | 0 | 0 | 2 | 1 | 8 | 9 | 0.57 | Y |
| 0 | 0 | 1 | 0 | 2 | 1 | 8 | 9 | 0.43 | Y |
| 0 | 0 | 1 | 0 | 2 | 1 | 7 | 8 | 0.56 | Y |
| 0 | 0 | 1 | 0 | 2 | 0 | 9 | 9 | 0.49 | N |
| 0 | 1 | 0 | 0 | 1 | 2 | 6 | 8 | 0.51 | Y |
| 0 | 0 | 2 | 1 | 1 | 3 | 8 | 11 | 0.51 | N |
| 0 | 0 | 0 | 0 | 2 | 0 | 9 | 9 | 0.46 | Y |
| 0 | 1 | 0 | 0 | 2 | 1 | 8 | 9 | 0.49 | Y |
| 0 | 1 | 0 | 0 | 2 | 2 | 7 | 9 | 0.58 | Y |
| 0 | 0 | 2 | 0 | 1 | 0 | 8 | 8 | 0.71 | Y |
| 0 | 0 | 2 | 0 | 2 | 0 | 11 | 11 | 0.45 | N |
| 0 | 0 | 2 | 0 | 2 | 1 | 10 | 11 | 0.50 | N |
| 0 | 2 | 0 | 0 | 2 | 3 | 8 | 11 | 0.60 | Y |
| 0 | 2 | 0 | 0 | 2 | 2 | 9 | 11 | 0.62 | Y |
| 0 | 2 | 0 | 0 | 2 | 2 | 9 | 11 | 0.48 | N |

| | | | | | | | | | |
|---|---|---|---|---|---|---|---|---|---|
| 0 | 2 | 0 | 2 | 0 | 5 | 5 | 10 | 0.56 | N |
| 0 | 2 | 0 | 0 | 2 | 2 | 9 | 11 | 0.60 | Y |
| 0 | 2 | 0 | 0 | 2 | 3 | 8 | 11 | 0.53 | Y |
| 0 | 2 | 0 | 1 | 1 | 5 | 6 | 11 | 0.63 | N |
| 1 | 0 | 2 | 0 | 2 | 3 | 9 | 12 | 0.61 | N |
| 0 | 0 | 0 | 0 | 1 | 0 | 3 | 3 | 0.78 | Y |
| 0 | 0 | 0 | 0 | 2 | 0 | 4 | 4 | 0.88 | N |
| 0 | 0 | 0 | 0 | 1 | 0 | 3 | 3 | 0.75 | N |
| 0 | 0 | 0 | 0 | 1 | 0 | 3 | 3 | 0.82 | Y |
| 0 | 0 | 0 | 0 | 1 | 0 | 3 | 3 | 0.66 | N |
| 0 | 0 | 0 | 0 | 2 | 0 | 4 | 4 | 0.88 | N |
| 0 | 0 | 0 | 0 | 2 | 0 | 4 | 4 | 1.00 | N |
| 0 | 0 | 0 | 0 | 2 | 0 | 4 | 4 | 0.84 | N |
| 0 | 0 | 0 | 0 | 2 | 0 | 4 | 4 | 0.81 | N |
| 0 | 0 | 0 | 0 | 2 | 0 | 4 | 4 | 0.92 | N |
| 1 | 0 | 0 | 0 | 1 | 0 | 4 | 4 | 0.54 | Y |
| 0 | 0 | 0 | 0 | 2 | 0 | 4 | 4 | 0.87 | N |
| 0 | 0 | 0 | 0 | 2 | 0 | 4 | 4 | 0.67 | N |
| 0 | 0 | 0 | 0 | 1 | 0 | 4 | 4 | 0.55 | N |
| 0 | 0 | 0 | 0 | 2 | 0 | 4 | 4 | 0.76 | N |
| 0 | 0 | 0 | 0 | 1 | 0 | 3 | 3 | 0.54 | N |
| 0 | 0 | 0 | 1 | 1 | 1 | 3 | 4 | 0.66 | N |
| 0 | 0 | 0 | 0 | 1 | 0 | 5 | 5 | 0.59 | N |
| 0 | 0 | 0 | 0 | 1 | 1 | 3 | 4 | 0.65 | N |
| 0 | 0 | 0 | 0 | 2 | 0 | 4 | 4 | 0.50 | N |
| 0 | 0 | 0 | 0 | 2 | 0 | 4 | 4 | 0.54 | N |
| 0 | 0 | 0 | 0 | 1 | 0 | 6 | 6 | 0.53 | Y |
| 0 | 0 | 0 | 0 | 2 | 0 | 5 | 5 | 0.54 | Y |
| 0 | 0 | 0 | 0 | 2 | 0 | 4 | 4 | 0.56 | N |
| 0 | 0 | 1 | 0 | 2 | 0 | 6 | 6 | 0.51 | Y |

| | | | | | | | | | |
|---|---|---|---|---|---|---|---|---|---|
| 0 | 0 | 1 | 0 | 2 | 2 | 6 | 8 | 0.57 | Y |
| 0 | 0 | 0 | 0 | 2 | 1 | 6 | 7 | 0.60 | N |
| 0 | 0 | 0 | 0 | 2 | 0 | 6 | 6 | 0.76 | N |
| 0 | 2 | 0 | 0 | 2 | 2 | 4 | 6 | 0.61 | N |
| 0 | 0 | 0 | 0 | 2 | 0 | 6 | 6 | 0.64 | Y |
| 0 | 0 | 1 | 0 | 2 | 0 | 6 | 6 | 0.51 | Y |
| 0 | 0 | 0 | 0 | 1 | 0 | 3 | 3 | 0.84 | Y |
| 0 | 0 | 0 | 0 | 2 | 0 | 7 | 7 | 0.55 | N |
| 0 | 0 | 0 | 0 | 2 | 0 | 4 | 4 | 0.90 | N |
| 0 | 0 | 0 | 0 | 1 | 0 | 3 | 3 | 0.74 | N |
| 0 | 0 | 2 | 0 | 2 | 0 | 6 | 6 | 0.59 | N |
| 0 | 0 | 0 | 0 | 2 | 0 | 6 | 6 | 0.66 | Y |
| 0 | 0 | 0 | 0 | 2 | 0 | 4 | 4 | 0.87 | N |
| 0 | 0 | 0 | 1 | 1 | 2 | 6 | 8 | 0.58 | N |
| 0 | 0 | 0 | 0 | 1 | 0 | 4 | 4 | 0.55 | Y |
| 0 | 0 | 0 | 0 | 2 | 0 | 8 | 8 | 0.59 | N |
| 0 | 0 | 2 | 0 | 1 | 0 | 7 | 7 | 0.59 | N |
| 0 | 2 | 0 | 1 | 0 | 3 | 4 | 7 | 0.67 | N |
| 0 | 0 | 0 | 0 | 1 | 0 | 3 | 3 | 0.53 | N |
| 0 | 0 | 2 | 0 | 2 | 0 | 9 | 9 | 0.50 | N |
| 0 | 0 | 2 | 0 | 2 | 0 | 9 | 9 | 0.55 | N |
| 0 | 0 | 0 | 0 | 2 | 0 | 4 | 4 | 0.90 | N |
| 0 | 0 | 0 | 0 | 2 | 0 | 4 | 4 | 0.69 | N |
| 0 | 0 | 2 | 0 | 2 | 0 | 8 | 8 | 0.52 | N |
| 0 | 0 | 0 | 0 | 2 | 0 | 4 | 4 | 0.81 | N |
| 0 | 2 | 0 | 0 | 2 | 2 | 7 | 9 | 0.50 | N |
| 0 | 0 | 2 | 0 | 2 | 0 | 8 | 8 | 0.61 | N |
| 0 | 0 | 2 | 0 | 2 | 0 | 8 | 8 | 0.61 | Y |
| 0 | 0 | 0 | 0 | 1 | 0 | 5 | 5 | 0.59 | N |
| 1 | 0 | 0 | 0 | 1 | 0 | 4 | 4 | 0.53 | Y |

| | | | | | | | | | |
|---|---|---|---|---|---|---|---|---|---|
| 0 | 0 | 0 | 0 | 2 | 0 | 4 | 4 | 0.49 | N |
| 0 | 0 | 1 | 0 | 2 | 0 | 6 | 6 | 0.52 | Y |
| 0 | 2 | 0 | 0 | 2 | 2 | 4 | 6 | 0.59 | N |
| 0 | 0 | 1 | 0 | 2 | 2 | 6 | 8 | 0.57 | Y |
| 0 | 0 | 0 | 0 | 2 | 0 | 8 | 8 | 0.64 | N |
| 0 | 0 | 2 | 0 | 2 | 0 | 8 | 8 | 0.54 | Y |
| 0 | 0 | 2 | 0 | 2 | 0 | 8 | 8 | 0.53 | Y |
| 0 | 2 | 0 | 0 | 2 | 2 | 6 | 8 | 0.56 | Y |
| 0 | 1 | 0 | 1 | 1 | 3 | 5 | 8 | 0.91 | N |
| 0 | 0 | 0 | 0 | 2 | 0 | 6 | 6 | 0.62 | Y |
| 0 | 0 | 0 | 0 | 2 | 0 | 7 | 7 | 0.55 | N |
| 0 | 0 | 2 | 0 | 2 | 0 | 9 | 9 | 0.59 | N |
| 0 | 0 | 1 | 0 | 2 | 0 | 6 | 6 | 0.52 | Y |
| 0 | 0 | 0 | 0 | 2 | 0 | 8 | 8 | 0.60 | N |
| 0 | 0 | 2 | 0 | 2 | 0 | 9 | 9 | 0.53 | N |
| 0 | 2 | 0 | 0 | 1 | 2 | 5 | 7 | 0.64 | N |
| 0 | 0 | 2 | 0 | 2 | 0 | 8 | 8 | 0.57 | N |
| 0 | 0 | 0 | 0 | 2 | 2 | 6 | 8 | 0.70 | N |
| 0 | 0 | 2 | 0 | 2 | 0 | 8 | 8 | 0.52 | Y |
| 0 | 1 | 0 | 1 | 1 | 4 | 5 | 9 | 0.84 | N |
| 0 | 0 | 2 | 0 | 2 | 0 | 9 | 9 | 0.58 | N |
| 0 | 0 | 2 | 0 | 2 | 0 | 8 | 8 | 0.56 | Y |
| 0 | 1 | 0 | 1 | 1 | 4 | 5 | 9 | 0.80 | N |

| SV-Humerus F | SV-Humerus D | SV-Forearm (Ulna & | SV-Forearm (Ulna & | SV-Olecranon F | SV-Olecranon D | SV-Ischium | SV-Femur F. | SV-Femur D. | SV-Tibia F. |
|---|---|---|---|---|---|---|---|---|---|
| 0 | 0 | 0 | 0 | 0 | 0 | 0 | 0 | 0 | 0 |
| 0 | 0 | 0 | 0 | 0 | 0 | 0 | 0 | 0 | 0 |
| 0 | 0 | 0 | 0 | 0 | 0 | 0 | 0 | 0 | 0 |
| 0 | 0 | 0 | 0 | 0 | 0 | 0 | 0 | 0 | 0 |
| 0 | 0 | 0 | 0 | 0 | 0 | 0 | 0 | 0 | 0 |
| 0 | 0 | 0 | 0 | 0 | 0 | 0 | 0 | 0 | 0 |
| 0 | 0 | 0 | 0 | 0 | 0 | 0 | 0 | 0 | 0 |
| 0 | 0 | 0 | 0 | 0 | 0 | 0 | 0 | 0 | 0 |
| 0 | 0 | 0 | 0 | 0 | 0 | 0 | 0 | 0 | 0 |
| 0 | 0 | 0 | 0 | 0 | 0 | 0 | 0 | 0 | 0 |
| 0 | 0 | 0 | 0 | 0 | 0 | 0 | 0 | 0 | 0 |
| 0 | 0 | 0 | 0 | 0 | 0 | 0 | 0 | 0 | 0 |
| 0 | 0 | 0 | 0 | 0 | 0 | 0 | 0 | 0 | 0 |
| 0 | 0 | 0 | 0 | 0 | 0 | 0 | 0 | 0 | 0 |
| 0 | 0 | 0 | 0 | 0 | 0 | 0 | 0 | 0 | 0 |
| 0 | 0 | 0 | 0 | 0 | 0 | 0 | 0 | 0 | 0 |
| 0 | 0 | 0 | 0 | 0 | 0 | 0 | 0 | 0 | 0 |
| 0 | 0 | 0 | 0 | 0 | 0 | 0 | 0 | 0 | 0 |
| 0 | 0 | 0 | 0 | 0 | 0 | 0 | 0 | 0 | 0 |
| 0 | 0 | 0 | 0 | 0 | 0 | 0 | 0 | 0 | 0 |
| 0 | 0 | 0 | 0 | 0 | 0 | 0 | 0 | 0 | 0 |
| 0 | 0 | 0 | 0 | 0 | 0 | 0 | 0 | 0 | 0 |
| 0 | 0 | 0 | 0 | 0 | 0 | 0 | 0 | 0 | 0 |
| 0 | 0 | 0 | 0 | 0 | 0 | 0 | 0 | 0 | 0 |
| 0 | 0 | 0 | 0 | 0 | 0 | 0 | 0 | 0 | 0 |
| 0 | 0 | 0 | 0 | 0 | 0 | 0 | 0 | 0 | 0 |
| 0 | 0 | 0 | 0 | 0 | 0 | 0 | 0 | 0 | 0 |
| 0 | 0 | 0 | 0 | 0 | 0 | 0 | 0 | 0 | 0 |
| 0 | 0 | 0 | 0 | 0 | 0 | 0 | 0 | 0 | 0 |

| | | | | | | | | | |
|---|---|---|---|---|---|---|---|---|---|
| 0 | 0 | 0 | 0 | 0 | 0 | 0 | 0 | 0 | 0 |
| 0 | 0 | 0 | 0 | 0 | 0 | 0 | 0 | 0 | 0 |
| 0 | 0 | 0 | 0 | 0 | 0 | 0 | 0 | 0 | 0 |
| 0 | 0 | 0 | 0 | 0 | 0 | 0 | 0 | 0 | 0 |
| 0 | 0 | 0 | 0 | 0 | 0 | 0 | 0 | 0 | 0 |
| 0 | 0 | 0 | 0 | 0 | 0 | 0 | 0 | 0 | 0 |
| 0 | 0 | 0 | 0 | 0 | 0 | 0 | 0 | 0 | 0 |
| 0 | 0 | 0 | 0 | 0 | 0 | 0 | 0 | 0 | 0 |
| 0 | 0 | 0 | 0 | 0 | 0 | 0 | 0 | 0 | 0 |
| 0 | 0 | 0 | 0 | 0 | 0 | 0 | 0 | 0 | 0 |
| 0 | 0 | 0 | 0 | 0 | 0 | 0 | 0 | 0 | 0 |
| 0 | 0 | 0 | 0 | 0 | 0 | 0 | 0 | 0 | 0 |
| 0 | 0 | 0 | 0 | 0 | 0 | 0 | 0 | 0 | 0 |
| 0 | 0 | 0 | 0 | 0 | 0 | 0 | 0 | 0 | 0 |
| 0 | 0 | 0 | 0 | 0 | 0 | 0 | 0 | 0 | 0 |
| 0 | 0 | 0 | 0 | 0 | 0 | 0 | 0 | 0 | 0 |
| 0 | 0 | 0 | 0 | 0 | 0 | 0 | 0 | 0 | 0 |
| 0 | 0 | 0 | 0 | 0 | 0 | 0 | 0 | 0 | 0 |
| 0 | 0 | 0 | 0 | 0 | 0 | 0 | 0 | 0 | 0 |
| 0 | 0 | 0 | 0 | 0 | 0 | 0 | 0 | 0 | 0 |
| 0 | 0 | 0 | 0 | 0 | 0 | 0 | 0 | 0 | 0 |
| 0 | 0 | 0 | 0 | 0 | 0 | 0 | 0 | 0 | 0 |
| 0 | 0 | 0 | 0 | 0 | 0 | 0 | 0 | 0 | 0 |
| 0 | 0 | 0 | 0 | 0 | 0 | 0 | 0 | 0 | 0 |
| 0 | 0 | 0 | 0 | 0 | 0 | 0 | 0 | 0 | 0 |
| 0 | 0 | 0 | 0 | 0 | 0 | 0 | 0 | 0 | 0 |
| 0 | 0 | 0 | 0 | 0 | 0 | 0 | 0 | 0 | 0 |
| 0 | 0 | 0 | 0 | 0 | 0 | 0 | 0 | 0 | 0 |
| 0 | 0 | 0 | 0 | 0 | 0 | 0 | 0 | 0 | 0 |
| 0 | 0 | 0 | 0 | 0 | 0 | 0 | 0 | 0 | 0 |

| | | | | | | | | | |
|---|---|---|---|---|---|---|---|---|---|
| 0 | 0 | 0 | 0 | 0 | 0 | 0 | 0 | 0 | 0 |
| 0 | 0 | 0 | 0 | 0 | 0 | 0 | 0 | 0 | 0 |
| 0 | 0 | 0 | 0 | 0 | 0 | 0 | 0 | 0 | 0 |
| 0 | 0 | 0 | 0 | 0 | 0 | 0 | 0 | 0 | 0 |
| 0 | 0 | 0 | 0 | 0 | 0 | 0 | 0 | 0 | 0 |
| 0 | 0 | 0 | 0 | 0 | 0 | 0 | 0 | 0 | 0 |
| 0 | 0 | 0 | 0 | 0 | 0 | 0 | 0 | 0 | 0 |
| 0 | 0 | 0 | 0 | 0 | 0 | 0 | 0 | 0 | 0 |
| 0 | 0 | 0 | 0 | 0 | 0 | 0 | 0 | 0 | 0 |
| 0 | 0 | 0 | 0 | 0 | 0 | 0 | 0 | 0 | 0 |
| 0 | 0 | 0 | 0 | 0 | 0 | 2 | 0 | 0 | 1 |
| 1 | 0 | 0 | 1 | 0 | 1 | 2 | 0 | 0 | 0 |
| 0 | 0 | 0 | 2 | 0 | 2 | 2 | 0 | 0 | 0 |
| 0 | 1 | 0 | 1 | 0 | 0 | 2 | 0 | 0 | 0 |
| 0 | 0 | 0 | 0 | 0 | 0 | 2 | 0 | 0 | 0 |
| 0 | 0 | 0 | 2 | 0 | 0 | 2 | 0 | 0 | 0 |
| 0 | 0 | 0 | 2 | 0 | 0 | 2 | 1 | 0 | 0 |
| 0 | 0 | 0 | 2 | 0 | 1 | 2 | 0 | 0 | 0 |
| 0 | 1 | 0 | 2 | 0 | 1 | 2 | 0 | 0 | 0 |
| 1 | 0 | 0 | 2 | 0 | 2 | 2 | 0 | 0 | 0 |
| 0 | 1 | 0 | 2 | 0 | 0 | 2 | 0 | 0 | 0 |
| 1 | 1 | 0 | 2 | 0 | 1 | 2 | 0 | 0 | 0 |
| 1 | 1 | 0 | 2 | 0 | 1 | 2 | 0 | 0 | 0 |
| 2 | 0 | 0 | 1 | 0 | 0 | 2 | 0 | 0 | 0 |
| 0 | 0 | 0 | 0 | 0 | 0 | 2 | 0 | 0 | 1 |
| 0 | 0 | 0 | 1 | 0 | 0 | 2 | 0 | 0 | 0 |
| 0 | 1 | 0 | 2 | 0 | 1 | 2 | 0 | 0 | 0 |
| 1 | 1 | 1 | 1 | 0 | 0 | 2 | 0 | 0 | 0 |
| 0 | 0 | 0 | 2 | 0 | 0 | 2 | 0 | 0 | 0 |
| 1 | 0 | 0 | 1 | 0 | 0 | 2 | 0 | 0 | 0 |

| | | | | | | | | | |
|---|---|---|---|---|---|---|---|---|---|
| 0 | 0 | 0 | 2 | 0 | 1 | 2 | 0 | 0 | 0 |
| 1 | 0 | 0 | 2 | 0 | 2 | 2 | 0 | 0 | 0 |
| 1 | 0 | 0 | 2 | 0 | 1 | 2 | 0 | 0 | 0 |
| 0 | 1 | 0 | 1 | 0 | 0 | 2 | 0 | 0 | 0 |
| 0 | 1 | 0 | 1 | 0 | 0 | 2 | 0 | 0 | 0 |
| 1 | 0 | 0 | 2 | 0 | 1 | 2 | 1 | 0 | 0 |
| 0 | 0 | 0 | 2 | 0 | 2 | 2 | 0 | 0 | 1 |
| 0 | 1 | 0 | 2 | 0 | 1 | 2 | 0 | 0 | 0 |
| 0 | 0 | 0 | 2 | 0 | 0 | 2 | 0 | 0 | 0 |
| 0 | 0 | 0 | 2 | 0 | 1 | 2 | 0 | 0 | 0 |
| 1 | 0 | 0 | 1 | 0 | 0 | 2 | 0 | 0 | 2 |
| 0 | 0 | 0 | 2 | 0 | 0 | 2 | 1 | 0 | 0 |
| 0 | 0 | 0 | 2 | 0 | 1 | 2 | 0 | 0 | 0 |
| 1 | 0 | 0 | 2 | 0 | 1 | 2 | 0 | 0 | 0 |
| 0 | 1 | 0 | 2 | 0 | 0 | 2 | 0 | 0 | 0 |
| 0 | 2 | 0 | 2 | 0 | 2 | 2 | 0 | 0 | 0 |
| 1 | 0 | 0 | 2 | 0 | 1 | 2 | 0 | 0 | 1 |
| 1 | 0 | 0 | 2 | 0 | 0 | 2 | 0 | 0 | 0 |
| 0 | 1 | 0 | 2 | 0 | 0 | 2 | 0 | 0 | 0 |
| 2 | 0 | 0 | 2 | 0 | 1 | 2 | 0 | 0 | 1 |
| 1 | 0 | 0 | 2 | 0 | 2 | 2 | 0 | 0 | 0 |
| 1 | 1 | 0 | 2 | 0 | 2 | 2 | 0 | 0 | 0 |
| 0 | 2 | 0 | 2 | 0 | 1 | 2 | 0 | 0 | 1 |
| 0 | 2 | 0 | 2 | 0 | 0 | 2 | 0 | 0 | 1 |
| 2 | 0 | 0 | 1 | 0 | 0 | 2 | 0 | 0 | 0 |
| 0 | 1 | 0 | 2 | 0 | 1 | 2 | 0 | 0 | 0 |
| 1 | 0 | 0 | 2 | 0 | 2 | 2 | 0 | 0 | 0 |
| 1 | 0 | 0 | 2 | 0 | 1 | 2 | 0 | 0 | 0 |
| 0 | 1 | 0 | 2 | 0 | 1 | 2 | 0 | 0 | 1 |
| 0 | 1 | 0 | 2 | 0 | 1 | 2 | 0 | 0 | 2 |

| | | | | | | | | | |
|---|---|---|---|---|---|---|---|---|---|
| 1 | 0 | 0 | 2 | 0 | 1 | 2 | 0 | 0 | 1 |
| 0 | 0 | 0 | 2 | 0 | 2 | 2 | 0 | 0 | 0 |
| 1 | 0 | 0 | 2 | 0 | 2 | 2 | 1 | 0 | 2 |
| 0 | 1 | 0 | 2 | 0 | 2 | 2 | 0 | 0 | 1 |
| 1 | 1 | 0 | 2 | 0 | 2 | 2 | 0 | 1 | 0 |
| 0 | 0 | 0 | 0 | 0 | 0 | 2 | 0 | 0 | 0 |
| 0 | 0 | 0 | 0 | 0 | 0 | 2 | 0 | 0 | 0 |
| 0 | 0 | 0 | 0 | 0 | 0 | 2 | 0 | 0 | 0 |
| 0 | 0 | 0 | 0 | 0 | 0 | 2 | 0 | 0 | 0 |
| 0 | 0 | 0 | 0 | 0 | 0 | 2 | 0 | 0 | 0 |
| 0 | 0 | 0 | 0 | 0 | 0 | 2 | 0 | 0 | 0 |
| 0 | 0 | 0 | 0 | | 0 | 0 | 0 | 0 | 0 |
| 0 | 0 | 0 | 0 | 0 | 0 | 1 | 0 | 0 | 0 |
| 0 | 0 | 0 | 0 | 0 | 0 | 2 | 0 | 0 | 0 |
| 0 | 0 | 0 | 0 | 0 | 0 | 1 | 0 | 0 | 1 |
| 0 | 0 | 0 | 0 | 0 | 0 | 2 | 0 | 1 | 0 |
| 0 | 0 | 0 | 0 | 0 | 0 | 0 | 0 | 0 | 0 |
| 0 | 0 | 0 | 0 | 0 | 0 | 2 | 0 | 0 | 0 |
| 1 | 0 | 0 | 0 | 0 | 0 | 2 | 0 | 0 | 0 |
| 0 | 0 | 0 | 0 | 0 | 0 | 2 | 0 | 0 | 0 |
| 0 | 0 | 0 | 0 | 0 | 0 | 2 | 0 | 0 | 0 |
| 0 | 0 | 0 | 0 | 0 | 0 | 2 | 0 | 0 | 0 |
| 0 | 0 | 0 | 0 | 0 | 0 | 2 | 0 | 0 | 0 |
| 0 | 0 | 0 | 0 | 0 | 0 | 2 | 1 | 0 | 0 |
| 0 | 0 | 0 | 0 | 0 | 0 | 2 | 0 | 0 | 0 |
| 0 | 0 | 0 | 0 | 0 | 0 | 2 | 0 | 0 | 0 |
| 0 | 0 | 0 | 0 | 0 | 0 | 2 | 0 | 0 | 0 |
| 0 | 0 | 0 | 0 | 0 | 0 | 2 | 0 | 0 | 0 |
| 0 | 0 | 0 | 0 | 0 | 0 | 2 | 0 | 0 | 0 |
| 1 | 0 | 0 | 0 | 0 | 0 | 2 | 0 | 0 | 0 |

| | | | | | | | | | |
|---|---|---|---|---|---|---|---|---|---|
| 0 | 0 | 0 | 0 | 0 | 0 | 2 | 0 | 0 | 0 |
| 1 | 0 | 0 | 1 | 0 | 1 | 2 | 0 | 0 | 0 |
| 0 | 0 | 0 | 2 | 0 | 0 | 2 | 0 | 0 | 0 |
| 0 | 0 | 0 | 0 | 0 | 0 | 2 | 0 | 0 | 0 |
| 0 | 0 | 0 | 2 | 0 | 0 | 2 | 0 | 0 | 0 |
| 0 | 0 | 0 | 0 | 0 | 2 | 2 | 0 | 0 | 0 |
| 0 | 0 | 0 | 0 | 0 | 0 | 2 | 0 | 0 | 0 |
| 0 | 0 | 0 | 1 | 0 | 0 | 2 | 0 | 0 | 0 |
| 0 | 0 | 0 | 0 | 0 | 0 | 1 | 0 | 0 | 0 |
| 0 | 0 | 0 | 0 | 0 | 0 | 2 | 0 | 0 | 0 |
| 0 | 0 | 0 | 0 | 0 | 1 | 2 | 0 | 0 | 0 |
| 0 | 0 | 0 | 2 | 0 | 2 | 2 | 0 | 0 | 0 |
| 0 | 0 | 0 | 0 | 0 | 0 | 2 | 0 | 0 | 0 |
| 1 | 0 | 0 | 2 | 0 | 2 | 2 | 0 | 0 | 0 |
| 0 | 0 | 0 | 1 | 0 | 1 | 2 | 0 | 0 | 0 |
| 0 | 0 | 0 | 2 | 0 | 1 | 2 | 0 | 0 | 0 |
| 0 | 0 | 0 | 2 | 0 | 0 | 2 | 0 | 0 | 0 |
| 0 | 0 | 0 | 2 | 0 | 0 | 2 | 0 | 0 | 0 |
| 0 | 0 | 0 | 0 | 0 | 0 | 2 | 0 | 0 | 0 |
| 0 | 0 | 0 | 2 | 0 | 1 | 2 | 0 | 0 | 2 |
| 0 | 0 | 0 | 2 | 0 | 1 | 2 | 0 | 0 | 2 |
| 0 | 0 | 0 | 0 | 0 | 0 | 0 | 0 | 0 | 0 |
| 0 | 0 | 0 | 0 | 0 | 0 | 2 | 0 | 0 | 0 |
| 0 | 0 | 0 | 2 | 0 | 2 | 2 | 0 | 0 | 0 |
| 0 | 0 | 0 | 0 | 0 | 0 | 2 | 0 | 0 | 0 |
| 0 | 0 | 0 | 2 | 0 | 2 | 2 | 0 | 0 | 2 |
| 0 | 0 | 0 | 2 | 0 | 0 | 2 | 0 | 0 | 0 |
| 0 | 0 | 0 | 2 | 0 | 0 | 2 | 0 | 0 | 0 |
| 0 | 0 | 0 | 0 | 0 | 0 | 2 | 0 | 0 | 0 |
| 0 | 0 | 0 | 0 | 0 | 0 | 2 | 0 | 1 | 0 |

| | | | | | | | | | |
|---|---|---|---|---|---|---|---|---|---|
| 0 | 0 | 0 | 0 | 0 | 0 | 2 | 0 | 0 | 0 |
| 1 | 0 | 0 | 0 | 0 | 0 | 2 | 0 | 0 | 0 |
| 0 | 0 | 0 | 0 | 0 | 0 | 2 | 0 | 0 | 0 |
| 1 | 1 | 0 | 1 | 0 | 0 | 2 | 0 | 0 | 0 |
| 1 | 0 | 0 | 2 | 0 | 1 | 2 | 0 | 0 | 0 |
| 0 | 0 | 0 | 2 | 0 | 0 | 2 | 0 | 0 | 0 |
| 0 | 0 | 0 | 2 | 0 | 2 | 2 | 0 | 0 | 0 |
| 0 | 0 | 0 | 2 | 0 | 1 | 2 | 0 | 0 | 2 |
| 1 | 0 | 0 | 2 | 0 | 1 | 1 | 0 | 0 | 0 |
| 0 | 0 | 0 | 2 | 0 | 0 | 2 | 0 | 0 | 0 |
| 1 | 0 | 0 | 1 | 0 | 1 | 2 | 0 | 0 | 0 |
| 0 | 0 | 0 | 2 | 0 | 1 | 2 | 0 | 0 | 2 |
| 0 | 0 | 0 | 1 | 0 | 0 | 2 | 0 | 0 | 0 |
| 0 | 1 | 0 | 2 | 0 | 2 | 2 | 0 | 0 | 0 |
| 0 | 0 | 0 | 2 | 0 | 2 | 2 | 0 | 0 | 0 |
| 0 | 0 | 0 | 2 | 0 | 0 | 2 | 0 | 0 | 0 |
| 0 | 0 | 0 | 2 | 0 | 0 | 2 | 0 | 0 | 0 |
| 0 | 0 | 0 | 2 | 0 | 0 | 2 | 0 | 0 | 0 |
| 0 | 0 | 0 | 2 | 0 | 0 | 2 | 0 | 0 | 0 |
| 1 | 1 | 0 | 2 | 0 | 1 | 1 | 0 | 0 | 0 |
| 1 | 0 | 0 | 2 | 0 | 2 | 2 | 0 | 0 | 0 |
| 0 | 0 | 0 | 2 | 0 | 0 | 2 | 0 | 0 | 0 |
| 1 | 1 | 0 | 2 | 0 | 1 | 1 | 0 | 0 | 1 |

| SV-Tibia D. | SV-Calcaneus F | SV-Calcaneus D | SV-Total Fractures | SV-Total Deformities | SV-Total Score | SV-Pelvis Ratio | SV-Scoliosis | TOTAL FRACTURES |
|---|---|---|---|---|---|---|---|---|
| 0 | 0 | 0 | 0 | 0 | 0 | 1 | N | 0.0 |
| 0 | 0 | 0 | 0 | 0 | 0 | 1 | N | 0.0 |
| 0 | 0 | 0 | 0 | 0 | 0 | 1 | N | 0.0 |
| 0 | 0 | 0 | 0 | 0 | 0 | 1 | N | 0.0 |
| 0 | 0 | 0 | 0 | 0 | 0 | 1 | N | 0.0 |
| 0 | 0 | 0 | 0 | 0 | 0 | 1 | N | 0.0 |
| 0 | 0 | 0 | 0 | 0 | 0 | 1 | N | 0.0 |
| 0 | 0 | 0 | 0 | 0 | 0 | 1 | N | 0.0 |
| 0 | 0 | 0 | 0 | 0 | 0 | 1 | N | 0.0 |
| 0 | 0 | 0 | 0 | 0 | 0 | 1 | N | 0.0 |
| 0 | 0 | 0 | 0 | 0 | 0 | 1 | N | 0.0 |
| 0 | 0 | 0 | 0 | 0 | 0 | 1 | N | 0.0 |
| 0 | 0 | 0 | 0 | 0 | 0 | 1 | N | 0.0 |
| 0 | 0 | 0 | 0 | 0 | 0 | 1 | N | 0.0 |
| 0 | 0 | 0 | 0 | 0 | 0 | 1 | N | 0.0 |
| 0 | 0 | 0 | 0 | 0 | 0 | 1 | N | 0.0 |
| 0 | 0 | 0 | 0 | 0 | 0 | 1 | N | 0.0 |
| 0 | 0 | 0 | 0 | 0 | 0 | 1 | N | 0.0 |
| 0 | 0 | 0 | 0 | 0 | 0 | 1 | N | 0.0 |
| 0 | 0 | 0 | 0 | 0 | 0 | 1 | N | 0.0 |
| 0 | 0 | 0 | 0 | 0 | 0 | 1 | N | 0.0 |
| 0 | 0 | 0 | 0 | 0 | 0 | 1 | N | 0.0 |
| 0 | 0 | 0 | 0 | 0 | 0 | 1 | N | 0.0 |
| 0 | 0 | 0 | 0 | 0 | 0 | 1 | N | 0.0 |
| 0 | 0 | 0 | 0 | 0 | 0 | 1 | N | 0.0 |
| 0 | 0 | 0 | 0 | 0 | 0 | 1 | N | 0.0 |
| 0 | 0 | 0 | 0 | 0 | 0 | 1 | N | 0.0 |
| 0 | 0 | 0 | 0 | 0 | 0 | 1 | N | 0.0 |
| 0 | 0 | 0 | 0 | 0 | 0 | 1 | N | 0.0 |

| | | | | | | | | |
|---|---|---|---|---|---|---|---|---|
| 0 | 0 | 0 | 0 | 0 | 0 | 1 | N | 0.0 |
| 0 | 0 | 0 | 0 | 0 | 0 | 1 | N | 0.0 |
| 0 | 0 | 0 | 0 | 0 | 0 | 1 | N | 0.0 |
| 0 | 0 | 0 | 0 | 0 | 0 | 1 | N | 0.0 |
| 0 | 0 | 0 | 0 | 0 | 0 | 1 | N | 0.0 |
| 0 | 0 | 0 | 0 | 0 | 0 | 1 | N | 0.0 |
| 0 | 0 | 0 | 0 | 0 | 0 | 1 | N | 0.0 |
| 0 | 0 | 0 | 0 | 0 | 0 | 1 | N | 0.0 |
| 0 | 0 | 0 | 0 | 0 | 0 | 1 | N | 0.0 |
| 0 | 0 | 0 | 0 | 0 | 0 | 1 | N | 0.0 |
| 0 | 0 | 0 | 0 | 0 | 0 | 1 | N | 0.0 |
| 0 | 0 | 0 | 0 | 0 | 0 | 1 | N | 0.0 |
| 0 | 0 | 0 | 0 | 0 | 0 | 1 | N | 0.0 |
| 0 | 0 | 0 | 0 | 0 | 0 | 1 | N | 0.0 |
| 0 | 0 | 0 | 0 | 0 | 0 | 1 | N | 0.0 |
| 0 | 0 | 0 | 0 | 0 | 0 | 1 | N | 0.0 |
| 0 | 0 | 0 | 0 | 0 | 0 | 1 | N | 0.0 |
| 0 | 0 | 0 | 0 | 0 | 0 | 1 | N | 0.0 |
| 0 | 0 | 0 | 0 | 0 | 0 | 1 | N | 0.0 |
| 0 | 0 | 0 | 0 | 0 | 0 | 1 | N | 0.0 |
| 0 | 0 | 0 | 0 | 0 | 0 | 1 | N | 0.0 |
| 0 | 0 | 0 | 0 | 0 | 0 | 1 | N | 0.0 |
| 0 | 0 | 0 | 0 | 0 | 0 | 1 | N | 0.0 |
| 0 | 0 | 0 | 0 | 0 | 0 | 1 | N | 0.0 |
| 0 | 0 | 0 | 0 | 0 | 0 | 1 | N | 0.0 |
| 0 | 0 | 0 | 0 | 0 | 0 | 1 | N | 0.0 |
| 0 | 0 | 0 | 0 | 0 | 0 | 1 | N | 0.0 |
| 0 | 0 | 0 | 0 | 0 | 0 | 1 | N | 0.0 |
| 0 | 0 | 0 | 0 | 0 | 0 | 1 | N | 0.0 |
| 0 | 0 | 0 | 0 | 0 | 0 | 1 | N | 0.0 |

| | | | | | | | | |
|---|---|---|---|---|---|---|---|---|
| 0 | 0 | 0 | 0 | 0 | 0 | 1 | N | 0.0 |
| 0 | 0 | 0 | 0 | 0 | 0 | 1 | N | 0.0 |
| 0 | 0 | 0 | 0 | 0 | 0 | 1 | N | 0.0 |
| 0 | 0 | 0 | 0 | 0 | 0 | 1 | N | 0.0 |
| 0 | 0 | 0 | 0 | 0 | 0 | 1 | N | 0.0 |
| 0 | 0 | 0 | 0 | 0 | 0 | 1 | N | 0.0 |
| 0 | 0 | 0 | 0 | 0 | 0 | 1 | N | 0.0 |
| 0 | 0 | 0 | 0 | 0 | 0 | 1 | N | 0.0 |
| 0 | 0 | 0 | 0 | 0 | 0 | 1 | N | 0.0 |
| 0 | 0 | 0 | 0 | 0 | 0 | 1 | N | 0.0 |
| 0 | 0 | 2 | 1 | 4 | 5 | 0.85 | N | 0.2 |
| 0 | 0 | 0 | 1 | 4 | 5 | 0.56 | N | 0.9 |
| 0 | 0 | 1 | 0 | 7 | 7 | 0.4 | Y | 0.0 |
| 1 | 0 | 2 | 0 | 7 | 7 | 0.69 | N | 0.0 |
| 1 | 0 | 2 | 0 | 5 | 5 | 0.93 | N | 0.6 |
| 0 | 0 | 2 | 0 | 6 | 6 | 0.61 | Y | 0.0 |
| 0 | 0 | 2 | 1 | 6 | 7 | 0.58 | N | 0.8 |
| 1 | 0 | 1 | 0 | 7 | 7 | 0.45 | Y | 1.0 |
| 0 | 0 | 2 | 0 | 8 | 8 | 0.55 | N | 0.3 |
| 0 | 0 | 2 | 1 | 8 | 9 | 0.45 | Y | 0.6 |
| 1 | 0 | 2 | 0 | 8 | 8 | 0.49 | Y | 0.4 |
| 1 | 0 | 2 | 1 | 9 | 10 | 0.51 | Y | 1.4 |
| 1 | 0 | 2 | 1 | 9 | 10 | 0.49 | Y | 1.2 |
| 2 | 0 | 1 | 2 | 6 | 8 | 0.59 | Y | 2.1 |
| 0 | 0 | 1 | 1 | 3 | 4 | 0.84 | N | 0.2 |
| 0 | 0 | 2 | 0 | 5 | 5 | 0.55 | N | 0.2 |
| 1 | 0 | 2 | 0 | 9 | 9 | 0.5 | N | 0.4 |
| 0 | 0 | 2 | 2 | 6 | 8 | 0.49 | N | 1.2 |
| 0 | 0 | 1 | 0 | 5 | 5 | 0.4 | Y | 0.0 |
| 0 | 0 | 1 | 1 | 4 | 5 | 0.58 | N | 0.9 |

| | | | | | | | | |
|---|---|---|---|---|---|---|---|---|
| 2 | 0 | 2 | 0 | 9 | 9 | 0.63 | Y | 0.9 |
| 2 | 0 | 2 | 1 | 10 | 11 | 0.57 | Y | 1.9 |
| 2 | 0 | 2 | 1 | 9 | 10 | 0.61 | N | 1.3 |
| 1 | 0 | 2 | 0 | 7 | 7 | 0.65 | N | 0.0 |
| 1 | 0 | 2 | 0 | 7 | 7 | 0.66 | N | 0.4 |
| 0 | 0 | 1 | 2 | 6 | 8 | 0.52 | Y | 0.9 |
| 0 | 0 | 1 | 1 | 7 | 8 | 0.57 | Y | 0.4 |
| 0 | 0 | 2 | 0 | 8 | 8 | 0.58 | Y | 0.4 |
| 0 | 0 | 2 | 0 | 6 | 6 | 0.61 | N | 0.8 |
| 0 | 0 | 2 | 0 | 7 | 7 | 0.56 | Y | 0.2 |
| 0 | 0 | 1 | 3 | 4 | 7 | 0.61 | Y | 1.9 |
| 0 | 0 | 2 | 1 | 6 | 7 | 0.52 | N | 0.4 |
| 0 | 0 | 2 | 0 | 7 | 7 | 0.53 | N | 0.1 |
| 0 | 0 | 2 | 1 | 7 | 8 | 0.54 | N | 1.1 |
| 1 | 0 | 2 | 0 | 8 | 8 | 0.49 | Y | 1.2 |
| 0 | 0 | 2 | 0 | 10 | 10 | 0.57 | Y | 0.8 |
| 0 | 0 | 0 | 2 | 5 | 7 | 0.43 | Y | 0.9 |
| 1 | 0 | 2 | 1 | 7 | 8 | 0.56 | Y | 1.0 |
| 1 | 0 | 2 | 0 | 8 | 8 | 0.51 | Y | 0.6 |
| 0 | 0 | 2 | 3 | 7 | 10 | 0.51 | Y | 1.6 |
| 0 | 0 | 2 | 1 | 8 | 9 | 0.51 | N | 1.4 |
| 0 | 0 | 2 | 1 | 9 | 10 | 0.46 | Y | 0.7 |
| 0 | 0 | 2 | 1 | 9 | 10 | 0.49 | Y | 0.6 |
| 1 | 0 | 1 | 1 | 8 | 9 | 0.58 | Y | 1.8 |
| 2 | 0 | 1 | 2 | 6 | 8 | 0.71 | Y | 1.6 |
| 0 | 0 | 2 | 0 | 8 | 8 | 0.45 | N | 0.6 |
| 2 | 0 | 2 | 1 | 10 | 11 | 0.5 | N | 1.0 |
| 2 | 0 | 2 | 1 | 9 | 10 | 0.63 | Y | 1.9 |
| 0 | 0 | 2 | 1 | 8 | 9 | 0.62 | Y | 1.7 |
| 0 | 0 | 2 | 2 | 8 | 10 | 0.48 | N | 0.9 |

| | | | | | | | | |
|---|---|---|---|---|---|---|---|---|
| 1 | 0 | 2 | 2 | 8 | 10 | 0.56 | Y | 1.9 |
| 2 | 1 | 1 | 1 | 9 | 9 | 0.43 | Y | 0.4 |
| 0 | 0 | 2 | 4 | 8 | 12 | 0.53 | Y | 1.9 |
| 1 | 0 | 2 | 1 | 10 | 11 | 0.63 | N | 1.4 |
| 1 | 0 | 2 | 2 | 11 | 13 | 0.73 | N | 1.9 |
| 0 | 0 | 1 | 0 | 3 | 3 | 0.78 | N | 0.0 |
| 0 | 0 | 2 | 0 | 4 | 4 | 0.8 | N | 0.1 |
| 0 | 0 | 1 | 0 | 3 | 3 | 0.7 | N | 0.0 |
| 0 | 0 | 1 | 0 | 3 | 3 | 0.82 | N | 0.0 |
| 0 | 0 | 1 | 0 | 3 | 3 | 0.7 | N | 0.0 |
| 0 | 0 | 2 | 0 | 4 | 4 | 0.88 | N | 0.3 |
| 0 | 0 | 2 | 0 | 2 | 2 | 1 | N | 0.0 |
| 0 | 0 | 2 | 0 | 3 | 3 | 0.77 | N | 0.0 |
| 0 | 0 | 2 | 0 | 4 | 4 | 0.77 | N | 0.0 |
| 0 | 0 | 2 | 1 | 3 | 4 | 0.88 | N | 0.6 |
| 0 | 0 | 1 | 0 | 4 | 4 | 0.54 | N | 0.0 |
| 0 | 0 | 2 | 0 | 2 | 2 | 1 | N | 0.0 |
| 0 | 0 | 2 | 0 | 4 | 4 | 0.67 | N | 0.1 |
| 0 | 0 | 1 | 1 | 3 | 4 | 0.55 | N | 0.7 |
| 0 | 0 | 2 | 0 | 4 | 4 | 0.88 | N | 0.0 |
| 0 | 0 | 1 | 0 | 3 | 3 | 0.62 | N | 0.0 |
| 0 | 0 | 2 | 0 | 4 | 4 | 0.66 | N | 0.1 |
| 0 | 0 | 1 | 0 | 3 | 3 | 0.59 | N | 0.0 |
| 0 | 0 | 1 | 1 | 3 | 4 | 0.65 | N | 1.0 |
| 0 | 0 | 2 | 0 | 4 | 4 | 0.45 | N | 0.1 |
| 0 | 0 | 2 | 0 | 4 | 4 | 0.52 | N | 0.0 |
| 0 | 0 | 2 | 0 | 4 | 4 | 0.53 | N | 0.3 |
| 0 | 0 | 2 | 0 | 4 | 4 | 0.54 | N | 0.1 |
| 0 | 0 | 2 | 0 | 4 | 4 | 0.67 | N | 0.4 |
| 0 | 0 | 2 | 1 | 4 | 5 | 0.59 | N | 0.7 |

| | | | | | | | | |
|---|---|---|---|---|---|---|---|---|
| 0 | 0 | 2 | 0 | 4 | 4 | 0.62 | N | 0.3 |
| 0 | 0 | 1 | 1 | 5 | 6 | 0.6 | N | 1.0 |
| 0 | 0 | 2 | 0 | 6 | 6 | 0.76 | N | 0.8 |
| 2 | 0 | 2 | 0 | 6 | 6 | 0.74 | Y | 0.7 |
| 0 | 0 | 2 | 0 | 6 | 6 | 0.58 | Y | 0.0 |
| 1 | 0 | 2 | 0 | 7 | 7 | 0.51 | Y | 0.2 |
| 0 | 0 | 1 | 0 | 3 | 3 | 0.84 | N | 0.0 |
| 0 | 0 | 2 | 0 | 5 | 5 | 0.55 | N | 0.0 |
| 0 | 0 | 2 | 0 | 3 | 3 | 0.9 | N | 0.0 |
| 0 | 0 | 1 | 0 | 3 | 3 | 0.71 | N | 0.0 |
| 1 | 0 | 1 | 0 | 5 | 5 | 0.59 | Y | 0.3 |
| 0 | 0 | 2 | 0 | 8 | 8 | 0.66 | Y | 0.4 |
| 0 | 0 | 2 | 0 | 4 | 4 | 0.85 | N | 0.0 |
| 0 | 0 | 1 | 1 | 7 | 8 | 0.53 | Y | 0.9 |
| 0 | 0 | 1 | 0 | 5 | 5 | 0.55 | N | 0.1 |
| 0 | 0 | 2 | 0 | 7 | 7 | 0.6 | Y | 0.0 |
| 1 | 0 | 1 | 0 | 6 | 6 | 0.59 | N | 0.1 |
| 2 | 0 | 1 | 0 | 7 | 7 | 0.61 | N | 1.2 |
| 0 | 0 | 1 | 0 | 3 | 3 | 0.58 | N | 0.0 |
| 0 | 0 | 0 | 2 | 5 | 7 | 0.5 | N | 0.7 |
| 0 | 0 | 0 | 2 | 5 | 7 | 0.55 | N | 0.9 |
| 0 | 0 | 2 | 0 | 2 | 2 | 0.9 | N | 0.1 |
| 0 | 0 | 2 | 0 | 4 | 4 | 0.69 | N | 0.0 |
| 2 | 0 | 2 | 0 | 8 | 8 | 0.45 | N | 0.0 |
| 0 | 0 | 2 | 0 | 4 | 4 | 0.81 | N | 0.2 |
| 0 | 0 | 1 | 2 | 7 | 9 | 0.5 | N | 0.7 |
| 2 | 0 | 2 | 0 | 8 | 8 | 0.57 | N | 0.0 |
| 2 | 0 | 2 | 0 | 8 | 8 | 0.53 | Y | 0.3 |
| 0 | 0 | 1 | 0 | 3 | 3 | 0.59 | N | 0.1 |
| 0 | 0 | 1 | 0 | 4 | 4 | 0.53 | N | 0.1 |

| | | | | | | | | |
|---|---|---|---|---|---|---|---|---|
| 0 | 0 | 2 | 0 | 4 | 4 | 0.5 | N | 0.0 |
| 0 | 0 | 2 | 1 | 4 | 5 | 0.55 | N | 0.7 |
| 2 | 0 | 2 | 0 | 6 | 6 | 0.59 | Y | 0.7 |
| 0 | 0 | 2 | 1 | 6 | 7 | 0.57 | Y | 1.0 |
| 0 | 0 | 2 | 1 | 7 | 8 | 0.64 | N | 0.7 |
| 2 | 0 | 2 | 0 | 8 | 8 | 0.54 | Y | 0.0 |
| 0 | 0 | 2 | 0 | 8 | 8 | 0.53 | Y | 0.0 |
| 0 | 0 | 2 | 2 | 7 | 9 | 0.67 | Y | 1.2 |
| 0 | 0 | 2 | 1 | 6 | 7 | 0.92 | N | 1.9 |
| 0 | 0 | 2 | 0 | 6 | 6 | 0.62 | Y | 0.0 |
| 0 | 0 | 2 | 1 | 6 | 7 | 0.55 | N | 0.1 |
| 0 | 0 | 1 | 2 | 6 | 8 | 0.6 | N | 0.4 |
| 0 | 0 | 2 | 0 | 5 | 5 | 0.52 | Y | 0.6 |
| 0 | 0 | 2 | 0 | 9 | 9 | 0.6 | Y | 0.1 |
| 0 | 0 | 0 | 0 | 6 | 6 | 0.53 | N | 0.6 |
| 1 | 0 | 1 | 0 | 6 | 6 | 0.64 | N | 0.3 |
| 2 | 0 | 1 | 0 | 7 | 7 | 0.57 | N | 0.0 |
| 0 | 0 | 2 | 0 | 6 | 6 | 0.7 | N | 1.1 |
| 0 | 0 | 2 | 0 | 6 | 6 | 0.52 | Y | 0.1 |
| 1 | 0 | 2 | 1 | 8 | 9 | 0.84 | N | 1.1 |
| 0 | 0 | 0 | 1 | 6 | 7 | 0.58 | N | 0.4 |
| 2 | 0 | 2 | 0 | 8 | 8 | 0.56 | Y | 0.2 |
| 0 | 0 | 2 | 2 | 7 | 9 | 0.86 | N | 1.6 |

| TOTAL DEFORMITIES | TOTAL SCORE | Scoliosis Y | Scoliosis N |
|---|---|---|---|
| 0.0 | 0.0 | 0 | 8 |
| 0.0 | 0.0 | 0 | 8 |
| 0.0 | 0.0 | 0 | 8 |
| 0.0 | 0.0 | 0 | 8 |
| 0.0 | 0.0 | 0 | 8 |
| 0.0 | 0.0 | 0 | 8 |
| 0.0 | 0.0 | 0 | 8 |
| 0.0 | 0.0 | 0 | 8 |
| 0.0 | 0.0 | 0 | 8 |
| 0.0 | 0.0 | 0 | 8 |
| 0.0 | 0.0 | 0 | 8 |
| 0.0 | 0.0 | 0 | 8 |
| 0.0 | 0.0 | 0 | 8 |
| 0.0 | 0.0 | 0 | 8 |
| 0.0 | 0.0 | 0 | 8 |
| 0.0 | 0.0 | 0 | 8 |
| 0.0 | 0.0 | 0 | 8 |
| 0.0 | 0.0 | 0 | 8 |
| 0.0 | 0.0 | 0 | 8 |
| 0.0 | 0.0 | 0 | 8 |
| 0.0 | 0.0 | 0 | 8 |
| 0.0 | 0.0 | 0 | 8 |
| 0.0 | 0.0 | 0 | 8 |
| 0.0 | 0.0 | 0 | 8 |
| 0.0 | 0.0 | 0 | 8 |
| 0.0 | 0.0 | 0 | 8 |
| 0.0 | 0.0 | 0 | 8 |
| 0.0 | 0.0 | 0 | 8 |
| 0.0 | 0.0 | 0 | 8 |

| | | | |
|---|---|---|---|
| 0.0 | 0.0 | 0 | 8 |
| 0.0 | 0.0 | 0 | 8 |
| 0.0 | 0.0 | 0 | 8 |
| 0.0 | 0.0 | 0 | 8 |
| 0.0 | 0.0 | 0 | 8 |
| 0.0 | 0.0 | 0 | 8 |
| 0.0 | 0.0 | 0 | 8 |
| 0.0 | 0.0 | 0 | 8 |
| 0.0 | 0.0 | 0 | 8 |
| 0.0 | 0.0 | 0 | 8 |
| 0.0 | 0.0 | 0 | 8 |
| 0.0 | 0.0 | 0 | 8 |
| 0.0 | 0.0 | 0 | 8 |
| 0.0 | 0.0 | 0 | 8 |
| 0.0 | 0.0 | 0 | 8 |
| 0.0 | 0.0 | 0 | 8 |
| 0.0 | 0.0 | 0 | 8 |
| 0.0 | 0.0 | 0 | 8 |
| 0.0 | 0.0 | 0 | 8 |
| 0.0 | 0.0 | 0 | 8 |
| 0.0 | 0.0 | 0 | 8 |
| 0.0 | 0.0 | 0 | 8 |
| 0.0 | 0.0 | 0 | 8 |
| 0.0 | 0.0 | 0 | 8 |
| 0.0 | 0.0 | 0 | 8 |
| 0.0 | 0.0 | 0 | 8 |
| 0.0 | 0.0 | 0 | 8 |
| 0.0 | 0.0 | 0 | 8 |
| 0.0 | 0.0 | 0 | 8 |
| 0.0 | 0.0 | 0 | 8 |

| | | | |
|---|---|---|---|
| 0.0 | 0.0 | 0 | 8 |
| 0.0 | 0.0 | 0 | 8 |
| 0.0 | 0.0 | 0 | 8 |
| 0.0 | 0.0 | 0 | 8 |
| 0.0 | 0.0 | 0 | 8 |
| 0.0 | 0.0 | 0 | 8 |
| 0.0 | 0.0 | 0 | 8 |
| 0.0 | 0.0 | 0 | 8 |
| 0.0 | 0.0 | 0 | 8 |
| 0.0 | 0.0 | 0 | 8 |
| 3.7 | 3.9 | 1 | 7 |
| 4.1 | 5.0 | 0 | 8 |
| 5.2 | 5.2 | 5 | 3 |
| 5.6 | 5.6 | 0 | 8 |
| 2.9 | 3.4 | 1 | 7 |
| 5.6 | 5.6 | 8 | 0 |
| 5.6 | 6.3 | 2 | 6 |
| 5.9 | 6.9 | 6 | 2 |
| 7.0 | 7.3 | 3 | 5 |
| 6.7 | 7.2 | 7 | 1 |
| 6.9 | 7.3 | 6 | 2 |
| 6.3 | 7.8 | 7 | 1 |
| 6.7 | 7.9 | 6 | 2 |
| 5.9 | 8.0 | 7 | 1 |
| 3.7 | 3.9 | 1 | 7 |
| 4.6 | 4.8 | 0 | 8 |
| 7.7 | 8.1 | 2 | 6 |
| 6.8 | 8.0 | 5 | 3 |
| 4.9 | 4.9 | 4 | 4 |
| 4.2 | 5.1 | 0 | 8 |

| | | | |
|---|---|---|---|
| 7.8 | 8.7 | 8 | 0 |
| 7.2 | 9.1 | 8 | 0 |
| 8.2 | 9.6 | 0 | 8 |
| 5.4 | 5.4 | 2 | 6 |
| 5.0 | 5.4 | 0 | 8 |
| 4.9 | 5.8 | 5 | 3 |
| 5.3 | 5.8 | 6 | 2 |
| 5.6 | 6.0 | 8 | 0 |
| 5.1 | 5.9 | 2 | 6 |
| 5.8 | 6.0 | 8 | 0 |
| 4.8 | 6.7 | 6 | 2 |
| 6.1 | 6.6 | 2 | 6 |
| 6.6 | 6.7 | 3 | 5 |
| 5.7 | 6.8 | 4 | 4 |
| 5.8 | 7.0 | 5 | 3 |
| 6.6 | 7.3 | 7 | 1 |
| 6.4 | 7.3 | 6 | 2 |
| 6.6 | 7.6 | 5 | 3 |
| 6.9 | 7.4 | 6 | 2 |
| 5.9 | 7.4 | 8 | 0 |
| 6.0 | 7.4 | 4 | 4 |
| 7.0 | 7.7 | 7 | 1 |
| 7.3 | 7.9 | 7 | 1 |
| 6.2 | 8.0 | 5 | 3 |
| 6.4 | 8.0 | 7 | 1 |
| 7.6 | 8.1 | 5 | 3 |
| 7.3 | 8.3 | 3 | 5 |
| 6.7 | 8.6 | 8 | 0 |
| 6.8 | 8.4 | 7 | 1 |
| 7.7 | 8.6 | 3 | 5 |

| | | | |
|---|---|---|---|
| 6.4 | 8.3 | 5 | 3 |
| 8.4 | 8.8 | 8 | 0 |
| 6.9 | 8.8 | 7 | 1 |
| 7.9 | 9.3 | 0 | 8 |
| 7.9 | 9.8 | 0 | 8 |
| 2.3 | 2.3 | 1 | 7 |
| 2.9 | 3.0 | 0 | 8 |
| 2.8 | 2.8 | 0 | 8 |
| 2.8 | 2.8 | 2 | 6 |
| 3.0 | 3.0 | 0 | 8 |
| 2.8 | 3.1 | 0 | 8 |
| 3.0 | 3.0 | 0 | 8 |
| 3.1 | 3.1 | 1 | 7 |
| 3.1 | 3.1 | 0 | 8 |
| 2.6 | 3.1 | 0 | 8 |
| 3.0 | 3.0 | 3 | 5 |
| 3.1 | 3.1 | 0 | 8 |
| 3.3 | 3.4 | 0 | 8 |
| 2.8 | 3.4 | 0 | 8 |
| 3.4 | 3.4 | 1 | 7 |
| 3.6 | 3.6 | 0 | 8 |
| 3.7 | 3.8 | 0 | 8 |
| 3.8 | 3.8 | 2 | 6 |
| 2.6 | 3.6 | 1 | 7 |
| 3.8 | 3.9 | 0 | 8 |
| 3.9 | 3.9 | 1 | 7 |
| 3.6 | 3.9 | 1 | 7 |
| 3.9 | 4.0 | 3 | 5 |
| 3.8 | 4.2 | 0 | 8 |
| 3.7 | 4.3 | 3 | 5 |

| | | | |
|---|---|---|---|
| 4.8 | 5.1 | 3 | 5 |
| 4.2 | 5.2 | 0 | 8 |
| 4.6 | 5.3 | 0 | 8 |
| 4.7 | 5.3 | 6 | 2 |
| 5.3 | 5.3 | 5 | 3 |
| 5.1 | 5.3 | 8 | 0 |
| 2.3 | 2.3 | 1 | 7 |
| 5.4 | 5.4 | 0 | 8 |
| 2.9 | 2.9 | 0 | 8 |
| 3.0 | 3.0 | 1 | 7 |
| 5.2 | 5.6 | 6 | 2 |
| 5.3 | 5.8 | 6 | 2 |
| 3.1 | 3.1 | 0 | 8 |
| 5.1 | 6.0 | 6 | 2 |
| 3.2 | 3.3 | 1 | 7 |
| 6.3 | 6.3 | 6 | 2 |
| 6.1 | 6.2 | 0 | 8 |
| 5.1 | 6.3 | 0 | 8 |
| 3.6 | 3.6 | 0 | 8 |
| 5.9 | 6.3 | 1 | 7 |
| 5.6 | 6.4 | 0 | 8 |
| 3.1 | 3.2 | 0 | 8 |
| 3.6 | 3.6 | 1 | 7 |
| 6.6 | 6.6 | 0 | 8 |
| 3.4 | 3.7 | 0 | 8 |
| 6.0 | 6.7 | 0 | 8 |
| 6.8 | 6.8 | 3 | 5 |
| 6.6 | 6.9 | 6 | 2 |
| 3.7 | 3.8 | 0 | 8 |
| 3.4 | 3.6 | 2 | 6 |

| | | | |
|---|---|---|---|
| 3.9 | 3.9 | 1 | 7 |
| 4.3 | 5.0 | 3 | 5 |
| 4.6 | 5.2 | 6 | 2 |
| 6.1 | 7.1 | 8 | 0 |
| 6.6 | 7.2 | 0 | 8 |
| 7.0 | 7.0 | 7 | 1 |
| 7.0 | 7.0 | 7 | 1 |
| 5.9 | 7.1 | 7 | 1 |
| 5.2 | 7.1 | 0 | 8 |
| 5.4 | 5.4 | 6 | 2 |
| 5.3 | 5.4 | 0 | 8 |
| 6.6 | 7.0 | 0 | 8 |
| 5.3 | 5.9 | 8 | 0 |
| 6.3 | 6.4 | 6 | 2 |
| 6.0 | 6.6 | 0 | 8 |
| 6.2 | 6.6 | 0 | 8 |
| 6.7 | 6.7 | 3 | 5 |
| 5.8 | 6.9 | 0 | 8 |
| 6.8 | 6.9 | 7 | 1 |
| 7.3 | 8.4 | 3 | 5 |
| 6.3 | 6.8 | 0 | 8 |
| 6.8 | 7.0 | 6 | 2 |
| 6.7 | 8.2 | 2 | 6 |
