## Supplementary material for "Enhancing Wnt signaling lowers fracture incidence in a severe mouse model of Osteogenesis Imperfecta": SupTable2-tab1

| ID | Genotype | Sex | Age (Weeks) | MW-Total Score | GM-Total Score | IH-Total Score | IB-Total Score |
|---|---|---|---|---|---|---|---|
| 164 | OI/AV | F | 13 | 8 | 9 | 11 | 10 |
| 164 | OI/AV | F | 5 | 6 | 8 | 11 | 8 |
| 164 | OI/AV | F | 9 | 8 | 9 | 11 | 9 |
| 165 | OI/AV | F | 13 | 3 | 3 | 3 | 4 |
| 165 | OI/AV | F | 5 | 3 | 4 | 3 | 3 |
| 165 | OI/AV | F | 9 | 4 | 4 | 3 | 4 |
| 1909 | OI/AV | F | 13 | 3 | 4 | 1 | 4 |
| 1909 | OI/AV | F | 5 | 3 | 4 | 1 | 3 |
| 1909 | OI/AV | F | 9 | 3 | 4 | 1 | 4 |
| 1930 | OI/AV | F | 13 | 7 | 8 | 8 | 8 |
| 1930 | OI/AV | F | 5 | 6 | 7 | 7 | 8 |
| 1930 | OI/AV | F | 9 | 7 | 9 | 6 | 8 |
| 1933 | OI/AV | F | 13 | 1 | 4 | 5 | 3 |
| 1933 | OI/AV | F | 5 | 2 | 1 | 3 | 3 |
| 1933 | OI/AV | F | 9 | 1 | 2 | 3 | 3 |
| 1934 | OI/AV | M | 13 | 3 | 6 | 6 | 2 |
| 1934 | OI/AV | M | 5 | 2 | 7 | 4 | 3 |
| 1934 | OI/AV | M | 9 | 3 | 7 | 5 | 3 |
| 1935 | OI/AV | M | 13 | 4 | 5 | 6 | 4 |
| 1935 | OI/AV | M | 5 | 4 | 5 | 6 | 4 |
| 1935 | OI/AV | M | 9 | 4 | 5 | 6 | 4 |
| 196 | OI/AV | F | 13 | 6 | 8 | 7 | 5 |
| 196 | OI/AV | F | 5 | 5 | 8 | 6 | 5 |
| 196 | OI/AV | F | 9 | 5 | 8 | 6 | 5 |
| 197 | OI/AV | F | 13 | 7 | 8 | 8 | 8 |
| 197 | OI/AV | F | 5 | 7 | 8 | 9 | 8 |
| 197 | OI/AV | F | 9 | 7 | 8 | 9 | 8 |
| 318 | OI/AV | F | 13 | 7 | 7 | 7 | 8 |
| 318 | OI/AV | F | 5 | 7 | 8 | 7 | 7 |
| 318 | OI/AV | F | 9 | 7 | 8 | 9 | 8 |
| 320 | OI/AV | M | 13 | 8 | 8 | 9 | 8 |
| 320 | OI/AV | M | 5 | 6 | 6 | 6 | 6 |
| 320 | OI/AV | M | 9 | 8 | 8 | 8 | 8 |
| 321 | OI/AV | M | 13 | 4 | 6 | 4 | 4 |
| 321 | OI/AV | M | 5 | 4 | 4 | 3 | 4 |
| 321 | OI/AV | M | 9 | 4 | 4 | 4 | 4 |
| 339 | OI/AV | F | 13 | 4 | 6 | 4 | 6 |
| 339 | OI/AV | F | 5 | 4 | 7 | 4 | 8 |
| 339 | OI/AV | F | 9 | 4 | 6 | 4 | 6 |
| 340 | OI/AV | F | 13 | 4 | 6 | 4 | 4 |

| | | | | | | | |
|---|---|---|---|---|---|---|---|
| 340 | OI/AV | F | 5 | 5 | 5 | 3 | 3 |
| 340 | OI/AV | F | 9 | 4 | 5 | 4 | 4 |
| 350 | OI/AV | M | 13 | 6 | 6 | 6 | 7 |
| 350 | OI/AV | M | 5 | 5 | 6 | 5 | 6 |
| 350 | OI/AV | M | 9 | 6 | 6 | 5 | 6 |
| 3515 | OI/AV | M | 13 | 6 | 6 | 6 | 6 |
| 3515 | OI/AV | M | 5 | 7 | 6 | 7 | 6 |
| 3515 | OI/AV | M | 9 | 6 | 7 | 6 | 6 |
| 3522 | OI/AV | M | 13 | 2 | 4 | 3 | 6 |
| 3522 | OI/AV | M | 5 | 2 | 4 | 3 | 3 |
| 3522 | OI/AV | M | 9 | 2 | 4 | 3 | 8 |
| 3523 | OI/AV | M | 13 | 3 | 3 | 3 | 4 |
| 3523 | OI/AV | M | 5 | 3 | 5 | 2 | 2 |
| 3523 | OI/AV | M | 9 | 3 | 2 | 3 | 3 |
| 4113 | OI/AV | F | 13 | 6 | 8 | 6 | 8 |
| 4113 | OI/AV | F | 5 | 6 | 8 | 6 | 8 |
| 4113 | OI/AV | F | 9 | 7 | 8 | 6 | 8 |
| 4117 | OI/AV | M | 13 | 3 | 4 | 4 | 4 |
| 4117 | OI/AV | M | 5 | 4 | 5 | 3 | 4 |
| 4117 | OI/AV | M | 9 | 3 | 5 | 4 | 4 |
| 4591 | OI/AV | F | 13 | 7 | 7 | 7 | 7 |
| 4591 | OI/AV | F | 5 | 6 | 7 | 6 | 5 |
| 4591 | OI/AV | F | 9 | 7 | 7 | 6 | 7 |
| 4592 | OI/AV | F | 13 | 8 | 8 | 7 | 8 |
| 4592 | OI/AV | F | 5 | 8 | 8 | 6 | 8 |
| 4592 | OI/AV | F | 9 | 8 | 8 | 7 | 8 |
| 4594 | OI/AV | F | 13 | 7 | 8 | 7 | 8 |
| 4594 | OI/AV | F | 5 | 7 | 9 | 6 | 6 |
| 4594 | OI/AV | F | 9 | 7 | 8 | 7 | 8 |
| 4598 | OI/AV | M | 13 | 9 | 8 | 7 | 9 |
| 4598 | OI/AV | M | 5 | 7 | 6 | 5 | 5 |
| 4598 | OI/AV | M | 9 | 7 | 7 | 7 | 7 |
| 4686 | OI/AV | F | 13 | 5 | 6 | 5 | 8 |
| 4686 | OI/AV | F | 5 | 4 | 6 | 3 | 5 |
| 4686 | OI/AV | F | 9 | 5 | 7 | 5 | 7 |
| 992 | OI/AV | F | 13 | 4 | 9 | 3 | 2 |
| 992 | OI/AV | F | 5 | 4 | 4 | 4 | 5 |
| 992 | OI/AV | F | 9 | 4 | 4 | 3 | 2 |
| 1163 | OI | F | 13 | 8 | 10 | 9 | 9 |
| 1163 | OI | F | 5 | 9 | 7 | 8 | 9 |
| 1163 | OI | F | 9 | 9 | 9 | 9 | 6 |

| | | | | | | | |
|---|---|---|---|---|---|---|---|
| 1210 | OI | M | 13 | 9 | 10 | 7 | 6 |
| 1210 | OI | M | 5 | 9 | 8 | 5 | 7 |
| 1210 | OI | M | 9 | 9 | 9 | 7 | 6 |
| 1211 | OI | M | 13 | 8 | 11 | 8 | 7 |
| 1211 | OI | M | 5 | 8 | 9 | 5 | 6 |
| 1211 | OI | M | 9 | 10 | 11 | 7 | 6 |
| 1214 | OI | M | 13 | 5 | 7 | 4 | 4 |
| 1214 | OI | M | 5 | 5 | 8 | 4 | 5 |
| 1214 | OI | M | 9 | 5 | 8 | 5 | 4 |
| 1910 | OI | F | 13 | 3 | 7 | 2 | 5 |
| 1910 | OI | F | 5 | 3 | 4 | 2 | 4 |
| 1910 | OI | F | 9 | 4 | 5 | 2 | 5 |
| 198 | OI | F | 13 | 10 | 9 | 9 | 11 |
| 198 | OI | F | 5 | 9 | 7 | 8 | 10 |
| 198 | OI | F | 9 | 8 | 11 | 8 | 11 |
| 319 | OI | M | 13 | 8 | 9 | 9 | 11 |
| 319 | OI | M | 5 | 6 | 9 | 8 | 8 |
| 319 | OI | M | 9 | 8 | 12 | 9 | 9 |
| 344 | OI | M | 13 | 9 | 8 | 7 | 9 |
| 344 | OI | M | 5 | 8 | 9 | 9 | 8 |
| 344 | OI | M | 9 | 9 | 9 | 7 | 9 |
| 345 | OI | F | 13 | 6 | 6 | 7 | 7 |
| 345 | OI | F | 5 | 7 | 7 | 7 | 6 |
| 345 | OI | F | 9 | 5 | 7 | 7 | 6 |
| 349 | OI | M | 13 | 8 | 8 | 5 | 7 |
| 349 | OI | M | 5 | 7 | 8 | 7 | 7 |
| 349 | OI | M | 9 | 8 | 9 | 8 | 8 |
| 351 | OI | M | 13 | 6 | 6 | 4 | 6 |
| 351 | OI | M | 5 | 7 | 6 | 4 | 6 |
| 351 | OI | M | 9 | 6 | 6 | 4 | 6 |
| 3520 | OI | F | 5 | 6 | 6 | 6 | 6 |
| 3520 | OI | F | 9 | 8 | 9 | 8 | 8 |
| 4115 | OI | F | 13 | 6 | 6 | 5 | 6 |
| 4115 | OI | F | 5 | 7 | 7 | 4 | 6 |
| 4115 | OI | F | 9 | 6 | 6 | 5 | 6 |
| 4596 | OI | M | 13 | 11 | 13 | 9 | 7 |
| 4596 | OI | M | 5 | 10 | 12 | 10 | 9 |
| 4596 | OI | M | 9 | 11 | 13 | 10 | 9 |
| 4690 | OI | M | 13 | 8 | 8 | 6 | 7 |
| 4690 | OI | M | 5 | 7 | 8 | 5 | 6 |
| 4690 | OI | M | 9 | 8 | 9 | 4 | 7 |

| | | | | | | | |
|---|---|---|---|---|---|---|---|
| 873 | OI | M | 13 | 8 | 12 | 7 | 8 |
| 873 | OI | M | 5 | 8 | 9 | 7 | 8 |
| 873 | OI | M | 9 | 8 | 9 | 8 | 9 |
| 941 | OI | F | 5 | 9 | 10 | 6 | 10 |
| 943 | OI | M | 5 | 9 | 10 | 6 | 10 |
| 987 | OI | F | 13 | 8 | 11 | 7 | 9 |
| 987 | OI | F | 5 | 10 | 9 | 8 | 9 |
| 987 | OI | F | 9 | 11 | 11 | 8 | 9 |
| 989 | OI | F | 13 | 8 | 9 | 8 | 8 |
| 989 | OI | F | 5 | 7 | 7 | 7 | 8 |
| 989 | OI | F | 9 | 8 | 8 | 7 | 7 |

| IE-Total Score | RC-Total Score | DN-Total Score | SV-Total Score |
|---|---|---|---|
| 10 | 10 | 9 | 9 |
| 8 | 8 | 8 | 7 |
| 9 | 10 | 9 | 9 |
| 3 | 3 | 3 | 3 |
| 4 | 4 | 3 | 3 |
| 3 | 3 | 3 | 3 |
| 4 | 4 | 4 | 4 |
| 4 | 4 | 4 | 4 |
| 4 | 4 | 4 | 4 |
| 8 | 7 | 9 | 8 |
| 8 | 6 | 9 | 9 |
| 8 | 7 | 9 | 7 |
| 3 | 3 | 3 | 3 |
| 3 | 3 | 3 | 3 |
| 3 | 3 | 3 | 3 |
| 3 | 2 | 4 | 2 |
| 3 | 2 | 4 | 2 |
| 3 | 2 | 4 | 2 |
| 4 | 4 | 4 | 4 |
| 4 | 4 | 4 | 4 |
| 4 | 4 | 4 | 4 |
| 6 | 7 | 6 | 5 |
| 6 | 6 | 6 | 6 |
| 5 | 6 | 6 | 6 |
| 8 | 8 | 8 | 8 |
| 7 | 8 | 8 | 9 |
| 8 | 8 | 8 | 6 |
| 7 | 7 | 7 | 6 |
| 7 | 7 | 7 | 7 |
| 7 | 7 | 7 | 6 |
| 8 | 8 | 8 | 8 |
| 6 | 6 | 6 | 6 |
| 8 | 8 | 8 | 6 |
| 4 | 4 | 4 | 4 |
| 4 | 4 | 4 | 4 |
| 4 | 4 | 4 | 4 |
| 3 | 3 | 3 | 3 |
| 4 | 3 | 4 | 4 |
| 3 | 3 | 3 | 3 |
| 5 | 3 | 5 | 3 |

| | | | |
|---|---|---|---|
| 4 | 3 | 4 | 4 |
| 6 | 3 | 5 | 3 |
| 6 | 6 | 7 | 5 |
| 6 | 6 | 7 | 6 |
| 6 | 6 | 7 | 7 |
| 6 | 6 | 6 | 6 |
| 6 | 6 | 6 | 8 |
| 6 | 6 | 6 | 6 |
| 4 | 4 | 4 | 4 |
| 4 | 4 | 4 | 4 |
| 4 | 4 | 4 | 4 |
| 4 | 4 | 4 | 3 |
| 4 | 4 | 4 | 4 |
| 4 | 4 | 4 | 3 |
| 7 | 6 | 9 | 7 |
| 8 | 6 | 9 | 7 |
| 9 | 6 | 9 | 6 |
| 4 | 4 | 4 | 4 |
| 4 | 4 | 4 | 4 |
| 4 | 4 | 4 | 4 |
| 7 | 7 | 8 | 7 |
| 7 | 7 | 8 | 8 |
| 7 | 7 | 8 | 9 |
| 8 | 8 | 8 | 8 |
| 8 | 8 | 8 | 8 |
| 8 | 8 | 8 | 8 |
| 8 | 7 | 8 | 8 |
| 8 | 7 | 8 | 8 |
| 8 | 7 | 8 | 7 |
| 8 | 8 | 8 | 7 |
| 6 | 6 | 6 | 7 |
| 7 | 7 | 6 | 5 |
| 5 | 5 | 8 | 4 |
| 5 | 5 | 6 | 5 |
| 5 | 5 | 6 | 5 |
| 5 | 4 | 5 | 4 |
| 4 | 4 | 6 | 4 |
| 4 | 4 | 4 | 5 |
| 10 | 10 | 11 | 10 |
| 10 | 10 | 11 | 11 |
| 10 | 10 | 11 | 9 |

| | | | |
|---|---|---|---|
| 9 | 9 | 9 | 8 |
| 9 | 8 | 9 | 8 |
| 9 | 9 | 9 | 8 |
| 10 | 10 | 11 | 8 |
| 9 | 10 | 11 | 9 |
| 9 | 10 | 11 | 8 |
| 7 | 7 | 8 | 7 |
| 7 | 7 | 6 | 7 |
| 7 | 7 | 7 | 7 |
| 5 | 4 | 5 | 4 |
| 5 | 4 | 4 | 5 |
| 5 | 4 | 5 | 5 |
| 11 | 9 | 11 | 9 |
| 11 | 11 | 11 | 9 |
| 11 | 10 | 10 | 9 |
| 10 | 9 | 8 | 8 |
| 8 | 6 | 8 | 7 |
| 9 | 9 | 8 | 8 |
| 9 | 8 | 9 | 7 |
| 9 | 9 | 8 | 8 |
| 9 | 9 | 9 | 10 |
| 7 | 7 | 7 | 7 |
| 6 | 6 | 7 | 8 |
| 6 | 6 | 7 | 6 |
| 8 | 8 | 9 | 7 |
| 8 | 8 | 8 | 8 |
| 8 | 8 | 9 | 8 |
| 6 | 4 | 6 | 5 |
| 6 | 6 | 6 | 5 |
| 6 | 6 | 6 | 5 |
| 7 | 6 | 7 | 8 |
| 7 | 7 | 8 | 7 |
| 5 | 5 | 6 | 5 |
| 7 | 6 | 7 | 8 |
| 6 | 5 | 6 | 7 |
| 11 | 11 | 11 | 11 |
| 11 | 11 | 12 | 13 |
| 11 | 11 | 11 | 10 |
| 9 | 7 | 7 | 7 |
| 8 | 6 | 7 | 6 |
| 8 | 7 | 7 | 7 |

| | | | |
|---|---|---|---|
| 9 | 8 | 9 | 10 |
| 9 | 8 | 8 | 10 |
| 9 | 8 | 9 | 10 |
| 10 | 10 | 10 | 10 |
| 10 | 9 | 9 | 9 |
| 10 | 11 | 11 | 12 |
| 11 | 9 | 11 | 10 |
| 11 | 10 | 11 | 11 |
| 9 | 8 | 9 | 10 |
| 9 | 9 | 9 | 10 |
| 9 | 8 | 9 | 9 |

Anova: Two-Factor Wit

| SUMMARY | Count | Sum | Average | Variance |
|---|---|---|---|---|
| 164-13 | 8 | 76 | 9.5 | 0.85714286 |
| 164-5 | 8 | 64 | 8 | 2 |
| 164-9 | 8 | 74 | 9.25 | 0.78571429 |
| 165-13 | 8 | 25 | 3.125 | 0.125 |
| 165-5 | 8 | 27 | 3.375 | 0.26785714 |
| 165-9 | 8 | 27 | 3.375 | 0.26785714 |
| 1909-13 | 8 | 28 | 3.5 | 1.14285714 |
| 1909-5 | 8 | 27 | 3.375 | 1.125 |
| 1909-9 | 8 | 28 | 3.5 | 1.14285714 |
| 1930-13 | 8 | 63 | 7.875 | 0.41071429 |
| 1930-5 | 8 | 60 | 7.5 | 1.42857143 |
| 1930-9 | 8 | 61 | 7.625 | 1.125 |
| 1933-13 | 8 | 25 | 3.125 | 1.26785714 |
| 1933-5 | 8 | 21 | 2.625 | 0.55357143 |
| 1933-9 | 8 | 21 | 2.625 | 0.55357143 |
| 1934-13 | 8 | 28 | 3.5 | 2.85714286 |
| 1934-5 | 8 | 27 | 3.375 | 2.83928571 |
| 1934-9 | 8 | 29 | 3.625 | 2.83928571 |
| 1935-13 | 8 | 35 | 4.375 | 0.55357143 |
| 1935-5 | 8 | 35 | 4.375 | 0.55357143 |
| 1935-9 | 8 | 35 | 4.375 | 0.55357143 |
| 196-13 | 8 | 50 | 6.25 | 1.07142857 |
| 196-5 | 8 | 48 | 6 | 0.85714286 |
| 196-9 | 8 | 47 | 5.875 | 0.98214286 |
| 197-13 | 8 | 63 | 7.875 | 0.125 |
| 197-5 | 8 | 64 | 8 | 0.57142857 |
| 197-9 | 8 | 62 | 7.75 | 0.78571429 |
| 318-13 | 8 | 56 | 7 | 0.28571429 |
| 318-5 | 8 | 57 | 7.125 | 0.125 |
| 318-9 | 8 | 59 | 7.375 | 0.83928571 |
| 320-13 | 8 | 65 | 8.125 | 0.125 |
| 320-5 | 8 | 48 | 6 | 0 |
| 320-9 | 8 | 62 | 7.75 | 0.5 |
| 321-13 | 8 | 34 | 4.25 | 0.5 |
| 321-5 | 8 | 31 | 3.875 | 0.125 |
| 321-9 | 8 | 32 | 4 | 0 |
| 339-13 | 8 | 32 | 4 | 1.71428571 |
| 339-5 | 8 | 38 | 4.75 | 3.07142857 |
| 339-9 | 8 | 32 | 4 | 1.71428571 |
| 340-13 | 8 | 34 | 4.25 | 1.07142857 |

| | | | | |
|---|---|---|---|---|
| 340-5 | 8 | 31 | 3.875 | 0.69642857 |
| 340-9 | 8 | 34 | 4.25 | 1.07142857 |
| 350-13 | 8 | 49 | 6.125 | 0.41071429 |
| 350-5 | 8 | 47 | 5.875 | 0.41071429 |
| 350-9 | 8 | 49 | 6.125 | 0.41071429 |
| 3515-13 | 8 | 48 | 6 | 0 |
| 3515-5 | 8 | 52 | 6.5 | 0.57142857 |
| 3515-9 | 8 | 49 | 6.125 | 0.125 |
| 3522-13 | 8 | 31 | 3.875 | 1.26785714 |
| 3522-5 | 8 | 28 | 3.5 | 0.57142857 |
| 3522-9 | 8 | 33 | 4.125 | 2.98214286 |
| 3523-13 | 8 | 28 | 3.5 | 0.28571429 |
| 3523-5 | 8 | 28 | 3.5 | 1.14285714 |
| 3523-9 | 8 | 26 | 3.25 | 0.5 |
| 4113-13 | 8 | 57 | 7.125 | 1.26785714 |
| 4113-5 | 8 | 58 | 7.25 | 1.35714286 |
| 4113-9 | 8 | 59 | 7.375 | 1.69642857 |
| 4117-13 | 8 | 31 | 3.875 | 0.125 |
| 4117-5 | 8 | 32 | 4 | 0.28571429 |
| 4117-9 | 8 | 32 | 4 | 0.28571429 |
| 4591-13 | 8 | 57 | 7.125 | 0.125 |
| 4591-5 | 8 | 54 | 6.75 | 1.07142857 |
| 4591-9 | 8 | 58 | 7.25 | 0.78571429 |
| 4592-13 | 8 | 63 | 7.875 | 0.125 |
| 4592-5 | 8 | 62 | 7.75 | 0.5 |
| 4592-9 | 8 | 63 | 7.875 | 0.125 |
| 4594-13 | 8 | 61 | 7.625 | 0.26785714 |
| 4594-5 | 8 | 59 | 7.375 | 1.125 |
| 4594-9 | 8 | 60 | 7.5 | 0.28571429 |
| 4598-13 | 8 | 64 | 8 | 0.57142857 |
| 4598-5 | 8 | 48 | 6 | 0.57142857 |
| 4598-9 | 8 | 53 | 6.625 | 0.55357143 |
| 4686-13 | 8 | 46 | 5.75 | 2.21428571 |
| 4686-5 | 8 | 39 | 4.875 | 0.98214286 |
| 4686-9 | 8 | 45 | 5.625 | 0.83928571 |
| 992-13 | 8 | 36 | 4.5 | 4.28571429 |
| 992-5 | 8 | 35 | 4.375 | 0.55357143 |
| 992-9 | 8 | 30 | 3.75 | 0.78571429 |
| 1163-13 | 8 | 77 | 9.625 | 0.83928571 |
| 1163-5 | 8 | 75 | 9.375 | 1.98214286 |
| 1163-9 | 8 | 73 | 9.125 | 2.125 |

| | | | | |
|---|---|---|---|---|
| 1210-13 | 8 | 67 | 8.375 | 1.69642857 |
| 1210-5 | 8 | 63 | 7.875 | 1.83928571 |
| 1210-9 | 8 | 66 | 8.25 | 1.35714286 |
| 1211-13 | 8 | 73 | 9.125 | 2.41071429 |
| 1211-5 | 8 | 67 | 8.375 | 3.98214286 |
| 1211-9 | 8 | 72 | 9 | 3.42857143 |
| 1214-13 | 8 | 49 | 6.125 | 2.41071429 |
| 1214-5 | 8 | 49 | 6.125 | 1.83928571 |
| 1214-9 | 8 | 50 | 6.25 | 1.92857143 |
| 1910-13 | 8 | 35 | 4.375 | 2.26785714 |
| 1910-5 | 8 | 31 | 3.875 | 0.98214286 |
| 1910-9 | 8 | 35 | 4.375 | 1.125 |
| 198-13 | 8 | 79 | 9.875 | 0.98214286 |
| 198-5 | 8 | 76 | 9.5 | 2.28571429 |
| 198-9 | 8 | 78 | 9.75 | 1.64285714 |
| 319-13 | 8 | 72 | 9 | 1.14285714 |
| 319-5 | 8 | 60 | 7.5 | 1.14285714 |
| 319-9 | 8 | 72 | 9 | 1.71428571 |
| 344-13 | 8 | 66 | 8.25 | 0.78571429 |
| 344-5 | 8 | 68 | 8.5 | 0.28571429 |
| 344-9 | 8 | 71 | 8.875 | 0.69642857 |
| 345-13 | 8 | 54 | 6.75 | 0.21428571 |
| 345-5 | 8 | 54 | 6.75 | 0.5 |
| 345-9 | 8 | 50 | 6.25 | 0.5 |
| 349-13 | 8 | 60 | 7.5 | 1.42857143 |
| 349-5 | 8 | 61 | 7.625 | 0.26785714 |
| 349-9 | 8 | 66 | 8.25 | 0.21428571 |
| 351-13 | 8 | 43 | 5.375 | 0.83928571 |
| 351-5 | 8 | 46 | 5.75 | 0.78571429 |
| 351-9 | 8 | 45 | 5.625 | 0.55357143 |
| 3520-5 | 8 | 52 | 6.5 | 0.57142857 |
| 3520-9 | 8 | 62 | 7.75 | 0.5 |
| 4115-13 | 8 | 44 | 5.5 | 0.28571429 |
| 4115-5 | 8 | 52 | 6.5 | 1.42857143 |
| 4115-9 | 8 | 47 | 5.875 | 0.41071429 |
| 4596-13 | 8 | 84 | 10.5 | 3.14285714 |
| 4596-5 | 8 | 88 | 11 | 1.71428571 |
| 4596-9 | 8 | 86 | 10.75 | 1.35714286 |
| 4690-13 | 8 | 59 | 7.375 | 0.83928571 |
| 4690-5 | 8 | 53 | 6.625 | 1.125 |
| 4690-9 | 8 | 57 | 7.125 | 2.125 |

| | | | | |
|---|---|---|---|---|
| 873-13 | 8 | 71 | 8.875 | 2.41071429 |
| 873-5 | 8 | 67 | 8.375 | 0.83928571 |
| 873-9 | 8 | 70 | 8.75 | 0.5 |
| 941-5 | 8 | 75 | 9.375 | 1.98214286 |
| 943-5 | 8 | 72 | 9 | 1.71428571 |
| 987-13 | 8 | 79 | 9.875 | 2.98214286 |
| 987-5 | 8 | 77 | 9.625 | 1.125 |
| 987-9 | 8 | 82 | 10.25 | 1.35714286 |
| 989-13 | 8 | 69 | 8.625 | 0.55357143 |
| 989-5 | 8 | 66 | 8.25 | 1.35714286 |
| 989-9 | 8 | 65 | 8.125 | 0.69642857 |
| MW-Total Sco | 133 | 817 | 6.14285714 | 5.27489177 |
| GM-Total Sco | 133 | 952 | 7.15789474 | 5.14912281 |
| IH-Total Score | 133 | 779 | 5.85714286 | 4.95670996 |
| IB-Total Score | 133 | 849 | 6.38345865 | 4.63214855 |
| IE-Total Score | 133 | 902 | 6.78195489 | 5.45967191 |
| KC-Total Scor | 133 | 855 | 6.42857143 | 5.47402597 |
| DN-Total Scor | 133 | 926 | 6.96240602 | 5.68796992 |
| SV-Total Scor | 133 | 865 | 6.5037594 | 5.78218273 |

ANOVA

| *Source of Variati* | *SS* | *df* | *MS* | *F* |
|---|---|---|---|---|
| Rows | 4760.08459 | 132 | 36.0612469 | 39.7182993 |
| Columns | 170.202068 | 7 | 24.3145811 | 26.7803776 |
| Error | 838.922932 | 924 | 0.90792525 | |
| Total | 5769.20959 | 1063 | | |

| | |
|---|---|
| ICC | 0.80213595 |

| *P-value* | *F crit* |
|---|---|
| 5.9375E-302 | 1.229826 |
| 1.44641E-33 | 2.01947239 |
