## Supplementary material for "Enhancing Wnt signaling lowers fracture incidence in a severe mouse model of Osteogenesis Imperfecta": SupTable2-tab2

| ID | Genotype | Sex | Age (Weeks) | MW-Scoliosis | GM-Scoliosis | IH-Scoliosis | IB-Scoliosis |
|---|---|---|---|---|---|---|---|
| 164 | OI/AV | F | 13 | Y | N | Y | Y |
| 164 | OI/AV | F | 5 | N | N | N | N |
| 164 | OI/AV | F | 9 | Y | N | N | Y |
| 165 | OI/AV | F | 13 | N | N | N | N |
| 165 | OI/AV | F | 5 | N | N | N | N |
| 165 | OI/AV | F | 9 | Y | N | N | N |
| 1909 | OI/AV | F | 13 | N | N | N | N |
| 1909 | OI/AV | F | 5 | N | N | N | N |
| 1909 | OI/AV | F | 9 | N | N | N | N |
| 1930 | OI/AV | F | 13 | N | N | N | N |
| 1930 | OI/AV | F | 5 | N | N | N | N |
| 1930 | OI/AV | F | 9 | N | N | N | N |
| 1933 | OI/AV | F | 13 | N | N | Y | N |
| 1933 | OI/AV | F | 5 | N | N | N | N |
| 1933 | OI/AV | F | 9 | N | N | N | N |
| 1934 | OI/AV | M | 13 | N | N | N | N |
| 1934 | OI/AV | M | 5 | N | N | N | N |
| 1934 | OI/AV | M | 9 | N | N | N | N |
| 1935 | OI/AV | M | 13 | N | N | Y | N |
| 1935 | OI/AV | M | 5 | N | N | N | N |
| 1935 | OI/AV | M | 9 | N | N | Y | N |
| 196 | OI/AV | F | 13 | Y | Y | Y | Y |
| 196 | OI/AV | F | 5 | Y | Y | Y | Y |
| 196 | OI/AV | F | 9 | Y | Y | Y | Y |
| 197 | OI/AV | F | 13 | Y | Y | Y | Y |
| 197 | OI/AV | F | 5 | Y | Y | Y | Y |
| 197 | OI/AV | F | 9 | Y | Y | Y | Y |
| 318 | OI/AV | F | 13 | N | N | N | N |
| 318 | OI/AV | F | 5 | N | N | N | N |
| 318 | OI/AV | F | 9 | N | N | N | N |
| 320 | OI/AV | M | 13 | N | N | N | N |
| 320 | OI/AV | M | 5 | N | N | N | N |
| 320 | OI/AV | M | 9 | N | N | N | N |
| 321 | OI/AV | M | 13 | N | N | N | N |
| 321 | OI/AV | M | 5 | N | N | N | N |
| 321 | OI/AV | M | 9 | N | Y | N | N |
| 339 | OI/AV | F | 13 | N | N | N | N |
| 339 | OI/AV | F | 5 | N | N | N | N |

| | | | | | | | |
|---|---|---|---|---|---|---|---|
| 339 | OI/AV | F | 9 | N | N | N | N |
| 340 | OI/AV | F | 13 | N | Y | Y | N |
| 340 | OI/AV | F | 5 | N | N | N | N |
| 340 | OI/AV | F | 9 | N | N | N | N |
| 350 | OI/AV | M | 13 | N | N | N | N |
| 350 | OI/AV | M | 5 | N | N | N | N |
| 350 | OI/AV | M | 9 | N | N | N | N |
| 3515 | OI/AV | M | 13 | Y | N | Y | Y |
| 3515 | OI/AV | M | 5 | Y | Y | Y | Y |
| 3515 | OI/AV | M | 9 | Y | Y | Y | Y |
| 3522 | OI/AV | M | 13 | N | N | N | Y |
| 3522 | OI/AV | M | 5 | N | N | N | N |
| 3522 | OI/AV | M | 9 | N | N | N | N |
| 3523 | OI/AV | M | 13 | N | N | Y | N |
| 3523 | OI/AV | M | 5 | N | N | N | N |
| 3523 | OI/AV | M | 9 | N | N | N | N |
| 4113 | OI/AV | F | 13 | N | N | Y | N |
| 4113 | OI/AV | F | 5 | N | N | N | N |
| 4113 | OI/AV | F | 9 | N | N | N | N |
| 4117 | OI/AV | M | 13 | N | N | Y | N |
| 4117 | OI/AV | M | 5 | N | N | N | N |
| 4117 | OI/AV | M | 9 | N | N | N | N |
| 4591 | OI/AV | F | 13 | Y | Y | Y | Y |
| 4591 | OI/AV | F | 5 | Y | Y | Y | Y |
| 4591 | OI/AV | F | 9 | Y | Y | Y | Y |
| 4592 | OI/AV | F | 13 | Y | Y | Y | N |
| 4592 | OI/AV | F | 5 | Y | Y | Y | N |
| 4592 | OI/AV | F | 9 | Y | Y | Y | N |
| 4594 | OI/AV | F | 13 | Y | Y | Y | N |
| 4594 | OI/AV | F | 5 | N | N | N | N |
| 4594 | OI/AV | F | 9 | Y | Y | Y | N |
| 4598 | OI/AV | M | 13 | Y | Y | Y | Y |
| 4598 | OI/AV | M | 5 | Y | Y | Y | Y |
| 4598 | OI/AV | M | 9 | Y | Y | Y | Y |
| 4686 | OI/AV | F | 13 | N | N | Y | N |
| 4686 | OI/AV | F | 5 | N | N | Y | N |
| 4686 | OI/AV | F | 9 | N | N | Y | N |
| 992 | OI/AV | F | 13 | N | Y | Y | N |

| | | | | | | | |
|---|---|---|---|---|---|---|---|
| 992 | OI/AV | F | 5 | N | N | N | N |
| 992 | OI/AV | F | 9 | N | N | N | N |
| 1163 | OI | F | 13 | Y | Y | Y | N |
| 1163 | OI | F | 5 | Y | Y | Y | N |
| 1163 | OI | F | 9 | N | Y | Y | N |
| 1210 | OI | M | 13 | Y | Y | Y | Y |
| 1210 | OI | M | 5 | Y | Y | Y | Y |
| 1210 | OI | M | 9 | Y | Y | Y | Y |
| 1211 | OI | M | 13 | Y | Y | Y | Y |
| 1211 | OI | M | 5 | Y | Y | Y | Y |
| 1211 | OI | M | 9 | Y | Y | Y | Y |
| 1214 | OI | M | 13 | Y | Y | N | N |
| 1214 | OI | M | 5 | N | N | N | N |
| 1214 | OI | M | 9 | N | N | N | N |
| 1910 | OI | F | 13 | N | N | N | N |
| 1910 | OI | F | 5 | N | N | N | N |
| 1910 | OI | F | 9 | N | N | N | N |
| 198 | OI | F | 13 | Y | Y | Y | Y |
| 198 | OI | F | 5 | Y | Y | Y | Y |
| 198 | OI | F | 9 | Y | Y | Y | Y |
| 319 | OI | M | 13 | Y | Y | Y | Y |
| 319 | OI | M | 5 | Y | Y | Y | N |
| 319 | OI | M | 9 | Y | Y | Y | Y |
| 344 | OI | M | 13 | Y | Y | Y | Y |
| 344 | OI | M | 5 | Y | Y | Y | Y |
| 344 | OI | M | 9 | Y | Y | Y | Y |
| 345 | OI | F | 13 | Y | Y | Y | Y |
| 345 | OI | F | 5 | Y | Y | Y | Y |
| 345 | OI | F | 9 | Y | Y | Y | Y |
| 349 | OI | M | 13 | Y | Y | Y | N |
| 349 | OI | M | 5 | Y | Y | Y | N |
| 349 | OI | M | 9 | Y | Y | Y | N |
| 351 | OI | M | 13 | N | N | N | N |
| 351 | OI | M | 5 | N | N | N | N |
| 351 | OI | M | 9 | N | N | N | N |
| 3520 | OI | F | 5 | Y | Y | Y | N |
| 3520 | OI | F | 9 | Y | Y | Y | N |
| 4115 | OI | F | 13 | N | N | Y | N |

| | | | | | | | |
|---|---|---|---|---|---|---|---|
| 4115 | OI | F | 5 | N | Y | Y | N |
| 4115 | OI | F | 9 | N | Y | Y | N |
| 4596 | OI | M | 13 | N | N | N | N |
| 4596 | OI | M | 5 | N | N | N | N |
| 4596 | OI | M | 9 | N | N | N | N |
| 4690 | OI | M | 13 | N | N | Y | N |
| 4690 | OI | M | 5 | N | N | Y | N |
| 4690 | OI | M | 9 | N | N | Y | N |
| 873 | OI | M | 13 | N | Y | Y | Y |
| 873 | OI | M | 5 | Y | Y | Y | Y |
| 873 | OI | M | 9 | N | Y | Y | Y |
| 941 | OI | F | 5 | Y | Y | Y | N |
| 943 | OI | M | 5 | Y | Y | Y | Y |
| 987 | OI | F | 13 | N | Y | Y | Y |
| 987 | OI | F | 5 | Y | Y | Y | N |
| 987 | OI | F | 9 | Y | Y | Y | Y |
| 989 | OI | F | 13 | N | Y | Y | Y |
| 989 | OI | F | 5 | Y | Y | Y | N |
| 989 | OI | F | 9 | N | Y | Y | Y |

| IE-Scoliosis | KC-Scoliosis | DN-Scoliosis | SV-Scoliosis | Scorers said yes |
|---|---|---|---|---|
| N | N | N | N | 3 |
| N | N | N | N | 0 |
| N | N | N | N | 2 |
| N | N | N | N | 0 |
| N | N | N | N | 0 |
| N | N | N | N | 1 |
| N | N | N | N | 0 |
| N | N | N | N | 0 |
| N | N | N | N | 0 |
| N | N | N | N | 0 |
| N | N | N | N | 0 |
| N | N | N | N | 0 |
| N | N | Y | N | 2 |
| N | N | Y | N | 1 |
| N | N | Y | N | 1 |
| N | N | N | N | 0 |
| N | N | N | N | 0 |
| N | N | N | N | 0 |
| N | N | N | N | 1 |
| N | N | N | N | 0 |
| N | N | N | N | 1 |
| N | Y | N | Y | 6 |
| N | Y | N | Y | 6 |
| N | Y | N | Y | 6 |
| N | Y | Y | Y | 7 |
| N | Y | Y | Y | 7 |
| N | Y | Y | Y | 7 |
| N | N | N | N | 0 |
| N | N | N | N | 0 |
| N | N | N | N | 0 |
| N | N | N | N | 0 |
| N | N | N | N | 0 |
| N | N | N | N | 0 |
| N | N | N | N | 0 |
| N | N | N | N | 0 |
| N | N | N | N | 1 |
| N | N | N | N | 0 |
| N | N | N | N | 0 |

| | | | | |
|---|---|---|---|---|
| N | N | N | N | 0 |
| N | N | N | N | 2 |
| N | N | N | N | 0 |
| N | N | N | N | 0 |
| N | N | N | N | 0 |
| N | N | N | N | 0 |
| N | N | N | N | 0 |
| N | N | Y | Y | 5 |
| N | N | Y | Y | 6 |
| N | N | Y | Y | 6 |
| N | N | N | N | 1 |
| N | N | N | N | 0 |
| N | N | N | N | 0 |
| N | N | N | N | 1 |
| N | N | N | N | 0 |
| N | N | N | N | 0 |
| N | N | N | N | 1 |
| N | N | N | N | 0 |
| N | N | N | N | 0 |
| N | Y | Y | N | 3 |
| N | Y | N | N | 1 |
| N | Y | Y | N | 2 |
| N | Y | N | Y | 6 |
| N | Y | N | Y | 6 |
| N | Y | N | Y | 6 |
| Y | Y | Y | Y | 7 |
| N | Y | Y | Y | 6 |
| N | Y | Y | Y | 6 |
| N | N | N | N | 3 |
| N | N | N | N | 0 |
| N | N | N | N | 3 |
| Y | Y | Y | Y | 8 |
| Y | Y | Y | Y | 8 |
| Y | Y | Y | Y | 8 |
| Y | N | Y | N | 3 |
| Y | N | Y | N | 3 |
| Y | N | Y | N | 3 |
| N | N | Y | N | 3 |

| | | | | |
|---|---|---|---|---|
| N | N | Y | N | 1 |
| N | N | Y | N | 1 |
| N | N | N | N | 3 |
| N | N | N | N | 3 |
| N | N | N | N | 2 |
| N | Y | N | Y | 6 |
| N | Y | N | Y | 6 |
| N | Y | N | Y | 6 |
| N | Y | N | N | 5 |
| N | Y | N | N | 5 |
| N | Y | N | N | 5 |
| N | N | N | N | 2 |
| N | N | N | N | 0 |
| N | N | N | N | 0 |
| N | Y | N | N | 1 |
| N | Y | N | N | 1 |
| N | Y | N | N | 1 |
| Y | Y | Y | Y | 8 |
| Y | Y | Y | Y | 8 |
| Y | Y | Y | Y | 8 |
| N | Y | Y | Y | 7 |
| N | Y | Y | Y | 6 |
| N | Y | Y | Y | 7 |
| N | N | Y | Y | 6 |
| N | N | Y | Y | 6 |
| N | N | Y | Y | 6 |
| Y | Y | Y | Y | 8 |
| Y | Y | Y | Y | 8 |
| Y | Y | Y | Y | 8 |
| N | N | N | N | 3 |
| N | N | N | N | 3 |
| N | N | N | N | 3 |
| N | N | N | N | 0 |
| N | N | N | N | 0 |
| N | N | N | N | 0 |
| N | Y | Y | Y | 6 |
| N | Y | Y | Y | 6 |
| N | Y | Y | Y | 4 |

| | | | | |
|---|---|---|---|---|
| N | Y | Y | Y | 5 |
| N | Y | Y | Y | 5 |
| N | N | N | N | 0 |
| N | N | N | N | 0 |
| N | N | N | N | 0 |
| N | Y | N | N | 2 |
| N | Y | N | N | 2 |
| N | Y | N | N | 2 |
| Y | Y | Y | Y | 7 |
| Y | Y | Y | Y | 8 |
| Y | Y | Y | Y | 7 |
| Y | N | N | Y | 5 |
| N | N | Y | Y | 6 |
| y | Y | Y | Y | 7 |
| Y | Y | Y | Y | 7 |
| Y | Y | Y | Y | 8 |
| Y | Y | Y | Y | 7 |
| Y | Y | Y | Y | 7 |
| Y | Y | Y | Y | 7 |

410
4.556
20.75

0.90413737

| Scorers said no | Sum | sum sq. c | sum sq. c/(9*8) |
|---|---|---|---|
| 5 | 8 | 26 | 0.361 |
| 8 | 8 | 56 | 0.778 |
| 6 | 8 | 32 | 0.444 |
| 8 | 8 | 56 | 0.778 |
| 8 | 8 | 56 | 0.778 |
| 7 | 8 | 42 | 0.583 |
| 8 | 8 | 56 | 0.778 |
| 8 | 8 | 56 | 0.778 |
| 8 | 8 | 56 | 0.778 |
| 8 | 8 | 56 | 0.778 |
| 8 | 8 | 56 | 0.778 |
| 8 | 8 | 56 | 0.778 |
| 6 | 8 | 32 | 0.444 |
| 7 | 8 | 42 | 0.583 |
| 7 | 8 | 42 | 0.583 |
| 8 | 8 | 56 | 0.778 |
| 8 | 8 | 56 | 0.778 |
| 8 | 8 | 56 | 0.778 |
| 7 | 8 | 42 | 0.583 |
| 8 | 8 | 56 | 0.778 |
| 7 | 8 | 42 | 0.583 |
| 2 | 8 | 32 | 0.444 |
| 2 | 8 | 32 | 0.444 |
| 2 | 8 | 32 | 0.444 |
| 1 | 8 | 42 | 0.583 |
| 1 | 8 | 42 | 0.583 |
| 1 | 8 | 42 | 0.583 |
| 8 | 8 | 56 | 0.778 |
| 8 | 8 | 56 | 0.778 |
| 8 | 8 | 56 | 0.778 |
| 8 | 8 | 56 | 0.778 |
| 8 | 8 | 56 | 0.778 |
| 8 | 8 | 56 | 0.778 |
| 8 | 8 | 56 | 0.778 |
| 8 | 8 | 56 | 0.778 |
| 7 | 8 | 42 | 0.583 |
| 8 | 8 | 56 | 0.778 |
| 8 | 8 | 56 | 0.778 |

| | | | |
|---|---|---|---|
| 8 | 8 | 56 | 0.778 |
| 6 | 8 | 32 | 0.444 |
| 8 | 8 | 56 | 0.778 |
| 8 | 8 | 56 | 0.778 |
| 8 | 8 | 56 | 0.778 |
| 8 | 8 | 56 | 0.778 |
| 8 | 8 | 56 | 0.778 |
| 3 | 8 | 26 | 0.361 |
| 2 | 8 | 32 | 0.444 |
| 2 | 8 | 32 | 0.444 |
| 7 | 8 | 42 | 0.583 |
| 8 | 8 | 56 | 0.778 |
| 8 | 8 | 56 | 0.778 |
| 7 | 8 | 42 | 0.583 |
| 8 | 8 | 56 | 0.778 |
| 8 | 8 | 56 | 0.778 |
| 7 | 8 | 42 | 0.583 |
| 8 | 8 | 56 | 0.778 |
| 8 | 8 | 56 | 0.778 |
| 5 | 8 | 26 | 0.361 |
| 7 | 8 | 42 | 0.583 |
| 6 | 8 | 32 | 0.444 |
| 2 | 8 | 32 | 0.444 |
| 2 | 8 | 32 | 0.444 |
| 2 | 8 | 32 | 0.444 |
| 1 | 8 | 42 | 0.583 |
| 2 | 8 | 32 | 0.444 |
| 2 | 8 | 32 | 0.444 |
| 5 | 8 | 26 | 0.361 |
| 8 | 8 | 56 | 0.778 |
| 5 | 8 | 26 | 0.361 |
| 0 | 8 | 56 | 0.778 |
| 0 | 8 | 56 | 0.778 |
| 0 | 8 | 56 | 0.778 |
| 5 | 8 | 26 | 0.361 |
| 5 | 8 | 26 | 0.361 |
| 5 | 8 | 26 | 0.361 |
| 5 | 8 | 26 | 0.361 |

| | | | |
|---|---|---|---|
| 7 | 8 | 42 | 0.583 |
| 7 | 8 | 42 | 0.583 |
| 5 | 8 | 26 | 0.361 |
| 5 | 8 | 26 | 0.361 |
| 6 | 8 | 32 | 0.444 |
| 2 | 8 | 32 | 0.444 |
| 2 | 8 | 32 | 0.444 |
| 2 | 8 | 32 | 0.444 |
| 3 | 8 | 26 | 0.361 |
| 3 | 8 | 26 | 0.361 |
| 3 | 8 | 26 | 0.361 |
| 6 | 8 | 32 | 0.444 |
| 8 | 8 | 56 | 0.778 |
| 8 | 8 | 56 | 0.778 |
| 7 | 8 | 42 | 0.583 |
| 7 | 8 | 42 | 0.583 |
| 7 | 8 | 42 | 0.583 |
| 0 | 8 | 56 | 0.778 |
| 0 | 8 | 56 | 0.778 |
| 0 | 8 | 56 | 0.778 |
| 1 | 8 | 42 | 0.583 |
| 2 | 8 | 32 | 0.444 |
| 1 | 8 | 42 | 0.583 |
| 2 | 8 | 32 | 0.444 |
| 2 | 8 | 32 | 0.444 |
| 2 | 8 | 32 | 0.444 |
| 0 | 8 | 56 | 0.778 |
| 0 | 8 | 56 | 0.778 |
| 0 | 8 | 56 | 0.778 |
| 5 | 8 | 26 | 0.361 |
| 5 | 8 | 26 | 0.361 |
| 5 | 8 | 26 | 0.361 |
| 8 | 8 | 56 | 0.778 |
| 8 | 8 | 56 | 0.778 |
| 8 | 8 | 56 | 0.778 |
| 2 | 8 | 32 | 0.444 |
| 2 | 8 | 32 | 0.444 |
| 4 | 8 | 24 | 0.333 |

| | | | | | |
|---|---|---|---|---|---|
| | 3 | 8 | 26 | 0.361 | |
| | 3 | 8 | 26 | 0.361 | |
| | 8 | 8 | 56 | 0.778 | |
| | 8 | 8 | 56 | 0.778 | |
| | 8 | 8 | 56 | 0.778 | |
| | 6 | 8 | 32 | 0.444 | |
| | 6 | 8 | 32 | 0.444 | |
| | 6 | 8 | 32 | 0.444 | |
| | 1 | 8 | 42 | 0.583 | |
| | 0 | 8 | 56 | 0.778 | |
| | 1 | 8 | 42 | 0.583 | |
| | 3 | 8 | 26 | 0.361 | |
| | 2 | 8 | 32 | 0.444 | |
| | 1 | 8 | 42 | 0.583 | |
| | 1 | 8 | 42 | 0.583 | |
| | 0 | 8 | 56 | 0.778 | |
| | 1 | 8 | 42 | 0.583 | |
| | 1 | 8 | 42 | 0.583 | |
| | 1 | 8 | 42 | 0.583 | |
| 654 | | | | 79.556 | sum |
| 7.267 | | | | **7.95555556** | sum / 10 |
| 52.80 | | 73.56 | | | |
