## Supplementary material for "Enhancing Wnt signaling lowers fracture incidence in a severe mouse model of Osteogenesis Imperfecta": SupTable2-tab3

| ID | Genotype | Sex | Age (Weeks) | MW-Pelvis Ratio | GM-Pelvis Ratio | IH-Pelvis Ratio | IB-Pelvis Ratio | IE-Pelvis Ratio |
|---|---|---|---|---|---|---|---|---|
| 989 | OI | F | 5 | 0.42 | 0.53 | 0.57 | 0.51 | 0.55 |
| 987 | OI | F | 5 | 0.57 | 0.56 | 0.56 | 0.53 | 0.65 |
| 941 | OI | F | 5 | 0.68 | 0.62 | 0.71 | 0.60 | 0.71 |
| 4115 | OI | F | 5 | 0.53 | 0.45 | 0.57 | 0.49 | 0.49 |
| 3520 | OI | F | 5 | 0.65 | 0.57 | 0.64 | 0.60 | 0.54 |
| 345 | OI | F | 5 | 0.49 | 0.58 | 0.65 | 0.67 | 0.54 |
| 198 | OI | F | 5 | 0.70 | 0.58 | 0.71 | 0.61 | 0.68 |
| 1910 | OI | F | 5 | 0.85 | 0.85 | 0.81 | 0.89 | 0.82 |
| 1163 | OI | F | 5 | 0.53 | 0.51 | 0.60 | 0.49 | 0.54 |
| 943 | OI | M | 5 | 0.57 | 0.62 | 0.69 | 0.67 | 0.62 |
| 873 | OI | M | 5 | 0.61 | 0.56 | 0.65 | 0.48 | 0.56 |
| 4690 | OI | M | 5 | 0.59 | 0.61 | 0.62 | 0.64 | 0.59 |
| 4596 | OI | M | 5 | 0.58 | 0.61 | 0.71 | 0.44 | 0.63 |
| 351 | OI | M | 5 | 0.56 | 0.64 | 0.68 | 0.57 | 0.61 |
| 349 | OI | M | 5 | 0.67 | 0.55 | 0.65 | 0.52 | 0.55 |
| 344 | OI | M | 5 | 0.43 | 0.57 | 0.64 | 0.59 | 0.58 |
| 319 | OI | M | 5 | 0.67 | 0.59 | 0.73 | 0.58 | 0.68 |
| 1214 | OI | M | 5 | 0.68 | 0.77 | 0.81 | 0.61 | 0.73 |
| 1211 | OI | M | 5 | 0.50 | 0.49 | 0.55 | 0.51 | 0.53 |
| 1210 | OI | M | 5 | 0.53 | 0.60 | 0.69 | 0.66 | 0.63 |
| 992 | OI/AV | F | 5 | 0.60 | 0.54 | 0.51 | 0.53 | 0.6 |
| 4686 | OI/AV | F | 5 | 0.55 | 0.66 | 0.66 | 0.63 | 0.58 |
| 4594 | OI/AV | F | 5 | 0.53 | 0.55 | 0.57 | 0.58 | 0.58 |
| 4592 | OI/AV | F | 5 | 0.44 | 0.60 | 0.70 | 0.55 | 0.53 |
| 4591 | OI/AV | F | 5 | 0.50 | 0.67 | 0.62 | 0.71 | 0.68 |
| 4113 | OI/AV | F | 5 | 0.57 | 0.54 | 0.57 | 0.59 | 0.59 |
| 340 | OI/AV | F | 5 | 0.62 | 0.68 | 0.66 | 0.68 | 0.66 |
| 339 | OI/AV | F | 5 | 0.60 | 0.58 | 0.75 | 0.58 | 0.61 |
| 318 | OI/AV | F | 5 | 0.68 | 0.54 | 0.68 | 0.59 | 0.59 |
| 197 | OI/AV | F | 5 | 0.67 | 0.52 | 0.59 | 0.51 | 0.67 |
| 196 | OI/AV | F | 5 | 0.54 | 0.59 | 0.54 | 0.59 | 0.55 |
| 1933 | OI/AV | F | 5 | 0.77 | 0.90 | 0.76 | 0.82 | 0.68 |
| 1930 | OI/AV | F | 5 | 0.51 | 0.53 | 0.63 | 0.53 | 0.56 |
| 1909 | OI/AV | F | 5 | 0.91 | 0.88 | 0.92 | 0.92 | 0.78 |
| 165 | OI/AV | F | 5 | 0.63 | 0.68 | 0.65 | 0.69 | 0.62 |
| 164 | OI/AV | F | 5 | 0.88 | 0.92 | 0.86 | 0.92 | 0.85 |
| 4598 | OI/AV | M | 5 | 0.63 | 0.67 | 0.62 | 0.70 | 0.64 |

| | | | | | | | | |
|---|---|---|---|---|---|---|---|---|
| 4117 | OI/AV | M | 5 | 0.67 | 0.65 | 0.70 | 0.73 | 0.56 |
| 3523 | OI/AV | M | 5 | 1.00 | 1.00 | 1.00 | 0.88 | 1 |
| 3522 | OI/AV | M | 5 | 0.85 | 0.92 | 0.93 | 0.88 | 0.88 |
| 3515 | OI/AV | M | 5 | 0.60 | 0.56 | 0.72 | 0.59 | 0.59 |
| 350 | OI/AV | M | 5 | 0.58 | 0.61 | 0.65 | 0.61 | 0.57 |
| 321 | OI/AV | M | 5 | 0.69 | 0.68 | 0.87 | 0.66 | 0.68 |
| 320 | OI/AV | M | 5 | 0.69 | 0.60 | 0.72 | 0.60 | 0.66 |
| 1935 | OI/AV | M | 5 | 0.47 | 0.53 | 0.56 | 0.53 | 0.59 |
| 1934 | OI/AV | M | 5 | 1.03 | 1.00 | 0.81 | 1.00 | 0.76 |
| 989 | OI | F | 9 | 0.49 | 0.48 | 0.53 | 0.52 | 0.63 |
| 987 | OI | F | 9 | 0.56 | 0.57 | 0.60 | 0.54 | 0.49 |
| 4115 | OI | F | 9 | 0.46 | 0.48 | 0.52 | 0.50 | 0.5 |
| 3520 | OI | F | 9 | 0.55 | 0.60 | 0.58 | 0.64 | 0.51 |
| 345 | OI | F | 9 | 0.53 | 0.55 | 0.63 | 0.63 | 0.53 |
| 198 | OI | F | 9 | 0.64 | 0.61 | 0.71 | 0.64 | 0.68 |
| 1910 | OI | F | 9 | 0.90 | 0.90 | 0.79 | 0.84 | 0.84 |
| 1163 | OI | F | 9 | 0.53 | 0.53 | 0.64 | 0.53 | 0.55 |
| 873 | OI | M | 9 | 0.55 | 0.53 | 0.63 | 0.48 | 0.5 |
| 4690 | OI | M | 9 | 0.46 | 0.57 | 0.59 | 0.60 | 0.46 |
| 4596 | OI | M | 9 | 0.54 | 0.64 | 0.64 | 0.61 | 0.63 |
| 351 | OI | M | 9 | 0.63 | 0.63 | 0.60 | 0.50 | 0.61 |
| 349 | OI | M | 9 | 0.51 | 0.54 | 0.57 | 0.53 | 0.63 |
| 344 | OI | M | 9 | 0.44 | 0.54 | 0.57 | 0.49 | 0.53 |
| 319 | OI | M | 9 | 0.59 | 0.61 | 0.69 | 0.59 | 0.61 |
| 1214 | OI | M | 9 | 0.58 | 0.73 | 0.82 | 0.69 | 0.67 |
| 1211 | OI | M | 9 | 0.47 | 0.50 | 0.53 | 0.47 | 0.52 |
| 1210 | OI | M | 9 | 0.42 | 0.53 | 0.64 | 0.51 | 0.52 |
| 992 | OI/AV | F | 9 | 0.59 | 0.61 | 0.63 | 0.59 | 0.58 |
| 4686 | OI/AV | F | 9 | 0.62 | 0.62 | 0.67 | 0.64 | 0.63 |
| 4594 | OI/AV | F | 9 | 0.49 | 0.60 | 0.58 | 0.70 | 0.57 |
| 4592 | OI/AV | F | 9 | 0.58 | 0.61 | 0.63 | 0.61 | 0.64 |
| 4591 | OI/AV | F | 9 | 0.58 | 0.62 | 0.62 | 0.63 | 0.63 |
| 4113 | OI/AV | F | 9 | 0.55 | 0.57 | 0.61 | 0.58 | 0.55 |
| 340 | OI/AV | F | 9 | 0.71 | 0.68 | 0.70 | 0.84 | 0.57 |
| 339 | OI/AV | F | 9 | 0.62 | 0.60 | 0.67 | 0.73 | 0.58 |
| 318 | OI/AV | F | 9 | 0.63 | 0.63 | 0.68 | 0.63 | 0.63 |
| 197 | OI/AV | F | 9 | 0.55 | 0.53 | 0.60 | 0.52 | 0.52 |
| 196 | OI/AV | F | 9 | 0.59 | 0.62 | 0.63 | 0.63 | 0.61 |

| | | | | | | | | |
|---|---|---|---|---|---|---|---|---|
| 1933 | OI/AV | F | 9 | 0.86 | 0.85 | 0.73 | 0.81 | 0.83 |
| 1930 | OI/AV | F | 9 | 0.60 | 0.57 | 0.56 | 0.56 | 0.61 |
| 1909 | OI/AV | F | 9 | 0.89 | 0.93 | 0.94 | 0.90 | 0.84 |
| 165 | OI/AV | F | 9 | 0.73 | 0.73 | 0.64 | 0.66 | 0.73 |
| 164 | OI/AV | F | 9 | 0.80 | 0.84 | 0.81 | 0.81 | 0.83 |
| 4598 | OI/AV | M | 9 | 0.65 | 0.63 | 0.66 | 0.64 | 0.64 |
| 4117 | OI/AV | M | 9 | 0.70 | 0.60 | 0.58 | 0.62 | 0.56 |
| 3523 | OI/AV | M | 9 | 0.89 | 0.93 | 0.93 | 0.90 | 0.89 |
| 3522 | OI/AV | M | 9 | 0.83 | 0.93 | 0.83 | 0.90 | 0.86 |
| 3515 | OI/AV | M | 9 | 0.60 | 0.57 | 0.64 | 0.57 | 0.54 |
| 350 | OI/AV | M | 9 | 0.50 | 0.60 | 0.68 | 0.60 | 0.55 |
| 321 | OI/AV | M | 9 | 0.67 | 0.69 | 0.80 | 0.62 | 0.63 |
| 320 | OI/AV | M | 9 | 0.65 | 0.60 | 0.70 | 0.57 | 0.64 |
| 1935 | OI/AV | M | 9 | 0.47 | 0.50 | 0.57 | 0.44 | 0.58 |
| 1934 | OI/AV | M | 9 | 0.97 | 1.10 | 0.90 | 1.00 | 0.77 |
| 989 | OI | F | 13 | 0.48 | 0.46 | 0.55 | 0.52 | 0.5 |
| 987 | OI | F | 13 | 0.47 | 0.53 | 0.51 | 0.50 | 0.48 |
| 4115 | OI | F | 13 | 0.44 | 0.43 | 0.52 | 0.52 | 0.41 |
| 345 | OI | F | 13 | 0.58 | 0.56 | 0.56 | 0.65 | 0.59 |
| 198 | OI | F | 13 | 0.65 | 0.62 | 0.73 | 0.60 | 0.68 |
| 1910 | OI | F | 13 | 0.87 | 0.84 | 0.79 | 0.79 | 0.8 |
| 1163 | OI | F | 13 | 0.48 | 0.48 | 0.56 | 0.50 | 0.57 |
| 873 | OI | M | 13 | 0.52 | 0.49 | 0.63 | 0.44 | 0.47 |
| 4690 | OI | M | 13 | 0.50 | 0.52 | 0.64 | 0.62 | 0.5 |
| 4596 | OI | M | 13 | 0.68 | 0.63 | 0.73 | 0.70 | 0.62 |
| 351 | OI | M | 13 | 0.57 | 0.60 | 0.65 | 0.60 | 0.6 |
| 349 | OI | M | 13 | 0.55 | 0.56 | 0.62 | 0.53 | 0.57 |
| 344 | OI | M | 13 | 0.43 | 0.43 | 0.56 | 0.53 | 0.51 |
| 319 | OI | M | 13 | 0.68 | 0.61 | 0.73 | 0.56 | 0.66 |
| 1214 | OI | M | 13 | 0.58 | 0.65 | 0.75 | 0.55 | 0.66 |
| 1211 | OI | M | 13 | 0.46 | 0.46 | 0.62 | 0.49 | 0.5 |
| 1210 | OI | M | 13 | 0.46 | 0.45 | 0.54 | 0.47 | 0.5 |
| 992 | OI/AV | F | 13 | 0.59 | 0.54 | 0.63 | 0.51 | 0.46 |
| 4686 | OI/AV | F | 13 | 0.62 | 0.62 | 0.71 | 0.68 | 0.63 |
| 4594 | OI/AV | F | 13 | 0.54 | 0.57 | 0.56 | 0.58 | 0.57 |
| 4592 | OI/AV | F | 13 | 0.54 | 0.56 | 0.64 | 0.60 | 0.6 |
| 4591 | OI/AV | F | 13 | 0.51 | 0.60 | 0.67 | 0.61 | 0.61 |
| 4113 | OI/AV | F | 13 | 0.56 | 0.50 | 0.62 | 0.58 | 0.56 |

| | | | | | | | | |
|---|---|---|---|---|---|---|---|---|
| 340 | OI/AV | F | 13 | 0.65 | 0.61 | 0.74 | 0.63 | 0.61 |
| 339 | OI/AV | F | 13 | 0.68 | 0.61 | 0.77 | 0.63 | 0.62 |
| 318 | OI/AV | F | 13 | 0.63 | 0.59 | 0.74 | 0.65 | 0.65 |
| 197 | OI/AV | F | 13 | 0.61 | 0.56 | 0.59 | 0.58 | 0.64 |
| 196 | OI/AV | F | 13 | 0.50 | 0.58 | 0.59 | 0.53 | 0.58 |
| 1933 | OI/AV | F | 13 | 0.84 | 0.82 | 0.69 | 0.94 | 0.81 |
| 1930 | OI/AV | F | 13 | 0.55 | 0.60 | 0.61 | 0.55 | 0.5 |
| 1909 | OI/AV | F | 13 | 0.94 | 0.93 | 0.94 | 0.88 | 0.81 |
| 165 | OI/AV | F | 13 | 0.66 | 0.57 | 0.68 | 0.59 | 0.68 |
| 164 | OI/AV | F | 13 | 0.78 | 0.84 | 0.79 | 0.78 | 0.78 |
| 4598 | OI/AV | M | 13 | 0.60 | 0.57 | 0.67 | 0.63 | 0.65 |
| 4117 | OI/AV | M | 13 | 0.62 | 0.59 | 0.68 | 0.49 | 0.56 |
| 3523 | OI/AV | M | 13 | 0.88 | 0.81 | 0.89 | 0.90 | 0.88 |
| 3522 | OI/AV | M | 13 | 0.86 | 0.88 | 0.90 | 0.90 | 0.86 |
| 3515 | OI/AV | M | 13 | 0.61 | 0.58 | 0.63 | 0.63 | 0.62 |
| 350 | OI/AV | M | 13 | 0.60 | 0.55 | 0.66 | 0.59 | 0.49 |
| 321 | OI/AV | M | 13 | 0.67 | 0.66 | 0.81 | 0.69 | 0.67 |
| 320 | OI/AV | M | 13 | 0.58 | 0.59 | 0.64 | 0.58 | 0.62 |
| 1935 | OI/AV | M | 13 | 0.45 | 0.50 | 0.58 | 0.51 | 0.58 |
| 1934 | OI/AV | M | 13 | 0.96 | 0.93 | 0.93 | 1.00 | 0.68 |

| KC-Pelvis Ratio | DN-Pelvis Ratio | SV-Pelvis Ratio | Anova: Two-Fa SUMMARY |
|---|---|---|---|
| 0.58 | 0.57 | 0.57 | 989-5 |
| 0.53 | 0.60 | 0.63 | 987-5 |
| 0.63 | 0.56 | 0.56 | 941-5 |
| 0.51 | 0.52 | 0.52 | 4115-5 |
| 0.5 | 0.50 | 0.57 | 3520-5 |
| 0.56 | 0.58 | 0.58 | 345-5 |
| 0.63 | 0.62 | 0.62 | 198-5 |
| 0.82 | 0.81 | 0.93 | 1910-5 |
| 0.49 | 0.50 | 0.5 | 1163-5 |
| 0.62 | 0.58 | 0.58 | 943-5 |
| 0.56 | 0.51 | 0.51 | 873-5 |
| 0.55 | 0.61 | 0.61 | 4690-5 |
| 0.58 | 0.61 | 0.73 | 4596-5 |
| 0.54 | 0.59 | 0.58 | 351-5 |
| 0.54 | 0.54 | 0.54 | 349-5 |
| 0.49 | 0.56 | 0.56 | 344-5 |
| 0.6 | 0.61 | 0.61 | 319-5 |
| 0.73 | 0.69 | 0.66 | 1214-5 |
| 0.49 | 0.51 | 0.51 | 1211-5 |
| 0.6 | 0.49 | 0.49 | 1210-5 |
| 0.65 | 0.53 | 0.53 | 992-5 |
| 0.63 | 0.51 | 0.59 | 4686-5 |
| 0.54 | 0.52 | 0.45 | 4594-5 |
| 0.61 | 0.61 | 0.53 | 4592-5 |
| 0.54 | 0.58 | 0.53 | 4591-5 |
| 0.52 | 0.55 | 0.55 | 4113-5 |
| 0.71 | 0.55 | 0.55 | 340-5 |
| 0.64 | 0.56 | 0.67 | 339-5 |
| 0.6 | 0.67 | 0.61 | 318-5 |
| 0.53 | 0.56 | 0.67 | 197-5 |
| 0.59 | 0.61 | 0.74 | 196-5 |
| 0.76 | 0.78 | 0.78 | 1933-5 |
| 0.54 | 0.50 | 0.5 | 1930-5 |
| 0.95 | 0.88 | 0.8 | 1909-5 |
| 0.73 | 0.66 | 0.7 | 165-5 |
| 0.94 | 0.91 | 0.92 | 164-5 |
| 0.64 | 0.51 | 0.51 | 4598-5 |

| | | | |
|---|---|---|---|
| 0.65 | 0.65 | 0.65 | 4117-5 |
| 0.9 | 0.92 | 0.88 | 3523-5 |
| 0.86 | 0.88 | 0.88 | 3522-5 |
| 0.52 | 0.66 | 0.66 | 3515-5 |
| 0.59 | 0.60 | 0.6 | 350-5 |
| 0.67 | 0.67 | 0.67 | 321-5 |
| 0.6 | 0.76 | 0.76 | 320-5 |
| 0.5 | 0.50 | 0.45 | 1935-5 |
| 1 | 1.00 | 1 | 1934-5 |
| 0.48 | 0.45 | 0.45 | 989-9 |
| 0.5 | 0.57 | 0.57 | 987-9 |
| 0.53 | 0.40 | 0.4 | 4115-9 |
| 0.53 | 0.67 | 0.45 | 3520-9 |
| 0.58 | 0.61 | 0.61 | 345-9 |
| 0.62 | 0.58 | 0.63 | 198-9 |
| 0.84 | 0.85 | 0.85 | 1910-9 |
| 0.56 | 0.50 | 0.5 | 1163-9 |
| 0.49 | 0.51 | 0.51 | 873-9 |
| 0.58 | 0.58 | 0.58 | 4690-9 |
| 0.61 | 0.61 | 0.61 | 4596-9 |
| 0.58 | 0.56 | 0.56 | 351-9 |
| 0.52 | 0.55 | 0.55 | 349-9 |
| 0.5 | 0.49 | 0.49 | 344-9 |
| 0.59 | 0.59 | 0.59 | 319-9 |
| 0.7 | 0.66 | 0.69 | 1214-9 |
| 0.48 | 0.45 | 0.49 | 1211-9 |
| 0.49 | 0.49 | 0.49 | 1210-9 |
| 0.58 | 0.55 | 0.55 | 992-9 |
| 0.63 | 0.52 | 0.55 | 4686-9 |
| 0.55 | 0.57 | 0.57 | 4594-9 |
| 0.57 | 0.56 | 0.56 | 4592-9 |
| 0.55 | 0.60 | 0.6 | 4591-9 |
| 0.45 | 0.53 | 0.53 | 4113-9 |
| 0.71 | 0.59 | 0.59 | 340-9 |
| 0.6 | 0.53 | 0.58 | 339-9 |
| 0.6 | 0.64 | 0.64 | 318-9 |
| 0.53 | 0.52 | 0.52 | 197-9 |
| 0.57 | 0.59 | 0.59 | 196-9 |

| | | | |
|---|---|---|---|
| 0.78 | 0.84 | 0.84 | 1933-9 |
| 0.56 | 0.58 | 0.58 | 1930-9 |
| 0.91 | 0.87 | 0.85 | 1909-9 |
| 0.75 | 0.74 | 0.71 | 165-9 |
| 0.78 | 0.80 | 0.86 | 164-9 |
| 0.64 | 0.52 | 0.52 | 4598-9 |
| 0.6 | 0.53 | 0.53 | 4117-9 |
| 0.86 | 0.90 | 0.9 | 3523-9 |
| 0.79 | 0.81 | 0.81 | 3522-9 |
| 0.55 | 0.62 | 0.62 | 3515-9 |
| 0.59 | 0.55 | 0.55 | 350-9 |
| 0.68 | 0.69 | 0.69 | 321-9 |
| 0.61 | 0.70 | 0.7 | 320-9 |
| 0.45 | 0.49 | 0.5 | 1935-9 |
| 1 | 0.90 | 0.9 | 1934-9 |
| 0.44 | 0.46 | 0.46 | 989-13 |
| 0.46 | 0.53 | 0.53 | 987-13 |
| 0.39 | 0.40 | 0.4 | 4115-13 |
| 0.57 | 0.56 | 0.56 | 345-13 |
| 0.61 | 0.60 | 0.43 | 198-13 |
| 0.81 | 0.84 | 0.84 | 1910-13 |
| 0.56 | 0.48 | 0.48 | 1163-13 |
| 0.47 | 0.49 | 0.49 | 873-13 |
| 0.59 | 0.53 | 0.52 | 4690-13 |
| 0.62 | 0.63 | 0.63 | 4596-13 |
| 0.6 | 0.55 | 0.55 | 351-13 |
| 0.5 | 0.53 | 0.53 | 349-13 |
| 0.48 | 0.43 | 0.43 | 344-13 |
| 0.53 | 0.71 | 0.71 | 319-13 |
| 0.71 | 0.66 | 0.65 | 1214-13 |
| 0.47 | 0.45 | 0.45 | 1211-13 |
| 0.52 | 0.49 | 0.51 | 1210-13 |
| 0.55 | 0.54 | 0.54 | 992-13 |
| 0.59 | 0.57 | 0.62 | 4686-13 |
| 0.53 | 0.61 | 0.57 | 4594-13 |
| 0.57 | 0.54 | 0.54 | 4592-13 |
| 0.55 | 0.59 | 0.6 | 4591-13 |
| 0.54 | 0.50 | 0.5 | 4113-13 |

| | | | |
|---|---|---|---|
| 0.68 | 0.59 | 0.59 | 340-13 |
| 0.6 | 0.54 | 0.62 | 339-13 |
| 0.6 | 0.59 | 0.59 | 318-13 |
| 0.53 | 0.53 | 0.53 | 197-13 |
| 0.53 | 0.59 | 0.59 | 196-13 |
| 0.78 | 0.82 | 0.82 | 1933-13 |
| 0.5 | 0.59 | 0.6 | 1930-13 |
| 0.83 | 0.81 | 0.77 | 1909-13 |
| 0.53 | 0.75 | 0.7 | 165-13 |
| 0.78 | 0.84 | 0.84 | 164-13 |
| 0.65 | 0.57 | 0.57 | 4598-13 |
| 0.5 | 0.54 | 0.54 | 4117-13 |
| 0.86 | 0.84 | 0.77 | 3523-13 |
| 0.88 | 0.76 | 0.88 | 3522-13 |
| 0.53 | 0.64 | 0.58 | 3515-13 |
| 0.57 | 0.55 | 0.55 | 350-13 |
| 0.62 | 0.66 | 0.66 | 321-13 |
| 0.57 | 0.64 | 0.64 | 320-13 |
| 0.5 | 0.54 | 0.52 | 1935-13 |
| 0.91 | 0.87 | 1 | 1934-13 |

MW-Pelvis Ra
GM-Pelvis Rat
IH-Pelvis Ratio
IB-Pelvis Ratio
IE-Pelvis Ratio
KC-Pelvis Rati
DN-Pelvis Rat
SV-Pelvis Rati

ANOVA

*Source of Variati*

Rows
Columns
Error

Total

| Count | Sum | Average | Variance |
|---|---|---|---|
| 8 | 4.29860465 | 0.54 | 0.00286852 |
| 8 | 4.62666667 | 0.58 | 0.00196508 |
| 8 | 5.06647059 | 0.63 | 0.00366742 |
| 8 | 4.08488372 | 0.51 | 0.00125946 |
| 8 | 4.57 | 0.57 | 0.00326964 |
| 8 | 4.64571429 | 0.58 | 0.00341224 |
| 8 | 5.1469697 | 0.64 | 0.00215066 |
| 8 | 6.77615385 | 0.85 | 0.0018491 |
| 8 | 4.155 | 0.52 | 0.00136027 |
| 8 | 4.95 | 0.62 | 0.00186964 |
| 8 | 4.43714286 | 0.55 | 0.00307041 |
| 8 | 4.81536585 | 0.60 | 0.00072638 |
| 8 | 4.89064516 | 0.61 | 0.00792108 |
| 8 | 4.7725 | 0.60 | 0.00205882 |
| 8 | 4.55666667 | 0.57 | 0.00310615 |
| 8 | 4.41857143 | 0.55 | 0.00421454 |
| 8 | 5.06666667 | 0.63 | 0.00279365 |
| 8 | 5.67741935 | 0.71 | 0.00405152 |
| 8 | 4.09 | 0.51 | 0.0004125 |
| 8 | 4.68941176 | 0.59 | 0.00575057 |
| 8 | 4.49 | 0.56 | 0.0024125 |
| 8 | 4.81263158 | 0.60 | 0.00283197 |
| 8 | 4.32333333 | 0.54 | 0.00182044 |
| 8 | 4.57444444 | 0.57 | 0.00576259 |
| 8 | 4.83 | 0.60 | 0.00608393 |
| 8 | 4.48142857 | 0.56 | 0.00060434 |
| 8 | 5.11068966 | 0.64 | 0.00363744 |
| 8 | 4.99 | 0.62 | 0.00385536 |
| 8 | 4.95741935 | 0.62 | 0.00258516 |
| 8 | 4.71666667 | 0.59 | 0.0049252 |
| 8 | 4.75054054 | 0.59 | 0.00418995 |
| 8 | 6.25419355 | 0.78 | 0.00382979 |
| 8 | 4.30020408 | 0.54 | 0.00181983 |
| 8 | 7.04304348 | 0.88 | 0.00365582 |
| 8 | 5.35857143 | 0.67 | 0.00138801 |
| 8 | 7.20461538 | 0.90 | 0.00103343 |
| 8 | 4.91857143 | 0.61 | 0.0048227 |

| | | | |
|---|---|---|---|
| 8 | 5.25666667 | 0.66 | 0.00241091 |
| 8 | 7.58 | 0.95 | 0.00330714 |
| 8 | 7.07615385 | 0.88 | 0.00078317 |
| 8 | 4.9 | 0.61 | 0.00407857 |
| 8 | 4.80692308 | 0.60 | 0.00060379 |
| 8 | 5.58965517 | 0.70 | 0.00487052 |
| 8 | 5.3875 | 0.67 | 0.00481596 |
| 8 | 4.13169811 | 0.52 | 0.00209042 |
| 8 | 7.60448276 | 0.95 | 0.01076212 |
| 8 | 4.03491525 | 0.50 | 0.0034105 |
| 8 | 4.4025 | 0.55 | 0.0014365 |
| 8 | 3.79 | 0.47 | 0.00254107 |
| 8 | 4.53263158 | 0.57 | 0.00510115 |
| 8 | 4.665 | 0.58 | 0.00187813 |
| 8 | 5.11102564 | 0.64 | 0.00164157 |
| 8 | 6.81322581 | 0.85 | 0.00131587 |
| 8 | 4.34 | 0.54 | 0.00199286 |
| 8 | 4.20465587 | 0.53 | 0.00232168 |
| 8 | 4.42341463 | 0.55 | 0.00324693 |
| 8 | 4.89285714 | 0.61 | 0.00095536 |
| 8 | 4.67 | 0.58 | 0.0019125 |
| 8 | 4.40428571 | 0.55 | 0.00135332 |
| 8 | 4.05117647 | 0.51 | 0.00154755 |
| 8 | 4.85974359 | 0.61 | 0.00119415 |
| 8 | 5.54 | 0.69 | 0.00456429 |
| 8 | 3.91 | 0.49 | 0.00072679 |
| 8 | 4.09 | 0.51 | 0.0038125 |
| 8 | 4.67695652 | 0.58 | 0.00073967 |
| 8 | 4.87904762 | 0.61 | 0.00245454 |
| 8 | 4.62648649 | 0.58 | 0.00353171 |
| 8 | 4.75777778 | 0.59 | 0.00101014 |
| 8 | 4.825 | 0.60 | 0.0008067 |
| 8 | 4.36761905 | 0.55 | 0.00222494 |
| 8 | 5.39052632 | 0.67 | 0.00797513 |
| 8 | 4.91162162 | 0.61 | 0.00377287 |
| 8 | 5.08157895 | 0.64 | 0.00048377 |
| 8 | 4.28761905 | 0.54 | 0.000761 |
| 8 | 4.82974359 | 0.60 | 0.00048494 |

| | | | |
|---|---|---|---|
| 8 | 6.54206897 | 0.82 | 0.00190423 |
| 8 | 4.6152381 | 0.58 | 0.00033651 |
| 8 | 7.13285714 | 0.89 | 0.00129821 |
| 8 | 5.68972973 | 0.71 | 0.00158249 |
| 8 | 6.53 | 0.82 | 0.00065536 |
| 8 | 4.9 | 0.61 | 0.00333571 |
| 8 | 4.7169697 | 0.59 | 0.00296305 |
| 8 | 7.20285714 | 0.90 | 0.00050714 |
| 8 | 6.75758621 | 0.84 | 0.00235393 |
| 8 | 4.71 | 0.59 | 0.00132679 |
| 8 | 4.62 | 0.58 | 0.00285 |
| 8 | 5.46666667 | 0.68 | 0.00298413 |
| 8 | 5.17 | 0.65 | 0.00256964 |
| 8 | 4.00169811 | 0.50 | 0.00261438 |
| 8 | 7.53666667 | 0.94 | 0.00959663 |
| 8 | 3.86766323 | 0.48 | 0.00137283 |
| 8 | 4.00850394 | 0.50 | 0.00082614 |
| 8 | 3.5075 | 0.44 | 0.00278382 |
| 8 | 4.62777778 | 0.58 | 0.00095518 |
| 8 | 4.92 | 0.62 | 0.00762857 |
| 8 | 6.58096774 | 0.82 | 0.00086325 |
| 8 | 4.11 | 0.51 | 0.00174107 |
| 8 | 3.99550388 | 0.50 | 0.00326255 |
| 8 | 4.42 | 0.55 | 0.00310714 |
| 8 | 5.23647059 | 0.65 | 0.00177635 |
| 8 | 4.72272727 | 0.59 | 0.00107106 |
| 8 | 4.38545455 | 0.55 | 0.00129917 |
| 8 | 3.79857143 | 0.47 | 0.00281862 |
| 8 | 5.19421053 | 0.65 | 0.0055666 |
| 8 | 5.21 | 0.65 | 0.00409821 |
| 8 | 3.9 | 0.49 | 0.00319286 |
| 8 | 3.94 | 0.49 | 0.00096429 |
| 8 | 4.36482759 | 0.55 | 0.00260784 |
| 8 | 5.03538462 | 0.63 | 0.00207299 |
| 8 | 4.53054054 | 0.57 | 0.0005942 |
| 8 | 4.59347826 | 0.57 | 0.00130904 |
| 8 | 4.74162791 | 0.59 | 0.00215482 |
| 8 | 4.35813953 | 0.54 | 0.00190674 |

| | | | |
|---|---|---|---|
| 8 | 5.09864865 | 0.64 | 0.0026454 |
| 8 | 5.06567568 | 0.63 | 0.00445769 |
| 8 | 5.04157895 | 0.63 | 0.00265745 |
| 8 | 4.57363636 | 0.57 | 0.00174013 |
| 8 | 4.49 | 0.56 | 0.00126964 |
| 8 | 6.51870968 | 0.81 | 0.00473385 |
| 8 | 4.49545455 | 0.56 | 0.00201167 |
| 8 | 6.90548387 | 0.86 | 0.00447381 |
| 8 | 5.15853659 | 0.64 | 0.00550828 |
| 8 | 6.43125 | 0.80 | 0.00090421 |
| 8 | 4.91465116 | 0.61 | 0.00169693 |
| 8 | 4.51764706 | 0.56 | 0.00396372 |
| 8 | 6.83461538 | 0.85 | 0.00203549 |
| 8 | 6.92206897 | 0.87 | 0.00202615 |
| 8 | 4.8197561 | 0.60 | 0.00136377 |
| 8 | 4.56 | 0.57 | 0.00242857 |
| 8 | 5.43666667 | 0.68 | 0.00315377 |
| 8 | 4.85894737 | 0.61 | 0.00094412 |
| 8 | 4.18454545 | 0.52 | 0.00181557 |
| 8 | 7.28428571 | 0.91 | 0.01063495 |
| 133 | 82.587541 | 0.62095895 | 0.01874678 |
| 133 | 83.84 | 0.63037594 | 0.01830668 |
| 133 | 89.81 | 0.67526316 | 0.01145088 |
| 133 | 84.08 | 0.63218045 | 0.01738233 |
| 133 | 83.05 | 0.62443609 | 0.0114582 |
| 133 | 81.64 | 0.61383459 | 0.01606776 |
| 133 | 81.5 | 0.61278195 | 0.01554751 |
| 133 | 81.8 | 0.61503759 | 0.01691913 |

| *SS* | *df* | *MS* | *F* |
|---|---|---|---|
| 14.4614253 | 132 | 0.10955625 | 46.9823871 |
| 0.38827789 | 7 | 0.05546827 | 23.7871563 |
| 2.15463673 | 924 | 0.00233186 | |
| 17.0043399 | 1063 | | |

0.8307111
