## Supplementary material for "Enhancing Wnt signaling lowers fracture incidence in a severe mouse model of Osteogenesis Imperfecta": SupTable3

| ID | Genotype | Sex | Age (Weeks) | Interval | Date | Session Info 0 |
|---|---|---|---|---|---|---|
| 164 | OI/AV | F | 5 | 1 | 20220321 | |
| 164 | OI/AV | F | 5 | 2 | 20220321 | |
| 164 | OI/AV | F | 5 | 3 | 20220321 | |
| 164 | OI/AV | F | 5 | 4 | 20220321 | |
| 164 | OI/AV | F | 5 | 5 | 20220321 | |
| 164 | OI/AV | F | 5 | 6 | 20220321 | |
| 164 | OI/AV | F | 5 | 7 | 20220321 | |
| 164 | OI/AV | F | 5 | 8 | 20220321 | |
| 164 | OI/AV | F | 5 | 9 | 20220321 | |
| 164 | OI/AV | F | 5 | 10 | 20220321 | |
| 164 | OI/AV | F | 5 | 11 | 20220321 | |
| 164 | OI/AV | F | 5 | 12 | 20220321 | |
| 164 | OI/AV | F | 5 | | | 60 |
| 164 | OI/AV | F | 9 | 1 | 20220420 | |
| 164 | OI/AV | F | 9 | 2 | 20220420 | |
| 164 | OI/AV | F | 9 | 3 | 20220420 | |
| 164 | OI/AV | F | 9 | 4 | 20220420 | |
| 164 | OI/AV | F | 9 | 5 | 20220420 | |
| 164 | OI/AV | F | 9 | 6 | 20220420 | |
| 164 | OI/AV | F | 9 | 7 | 20220420 | |
| 164 | OI/AV | F | 9 | 8 | 20220420 | |
| 164 | OI/AV | F | 9 | 9 | 20220420 | |
| 164 | OI/AV | F | 9 | 10 | 20220420 | |
| 164 | OI/AV | F | 9 | 11 | 20220420 | |
| 164 | OI/AV | F | 9 | 12 | 20220420 | |
| 164 | OI/AV | F | 9 | | | 60 |
| 164 | OI/AV | F | 13 | 1 | 20220516 | |
| 164 | OI/AV | F | 13 | 2 | 20220516 | |
| 164 | OI/AV | F | 13 | 3 | 20220516 | |
| 164 | OI/AV | F | 13 | 4 | 20220516 | |
| 164 | OI/AV | F | 13 | 5 | 20220516 | |
| 164 | OI/AV | F | 13 | 6 | 20220516 | |
| 164 | OI/AV | F | 13 | 7 | 20220516 | |
| 164 | OI/AV | F | 13 | 8 | 20220516 | |
| 164 | OI/AV | F | 13 | 9 | 20220516 | |
| 164 | OI/AV | F | 13 | 10 | 20220516 | |
| 164 | OI/AV | F | 13 | 11 | 20220516 | |
| 164 | OI/AV | F | 13 | 12 | 20220516 | |
| 164 | OI/AV | F | 13 | | | 60 |
| 165 | OI/AV | F | 5 | 1 | 20220321 | |

| | | | | | | |
|---|---|---|---|---|---|---|
| 165 | OI/AV | F | 5 | 2 | 20220321 | |
| 165 | OI/AV | F | 5 | 3 | 20220321 | |
| 165 | OI/AV | F | 5 | 4 | 20220321 | |
| 165 | OI/AV | F | 5 | 5 | 20220321 | |
| 165 | OI/AV | F | 5 | 6 | 20220321 | |
| 165 | OI/AV | F | 5 | 7 | 20220321 | |
| 165 | OI/AV | F | 5 | 8 | 20220321 | |
| 165 | OI/AV | F | 5 | 9 | 20220321 | |
| 165 | OI/AV | F | 5 | 10 | 20220321 | |
| 165 | OI/AV | F | 5 | 11 | 20220321 | |
| 165 | OI/AV | F | 5 | 12 | 20220321 | |
| 165 | OI/AV | F | 5 | | | 60 |
| 165 | OI/AV | F | 9 | 1 | 20220420 | |
| 165 | OI/AV | F | 9 | 2 | 20220420 | |
| 165 | OI/AV | F | 9 | 3 | 20220420 | |
| 165 | OI/AV | F | 9 | 4 | 20220420 | |
| 165 | OI/AV | F | 9 | 5 | 20220420 | |
| 165 | OI/AV | F | 9 | 6 | 20220420 | |
| 165 | OI/AV | F | 9 | 7 | 20220420 | |
| 165 | OI/AV | F | 9 | 8 | 20220420 | |
| 165 | OI/AV | F | 9 | 9 | 20220420 | |
| 165 | OI/AV | F | 9 | 10 | 20220420 | |
| 165 | OI/AV | F | 9 | 11 | 20220420 | |
| 165 | OI/AV | F | 9 | 12 | 20220420 | |
| 165 | OI/AV | F | 9 | | | 60 |
| 165 | OI/AV | F | 13 | 1 | 20220516 | |
| 165 | OI/AV | F | 13 | 2 | 20220516 | |
| 165 | OI/AV | F | 13 | 3 | 20220516 | |
| 165 | OI/AV | F | 13 | 4 | 20220516 | |
| 165 | OI/AV | F | 13 | 5 | 20220516 | |
| 165 | OI/AV | F | 13 | 6 | 20220516 | |
| 165 | OI/AV | F | 13 | 7 | 20220516 | |
| 165 | OI/AV | F | 13 | 8 | 20220516 | |
| 165 | OI/AV | F | 13 | 9 | 20220516 | |
| 165 | OI/AV | F | 13 | 10 | 20220516 | |
| 165 | OI/AV | F | 13 | 11 | 20220516 | |
| 165 | OI/AV | F | 13 | 12 | 20220516 | |
| 165 | OI/AV | F | 13 | | | 60 |
| 166 | WT | M | 5 | 1 | 20220321 | |
| 166 | WT | M | 5 | 2 | 20220321 | |
| 166 | WT | M | 5 | 3 | 20220321 | |

| | | | | | | |
|---|---|---|---|---|---|---|
| 166 | WT | M | 5 | 4 | 20220321 | |
| 166 | WT | M | 5 | 5 | 20220321 | |
| 166 | WT | M | 5 | 6 | 20220321 | |
| 166 | WT | M | 5 | 7 | 20220321 | |
| 166 | WT | M | 5 | 8 | 20220321 | |
| 166 | WT | M | 5 | 9 | 20220321 | |
| 166 | WT | M | 5 | 10 | 20220321 | |
| 166 | WT | M | 5 | 11 | 20220321 | |
| 166 | WT | M | 5 | 12 | 20220321 | |
| 166 | WT | M | 5 | | | 60 |
| 166 | WT | M | 9 | 1 | 20220420 | |
| 166 | WT | M | 9 | 2 | 20220420 | |
| 166 | WT | M | 9 | 3 | 20220420 | |
| 166 | WT | M | 9 | 4 | 20220420 | |
| 166 | WT | M | 9 | 5 | 20220420 | |
| 166 | WT | M | 9 | 6 | 20220420 | |
| 166 | WT | M | 9 | 7 | 20220420 | |
| 166 | WT | M | 9 | 8 | 20220420 | |
| 166 | WT | M | 9 | 9 | 20220420 | |
| 166 | WT | M | 9 | 10 | 20220420 | |
| 166 | WT | M | 9 | 11 | 20220420 | |
| 166 | WT | M | 9 | 12 | 20220420 | |
| 166 | WT | M | 9 | | | 60 |
| 167 | WT/AV | F | 5 | 1 | 20220321 | |
| 167 | WT/AV | F | 5 | 2 | 20220321 | |
| 167 | WT/AV | F | 5 | 3 | 20220321 | |
| 167 | WT/AV | F | 5 | 4 | 20220321 | |
| 167 | WT/AV | F | 5 | 5 | 20220321 | |
| 167 | WT/AV | F | 5 | 6 | 20220321 | |
| 167 | WT/AV | F | 5 | 7 | 20220321 | |
| 167 | WT/AV | F | 5 | 8 | 20220321 | |
| 167 | WT/AV | F | 5 | 9 | 20220321 | |
| 167 | WT/AV | F | 5 | 10 | 20220321 | |
| 167 | WT/AV | F | 5 | 11 | 20220321 | |
| 167 | WT/AV | F | 5 | 12 | 20220321 | |
| 167 | WT/AV | F | 5 | | | 60 |
| 167 | WT/AV | F | 9 | 1 | 20220420 | |
| 167 | WT/AV | F | 9 | 2 | 20220420 | |
| 167 | WT/AV | F | 9 | 3 | 20220420 | |
| 167 | WT/AV | F | 9 | 4 | 20220420 | |
| 167 | WT/AV | F | 9 | 5 | 20220420 | |

| | | | | | | |
|---|---|---|---|---|---|---|
| 167 | WT/AV | F | 9 | 6 | 20220420 | |
| 167 | WT/AV | F | 9 | 7 | 20220420 | |
| 167 | WT/AV | F | 9 | 8 | 20220420 | |
| 167 | WT/AV | F | 9 | 9 | 20220420 | |
| 167 | WT/AV | F | 9 | 10 | 20220420 | |
| 167 | WT/AV | F | 9 | 11 | 20220420 | |
| 167 | WT/AV | F | 9 | 12 | 20220420 | |
| 167 | WT/AV | F | 9 | | | 60 |
| 167 | WT/AV | F | 13 | 1 | 20220516 | |
| 167 | WT/AV | F | 13 | 2 | 20220516 | |
| 167 | WT/AV | F | 13 | 3 | 20220516 | |
| 167 | WT/AV | F | 13 | 4 | 20220516 | |
| 167 | WT/AV | F | 13 | 5 | 20220516 | |
| 167 | WT/AV | F | 13 | 6 | 20220516 | |
| 167 | WT/AV | F | 13 | 7 | 20220516 | |
| 167 | WT/AV | F | 13 | 8 | 20220516 | |
| 167 | WT/AV | F | 13 | 9 | 20220516 | |
| 167 | WT/AV | F | 13 | 10 | 20220516 | |
| 167 | WT/AV | F | 13 | 11 | 20220516 | |
| 167 | WT/AV | F | 13 | 12 | 20220516 | |
| 167 | WT/AV | F | 13 | | | 60 |
| 168 | WT/AV | M | 5 | 1 | 20220321 | |
| 168 | WT/AV | M | 5 | 2 | 20220321 | |
| 168 | WT/AV | M | 5 | 3 | 20220321 | |
| 168 | WT/AV | M | 5 | 4 | 20220321 | |
| 168 | WT/AV | M | 5 | 5 | 20220321 | |
| 168 | WT/AV | M | 5 | 6 | 20220321 | |
| 168 | WT/AV | M | 5 | 7 | 20220321 | |
| 168 | WT/AV | M | 5 | 8 | 20220321 | |
| 168 | WT/AV | M | 5 | 9 | 20220321 | |
| 168 | WT/AV | M | 5 | 10 | 20220321 | |
| 168 | WT/AV | M | 5 | 11 | 20220321 | |
| 168 | WT/AV | M | 5 | 12 | 20220321 | |
| 168 | WT/AV | M | 5 | | | 60 |
| 168 | WT/AV | M | 9 | 1 | 20220420 | |
| 168 | WT/AV | M | 9 | 2 | 20220420 | |
| 168 | WT/AV | M | 9 | 3 | 20220420 | |
| 168 | WT/AV | M | 9 | 4 | 20220420 | |
| 168 | WT/AV | M | 9 | 5 | 20220420 | |
| 168 | WT/AV | M | 9 | 6 | 20220420 | |
| 168 | WT/AV | M | 9 | 7 | 20220420 | |

| | | | | | | |
|---|---|---|---|---|---|---|
| 168 | WT/AV | M | 9 | 8 | 20220420 | |
| 168 | WT/AV | M | 9 | 9 | 20220420 | |
| 168 | WT/AV | M | 9 | 10 | 20220420 | |
| 168 | WT/AV | M | 9 | 11 | 20220420 | |
| 168 | WT/AV | M | 9 | 12 | 20220420 | |
| 168 | WT/AV | M | 9 | | | 60 |
| 169 | WT | M | 5 | 1 | 20220321 | |
| 169 | WT | M | 5 | 2 | 20220321 | |
| 169 | WT | M | 5 | 3 | 20220321 | |
| 169 | WT | M | 5 | 4 | 20220321 | |
| 169 | WT | M | 5 | 5 | 20220321 | |
| 169 | WT | M | 5 | 6 | 20220321 | |
| 169 | WT | M | 5 | 7 | 20220321 | |
| 169 | WT | M | 5 | 8 | 20220321 | |
| 169 | WT | M | 5 | 9 | 20220321 | |
| 169 | WT | M | 5 | 10 | 20220321 | |
| 169 | WT | M | 5 | 11 | 20220321 | |
| 169 | WT | M | 5 | 12 | 20220321 | |
| 169 | WT | M | 5 | | | 60 |
| 169 | WT | M | 9 | 1 | 20220420 | |
| 169 | WT | M | 9 | 2 | 20220420 | |
| 169 | WT | M | 9 | 3 | 20220420 | |
| 169 | WT | M | 9 | 4 | 20220420 | |
| 169 | WT | M | 9 | 5 | 20220420 | |
| 169 | WT | M | 9 | 6 | 20220420 | |
| 169 | WT | M | 9 | 7 | 20220420 | |
| 169 | WT | M | 9 | 8 | 20220420 | |
| 169 | WT | M | 9 | 9 | 20220420 | |
| 169 | WT | M | 9 | 10 | 20220420 | |
| 169 | WT | M | 9 | 11 | 20220420 | |
| 169 | WT | M | 9 | 12 | 20220420 | |
| 169 | WT | M | 9 | | | 60 |
| 195 | WT | F | 5 | 1 | 20220404 | |
| 195 | WT | F | 5 | 2 | 20220404 | |
| 195 | WT | F | 5 | 3 | 20220404 | |
| 195 | WT | F | 5 | 4 | 20220404 | |
| 195 | WT | F | 5 | 5 | 20220404 | |
| 195 | WT | F | 5 | 6 | 20220404 | |
| 195 | WT | F | 5 | 7 | 20220404 | |
| 195 | WT | F | 5 | 8 | 20220404 | |
| 195 | WT | F | 5 | 9 | 20220404 | |

| | | | | | | |
|---|---|---|---|---|---|---|
| 195 | WT | F | 5 | 10 | 20220404 | |
| 195 | WT | F | 5 | 11 | 20220404 | |
| 195 | WT | F | 5 | 12 | 20220404 | |
| 195 | WT | F | 5 | | | 60 |
| 195 | WT | F | 9 | 1 | 20220502 | |
| 195 | WT | F | 9 | 2 | 20220502 | |
| 195 | WT | F | 9 | 3 | 20220502 | |
| 195 | WT | F | 9 | 4 | 20220502 | |
| 195 | WT | F | 9 | 5 | 20220502 | |
| 195 | WT | F | 9 | 6 | 20220502 | |
| 195 | WT | F | 9 | 7 | 20220502 | |
| 195 | WT | F | 9 | 8 | 20220502 | |
| 195 | WT | F | 9 | 9 | 20220502 | |
| 195 | WT | F | 9 | 10 | 20220502 | |
| 195 | WT | F | 9 | 11 | 20220502 | |
| 195 | WT | F | 9 | 12 | 20220502 | |
| 195 | WT | F | 9 | | | 60 |
| 195 | WT | F | 13 | | 20220531 | |
| 195 | WT | F | 13 | | 20220531 | |
| 195 | WT | F | 13 | | 20220531 | |
| 195 | WT | F | 13 | | 20220531 | |
| 195 | WT | F | 13 | | 20220531 | |
| 195 | WT | F | 13 | | 20220531 | |
| 195 | WT | F | 13 | | 20220531 | |
| 195 | WT | F | 13 | | 20220531 | |
| 195 | WT | F | 13 | | 20220531 | |
| 195 | WT | F | 13 | | 20220531 | |
| 195 | WT | F | 13 | | 20220531 | |
| 195 | WT | F | 13 | | 20220531 | |
| 195 | WT | F | 13 | | | 60 |
| 196 | OI/AV | F | 5 | 1 | 20220404 | |
| 196 | OI/AV | F | 5 | 2 | 20220404 | |
| 196 | OI/AV | F | 5 | 3 | 20220404 | |
| 196 | OI/AV | F | 5 | 4 | 20220404 | |
| 196 | OI/AV | F | 5 | 5 | 20220404 | |
| 196 | OI/AV | F | 5 | 6 | 20220404 | |
| 196 | OI/AV | F | 5 | 7 | 20220404 | |
| 196 | OI/AV | F | 5 | 8 | 20220404 | |
| 196 | OI/AV | F | 5 | 9 | 20220404 | |
| 196 | OI/AV | F | 5 | 10 | 20220404 | |
| 196 | OI/AV | F | 5 | 11 | 20220404 | |

| | | | | | | |
|---|---|---|---|---|---|---|
| 196 | OI/AV | F | 5 | 12 | 20220404 | |
| 196 | OI/AV | F | 5 | | | 60 |
| 196 | OI/AV | F | 9 | 1 | 20220502 | |
| 196 | OI/AV | F | 9 | 2 | 20220502 | |
| 196 | OI/AV | F | 9 | 3 | 20220502 | |
| 196 | OI/AV | F | 9 | 4 | 20220502 | |
| 196 | OI/AV | F | 9 | 5 | 20220502 | |
| 196 | OI/AV | F | 9 | 6 | 20220502 | |
| 196 | OI/AV | F | 9 | 7 | 20220502 | |
| 196 | OI/AV | F | 9 | 8 | 20220502 | |
| 196 | OI/AV | F | 9 | 9 | 20220502 | |
| 196 | OI/AV | F | 9 | 10 | 20220502 | |
| 196 | OI/AV | F | 9 | 11 | 20220502 | |
| 196 | OI/AV | F | 9 | 12 | 20220502 | |
| 196 | OI/AV | F | 9 | | | 60 |
| 196 | OI/AV | F | 13 | 122127 | | |
| 196 | OI/AV | F | 13 | 122627 | | |
| 196 | OI/AV | F | 13 | 123127 | | |
| 196 | OI/AV | F | 13 | 123627 | | |
| 196 | OI/AV | F | 13 | 124127 | | |
| 196 | OI/AV | F | 13 | 124627 | | |
| 196 | OI/AV | F | 13 | 125127 | | |
| 196 | OI/AV | F | 13 | 125627 | | |
| 196 | OI/AV | F | 13 | 130127 | | |
| 196 | OI/AV | F | 13 | 130627 | | |
| 196 | OI/AV | F | 13 | 131127 | | |
| 196 | OI/AV | F | 13 | 131627 | | |
| 196 | OI/AV | F | 13 | | | 60 |
| 197 | OI/AV | F | 5 | 1 | 20220404 | |
| 197 | OI/AV | F | 5 | 2 | 20220404 | |
| 197 | OI/AV | F | 5 | 3 | 20220404 | |
| 197 | OI/AV | F | 5 | 4 | 20220404 | |
| 197 | OI/AV | F | 5 | 5 | 20220404 | |
| 197 | OI/AV | F | 5 | 6 | 20220404 | |
| 197 | OI/AV | F | 5 | 7 | 20220404 | |
| 197 | OI/AV | F | 5 | 8 | 20220404 | |
| 197 | OI/AV | F | 5 | 9 | 20220404 | |
| 197 | OI/AV | F | 5 | 10 | 20220404 | |
| 197 | OI/AV | F | 5 | 11 | 20220404 | |
| 197 | OI/AV | F | 5 | 12 | 20220404 | |
| 197 | OI/AV | F | 5 | | | 60 |

| | | | | | |
|---|---|---|---|---|---|
| 197 OI/AV | F | 9 | 1 | 20220502 | |
| 197 OI/AV | F | 9 | 2 | 20220502 | |
| 197 OI/AV | F | 9 | 3 | 20220502 | |
| 197 OI/AV | F | 9 | 4 | 20220502 | |
| 197 OI/AV | F | 9 | 5 | 20220502 | |
| 197 OI/AV | F | 9 | 6 | 20220502 | |
| 197 OI/AV | F | 9 | 7 | 20220502 | |
| 197 OI/AV | F | 9 | 8 | 20220502 | |
| 197 OI/AV | F | 9 | 9 | 20220502 | |
| 197 OI/AV | F | 9 | 10 | 20220502 | |
| 197 OI/AV | F | 9 | 11 | 20220502 | |
| 197 OI/AV | F | 9 | 12 | 20220502 | |
| 197 OI/AV | F | 9 | | | 60 |
| 197 OI/AV | F | 13 | | 20220531 | |
| 197 OI/AV | F | 13 | | 20220531 | |
| 197 OI/AV | F | 13 | | 20220531 | |
| 197 OI/AV | F | 13 | | 20220531 | |
| 197 OI/AV | F | 13 | | 20220531 | |
| 197 OI/AV | F | 13 | | 20220531 | |
| 197 OI/AV | F | 13 | | 20220531 | |
| 197 OI/AV | F | 13 | | 20220531 | |
| 197 OI/AV | F | 13 | | 20220531 | |
| 197 OI/AV | F | 13 | | 20220531 | |
| 197 OI/AV | F | 13 | | 20220531 | |
| 197 OI/AV | F | 13 | | 20220531 | |
| 197 OI/AV | F | 13 | | | 60 |
| 198 OI | F | 5 | 1 | 20220404 | |
| 198 OI | F | 5 | 2 | 20220404 | |
| 198 OI | F | 5 | 3 | 20220404 | |
| 198 OI | F | 5 | 4 | 20220404 | |
| 198 OI | F | 5 | 5 | 20220404 | |
| 198 OI | F | 5 | 6 | 20220404 | |
| 198 OI | F | 5 | 7 | 20220404 | |
| 198 OI | F | 5 | 8 | 20220404 | |
| 198 OI | F | 5 | 9 | 20220404 | |
| 198 OI | F | 5 | 10 | 20220404 | |
| 198 OI | F | 5 | 11 | 20220404 | |
| 198 OI | F | 5 | 12 | 20220404 | |
| 198 OI | F | 5 | | | 60 |
| 198 OI | F | 9 | 1 | 20220502 | |
| 198 OI | F | 9 | 2 | 20220502 | |

| | | | | | | |
|---|---|---|---|---|---|---|
| 198 | OI | F | 9 | 3 | 20220502 | |
| 198 | OI | F | 9 | 4 | 20220502 | |
| 198 | OI | F | 9 | 5 | 20220502 | |
| 198 | OI | F | 9 | 6 | 20220502 | |
| 198 | OI | F | 9 | 7 | 20220502 | |
| 198 | OI | F | 9 | 8 | 20220502 | |
| 198 | OI | F | 9 | 9 | 20220502 | |
| 198 | OI | F | 9 | 10 | 20220502 | |
| 198 | OI | F | 9 | 11 | 20220502 | |
| 198 | OI | F | 9 | 12 | 20220502 | |
| 198 | OI | F | 9 | | | 60 |
| 198 | OI | F | 13 | | 20220531 | |
| 198 | OI | F | 13 | | 20220531 | |
| 198 | OI | F | 13 | | 20220531 | |
| 198 | OI | F | 13 | | 20220531 | |
| 198 | OI | F | 13 | | 20220531 | |
| 198 | OI | F | 13 | | 20220531 | |
| 198 | OI | F | 13 | | 20220531 | |
| 198 | OI | F | 13 | | 20220531 | |
| 198 | OI | F | 13 | | 20220531 | |
| 198 | OI | F | 13 | | 20220531 | |
| 198 | OI | F | 13 | | 20220531 | |
| 198 | OI | F | 13 | | 20220531 | |
| 198 | OI | F | 13 | | | 60 |
| 301 | WT/AV | F | 5 | 1 | 20220404 | |
| 301 | WT/AV | F | 5 | 2 | 20220404 | |
| 301 | WT/AV | F | 5 | 3 | 20220404 | |
| 301 | WT/AV | F | 5 | 4 | 20220404 | |
| 301 | WT/AV | F | 5 | 5 | 20220404 | |
| 301 | WT/AV | F | 5 | 6 | 20220404 | |
| 301 | WT/AV | F | 5 | 7 | 20220404 | |
| 301 | WT/AV | F | 5 | 8 | 20220404 | |
| 301 | WT/AV | F | 5 | 9 | 20220404 | |
| 301 | WT/AV | F | 5 | 10 | 20220404 | |
| 301 | WT/AV | F | 5 | 11 | 20220404 | |
| 301 | WT/AV | F | 5 | 12 | 20220404 | |
| 301 | WT/AV | F | 5 | | | 60 |
| 301 | WT/AV | F | 9 | 1 | 20220502 | |
| 301 | WT/AV | F | 9 | 2 | 20220502 | |
| 301 | WT/AV | F | 9 | 3 | 20220502 | |
| 301 | WT/AV | F | 9 | 4 | 20220502 | |

| | | | | | | |
|---|---|---|---|---|---|---|
| 301 | WT/AV | F | 9 | 5 | 20220502 | |
| 301 | WT/AV | F | 9 | 6 | 20220502 | |
| 301 | WT/AV | F | 9 | 7 | 20220502 | |
| 301 | WT/AV | F | 9 | 8 | 20220502 | |
| 301 | WT/AV | F | 9 | 9 | 20220502 | |
| 301 | WT/AV | F | 9 | 10 | 20220502 | |
| 301 | WT/AV | F | 9 | 11 | 20220502 | |
| 301 | WT/AV | F | 9 | 12 | 20220502 | |
| 301 | WT/AV | F | 9 | | | 60 |
| 301 | WT/AV | F | 13 | | 20220531 | |
| 301 | WT/AV | F | 13 | | 20220531 | |
| 301 | WT/AV | F | 13 | | 20220531 | |
| 301 | WT/AV | F | 13 | | 20220531 | |
| 301 | WT/AV | F | 13 | | 20220531 | |
| 301 | WT/AV | F | 13 | | 20220531 | |
| 301 | WT/AV | F | 13 | | 20220531 | |
| 301 | WT/AV | F | 13 | | 20220531 | |
| 301 | WT/AV | F | 13 | | 20220531 | |
| 301 | WT/AV | F | 13 | | 20220531 | |
| 301 | WT/AV | F | 13 | | 20220531 | |
| 301 | WT/AV | F | 13 | | 20220531 | |
| 301 | WT/AV | F | 13 | | | 60 |
| 302 | WT | F | 5 | 1 | 20220404 | |
| 302 | WT | F | 5 | 2 | 20220404 | |
| 302 | WT | F | 5 | 3 | 20220404 | |
| 302 | WT | F | 5 | 4 | 20220404 | |
| 302 | WT | F | 5 | 5 | 20220404 | |
| 302 | WT | F | 5 | 6 | 20220404 | |
| 302 | WT | F | 5 | 7 | 20220404 | |
| 302 | WT | F | 5 | 8 | 20220404 | |
| 302 | WT | F | 5 | 9 | 20220404 | |
| 302 | WT | F | 5 | 10 | 20220404 | |
| 302 | WT | F | 5 | 11 | 20220404 | |
| 302 | WT | F | 5 | 12 | 20220404 | |
| 302 | WT | F | 5 | | | 60 |
| 302 | WT | F | 9 | 1 | 20220502 | |
| 302 | WT | F | 9 | 2 | 20220502 | |
| 302 | WT | F | 9 | 3 | 20220502 | |
| 302 | WT | F | 9 | 4 | 20220502 | |
| 302 | WT | F | 9 | 5 | 20220502 | |
| 302 | WT | F | 9 | 6 | 20220502 | |

| | | | | | | |
|---|---|---|---|---|---|---|
| 302 | WT | F | 9 | 7 | 20220502 | |
| 302 | WT | F | 9 | 8 | 20220502 | |
| 302 | WT | F | 9 | 9 | 20220502 | |
| 302 | WT | F | 9 | 10 | 20220502 | |
| 302 | WT | F | 9 | 11 | 20220502 | |
| 302 | WT | F | 9 | 12 | 20220502 | |
| 302 | WT | F | 9 | | | 60 |
| 302 | WT | F | 13 | | 20220531 | |
| 302 | WT | F | 13 | | 20220531 | |
| 302 | WT | F | 13 | | 20220531 | |
| 302 | WT | F | 13 | | 20220531 | |
| 302 | WT | F | 13 | | 20220531 | |
| 302 | WT | F | 13 | | 20220531 | |
| 302 | WT | F | 13 | | 20220531 | |
| 302 | WT | F | 13 | | 20220531 | |
| 302 | WT | F | 13 | | 20220531 | |
| 302 | WT | F | 13 | | 20220531 | |
| 302 | WT | F | 13 | | 20220531 | |
| 302 | WT | F | 13 | | 20220531 | |
| 302 | WT | F | 13 | | | 60 |
| 303 | WT | M | 5 | | 20220405 | |
| 303 | WT | M | 5 | | 20220405 | |
| 303 | WT | M | 5 | | 20220405 | |
| 303 | WT | M | 5 | | 20220405 | |
| 303 | WT | M | 5 | | 20220405 | |
| 303 | WT | M | 5 | | 20220405 | |
| 303 | WT | M | 5 | | 20220405 | |
| 303 | WT | M | 5 | | 20220405 | |
| 303 | WT | M | 5 | | 20220405 | |
| 303 | WT | M | 5 | | 20220405 | |
| 303 | WT | M | 5 | | 20220405 | |
| 303 | WT | M | 5 | | 20220405 | |
| 303 | WT | M | 5 | | | 60 |
| 303 | WT | M | 9 | 1 | 20220502 | |
| 303 | WT | M | 9 | 2 | 20220502 | |
| 303 | WT | M | 9 | 3 | 20220502 | |
| 303 | WT | M | 9 | 4 | 20220502 | |
| 303 | WT | M | 9 | 5 | 20220502 | |
| 303 | WT | M | 9 | 6 | 20220502 | |
| 303 | WT | M | 9 | 7 | 20220502 | |
| 303 | WT | M | 9 | 8 | 20220502 | |

| | | | | | | |
|---|---|---|---|---|---|---|
| 303 | WT | M | 9 | 9 | 20220502 | |
| 303 | WT | M | 9 | 10 | 20220502 | |
| 303 | WT | M | 9 | 11 | 20220502 | |
| 303 | WT | M | 9 | 12 | 20220502 | |
| 303 | WT | M | 9 | | | 60 |
| 303 | WT | M | 13 | | 20220531 | |
| 303 | WT | M | 13 | | 20220531 | |
| 303 | WT | M | 13 | | 20220531 | |
| 303 | WT | M | 13 | | 20220531 | |
| 303 | WT | M | 13 | | 20220531 | |
| 303 | WT | M | 13 | | 20220531 | |
| 303 | WT | M | 13 | | 20220531 | |
| 303 | WT | M | 13 | | 20220531 | |
| 303 | WT | M | 13 | | 20220531 | |
| 303 | WT | M | 13 | | 20220531 | |
| 303 | WT | M | 13 | | 20220531 | |
| 303 | WT | M | 13 | | 20220531 | |
| 303 | WT | M | 13 | | | 60 |
| 304 | WT/AV | M | 5 | | 20220405 | |
| 304 | WT/AV | M | 5 | | 20220405 | |
| 304 | WT/AV | M | 5 | | 20220405 | |
| 304 | WT/AV | M | 5 | | 20220405 | |
| 304 | WT/AV | M | 5 | | 20220405 | |
| 304 | WT/AV | M | 5 | | 20220405 | |
| 304 | WT/AV | M | 5 | | 20220405 | |
| 304 | WT/AV | M | 5 | | 20220405 | |
| 304 | WT/AV | M | 5 | | 20220405 | |
| 304 | WT/AV | M | 5 | | 20220405 | |
| 304 | WT/AV | M | 5 | | 20220405 | |
| 304 | WT/AV | M | 5 | | 20220405 | |
| 304 | WT/AV | M | 5 | | | 60 |
| 304 | WT/AV | M | 9 | 1 | 20220502 | |
| 304 | WT/AV | M | 9 | 2 | 20220502 | |
| 304 | WT/AV | M | 9 | 3 | 20220502 | |
| 304 | WT/AV | M | 9 | 4 | 20220502 | |
| 304 | WT/AV | M | 9 | 5 | 20220502 | |
| 304 | WT/AV | M | 9 | 6 | 20220502 | |
| 304 | WT/AV | M | 9 | 7 | 20220502 | |
| 304 | WT/AV | M | 9 | 8 | 20220502 | |
| 304 | WT/AV | M | 9 | 9 | 20220502 | |
| 304 | WT/AV | M | 9 | 10 | 20220502 | |

| | | | | | | |
|---|---|---|---|---|---|---|
| 304 | WT/AV | M | 9 | 11 | 20220502 | |
| 304 | WT/AV | M | 9 | 12 | 20220502 | |
| 304 | WT/AV | M | 9 | | | 60 |
| 304 | WT/AV | M | 13 | | 20220531 | |
| 304 | WT/AV | M | 13 | | 20220531 | |
| 304 | WT/AV | M | 13 | | 20220531 | |
| 304 | WT/AV | M | 13 | | 20220531 | |
| 304 | WT/AV | M | 13 | | 20220531 | |
| 304 | WT/AV | M | 13 | | 20220531 | |
| 304 | WT/AV | M | 13 | | 20220531 | |
| 304 | WT/AV | M | 13 | | 20220531 | |
| 304 | WT/AV | M | 13 | | 20220531 | |
| 304 | WT/AV | M | 13 | | 20220531 | |
| 304 | WT/AV | M | 13 | | 20220531 | |
| 304 | WT/AV | M | 13 | | 20220531 | |
| 304 | WT/AV | M | 13 | | | 60 |
| 317 | WT | M | 5 | 1 | 20220405 | |
| 317 | WT | M | 5 | 2 | 20220405 | |
| 317 | WT | M | 5 | 3 | 20220405 | |
| 317 | WT | M | 5 | 4 | 20220405 | |
| 317 | WT | M | 5 | 5 | 20220405 | |
| 317 | WT | M | 5 | 6 | 20220405 | |
| 317 | WT | M | 5 | 7 | 20220405 | |
| 317 | WT | M | 5 | 8 | 20220405 | |
| 317 | WT | M | 5 | 9 | 20220405 | |
| 317 | WT | M | 5 | 10 | 20220405 | |
| 317 | WT | M | 5 | 11 | 20220405 | |
| 317 | WT | M | 5 | 12 | 20220405 | |
| 317 | WT | M | 5 | | | 60 |
| 317 | WT | M | 9 | 1 | 20220503 | |
| 317 | WT | M | 9 | 2 | 20220503 | |
| 317 | WT | M | 9 | 3 | 20220503 | |
| 317 | WT | M | 9 | 4 | 20220503 | |
| 317 | WT | M | 9 | 5 | 20220503 | |
| 317 | WT | M | 9 | 6 | 20220503 | |
| 317 | WT | M | 9 | 7 | 20220503 | |
| 317 | WT | M | 9 | 8 | 20220503 | |
| 317 | WT | M | 9 | 9 | 20220503 | |
| 317 | WT | M | 9 | 10 | 20220503 | |
| 317 | WT | M | 9 | 11 | 20220503 | |
| 317 | WT | M | 9 | 12 | 20220503 | |

| | | | | | | |
|---|---|---|---|---|---|---|
| 317 | WT | M | 9 | | | 60 |
| 317 | WT | M | 13 | 1 | 20220601 | |
| 317 | WT | M | 13 | 2 | 20220601 | |
| 317 | WT | M | 13 | 3 | 20220601 | |
| 317 | WT | M | 13 | 4 | 20220601 | |
| 317 | WT | M | 13 | 5 | 20220601 | |
| 317 | WT | M | 13 | 6 | 20220601 | |
| 317 | WT | M | 13 | 7 | 20220601 | |
| 317 | WT | M | 13 | 8 | 20220601 | |
| 317 | WT | M | 13 | 9 | 20220601 | |
| 317 | WT | M | 13 | 10 | 20220601 | |
| 317 | WT | M | 13 | 11 | 20220601 | |
| 317 | WT | M | 13 | 12 | 20220601 | |
| 317 | WT | M | 13 | | | 60 |
| 318 | OI/AV | F | 5 | 1 | 20220404 | |
| 318 | OI/AV | F | 5 | 2 | 20220404 | |
| 318 | OI/AV | F | 5 | 3 | 20220404 | |
| 318 | OI/AV | F | 5 | 4 | 20220404 | |
| 318 | OI/AV | F | 5 | 5 | 20220404 | |
| 318 | OI/AV | F | 5 | 6 | 20220404 | |
| 318 | OI/AV | F | 5 | 7 | 20220404 | |
| 318 | OI/AV | F | 5 | 8 | 20220404 | |
| 318 | OI/AV | F | 5 | 9 | 20220404 | |
| 318 | OI/AV | F | 5 | 10 | 20220404 | |
| 318 | OI/AV | F | 5 | 11 | 20220404 | |
| 318 | OI/AV | F | 5 | 12 | 20220404 | |
| 318 | OI/AV | F | 5 | | | 60 |
| 318 | OI/AV | F | 9 | 1 | 20220503 | |
| 318 | OI/AV | F | 9 | 2 | 20220503 | |
| 318 | OI/AV | F | 9 | 3 | 20220503 | |
| 318 | OI/AV | F | 9 | 4 | 20220503 | |
| 318 | OI/AV | F | 9 | 5 | 20220503 | |
| 318 | OI/AV | F | 9 | 6 | 20220503 | |
| 318 | OI/AV | F | 9 | 7 | 20220503 | |
| 318 | OI/AV | F | 9 | 8 | 20220503 | |
| 318 | OI/AV | F | 9 | 9 | 20220503 | |
| 318 | OI/AV | F | 9 | 10 | 20220503 | |
| 318 | OI/AV | F | 9 | 11 | 20220503 | |
| 318 | OI/AV | F | 9 | 12 | 20220503 | |
| 318 | OI/AV | F | 9 | | | 60 |
| 318 | OI/AV | F | 13 | 1 | 20220601 | |

| | | | | | |
|---|---|---|---|---|---|
| 318 | OI/AV | F | 13 | 2 | 20220601 | |
| 318 | OI/AV | F | 13 | 3 | 20220601 | |
| 318 | OI/AV | F | 13 | 4 | 20220601 | |
| 318 | OI/AV | F | 13 | 5 | 20220601 | |
| 318 | OI/AV | F | 13 | 6 | 20220601 | |
| 318 | OI/AV | F | 13 | 7 | 20220601 | |
| 318 | OI/AV | F | 13 | 8 | 20220601 | |
| 318 | OI/AV | F | 13 | 9 | 20220601 | |
| 318 | OI/AV | F | 13 | 10 | 20220601 | |
| 318 | OI/AV | F | 13 | 11 | 20220601 | |
| 318 | OI/AV | F | 13 | 12 | 20220601 | |
| 318 | OI/AV | F | 13 | | | 60 |
| 319 | OI | M | 5 | 1 | 20220405 | |
| 319 | OI | M | 5 | 2 | 20220405 | |
| 319 | OI | M | 5 | 3 | 20220405 | |
| 319 | OI | M | 5 | 4 | 20220405 | |
| 319 | OI | M | 5 | 5 | 20220405 | |
| 319 | OI | M | 5 | 6 | 20220405 | |
| 319 | OI | M | 5 | 7 | 20220405 | |
| 319 | OI | M | 5 | 8 | 20220405 | |
| 319 | OI | M | 5 | 9 | 20220405 | |
| 319 | OI | M | 5 | 10 | 20220405 | |
| 319 | OI | M | 5 | 11 | 20220405 | |
| 319 | OI | M | 5 | 12 | 20220405 | |
| 319 | OI | M | 5 | | | 60 |
| 319 | OI | M | 9 | 1 | 20220503 | |
| 319 | OI | M | 9 | 2 | 20220503 | |
| 319 | OI | M | 9 | 3 | 20220503 | |
| 319 | OI | M | 9 | 4 | 20220503 | |
| 319 | OI | M | 9 | 5 | 20220503 | |
| 319 | OI | M | 9 | 6 | 20220503 | |
| 319 | OI | M | 9 | 7 | 20220503 | |
| 319 | OI | M | 9 | 8 | 20220503 | |
| 319 | OI | M | 9 | 9 | 20220503 | |
| 319 | OI | M | 9 | 10 | 20220503 | |
| 319 | OI | M | 9 | 11 | 20220503 | |
| 319 | OI | M | 9 | 12 | 20220503 | |
| 319 | OI | M | 9 | | | 60 |
| 319 | OI | M | 13 | 1 | 20220601 | |
| 319 | OI | M | 13 | 2 | 20220601 | |
| 319 | OI | M | 13 | 3 | 20220601 | |

| | | | | | |
|---|---|---|---|---|---|
| 319 OI | M | 13 | 4 | 20220601 | |
| 319 OI | M | 13 | 5 | 20220601 | |
| 319 OI | M | 13 | 6 | 20220601 | |
| 319 OI | M | 13 | 7 | 20220601 | |
| 319 OI | M | 13 | 8 | 20220601 | |
| 319 OI | M | 13 | 9 | 20220601 | |
| 319 OI | M | 13 | 10 | 20220601 | |
| 319 OI | M | 13 | 11 | 20220601 | |
| 319 OI | M | 13 | 12 | 20220601 | |
| 319 OI | M | 13 | | | 60 |
| 320 OI/AV | M | 5 | 1 | 20220405 | |
| 320 OI/AV | M | 5 | 2 | 20220405 | |
| 320 OI/AV | M | 5 | 3 | 20220405 | |
| 320 OI/AV | M | 5 | 4 | 20220405 | |
| 320 OI/AV | M | 5 | 5 | 20220405 | |
| 320 OI/AV | M | 5 | 6 | 20220405 | |
| 320 OI/AV | M | 5 | 7 | 20220405 | |
| 320 OI/AV | M | 5 | 8 | 20220405 | |
| 320 OI/AV | M | 5 | 9 | 20220405 | |
| 320 OI/AV | M | 5 | 10 | 20220405 | |
| 320 OI/AV | M | 5 | 11 | 20220405 | |
| 320 OI/AV | M | 5 | 12 | 20220405 | |
| 320 OI/AV | M | 5 | | | 60 |
| 320 OI/AV | M | 9 | 1 | 20220503 | |
| 320 OI/AV | M | 9 | 2 | 20220503 | |
| 320 OI/AV | M | 9 | 3 | 20220503 | |
| 320 OI/AV | M | 9 | 4 | 20220503 | |
| 320 OI/AV | M | 9 | 5 | 20220503 | |
| 320 OI/AV | M | 9 | 6 | 20220503 | |
| 320 OI/AV | M | 9 | 7 | 20220503 | |
| 320 OI/AV | M | 9 | 8 | 20220503 | |
| 320 OI/AV | M | 9 | 9 | 20220503 | |
| 320 OI/AV | M | 9 | 10 | 20220503 | |
| 320 OI/AV | M | 9 | 11 | 20220503 | |
| 320 OI/AV | M | 9 | 12 | 20220503 | |
| 320 OI/AV | M | 9 | | | 60 |
| 320 OI/AV | M | 13 | 1 | 20220601 | |
| 320 OI/AV | M | 13 | 2 | 20220601 | |
| 320 OI/AV | M | 13 | 3 | 20220601 | |
| 320 OI/AV | M | 13 | 4 | 20220601 | |
| 320 OI/AV | M | 13 | 5 | 20220601 | |

| | | | | | |
|---|---|---|---|---|---|
| 320 OI/AV | M | 13 | 6 | 20220601 | |
| 320 OI/AV | M | 13 | 7 | 20220601 | |
| 320 OI/AV | M | 13 | 8 | 20220601 | |
| 320 OI/AV | M | 13 | 9 | 20220601 | |
| 320 OI/AV | M | 13 | 10 | 20220601 | |
| 320 OI/AV | M | 13 | 11 | 20220601 | |
| 320 OI/AV | M | 13 | 12 | 20220601 | |
| 320 OI/AV | M | 13 | | | 60 |
| 321 OI/AV | M | 5 | 1 | 20220405 | |
| 321 OI/AV | M | 5 | 2 | 20220405 | |
| 321 OI/AV | M | 5 | 3 | 20220405 | |
| 321 OI/AV | M | 5 | 4 | 20220405 | |
| 321 OI/AV | M | 5 | 5 | 20220405 | |
| 321 OI/AV | M | 5 | 6 | 20220405 | |
| 321 OI/AV | M | 5 | 7 | 20220405 | |
| 321 OI/AV | M | 5 | 8 | 20220405 | |
| 321 OI/AV | M | 5 | 9 | 20220405 | |
| 321 OI/AV | M | 5 | 10 | 20220405 | |
| 321 OI/AV | M | 5 | 11 | 20220405 | |
| 321 OI/AV | M | 5 | 12 | 20220405 | |
| 321 OI/AV | M | 5 | | | 60 |
| 321 OI/AV | M | 9 | 1 | 20220503 | |
| 321 OI/AV | M | 9 | 2 | 20220503 | |
| 321 OI/AV | M | 9 | 3 | 20220503 | |
| 321 OI/AV | M | 9 | 4 | 20220503 | |
| 321 OI/AV | M | 9 | 5 | 20220503 | |
| 321 OI/AV | M | 9 | 6 | 20220503 | |
| 321 OI/AV | M | 9 | 7 | 20220503 | |
| 321 OI/AV | M | 9 | 8 | 20220503 | |
| 321 OI/AV | M | 9 | 9 | 20220503 | |
| 321 OI/AV | M | 9 | 10 | 20220503 | |
| 321 OI/AV | M | 9 | 11 | 20220503 | |
| 321 OI/AV | M | 9 | 12 | 20220503 | |
| 321 OI/AV | M | 9 | | | 60 |
| 321 OI/AV | M | 13 | 1 | 20220601 | |
| 321 OI/AV | M | 13 | 2 | 20220601 | |
| 321 OI/AV | M | 13 | 3 | 20220601 | |
| 321 OI/AV | M | 13 | 4 | 20220601 | |
| 321 OI/AV | M | 13 | 5 | 20220601 | |
| 321 OI/AV | M | 13 | 6 | 20220601 | |
| 321 OI/AV | M | 13 | 7 | 20220601 | |

| | | | | | | |
|---|---|---|---|---|---|---|
| 321 | OI/AV | M | 13 | 8 | 20220601 | |
| 321 | OI/AV | M | 13 | 9 | 20220601 | |
| 321 | OI/AV | M | 13 | 10 | 20220601 | |
| 321 | OI/AV | M | 13 | 11 | 20220601 | |
| 321 | OI/AV | M | 13 | 12 | 20220601 | |
| 321 | OI/AV | M | 13 | | | 60 |
| 322 | WT | M | 5 | | 20220405 | |
| 322 | WT | M | 5 | | 20220405 | |
| 322 | WT | M | 5 | | 20220405 | |
| 322 | WT | M | 5 | | 20220405 | |
| 322 | WT | M | 5 | | 20220405 | |
| 322 | WT | M | 5 | | 20220405 | |
| 322 | WT | M | 5 | | 20220405 | |
| 322 | WT | M | 5 | | 20220405 | |
| 322 | WT | M | 5 | | 20220405 | |
| 322 | WT | M | 5 | | 20220405 | |
| 322 | WT | M | 5 | | 20220405 | |
| 322 | WT | M | 5 | | 20220405 | |
| 322 | WT | M | 5 | | | 60 |
| 322 | WT | M | 9 | 1 | 20220503 | |
| 322 | WT | M | 9 | 2 | 20220503 | |
| 322 | WT | M | 9 | 3 | 20220503 | |
| 322 | WT | M | 9 | 4 | 20220503 | |
| 322 | WT | M | 9 | 5 | 20220503 | |
| 322 | WT | M | 9 | 6 | 20220503 | |
| 322 | WT | M | 9 | 7 | 20220503 | |
| 322 | WT | M | 9 | 8 | 20220503 | |
| 322 | WT | M | 9 | 9 | 20220503 | |
| 322 | WT | M | 9 | 10 | 20220503 | |
| 322 | WT | M | 9 | 11 | 20220503 | |
| 322 | WT | M | 9 | 12 | 20220503 | |
| 322 | WT | M | 9 | | | 60 |
| 322 | WT | M | 13 | 1 | 20220601 | |
| 322 | WT | M | 13 | 2 | 20220601 | |
| 322 | WT | M | 13 | 3 | 20220601 | |
| 322 | WT | M | 13 | 4 | 20220601 | |
| 322 | WT | M | 13 | 5 | 20220601 | |
| 322 | WT | M | 13 | 6 | 20220601 | |
| 322 | WT | M | 13 | 7 | 20220601 | |
| 322 | WT | M | 13 | 8 | 20220601 | |
| 322 | WT | M | 13 | 9 | 20220601 | |

| | | | | | | |
|---|---|---|---|---|---|---|
| 322 | WT | M | 13 | 10 | 20220601 | |
| 322 | WT | M | 13 | 11 | 20220601 | |
| 322 | WT | M | 13 | 12 | 20220601 | |
| 322 | WT | M | 13 | | | 60 |
| 332 | WT/AV | F | 5 | 1 | 20220404 | |
| 332 | WT/AV | F | 5 | 2 | 20220404 | |
| 332 | WT/AV | F | 5 | 3 | 20220404 | |
| 332 | WT/AV | F | 5 | 4 | 20220404 | |
| 332 | WT/AV | F | 5 | 5 | 20220404 | |
| 332 | WT/AV | F | 5 | 6 | 20220404 | |
| 332 | WT/AV | F | 5 | 7 | 20220404 | |
| 332 | WT/AV | F | 5 | 8 | 20220404 | |
| 332 | WT/AV | F | 5 | 9 | 20220404 | |
| 332 | WT/AV | F | 5 | 10 | 20220404 | |
| 332 | WT/AV | F | 5 | 11 | 20220404 | |
| 332 | WT/AV | F | 5 | 12 | 20220404 | |
| 332 | WT/AV | F | 5 | | | 60 |
| 332 | WT/AV | F | 9 | 1 | 20220503 | |
| 332 | WT/AV | F | 9 | 2 | 20220503 | |
| 332 | WT/AV | F | 9 | 3 | 20220503 | |
| 332 | WT/AV | F | 9 | 4 | 20220503 | |
| 332 | WT/AV | F | 9 | 5 | 20220503 | |
| 332 | WT/AV | F | 9 | 6 | 20220503 | |
| 332 | WT/AV | F | 9 | 7 | 20220503 | |
| 332 | WT/AV | F | 9 | 8 | 20220503 | |
| 332 | WT/AV | F | 9 | 9 | 20220503 | |
| 332 | WT/AV | F | 9 | 10 | 20220503 | |
| 332 | WT/AV | F | 9 | 11 | 20220503 | |
| 332 | WT/AV | F | 9 | 12 | 20220503 | |
| 332 | WT/AV | F | 9 | | | 60 |
| 332 | WT/AV | F | 13 | 1 | 20220601 | |
| 332 | WT/AV | F | 13 | 2 | 20220601 | |
| 332 | WT/AV | F | 13 | 3 | 20220601 | |
| 332 | WT/AV | F | 13 | 4 | 20220601 | |
| 332 | WT/AV | F | 13 | 5 | 20220601 | |
| 332 | WT/AV | F | 13 | 6 | 20220601 | |
| 332 | WT/AV | F | 13 | 7 | 20220601 | |
| 332 | WT/AV | F | 13 | 8 | 20220601 | |
| 332 | WT/AV | F | 13 | 9 | 20220601 | |
| 332 | WT/AV | F | 13 | 10 | 20220601 | |
| 332 | WT/AV | F | 13 | 11 | 20220601 | |

| | | | | | |
|---|---|---|---|---|---|
| 332 | WT/AV | F | 13 | 12 | 20220601 | |
| 332 | WT/AV | F | 13 | | | 60 |
| 339 | OI/AV | F | 5 | 1 | 20220411 | |
| 339 | OI/AV | F | 5 | 2 | 20220411 | |
| 339 | OI/AV | F | 5 | 3 | 20220411 | |
| 339 | OI/AV | F | 5 | 4 | 20220411 | |
| 339 | OI/AV | F | 5 | 5 | 20220411 | |
| 339 | OI/AV | F | 5 | 6 | 20220411 | |
| 339 | OI/AV | F | 5 | 7 | 20220411 | |
| 339 | OI/AV | F | 5 | 8 | 20220411 | |
| 339 | OI/AV | F | 5 | 9 | 20220411 | |
| 339 | OI/AV | F | 5 | 10 | 20220411 | |
| 339 | OI/AV | F | 5 | 11 | 20220411 | |
| 339 | OI/AV | F | 5 | 12 | 20220411 | |
| 339 | OI/AV | F | 5 | | | 60 |
| 339 | OI/AV | F | 9 | 1 | 20220509 | |
| 339 | OI/AV | F | 9 | 2 | 20220509 | |
| 339 | OI/AV | F | 9 | 3 | 20220509 | |
| 339 | OI/AV | F | 9 | 4 | 20220509 | |
| 339 | OI/AV | F | 9 | 5 | 20220509 | |
| 339 | OI/AV | F | 9 | 6 | 20220509 | |
| 339 | OI/AV | F | 9 | 7 | 20220509 | |
| 339 | OI/AV | F | 9 | 8 | 20220509 | |
| 339 | OI/AV | F | 9 | 9 | 20220509 | |
| 339 | OI/AV | F | 9 | 10 | 20220509 | |
| 339 | OI/AV | F | 9 | 11 | 20220509 | |
| 339 | OI/AV | F | 9 | 12 | 20220509 | |
| 339 | OI/AV | F | 9 | | | 60 |
| 339 | OI/AV | F | 13 | 1 | 20220606 | |
| 339 | OI/AV | F | 13 | 2 | 20220606 | |
| 339 | OI/AV | F | 13 | 3 | 20220606 | |
| 339 | OI/AV | F | 13 | 4 | 20220606 | |
| 339 | OI/AV | F | 13 | 5 | 20220606 | |
| 339 | OI/AV | F | 13 | 6 | 20220606 | |
| 339 | OI/AV | F | 13 | 7 | 20220606 | |
| 339 | OI/AV | F | 13 | 8 | 20220606 | |
| 339 | OI/AV | F | 13 | 9 | 20220606 | |
| 339 | OI/AV | F | 13 | 10 | 20220606 | |
| 339 | OI/AV | F | 13 | 11 | 20220606 | |
| 339 | OI/AV | F | 13 | 12 | 20220606 | |
| 339 | OI/AV | F | 13 | | | 60 |

| | | | | | |
|---|---|---|---|---|---|
| 340 | OI/AV | F | 5 | 1 | 20220411 | |
| 340 | OI/AV | F | 5 | 2 | 20220411 | |
| 340 | OI/AV | F | 5 | 3 | 20220411 | |
| 340 | OI/AV | F | 5 | 4 | 20220411 | |
| 340 | OI/AV | F | 5 | 5 | 20220411 | |
| 340 | OI/AV | F | 5 | 6 | 20220411 | |
| 340 | OI/AV | F | 5 | 7 | 20220411 | |
| 340 | OI/AV | F | 5 | 8 | 20220411 | |
| 340 | OI/AV | F | 5 | 9 | 20220411 | |
| 340 | OI/AV | F | 5 | 10 | 20220411 | |
| 340 | OI/AV | F | 5 | 11 | 20220411 | |
| 340 | OI/AV | F | 5 | 12 | 20220411 | |
| 340 | OI/AV | F | 5 | | | 60 |
| 340 | OI/AV | F | 9 | 1 | 20220509 | |
| 340 | OI/AV | F | 9 | 2 | 20220509 | |
| 340 | OI/AV | F | 9 | 3 | 20220509 | |
| 340 | OI/AV | F | 9 | 4 | 20220509 | |
| 340 | OI/AV | F | 9 | 5 | 20220509 | |
| 340 | OI/AV | F | 9 | 6 | 20220509 | |
| 340 | OI/AV | F | 9 | 7 | 20220509 | |
| 340 | OI/AV | F | 9 | 8 | 20220509 | |
| 340 | OI/AV | F | 9 | 9 | 20220509 | |
| 340 | OI/AV | F | 9 | 10 | 20220509 | |
| 340 | OI/AV | F | 9 | 11 | 20220509 | |
| 340 | OI/AV | F | 9 | 12 | 20220509 | |
| 340 | OI/AV | F | 9 | | | 60 |
| 340 | OI/AV | F | 13 | 1 | 20220606 | |
| 340 | OI/AV | F | 13 | 2 | 20220606 | |
| 340 | OI/AV | F | 13 | 3 | 20220606 | |
| 340 | OI/AV | F | 13 | 4 | 20220606 | |
| 340 | OI/AV | F | 13 | 5 | 20220606 | |
| 340 | OI/AV | F | 13 | 6 | 20220606 | |
| 340 | OI/AV | F | 13 | 7 | 20220606 | |
| 340 | OI/AV | F | 13 | 8 | 20220606 | |
| 340 | OI/AV | F | 13 | 9 | 20220606 | |
| 340 | OI/AV | F | 13 | 10 | 20220606 | |
| 340 | OI/AV | F | 13 | 11 | 20220606 | |
| 340 | OI/AV | F | 13 | 12 | 20220606 | |
| 340 | OI/AV | F | 13 | | | 60 |
| 341 | WT/AV | M | 5 | 1 | 20220411 | |
| 341 | WT/AV | M | 5 | 2 | 20220411 | |

| | | | | | |
|---|---|---|---|---|---|
| 341 | WT/AV | M | 5 | 3 | 20220411 | |
| 341 | WT/AV | M | 5 | 4 | 20220411 | |
| 341 | WT/AV | M | 5 | 5 | 20220411 | |
| 341 | WT/AV | M | 5 | 6 | 20220411 | |
| 341 | WT/AV | M | 5 | 7 | 20220411 | |
| 341 | WT/AV | M | 5 | 8 | 20220411 | |
| 341 | WT/AV | M | 5 | 9 | 20220411 | |
| 341 | WT/AV | M | 5 | 10 | 20220411 | |
| 341 | WT/AV | M | 5 | 11 | 20220411 | |
| 341 | WT/AV | M | 5 | 12 | 20220411 | |
| 341 | WT/AV | M | 5 | | | 60 |
| 341 | WT/AV | M | 9 | 1 | 20220509 | |
| 341 | WT/AV | M | 9 | 2 | 20220509 | |
| 341 | WT/AV | M | 9 | 3 | 20220509 | |
| 341 | WT/AV | M | 9 | 4 | 20220509 | |
| 341 | WT/AV | M | 9 | 5 | 20220509 | |
| 341 | WT/AV | M | 9 | 6 | 20220509 | |
| 341 | WT/AV | M | 9 | 7 | 20220509 | |
| 341 | WT/AV | M | 9 | 8 | 20220509 | |
| 341 | WT/AV | M | 9 | 9 | 20220509 | |
| 341 | WT/AV | M | 9 | 10 | 20220509 | |
| 341 | WT/AV | M | 9 | 11 | 20220509 | |
| 341 | WT/AV | M | 9 | 12 | 20220509 | |
| 341 | WT/AV | M | 9 | | | 60 |
| 341 | WT/AV | M | 13 | 1 | 20220606 | |
| 341 | WT/AV | M | 13 | 2 | 20220606 | |
| 341 | WT/AV | M | 13 | 3 | 20220606 | |
| 341 | WT/AV | M | 13 | 4 | 20220606 | |
| 341 | WT/AV | M | 13 | 5 | 20220606 | |
| 341 | WT/AV | M | 13 | 6 | 20220606 | |
| 341 | WT/AV | M | 13 | 7 | 20220606 | |
| 341 | WT/AV | M | 13 | 8 | 20220606 | |
| 341 | WT/AV | M | 13 | 9 | 20220606 | |
| 341 | WT/AV | M | 13 | 10 | 20220606 | |
| 341 | WT/AV | M | 13 | 11 | 20220606 | |
| 341 | WT/AV | M | 13 | 12 | 20220606 | |
| 341 | WT/AV | M | 13 | | | 60 |
| 342 | WT | M | 5 | 1 | 20220412 | |
| 342 | WT | M | 5 | 2 | 20220412 | |
| 342 | WT | M | 5 | 3 | 20220412 | |
| 342 | WT | M | 5 | 4 | 20220412 | |

| | | | | | | |
|---|---|---|---|---|---|---|
| 342 | WT | M | 5 | 5 | 20220412 | |
| 342 | WT | M | 5 | 6 | 20220412 | |
| 342 | WT | M | 5 | 7 | 20220412 | |
| 342 | WT | M | 5 | 8 | 20220412 | |
| 342 | WT | M | 5 | 9 | 20220412 | |
| 342 | WT | M | 5 | 10 | 20220412 | |
| 342 | WT | M | 5 | 11 | 20220412 | |
| 342 | WT | M | 5 | 12 | 20220412 | |
| 342 | WT | M | 5 | | | 60 |
| 342 | WT | M | 9 | 1 | 20220509 | |
| 342 | WT | M | 9 | 2 | 20220509 | |
| 342 | WT | M | 9 | 3 | 20220509 | |
| 342 | WT | M | 9 | 4 | 20220509 | |
| 342 | WT | M | 9 | 5 | 20220509 | |
| 342 | WT | M | 9 | 6 | 20220509 | |
| 342 | WT | M | 9 | 7 | 20220509 | |
| 342 | WT | M | 9 | 8 | 20220509 | |
| 342 | WT | M | 9 | 9 | 20220509 | |
| 342 | WT | M | 9 | 10 | 20220509 | |
| 342 | WT | M | 9 | 11 | 20220509 | |
| 342 | WT | M | 9 | 12 | 20220509 | |
| 342 | WT | M | 9 | | | 60 |
| 342 | WT | M | 13 | 1 | 20220606 | |
| 342 | WT | M | 13 | 2 | 20220606 | |
| 342 | WT | M | 13 | 3 | 20220606 | |
| 342 | WT | M | 13 | 4 | 20220606 | |
| 342 | WT | M | 13 | 5 | 20220606 | |
| 342 | WT | M | 13 | 6 | 20220606 | |
| 342 | WT | M | 13 | 7 | 20220606 | |
| 342 | WT | M | 13 | 8 | 20220606 | |
| 342 | WT | M | 13 | 9 | 20220606 | |
| 342 | WT | M | 13 | 10 | 20220606 | |
| 342 | WT | M | 13 | 11 | 20220606 | |
| 342 | WT | M | 13 | 12 | 20220606 | |
| 342 | WT | M | 13 | | | 60 |
| 343 | WT/AV | M | 5 | 1 | 20220412 | |
| 343 | WT/AV | M | 5 | 2 | 20220412 | |
| 343 | WT/AV | M | 5 | 3 | 20220412 | |
| 343 | WT/AV | M | 5 | 4 | 20220412 | |
| 343 | WT/AV | M | 5 | 5 | 20220412 | |
| 343 | WT/AV | M | 5 | 6 | 20220412 | |

| | | | | | |
|---|---|---|---|---|---|
| 343 | WT/AV | M | 5 | 7 | 20220412 | |
| 343 | WT/AV | M | 5 | 8 | 20220412 | |
| 343 | WT/AV | M | 5 | 9 | 20220412 | |
| 343 | WT/AV | M | 5 | 10 | 20220412 | |
| 343 | WT/AV | M | 5 | 11 | 20220412 | |
| 343 | WT/AV | M | 5 | 12 | 20220412 | |
| 343 | WT/AV | M | 5 | | | 60 |
| 343 | WT/AV | M | 9 | 1 | 20220509 | |
| 343 | WT/AV | M | 9 | 2 | 20220509 | |
| 343 | WT/AV | M | 9 | 3 | 20220509 | |
| 343 | WT/AV | M | 9 | 4 | 20220509 | |
| 343 | WT/AV | M | 9 | 5 | 20220509 | |
| 343 | WT/AV | M | 9 | 6 | 20220509 | |
| 343 | WT/AV | M | 9 | 7 | 20220509 | |
| 343 | WT/AV | M | 9 | 8 | 20220509 | |
| 343 | WT/AV | M | 9 | 9 | 20220509 | |
| 343 | WT/AV | M | 9 | 10 | 20220509 | |
| 343 | WT/AV | M | 9 | 11 | 20220509 | |
| 343 | WT/AV | M | 9 | 12 | 20220509 | |
| 343 | WT/AV | M | 9 | | | 60 |
| 343 | WT/AV | M | 13 | 1 | 20220606 | |
| 343 | WT/AV | M | 13 | 2 | 20220606 | |
| 343 | WT/AV | M | 13 | 3 | 20220606 | |
| 343 | WT/AV | M | 13 | 4 | 20220606 | |
| 343 | WT/AV | M | 13 | 5 | 20220606 | |
| 343 | WT/AV | M | 13 | 6 | 20220606 | |
| 343 | WT/AV | M | 13 | 7 | 20220606 | |
| 343 | WT/AV | M | 13 | 8 | 20220606 | |
| 343 | WT/AV | M | 13 | 9 | 20220606 | |
| 343 | WT/AV | M | 13 | 10 | 20220606 | |
| 343 | WT/AV | M | 13 | 11 | 20220606 | |
| 343 | WT/AV | M | 13 | 12 | 20220606 | |
| 343 | WT/AV | M | 13 | | | 60 |
| 344 | OI | M | 5 | 1 | 20220412 | |
| 344 | OI | M | 5 | 2 | 20220412 | |
| 344 | OI | M | 5 | 3 | 20220412 | |
| 344 | OI | M | 5 | 4 | 20220412 | |
| 344 | OI | M | 5 | 5 | 20220412 | |
| 344 | OI | M | 5 | 6 | 20220412 | |
| 344 | OI | M | 5 | 7 | 20220412 | |
| 344 | OI | M | 5 | 8 | 20220412 | |

| | | | | | | |
|---|---|---|---|---|---|---|
| 344 | OI | M | 5 | 9 | 20220412 | |
| 344 | OI | M | 5 | 10 | 20220412 | |
| 344 | OI | M | 5 | 11 | 20220412 | |
| 344 | OI | M | 5 | 12 | 20220412 | |
| 344 | OI | M | 5 | | | 60 |
| 344 | OI | M | 9 | 1 | 20220509 | |
| 344 | OI | M | 9 | 2 | 20220509 | |
| 344 | OI | M | 9 | 3 | 20220509 | |
| 344 | OI | M | 9 | 4 | 20220509 | |
| 344 | OI | M | 9 | 5 | 20220509 | |
| 344 | OI | M | 9 | 6 | 20220509 | |
| 344 | OI | M | 9 | 7 | 20220509 | |
| 344 | OI | M | 9 | 8 | 20220509 | |
| 344 | OI | M | 9 | 9 | 20220509 | |
| 344 | OI | M | 9 | 10 | 20220509 | |
| 344 | OI | M | 9 | 11 | 20220509 | |
| 344 | OI | M | 9 | 12 | 20220509 | |
| 344 | OI | M | 9 | | | 60 |
| 344 | OI | M | 13 | 1 | 20220606 | |
| 344 | OI | M | 13 | 2 | 20220606 | |
| 344 | OI | M | 13 | 3 | 20220606 | |
| 344 | OI | M | 13 | 4 | 20220606 | |
| 344 | OI | M | 13 | 5 | 20220606 | |
| 344 | OI | M | 13 | 6 | 20220606 | |
| 344 | OI | M | 13 | 7 | 20220606 | |
| 344 | OI | M | 13 | 8 | 20220606 | |
| 344 | OI | M | 13 | 9 | 20220606 | |
| 344 | OI | M | 13 | 10 | 20220606 | |
| 344 | OI | M | 13 | 11 | 20220606 | |
| 344 | OI | M | 13 | 12 | 20220606 | |
| 344 | OI | M | 13 | | | 60 |
| 345 | OI | F | 5 | 1 | 20220418 | |
| 345 | OI | F | 5 | 2 | 20220418 | |
| 345 | OI | F | 5 | 3 | 20220418 | |
| 345 | OI | F | 5 | 4 | 20220418 | |
| 345 | OI | F | 5 | 5 | 20220418 | |
| 345 | OI | F | 5 | 6 | 20220418 | |
| 345 | OI | F | 5 | 7 | 20220418 | |
| 345 | OI | F | 5 | 8 | 20220418 | |
| 345 | OI | F | 5 | 9 | 20220418 | |
| 345 | OI | F | 5 | 10 | 20220418 | |

| | | | | | | |
|---|---|---|---|---|---|---|
| 345 | OI | F | 5 | 11 | 20220418 | |
| 345 | OI | F | 5 | 12 | 20220418 | |
| 345 | OI | F | 5 | | | 60 |
| 345 | OI | F | 9 | 1 | 20220517 | |
| 345 | OI | F | 9 | 2 | 20220517 | |
| 345 | OI | F | 9 | 3 | 20220517 | |
| 345 | OI | F | 9 | 4 | 20220517 | |
| 345 | OI | F | 9 | 5 | 20220517 | |
| 345 | OI | F | 9 | 6 | 20220517 | |
| 345 | OI | F | 9 | 7 | 20220517 | |
| 345 | OI | F | 9 | 8 | 20220517 | |
| 345 | OI | F | 9 | 9 | 20220517 | |
| 345 | OI | F | 9 | 10 | 20220517 | |
| 345 | OI | F | 9 | 11 | 20220517 | |
| 345 | OI | F | 9 | 12 | 20220517 | |
| 345 | OI | F | 9 | | | 60 |
| 345 | OI | F | 13 | | 20220614 | |
| 345 | OI | F | 13 | | 20220614 | |
| 345 | OI | F | 13 | | 20220614 | |
| 345 | OI | F | 13 | | 20220614 | |
| 345 | OI | F | 13 | | 20220614 | |
| 345 | OI | F | 13 | | 20220614 | |
| 345 | OI | F | 13 | | 20220614 | |
| 345 | OI | F | 13 | | 20220614 | |
| 345 | OI | F | 13 | | 20220614 | |
| 345 | OI | F | 13 | | 20220614 | |
| 345 | OI | F | 13 | | 20220614 | |
| 345 | OI | F | 13 | | 20220614 | |
| 345 | OI | F | 13 | | | 60 |
| 349 | OI | M | 5 | 1 | 20220418 | |
| 349 | OI | M | 5 | 2 | 20220418 | |
| 349 | OI | M | 5 | 3 | 20220418 | |
| 349 | OI | M | 5 | 4 | 20220418 | |
| 349 | OI | M | 5 | 5 | 20220418 | |
| 349 | OI | M | 5 | 6 | 20220418 | |
| 349 | OI | M | 5 | 7 | 20220418 | |
| 349 | OI | M | 5 | 8 | 20220418 | |
| 349 | OI | M | 5 | 9 | 20220418 | |
| 349 | OI | M | 5 | 10 | 20220418 | |
| 349 | OI | M | 5 | 11 | 20220418 | |
| 349 | OI | M | 5 | 12 | 20220418 | |

| | | | | | | |
|---|---|---|---|---|---|---|
| 349 | OI | M | 5 | | | 60 |
| 349 | OI | M | 9 | 1 | 20220517 | |
| 349 | OI | M | 9 | 2 | 20220517 | |
| 349 | OI | M | 9 | 3 | 20220517 | |
| 349 | OI | M | 9 | 4 | 20220517 | |
| 349 | OI | M | 9 | 5 | 20220517 | |
| 349 | OI | M | 9 | 6 | 20220517 | |
| 349 | OI | M | 9 | 7 | 20220517 | |
| 349 | OI | M | 9 | 8 | 20220517 | |
| 349 | OI | M | 9 | 9 | 20220517 | |
| 349 | OI | M | 9 | 10 | 20220517 | |
| 349 | OI | M | 9 | 11 | 20220517 | |
| 349 | OI | M | 9 | 12 | 20220517 | |
| 349 | OI | M | 9 | | | 60 |
| 349 | OI | M | 13 | 1 | 20220614 | |
| 349 | OI | M | 13 | 2 | 20220614 | |
| 349 | OI | M | 13 | 3 | 20220614 | |
| 349 | OI | M | 13 | 4 | 20220614 | |
| 349 | OI | M | 13 | 5 | 20220614 | |
| 349 | OI | M | 13 | 6 | 20220614 | |
| 349 | OI | M | 13 | 7 | 20220614 | |
| 349 | OI | M | 13 | 8 | 20220614 | |
| 349 | OI | M | 13 | 9 | 20220614 | |
| 349 | OI | M | 13 | 10 | 20220614 | |
| 349 | OI | M | 13 | 11 | 20220614 | |
| 349 | OI | M | 13 | 12 | 20220614 | |
| 349 | OI | M | 13 | | | 60 |
| 350 | OI/AV | M | 5 | 1 | 20220418 | |
| 350 | OI/AV | M | 5 | 2 | 20220418 | |
| 350 | OI/AV | M | 5 | 3 | 20220418 | |
| 350 | OI/AV | M | 5 | 4 | 20220418 | |
| 350 | OI/AV | M | 5 | 5 | 20220418 | |
| 350 | OI/AV | M | 5 | 6 | 20220418 | |
| 350 | OI/AV | M | 5 | 7 | 20220418 | |
| 350 | OI/AV | M | 5 | 8 | 20220418 | |
| 350 | OI/AV | M | 5 | 9 | 20220418 | |
| 350 | OI/AV | M | 5 | 10 | 20220418 | |
| 350 | OI/AV | M | 5 | 11 | 20220418 | |
| 350 | OI/AV | M | 5 | 12 | 20220418 | |
| 350 | OI/AV | M | 5 | | | 60 |
| 350 | OI/AV | M | 9 | 1 | 20220517 | |

| | | | | | |
|---|---|---|---|---|---|
| 350 OI/AV | M | 9 | 2 | 20220517 | |
| 350 OI/AV | M | 9 | 3 | 20220517 | |
| 350 OI/AV | M | 9 | 4 | 20220517 | |
| 350 OI/AV | M | 9 | 5 | 20220517 | |
| 350 OI/AV | M | 9 | 6 | 20220517 | |
| 350 OI/AV | M | 9 | 7 | 20220517 | |
| 350 OI/AV | M | 9 | 8 | 20220517 | |
| 350 OI/AV | M | 9 | 9 | 20220517 | |
| 350 OI/AV | M | 9 | 10 | 20220517 | |
| 350 OI/AV | M | 9 | 11 | 20220517 | |
| 350 OI/AV | M | 9 | 12 | 20220517 | |
| 350 OI/AV | M | 9 | | | 60 |
| 350 OI/AV | M | 13 | 1 | 20220614 | |
| 350 OI/AV | M | 13 | 2 | 20220614 | |
| 350 OI/AV | M | 13 | 3 | 20220614 | |
| 350 OI/AV | M | 13 | 4 | 20220614 | |
| 350 OI/AV | M | 13 | 5 | 20220614 | |
| 350 OI/AV | M | 13 | 6 | 20220614 | |
| 350 OI/AV | M | 13 | 7 | 20220614 | |
| 350 OI/AV | M | 13 | 8 | 20220614 | |
| 350 OI/AV | M | 13 | 9 | 20220614 | |
| 350 OI/AV | M | 13 | 10 | 20220614 | |
| 350 OI/AV | M | 13 | 11 | 20220614 | |
| 350 OI/AV | M | 13 | 12 | 20220614 | |
| 350 OI/AV | M | 13 | | | 60 |
| 351 OI | M | 5 | 1 | 20220418 | |
| 351 OI | M | 5 | 2 | 20220418 | |
| 351 OI | M | 5 | 3 | 20220418 | |
| 351 OI | M | 5 | 4 | 20220418 | |
| 351 OI | M | 5 | 5 | 20220418 | |
| 351 OI | M | 5 | 6 | 20220418 | |
| 351 OI | M | 5 | 7 | 20220418 | |
| 351 OI | M | 5 | 8 | 20220418 | |
| 351 OI | M | 5 | 9 | 20220418 | |
| 351 OI | M | 5 | 10 | 20220418 | |
| 351 OI | M | 5 | 11 | 20220418 | |
| 351 OI | M | 5 | 12 | 20220418 | |
| 351 OI | M | 5 | | | 60 |
| 351 OI | M | 9 | 1 | 20220517 | |
| 351 OI | M | 9 | 2 | 20220517 | |
| 351 OI | M | 9 | 3 | 20220517 | |

| | | | | | |
|---|---|---|---|---|---|
| 351 OI | M | 9 | 4 | 20220517 | |
| 351 OI | M | 9 | 5 | 20220517 | |
| 351 OI | M | 9 | 6 | 20220517 | |
| 351 OI | M | 9 | 7 | 20220517 | |
| 351 OI | M | 9 | 8 | 20220517 | |
| 351 OI | M | 9 | 9 | 20220517 | |
| 351 OI | M | 9 | 10 | 20220517 | |
| 351 OI | M | 9 | 11 | 20220517 | |
| 351 OI | M | 9 | 12 | 20220517 | |
| 351 OI | M | 9 | | | 60 |
| 351 OI | M | 13 | | 20220614 | |
| 351 OI | M | 13 | | 20220614 | |
| 351 OI | M | 13 | | 20220614 | |
| 351 OI | M | 13 | | 20220614 | |
| 351 OI | M | 13 | | 20220614 | |
| 351 OI | M | 13 | | 20220614 | |
| 351 OI | M | 13 | | 20220614 | |
| 351 OI | M | 13 | | 20220614 | |
| 351 OI | M | 13 | | 20220614 | |
| 351 OI | M | 13 | | 20220614 | |
| 351 OI | M | 13 | | 20220614 | |
| 351 OI | M | 13 | | 20220614 | |
| 351 OI | M | 13 | | | 60 |
| 814 WT/AV | M | 5 | 1 | 20220523 | |
| 814 WT/AV | M | 5 | 2 | 20220523 | |
| 814 WT/AV | M | 5 | 3 | 20220523 | |
| 814 WT/AV | M | 5 | 4 | 20220523 | |
| 814 WT/AV | M | 5 | 5 | 20220523 | |
| 814 WT/AV | M | 5 | 6 | 20220523 | |
| 814 WT/AV | M | 5 | 7 | 20220523 | |
| 814 WT/AV | M | 5 | 8 | 20220523 | |
| 814 WT/AV | M | 5 | 9 | 20220523 | |
| 814 WT/AV | M | 5 | 10 | 20220523 | |
| 814 WT/AV | M | 5 | 11 | 20220523 | |
| 814 WT/AV | M | 5 | 12 | 20220523 | |
| 814 WT/AV | M | 5 | | | 60 |
| 814 WT/AV | M | 9 | 1 | 20220621 | |
| 814 WT/AV | M | 9 | 2 | 20220621 | |
| 814 WT/AV | M | 9 | 3 | 20220621 | |
| 814 WT/AV | M | 9 | 4 | 20220621 | |
| 814 WT/AV | M | 9 | 5 | 20220621 | |

| | | | | | | |
|---|---|---|---|---|---|---|
| 814 | WT/AV | M | 9 | 6 | 20220621 | |
| 814 | WT/AV | M | 9 | 7 | 20220621 | |
| 814 | WT/AV | M | 9 | 8 | 20220621 | |
| 814 | WT/AV | M | 9 | 9 | 20220621 | |
| 814 | WT/AV | M | 9 | 10 | 20220621 | |
| 814 | WT/AV | M | 9 | 11 | 20220621 | |
| 814 | WT/AV | M | 9 | 12 | 20220621 | |
| 814 | WT/AV | M | 9 | | | 60 |
| 814 | WT/AV | M | 13 | 1 | 20220719 | |
| 814 | WT/AV | M | 13 | 2 | 20220719 | |
| 814 | WT/AV | M | 13 | 3 | 20220719 | |
| 814 | WT/AV | M | 13 | 4 | 20220719 | |
| 814 | WT/AV | M | 13 | 5 | 20220719 | |
| 814 | WT/AV | M | 13 | 6 | 20220719 | |
| 814 | WT/AV | M | 13 | 7 | 20220719 | |
| 814 | WT/AV | M | 13 | 8 | 20220719 | |
| 814 | WT/AV | M | 13 | 9 | 20220719 | |
| 814 | WT/AV | M | 13 | 10 | 20220719 | |
| 814 | WT/AV | M | 13 | 11 | 20220719 | |
| 814 | WT/AV | M | 13 | 12 | 20220719 | |
| 814 | WT/AV | M | 13 | | | 60 |
| 816 | WT | F | 5 | 1 | 20220523 | |
| 816 | WT | F | 5 | 2 | 20220523 | |
| 816 | WT | F | 5 | 3 | 20220523 | |
| 816 | WT | F | 5 | 4 | 20220523 | |
| 816 | WT | F | 5 | 5 | 20220523 | |
| 816 | WT | F | 5 | 6 | 20220523 | |
| 816 | WT | F | 5 | 7 | 20220523 | |
| 816 | WT | F | 5 | 8 | 20220523 | |
| 816 | WT | F | 5 | 9 | 20220523 | |
| 816 | WT | F | 5 | 10 | 20220523 | |
| 816 | WT | F | 5 | 11 | 20220523 | |
| 816 | WT | F | 5 | 12 | 20220523 | |
| 816 | WT | F | 5 | | | 60 |
| 816 | WT | F | 9 | 1 | 20220621 | |
| 816 | WT | F | 9 | 2 | 20220621 | |
| 816 | WT | F | 9 | 3 | 20220621 | |
| 816 | WT | F | 9 | 4 | 20220621 | |
| 816 | WT | F | 9 | 5 | 20220621 | |
| 816 | WT | F | 9 | 6 | 20220621 | |
| 816 | WT | F | 9 | 7 | 20220621 | |

| | | | | | |
|---|---|---|---|---|---|
| 816 WT | F | 9 | 8 | 20220621 | |
| 816 WT | F | 9 | 9 | 20220621 | |
| 816 WT | F | 9 | 10 | 20220621 | |
| 816 WT | F | 9 | 11 | 20220621 | |
| 816 WT | F | 9 | 12 | 20220621 | |
| 816 WT | F | 9 | | | 60 |
| 816 WT | F | 13 | 1 | 20220719 | |
| 816 WT | F | 13 | 2 | 20220719 | |
| 816 WT | F | 13 | 3 | 20220719 | |
| 816 WT | F | 13 | 4 | 20220719 | |
| 816 WT | F | 13 | 5 | 20220719 | |
| 816 WT | F | 13 | 6 | 20220719 | |
| 816 WT | F | 13 | 7 | 20220719 | |
| 816 WT | F | 13 | 8 | 20220719 | |
| 816 WT | F | 13 | 9 | 20220719 | |
| 816 WT | F | 13 | 10 | 20220719 | |
| 816 WT | F | 13 | 11 | 20220719 | |
| 816 WT | F | 13 | 12 | 20220719 | |
| 816 WT | F | 13 | | | 60 |
| 857 WT/AV | M | 5 | 1 | 20220621 | |
| 857 WT/AV | M | 5 | 2 | 20220621 | |
| 857 WT/AV | M | 5 | 3 | 20220621 | |
| 857 WT/AV | M | 5 | 4 | 20220621 | |
| 857 WT/AV | M | 5 | 5 | 20220621 | |
| 857 WT/AV | M | 5 | 6 | 20220621 | |
| 857 WT/AV | M | 5 | 7 | 20220621 | |
| 857 WT/AV | M | 5 | 8 | 20220621 | |
| 857 WT/AV | M | 5 | 9 | 20220621 | |
| 857 WT/AV | M | 5 | 10 | 20220621 | |
| 857 WT/AV | M | 5 | 11 | 20220621 | |
| 857 WT/AV | M | 5 | 12 | 20220621 | |
| 857 WT/AV | M | 5 | | | 60 |
| 857 WT/AV | M | 9 | 1 | 20220719 | |
| 857 WT/AV | M | 9 | 2 | 20220719 | |
| 857 WT/AV | M | 9 | 3 | 20220719 | |
| 857 WT/AV | M | 9 | 4 | 20220719 | |
| 857 WT/AV | M | 9 | 5 | 20220719 | |
| 857 WT/AV | M | 9 | 6 | 20220719 | |
| 857 WT/AV | M | 9 | 7 | 20220719 | |
| 857 WT/AV | M | 9 | 8 | 20220719 | |
| 857 WT/AV | M | 9 | 9 | 20220719 | |

| | | | | | | |
|---|---|---|---|---|---|---|
| 857 | WT/AV | M | 9 | 10 | 20220719 | |
| 857 | WT/AV | M | 9 | 11 | 20220719 | |
| 857 | WT/AV | M | 9 | 12 | 20220719 | |
| 857 | WT/AV | M | 9 | | | 60 |
| 858 | WT | F | 5 | 1 | 20220621 | |
| 858 | WT | F | 5 | 2 | 20220621 | |
| 858 | WT | F | 5 | 3 | 20220621 | |
| 858 | WT | F | 5 | 4 | 20220621 | |
| 858 | WT | F | 5 | 5 | 20220621 | |
| 858 | WT | F | 5 | 6 | 20220621 | |
| 858 | WT | F | 5 | 7 | 20220621 | |
| 858 | WT | F | 5 | 8 | 20220621 | |
| 858 | WT | F | 5 | 9 | 20220621 | |
| 858 | WT | F | 5 | 10 | 20220621 | |
| 858 | WT | F | 5 | 11 | 20220621 | |
| 858 | WT | F | 5 | 12 | 20220621 | |
| 858 | WT | F | 5 | | | 60 |
| 858 | WT | F | 9 | 1 | 20220719 | |
| 858 | WT | F | 9 | 2 | 20220719 | |
| 858 | WT | F | 9 | 3 | 20220719 | |
| 858 | WT | F | 9 | 4 | 20220719 | |
| 858 | WT | F | 9 | 5 | 20220719 | |
| 858 | WT | F | 9 | 6 | 20220719 | |
| 858 | WT | F | 9 | 7 | 20220719 | |
| 858 | WT | F | 9 | 8 | 20220719 | |
| 858 | WT | F | 9 | 9 | 20220719 | |
| 858 | WT | F | 9 | 10 | 20220719 | |
| 858 | WT | F | 9 | 11 | 20220719 | |
| 858 | WT | F | 9 | 12 | 20220719 | |
| 858 | WT | F | 9 | | | 60 |
| 872 | WT/AV | M | 5 | | 20220705 | |
| 872 | WT/AV | M | 5 | | 20220705 | |
| 872 | WT/AV | M | 5 | | 20220705 | |
| 872 | WT/AV | M | 5 | | 20220705 | |
| 872 | WT/AV | M | 5 | | 20220705 | |
| 872 | WT/AV | M | 5 | | 20220705 | |
| 872 | WT/AV | M | 5 | | 20220705 | |
| 872 | WT/AV | M | 5 | | 20220705 | |
| 872 | WT/AV | M | 5 | | 20220705 | |
| 872 | WT/AV | M | 5 | | 20220705 | |
| 872 | WT/AV | M | 5 | | 20220705 | |

| | | | | | |
|---|---|---|---|---|---|
| 872 | WT/AV | M | 5 | 20220705 | |
| 872 | WT/AV | M | 5 | | 60 |
| 872 | WT/AV | M | 9 | 20220802 | |
| 872 | WT/AV | M | 9 | 20220802 | |
| 872 | WT/AV | M | 9 | 20220802 | |
| 872 | WT/AV | M | 9 | 20220802 | |
| 872 | WT/AV | M | 9 | 20220802 | |
| 872 | WT/AV | M | 9 | 20220802 | |
| 872 | WT/AV | M | 9 | 20220802 | |
| 872 | WT/AV | M | 9 | 20220802 | |
| 872 | WT/AV | M | 9 | 20220802 | |
| 872 | WT/AV | M | 9 | 20220802 | |
| 872 | WT/AV | M | 9 | 20220802 | |
| 872 | WT/AV | M | 9 | 20220802 | |
| 872 | WT/AV | M | 9 | | 60 |
| 872 | WT/AV | M | 13 | 20220829 | |
| 872 | WT/AV | M | 13 | 20220829 | |
| 872 | WT/AV | M | 13 | 20220829 | |
| 872 | WT/AV | M | 13 | 20220829 | |
| 872 | WT/AV | M | 13 | 20220829 | |
| 872 | WT/AV | M | 13 | 20220829 | |
| 872 | WT/AV | M | 13 | 20220829 | |
| 872 | WT/AV | M | 13 | 20220829 | |
| 872 | WT/AV | M | 13 | 20220829 | |
| 872 | WT/AV | M | 13 | 20220829 | |
| 872 | WT/AV | M | 13 | 20220829 | |
| 872 | WT/AV | M | 13 | 20220829 | |
| 872 | WT/AV | M | 13 | | 60 |
| 873 | OI | M | 5 | 20220705 | |
| 873 | OI | M | 5 | 20220705 | |
| 873 | OI | M | 5 | 20220705 | |
| 873 | OI | M | 5 | 20220705 | |
| 873 | OI | M | 5 | 20220705 | |
| 873 | OI | M | 5 | 20220705 | |
| 873 | OI | M | 5 | 20220705 | |
| 873 | OI | M | 5 | 20220705 | |
| 873 | OI | M | 5 | 20220705 | |
| 873 | OI | M | 5 | 20220705 | |
| 873 | OI | M | 5 | 20220705 | |
| 873 | OI | M | 5 | 20220705 | |
| 873 | OI | M | 5 | | 60 |

| | | | | | | |
|---|---|---|---|---|---|---|
| 873 | OI | M | 9 | 1 | 20220802 | |
| 873 | OI | M | 9 | 2 | 20220802 | |
| 873 | OI | M | 9 | 3 | 20220802 | |
| 873 | OI | M | 9 | 4 | 20220802 | |
| 873 | OI | M | 9 | 5 | 20220802 | |
| 873 | OI | M | 9 | 6 | 20220802 | |
| 873 | OI | M | 9 | 7 | 20220802 | |
| 873 | OI | M | 9 | 8 | 20220802 | |
| 873 | OI | M | 9 | 9 | 20220802 | |
| 873 | OI | M | 9 | 10 | 20220802 | |
| 873 | OI | M | 9 | 11 | 20220802 | |
| 873 | OI | M | 9 | 12 | 20220802 | |
| 873 | OI | M | 9 | | | 60 |
| 873 | OI | M | 13 | | 20220829 | |
| 873 | OI | M | 13 | | 20220829 | |
| 873 | OI | M | 13 | | 20220829 | |
| 873 | OI | M | 13 | | 20220829 | |
| 873 | OI | M | 13 | | 20220829 | |
| 873 | OI | M | 13 | | 20220829 | |
| 873 | OI | M | 13 | | 20220829 | |
| 873 | OI | M | 13 | | 20220829 | |
| 873 | OI | M | 13 | | 20220829 | |
| 873 | OI | M | 13 | | 20220829 | |
| 873 | OI | M | 13 | | 20220829 | |
| 873 | OI | M | 13 | | 20220829 | |
| 873 | OI | M | 13 | | | 60 |
| 941 | OI | F | 5 | 1 | 20220606 | |
| 941 | OI | F | 5 | 2 | 20220606 | |
| 941 | OI | F | 5 | 3 | 20220606 | |
| 941 | OI | F | 5 | 4 | 20220606 | |
| 941 | OI | F | 5 | 5 | 20220606 | |
| 941 | OI | F | 5 | 6 | 20220606 | |
| 941 | OI | F | 5 | 7 | 20220606 | |
| 941 | OI | F | 5 | 8 | 20220606 | |
| 941 | OI | F | 5 | 9 | 20220606 | |
| 941 | OI | F | 5 | 10 | 20220606 | |
| 941 | OI | F | 5 | 11 | 20220606 | |
| 941 | OI | F | 5 | 12 | 20220606 | |
| 941 | OI | F | 5 | | | 60 |
| 943 | OI | M | 5 | 1 | 20220606 | |
| 943 | OI | M | 5 | 2 | 20220606 | |

| | | | | | |
|---|---|---|---|---|---|
| 943 | OI | M | 5 | 3 | 20220606 | |
| 943 | OI | M | 5 | 4 | 20220606 | |
| 943 | OI | M | 5 | 5 | 20220606 | |
| 943 | OI | M | 5 | 6 | 20220606 | |
| 943 | OI | M | 5 | 7 | 20220606 | |
| 943 | OI | M | 5 | 8 | 20220606 | |
| 943 | OI | M | 5 | 9 | 20220606 | |
| 943 | OI | M | 5 | 10 | 20220606 | |
| 943 | OI | M | 5 | 11 | 20220606 | |
| 943 | OI | M | 5 | 12 | 20220606 | |
| 943 | OI | M | 5 | | | 60 |
| 944 | WT/AV | M | 5 | 1 | 20220606 | |
| 944 | WT/AV | M | 5 | 2 | 20220606 | |
| 944 | WT/AV | M | 5 | 3 | 20220606 | |
| 944 | WT/AV | M | 5 | 4 | 20220606 | |
| 944 | WT/AV | M | 5 | 5 | 20220606 | |
| 944 | WT/AV | M | 5 | 6 | 20220606 | |
| 944 | WT/AV | M | 5 | 7 | 20220606 | |
| 944 | WT/AV | M | 5 | 8 | 20220606 | |
| 944 | WT/AV | M | 5 | 9 | 20220606 | |
| 944 | WT/AV | M | 5 | 10 | 20220606 | |
| 944 | WT/AV | M | 5 | 11 | 20220606 | |
| 944 | WT/AV | M | 5 | 12 | 20220606 | |
| 944 | WT/AV | M | 5 | | | 60 |
| 944 | WT/AV | M | 9 | 1 | 20220705 | |
| 944 | WT/AV | M | 9 | 2 | 20220705 | |
| 944 | WT/AV | M | 9 | 3 | 20220705 | |
| 944 | WT/AV | M | 9 | 4 | 20220705 | |
| 944 | WT/AV | M | 9 | 5 | 20220705 | |
| 944 | WT/AV | M | 9 | 6 | 20220705 | |
| 944 | WT/AV | M | 9 | 7 | 20220705 | |
| 944 | WT/AV | M | 9 | 8 | 20220705 | |
| 944 | WT/AV | M | 9 | 9 | 20220705 | |
| 944 | WT/AV | M | 9 | 10 | 20220705 | |
| 944 | WT/AV | M | 9 | 11 | 20220705 | |
| 944 | WT/AV | M | 9 | 12 | 20220705 | |
| 944 | WT/AV | M | 9 | | | 60 |
| 944 | WT/AV | M | 13 | 1 | 20220802 | |
| 944 | WT/AV | M | 13 | 2 | 20220802 | |
| 944 | WT/AV | M | 13 | 3 | 20220802 | |
| 944 | WT/AV | M | 13 | 4 | 20220802 | |

| | | | | | |
|---|---|---|---|---|---|
| 944 | WT/AV | M | 13 | 5 | 20220802 | |
| 944 | WT/AV | M | 13 | 6 | 20220802 | |
| 944 | WT/AV | M | 13 | 7 | 20220802 | |
| 944 | WT/AV | M | 13 | 8 | 20220802 | |
| 944 | WT/AV | M | 13 | 9 | 20220802 | |
| 944 | WT/AV | M | 13 | 10 | 20220802 | |
| 944 | WT/AV | M | 13 | 11 | 20220802 | |
| 944 | WT/AV | M | 13 | 12 | 20220802 | |
| 944 | WT/AV | M | 13 | | | 60 |
| 987 | OI | F | 5 | | 20220621 | |
| 987 | OI | F | 5 | | 20220621 | |
| 987 | OI | F | 5 | | 20220621 | |
| 987 | OI | F | 5 | | 20220621 | |
| 987 | OI | F | 5 | | 20220621 | |
| 987 | OI | F | 5 | | 20220621 | |
| 987 | OI | F | 5 | | 20220621 | |
| 987 | OI | F | 5 | | 20220621 | |
| 987 | OI | F | 5 | | 20220621 | |
| 987 | OI | F | 5 | | 20220621 | |
| 987 | OI | F | 5 | | 20220621 | |
| 987 | OI | F | 5 | | 20220621 | |
| 987 | OI | F | 5 | | | 60 |
| 987 | OI | F | 9 | | 20220719 | |
| 987 | OI | F | 9 | | 20220719 | |
| 987 | OI | F | 9 | | 20220719 | |
| 987 | OI | F | 9 | | 20220719 | |
| 987 | OI | F | 9 | | 20220719 | |
| 987 | OI | F | 9 | | 20220719 | |
| 987 | OI | F | 9 | | 20220719 | |
| 987 | OI | F | 9 | | 20220719 | |
| 987 | OI | F | 9 | | 20220719 | |
| 987 | OI | F | 9 | | 20220719 | |
| 987 | OI | F | 9 | | 20220719 | |
| 987 | OI | F | 9 | | 20220719 | |
| 987 | OI | F | 9 | | | 60 |
| 987 | OI | F | 13 | | 20220816 | |
| 987 | OI | F | 13 | | 20220816 | |
| 987 | OI | F | 13 | | 20220816 | |
| 987 | OI | F | 13 | | 20220816 | |
| 987 | OI | F | 13 | | 20220816 | |
| 987 | OI | F | 13 | | 20220816 | |

| | | | | |
|---|---|---|---|---|
| 987 OI | F | 13 | 20220816 | |
| 987 OI | F | 13 | 20220816 | |
| 987 OI | F | 13 | 20220816 | |
| 987 OI | F | 13 | 20220816 | |
| 987 OI | F | 13 | 20220816 | |
| 987 OI | F | 13 | 20220816 | |
| 987 OI | F | 13 | | 60 |
| 989 OI | F | 5 | 20220621 | |
| 989 OI | F | 5 | 20220621 | |
| 989 OI | F | 5 | 20220621 | |
| 989 OI | F | 5 | 20220621 | |
| 989 OI | F | 5 | 20220621 | |
| 989 OI | F | 5 | 20220621 | |
| 989 OI | F | 5 | 20220621 | |
| 989 OI | F | 5 | 20220621 | |
| 989 OI | F | 5 | 20220621 | |
| 989 OI | F | 5 | 20220621 | |
| 989 OI | F | 5 | 20220621 | |
| 989 OI | F | 5 | 20220621 | |
| 989 OI | F | 5 | | 60 |
| 989 OI | F | 9 | 20220719 | |
| 989 OI | F | 9 | 20220719 | |
| 989 OI | F | 9 | 20220719 | |
| 989 OI | F | 9 | 20220719 | |
| 989 OI | F | 9 | 20220719 | |
| 989 OI | F | 9 | 20220719 | |
| 989 OI | F | 9 | 20220719 | |
| 989 OI | F | 9 | 20220719 | |
| 989 OI | F | 9 | 20220719 | |
| 989 OI | F | 9 | 20220719 | |
| 989 OI | F | 9 | 20220719 | |
| 989 OI | F | 9 | 20220719 | |
| 989 OI | F | 9 | | 60 |
| 989 OI | F | 13 | 20220816 | |
| 989 OI | F | 13 | 20220816 | |
| 989 OI | F | 13 | 20220816 | |
| 989 OI | F | 13 | 20220816 | |
| 989 OI | F | 13 | 20220816 | |
| 989 OI | F | 13 | 20220816 | |
| 989 OI | F | 13 | 20220816 | |
| 989 OI | F | 13 | 20220816 | |

| | | | | | |
|---|---|---|---|---|---|
| 989 | OI | F | 13 | | 20220816 | |
| 989 | OI | F | 13 | | 20220816 | |
| 989 | OI | F | 13 | | 20220816 | |
| 989 | OI | F | 13 | | 20220816 | |
| 989 | OI | F | 13 | | | 60 |
| 992 | OI/AV | F | 5 | | 20220621 | |
| 992 | OI/AV | F | 5 | | 20220621 | |
| 992 | OI/AV | F | 5 | | 20220621 | |
| 992 | OI/AV | F | 5 | | 20220621 | |
| 992 | OI/AV | F | 5 | | 20220621 | |
| 992 | OI/AV | F | 5 | | 20220621 | |
| 992 | OI/AV | F | 5 | | 20220621 | |
| 992 | OI/AV | F | 5 | | 20220621 | |
| 992 | OI/AV | F | 5 | | 20220621 | |
| 992 | OI/AV | F | 5 | | 20220621 | |
| 992 | OI/AV | F | 5 | | 20220621 | |
| 992 | OI/AV | F | 5 | | 20220621 | |
| 992 | OI/AV | F | 5 | | | 60 |
| 992 | OI/AV | F | 9 | 1 | 20220719 | |
| 992 | OI/AV | F | 9 | 2 | 20220719 | |
| 992 | OI/AV | F | 9 | 3 | 20220719 | |
| 992 | OI/AV | F | 9 | 4 | 20220719 | |
| 992 | OI/AV | F | 9 | 5 | 20220719 | |
| 992 | OI/AV | F | 9 | 6 | 20220719 | |
| 992 | OI/AV | F | 9 | 7 | 20220719 | |
| 992 | OI/AV | F | 9 | 8 | 20220719 | |
| 992 | OI/AV | F | 9 | 9 | 20220719 | |
| 992 | OI/AV | F | 9 | 10 | 20220719 | |
| 992 | OI/AV | F | 9 | 11 | 20220719 | |
| 992 | OI/AV | F | 9 | 12 | 20220719 | |
| 992 | OI/AV | F | 9 | | | 60 |
| 992 | OI/AV | F | 13 | | | |
| 992 | OI/AV | F | 13 | | 20220816 | |
| 992 | OI/AV | F | 13 | | 20220816 | |
| 992 | OI/AV | F | 13 | | 20220816 | |
| 992 | OI/AV | F | 13 | | 20220816 | |
| 992 | OI/AV | F | 13 | | 20220816 | |
| 992 | OI/AV | F | 13 | | 20220816 | |
| 992 | OI/AV | F | 13 | | 20220816 | |
| 992 | OI/AV | F | 13 | | 20220816 | |
| 992 | OI/AV | F | 13 | | 20220816 | |

| | | | | | | |
|---|---|---|---|---|---|---|
| 992 | OI/AV | F | 13 | | 20220816 | |
| 992 | OI/AV | F | 13 | | 20220816 | |
| 992 | OI/AV | F | 13 | | | 60 |
| 994 | WT/AV | F | 5 | | 20220621 | |
| 994 | WT/AV | F | 5 | | 20220621 | |
| 994 | WT/AV | F | 5 | | 20220621 | |
| 994 | WT/AV | F | 5 | | 20220621 | |
| 994 | WT/AV | F | 5 | | 20220621 | |
| 994 | WT/AV | F | 5 | | 20220621 | |
| 994 | WT/AV | F | 5 | | 20220621 | |
| 994 | WT/AV | F | 5 | | 20220621 | |
| 994 | WT/AV | F | 5 | | 20220621 | |
| 994 | WT/AV | F | 5 | | 20220621 | |
| 994 | WT/AV | F | 5 | | 20220621 | |
| 994 | WT/AV | F | 5 | | 20220621 | |
| 994 | WT/AV | F | 5 | | | 60 |
| 994 | WT/AV | F | 9 | 1 | 20220719 | |
| 994 | WT/AV | F | 9 | 2 | 20220719 | |
| 994 | WT/AV | F | 9 | 3 | 20220719 | |
| 994 | WT/AV | F | 9 | 4 | 20220719 | |
| 994 | WT/AV | F | 9 | 5 | 20220719 | |
| 994 | WT/AV | F | 9 | 6 | 20220719 | |
| 994 | WT/AV | F | 9 | 7 | 20220719 | |
| 994 | WT/AV | F | 9 | 8 | 20220719 | |
| 994 | WT/AV | F | 9 | 9 | 20220719 | |
| 994 | WT/AV | F | 9 | 10 | 20220719 | |
| 994 | WT/AV | F | 9 | 11 | 20220719 | |
| 994 | WT/AV | F | 9 | 12 | 20220719 | |
| 994 | WT/AV | F | 9 | | | 60 |
| 994 | WT/AV | F | 13 | | 20220816 | |
| 994 | WT/AV | F | 13 | | 20220816 | |
| 994 | WT/AV | F | 13 | | 20220816 | |
| 994 | WT/AV | F | 13 | | 20220816 | |
| 994 | WT/AV | F | 13 | | 20220816 | |
| 994 | WT/AV | F | 13 | | 20220816 | |
| 994 | WT/AV | F | 13 | | 20220816 | |
| 994 | WT/AV | F | 13 | | 20220816 | |
| 994 | WT/AV | F | 13 | | 20220816 | |
| 994 | WT/AV | F | 13 | | 20220816 | |
| 994 | WT/AV | F | 13 | | 20220816 | |
| 994 | WT/AV | F | 13 | | 20220816 | |

| | | | | | |
|---|---|---|---|---|---|
| 994 | WT/AV | F | 13 | | 60 |
| 1163 | OI | F | 5 | 20220726 | |
| 1163 | OI | F | 5 | 20220726 | |
| 1163 | OI | F | 5 | 20220726 | |
| 1163 | OI | F | 5 | 20220726 | |
| 1163 | OI | F | 5 | 20220726 | |
| 1163 | OI | F | 5 | 20220726 | |
| 1163 | OI | F | 5 | 20220726 | |
| 1163 | OI | F | 5 | 20220726 | |
| 1163 | OI | F | 5 | 20220726 | |
| 1163 | OI | F | 5 | 20220726 | |
| 1163 | OI | F | 5 | 20220726 | |
| 1163 | OI | F | 5 | 20220726 | |
| 1163 | OI | F | 5 | | 60 |
| 1163 | OI | F | 9 | 20220823 | |
| 1163 | OI | F | 9 | 20220823 | |
| 1163 | OI | F | 9 | 20220823 | |
| 1163 | OI | F | 9 | 20220823 | |
| 1163 | OI | F | 9 | 20220823 | |
| 1163 | OI | F | 9 | 20220823 | |
| 1163 | OI | F | 9 | 20220823 | |
| 1163 | OI | F | 9 | 20220823 | |
| 1163 | OI | F | 9 | 20220823 | |
| 1163 | OI | F | 9 | 20220823 | |
| 1163 | OI | F | 9 | 20220823 | |
| 1163 | OI | F | 9 | 20220823 | |
| 1163 | OI | F | 9 | | 60 |
| 1163 | OI | F | 13 | 20220920 | |
| 1163 | OI | F | 13 | 20220920 | |
| 1163 | OI | F | 13 | 20220920 | |
| 1163 | OI | F | 13 | 20220920 | |
| 1163 | OI | F | 13 | 20220920 | |
| 1163 | OI | F | 13 | 20220920 | |
| 1163 | OI | F | 13 | 20220920 | |
| 1163 | OI | F | 13 | 20220920 | |
| 1163 | OI | F | 13 | 20220920 | |
| 1163 | OI | F | 13 | 20220920 | |
| 1163 | OI | F | 13 | 20220920 | |
| 1163 | OI | F | 13 | 20220920 | |
| 1163 | OI | F | 13 | | 60 |
| 1164 | WT/AV | F | 5 | 20220726 | |

| | | | | | |
|---|---|---|---|---|---|
| 1164 | WT/AV | F | 5 | 20220726 | |
| 1164 | WT/AV | F | 5 | 20220726 | |
| 1164 | WT/AV | F | 5 | 20220726 | |
| 1164 | WT/AV | F | 5 | 20220726 | |
| 1164 | WT/AV | F | 5 | 20220726 | |
| 1164 | WT/AV | F | 5 | 20220726 | |
| 1164 | WT/AV | F | 5 | 20220726 | |
| 1164 | WT/AV | F | 5 | 20220726 | |
| 1164 | WT/AV | F | 5 | 20220726 | |
| 1164 | WT/AV | F | 5 | 20220726 | |
| 1164 | WT/AV | F | 5 | 20220726 | |
| 1164 | WT/AV | F | 5 | | 60 |
| 1164 | WT/AV | F | 9 | 20220823 | |
| 1164 | WT/AV | F | 9 | 20220823 | |
| 1164 | WT/AV | F | 9 | 20220823 | |
| 1164 | WT/AV | F | 9 | 20220823 | |
| 1164 | WT/AV | F | 9 | 20220823 | |
| 1164 | WT/AV | F | 9 | 20220823 | |
| 1164 | WT/AV | F | 9 | 20220823 | |
| 1164 | WT/AV | F | 9 | 20220823 | |
| 1164 | WT/AV | F | 9 | 20220823 | |
| 1164 | WT/AV | F | 9 | 20220823 | |
| 1164 | WT/AV | F | 9 | 20220823 | |
| 1164 | WT/AV | F | 9 | 20220823 | |
| 1164 | WT/AV | F | 9 | | 60 |
| 1164 | WT/AV | F | 13 | 20220920 | |
| 1164 | WT/AV | F | 13 | 20220920 | |
| 1164 | WT/AV | F | 13 | 20220920 | |
| 1164 | WT/AV | F | 13 | 20220920 | |
| 1164 | WT/AV | F | 13 | 20220920 | |
| 1164 | WT/AV | F | 13 | 20220920 | |
| 1164 | WT/AV | F | 13 | 20220920 | |
| 1164 | WT/AV | F | 13 | 20220920 | |
| 1164 | WT/AV | F | 13 | 20220920 | |
| 1164 | WT/AV | F | 13 | 20220920 | |
| 1164 | WT/AV | F | 13 | 20220920 | |
| 1164 | WT/AV | F | 13 | 20220920 | |
| 1164 | WT/AV | F | 13 | | 60 |
| 1164 | WT/AV | F | 5 | 20220829 | |
| 1195 | WT/AV | F | 5 | 20220829 | |
| 1195 | WT/AV | F | 5 | 20220829 | |

| | | | | | | |
|---|---|---|---|---|---|---|
| 1195 | WT/AV | F | 5 | | 20220829 | |
| 1195 | WT/AV | F | 5 | | 20220829 | |
| 1195 | WT/AV | F | 5 | | 20220829 | |
| 1195 | WT/AV | F | 5 | | 20220829 | |
| 1195 | WT/AV | F | 5 | | 20220829 | |
| 1195 | WT/AV | F | 5 | | 20220829 | |
| 1195 | WT/AV | F | 5 | | 20220829 | |
| 1195 | WT/AV | F | 5 | | 20220829 | |
| 1195 | WT/AV | F | 5 | | 20220829 | |
| 1195 | WT/AV | F | 5 | | | 60 |
| 1195 | WT/AV | F | 9 | | 20220919 | |
| 1195 | WT/AV | F | 9 | | 20220919 | |
| 1195 | WT/AV | F | 9 | | 20220919 | |
| 1195 | WT/AV | F | 9 | | 20220919 | |
| 1195 | WT/AV | F | 9 | | 20220919 | |
| 1195 | WT/AV | F | 9 | | 20220919 | |
| 1195 | WT/AV | F | 9 | | 20220919 | |
| 1195 | WT/AV | F | 9 | | 20220919 | |
| 1195 | WT/AV | F | 9 | | 20220919 | |
| 1195 | WT/AV | F | 9 | | 20220919 | |
| 1195 | WT/AV | F | 9 | | 20220919 | |
| 1195 | WT/AV | F | 9 | | 20220919 | |
| 1195 | WT/AV | F | 9 | | | 60 |
| 1195 | WT/AV | F | 13 | 1 | 20221021 | |
| 1195 | WT/AV | F | 13 | 2 | 20221021 | |
| 1195 | WT/AV | F | 13 | 3 | 20221021 | |
| 1195 | WT/AV | F | 13 | 4 | 20221021 | |
| 1195 | WT/AV | F | 13 | 5 | 20221021 | |
| 1195 | WT/AV | F | 13 | 6 | 20221021 | |
| 1195 | WT/AV | F | 13 | 7 | 20221021 | |
| 1195 | WT/AV | F | 13 | 8 | 20221021 | |
| 1195 | WT/AV | F | 13 | 9 | 20221021 | |
| 1195 | WT/AV | F | 13 | 10 | 20221021 | |
| 1195 | WT/AV | F | 13 | 11 | 20221021 | |
| 1195 | WT/AV | F | 13 | 12 | 20221021 | |
| 1195 | WT/AV | F | 13 | | | 60 |
| 1199 | WT | M | 5 | | 20220829 | |
| 1199 | WT | M | 5 | | 20220829 | |
| 1199 | WT | M | 5 | | 20220829 | |
| 1199 | WT | M | 5 | | 20220829 | |
| 1199 | WT | M | 5 | | 20220829 | |

| | | | | | | |
|---|---|---|---|---|---|---|
| 1199 | WT | M | 5 | | 20220829 | |
| 1199 | WT | M | 5 | | 20220829 | |
| 1199 | WT | M | 5 | | 20220829 | |
| 1199 | WT | M | 5 | | 20220829 | |
| 1199 | WT | M | 5 | | 20220829 | |
| 1199 | WT | M | 5 | | 20220829 | |
| 1199 | WT | M | 5 | | 20220829 | |
| 1199 | WT | M | 5 | | | 60 |
| 1199 | WT | M | 9 | | 20220919 | |
| 1199 | WT | M | 9 | | 20220919 | |
| 1199 | WT | M | 9 | | 20220919 | |
| 1199 | WT | M | 9 | | 20220919 | |
| 1199 | WT | M | 9 | | 20220919 | |
| 1199 | WT | M | 9 | | 20220919 | |
| 1199 | WT | M | 9 | | 20220919 | |
| 1199 | WT | M | 9 | | 20220919 | |
| 1199 | WT | M | 9 | | 20220919 | |
| 1199 | WT | M | 9 | | 20220919 | |
| 1199 | WT | M | 9 | | 20220919 | |
| 1199 | WT | M | 9 | | 20220919 | |
| 1199 | WT | M | 9 | | | 60 |
| 1199 | WT | M | 13 | 1 | 20221021 | |
| 1199 | WT | M | 13 | 2 | 20221021 | |
| 1199 | WT | M | 13 | 3 | 20221021 | |
| 1199 | WT | M | 13 | 4 | 20221021 | |
| 1199 | WT | M | 13 | 5 | 20221021 | |
| 1199 | WT | M | 13 | 6 | 20221021 | |
| 1199 | WT | M | 13 | 7 | 20221021 | |
| 1199 | WT | M | 13 | 8 | 20221021 | |
| 1199 | WT | M | 13 | 9 | 20221021 | |
| 1199 | WT | M | 13 | 10 | 20221021 | |
| 1199 | WT | M | 13 | 11 | 20221021 | |
| 1199 | WT | M | 13 | 12 | 20221021 | |
| 1199 | WT | M | 13 | | | 60 |
| 1210 | OI | M | 5 | | 20220823 | |
| 1210 | OI | M | 5 | | 20220823 | |
| 1210 | OI | M | 5 | | 20220823 | |
| 1210 | OI | M | 5 | | 20220823 | |
| 1210 | OI | M | 5 | | 20220823 | |
| 1210 | OI | M | 5 | | 20220823 | |
| 1210 | OI | M | 5 | | 20220823 | |

| | | | | | |
|---|---|---|---|---|---|
| 1210 | OI | M | 5 | | 20220823 | |
| 1210 | OI | M | 5 | | 20220823 | |
| 1210 | OI | M | 5 | | 20220823 | |
| 1210 | OI | M | 5 | | 20220823 | |
| 1210 | OI | M | 5 | | 20220823 | |
| 1210 | OI | M | 5 | | | 60 |
| 1210 | OI | M | 9 | | 20220920 | |
| 1210 | OI | M | 9 | | 20220920 | |
| 1210 | OI | M | 9 | | 20220920 | |
| 1210 | OI | M | 9 | | 20220920 | |
| 1210 | OI | M | 9 | | 20220920 | |
| 1210 | OI | M | 9 | | 20220920 | |
| 1210 | OI | M | 9 | | 20220920 | |
| 1210 | OI | M | 9 | | 20220920 | |
| 1210 | OI | M | 9 | | 20220920 | |
| 1210 | OI | M | 9 | | 20220920 | |
| 1210 | OI | M | 9 | | 20220920 | |
| 1210 | OI | M | 9 | | 20220920 | |
| 1210 | OI | M | 9 | | | 60 |
| 1210 | OI | M | 13 | 1 | 20221017 | |
| 1210 | OI | M | 13 | 2 | 20221017 | |
| 1210 | OI | M | 13 | 3 | 20221017 | |
| 1210 | OI | M | 13 | 4 | 20221017 | |
| 1210 | OI | M | 13 | 5 | 20221017 | |
| 1210 | OI | M | 13 | 6 | 20221017 | |
| 1210 | OI | M | 13 | 7 | 20221017 | |
| 1210 | OI | M | 13 | 8 | 20221017 | |
| 1210 | OI | M | 13 | 9 | 20221017 | |
| 1210 | OI | M | 13 | 10 | 20221017 | |
| 1210 | OI | M | 13 | 11 | 20221017 | |
| 1210 | OI | M | 13 | 12 | 20221017 | |
| 1210 | OI | M | 13 | | | 60 |
| 1211 | OI | M | 5 | | 20220823 | |
| 1211 | OI | M | 5 | | 20220823 | |
| 1211 | OI | M | 5 | | 20220823 | |
| 1211 | OI | M | 5 | | 20220823 | |
| 1211 | OI | M | 5 | | 20220823 | |
| 1211 | OI | M | 5 | | 20220823 | |
| 1211 | OI | M | 5 | | 20220823 | |
| 1211 | OI | M | 5 | | 20220823 | |
| 1211 | OI | M | 5 | | 20220823 | |

| | | | | | |
|---|---|---|---|---|---|
| 1211 OI | M | 5 | | 20220823 | |
| 1211 OI | M | 5 | | 20220823 | |
| 1211 OI | M | 5 | | 20220823 | |
| 1211 OI | M | 5 | | | 60 |
| 1211 OI | M | 9 | | 20220920 | |
| 1211 OI | M | 9 | | 20220920 | |
| 1211 OI | M | 9 | | 20220920 | |
| 1211 OI | M | 9 | | 20220920 | |
| 1211 OI | M | 9 | | 20220920 | |
| 1211 OI | M | 9 | | 20220920 | |
| 1211 OI | M | 9 | | 20220920 | |
| 1211 OI | M | 9 | | 20220920 | |
| 1211 OI | M | 9 | | 20220920 | |
| 1211 OI | M | 9 | | 20220920 | |
| 1211 OI | M | 9 | | 20220920 | |
| 1211 OI | M | 9 | | 20220920 | |
| 1211 OI | M | 9 | | | 60 |
| 1211 OI | M | 13 | 1 | 20221017 | |
| 1211 OI | M | 13 | 2 | 20221017 | |
| 1211 OI | M | 13 | 3 | 20221017 | |
| 1211 OI | M | 13 | 4 | 20221017 | |
| 1211 OI | M | 13 | 5 | 20221017 | |
| 1211 OI | M | 13 | 6 | 20221017 | |
| 1211 OI | M | 13 | 7 | 20221017 | |
| 1211 OI | M | 13 | 8 | 20221017 | |
| 1211 OI | M | 13 | 9 | 20221017 | |
| 1211 OI | M | 13 | 10 | 20221017 | |
| 1211 OI | M | 13 | 11 | 20221017 | |
| 1211 OI | M | 13 | 12 | 20221017 | |
| 1211 OI | M | 13 | | | 60 |
| 1212 WT | M | 5 | | 20220823 | |
| 1212 WT | M | 5 | | 20220823 | |
| 1212 WT | M | 5 | | 20220823 | |
| 1212 WT | M | 5 | | 20220823 | |
| 1212 WT | M | 5 | | 20220823 | |
| 1212 WT | M | 5 | | 20220823 | |
| 1212 WT | M | 5 | | 20220823 | |
| 1212 WT | M | 5 | | 20220823 | |
| 1212 WT | M | 5 | | 20220823 | |
| 1212 WT | M | 5 | | 20220823 | |
| 1212 WT | M | 5 | | 20220823 | |

| | | | | | | |
|---|---|---|---|---|---|---|
| 1212 | WT | M | 5 | | 20220823 | |
| 1212 | WT | M | 5 | | | 60 |
| 1212 | WT | M | 9 | | 20220920 | |
| 1212 | WT | M | 9 | | 20220920 | |
| 1212 | WT | M | 9 | | 20220920 | |
| 1212 | WT | M | 9 | | 20220920 | |
| 1212 | WT | M | 9 | | 20220920 | |
| 1212 | WT | M | 9 | | 20220920 | |
| 1212 | WT | M | 9 | | 20220920 | |
| 1212 | WT | M | 9 | | 20220920 | |
| 1212 | WT | M | 9 | | 20220920 | |
| 1212 | WT | M | 9 | | 20220920 | |
| 1212 | WT | M | 9 | | 20220920 | |
| 1212 | WT | M | 9 | | 20220920 | |
| 1212 | WT | M | 9 | | | 60 |
| 1212 | WT | M | 13 | 1 | 20221017 | |
| 1212 | WT | M | 13 | 2 | 20221017 | |
| 1212 | WT | M | 13 | 3 | 20221017 | |
| 1212 | WT | M | 13 | 4 | 20221017 | |
| 1212 | WT | M | 13 | 5 | 20221017 | |
| 1212 | WT | M | 13 | 6 | 20221017 | |
| 1212 | WT | M | 13 | 7 | 20221017 | |
| 1212 | WT | M | 13 | 8 | 20221017 | |
| 1212 | WT | M | 13 | 9 | 20221017 | |
| 1212 | WT | M | 13 | 10 | 20221017 | |
| 1212 | WT | M | 13 | 11 | 20221017 | |
| 1212 | WT | M | 13 | 12 | 20221017 | |
| 1212 | WT | M | 13 | | | 60 |
| 1214 | OI | M | 5 | | 20220823 | |
| 1214 | OI | M | 5 | | 20220823 | |
| 1214 | OI | M | 5 | | 20220823 | |
| 1214 | OI | M | 5 | | 20220823 | |
| 1214 | OI | M | 5 | | 20220823 | |
| 1214 | OI | M | 5 | | 20220823 | |
| 1214 | OI | M | 5 | | 20220823 | |
| 1214 | OI | M | 5 | | 20220823 | |
| 1214 | OI | M | 5 | | 20220823 | |
| 1214 | OI | M | 5 | | 20220823 | |
| 1214 | OI | M | 5 | | 20220823 | |
| 1214 | OI | M | 5 | | 20220823 | |
| 1214 | OI | M | 5 | | | 60 |

| | | | | | | |
|---|---|---|---|---|---|---|
| 1214 | OI | M | 9 | | 20220920 | |
| 1214 | OI | M | 9 | | 20220920 | |
| 1214 | OI | M | 9 | | 20220920 | |
| 1214 | OI | M | 9 | | 20220920 | |
| 1214 | OI | M | 9 | | 20220920 | |
| 1214 | OI | M | 9 | | 20220920 | |
| 1214 | OI | M | 9 | | 20220920 | |
| 1214 | OI | M | 9 | | 20220920 | |
| 1214 | OI | M | 9 | | 20220920 | |
| 1214 | OI | M | 9 | | 20220920 | |
| 1214 | OI | M | 9 | | 20220920 | |
| 1214 | OI | M | 9 | | 20220920 | |
| 1214 | OI | M | 9 | | | 60 |
| 1214 | OI | M | 13 | 1 | 20221017 | |
| 1214 | OI | M | 13 | 2 | 20221017 | |
| 1214 | OI | M | 13 | 3 | 20221017 | |
| 1214 | OI | M | 13 | 4 | 20221017 | |
| 1214 | OI | M | 13 | 5 | 20221017 | |
| 1214 | OI | M | 13 | 6 | 20221017 | |
| 1214 | OI | M | 13 | 7 | 20221017 | |
| 1214 | OI | M | 13 | 8 | 20221017 | |
| 1214 | OI | M | 13 | 9 | 20221017 | |
| 1214 | OI | M | 13 | 10 | 20221017 | |
| 1214 | OI | M | 13 | 11 | 20221017 | |
| 1214 | OI | M | 13 | 12 | 20221017 | |
| 1214 | OI | M | 13 | | | 60 |
| 1652 | WT | F | 5 | 1 | 20210405 | |
| 1652 | WT | F | 5 | 2 | 20210405 | |
| 1652 | WT | F | 5 | 3 | 20210405 | |
| 1652 | WT | F | 5 | 4 | 20210405 | |
| 1652 | WT | F | 5 | 5 | 20210405 | |
| 1652 | WT | F | 5 | 6 | 20210405 | |
| 1652 | WT | F | 5 | 7 | 20210405 | |
| 1652 | WT | F | 5 | 8 | 20210405 | |
| 1652 | WT | F | 5 | 9 | 20210405 | |
| 1652 | WT | F | 5 | 10 | 20210405 | |
| 1652 | WT | F | 5 | 11 | 20210405 | |
| 1652 | WT | F | 5 | 12 | 20210405 | |
| 1652 | WT | F | 5 | | | 60 |
| 1652 | WT | F | 9 | 1 | 20210505 | |
| 1652 | WT | F | 9 | 2 | 20210505 | |

| | | | | | | |
|---|---|---|---|---|---|---|
| 1652 | WT | F | 9 | 3 | 20210505 | |
| 1652 | WT | F | 9 | 4 | 20210505 | |
| 1652 | WT | F | 9 | 5 | 20210505 | |
| 1652 | WT | F | 9 | 6 | 20210505 | |
| 1652 | WT | F | 9 | 7 | 20210505 | |
| 1652 | WT | F | 9 | 8 | 20210505 | |
| 1652 | WT | F | 9 | 9 | 20210505 | |
| 1652 | WT | F | 9 | 10 | 20210505 | |
| 1652 | WT | F | 9 | 11 | 20210505 | |
| 1652 | WT | F | 9 | 12 | 20210505 | |
| 1652 | WT | F | 9 | | | 60 |
| 1652 | WT | F | 13 | 1 | 20210531 | |
| 1652 | WT | F | 13 | 2 | 20210531 | |
| 1652 | WT | F | 13 | 3 | 20210531 | |
| 1652 | WT | F | 13 | 4 | 20210531 | |
| 1652 | WT | F | 13 | 5 | 20210531 | |
| 1652 | WT | F | 13 | 6 | 20210531 | |
| 1652 | WT | F | 13 | 7 | 20210531 | |
| 1652 | WT | F | 13 | 8 | 20210531 | |
| 1652 | WT | F | 13 | 9 | 20210531 | |
| 1652 | WT | F | 13 | 10 | 20210531 | |
| 1652 | WT | F | 13 | 11 | 20210531 | |
| 1652 | WT | F | 13 | 12 | 20210531 | |
| 1652 | WT | F | 13 | | | 60 |
| 1654 | WT | F | 5 | 1 | 20210405 | |
| 1654 | WT | F | 5 | 2 | 20210405 | |
| 1654 | WT | F | 5 | 3 | 20210405 | |
| 1654 | WT | F | 5 | 4 | 20210405 | |
| 1654 | WT | F | 5 | 5 | 20210405 | |
| 1654 | WT | F | 5 | 6 | 20210405 | |
| 1654 | WT | F | 5 | 7 | 20210405 | |
| 1654 | WT | F | 5 | 8 | 20210405 | |
| 1654 | WT | F | 5 | 9 | 20210405 | |
| 1654 | WT | F | 5 | 10 | 20210405 | |
| 1654 | WT | F | 5 | 11 | 20210405 | |
| 1654 | WT | F | 5 | 12 | 20210405 | |
| 1654 | WT | F | 5 | | | 60 |
| 1654 | WT | F | 9 | 1 | 20210505 | |
| 1654 | WT | F | 9 | 2 | 20210505 | |
| 1654 | WT | F | 9 | 3 | 20210505 | |
| 1654 | WT | F | 9 | 4 | 20210505 | |

| | | | | | | |
|---|---|---|---|---|---|---|
| 1654 | WT | F | 9 | 5 | 20210505 | |
| 1654 | WT | F | 9 | 6 | 20210505 | |
| 1654 | WT | F | 9 | 7 | 20210505 | |
| 1654 | WT | F | 9 | 8 | 20210505 | |
| 1654 | WT | F | 9 | 9 | 20210505 | |
| 1654 | WT | F | 9 | 10 | 20210505 | |
| 1654 | WT | F | 9 | 11 | 20210505 | |
| 1654 | WT | F | 9 | 12 | 20210505 | |
| 1654 | WT | F | 9 | | | 60 |
| 1654 | WT | F | 13 | 1 | 20210531 | |
| 1654 | WT | F | 13 | 2 | 20210531 | |
| 1654 | WT | F | 13 | 3 | 20210531 | |
| 1654 | WT | F | 13 | 4 | 20210531 | |
| 1654 | WT | F | 13 | 5 | 20210531 | |
| 1654 | WT | F | 13 | 6 | 20210531 | |
| 1654 | WT | F | 13 | 7 | 20210531 | |
| 1654 | WT | F | 13 | 8 | 20210531 | |
| 1654 | WT | F | 13 | 9 | 20210531 | |
| 1654 | WT | F | 13 | 10 | 20210531 | |
| 1654 | WT | F | 13 | 11 | 20210531 | |
| 1654 | WT | F | 13 | 12 | 20210531 | |
| 1654 | WT | F | 13 | | | 60 |
| 1909 | OI/AV | F | 5 | 1 | 20210405 | |
| 1909 | OI/AV | F | 5 | 2 | 20210405 | |
| 1909 | OI/AV | F | 5 | 3 | 20210405 | |
| 1909 | OI/AV | F | 5 | 4 | 20210405 | |
| 1909 | OI/AV | F | 5 | 5 | 20210405 | |
| 1909 | OI/AV | F | 5 | 6 | 20210405 | |
| 1909 | OI/AV | F | 5 | 7 | 20210405 | |
| 1909 | OI/AV | F | 5 | 8 | 20210405 | |
| 1909 | OI/AV | F | 5 | 9 | 20210405 | |
| 1909 | OI/AV | F | 5 | 10 | 20210405 | |
| 1909 | OI/AV | F | 5 | 11 | 20210405 | |
| 1909 | OI/AV | F | 5 | 12 | 20210405 | |
| 1909 | OI/AV | F | 5 | | | 60 |
| 1909 | OI/AV | F | 9 | 1 | 20210505 | |
| 1909 | OI/AV | F | 9 | 2 | 20210505 | |
| 1909 | OI/AV | F | 9 | 3 | 20210505 | |
| 1909 | OI/AV | F | 9 | 4 | 20210505 | |
| 1909 | OI/AV | F | 9 | 5 | 20210505 | |
| 1909 | OI/AV | F | 9 | 6 | 20210505 | |

| | | | | | |
|---|---|---|---|---|---|
| 1909 | OI/AV | F | 9 | 7 | 20210505 | |
| 1909 | OI/AV | F | 9 | 8 | 20210505 | |
| 1909 | OI/AV | F | 9 | 9 | 20210505 | |
| 1909 | OI/AV | F | 9 | 10 | 20210505 | |
| 1909 | OI/AV | F | 9 | 11 | 20210505 | |
| 1909 | OI/AV | F | 9 | 12 | 20210505 | |
| 1909 | OI/AV | F | 9 | | | 60 |
| 1909 | OI/AV | F | 13 | 1 | 20210531 | |
| 1909 | OI/AV | F | 13 | 2 | 20210531 | |
| 1909 | OI/AV | F | 13 | 3 | 20210531 | |
| 1909 | OI/AV | F | 13 | 4 | 20210531 | |
| 1909 | OI/AV | F | 13 | 5 | 20210531 | |
| 1909 | OI/AV | F | 13 | 6 | 20210531 | |
| 1909 | OI/AV | F | 13 | 7 | 20210531 | |
| 1909 | OI/AV | F | 13 | 8 | 20210531 | |
| 1909 | OI/AV | F | 13 | 9 | 20210531 | |
| 1909 | OI/AV | F | 13 | 10 | 20210531 | |
| 1909 | OI/AV | F | 13 | 11 | 20210531 | |
| 1909 | OI/AV | F | 13 | 12 | 20210531 | |
| 1909 | OI/AV | F | 13 | | | 60 |
| 1910 | OI | F | 5 | 1 | 20210405 | |
| 1910 | OI | F | 5 | 2 | 20210405 | |
| 1910 | OI | F | 5 | 3 | 20210405 | |
| 1910 | OI | F | 5 | 4 | 20210405 | |
| 1910 | OI | F | 5 | 5 | 20210405 | |
| 1910 | OI | F | 5 | 6 | 20210405 | |
| 1910 | OI | F | 5 | 7 | 20210405 | |
| 1910 | OI | F | 5 | 8 | 20210405 | |
| 1910 | OI | F | 5 | 9 | 20210405 | |
| 1910 | OI | F | 5 | 10 | 20210405 | |
| 1910 | OI | F | 5 | 11 | 20210405 | |
| 1910 | OI | F | 5 | 12 | 20210405 | |
| 1910 | OI | F | 5 | | | 60 |
| 1910 | OI | F | 9 | 1 | 20210505 | |
| 1910 | OI | F | 9 | 2 | 20210505 | |
| 1910 | OI | F | 9 | 3 | 20210505 | |
| 1910 | OI | F | 9 | 4 | 20210505 | |
| 1910 | OI | F | 9 | 5 | 20210505 | |
| 1910 | OI | F | 9 | 6 | 20210505 | |
| 1910 | OI | F | 9 | 7 | 20210505 | |
| 1910 | OI | F | 9 | 8 | 20210505 | |

| | | | | | |
|---|---|---|---|---|---|
| 1910 OI | F | 9 | 9 | 20210505 | |
| 1910 OI | F | 9 | 10 | 20210505 | |
| 1910 OI | F | 9 | 11 | 20210505 | |
| 1910 OI | F | 9 | 12 | 20210505 | |
| 1910 OI | F | 9 | | | 60 |
| 1910 OI | F | 13 | 1 | 20210531 | |
| 1910 OI | F | 13 | 2 | 20210531 | |
| 1910 OI | F | 13 | 3 | 20210531 | |
| 1910 OI | F | 13 | 4 | 20210531 | |
| 1910 OI | F | 13 | 5 | 20210531 | |
| 1910 OI | F | 13 | 6 | 20210531 | |
| 1910 OI | F | 13 | 7 | 20210531 | |
| 1910 OI | F | 13 | 8 | 20210531 | |
| 1910 OI | F | 13 | 9 | 20210531 | |
| 1910 OI | F | 13 | 10 | 20210531 | |
| 1910 OI | F | 13 | 11 | 20210531 | |
| 1910 OI | F | 13 | 12 | 20210531 | |
| 1910 OI | F | 13 | | | 60 |
| 1930 OI/AV | F | 5 | 1 | 20210426 | |
| 1930 OI/AV | F | 5 | 2 | 20210426 | |
| 1930 OI/AV | F | 5 | 3 | 20210426 | |
| 1930 OI/AV | F | 5 | 4 | 20210426 | |
| 1930 OI/AV | F | 5 | 5 | 20210426 | |
| 1930 OI/AV | F | 5 | 6 | 20210426 | |
| 1930 OI/AV | F | 5 | 7 | 20210426 | |
| 1930 OI/AV | F | 5 | 8 | 20210426 | |
| 1930 OI/AV | F | 5 | 9 | 20210426 | |
| 1930 OI/AV | F | 5 | 10 | 20210426 | |
| 1930 OI/AV | F | 5 | 11 | 20210426 | |
| 1930 OI/AV | F | 5 | 12 | 20210426 | |
| 1930 OI/AV | F | 5 | | | 60 |
| 1930 OI/AV | F | 9 | 1 | 20210524 | |
| 1930 OI/AV | F | 9 | 2 | 20210524 | |
| 1930 OI/AV | F | 9 | 3 | 20210524 | |
| 1930 OI/AV | F | 9 | 4 | 20210524 | |
| 1930 OI/AV | F | 9 | 5 | 20210524 | |
| 1930 OI/AV | F | 9 | 6 | 20210524 | |
| 1930 OI/AV | F | 9 | 7 | 20210524 | |
| 1930 OI/AV | F | 9 | 8 | 20210524 | |
| 1930 OI/AV | F | 9 | 9 | 20210524 | |
| 1930 OI/AV | F | 9 | 10 | 20210524 | |

| | | | | | |
|---|---|---|---|---|---|
| 1930 OI/AV | F | 9 | 11 | 20210524 | |
| 1930 OI/AV | F | 9 | 12 | 20210524 | |
| 1930 OI/AV | F | 9 | | | 60 |
| 1930 OI/AV | F | 13 | 1 | 20210621 | |
| 1930 OI/AV | F | 13 | 2 | 20210621 | |
| 1930 OI/AV | F | 13 | 3 | 20210621 | |
| 1930 OI/AV | F | 13 | 4 | 20210621 | |
| 1930 OI/AV | F | 13 | 5 | 20210621 | |
| 1930 OI/AV | F | 13 | 6 | 20210621 | |
| 1930 OI/AV | F | 13 | 7 | 20210621 | |
| 1930 OI/AV | F | 13 | 8 | 20210621 | |
| 1930 OI/AV | F | 13 | 9 | 20210621 | |
| 1930 OI/AV | F | 13 | 10 | 20210621 | |
| 1930 OI/AV | F | 13 | 11 | 20210621 | |
| 1930 OI/AV | F | 13 | 12 | 20210621 | |
| 1930 OI/AV | F | 13 | | | 60 |
| 1931 WT | F | 5 | 1 | 20210426 | |
| 1931 WT | F | 5 | 2 | 20210426 | |
| 1931 WT | F | 5 | 3 | 20210426 | |
| 1931 WT | F | 5 | 4 | 20210426 | |
| 1931 WT | F | 5 | 5 | 20210426 | |
| 1931 WT | F | 5 | 6 | 20210426 | |
| 1931 WT | F | 5 | 7 | 20210426 | |
| 1931 WT | F | 5 | 8 | 20210426 | |
| 1931 WT | F | 5 | 9 | 20210426 | |
| 1931 WT | F | 5 | 10 | 20210426 | |
| 1931 WT | F | 5 | 11 | 20210426 | |
| 1931 WT | F | 5 | 12 | 20210426 | |
| 1931 WT | F | 5 | | | 60 |
| 1931 WT | F | 9 | 1 | 20210524 | |
| 1931 WT | F | 9 | 2 | 20210524 | |
| 1931 WT | F | 9 | 3 | 20210524 | |
| 1931 WT | F | 9 | 4 | 20210524 | |
| 1931 WT | F | 9 | 5 | 20210524 | |
| 1931 WT | F | 9 | 6 | 20210524 | |
| 1931 WT | F | 9 | 7 | 20210524 | |
| 1931 WT | F | 9 | 8 | 20210524 | |
| 1931 WT | F | 9 | 9 | 20210524 | |
| 1931 WT | F | 9 | 10 | 20210524 | |
| 1931 WT | F | 9 | 11 | 20210524 | |
| 1931 WT | F | 9 | 12 | 20210524 | |

| | | | | | | |
|---|---|---|---|---|---|---|
| 1931 | WT | F | 9 | | | 60 |
| 1931 | WT | F | 13 | 1 | 20210621 | |
| 1931 | WT | F | 13 | 2 | 20210621 | |
| 1931 | WT | F | 13 | 3 | 20210621 | |
| 1931 | WT | F | 13 | 4 | 20210621 | |
| 1931 | WT | F | 13 | 5 | 20210621 | |
| 1931 | WT | F | 13 | 6 | 20210621 | |
| 1931 | WT | F | 13 | 7 | 20210621 | |
| 1931 | WT | F | 13 | 8 | 20210621 | |
| 1931 | WT | F | 13 | 9 | 20210621 | |
| 1931 | WT | F | 13 | 10 | 20210621 | |
| 1931 | WT | F | 13 | 11 | 20210621 | |
| 1931 | WT | F | 13 | 12 | 20210621 | |
| 1931 | WT | F | 13 | | | 60 |
| 1932 | WT | F | 5 | 1 | 20210426 | |
| 1932 | WT | F | 5 | 2 | 20210426 | |
| 1932 | WT | F | 5 | 3 | 20210426 | |
| 1932 | WT | F | 5 | 4 | 20210426 | |
| 1932 | WT | F | 5 | 5 | 20210426 | |
| 1932 | WT | F | 5 | 6 | 20210426 | |
| 1932 | WT | F | 5 | 7 | 20210426 | |
| 1932 | WT | F | 5 | 8 | 20210426 | |
| 1932 | WT | F | 5 | 9 | 20210426 | |
| 1932 | WT | F | 5 | 10 | 20210426 | |
| 1932 | WT | F | 5 | 11 | 20210426 | |
| 1932 | WT | F | 5 | 12 | 20210426 | |
| 1932 | WT | F | 5 | | | 60 |
| 1932 | WT | F | 9 | 1 | 20210524 | |
| 1932 | WT | F | 9 | 2 | 20210524 | |
| 1932 | WT | F | 9 | 3 | 20210524 | |
| 1932 | WT | F | 9 | 4 | 20210524 | |
| 1932 | WT | F | 9 | 5 | 20210524 | |
| 1932 | WT | F | 9 | 6 | 20210524 | |
| 1932 | WT | F | 9 | 7 | 20210524 | |
| 1932 | WT | F | 9 | 8 | 20210524 | |
| 1932 | WT | F | 9 | 9 | 20210524 | |
| 1932 | WT | F | 9 | 10 | 20210524 | |
| 1932 | WT | F | 9 | 11 | 20210524 | |
| 1932 | WT | F | 9 | 12 | 20210524 | |
| 1932 | WT | F | 9 | | | 60 |
| 1932 | WT | F | 13 | 1 | 20210621 | |

| | | | | | | |
|---|---|---|---|---|---|---|
| 1932 | WT | F | 13 | 2 | 20210621 | |
| 1932 | WT | F | 13 | 3 | 20210621 | |
| 1932 | WT | F | 13 | 4 | 20210621 | |
| 1932 | WT | F | 13 | 5 | 20210621 | |
| 1932 | WT | F | 13 | 6 | 20210621 | |
| 1932 | WT | F | 13 | 7 | 20210621 | |
| 1932 | WT | F | 13 | 8 | 20210621 | |
| 1932 | WT | F | 13 | 9 | 20210621 | |
| 1932 | WT | F | 13 | 10 | 20210621 | |
| 1932 | WT | F | 13 | 11 | 20210621 | |
| 1932 | WT | F | 13 | 12 | 20210621 | |
| 1932 | WT | F | 13 | | | 60 |
| 1933 | OI/AV | F | 5 | 1 | 20210426 | |
| 1933 | OI/AV | F | 5 | 2 | 20210426 | |
| 1933 | OI/AV | F | 5 | 3 | 20210426 | |
| 1933 | OI/AV | F | 5 | 4 | 20210426 | |
| 1933 | OI/AV | F | 5 | 5 | 20210426 | |
| 1933 | OI/AV | F | 5 | 6 | 20210426 | |
| 1933 | OI/AV | F | 5 | 7 | 20210426 | |
| 1933 | OI/AV | F | 5 | 8 | 20210426 | |
| 1933 | OI/AV | F | 5 | 9 | 20210426 | |
| 1933 | OI/AV | F | 5 | 10 | 20210426 | |
| 1933 | OI/AV | F | 5 | 11 | 20210426 | |
| 1933 | OI/AV | F | 5 | 12 | 20210426 | |
| 1933 | OI/AV | F | 5 | | | 60 |
| 1933 | OI/AV | F | 9 | 1 | 20210524 | |
| 1933 | OI/AV | F | 9 | 2 | 20210524 | |
| 1933 | OI/AV | F | 9 | 3 | 20210524 | |
| 1933 | OI/AV | F | 9 | 4 | 20210524 | |
| 1933 | OI/AV | F | 9 | 5 | 20210524 | |
| 1933 | OI/AV | F | 9 | 6 | 20210524 | |
| 1933 | OI/AV | F | 9 | 7 | 20210524 | |
| 1933 | OI/AV | F | 9 | 8 | 20210524 | |
| 1933 | OI/AV | F | 9 | 9 | 20210524 | |
| 1933 | OI/AV | F | 9 | 10 | 20210524 | |
| 1933 | OI/AV | F | 9 | 11 | 20210524 | |
| 1933 | OI/AV | F | 9 | 12 | 20210524 | |
| 1933 | OI/AV | F | 9 | | | 60 |
| 1933 | OI/AV | F | 13 | 1 | 20210621 | |
| 1933 | OI/AV | F | 13 | 2 | 20210621 | |
| 1933 | OI/AV | F | 13 | 3 | 20210621 | |

| | | | | | | |
|---|---|---|---|---|---|---|
| 1933 | OI/AV | F | 13 | 4 | 20210621 | |
| 1933 | OI/AV | F | 13 | 5 | 20210621 | |
| 1933 | OI/AV | F | 13 | 6 | 20210621 | |
| 1933 | OI/AV | F | 13 | 7 | 20210621 | |
| 1933 | OI/AV | F | 13 | 8 | 20210621 | |
| 1933 | OI/AV | F | 13 | 9 | 20210621 | |
| 1933 | OI/AV | F | 13 | 10 | 20210621 | |
| 1933 | OI/AV | F | 13 | 11 | 20210621 | |
| 1933 | OI/AV | F | 13 | 12 | 20210621 | |
| 1933 | OI/AV | F | 13 | | | 60 |
| 1934 | OI/AV | M | 5 | 1 | 20210426 | |
| 1934 | OI/AV | M | 5 | 2 | 20210426 | |
| 1934 | OI/AV | M | 5 | 3 | 20210426 | |
| 1934 | OI/AV | M | 5 | 4 | 20210426 | |
| 1934 | OI/AV | M | 5 | 5 | 20210426 | |
| 1934 | OI/AV | M | 5 | 6 | 20210426 | |
| 1934 | OI/AV | M | 5 | 7 | 20210426 | |
| 1934 | OI/AV | M | 5 | 8 | 20210426 | |
| 1934 | OI/AV | M | 5 | 9 | 20210426 | |
| 1934 | OI/AV | M | 5 | 10 | 20210426 | |
| 1934 | OI/AV | M | 5 | 11 | 20210426 | |
| 1934 | OI/AV | M | 5 | 12 | 20210426 | |
| 1934 | OI/AV | M | 5 | | | 60 |
| 1934 | OI/AV | M | 9 | 1 | 20210524 | |
| 1934 | OI/AV | M | 9 | 2 | 20210524 | |
| 1934 | OI/AV | M | 9 | 3 | 20210524 | |
| 1934 | OI/AV | M | 9 | 4 | 20210524 | |
| 1934 | OI/AV | M | 9 | 5 | 20210524 | |
| 1934 | OI/AV | M | 9 | 6 | 20210524 | |
| 1934 | OI/AV | M | 9 | 7 | 20210524 | |
| 1934 | OI/AV | M | 9 | 8 | 20210524 | |
| 1934 | OI/AV | M | 9 | 9 | 20210524 | |
| 1934 | OI/AV | M | 9 | 10 | 20210524 | |
| 1934 | OI/AV | M | 9 | 11 | 20210524 | |
| 1934 | OI/AV | M | 9 | 12 | 20210524 | |
| 1934 | OI/AV | M | 9 | | | 60 |
| 1934 | OI/AV | M | 13 | 1 | 20210621 | |
| 1934 | OI/AV | M | 13 | 2 | 20210621 | |
| 1934 | OI/AV | M | 13 | 3 | 20210621 | |
| 1934 | OI/AV | M | 13 | 4 | 20210621 | |
| 1934 | OI/AV | M | 13 | 5 | 20210621 | |

| | | | | | | |
|---|---|---|---|---|---|---|
| 1934 | OI/AV | M | 13 | 6 | 20210621 | |
| 1934 | OI/AV | M | 13 | 7 | 20210621 | |
| 1934 | OI/AV | M | 13 | 8 | 20210621 | |
| 1934 | OI/AV | M | 13 | 9 | 20210621 | |
| 1934 | OI/AV | M | 13 | 10 | 20210621 | |
| 1934 | OI/AV | M | 13 | 11 | 20210621 | |
| 1934 | OI/AV | M | 13 | 12 | 20210621 | |
| 1934 | OI/AV | M | 13 | | | 60 |
| 1935 | OI/AV | M | 5 | 1 | 20210426 | |
| 1935 | OI/AV | M | 5 | 2 | 20210426 | |
| 1935 | OI/AV | M | 5 | 3 | 20210426 | |
| 1935 | OI/AV | M | 5 | 4 | 20210426 | |
| 1935 | OI/AV | M | 5 | 5 | 20210426 | |
| 1935 | OI/AV | M | 5 | 6 | 20210426 | |
| 1935 | OI/AV | M | 5 | 7 | 20210426 | |
| 1935 | OI/AV | M | 5 | 8 | 20210426 | |
| 1935 | OI/AV | M | 5 | 9 | 20210426 | |
| 1935 | OI/AV | M | 5 | 10 | 20210426 | |
| 1935 | OI/AV | M | 5 | 11 | 20210426 | |
| 1935 | OI/AV | M | 5 | 12 | 20210426 | |
| 1935 | OI/AV | M | 5 | | | 60 |
| 1935 | OI/AV | M | 9 | 1 | 20210524 | |
| 1935 | OI/AV | M | 9 | 2 | 20210524 | |
| 1935 | OI/AV | M | 9 | 3 | 20210524 | |
| 1935 | OI/AV | M | 9 | 4 | 20210524 | |
| 1935 | OI/AV | M | 9 | 5 | 20210524 | |
| 1935 | OI/AV | M | 9 | 6 | 20210524 | |
| 1935 | OI/AV | M | 9 | 7 | 20210524 | |
| 1935 | OI/AV | M | 9 | 8 | 20210524 | |
| 1935 | OI/AV | M | 9 | 9 | 20210524 | |
| 1935 | OI/AV | M | 9 | 10 | 20210524 | |
| 1935 | OI/AV | M | 9 | 11 | 20210524 | |
| 1935 | OI/AV | M | 9 | 12 | 20210524 | |
| 1935 | OI/AV | M | 9 | | | 60 |
| 1935 | OI/AV | M | 13 | 1 | 20210621 | |
| 1935 | OI/AV | M | 13 | 2 | 20210621 | |
| 1935 | OI/AV | M | 13 | 3 | 20210621 | |
| 1935 | OI/AV | M | 13 | 4 | 20210621 | |
| 1935 | OI/AV | M | 13 | 5 | 20210621 | |
| 1935 | OI/AV | M | 13 | 6 | 20210621 | |
| 1935 | OI/AV | M | 13 | 7 | 20210621 | |

| | | | | | |
|---|---|---|---|---|---|
| 1935 | OI/AV | M | 13 | 8 | 20210621 | |
| 1935 | OI/AV | M | 13 | 9 | 20210621 | |
| 1935 | OI/AV | M | 13 | 10 | 20210621 | |
| 1935 | OI/AV | M | 13 | 11 | 20210621 | |
| 1935 | OI/AV | M | 13 | 12 | 20210621 | |
| 1935 | OI/AV | M | 13 | | | 60 |
| 3514 | WT | F | 5 | 1 | 20211213 | |
| 3514 | WT | F | 5 | 2 | 20211213 | |
| 3514 | WT | F | 5 | 3 | 20211213 | |
| 3514 | WT | F | 5 | 4 | 20211213 | |
| 3514 | WT | F | 5 | 5 | 20211213 | |
| 3514 | WT | F | 5 | 6 | 20211213 | |
| 3514 | WT | F | 5 | 7 | 20211213 | |
| 3514 | WT | F | 5 | 8 | 20211213 | |
| 3514 | WT | F | 5 | 9 | 20211213 | |
| 3514 | WT | F | 5 | 10 | 20211213 | |
| 3514 | WT | F | 5 | 11 | 20211213 | |
| 3514 | WT | F | 5 | 12 | 20211213 | |
| 3514 | WT | F | 5 | | | 60 |
| 3514 | WT | F | 9 | 1 | 20220110 | |
| 3514 | WT | F | 9 | 2 | 20220110 | |
| 3514 | WT | F | 9 | 3 | 20220110 | |
| 3514 | WT | F | 9 | 4 | 20220110 | |
| 3514 | WT | F | 9 | 5 | 20220110 | |
| 3514 | WT | F | 9 | 6 | 20220110 | |
| 3514 | WT | F | 9 | 7 | 20220110 | |
| 3514 | WT | F | 9 | 8 | 20220110 | |
| 3514 | WT | F | 9 | 9 | 20220110 | |
| 3514 | WT | F | 9 | 10 | 20220110 | |
| 3514 | WT | F | 9 | 11 | 20220110 | |
| 3514 | WT | F | 9 | 12 | 20220110 | |
| 3514 | WT | F | 9 | | | 60 |
| 3515 | OI/AV | M | 5 | 1 | 20211213 | |
| 3515 | OI/AV | M | 5 | 2 | 20211213 | |
| 3515 | OI/AV | M | 5 | 3 | 20211213 | |
| 3515 | OI/AV | M | 5 | 4 | 20211213 | |
| 3515 | OI/AV | M | 5 | 5 | 20211213 | |
| 3515 | OI/AV | M | 5 | 6 | 20211213 | |
| 3515 | OI/AV | M | 5 | 7 | 20211213 | |
| 3515 | OI/AV | M | 5 | 8 | 20211213 | |
| 3515 | OI/AV | M | 5 | 9 | 20211213 | |

| | | | | | |
|---|---|---|---|---|---|
| 3515 OI/AV | M | 5 | 10 | 20211213 | |
| 3515 OI/AV | M | 5 | 11 | 20211213 | |
| 3515 OI/AV | M | 5 | 12 | 20211213 | |
| 3515 OI/AV | M | 5 | | | 60 |
| 3515 OI/AV | M | 9 | 1 | 20220110 | |
| 3515 OI/AV | M | 9 | 2 | 20220110 | |
| 3515 OI/AV | M | 9 | 3 | 20220110 | |
| 3515 OI/AV | M | 9 | 4 | 20220110 | |
| 3515 OI/AV | M | 9 | 5 | 20220110 | |
| 3515 OI/AV | M | 9 | 6 | 20220110 | |
| 3515 OI/AV | M | 9 | 7 | 20220110 | |
| 3515 OI/AV | M | 9 | 8 | 20220110 | |
| 3515 OI/AV | M | 9 | 9 | 20220110 | |
| 3515 OI/AV | M | 9 | 10 | 20220110 | |
| 3515 OI/AV | M | 9 | 11 | 20220110 | |
| 3515 OI/AV | M | 9 | 12 | 20220110 | |
| 3515 OI/AV | M | 9 | | | 60 |
| 3520 OI | F | 5 | 1 | 20211213 | |
| 3520 OI | F | 5 | 2 | 20211213 | |
| 3520 OI | F | 5 | 3 | 20211213 | |
| 3520 OI | F | 5 | 4 | 20211213 | |
| 3520 OI | F | 5 | 5 | 20211213 | |
| 3520 OI | F | 5 | 6 | 20211213 | |
| 3520 OI | F | 5 | 7 | 20211213 | |
| 3520 OI | F | 5 | 8 | 20211213 | |
| 3520 OI | F | 5 | 9 | 20211213 | |
| 3520 OI | F | 5 | 10 | 20211213 | |
| 3520 OI | F | 5 | 11 | 20211213 | |
| 3520 OI | F | 5 | 12 | 20211213 | |
| 3520 OI | F | 5 | | | 60 |
| 3520 OI | F | 9 | 1 | 20220110 | |
| 3520 OI | F | 9 | 2 | 20220110 | |
| 3520 OI | F | 9 | 3 | 20220110 | |
| 3520 OI | F | 9 | 4 | 20220110 | |
| 3520 OI | F | 9 | 5 | 20220110 | |
| 3520 OI | F | 9 | 6 | 20220110 | |
| 3520 OI | F | 9 | 7 | 20220110 | |
| 3520 OI | F | 9 | 8 | 20220110 | |
| 3520 OI | F | 9 | 9 | 20220110 | |
| 3520 OI | F | 9 | 10 | 20220110 | |
| 3520 OI | F | 9 | 11 | 20220110 | |

| | | | | | | |
|---|---|---|---|---|---|---|
| 3520 | OI | F | 9 | 12 | 20220110 | |
| 3520 | OI | F | 9 | | | 60 |
| 3521 | WT | F | 5 | 1 | 20211213 | |
| 3521 | WT | F | 5 | 2 | 20211213 | |
| 3521 | WT | F | 5 | 3 | 20211213 | |
| 3521 | WT | F | 5 | 4 | 20211213 | |
| 3521 | WT | F | 5 | 5 | 20211213 | |
| 3521 | WT | F | 5 | 6 | 20211213 | |
| 3521 | WT | F | 5 | 7 | 20211213 | |
| 3521 | WT | F | 5 | 8 | 20211213 | |
| 3521 | WT | F | 5 | 9 | 20211213 | |
| 3521 | WT | F | 5 | 10 | 20211213 | |
| 3521 | WT | F | 5 | 11 | 20211213 | |
| 3521 | WT | F | 5 | 12 | 20211213 | |
| 3521 | WT | F | 5 | | | 60 |
| 3521 | WT | F | 9 | 1 | 20220110 | |
| 3521 | WT | F | 9 | 2 | 20220110 | |
| 3521 | WT | F | 9 | 3 | 20220110 | |
| 3521 | WT | F | 9 | 4 | 20220110 | |
| 3521 | WT | F | 9 | 5 | 20220110 | |
| 3521 | WT | F | 9 | 6 | 20220110 | |
| 3521 | WT | F | 9 | 7 | 20220110 | |
| 3521 | WT | F | 9 | 8 | 20220110 | |
| 3521 | WT | F | 9 | 9 | 20220110 | |
| 3521 | WT | F | 9 | 10 | 20220110 | |
| 3521 | WT | F | 9 | 11 | 20220110 | |
| 3521 | WT | F | 9 | 12 | 20220110 | |
| 3521 | WT | F | 9 | | | 60 |
| 3522 | OI/AV | M | 5 | 1 | 20211213 | |
| 3522 | OI/AV | M | 5 | 2 | 20211213 | |
| 3522 | OI/AV | M | 5 | 3 | 20211213 | |
| 3522 | OI/AV | M | 5 | 4 | 20211213 | |
| 3522 | OI/AV | M | 5 | 5 | 20211213 | |
| 3522 | OI/AV | M | 5 | 6 | 20211213 | |
| 3522 | OI/AV | M | 5 | 7 | 20211213 | |
| 3522 | OI/AV | M | 5 | 8 | 20211213 | |
| 3522 | OI/AV | M | 5 | 9 | 20211213 | |
| 3522 | OI/AV | M | 5 | 10 | 20211213 | |
| 3522 | OI/AV | M | 5 | 11 | 20211213 | |
| 3522 | OI/AV | M | 5 | 12 | 20211213 | |
| 3522 | OI/AV | M | 5 | | | 60 |

| | | | | | | |
|---|---|---|---|---|---|---|
| 3522 | OI/AV | M | 9 | 1 | 20220110 | |
| 3522 | OI/AV | M | 9 | 2 | 20220110 | |
| 3522 | OI/AV | M | 9 | 3 | 20220110 | |
| 3522 | OI/AV | M | 9 | 4 | 20220110 | |
| 3522 | OI/AV | M | 9 | 5 | 20220110 | |
| 3522 | OI/AV | M | 9 | 6 | 20220110 | |
| 3522 | OI/AV | M | 9 | 7 | 20220110 | |
| 3522 | OI/AV | M | 9 | 8 | 20220110 | |
| 3522 | OI/AV | M | 9 | 9 | 20220110 | |
| 3522 | OI/AV | M | 9 | 10 | 20220110 | |
| 3522 | OI/AV | M | 9 | 11 | 20220110 | |
| 3522 | OI/AV | M | 9 | 12 | 20220110 | |
| 3522 | OI/AV | M | 9 | | | 60 |
| 3523 | OI/AV | M | 5 | 1 | 20211213 | |
| 3523 | OI/AV | M | 5 | 2 | 20211213 | |
| 3523 | OI/AV | M | 5 | 3 | 20211213 | |
| 3523 | OI/AV | M | 5 | 4 | 20211213 | |
| 3523 | OI/AV | M | 5 | 5 | 20211213 | |
| 3523 | OI/AV | M | 5 | 6 | 20211213 | |
| 3523 | OI/AV | M | 5 | 7 | 20211213 | |
| 3523 | OI/AV | M | 5 | 8 | 20211213 | |
| 3523 | OI/AV | M | 5 | 9 | 20211213 | |
| 3523 | OI/AV | M | 5 | 10 | 20211213 | |
| 3523 | OI/AV | M | 5 | 11 | 20211213 | |
| 3523 | OI/AV | M | 5 | 12 | 20211213 | |
| 3523 | OI/AV | M | 5 | | | 60 |
| 3523 | OI/AV | M | 9 | 1 | 20220110 | |
| 3523 | OI/AV | M | 9 | 2 | 20220110 | |
| 3523 | OI/AV | M | 9 | 3 | 20220110 | |
| 3523 | OI/AV | M | 9 | 4 | 20220110 | |
| 3523 | OI/AV | M | 9 | 5 | 20220110 | |
| 3523 | OI/AV | M | 9 | 6 | 20220110 | |
| 3523 | OI/AV | M | 9 | 7 | 20220110 | |
| 3523 | OI/AV | M | 9 | 8 | 20220110 | |
| 3523 | OI/AV | M | 9 | 9 | 20220110 | |
| 3523 | OI/AV | M | 9 | 10 | 20220110 | |
| 3523 | OI/AV | M | 9 | 11 | 20220110 | |
| 3523 | OI/AV | M | 9 | 12 | 20220110 | |
| 3523 | OI/AV | M | 9 | | | 60 |
| 4113 | OI/AV | F | 5 | 1 | 20210607 | |
| 4113 | OI/AV | F | 5 | 2 | 20210607 | |

| | | | | | | |
|---|---|---|---|---|---|---|
| 4113 | OI/AV | F | 5 | 3 | 20210607 | |
| 4113 | OI/AV | F | 5 | 4 | 20210607 | |
| 4113 | OI/AV | F | 5 | 5 | 20210607 | |
| 4113 | OI/AV | F | 5 | 6 | 20210607 | |
| 4113 | OI/AV | F | 5 | 7 | 20210607 | |
| 4113 | OI/AV | F | 5 | 8 | 20210607 | |
| 4113 | OI/AV | F | 5 | 9 | 20210607 | |
| 4113 | OI/AV | F | 5 | 10 | 20210607 | |
| 4113 | OI/AV | F | 5 | 11 | 20210607 | |
| 4113 | OI/AV | F | 5 | 12 | 20210607 | |
| 4113 | OI/AV | F | 5 | | | 60 |
| 4113 | OI/AV | F | 9 | 1 | 20210705 | |
| 4113 | OI/AV | F | 9 | 2 | 20210705 | |
| 4113 | OI/AV | F | 9 | 3 | 20210705 | |
| 4113 | OI/AV | F | 9 | 4 | 20210705 | |
| 4113 | OI/AV | F | 9 | 5 | 20210705 | |
| 4113 | OI/AV | F | 9 | 6 | 20210705 | |
| 4113 | OI/AV | F | 9 | 7 | 20210705 | |
| 4113 | OI/AV | F | 9 | 8 | 20210705 | |
| 4113 | OI/AV | F | 9 | 9 | 20210705 | |
| 4113 | OI/AV | F | 9 | 10 | 20210705 | |
| 4113 | OI/AV | F | 9 | 11 | 20210705 | |
| 4113 | OI/AV | F | 9 | 12 | 20210705 | |
| 4113 | OI/AV | F | 9 | | | 60 |
| 4113 | OI/AV | F | 13 | | 20210802 | |
| 4113 | OI/AV | F | 13 | | 20210802 | |
| 4113 | OI/AV | F | 13 | | 20210802 | |
| 4113 | OI/AV | F | 13 | | 20210802 | |
| 4113 | OI/AV | F | 13 | | 20210802 | |
| 4113 | OI/AV | F | 13 | | 20210802 | |
| 4113 | OI/AV | F | 13 | | 20210802 | |
| 4113 | OI/AV | F | 13 | | 20210802 | |
| 4113 | OI/AV | F | 13 | | 20210802 | |
| 4113 | OI/AV | F | 13 | | 20210802 | |
| 4113 | OI/AV | F | 13 | | 20210802 | |
| 4113 | OI/AV | F | 13 | | 20210802 | |
| 4113 | OI/AV | F | 13 | | | 60 |
| 4114 | WT | F | 5 | | 20210607 | |
| 4114 | WT | F | 5 | | 20210607 | |
| 4114 | WT | F | 5 | | 20210607 | |
| 4114 | WT | F | 5 | | 20210607 | |

| | | | | | | |
|---|---|---|---|---|---|---|
| 4114 | WT | F | 5 | | 20210607 | |
| 4114 | WT | F | 5 | | 20210607 | |
| 4114 | WT | F | 5 | | 20210607 | |
| 4114 | WT | F | 5 | | 20210607 | |
| 4114 | WT | F | 5 | | 20210607 | |
| 4114 | WT | F | 5 | | 20210607 | |
| 4114 | WT | F | 5 | | 20210607 | |
| 4114 | WT | F | 5 | | 20210607 | |
| 4114 | WT | F | 5 | | | 60 |
| 4114 | WT | F | 9 | 1 | 20210705 | |
| 4114 | WT | F | 9 | 2 | 20210705 | |
| 4114 | WT | F | 9 | 3 | 20210705 | |
| 4114 | WT | F | 9 | 4 | 20210705 | |
| 4114 | WT | F | 9 | 5 | 20210705 | |
| 4114 | WT | F | 9 | 6 | 20210705 | |
| 4114 | WT | F | 9 | 7 | 20210705 | |
| 4114 | WT | F | 9 | 8 | 20210705 | |
| 4114 | WT | F | 9 | 9 | 20210705 | |
| 4114 | WT | F | 9 | 10 | 20210705 | |
| 4114 | WT | F | 9 | 11 | 20210705 | |
| 4114 | WT | F | 9 | 12 | 20210705 | |
| 4114 | WT | F | 9 | | | 60 |
| 4114 | WT | F | 13 | 1 | 20210802 | |
| 4114 | WT | F | 13 | 2 | 20210802 | |
| 4114 | WT | F | 13 | 3 | 20210802 | |
| 4114 | WT | F | 13 | 4 | 20210802 | |
| 4114 | WT | F | 13 | 5 | 20210802 | |
| 4114 | WT | F | 13 | 6 | 20210802 | |
| 4114 | WT | F | 13 | 7 | 20210802 | |
| 4114 | WT | F | 13 | 8 | 20210802 | |
| 4114 | WT | F | 13 | 9 | 20210802 | |
| 4114 | WT | F | 13 | 10 | 20210802 | |
| 4114 | WT | F | 13 | 11 | 20210802 | |
| 4114 | WT | F | 13 | 12 | 20210802 | |
| 4114 | WT | F | 13 | | | 60 |
| 4115 | OI | F | 5 | 1 | 20210607 | |
| 4115 | OI | F | 5 | 2 | 20210607 | |
| 4115 | OI | F | 5 | 3 | 20210607 | |
| 4115 | OI | F | 5 | 4 | 20210607 | |
| 4115 | OI | F | 5 | 5 | 20210607 | |
| 4115 | OI | F | 5 | 6 | 20210607 | |

| | | | | | | |
|---|---|---|---|---|---|---|
| 4115 | OI | F | 5 | 7 | 20210607 | |
| 4115 | OI | F | 5 | 8 | 20210607 | |
| 4115 | OI | F | 5 | 9 | 20210607 | |
| 4115 | OI | F | 5 | 10 | 20210607 | |
| 4115 | OI | F | 5 | 11 | 20210607 | |
| 4115 | OI | F | 5 | 12 | 20210607 | |
| 4115 | OI | F | 5 | | | 60 |
| 4115 | OI | F | 9 | 1 | 20210705 | |
| 4115 | OI | F | 9 | 2 | 20210705 | |
| 4115 | OI | F | 9 | 3 | 20210705 | |
| 4115 | OI | F | 9 | 4 | 20210705 | |
| 4115 | OI | F | 9 | 5 | 20210705 | |
| 4115 | OI | F | 9 | 6 | 20210705 | |
| 4115 | OI | F | 9 | 7 | 20210705 | |
| 4115 | OI | F | 9 | 8 | 20210705 | |
| 4115 | OI | F | 9 | 9 | 20210705 | |
| 4115 | OI | F | 9 | 10 | 20210705 | |
| 4115 | OI | F | 9 | 11 | 20210705 | |
| 4115 | OI | F | 9 | 12 | 20210705 | |
| 4115 | OI | F | 9 | | | 60 |
| 4115 | OI | F | 13 | | 20210802 | |
| 4115 | OI | F | 13 | | 20210802 | |
| 4115 | OI | F | 13 | | 20210802 | |
| 4115 | OI | F | 13 | | 20210802 | |
| 4115 | OI | F | 13 | | 20210802 | |
| 4115 | OI | F | 13 | | 20210802 | |
| 4115 | OI | F | 13 | | 20210802 | |
| 4115 | OI | F | 13 | | 20210802 | |
| 4115 | OI | F | 13 | | 20210802 | |
| 4115 | OI | F | 13 | | 20210802 | |
| 4115 | OI | F | 13 | | 20210802 | |
| 4115 | OI | F | 13 | | 20210802 | |
| 4115 | OI | F | 13 | | | 60 |
| 4117 | OI/AV | M | 5 | | 20210607 | |
| 4117 | OI/AV | M | 5 | | 20210607 | |
| 4117 | OI/AV | M | 5 | | 20210607 | |
| 4117 | OI/AV | M | 5 | | 20210607 | |
| 4117 | OI/AV | M | 5 | | 20210607 | |
| 4117 | OI/AV | M | 5 | | 20210607 | |
| 4117 | OI/AV | M | 5 | | 20210607 | |
| 4117 | OI/AV | M | 5 | | 20210607 | |

| | | | | | |
|---|---|---|---|---|---|
| 4117 | OI/AV | M | 5 | | 20210607 | |
| 4117 | OI/AV | M | 5 | | 20210607 | |
| 4117 | OI/AV | M | 5 | | 20210607 | |
| 4117 | OI/AV | M | 5 | | 20210607 | |
| 4117 | OI/AV | M | 5 | | | 60 |
| 4117 | OI/AV | M | 9 | 1 | 20210705 | |
| 4117 | OI/AV | M | 9 | 2 | 20210705 | |
| 4117 | OI/AV | M | 9 | 3 | 20210705 | |
| 4117 | OI/AV | M | 9 | 4 | 20210705 | |
| 4117 | OI/AV | M | 9 | 5 | 20210705 | |
| 4117 | OI/AV | M | 9 | 6 | 20210705 | |
| 4117 | OI/AV | M | 9 | 7 | 20210705 | |
| 4117 | OI/AV | M | 9 | 8 | 20210705 | |
| 4117 | OI/AV | M | 9 | 9 | 20210705 | |
| 4117 | OI/AV | M | 9 | 10 | 20210705 | |
| 4117 | OI/AV | M | 9 | 11 | 20210705 | |
| 4117 | OI/AV | M | 9 | 12 | 20210705 | |
| 4117 | OI/AV | M | 9 | | | 60 |
| 4117 | OI/AV | M | 13 | 1 | 20210802 | |
| 4117 | OI/AV | M | 13 | 2 | 20210802 | |
| 4117 | OI/AV | M | 13 | 3 | 20210802 | |
| 4117 | OI/AV | M | 13 | 4 | 20210802 | |
| 4117 | OI/AV | M | 13 | 5 | 20210802 | |
| 4117 | OI/AV | M | 13 | 6 | 20210802 | |
| 4117 | OI/AV | M | 13 | 7 | 20210802 | |
| 4117 | OI/AV | M | 13 | 8 | 20210802 | |
| 4117 | OI/AV | M | 13 | 9 | 20210802 | |
| 4117 | OI/AV | M | 13 | 10 | 20210802 | |
| 4117 | OI/AV | M | 13 | 11 | 20210802 | |
| 4117 | OI/AV | M | 13 | 12 | 20210802 | |
| 4117 | OI/AV | M | 13 | | | 60 |
| 4591 | OI/AV | F | 5 | 1 | 20210920 | |
| 4591 | OI/AV | F | 5 | 2 | 20210920 | |
| 4591 | OI/AV | F | 5 | 3 | 20210920 | |
| 4591 | OI/AV | F | 5 | 4 | 20210920 | |
| 4591 | OI/AV | F | 5 | 5 | 20210920 | |
| 4591 | OI/AV | F | 5 | 6 | 20210920 | |
| 4591 | OI/AV | F | 5 | 7 | 20210920 | |
| 4591 | OI/AV | F | 5 | 8 | 20210920 | |
| 4591 | OI/AV | F | 5 | 9 | 20210920 | |
| 4591 | OI/AV | F | 5 | 10 | 20210920 | |

| | | | | | | |
|---|---|---|---|---|---|---|
| 4591 | OI/AV | F | 5 | 11 | 20210920 | |
| 4591 | OI/AV | F | 5 | 12 | 20210920 | |
| 4591 | OI/AV | F | 5 | | | 60 |
| 4591 | OI/AV | F | 9 | 1 | 20211019 | |
| 4591 | OI/AV | F | 9 | 2 | 20211019 | |
| 4591 | OI/AV | F | 9 | 3 | 20211019 | |
| 4591 | OI/AV | F | 9 | 4 | 20211019 | |
| 4591 | OI/AV | F | 9 | 5 | 20211019 | |
| 4591 | OI/AV | F | 9 | 6 | 20211019 | |
| 4591 | OI/AV | F | 9 | 7 | 20211019 | |
| 4591 | OI/AV | F | 9 | 8 | 20211019 | |
| 4591 | OI/AV | F | 9 | 9 | 20211019 | |
| 4591 | OI/AV | F | 9 | 10 | 20211019 | |
| 4591 | OI/AV | F | 9 | 11 | 20211019 | |
| 4591 | OI/AV | F | 9 | 12 | 20211019 | |
| 4591 | OI/AV | F | 9 | | | 60 |
| 4591 | OI/AV | F | 13 | 1 | 20211115 | |
| 4591 | OI/AV | F | 13 | 2 | 20211115 | |
| 4591 | OI/AV | F | 13 | 3 | 20211115 | |
| 4591 | OI/AV | F | 13 | 4 | 20211115 | |
| 4591 | OI/AV | F | 13 | 5 | 20211115 | |
| 4591 | OI/AV | F | 13 | 6 | 20211115 | |
| 4591 | OI/AV | F | 13 | 7 | 20211115 | |
| 4591 | OI/AV | F | 13 | 8 | 20211115 | |
| 4591 | OI/AV | F | 13 | 9 | 20211115 | |
| 4591 | OI/AV | F | 13 | 10 | 20211115 | |
| 4591 | OI/AV | F | 13 | 11 | 20211115 | |
| 4591 | OI/AV | F | 13 | 12 | 20211115 | |
| 4591 | OI/AV | F | 13 | | | 60 |
| 4592 | OI/AV | F | 5 | 1 | 20210920 | |
| 4592 | OI/AV | F | 5 | 2 | 20210920 | |
| 4592 | OI/AV | F | 5 | 3 | 20210920 | |
| 4592 | OI/AV | F | 5 | 4 | 20210920 | |
| 4592 | OI/AV | F | 5 | 5 | 20210920 | |
| 4592 | OI/AV | F | 5 | 6 | 20210920 | |
| 4592 | OI/AV | F | 5 | 7 | 20210920 | |
| 4592 | OI/AV | F | 5 | 8 | 20210920 | |
| 4592 | OI/AV | F | 5 | 9 | 20210920 | |
| 4592 | OI/AV | F | 5 | 10 | 20210920 | |
| 4592 | OI/AV | F | 5 | 11 | 20210920 | |
| 4592 | OI/AV | F | 5 | 12 | 20210920 | |

| | | | | | | |
|---|---|---|---|---|---|---|
| 4592 | OI/AV | F | 5 | | | 60 |
| 4592 | OI/AV | F | 9 | 1 | 20211019 | |
| 4592 | OI/AV | F | 9 | 2 | 20211019 | |
| 4592 | OI/AV | F | 9 | 3 | 20211019 | |
| 4592 | OI/AV | F | 9 | 4 | 20211019 | |
| 4592 | OI/AV | F | 9 | 5 | 20211019 | |
| 4592 | OI/AV | F | 9 | 6 | 20211019 | |
| 4592 | OI/AV | F | 9 | 7 | 20211019 | |
| 4592 | OI/AV | F | 9 | 8 | 20211019 | |
| 4592 | OI/AV | F | 9 | 9 | 20211019 | |
| 4592 | OI/AV | F | 9 | 10 | 20211019 | |
| 4592 | OI/AV | F | 9 | 11 | 20211019 | |
| 4592 | OI/AV | F | 9 | 12 | 20211019 | |
| 4592 | OI/AV | F | 9 | | | 60 |
| 4592 | OI/AV | F | 13 | 1 | 20211115 | |
| 4592 | OI/AV | F | 13 | 2 | 20211115 | |
| 4592 | OI/AV | F | 13 | 3 | 20211115 | |
| 4592 | OI/AV | F | 13 | 4 | 20211115 | |
| 4592 | OI/AV | F | 13 | 5 | 20211115 | |
| 4592 | OI/AV | F | 13 | 6 | 20211115 | |
| 4592 | OI/AV | F | 13 | 7 | 20211115 | |
| 4592 | OI/AV | F | 13 | 8 | 20211115 | |
| 4592 | OI/AV | F | 13 | 9 | 20211115 | |
| 4592 | OI/AV | F | 13 | 10 | 20211115 | |
| 4592 | OI/AV | F | 13 | 11 | 20211115 | |
| 4592 | OI/AV | F | 13 | 12 | 20211115 | |
| 4592 | OI/AV | F | 13 | | | 60 |
| 4594 | OI/AV | F | 5 | 1 | 20210920 | |
| 4594 | OI/AV | F | 5 | 2 | 20210920 | |
| 4594 | OI/AV | F | 5 | 3 | 20210920 | |
| 4594 | OI/AV | F | 5 | 4 | 20210920 | |
| 4594 | OI/AV | F | 5 | 5 | 20210920 | |
| 4594 | OI/AV | F | 5 | 6 | 20210920 | |
| 4594 | OI/AV | F | 5 | 7 | 20210920 | |
| 4594 | OI/AV | F | 5 | 8 | 20210920 | |
| 4594 | OI/AV | F | 5 | 9 | 20210920 | |
| 4594 | OI/AV | F | 5 | 10 | 20210920 | |
| 4594 | OI/AV | F | 5 | 11 | 20210920 | |
| 4594 | OI/AV | F | 5 | 12 | 20210920 | |
| 4594 | OI/AV | F | 5 | | | 60 |
| 4594 | OI/AV | F | 9 | 1 | 20211019 | |

| | | | | | | |
|---|---|---|---|---|---|---|
| 4594 | OI/AV | F | 9 | 2 | 20211019 | |
| 4594 | OI/AV | F | 9 | 3 | 20211019 | |
| 4594 | OI/AV | F | 9 | 4 | 20211019 | |
| 4594 | OI/AV | F | 9 | 5 | 20211019 | |
| 4594 | OI/AV | F | 9 | 6 | 20211019 | |
| 4594 | OI/AV | F | 9 | 7 | 20211019 | |
| 4594 | OI/AV | F | 9 | 8 | 20211019 | |
| 4594 | OI/AV | F | 9 | 9 | 20211019 | |
| 4594 | OI/AV | F | 9 | 10 | 20211019 | |
| 4594 | OI/AV | F | 9 | 11 | 20211019 | |
| 4594 | OI/AV | F | 9 | 12 | 20211019 | |
| 4594 | OI/AV | F | 9 | | | 60 |
| 4594 | OI/AV | F | 13 | 1 | 20211115 | |
| 4594 | OI/AV | F | 13 | 2 | 20211115 | |
| 4594 | OI/AV | F | 13 | 3 | 20211115 | |
| 4594 | OI/AV | F | 13 | 4 | 20211115 | |
| 4594 | OI/AV | F | 13 | 5 | 20211115 | |
| 4594 | OI/AV | F | 13 | 6 | 20211115 | |
| 4594 | OI/AV | F | 13 | 7 | 20211115 | |
| 4594 | OI/AV | F | 13 | 8 | 20211115 | |
| 4594 | OI/AV | F | 13 | 9 | 20211115 | |
| 4594 | OI/AV | F | 13 | 10 | 20211115 | |
| 4594 | OI/AV | F | 13 | 11 | 20211115 | |
| 4594 | OI/AV | F | 13 | 12 | 20211115 | |
| 4594 | OI/AV | F | 13 | | | 60 |
| 4596 | OI | M | 5 | 1 | 20210920 | |
| 4596 | OI | M | 5 | 2 | 20210920 | |
| 4596 | OI | M | 5 | 3 | 20210920 | |
| 4596 | OI | M | 5 | 4 | 20210920 | |
| 4596 | OI | M | 5 | 5 | 20210920 | |
| 4596 | OI | M | 5 | 6 | 20210920 | |
| 4596 | OI | M | 5 | 7 | 20210920 | |
| 4596 | OI | M | 5 | 8 | 20210920 | |
| 4596 | OI | M | 5 | 9 | 20210920 | |
| 4596 | OI | M | 5 | 10 | 20210920 | |
| 4596 | OI | M | 5 | 11 | 20210920 | |
| 4596 | OI | M | 5 | 12 | 20210920 | |
| 4596 | OI | M | 5 | | | 60 |
| 4596 | OI | M | 9 | 1 | 20211019 | |
| 4596 | OI | M | 9 | 2 | 20211019 | |
| 4596 | OI | M | 9 | 3 | 20211019 | |

| | | | | | |
|---|---|---|---|---|---|
| 4596 | OI | M | 9 | 4 | 20211019 | |
| 4596 | OI | M | 9 | 5 | 20211019 | |
| 4596 | OI | M | 9 | 6 | 20211019 | |
| 4596 | OI | M | 9 | 7 | 20211019 | |
| 4596 | OI | M | 9 | 8 | 20211019 | |
| 4596 | OI | M | 9 | 9 | 20211019 | |
| 4596 | OI | M | 9 | 10 | 20211019 | |
| 4596 | OI | M | 9 | 11 | 20211019 | |
| 4596 | OI | M | 9 | 12 | 20211019 | |
| 4596 | OI | M | 9 | | | 60 |
| 4596 | OI | M | 13 | 1 | 20211115 | |
| 4596 | OI | M | 13 | 2 | 20211115 | |
| 4596 | OI | M | 13 | 3 | 20211115 | |
| 4596 | OI | M | 13 | 4 | 20211115 | |
| 4596 | OI | M | 13 | 5 | 20211115 | |
| 4596 | OI | M | 13 | 6 | 20211115 | |
| 4596 | OI | M | 13 | 7 | 20211115 | |
| 4596 | OI | M | 13 | 8 | 20211115 | |
| 4596 | OI | M | 13 | 9 | 20211115 | |
| 4596 | OI | M | 13 | 10 | 20211115 | |
| 4596 | OI | M | 13 | 11 | 20211115 | |
| 4596 | OI | M | 13 | 12 | 20211115 | |
| 4596 | OI | M | 13 | | | 60 |
| 4598 | OI/AV | M | 5 | 1 | 20210920 | |
| 4598 | OI/AV | M | 5 | 2 | 20210920 | |
| 4598 | OI/AV | M | 5 | 3 | 20210920 | |
| 4598 | OI/AV | M | 5 | 4 | 20210920 | |
| 4598 | OI/AV | M | 5 | 5 | 20210920 | |
| 4598 | OI/AV | M | 5 | 6 | 20210920 | |
| 4598 | OI/AV | M | 5 | 7 | 20210920 | |
| 4598 | OI/AV | M | 5 | 8 | 20210920 | |
| 4598 | OI/AV | M | 5 | 9 | 20210920 | |
| 4598 | OI/AV | M | 5 | 10 | 20210920 | |
| 4598 | OI/AV | M | 5 | 11 | 20210920 | |
| 4598 | OI/AV | M | 5 | 12 | 20210920 | |
| 4598 | OI/AV | M | 5 | | | 60 |
| 4598 | OI/AV | M | 9 | 1 | 20211019 | |
| 4598 | OI/AV | M | 9 | 2 | 20211019 | |
| 4598 | OI/AV | M | 9 | 3 | 20211019 | |
| 4598 | OI/AV | M | 9 | 4 | 20211019 | |
| 4598 | OI/AV | M | 9 | 5 | 20211019 | |

| | | | | | |
|---|---|---|---|---|---|
| 4598 | OI/AV | M | 9 | 6 | 20211019 | |
| 4598 | OI/AV | M | 9 | 7 | 20211019 | |
| 4598 | OI/AV | M | 9 | 8 | 20211019 | |
| 4598 | OI/AV | M | 9 | 9 | 20211019 | |
| 4598 | OI/AV | M | 9 | 10 | 20211019 | |
| 4598 | OI/AV | M | 9 | 11 | 20211019 | |
| 4598 | OI/AV | M | 9 | 12 | 20211019 | |
| 4598 | OI/AV | M | 9 | | | 60 |
| 4598 | OI/AV | M | 13 | 1 | 20211115 | |
| 4598 | OI/AV | M | 13 | 2 | 20211115 | |
| 4598 | OI/AV | M | 13 | 3 | 20211115 | |
| 4598 | OI/AV | M | 13 | 4 | 20211115 | |
| 4598 | OI/AV | M | 13 | 5 | 20211115 | |
| 4598 | OI/AV | M | 13 | 6 | 20211115 | |
| 4598 | OI/AV | M | 13 | 7 | 20211115 | |
| 4598 | OI/AV | M | 13 | 8 | 20211115 | |
| 4598 | OI/AV | M | 13 | 9 | 20211115 | |
| 4598 | OI/AV | M | 13 | 10 | 20211115 | |
| 4598 | OI/AV | M | 13 | 11 | 20211115 | |
| 4598 | OI/AV | M | 13 | 12 | 20211115 | |
| 4598 | OI/AV | M | 13 | | | 60 |
| 4599 | WT | M | 5 | 1 | 20210920 | |
| 4599 | WT | M | 5 | 2 | 20210920 | |
| 4599 | WT | M | 5 | 3 | 20210920 | |
| 4599 | WT | M | 5 | 4 | 20210920 | |
| 4599 | WT | M | 5 | 5 | 20210920 | |
| 4599 | WT | M | 5 | 6 | 20210920 | |
| 4599 | WT | M | 5 | 7 | 20210920 | |
| 4599 | WT | M | 5 | 8 | 20210920 | |
| 4599 | WT | M | 5 | 9 | 20210920 | |
| 4599 | WT | M | 5 | 10 | 20210920 | |
| 4599 | WT | M | 5 | 11 | 20210920 | |
| 4599 | WT | M | 5 | 12 | 20210920 | |
| 4599 | WT | M | 5 | | | 60 |
| 4599 | WT | M | 9 | 1 | 20211019 | |
| 4599 | WT | M | 9 | 2 | 20211019 | |
| 4599 | WT | M | 9 | 3 | 20211019 | |
| 4599 | WT | M | 9 | 4 | 20211019 | |
| 4599 | WT | M | 9 | 5 | 20211019 | |
| 4599 | WT | M | 9 | 6 | 20211019 | |
| 4599 | WT | M | 9 | 7 | 20211019 | |

| | | | | | | |
|---|---|---|---|---|---|---|
| 4599 | WT | M | 9 | 8 | 20211019 | |
| 4599 | WT | M | 9 | 9 | 20211019 | |
| 4599 | WT | M | 9 | 10 | 20211019 | |
| 4599 | WT | M | 9 | 11 | 20211019 | |
| 4599 | WT | M | 9 | 12 | 20211019 | |
| 4599 | WT | M | 9 | | | 60 |
| 4599 | WT | M | 13 | 1 | 20211115 | |
| 4599 | WT | M | 13 | 2 | 20211115 | |
| 4599 | WT | M | 13 | 3 | 20211115 | |
| 4599 | WT | M | 13 | 4 | 20211115 | |
| 4599 | WT | M | 13 | 5 | 20211115 | |
| 4599 | WT | M | 13 | 6 | 20211115 | |
| 4599 | WT | M | 13 | 7 | 20211115 | |
| 4599 | WT | M | 13 | 8 | 20211115 | |
| 4599 | WT | M | 13 | 9 | 20211115 | |
| 4599 | WT | M | 13 | 10 | 20211115 | |
| 4599 | WT | M | 13 | 11 | 20211115 | |
| 4599 | WT | M | 13 | 12 | 20211115 | |
| 4599 | WT | M | 13 | | | 60 |
| 4685 | WT | F | 5 | 1 | 20211122 | |
| 4685 | WT | F | 5 | 2 | 20211122 | |
| 4685 | WT | F | 5 | 3 | 20211122 | |
| 4685 | WT | F | 5 | 4 | 20211122 | |
| 4685 | WT | F | 5 | 5 | 20211122 | |
| 4685 | WT | F | 5 | 6 | 20211122 | |
| 4685 | WT | F | 5 | 7 | 20211122 | |
| 4685 | WT | F | 5 | 8 | 20211122 | |
| 4685 | WT | F | 5 | 9 | 20211122 | |
| 4685 | WT | F | 5 | 10 | 20211122 | |
| 4685 | WT | F | 5 | 11 | 20211122 | |
| 4685 | WT | F | 5 | 12 | 20211122 | |
| 4685 | WT | F | 5 | | | 60 |
| 4685 | WT | F | 9 | 1 | 20211215 | |
| 4685 | WT | F | 9 | 2 | 20211215 | |
| 4685 | WT | F | 9 | 3 | 20211215 | |
| 4685 | WT | F | 9 | 4 | 20211215 | |
| 4685 | WT | F | 9 | 5 | 20211215 | |
| 4685 | WT | F | 9 | 6 | 20211215 | |
| 4685 | WT | F | 9 | 7 | 20211215 | |
| 4685 | WT | F | 9 | 8 | 20211215 | |
| 4685 | WT | F | 9 | 9 | 20211215 | |

| | | | | | | |
|---|---|---|---|---|---|---|
| 4685 | WT | F | 9 | 10 | 20211215 | |
| 4685 | WT | F | 9 | 11 | 20211215 | |
| 4685 | WT | F | 9 | 12 | 20211215 | |
| 4685 | WT | F | 9 | | | 60 |
| 4685 | WT | F | 13 | 1 | 20220119 | |
| 4685 | WT | F | 13 | 2 | 20220119 | |
| 4685 | WT | F | 13 | 3 | 20220119 | |
| 4685 | WT | F | 13 | 4 | 20220119 | |
| 4685 | WT | F | 13 | 5 | 20220119 | |
| 4685 | WT | F | 13 | 6 | 20220119 | |
| 4685 | WT | F | 13 | 7 | 20220119 | |
| 4685 | WT | F | 13 | 8 | 20220119 | |
| 4685 | WT | F | 13 | 9 | 20220119 | |
| 4685 | WT | F | 13 | 10 | 20220119 | |
| 4685 | WT | F | 13 | 11 | 20220119 | |
| 4685 | WT | F | 13 | 12 | 20220119 | |
| 4685 | WT | F | 13 | | | 60 |
| 4686 | OI/AV | F | 5 | 1 | 20211122 | |
| 4686 | OI/AV | F | 5 | 2 | 20211122 | |
| 4686 | OI/AV | F | 5 | 3 | 20211122 | |
| 4686 | OI/AV | F | 5 | 4 | 20211122 | |
| 4686 | OI/AV | F | 5 | 5 | 20211122 | |
| 4686 | OI/AV | F | 5 | 6 | 20211122 | |
| 4686 | OI/AV | F | 5 | 7 | 20211122 | |
| 4686 | OI/AV | F | 5 | 8 | 20211122 | |
| 4686 | OI/AV | F | 5 | 9 | 20211122 | |
| 4686 | OI/AV | F | 5 | 10 | 20211122 | |
| 4686 | OI/AV | F | 5 | 11 | 20211122 | |
| 4686 | OI/AV | F | 5 | 12 | 20211122 | |
| 4686 | OI/AV | F | 5 | | | 60 |
| 4686 | OI/AV | F | 9 | 1 | 20211215 | |
| 4686 | OI/AV | F | 9 | 2 | 20211215 | |
| 4686 | OI/AV | F | 9 | 3 | 20211215 | |
| 4686 | OI/AV | F | 9 | 4 | 20211215 | |
| 4686 | OI/AV | F | 9 | 5 | 20211215 | |
| 4686 | OI/AV | F | 9 | 6 | 20211215 | |
| 4686 | OI/AV | F | 9 | 7 | 20211215 | |
| 4686 | OI/AV | F | 9 | 8 | 20211215 | |
| 4686 | OI/AV | F | 9 | 9 | 20211215 | |
| 4686 | OI/AV | F | 9 | 10 | 20211215 | |
| 4686 | OI/AV | F | 9 | 11 | 20211215 | |

| | | | | | | |
|---|---|---|---|---|---|---|
| 4686 | OI/AV | F | 9 | 12 | 20211215 | |
| 4686 | OI/AV | F | 9 | | | 60 |
| 4686 | OI/AV | F | 13 | 1 | 20220119 | |
| 4686 | OI/AV | F | 13 | 2 | 20220119 | |
| 4686 | OI/AV | F | 13 | 3 | 20220119 | |
| 4686 | OI/AV | F | 13 | 4 | 20220119 | |
| 4686 | OI/AV | F | 13 | 5 | 20220119 | |
| 4686 | OI/AV | F | 13 | 6 | 20220119 | |
| 4686 | OI/AV | F | 13 | 7 | 20220119 | |
| 4686 | OI/AV | F | 13 | 8 | 20220119 | |
| 4686 | OI/AV | F | 13 | 9 | 20220119 | |
| 4686 | OI/AV | F | 13 | 10 | 20220119 | |
| 4686 | OI/AV | F | 13 | 11 | 20220119 | |
| 4686 | OI/AV | F | 13 | 12 | 20220119 | |
| 4686 | OI/AV | F | 13 | | | 60 |
| 4689 | WT/AV | M | 5 | 1 | 20211122 | |
| 4689 | WT/AV | M | 5 | 2 | 20211122 | |
| 4689 | WT/AV | M | 5 | 3 | 20211122 | |
| 4689 | WT/AV | M | 5 | 4 | 20211122 | |
| 4689 | WT/AV | M | 5 | 5 | 20211122 | |
| 4689 | WT/AV | M | 5 | 6 | 20211122 | |
| 4689 | WT/AV | M | 5 | 7 | 20211122 | |
| 4689 | WT/AV | M | 5 | 8 | 20211122 | |
| 4689 | WT/AV | M | 5 | 9 | 20211122 | |
| 4689 | WT/AV | M | 5 | 10 | 20211122 | |
| 4689 | WT/AV | M | 5 | 11 | 20211122 | |
| 4689 | WT/AV | M | 5 | 12 | 20211122 | |
| 4689 | WT/AV | M | 5 | | | 60 |
| 4689 | WT/AV | M | 9 | 1 | 20211215 | |
| 4689 | WT/AV | M | 9 | 2 | 20211215 | |
| 4689 | WT/AV | M | 9 | 3 | 20211215 | |
| 4689 | WT/AV | M | 9 | 4 | 20211215 | |
| 4689 | WT/AV | M | 9 | 5 | 20211215 | |
| 4689 | WT/AV | M | 9 | 6 | 20211215 | |
| 4689 | WT/AV | M | 9 | 7 | 20211215 | |
| 4689 | WT/AV | M | 9 | 8 | 20211215 | |
| 4689 | WT/AV | M | 9 | 9 | 20211215 | |
| 4689 | WT/AV | M | 9 | 10 | 20211215 | |
| 4689 | WT/AV | M | 9 | 11 | 20211215 | |
| 4689 | WT/AV | M | 9 | 12 | 20211215 | |
| 4689 | WT/AV | M | 9 | | | 60 |

| | | | | | | |
|---|---|---|---|---|---|---|
| 4689 | WT/AV | M | 13 | 1 | 20220119 | |
| 4689 | WT/AV | M | 13 | 2 | 20220119 | |
| 4689 | WT/AV | M | 13 | 3 | 20220119 | |
| 4689 | WT/AV | M | 13 | 4 | 20220119 | |
| 4689 | WT/AV | M | 13 | 5 | 20220119 | |
| 4689 | WT/AV | M | 13 | 6 | 20220119 | |
| 4689 | WT/AV | M | 13 | 7 | 20220119 | |
| 4689 | WT/AV | M | 13 | 8 | 20220119 | |
| 4689 | WT/AV | M | 13 | 9 | 20220119 | |
| 4689 | WT/AV | M | 13 | 10 | 20220119 | |
| 4689 | WT/AV | M | 13 | 11 | 20220119 | |
| 4689 | WT/AV | M | 13 | 12 | 20220119 | |
| 4689 | WT/AV | M | 13 | | | 60 |
| 4690 | OI | M | 5 | 1 | 20211122 | |
| 4690 | OI | M | 5 | 2 | 20211122 | |
| 4690 | OI | M | 5 | 3 | 20211122 | |
| 4690 | OI | M | 5 | 4 | 20211122 | |
| 4690 | OI | M | 5 | 5 | 20211122 | |
| 4690 | OI | M | 5 | 6 | 20211122 | |
| 4690 | OI | M | 5 | 7 | 20211122 | |
| 4690 | OI | M | 5 | 8 | 20211122 | |
| 4690 | OI | M | 5 | 9 | 20211122 | |
| 4690 | OI | M | 5 | 10 | 20211122 | |
| 4690 | OI | M | 5 | 11 | 20211122 | |
| 4690 | OI | M | 5 | 12 | 20211122 | |
| 4690 | OI | M | 5 | | | 60 |
| 4690 | OI | M | 9 | 1 | 20211215 | |
| 4690 | OI | M | 9 | 2 | 20211215 | |
| 4690 | OI | M | 9 | 3 | 20211215 | |
| 4690 | OI | M | 9 | 4 | 20211215 | |
| 4690 | OI | M | 9 | 5 | 20211215 | |
| 4690 | OI | M | 9 | 6 | 20211215 | |
| 4690 | OI | M | 9 | 7 | 20211215 | |
| 4690 | OI | M | 9 | 8 | 20211215 | |
| 4690 | OI | M | 9 | 9 | 20211215 | |
| 4690 | OI | M | 9 | 10 | 20211215 | |
| 4690 | OI | M | 9 | 11 | 20211215 | |
| 4690 | OI | M | 9 | 12 | 20211215 | |
| 4690 | OI | M | 9 | | | 60 |
| 4690 | OI | M | 13 | 1 | 20220119 | |
| 4690 | OI | M | 13 | 2 | 20220119 | |

| | | | | | | |
|---|---|---|---|---|---|---|
| 4690 | OI | M | 13 | 3 | 20220119 | |
| 4690 | OI | M | 13 | 4 | 20220119 | |
| 4690 | OI | M | 13 | 5 | 20220119 | |
| 4690 | OI | M | 13 | 6 | 20220119 | |
| 4690 | OI | M | 13 | 7 | 20220119 | |
| 4690 | OI | M | 13 | 8 | 20220119 | |
| 4690 | OI | M | 13 | 9 | 20220119 | |
| 4690 | OI | M | 13 | 10 | 20220119 | |
| 4690 | OI | M | 13 | 11 | 20220119 | |
| 4690 | OI | M | 13 | 12 | 20220119 | |
| 4690 | OI | M | 13 | | | 60 |
| all (9) | WT | M | 5 | | | 60 |
| All (12) | WT | F | 5 | | | 60 |
| All (9) | OI | F | 5 | | | 60 |
| All (11) | OI | M | 5 | | | 60 |

| Basic Movem | poke 1 | poke 2 | poke 3 | poke 4 | poke 5 | poke 6 |
|---|---|---|---|---|---|---|
| 551 | 5 | 6 | 13 | 6 | 2 | 2 |
| 386 | 2 | 0 | 3 | 5 | 1 | 1 |
| 345 | 11 | 3 | 15 | 3 | 1 | 2 |
| 264 | 7 | 5 | 2 | 0 | 0 | 0 |
| 152 | 7 | 2 | 4 | 0 | 1 | 0 |
| 346 | 9 | 5 | 3 | 1 | 1 | 1 |
| 175 | 3 | 0 | 0 | 0 | 0 | 0 |
| 95 | 3 | 0 | 0 | 0 | 0 | 0 |
| 31 | 2 | 0 | 0 | 0 | 0 | 0 |
| 86 | 2 | 0 | 0 | 0 | 0 | 0 |
| 39 | 0 | 0 | 0 | 0 | 0 | 0 |
| 68 | 0 | 0 | 0 | 0 | 0 | 0 |
| 2538 | 190 | | | | | |
| 622 | 9 | 5 | 13 | 0 | 0 | 3 |
| 452 | 4 | 3 | 5 | 0 | 0 | 1 |
| 294 | 5 | 0 | 19 | 0 | 0 | 0 |
| 131 | 0 | 0 | 0 | 0 | 0 | 0 |
| 96 | 0 | 0 | 0 | 0 | 0 | 0 |
| 493 | 1 | 3 | 0 | 4 | 4 | 6 |
| 139 | 0 | 0 | 1 | 0 | 0 | 0 |
| 91 | 0 | 0 | 0 | 0 | 0 | 0 |
| 27 | 0 | 0 | 0 | 0 | 0 | 0 |
| 102 | 0 | 0 | 0 | 0 | 0 | 0 |
| 35 | 0 | 0 | 0 | 0 | 0 | 0 |
| 21 | 0 | 0 | 0 | 0 | 0 | 0 |
| 2503 | 110 | | | | | |
| 686 | 6 | 0 | 2 | 0 | 0 | 0 |
| 348 | 0 | 0 | 2 | 0 | 0 | 0 |
| 360 | 3 | 0 | 0 | 0 | 0 | 0 |
| 117 | 0 | 0 | 0 | 0 | 0 | 0 |
| 113 | 0 | 0 | 0 | 0 | 0 | 0 |
| 110 | 0 | 0 | 0 | 0 | 0 | 0 |
| 29 | 0 | 0 | 0 | 0 | 0 | 0 |
| 24 | 0 | 0 | 0 | 0 | 0 | 0 |
| 24 | 0 | 0 | 0 | 0 | 0 | 0 |
| 64 | 0 | 0 | 0 | 0 | 0 | 0 |
| 445 | 3 | 3 | 0 | 2 | 3 | 5 |
| 335 | 0 | 0 | 2 | 0 | 0 | 0 |
| 2655 | 40 | | | | | |
| 618 | 5 | 0 | 4 | 4 | 1 | 3 |

| | | | | | | |
|---|---|---|---|---|---|---|
| 549 | 10 | 6 | 7 | 1 | 1 | 1 |
| 462 | 4 | 4 | 3 | 0 | 3 | 5 |
| 297 | 6 | 0 | 5 | 4 | 0 | 0 |
| 463 | 1 | 7 | 2 | 2 | 2 | 0 |
| 180 | 1 | 0 | 0 | 0 | 0 | 0 |
| 421 | 11 | 1 | 3 | 2 | 3 | 1 |
| 113 | 6 | 0 | 0 | 0 | 0 | 0 |
| 260 | 1 | 0 | 2 | 0 | 0 | 0 |
| 104 | 0 | 0 | 0 | 0 | 0 | 0 |
| 109 | 0 | 0 | 0 | 0 | 0 | 0 |
| 12 | 0 | 0 | 0 | 0 | 0 | 0 |
| 3588 | 166 | | | | | |
| 538 | 3 | 4 | 4 | 0 | 0 | 2 |
| 523 | 3 | 2 | 4 | 2 | 4 | 9 |
| 386 | 0 | 3 | 2 | 0 | 0 | 0 |
| 385 | 6 | 0 | 1 | 2 | 0 | 0 |
| 267 | 3 | 4 | 2 | 0 | 0 | 0 |
| 282 | 2 | 0 | 2 | 0 | 0 | 0 |
| 225 | 1 | 0 | 0 | 0 | 0 | 0 |
| 316 | 0 | 0 | 3 | 2 | 0 | 0 |
| 275 | 0 | 0 | 1 | 0 | 0 | 0 |
| 184 | 0 | 0 | 0 | 0 | 0 | 0 |
| 258 | 0 | 0 | 0 | 0 | 0 | 0 |
| 87 | 0 | 0 | 0 | 0 | 0 | 0 |
| 3726 | 109 | | | | | |
| 446 | 3 | 3 | 0 | 5 | 1 | 1 |
| 306 | 0 | 0 | 0 | 0 | 0 | 0 |
| 317 | 2 | 0 | 6 | 3 | 4 | 0 |
| 245 | 0 | 0 | 1 | 0 | 0 | 0 |
| 189 | 0 | 0 | 0 | 0 | 0 | 0 |
| 80 | 0 | 0 | 0 | 0 | 0 | 0 |
| 79 | 4 | 0 | 0 | 0 | 0 | 0 |
| 74 | 0 | 0 | 0 | 0 | 0 | 0 |
| 8 | 0 | 0 | 0 | 0 | 0 | 0 |
| 7 | 0 | 0 | 0 | 0 | 0 | 0 |
| 2 | 0 | 0 | 0 | 0 | 0 | 0 |
| 288 | 0 | 0 | 1 | 0 | 0 | 0 |
| 2041 | 46 | | | | | |
| 732 | 9 | 5 | 6 | 2 | 0 | 3 |
| 443 | 4 | 0 | 2 | 0 | 0 | 0 |
| 337 | 4 | 2 | 9 | 0 | 2 | 0 |

| | | | | | | |
|---|---|---|---|---|---|---|
| 136 | 5 | 0 | 0 | 0 | 0 | 0 |
| 260 | 1 | 0 | 2 | 2 | 0 | 0 |
| 126 | 7 | 0 | 0 | 0 | 0 | 0 |
| 107 | 0 | 0 | 0 | 0 | 0 | 0 |
| 43 | 1 | 0 | 0 | 0 | 0 | 0 |
| 97 | 3 | 0 | 0 | 0 | 0 | 0 |
| 170 | 2 | 0 | 4 | 0 | 0 | 0 |
| 58 | 0 | 0 | 0 | 0 | 0 | 0 |
| 0 | 0 | 0 | 0 | 0 | 0 | 0 |
| 2509 | 91 | | | | | |
| 439 | 1 | 0 | 3 | 0 | 2 | 0 |
| 501 | 3 | 1 | 2 | 1 | 0 | 2 |
| 424 | 0 | 0 | 2 | 0 | 0 | 0 |
| 436 | 2 | 0 | 0 | 0 | 0 | 1 |
| 384 | 0 | 0 | 1 | 0 | 0 | 0 |
| 388 | 0 | 2 | 0 | 0 | 0 | 0 |
| 362 | 0 | 0 | 3 | 0 | 0 | 0 |
| 194 | 0 | 0 | 0 | 0 | 0 | 0 |
| 86 | 0 | 0 | 0 | 0 | 0 | 0 |
| 91 | 0 | 0 | 0 | 0 | 0 | 0 |
| 374 | 0 | 0 | 0 | 0 | 0 | 0 |
| 296 | 0 | 0 | 4 | 0 | 0 | 0 |
| 3975 | 63 | | | | | |
| 686 | 10 | 3 | 4 | 7 | 0 | 2 |
| 508 | 6 | 2 | 7 | 6 | 0 | 1 |
| 605 | 15 | 3 | 7 | 5 | 0 | 2 |
| 481 | 5 | 0 | 3 | 0 | 0 | 0 |
| 445 | 5 | 6 | 2 | 7 | 10 | 7 |
| 482 | 6 | 2 | 5 | 2 | 2 | 4 |
| 442 | 13 | 1 | 2 | 9 | 2 | 1 |
| 594 | 4 | 3 | 12 | 1 | 1 | 2 |
| 425 | 1 | 0 | 2 | 1 | 0 | 1 |
| 262 | 2 | 1 | 4 | 0 | 1 | 0 |
| 266 | 0 | 0 | 0 | 1 | 0 | 0 |
| 293 | 2 | 1 | 1 | 0 | 1 | 0 |
| 5489 | 372 | | | | | |
| 725 | 7 | 4 | 3 | 2 | 1 | 0 |
| 646 | 0 | 2 | 9 | 0 | 2 | 2 |
| 578 | 4 | 0 | 2 | 2 | 0 | 4 |
| 558 | 8 | 5 | 4 | 1 | 2 | 0 |
| 548 | 3 | 1 | 7 | 1 | 1 | 2 |

| | | | | | | |
|---|---|---|---|---|---|---|
| 394 | 1 | 2 | 4 | 3 | 1 | 1 |
| 535 | 3 | 1 | 1 | 0 | 1 | 1 |
| 271 | 0 | 0 | 0 | 0 | 0 | 0 |
| 415 | 4 | 0 | 4 | 0 | 0 | 0 |
| 360 | 0 | 0 | 8 | 0 | 0 | 1 |
| 365 | 0 | 1 | 1 | 0 | 0 | 0 |
| 136 | 0 | 0 | 0 | 0 | 0 | 0 |
| 5531 | 183 | | | | | |
| 528 | 2 | 1 | 2 | 2 | 0 | 1 |
| 503 | 3 | 3 | 4 | 0 | 0 | 1 |
| 400 | 2 | 1 | 1 | 0 | 2 | 0 |
| 369 | 1 | 0 | 0 | 0 | 0 | 0 |
| 487 | 3 | 0 | 4 | 0 | 0 | 0 |
| 256 | 0 | 0 | 1 | 0 | 0 | 0 |
| 326 | 4 | 0 | 1 | 2 | 0 | 0 |
| 240 | 0 | 1 | 1 | 0 | 0 | 0 |
| 162 | 0 | 0 | 0 | 0 | 0 | 0 |
| 118 | 0 | 0 | 0 | 0 | 0 | 0 |
| 189 | 1 | 0 | 0 | 0 | 0 | 0 |
| 12 | 0 | 0 | 0 | 0 | 0 | 0 |
| 3590 | 105 | | | | | |
| 548 | 11 | 7 | 8 | 5 | 0 | 5 |
| 376 | 2 | 3 | 9 | 7 | 0 | 5 |
| 424 | 3 | 4 | 5 | 0 | 0 | 3 |
| 445 | 7 | 4 | 10 | 3 | 4 | 2 |
| 434 | 5 | 0 | 5 | 2 | 4 | 4 |
| 197 | 2 | 1 | 7 | 0 | 0 | 3 |
| 274 | 1 | 5 | 3 | 1 | 0 | 1 |
| 322 | 10 | 4 | 3 | 2 | 1 | 0 |
| 250 | 4 | 1 | 10 | 0 | 0 | 3 |
| 180 | 9 | 1 | 1 | 1 | 0 | 0 |
| 188 | 1 | 0 | 8 | 0 | 0 | 0 |
| 113 | 1 | 0 | 2 | 0 | 0 | 0 |
| 3751 | 300 | | | | | |
| 628 | 4 | 2 | 6 | 5 | 3 | 6 |
| 462 | 10 | 4 | 3 | 6 | 1 | 0 |
| 472 | 1 | 4 | 2 | 5 | 1 | 3 |
| 363 | 6 | 2 | 2 | 2 | 1 | 5 |
| 345 | 7 | 1 | 1 | 3 | 2 | 1 |
| 369 | 3 | 4 | 3 | 2 | 4 | 1 |
| 402 | 0 | 3 | 1 | 2 | 0 | 0 |

| | | | | | | |
|---|---|---|---|---|---|---|
| 289 | 4 | 1 | 2 | 4 | 2 | 4 |
| 306 | 3 | 4 | 6 | 2 | 0 | 0 |
| 259 | 1 | 1 | 8 | 8 | 0 | 2 |
| 345 | 1 | 1 | 0 | 0 | 0 | 1 |
| 183 | 6 | 0 | 0 | 0 | 0 | 0 |
| 4423 | 259 | | | | | |
| 632 | 16 | 3 | 9 | 2 | 1 | 9 |
| 462 | 3 | 3 | 3 | 1 | 4 | 7 |
| 510 | 4 | 2 | 1 | 1 | 5 | 0 |
| 382 | 5 | 4 | 2 | 4 | 0 | 1 |
| 488 | 3 | 5 | 3 | 0 | 0 | 2 |
| 498 | 6 | 4 | 3 | 2 | 2 | 3 |
| 452 | 1 | 4 | 2 | 0 | 2 | 2 |
| 385 | 4 | 1 | 3 | 1 | 1 | 0 |
| 251 | 2 | 1 | 2 | 0 | 0 | 1 |
| 293 | 1 | 1 | 0 | 1 | 0 | 0 |
| 104 | 0 | 0 | 0 | 0 | 0 | 0 |
| 113 | 0 | 0 | 0 | 0 | 0 | 0 |
| 4570 | 224 | | | | | |
| 655 | 6 | 5 | 2 | 4 | 2 | 0 |
| 438 | 7 | 8 | 5 | 1 | 7 | 2 |
| 385 | 4 | 4 | 3 | 1 | 0 | 2 |
| 381 | 3 | 1 | 3 | 2 | 1 | 1 |
| 423 | 10 | 3 | 2 | 4 | 4 | 3 |
| 341 | 3 | 3 | 8 | 0 | 2 | 0 |
| 376 | 3 | 4 | 2 | 1 | 0 | 0 |
| 369 | 2 | 0 | 3 | 0 | 0 | 2 |
| 278 | 1 | 0 | 5 | 0 | 0 | 2 |
| 299 | 0 | 0 | 2 | 2 | 2 | 0 |
| 338 | 15 | 1 | 3 | 2 | 1 | 0 |
| 298 | 3 | 3 | 3 | 0 | 0 | 1 |
| 4581 | 239 | | | | | |
| 456 | 5 | 1 | 18 | 0 | 0 | 0 |
| 345 | 5 | 7 | 8 | 0 | 0 | 0 |
| 460 | 9 | 7 | 12 | 2 | 0 | 0 |
| 364 | 2 | 5 | 18 | 1 | 0 | 3 |
| 406 | 7 | 4 | 8 | 2 | 0 | 4 |
| 279 | 10 | 6 | 3 | 1 | 0 | 2 |
| 316 | 1 | 1 | 9 | 1 | 0 | 3 |
| 229 | 1 | 3 | 9 | 0 | 0 | 1 |
| 229 | 1 | 0 | 4 | 0 | 0 | 1 |

| | | | | | | |
|---|---|---|---|---|---|---|
| 214 | 1 | 1 | 6 | 0 | 0 | 0 |
| 130 | 0 | 0 | 3 | 0 | 0 | 0 |
| 76 | 0 | 0 | 1 | 0 | 0 | 0 |
| 3504 | 260 | | | | | |
| 428 | 3 | 5 | 4 | 3 | 0 | 1 |
| 483 | 3 | 3 | 7 | 2 | 0 | 2 |
| 412 | 3 | 2 | 4 | 0 | 0 | 2 |
| 368 | 3 | 2 | 3 | 2 | 0 | 0 |
| 460 | 1 | 3 | 4 | 2 | 6 | 0 |
| 276 | 0 | 0 | 1 | 0 | 1 | 1 |
| 161 | 0 | 0 | 3 | 0 | 0 | 0 |
| 108 | 0 | 0 | 2 | 0 | 0 | 0 |
| 274 | 0 | 0 | 6 | 0 | 0 | 0 |
| 263 | 0 | 0 | 0 | 0 | 2 | 0 |
| 186 | 0 | 0 | 1 | 0 | 0 | 0 |
| 7 | 0 | 0 | 0 | 0 | 0 | 0 |
| 3426 | 132 | | | | | |
| 406 | 1 | 1 | 0 | 0 | 0 | 1 |
| 554 | 5 | 6 | 5 | 1 | 1 | 0 |
| 442 | 3 | 10 | 5 | 3 | 1 | 4 |
| 547 | 6 | 7 | 3 | 2 | 1 | 0 |
| 390 | 6 | 2 | 0 | 0 | 2 | 0 |
| 299 | 3 | 2 | 1 | 0 | 0 | 0 |
| 269 | 1 | 0 | 0 | 0 | 1 | 1 |
| 214 | 0 | 1 | 1 | 0 | 0 | 0 |
| 274 | 2 | 0 | 1 | 0 | 0 | 4 |
| 155 | 2 | 0 | 0 | 0 | 0 | 0 |
| 234 | 1 | 3 | 0 | 0 | 1 | 0 |
| 413 | 2 | 0 | 0 | 6 | 1 | 1 |
| 4197 | 141 | | | | | |
| 580 | 10 | 2 | 8 | 1 | 0 | 0 |
| 441 | 8 | 3 | 7 | 2 | 3 | 6 |
| 431 | 3 | 12 | 10 | 0 | 2 | 3 |
| 437 | 1 | 0 | 2 | 0 | 0 | 0 |
| 360 | 2 | 1 | 2 | 2 | 0 | 0 |
| 357 | 7 | 3 | 7 | 0 | 1 | 4 |
| 276 | 9 | 3 | 10 | 0 | 0 | 0 |
| 117 | 13 | 0 | 0 | 0 | 0 | 0 |
| 78 | 6 | 0 | 0 | 0 | 0 | 0 |
| 70 | 2 | 0 | 0 | 0 | 0 | 0 |
| 9 | 0 | 0 | 0 | 0 | 0 | 0 |

| | | | | | | |
|---|---|---|---|---|---|---|
| 5 | 0 | 0 | 0 | 0 | 0 | 0 |
| 3161 | 185 | | | | | |
| 442 | 5 | 1 | 4 | 1 | 2 | 0 |
| 374 | 3 | 3 | 0 | 2 | 0 | 2 |
| 490 | 9 | 1 | 1 | 6 | 5 | 0 |
| 284 | 2 | 0 | 4 | 0 | 0 | 0 |
| 106 | 0 | 0 | 0 | 0 | 0 | 0 |
| 65 | 0 | 0 | 0 | 0 | 0 | 0 |
| 17 | 0 | 0 | 0 | 0 | 0 | 0 |
| 9 | 0 | 0 | 0 | 0 | 0 | 0 |
| 10 | 0 | 0 | 0 | 0 | 0 | 0 |
| 57 | 0 | 0 | 0 | 0 | 0 | 0 |
| 0 | 0 | 0 | 0 | 0 | 0 | 0 |
| 0 | 0 | 0 | 0 | 0 | 0 | 0 |
| 1854 | 65 | | | | | |
| 449 | 2 | 3 | 3 | 0 | 0 | 0 |
| 245 | 1 | 0 | 0 | 0 | 0 | 1 |
| 234 | 0 | 0 | 0 | 1 | 2 | 0 |
| 164 | 4 | 0 | 0 | 0 | 0 | 0 |
| 5 | 0 | 0 | 0 | 0 | 0 | 0 |
| 53 | 0 | 0 | 0 | 0 | 0 | 0 |
| 17 | 0 | 0 | 0 | 0 | 0 | 0 |
| 0 | 0 | 0 | 0 | 0 | 0 | 0 |
| 21 | 0 | 0 | 0 | 0 | 0 | 0 |
| 87 | 0 | 0 | 0 | 0 | 0 | 0 |
| 114 | 0 | 0 | 0 | 0 | 0 | 0 |
| 153 | 2 | 1 | 2 | 0 | 0 | 0 |
| 1542 | 36 | | | | | |
| 569 | 7 | 6 | 8 | 3 | 0 | 0 |
| 536 | 4 | 3 | 8 | 2 | 2 | 2 |
| 409 | 5 | 2 | 5 | 3 | 0 | 0 |
| 420 | 2 | 1 | 3 | 0 | 3 | 0 |
| 502 | 13 | 2 | 7 | 4 | 1 | 0 |
| 373 | 3 | 2 | 0 | 5 | 1 | 0 |
| 380 | 2 | 0 | 0 | 0 | 0 | 0 |
| 263 | 0 | 0 | 0 | 0 | 0 | 0 |
| 335 | 7 | 1 | 0 | 2 | 0 | 0 |
| 214 | 1 | 0 | 0 | 0 | 0 | 0 |
| 154 | 0 | 0 | 1 | 0 | 0 | 3 |
| 96 | 0 | 0 | 0 | 0 | 0 | 0 |
| 4251 | 173 | | | | | |

| | | | | | | |
|---|---|---|---|---|---|---|
| 455 | 3 | 0 | 1 | 3 | 3 | 0 |
| 397 | 0 | 2 | 2 | 0 | 0 | 1 |
| 388 | 4 | 0 | 7 | 0 | 0 | 0 |
| 396 | 1 | 0 | 6 | 0 | 0 | 3 |
| 154 | 0 | 0 | 0 | 0 | 0 | 0 |
| 104 | 0 | 0 | 0 | 0 | 0 | 0 |
| 50 | 0 | 0 | 0 | 0 | 0 | 0 |
| 70 | 0 | 0 | 0 | 0 | 0 | 0 |
| 11 | 0 | 0 | 0 | 0 | 0 | 0 |
| 53 | 0 | 0 | 0 | 0 | 0 | 0 |
| 8 | 0 | 0 | 0 | 0 | 0 | 0 |
| 45 | 0 | 0 | 0 | 0 | 0 | 0 |
| 2131 | 45 | | | | | |
| 486 | 4 | 2 | 1 | 1 | 0 | 1 |
| 442 | 1 | 0 | 0 | 0 | 0 | 0 |
| 410 | 1 | 2 | 11 | 0 | 0 | 0 |
| 344 | 0 | 0 | 1 | 0 | 0 | 0 |
| 121 | 9 | 0 | 0 | 0 | 0 | 0 |
| 151 | 0 | 0 | 2 | 0 | 0 | 0 |
| 274 | 1 | 2 | 1 | 0 | 0 | 3 |
| 43 | 0 | 0 | 0 | 0 | 0 | 0 |
| 8 | 0 | 0 | 0 | 0 | 0 | 0 |
| 12 | 0 | 0 | 0 | 0 | 0 | 0 |
| 186 | 2 | 0 | 0 | 2 | 0 | 0 |
| 222 | 0 | 0 | 3 | 0 | 0 | 0 |
| 2699 | 65 | | | | | |
| 310 | 8 | 0 | 23 | 0 | 0 | 0 |
| 257 | 4 | 0 | 6 | 0 | 0 | 7 |
| 264 | 2 | 7 | 4 | 5 | 4 | 5 |
| 288 | 3 | 0 | 0 | 1 | 3 | 0 |
| 270 | 10 | 0 | 5 | 0 | 0 | 0 |
| 239 | 5 | 1 | 3 | 3 | 1 | 2 |
| 228 | 0 | 0 | 11 | 1 | 1 | 1 |
| 161 | 4 | 4 | 0 | 5 | 0 | 2 |
| 167 | 0 | 0 | 6 | 0 | 0 | 0 |
| 208 | 0 | 0 | 0 | 3 | 0 | 0 |
| 254 | 3 | 0 | 3 | 5 | 1 | 0 |
| 66 | 0 | 0 | 1 | 0 | 0 | 0 |
| 2712 | 307 | | | | | |
| 315 | 2 | 3 | 12 | 0 | 3 | 2 |
| 281 | 10 | 0 | 4 | 0 | 0 | 0 |

| | | | | | | |
|---|---|---|---|---|---|---|
| 187 | 0 | 0 | 6 | 0 | 0 | 2 |
| 76 | 0 | 0 | 11 | 0 | 0 | 0 |
| 18 | 0 | 0 | 2 | 0 | 0 | 0 |
| 24 | 0 | 0 | 0 | 0 | 0 | 0 |
| 13 | 0 | 0 | 4 | 0 | 0 | 0 |
| 32 | 0 | 0 | 6 | 0 | 0 | 0 |
| 26 | 0 | 0 | 0 | 0 | 0 | 0 |
| 37 | 0 | 0 | 0 | 0 | 0 | 0 |
| 3 | 0 | 0 | 0 | 0 | 0 | 0 |
| 38 | 0 | 0 | 0 | 0 | 0 | 0 |
| 1050 | 83 | | | | | |
| 470 | 4 | 2 | 5 | 0 | 2 | 2 |
| 319 | 2 | 1 | 2 | 3 | 1 | 0 |
| 262 | 11 | 4 | 12 | 4 | 6 | 4 |
| 68 | 2 | 0 | 0 | 0 | 0 | 1 |
| 156 | 4 | 0 | 0 | 0 | 0 | 0 |
| 267 | 0 | 0 | 0 | 0 | 2 | 1 |
| 215 | 2 | 0 | 3 | 0 | 0 | 0 |
| 108 | 0 | 0 | 0 | 0 | 0 | 0 |
| 56 | 0 | 0 | 0 | 0 | 0 | 0 |
| 37 | 0 | 0 | 0 | 0 | 0 | 0 |
| 52 | 3 | 0 | 0 | 0 | 0 | 0 |
| 37 | 0 | 0 | 0 | 0 | 0 | 0 |
| 2047 | 149 | | | | | |
| 687 | 6 | 0 | 14 | 0 | 1 | 6 |
| 405 | 1 | 5 | 7 | 3 | 0 | 6 |
| 439 | 5 | 2 | 6 | 0 | 0 | 4 |
| 491 | 4 | 16 | 3 | 2 | 11 | 5 |
| 518 | 4 | 11 | 2 | 4 | 1 | 0 |
| 529 | 5 | 7 | 11 | 0 | 4 | 4 |
| 408 | 4 | 0 | 3 | 2 | 0 | 0 |
| 376 | 1 | 3 | 12 | 0 | 0 | 1 |
| 252 | 0 | 0 | 2 | 0 | 0 | 0 |
| 258 | 4 | 3 | 7 | 3 | 0 | 1 |
| 275 | 0 | 0 | 2 | 0 | 0 | 0 |
| 111 | 0 | 0 | 0 | 0 | 0 | 0 |
| 4749 | 269 | | | | | |
| 503 | 4 | 2 | 7 | 1 | 1 | 1 |
| 462 | 5 | 2 | 2 | 4 | 1 | 0 |
| 398 | 3 | 0 | 0 | 0 | 1 | 1 |
| 193 | 0 | 0 | 2 | 0 | 0 | 0 |

| | | | | | | |
|---|---|---|---|---|---|---|
| 121 | 0 | 0 | 0 | 0 | 0 | 0 |
| 404 | 1 | 2 | 2 | 0 | 0 | 0 |
| 289 | 3 | 0 | 0 | 0 | 0 | 0 |
| 215 | 0 | 0 | 0 | 0 | 0 | 2 |
| 49 | 0 | 0 | 0 | 0 | 0 | 0 |
| 83 | 0 | 0 | 0 | 0 | 0 | 0 |
| 51 | 0 | 0 | 0 | 0 | 0 | 0 |
| 46 | 0 | 0 | 0 | 0 | 0 | 0 |
| 2814 | 91 | | | | | |
| 498 | 0 | 0 | 2 | 0 | 0 | 2 |
| 416 | 5 | 1 | 1 | 2 | 2 | 2 |
| 498 | 3 | 8 | 13 | 5 | 0 | 3 |
| 535 | 2 | 12 | 1 | 3 | 0 | 0 |
| 455 | 1 | 19 | 3 | 2 | 1 | 1 |
| 484 | 1 | 5 | 2 | 1 | 3 | 5 |
| 579 | 3 | 4 | 0 | 0 | 0 | 1 |
| 456 | 2 | 1 | 2 | 2 | 1 | 0 |
| 609 | 0 | 7 | 2 | 1 | 0 | 1 |
| 460 | 7 | 2 | 1 | 2 | 0 | 0 |
| 379 | 4 | 1 | 1 | 1 | 0 | 7 |
| 363 | 0 | 0 | 1 | 0 | 0 | 0 |
| 5732 | 213 | | | | | |
| 662 | 20 | 4 | 12 | 0 | 0 | 2 |
| 433 | 10 | 4 | 11 | 1 | 0 | 2 |
| 443 | 6 | 5 | 8 | 3 | 0 | 1 |
| 419 | 6 | 11 | 6 | 2 | 6 | 0 |
| 443 | 4 | 1 | 14 | 1 | 5 | 1 |
| 303 | 2 | 5 | 3 | 0 | 3 | 3 |
| 265 | 0 | 3 | 7 | 0 | 0 | 1 |
| 133 | 0 | 0 | 1 | 0 | 0 | 0 |
| 160 | 0 | 0 | 3 | 0 | 0 | 0 |
| 50 | 0 | 0 | 0 | 0 | 0 | 0 |
| 33 | 0 | 0 | 0 | 0 | 0 | 0 |
| 3 | 0 | 0 | 0 | 0 | 0 | 0 |
| 3347 | 202 | | | | | |
| 632 | 6 | 2 | 4 | 2 | 3 | 5 |
| 528 | 2 | 3 | 4 | 1 | 1 | 2 |
| 368 | 0 | 0 | 0 | 0 | 1 | 1 |
| 401 | 9 | 1 | 7 | 0 | 0 | 2 |
| 321 | 0 | 0 | 0 | 0 | 0 | 0 |
| 486 | 2 | 2 | 4 | 4 | 3 | 3 |

| | | | | | | |
|---|---|---|---|---|---|---|
| 269 | 0 | 1 | 2 | 0 | 4 | 3 |
| 323 | 4 | 0 | 7 | 0 | 0 | 0 |
| 148 | 0 | 0 | 0 | 0 | 0 | 0 |
| 85 | 0 | 0 | 0 | 0 | 0 | 0 |
| 230 | 2 | 0 | 2 | 0 | 0 | 0 |
| 253 | 1 | 2 | 0 | 0 | 0 | 0 |
| 4044 | 138 | | | | | |
| 577 | 2 | 3 | 0 | 1 | 1 | 0 |
| 510 | 3 | 2 | 2 | 1 | 2 | 0 |
| 392 | 5 | 1 | 1 | 1 | 3 | 1 |
| 500 | 3 | 0 | 2 | 2 | 0 | 1 |
| 480 | 1 | 3 | 3 | 1 | 2 | 2 |
| 367 | 1 | 1 | 0 | 0 | 0 | 1 |
| 400 | 6 | 1 | 0 | 2 | 1 | 0 |
| 522 | 0 | 3 | 2 | 2 | 2 | 0 |
| 601 | 3 | 3 | 3 | 1 | 0 | 2 |
| 353 | 2 | 0 | 0 | 0 | 0 | 0 |
| 435 | 7 | 0 | 2 | 1 | 0 | 0 |
| 410 | 2 | 1 | 1 | 0 | 1 | 1 |
| 5547 | 142 | | | | | |
| 553 | 20 | 3 | 7 | 3 | 0 | 1 |
| 541 | 7 | 1 | 7 | 4 | 0 | 4 |
| 333 | 10 | 9 | 3 | 2 | 0 | 6 |
| 425 | 8 | 4 | 7 | 1 | 0 | 4 |
| 317 | 5 | 4 | 5 | 2 | 0 | 4 |
| 468 | 6 | 4 | 5 | 4 | 5 | 2 |
| 357 | 3 | 6 | 5 | 1 | 2 | 0 |
| 344 | 10 | 4 | 4 | 4 | 0 | 0 |
| 335 | 5 | 4 | 3 | 1 | 2 | 2 |
| 346 | 2 | 8 | 6 | 0 | 1 | 0 |
| 272 | 6 | 0 | 2 | 0 | 2 | 1 |
| 345 | 6 | 2 | 3 | 0 | 2 | 2 |
| 4636 | 343 | | | | | |
| 783 | 4 | 6 | 4 | 4 | 3 | 3 |
| 539 | 6 | 3 | 9 | 1 | 2 | 1 |
| 513 | 8 | 3 | 3 | 4 | 2 | 4 |
| 460 | 3 | 1 | 2 | 1 | 1 | 0 |
| 440 | 0 | 0 | 1 | 1 | 0 | 0 |
| 441 | 5 | 3 | 2 | 3 | 2 | 4 |
| 369 | 2 | 2 | 3 | 1 | 1 | 0 |
| 339 | 4 | 3 | 3 | 1 | 0 | 0 |

| | | | | | | |
|---|---|---|---|---|---|---|
| 321 | 0 | 0 | 0 | 0 | 0 | 0 |
| 320 | 1 | 1 | 4 | 3 | 2 | 1 |
| 315 | 1 | 2 | 2 | 1 | 0 | 1 |
| 175 | 0 | 0 | 1 | 0 | 0 | 0 |
| 5015 | 181 | | | | | |
| 694 | 6 | 3 | 1 | 1 | 3 | 2 |
| 479 | 0 | 2 | 3 | 3 | 3 | 2 |
| 376 | 3 | 7 | 3 | 0 | 1 | 1 |
| 433 | 2 | 3 | 1 | 2 | 0 | 0 |
| 384 | 0 | 0 | 1 | 0 | 0 | 0 |
| 335 | 0 | 2 | 1 | 0 | 0 | 0 |
| 404 | 4 | 1 | 5 | 0 | 2 | 1 |
| 412 | 0 | 1 | 2 | 3 | 0 | 2 |
| 321 | 3 | 4 | 3 | 0 | 3 | 1 |
| 289 | 0 | 1 | 0 | 5 | 0 | 0 |
| 352 | 1 | 0 | 2 | 0 | 0 | 0 |
| 334 | 0 | 2 | 0 | 1 | 0 | 0 |
| 4813 | 133 | | | | | |
| 852 | 3 | 7 | 5 | 2 | 1 | 3 |
| 644 | 3 | 0 | 6 | 1 | 5 | 2 |
| 543 | 6 | 2 | 7 | 0 | 0 | 2 |
| 520 | 10 | 4 | 6 | 8 | 0 | 5 |
| 483 | 2 | 0 | 3 | 0 | 0 | 0 |
| 513 | 3 | 2 | 6 | 2 | 0 | 0 |
| 473 | 2 | 1 | 6 | 0 | 0 | 0 |
| 306 | 8 | 0 | 5 | 1 | 0 | 0 |
| 316 | 1 | 0 | 0 | 0 | 0 | 0 |
| 199 | 0 | 0 | 0 | 0 | 0 | 0 |
| 377 | 0 | 0 | 0 | 0 | 0 | 0 |
| 145 | 2 | 0 | 0 | 0 | 0 | 0 |
| 5371 | 210 | | | | | |
| 708 | 7 | 1 | 5 | 1 | 0 | 3 |
| 527 | 0 | 0 | 4 | 2 | 0 | 2 |
| 448 | 1 | 1 | 3 | 0 | 0 | 3 |
| 455 | 1 | 1 | 20 | 1 | 2 | 2 |
| 395 | 0 | 0 | 2 | 0 | 0 | 0 |
| 422 | 1 | 2 | 2 | 1 | 0 | 2 |
| 367 | 2 | 1 | 5 | 0 | 1 | 2 |
| 171 | 0 | 0 | 0 | 0 | 0 | 0 |
| 165 | 1 | 0 | 4 | 0 | 0 | 0 |
| 80 | 0 | 0 | 0 | 0 | 0 | 0 |

| | | | | | | |
|---|---|---|---|---|---|---|
| 147 | 0 | 0 | 1 | 0 | 0 | 0 |
| 5 | 0 | 0 | 0 | 0 | 0 | 0 |
| 3890 | 133 | | | | | |
| 738 | 2 | 2 | 3 | 0 | 1 | 2 |
| 537 | 3 | 1 | 2 | 3 | 0 | 1 |
| 496 | 4 | 7 | 2 | 3 | 0 | 1 |
| 317 | 0 | 0 | 1 | 0 | 0 | 2 |
| 224 | 0 | 1 | 4 | 0 | 0 | 0 |
| 212 | 3 | 0 | 0 | 4 | 0 | 0 |
| 166 | 0 | 0 | 13 | 0 | 0 | 0 |
| 57 | 0 | 0 | 0 | 0 | 0 | 0 |
| 29 | 0 | 0 | 0 | 0 | 0 | 0 |
| 33 | 0 | 0 | 0 | 0 | 0 | 0 |
| 204 | 0 | 0 | 0 | 0 | 0 | 0 |
| 56 | 0 | 0 | 0 | 0 | 0 | 0 |
| 3069 | 79 | | | | | |
| 621 | 3 | 7 | 14 | 0 | 0 | 2 |
| 515 | 4 | 1 | 4 | 2 | 3 | 1 |
| 326 | 3 | 1 | 7 | 2 | 3 | 2 |
| 497 | 13 | 3 | 10 | 1 | 4 | 0 |
| 345 | 1 | 1 | 7 | 0 | 0 | 3 |
| 305 | 20 | 5 | 4 | 3 | 0 | 1 |
| 489 | 7 | 3 | 4 | 2 | 1 | 2 |
| 340 | 3 | 1 | 2 | 0 | 2 | 1 |
| 336 | 10 | 4 | 3 | 2 | 2 | 18 |
| 508 | 8 | 4 | 5 | 1 | 2 | 2 |
| 337 | 3 | 4 | 3 | 1 | 0 | 1 |
| 314 | 10 | 2 | 1 | 2 | 2 | 0 |
| 4933 | 319 | | | | | |
| 738 | 9 | 4 | 7 | 4 | 4 | 1 |
| 539 | 2 | 2 | 4 | 1 | 1 | 0 |
| 598 | 1 | 2 | 8 | 0 | 2 | 1 |
| 481 | 7 | 3 | 0 | 0 | 3 | 1 |
| 533 | 1 | 3 | 3 | 0 | 1 | 0 |
| 501 | 2 | 0 | 2 | 1 | 0 | 3 |
| 641 | 1 | 0 | 0 | 0 | 0 | 0 |
| 507 | 1 | 3 | 3 | 0 | 1 | 3 |
| 667 | 2 | 2 | 2 | 3 | 5 | 0 |
| 565 | 3 | 1 | 1 | 1 | 5 | 1 |
| 445 | 5 | 1 | 0 | 0 | 0 | 0 |
| 582 | 3 | 2 | 2 | 0 | 6 | 0 |

| | | | | | | |
|---|---|---|---|---|---|---|
| 6797 | 185 | | | | | |
| 689 | 5 | 1 | 4 | 2 | 0 | 3 |
| 584 | 3 | 0 | 3 | 2 | 3 | 1 |
| 469 | 2 | 1 | 8 | 2 | 0 | 4 |
| 412 | 9 | 3 | 2 | 1 | 0 | 2 |
| 381 | 4 | 0 | 4 | 2 | 1 | 0 |
| 471 | 2 | 0 | 2 | 0 | 0 | 2 |
| 431 | 2 | 0 | 4 | 0 | 0 | 3 |
| 510 | 2 | 0 | 2 | 0 | 3 | 6 |
| 337 | 0 | 0 | 0 | 0 | 0 | 1 |
| 405 | 2 | 0 | 2 | 3 | 0 | 3 |
| 684 | 0 | 2 | 2 | 0 | 0 | 0 |
| 688 | 0 | 1 | 0 | 4 | 0 | 0 |
| 6061 | 185 | | | | | |
| 800 | 8 | 1 | 7 | 1 | 0 | 1 |
| 486 | 9 | 6 | 10 | 1 | 0 | 0 |
| 533 | 3 | 4 | 6 | 1 | 2 | 0 |
| 441 | 3 | 0 | 3 | 0 | 3 | 4 |
| 244 | 10 | 0 | 4 | 0 | 0 | 0 |
| 218 | 0 | 0 | 2 | 1 | 0 | 0 |
| 126 | 0 | 0 | 0 | 0 | 0 | 0 |
| 73 | 0 | 0 | 0 | 0 | 0 | 0 |
| 16 | 0 | 0 | 0 | 0 | 0 | 0 |
| 145 | 1 | 0 | 1 | 0 | 0 | 0 |
| 283 | 4 | 0 | 0 | 0 | 0 | 0 |
| 61 | 0 | 0 | 0 | 0 | 0 | 0 |
| 3426 | 148 | | | | | |
| 662 | 1 | 6 | 3 | 5 | 1 | 2 |
| 699 | 6 | 1 | 4 | 0 | 2 | 11 |
| 698 | 4 | 6 | 4 | 5 | 3 | 1 |
| 563 | 0 | 1 | 2 | 3 | 1 | 8 |
| 644 | 3 | 4 | 1 | 4 | 2 | 2 |
| 553 | 0 | 4 | 1 | 1 | 1 | 1 |
| 642 | 1 | 0 | 3 | 0 | 0 | 0 |
| 541 | 0 | 0 | 0 | 0 | 0 | 0 |
| 588 | 0 | 1 | 1 | 0 | 0 | 1 |
| 488 | 1 | 2 | 0 | 0 | 0 | 3 |
| 234 | 0 | 1 | 1 | 0 | 0 | 0 |
| 165 | 0 | 0 | 0 | 0 | 0 | 0 |
| 6477 | 195 | | | | | |
| 573 | 4 | 3 | 3 | 1 | 1 | 1 |

| | | | | | | |
|---|---|---|---|---|---|---|
| 298 | 0 | 1 | 1 | 0 | 0 | 0 |
| 350 | 2 | 0 | 1 | 3 | 1 | 0 |
| 338 | 2 | 6 | 1 | 0 | 0 | 1 |
| 196 | 0 | 0 | 1 | 0 | 0 | 0 |
| 163 | 1 | 0 | 2 | 0 | 0 | 0 |
| 61 | 0 | 0 | 0 | 0 | 0 | 0 |
| 201 | 3 | 2 | 1 | 0 | 0 | 0 |
| 150 | 3 | 0 | 2 | 0 | 0 | 0 |
| 66 | 0 | 0 | 0 | 0 | 0 | 0 |
| 99 | 0 | 0 | 1 | 0 | 0 | 0 |
| 23 | 0 | 0 | 0 | 0 | 0 | 0 |
| 2518 | 82 | | | | | |
| 322 | 24 | 0 | 0 | 0 | 0 | 0 |
| 308 | 10 | 0 | 7 | 0 | 0 | 0 |
| 561 | 9 | 9 | 4 | 3 | 0 | 3 |
| 467 | 5 | 1 | 3 | 3 | 0 | 0 |
| 489 | 9 | 1 | 3 | 5 | 5 | 4 |
| 342 | 4 | 6 | 3 | 3 | 1 | 3 |
| 228 | 0 | 0 | 2 | 0 | 0 | 0 |
| 175 | 0 | 0 | 1 | 0 | 0 | 0 |
| 384 | 1 | 3 | 9 | 3 | 3 | 3 |
| 162 | 0 | 0 | 3 | 0 | 0 | 0 |
| 36 | 0 | 0 | 1 | 0 | 0 | 0 |
| 0 | 0 | 0 | 0 | 0 | 0 | 0 |
| 3474 | 210 | | | | | |
| 55 | 0 | 0 | 0 | 0 | 3 | 0 |
| 35 | 0 | 0 | 0 | 0 | 0 | 0 |
| 54 | 0 | 0 | 0 | 0 | 0 | 0 |
| 4 | 0 | 0 | 0 | 0 | 0 | 0 |
| 12 | 0 | 0 | 0 | 0 | 0 | 0 |
| 7 | 0 | 0 | 0 | 0 | 0 | 0 |
| 16 | 0 | 0 | 0 | 0 | 0 | 0 |
| 7 | 0 | 0 | 0 | 0 | 0 | 0 |
| 12 | 0 | 0 | 0 | 0 | 0 | 0 |
| 3 | 0 | 0 | 0 | 0 | 0 | 0 |
| 13 | 0 | 0 | 0 | 0 | 0 | 0 |
| 8 | 0 | 0 | 0 | 0 | 0 | 0 |
| 226 | 209 | | | | | |
| 331 | 2 | 1 | 1 | 0 | 3 | 1 |
| 325 | 2 | 1 | 3 | 0 | 0 | 1 |
| 270 | 0 | 0 | 0 | 0 | 0 | 0 |

| | | | | | | |
|---|---|---|---|---|---|---|
| 191 | 0 | 0 | 0 | 0 | 0 | 0 |
| 44 | 0 | 0 | 0 | 0 | 0 | 0 |
| 21 | 0 | 0 | 0 | 0 | 0 | 0 |
| 33 | 0 | 0 | 0 | 0 | 0 | 0 |
| 48 | 0 | 0 | 0 | 0 | 0 | 0 |
| 14 | 0 | 0 | 0 | 0 | 0 | 0 |
| 4 | 0 | 0 | 0 | 0 | 0 | 0 |
| 8 | 0 | 0 | 0 | 0 | 0 | 0 |
| 28 | 0 | 0 | 0 | 0 | 0 | 0 |
| 1317 | 48 | | | | | |
| 703 | 9 | 2 | 8 | 2 | 0 | 0 |
| 626 | 8 | 2 | 8 | 0 | 6 | 1 |
| 627 | 11 | 1 | 2 | 2 | 0 | 0 |
| 476 | 6 | 6 | 10 | 2 | 5 | 4 |
| 439 | 9 | 2 | 4 | 3 | 0 | 1 |
| 381 | 2 | 1 | 2 | 1 | 0 | 3 |
| 519 | 12 | 2 | 2 | 8 | 2 | 1 |
| 470 | 6 | 1 | 2 | 0 | 0 | 2 |
| 293 | 1 | 0 | 0 | 0 | 0 | 0 |
| 278 | 5 | 1 | 0 | 1 | 0 | 3 |
| 22 | 0 | 0 | 0 | 0 | 0 | 0 |
| 45 | 0 | 0 | 0 | 0 | 0 | 0 |
| 4879 | 263 | | | | | |
| 579 | 2 | 3 | 1 | 3 | 1 | 2 |
| 418 | 2 | 0 | 1 | 5 | 3 | 2 |
| 408 | 1 | 0 | 0 | 0 | 0 | 0 |
| 154 | 0 | 0 | 0 | 0 | 0 | 0 |
| 51 | 0 | 0 | 0 | 0 | 0 | 0 |
| 60 | 0 | 0 | 0 | 0 | 0 | 0 |
| 43 | 0 | 0 | 0 | 0 | 0 | 0 |
| 52 | 0 | 0 | 0 | 0 | 0 | 0 |
| 496 | 0 | 1 | 1 | 1 | 0 | 0 |
| 184 | 0 | 0 | 0 | 0 | 0 | 0 |
| 53 | 0 | 0 | 0 | 0 | 0 | 0 |
| 9 | 0 | 0 | 0 | 0 | 0 | 0 |
| 2507 | 50 | | | | | |
| 497 | 3 | 5 | 6 | 1 | 2 | 0 |
| 463 | 2 | 1 | 0 | 4 | 0 | 2 |
| 441 | 0 | 3 | 2 | 0 | 2 | 0 |
| 288 | 0 | 3 | 0 | 0 | 1 | 0 |
| 220 | 0 | 0 | 0 | 0 | 0 | 0 |

| | | | | | | |
|---|---|---|---|---|---|---|
| 99 | 0 | 0 | 1 | 0 | 0 | 0 |
| 138 | 0 | 0 | 0 | 0 | 0 | 2 |
| 399 | 8 | 10 | 13 | 5 | 3 | 7 |
| 200 | 0 | 0 | 3 | 0 | 0 | 1 |
| 97 | 0 | 0 | 2 | 0 | 0 | 0 |
| 191 | 0 | 0 | 3 | 1 | 0 | 0 |
| 90 | 0 | 0 | 1 | 0 | 0 | 0 |
| 3123 | 137 | | | | | |
| 747 | 25 | 0 | 9 | 4 | 0 | 2 |
| 555 | 6 | 10 | 13 | 1 | 4 | 0 |
| 581 | 11 | 9 | 9 | 2 | 4 | 3 |
| 450 | 8 | 2 | 6 | 1 | 2 | 0 |
| 448 | 7 | 2 | 7 | 3 | 0 | 1 |
| 404 | 9 | 6 | 2 | 0 | 0 | 0 |
| 441 | 7 | 2 | 5 | 3 | 1 | 1 |
| 441 | 0 | 0 | 1 | 0 | 0 | 2 |
| 219 | 0 | 0 | 0 | 2 | 0 | 0 |
| 213 | 0 | 0 | 1 | 0 | 0 | 0 |
| 199 | 0 | 0 | 0 | 0 | 0 | 0 |
| 248 | 5 | 0 | 2 | 2 | 0 | 0 |
| 4946 | 295 | | | | | |
| 579 | 4 | 7 | 4 | 4 | 2 | 1 |
| 532 | 9 | 2 | 0 | 0 | 0 | 2 |
| 504 | 2 | 1 | 2 | 2 | 2 | 0 |
| 458 | 5 | 0 | 1 | 4 | 0 | 2 |
| 302 | 1 | 0 | 0 | 0 | 0 | 0 |
| 162 | 0 | 0 | 0 | 0 | 0 | 0 |
| 134 | 0 | 0 | 0 | 0 | 0 | 0 |
| 113 | 2 | 0 | 0 | 0 | 0 | 0 |
| 9 | 0 | 0 | 0 | 0 | 0 | 0 |
| 152 | 0 | 0 | 0 | 0 | 0 | 0 |
| 136 | 0 | 0 | 0 | 0 | 0 | 0 |
| 9 | 0 | 0 | 0 | 0 | 0 | 0 |
| 3090 | 100 | | | | | |
| 594 | 1 | 6 | 4 | 2 | 2 | 2 |
| 444 | 1 | 1 | 5 | 1 | 1 | 1 |
| 345 | 4 | 0 | 8 | 0 | 2 | 2 |
| 415 | 0 | 6 | 8 | 0 | 0 | 3 |
| 288 | 0 | 0 | 2 | 1 | 2 | 0 |
| 145 | 1 | 0 | 1 | 0 | 0 | 0 |
| 243 | 0 | 0 | 1 | 0 | 0 | 0 |

| | | | | | | |
|---|---|---|---|---|---|---|
| 396 | 3 | 1 | 13 | 0 | 0 | 1 |
| 50 | 0 | 0 | 0 | 0 | 0 | 0 |
| 10 | 0 | 0 | 0 | 0 | 0 | 0 |
| 13 | 0 | 0 | 0 | 0 | 0 | 0 |
| 16 | 0 | 0 | 0 | 0 | 0 | 0 |
| 2959 | 113 | | | | | |
| 445 | 6 | 4 | 14 | 2 | 0 | 1 |
| 318 | 2 | 2 | 5 | 0 | 1 | 7 |
| 397 | 0 | 0 | 5 | 0 | 1 | 1 |
| 278 | 3 | 6 | 36 | 0 | 1 | 4 |
| 182 | 0 | 4 | 1 | 0 | 0 | 3 |
| 78 | 0 | 0 | 0 | 0 | 0 | 0 |
| 196 | 4 | 0 | 1 | 0 | 0 | 0 |
| 54 | 0 | 0 | 0 | 0 | 0 | 0 |
| 149 | 0 | 0 | 4 | 0 | 0 | 0 |
| 344 | 8 | 0 | 1 | 4 | 4 | 2 |
| 231 | 0 | 3 | 1 | 0 | 0 | 1 |
| 197 | 0 | 0 | 0 | 0 | 0 | 0 |
| 2869 | 204 | | | | | |
| 620 | 5 | 9 | 8 | 6 | 3 | 4 |
| 523 | 9 | 6 | 7 | 5 | 1 | 0 |
| 296 | 1 | 3 | 2 | 0 | 1 | 0 |
| 295 | 1 | 0 | 3 | 0 | 0 | 3 |
| 249 | 2 | 2 | 2 | 1 | 0 | 0 |
| 414 | 5 | 2 | 3 | 7 | 0 | 2 |
| 205 | 0 | 0 | 1 | 0 | 1 | 0 |
| 46 | 0 | 0 | 2 | 0 | 0 | 0 |
| 119 | 0 | 0 | 0 | 0 | 0 | 0 |
| 134 | 0 | 0 | 1 | 0 | 0 | 0 |
| 33 | 0 | 0 | 0 | 0 | 0 | 0 |
| 6 | 0 | 0 | 0 | 0 | 0 | 0 |
| 2940 | 159 | | | | | |
| 421 | 2 | 1 | 1 | 2 | 0 | 0 |
| 189 | 1 | 1 | 0 | 0 | 0 | 0 |
| 86 | 0 | 0 | 0 | 0 | 0 | 0 |
| 424 | 3 | 1 | 6 | 2 | 2 | 1 |
| 142 | 0 | 0 | 0 | 2 | 0 | 0 |
| 129 | 2 | 0 | 0 | 0 | 0 | 0 |
| 113 | 0 | 0 | 0 | 0 | 0 | 0 |
| 25 | 0 | 0 | 0 | 0 | 0 | 0 |
| 25 | 0 | 0 | 0 | 0 | 0 | 0 |

| | | | | | | |
|---|---|---|---|---|---|---|
| 76 | 0 | 0 | 0 | 0 | 0 | 0 |
| 62 | 0 | 0 | 0 | 0 | 0 | 0 |
| 5 | 0 | 0 | 0 | 0 | 0 | 0 |
| 1697 | 54 | | | | | |
| 779 | 4 | 1 | 10 | 2 | 6 | 3 |
| 721 | 1 | 7 | 8 | 3 | 4 | 2 |
| 513 | 0 | 2 | 4 | 0 | 0 | 6 |
| 564 | 4 | 3 | 5 | 4 | 1 | 3 |
| 354 | 1 | 0 | 6 | 0 | 4 | 2 |
| 463 | 3 | 2 | 3 | 2 | 1 | 1 |
| 256 | 0 | 3 | 8 | 1 | 0 | 0 |
| 366 | 4 | 2 | 21 | 4 | 0 | 4 |
| 190 | 0 | 0 | 2 | 0 | 2 | 0 |
| 175 | 1 | 2 | 6 | 0 | 0 | 1 |
| 82 | 0 | 0 | 1 | 0 | 0 | 0 |
| 25 | 0 | 0 | 0 | 0 | 0 | 0 |
| 4488 | 292 | | | | | |
| 905 | 7 | 3 | 6 | 7 | 5 | 2 |
| 791 | 8 | 4 | 4 | 4 | 2 | 3 |
| 758 | 6 | 4 | 2 | 3 | 3 | 3 |
| 598 | 4 | 3 | 3 | 3 | 4 | 1 |
| 531 | 5 | 3 | 5 | 6 | 3 | 5 |
| 543 | 2 | 5 | 4 | 3 | 4 | 1 |
| 644 | 7 | 2 | 2 | 3 | 0 | 2 |
| 558 | 4 | 3 | 7 | 4 | 2 | 1 |
| 297 | 3 | 5 | 1 | 0 | 2 | 0 |
| 529 | 1 | 1 | 1 | 2 | 0 | 0 |
| 530 | 0 | 1 | 1 | 3 | 1 | 1 |
| 334 | 2 | 2 | 1 | 0 | 1 | 2 |
| 7018 | 338 | | | | | |
| 888 | 4 | 5 | 10 | 6 | 4 | 5 |
| 676 | 2 | 7 | 3 | 3 | 2 | 2 |
| 660 | 3 | 2 | 7 | 8 | 6 | 2 |
| 621 | 1 | 1 | 4 | 4 | 2 | 2 |
| 567 | 4 | 5 | 5 | 2 | 4 | 7 |
| 716 | 1 | 3 | 2 | 2 | 2 | 3 |
| 613 | 1 | 1 | 6 | 4 | 2 | 1 |
| 592 | 1 | 4 | 4 | 1 | 3 | 4 |
| 531 | 1 | 0 | 2 | 4 | 2 | 4 |
| 454 | 7 | 3 | 1 | 1 | 2 | 2 |
| 515 | 4 | 2 | 2 | 3 | 2 | 2 |

| | | | | | | |
|---|---|---|---|---|---|---|
| 409 | 3 | 5 | 1 | 1 | 3 | 1 |
| 7242 | 351 | | | | | |
| 550 | 6 | 0 | 9 | 1 | 1 | 5 |
| 550 | 4 | 0 | 2 | 2 | 3 | 0 |
| 476 | 6 | 4 | 11 | 1 | 3 | 2 |
| 361 | 8 | 1 | 1 | 2 | 1 | 0 |
| 408 | 5 | 1 | 6 | 1 | 0 | 4 |
| 373 | 6 | 2 | 5 | 0 | 1 | 0 |
| 423 | 4 | 0 | 3 | 4 | 0 | 1 |
| 237 | 2 | 0 | 1 | 0 | 0 | 1 |
| 239 | 2 | 0 | 0 | 0 | 0 | 1 |
| 132 | 0 | 0 | 0 | 0 | 0 | 0 |
| 112 | 0 | 0 | 0 | 0 | 0 | 0 |
| 101 | 0 | 0 | 0 | 0 | 0 | 0 |
| 3962 | 213 | | | | | |
| 366 | 3 | 0 | 2 | 0 | 0 | 0 |
| 373 | 3 | 1 | 5 | 3 | 0 | 1 |
| 175 | 1 | 0 | 3 | 0 | 0 | 0 |
| 114 | 0 | 0 | 1 | 0 | 0 | 0 |
| 143 | 0 | 0 | 0 | 0 | 0 | 0 |
| 29 | 0 | 0 | 0 | 0 | 0 | 0 |
| 103 | 1 | 0 | 0 | 0 | 0 | 0 |
| 9 | 0 | 0 | 0 | 0 | 0 | 0 |
| 23 | 0 | 0 | 0 | 0 | 0 | 0 |
| 5 | 0 | 0 | 0 | 0 | 0 | 0 |
| 94 | 0 | 0 | 0 | 0 | 0 | 0 |
| 62 | 0 | 0 | 0 | 0 | 0 | 0 |
| 1496 | 54 | | | | | |
| 349 | 5 | 0 | 1 | 0 | 0 | 1 |
| 239 | 1 | 0 | 2 | 1 | 0 | 0 |
| 323 | 4 | 10 | 0 | 0 | 0 | 2 |
| 208 | 0 | 0 | 0 | 0 | 0 | 0 |
| 61 | 0 | 0 | 0 | 0 | 0 | 0 |
| 32 | 0 | 0 | 0 | 0 | 0 | 0 |
| 13 | 0 | 0 | 0 | 0 | 0 | 0 |
| 189 | 3 | 0 | 0 | 0 | 0 | 0 |
| 128 | 0 | 0 | 0 | 0 | 0 | 0 |
| 11 | 0 | 0 | 0 | 0 | 0 | 0 |
| 11 | 0 | 0 | 0 | 0 | 0 | 0 |
| 6 | 0 | 0 | 0 | 0 | 0 | 0 |
| 1570 | 13 | | | | | |

| | | | | | | |
|---|---|---|---|---|---|---|
| 549 | 26 | 4 | 7 | 13 | 0 | 5 |
| 477 | 6 | 3 | 12 | 1 | 3 | 2 |
| 517 | 10 | 2 | 3 | 7 | 0 | 3 |
| 349 | 5 | 1 | 3 | 5 | 0 | 0 |
| 376 | 6 | 1 | 2 | 2 | 1 | 0 |
| 238 | 5 | 0 | 1 | 2 | 0 | 1 |
| 114 | 0 | 0 | 0 | 0 | 0 | 0 |
| 48 | 0 | 0 | 0 | 0 | 0 | 0 |
| 17 | 0 | 0 | 0 | 0 | 0 | 0 |
| 39 | 0 | 0 | 0 | 0 | 0 | 0 |
| 85 | 0 | 0 | 0 | 0 | 0 | 0 |
| 49 | 0 | 0 | 0 | 0 | 0 | 0 |
| 2858 | 172 | | | | | |
| 536 | 4 | 2 | 2 | 2 | 0 | 0 |
| 353 | 4 | 1 | 3 | 0 | 0 | 0 |
| 212 | 0 | 0 | 0 | 0 | 0 | 0 |
| 141 | 2 | 0 | 0 | 0 | 0 | 0 |
| 123 | 2 | 0 | 1 | 0 | 0 | 2 |
| 29 | 0 | 0 | 0 | 0 | 0 | 0 |
| 24 | 0 | 0 | 0 | 0 | 0 | 0 |
| 200 | 0 | 0 | 0 | 0 | 0 | 0 |
| 129 | 0 | 0 | 0 | 1 | 0 | 0 |
| 197 | 0 | 0 | 0 | 0 | 0 | 0 |
| 199 | 0 | 0 | 0 | 0 | 0 | 0 |
| 93 | 0 | 0 | 0 | 0 | 0 | 0 |
| 2236 | 40 | | | | | |
| 344 | 0 | 0 | 1 | 0 | 0 | 0 |
| 375 | 6 | 6 | 2 | 1 | 0 | 2 |
| 175 | 0 | 0 | 0 | 0 | 0 | 0 |
| 81 | 0 | 0 | 0 | 0 | 0 | 0 |
| 18 | 0 | 0 | 0 | 0 | 0 | 0 |
| 9 | 0 | 0 | 0 | 0 | 0 | 0 |
| 441 | 7 | 6 | 4 | 1 | 0 | 0 |
| 219 | 0 | 0 | 0 | 0 | 0 | 3 |
| 58 | 0 | 0 | 0 | 0 | 0 | 0 |
| 7 | 0 | 0 | 0 | 0 | 0 | 0 |
| 2 | 0 | 0 | 0 | 0 | 0 | 0 |
| 18 | 0 | 0 | 0 | 0 | 0 | 0 |
| 1747 | 13 | | | | | |
| 911 | 4 | 4 | 4 | 2 | 1 | 3 |
| 745 | 6 | 2 | 5 | 1 | 1 | 1 |

| | | | | | | |
|---|---|---|---|---|---|---|
| 525 | 7 | 5 | 13 | 5 | 2 | 0 |
| 566 | 2 | 3 | 9 | 4 | 1 | 4 |
| 618 | 3 | 1 | 1 | 0 | 0 | 0 |
| 505 | 5 | 3 | 9 | 6 | 2 | 2 |
| 522 | 2 | 2 | 1 | 0 | 0 | 1 |
| 442 | 4 | 3 | 8 | 2 | 2 | 1 |
| 888 | 2 | 3 | 2 | 1 | 0 | 4 |
| 720 | 0 | 1 | 8 | 0 | 0 | 2 |
| 507 | 0 | 2 | 12 | 3 | 2 | 3 |
| 467 | 2 | 0 | 16 | 0 | 0 | 1 |
| 7416 | 333 | | | | | |
| 985 | 1 | 3 | 10 | 5 | 2 | 4 |
| 647 | 5 | 3 | 3 | 6 | 2 | 3 |
| 658 | 1 | 6 | 2 | 2 | 2 | 1 |
| 540 | 1 | 0 | 3 | 0 | 0 | 2 |
| 491 | 4 | 1 | 8 | 3 | 1 | 5 |
| 483 | 7 | 2 | 3 | 1 | 1 | 0 |
| 549 | 3 | 1 | 4 | 0 | 0 | 0 |
| 489 | 2 | 3 | 7 | 2 | 1 | 13 |
| 455 | 4 | 1 | 2 | 0 | 0 | 7 |
| 227 | 0 | 0 | 2 | 0 | 0 | 2 |
| 39 | 0 | 0 | 0 | 0 | 0 | 0 |
| 73 | 0 | 0 | 0 | 0 | 0 | 0 |
| 5636 | 220 | | | | | |
| 625 | 2 | 2 | 2 | 2 | 0 | 3 |
| 452 | 0 | 3 | 3 | 0 | 0 | 2 |
| 411 | 0 | 0 | 1 | 3 | 0 | 0 |
| 278 | 0 | 0 | 3 | 0 | 0 | 1 |
| 281 | 0 | 2 | 3 | 0 | 0 | 0 |
| 305 | 2 | 0 | 0 | 1 | 0 | 1 |
| 299 | 0 | 0 | 1 | 0 | 0 | 0 |
| 158 | 0 | 0 | 0 | 0 | 0 | 0 |
| 190 | 0 | 0 | 0 | 0 | 0 | 0 |
| 104 | 0 | 0 | 0 | 0 | 0 | 0 |
| 32 | 0 | 0 | 0 | 0 | 0 | 0 |
| 25 | 0 | 0 | 0 | 0 | 0 | 0 |
| 3160 | 80 | | | | | |
| 648 | 8 | 9 | 6 | 2 | 3 | 6 |
| 372 | 3 | 3 | 10 | 4 | 3 | 4 |
| 324 | 6 | 0 | 2 | 4 | 1 | 0 |
| 318 | 9 | 6 | 3 | 1 | 0 | 2 |

| | | | | | | |
|---|---|---|---|---|---|---|
| 388 | 9 | 5 | 6 | 0 | 1 | 3 |
| 242 | 0 | 0 | 0 | 0 | 0 | 1 |
| 211 | 6 | 0 | 0 | 1 | 0 | 0 |
| 85 | 0 | 0 | 0 | 0 | 0 | 0 |
| 49 | 0 | 0 | 0 | 0 | 0 | 0 |
| 11 | 0 | 0 | 0 | 0 | 0 | 0 |
| 65 | 0 | 0 | 0 | 0 | 0 | 0 |
| 105 | 3 | 0 | 0 | 0 | 0 | 0 |
| 2818 | 184 | | | | | |
| 750 | 2 | 3 | 3 | 4 | 1 | 1 |
| 725 | 8 | 4 | 3 | 3 | 2 | 3 |
| 675 | 1 | 1 | 2 | 3 | 3 | 1 |
| 679 | 3 | 4 | 2 | 2 | 0 | 1 |
| 655 | 6 | 0 | 3 | 2 | 3 | 1 |
| 484 | 1 | 2 | 1 | 0 | 0 | 2 |
| 460 | 2 | 0 | 0 | 0 | 0 | 0 |
| 490 | 0 | 2 | 2 | 1 | 0 | 3 |
| 189 | 3 | 0 | 0 | 0 | 0 | 0 |
| 284 | 3 | 0 | 1 | 0 | 0 | 0 |
| 135 | 1 | 0 | 0 | 0 | 0 | 0 |
| 113 | 3 | 0 | 0 | 0 | 0 | 0 |
| 5639 | 160 | | | | | |
| 630 | 1 | 1 | 7 | 3 | 2 | 0 |
| 481 | 3 | 0 | 2 | 2 | 2 | 3 |
| 339 | 9 | 4 | 2 | 2 | 0 | 0 |
| 237 | 0 | 0 | 0 | 0 | 0 | 0 |
| 275 | 5 | 0 | 2 | 0 | 0 | 0 |
| 224 | 1 | 0 | 1 | 0 | 0 | 0 |
| 158 | 1 | 0 | 1 | 1 | 0 | 0 |
| 161 | 4 | 0 | 2 | 0 | 0 | 0 |
| 119 | 0 | 0 | 0 | 0 | 0 | 0 |
| 54 | 0 | 0 | 0 | 0 | 0 | 0 |
| 82 | 0 | 0 | 0 | 0 | 0 | 0 |
| 24 | 0 | 0 | 0 | 0 | 0 | 0 |
| 2784 | 24 | | | | | |
| 748 | 19 | 8 | 10 | 4 | 1 | 4 |
| 542 | 5 | 2 | 5 | 3 | 1 | 1 |
| 504 | 7 | 5 | 5 | 5 | 2 | 1 |
| 517 | 9 | 6 | 3 | 2 | 1 | 1 |
| 289 | 2 | 1 | 1 | 0 | 0 | 2 |
| 312 | 2 | 0 | 2 | 1 | 0 | 0 |

| | | | | | | |
|---|---|---|---|---|---|---|
| 129 | 0 | 0 | 0 | 0 | 0 | 0 |
| 57 | 3 | 0 | 0 | 0 | 0 | 0 |
| 9 | 0 | 0 | 0 | 0 | 0 | 0 |
| 1 | 0 | 0 | 0 | 0 | 0 | 0 |
| 96 | 0 | 0 | 0 | 0 | 0 | 0 |
| 65 | 0 | 0 | 0 | 0 | 0 | 0 |
| 3269 | 186 | | | | | |
| 1046 | 5 | 2 | 13 | 3 | 1 | 1 |
| 880 | 3 | 6 | 7 | 2 | 4 | 4 |
| 707 | 5 | 4 | 4 | 0 | 3 | 2 |
| 803 | 2 | 2 | 3 | 2 | 1 | 0 |
| 633 | 1 | 2 | 2 | 1 | 1 | 0 |
| 698 | 2 | 2 | 3 | 3 | 1 | 0 |
| 828 | 6 | 3 | 2 | 0 | 2 | 2 |
| 681 | 7 | 1 | 2 | 0 | 0 | 3 |
| 483 | 0 | 1 | 0 | 0 | 0 | 0 |
| 403 | 5 | 1 | 0 | 2 | 0 | 1 |
| 528 | 7 | 0 | 1 | 1 | 1 | 0 |
| 245 | 0 | 1 | 0 | 0 | 0 | 0 |
| 7935 | 214 | | | | | |
| 698 | 0 | 6 | 5 | 2 | 1 | 1 |
| 543 | 4 | 4 | 2 | 3 | 1 | 2 |
| 489 | 7 | 0 | 6 | 2 | 0 | 2 |
| 448 | 4 | 1 | 2 | 3 | 1 | 2 |
| 484 | 4 | 5 | 3 | 2 | 1 | 6 |
| 462 | 3 | 2 | 2 | 1 | 1 | 0 |
| 442 | 4 | 0 | 2 | 0 | 0 | 0 |
| 547 | 1 | 0 | 0 | 0 | 2 | 0 |
| 337 | 8 | 0 | 0 | 3 | 0 | 0 |
| 253 | 2 | 0 | 0 | 1 | 0 | 1 |
| 257 | 1 | 0 | 0 | 0 | 0 | 0 |
| 126 | 0 | 0 | 0 | 0 | 0 | 0 |
| 5086 | 200 | | | | | |
| 117 | 13 | 0 | 0 | 0 | 1 | 0 |
| 88 | 8 | 0 | 0 | 0 | 0 | 0 |
| 71 | 16 | 0 | 0 | 0 | 0 | 0 |
| 54 | 13 | 0 | 0 | 0 | 0 | 0 |
| 122 | 8 | 3 | 0 | 0 | 0 | 0 |
| 140 | 6 | 0 | 0 | 0 | 0 | 0 |
| 35 | 12 | 0 | 0 | 0 | 0 | 0 |
| 120 | 3 | 0 | 0 | 0 | 0 | 2 |

| | | | | | | |
|---|---|---|---|---|---|---|
| 306 | 0 | 2 | 17 | 4 | 0 | 2 |
| 141 | 1 | 0 | 3 | 0 | 0 | 0 |
| 56 | 0 | 0 | 0 | 0 | 0 | 0 |
| 53 | 3 | 0 | 0 | 0 | 0 | 0 |
| 1303 | 127 | | | | | |
| 405 | 1 | 1 | 1 | 1 | 7 | 1 |
| 482 | 2 | 1 | 8 | 1 | 4 | 5 |
| 304 | 3 | 0 | 3 | 4 | 1 | 1 |
| 333 | 0 | 0 | 1 | 1 | 5 | 0 |
| 283 | 2 | 1 | 6 | 0 | 0 | 1 |
| 166 | 0 | 0 | 1 | 0 | 0 | 1 |
| 33 | 0 | 0 | 0 | 0 | 0 | 0 |
| 8 | 0 | 0 | 0 | 0 | 0 | 0 |
| 106 | 0 | 0 | 2 | 0 | 0 | 0 |
| 56 | 0 | 0 | 0 | 0 | 0 | 0 |
| 31 | 0 | 0 | 0 | 0 | 0 | 0 |
| 36 | 0 | 0 | 0 | 0 | 0 | 0 |
| 2243 | 112 | | | | | |
| 424 | 1 | 2 | 4 | 3 | 0 | 0 |
| 301 | 0 | 0 | 0 | 0 | 0 | 0 |
| 366 | 2 | 0 | 4 | 0 | 0 | 1 |
| 286 | 0 | 3 | 3 | 0 | 0 | 2 |
| 121 | 0 | 0 | 0 | 0 | 0 | 0 |
| 96 | 0 | 0 | 0 | 0 | 0 | 0 |
| 56 | 0 | 0 | 0 | 0 | 0 | 0 |
| 47 | 0 | 0 | 0 | 0 | 0 | 0 |
| 36 | 0 | 0 | 0 | 0 | 0 | 0 |
| 3 | 0 | 0 | 0 | 0 | 0 | 0 |
| 168 | 0 | 0 | 2 | 0 | 0 | 0 |
| 152 | 0 | 0 | 0 | 0 | 0 | 0 |
| 2056 | 3 | | | | | |
| 292 | 0 | 0 | 0 | 0 | 0 | 0 |
| 259 | 2 | 0 | 0 | 0 | 0 | 1 |
| 271 | 0 | 0 | 6 | 2 | 0 | 0 |
| 266 | 1 | 2 | 4 | 1 | 0 | 0 |
| 199 | 2 | 3 | 2 | 0 | 0 | 0 |
| 250 | 2 | 3 | 1 | 1 | 1 | 5 |
| 223 | 0 | 4 | 5 | 0 | 1 | 0 |
| 202 | 4 | 0 | 1 | 1 | 1 | 1 |
| 185 | 2 | 0 | 1 | 1 | 0 | 1 |
| 158 | 0 | 0 | 6 | 1 | 0 | 1 |

| | | | | | | |
|---|---|---|---|---|---|---|
| 116 | 0 | 0 | 0 | 0 | 0 | 0 |
| 170 | 0 | 1 | 0 | 1 | 0 | 0 |
| 2591 | 138 | | | | | |
| 267 | 0 | 1 | 1 | 1 | 0 | 0 |
| 263 | 0 | 0 | 4 | 4 | 3 | 4 |
| 248 | 1 | 1 | 3 | 2 | 3 | 1 |
| 200 | 1 | 0 | 0 | 0 | 0 | 0 |
| 256 | 0 | 3 | 4 | 1 | 1 | 1 |
| 206 | 1 | 0 | 0 | 0 | 0 | 1 |
| 139 | 0 | 0 | 1 | 0 | 0 | 3 |
| 244 | 1 | 4 | 8 | 1 | 0 | 0 |
| 165 | 0 | 0 | 0 | 0 | 0 | 0 |
| 176 | 1 | 0 | 1 | 2 | 0 | 0 |
| 139 | 11 | 2 | 0 | 0 | 0 | 0 |
| 203 | 0 | 0 | 4 | 3 | 1 | 0 |
| 2506 | 134 | | | | | |
| 315 | 2 | 0 | 2 | 2 | 0 | 0 |
| 259 | 0 | 0 | 2 | 5 | 1 | 0 |
| 252 | 2 | 0 | 5 | 0 | 0 | 3 |
| 158 | 6 | 0 | 8 | 0 | 1 | 0 |
| 177 | 0 | 1 | 0 | 0 | 0 | 0 |
| 195 | 1 | 1 | 0 | 0 | 1 | 1 |
| 129 | 0 | 0 | 1 | 0 | 0 | 0 |
| 116 | 0 | 0 | 8 | 0 | 0 | 0 |
| 102 | 0 | 0 | 3 | 0 | 0 | 0 |
| 20 | 0 | 0 | 3 | 0 | 0 | 0 |
| 69 | 1 | 0 | 0 | 0 | 0 | 0 |
| 264 | 3 | 1 | 1 | 2 | 2 | 0 |
| 2056 | 104 | | | | | |
| 733 | 10 | 2 | 6 | 3 | 1 | 3 |
| 549 | 3 | 2 | 6 | 2 | 5 | 1 |
| 405 | 0 | 3 | 2 | 0 | 0 | 5 |
| 374 | 6 | 1 | 2 | 5 | 0 | 0 |
| 360 | 1 | 2 | 2 | 0 | 0 | 0 |
| 291 | 1 | 1 | 2 | 0 | 0 | 0 |
| 171 | 0 | 0 | 1 | 0 | 0 | 0 |
| 83 | 0 | 0 | 0 | 0 | 0 | 0 |
| 36 | 0 | 0 | 0 | 0 | 0 | 0 |
| 9 | 0 | 0 | 0 | 0 | 0 | 0 |
| 171 | 0 | 0 | 0 | 0 | 0 | 0 |
| 38 | 0 | 0 | 0 | 0 | 0 | 0 |

| | | | | | | |
|---|---|---|---|---|---|---|
| 3220 | 129 | | | | | |
| 718 | 7 | 2 | 4 | 0 | 1 | 1 |
| 615 | 5 | 4 | 2 | 3 | 0 | 0 |
| 423 | 0 | 1 | 2 | 0 | 0 | 0 |
| 364 | 0 | 0 | 1 | 0 | 0 | 0 |
| 243 | 7 | 0 | 0 | 0 | 0 | 0 |
| 125 | 1 | 0 | 0 | 0 | 0 | 0 |
| 86 | 0 | 0 | 0 | 0 | 0 | 0 |
| 270 | 0 | 0 | 0 | 0 | 0 | 0 |
| 446 | 0 | 0 | 5 | 1 | 3 | 3 |
| 183 | 1 | 0 | 0 | 0 | 0 | 0 |
| 68 | 2 | 0 | 0 | 0 | 0 | 0 |
| 22 | 0 | 0 | 0 | 0 | 0 | 0 |
| 3563 | 91 | | | | | |
| 387 | 2 | 0 | 4 | 1 | 0 | 1 |
| 346 | 2 | 0 | 1 | 0 | 0 | 0 |
| 280 | 3 | 0 | 0 | 0 | 0 | 0 |
| 342 | 1 | 2 | 2 | 2 | 0 | 1 |
| 214 | 0 | 0 | 0 | 0 | 0 | 0 |
| 240 | 2 | 0 | 3 | 0 | 0 | 0 |
| 212 | 0 | 0 | 0 | 0 | 0 | 0 |
| 104 | 0 | 0 | 0 | 0 | 0 | 0 |
| 119 | 0 | 0 | 0 | 0 | 0 | 0 |
| 106 | 0 | 0 | 0 | 0 | 0 | 0 |
| 45 | 0 | 0 | 0 | 0 | 0 | 0 |
| 17 | 0 | 0 | 0 | 0 | 0 | 0 |
| 2412 | 46 | | | | | |
| 1074 | 9 | 6 | 5 | 4 | 8 | 5 |
| 807 | 15 | 0 | 12 | 2 | 2 | 2 |
| 769 | 5 | 4 | 5 | 1 | 0 | 5 |
| 625 | 0 | 2 | 5 | 1 | 0 | 2 |
| 659 | 4 | 3 | 5 | 0 | 3 | 0 |
| 391 | 0 | 2 | 9 | 0 | 0 | 0 |
| 423 | 3 | 0 | 13 | 1 | 4 | 6 |
| 516 | 1 | 0 | 9 | 0 | 0 | 2 |
| 409 | 2 | 5 | 3 | 0 | 0 | 0 |
| 248 | 0 | 0 | 0 | 0 | 0 | 0 |
| 264 | 0 | 0 | 3 | 0 | 0 | 0 |
| 542 | 0 | 3 | 12 | 0 | 1 | 0 |
| 6727 | 280 | | | | | |
| 756 | 1 | 1 | 6 | 2 | 0 | 1 |

| | | | | | | |
|---|---|---|---|---|---|---|
| 734 | 2 | 1 | 6 | 0 | 1 | 0 |
| 525 | 9 | 0 | 1 | 1 | 0 | 1 |
| 311 | 0 | 0 | 5 | 0 | 0 | 2 |
| 182 | 0 | 0 | 3 | 0 | 0 | 0 |
| 68 | 0 | 0 | 0 | 0 | 0 | 0 |
| 59 | 0 | 0 | 0 | 0 | 0 | 0 |
| 246 | 0 | 0 | 1 | 0 | 0 | 0 |
| 206 | 0 | 0 | 0 | 0 | 0 | 0 |
| 36 | 0 | 0 | 0 | 0 | 0 | 0 |
| 29 | 0 | 0 | 0 | 0 | 0 | 0 |
| 45 | 0 | 0 | 0 | 0 | 0 | 0 |
| 3197 | 58 | | | | | |
| 551 | 5 | 0 | 1 | 2 | 0 | 1 |
| 453 | 0 | 0 | 2 | 0 | 0 | 0 |
| 455 | 0 | 2 | 2 | 0 | 0 | 1 |
| 267 | 2 | 0 | 0 | 0 | 0 | 0 |
| 188 | 0 | 0 | 0 | 0 | 0 | 0 |
| 59 | 0 | 0 | 0 | 0 | 0 | 0 |
| 39 | 0 | 0 | 0 | 0 | 0 | 0 |
| 16 | 0 | 0 | 0 | 0 | 0 | 0 |
| 32 | 0 | 0 | 0 | 0 | 0 | 0 |
| 8 | 0 | 0 | 0 | 0 | 0 | 0 |
| 0 | 0 | 0 | 0 | 0 | 0 | 0 |
| 15 | 0 | 0 | 0 | 0 | 0 | 0 |
| 2083 | 35 | | | | | |
| 581 | 6 | 7 | 15 | 0 | 0 | 2 |
| 523 | 4 | 1 | 6 | 1 | 3 | 2 |
| 409 | 4 | 4 | 14 | 4 | 2 | 2 |
| 335 | 2 | 2 | 5 | 1 | 1 | 0 |
| 268 | 3 | 1 | 0 | 3 | 0 | 2 |
| 327 | 1 | 1 | 6 | 0 | 0 | 3 |
| 164 | 0 | 0 | 1 | 0 | 0 | 0 |
| 232 | 0 | 0 | 8 | 0 | 0 | 0 |
| 143 | 0 | 0 | 0 | 0 | 0 | 0 |
| 32 | 0 | 0 | 0 | 0 | 0 | 0 |
| 9 | 0 | 0 | 0 | 0 | 0 | 0 |
| 383 | 0 | 0 | 3 | 0 | 0 | 0 |
| 3406 | 193 | | | | | |
| 699 | 5 | 1 | 2 | 0 | 2 | 0 |
| 620 | 1 | 1 | 2 | 1 | 1 | 0 |
| 474 | 0 | 0 | 6 | 0 | 2 | 1 |

| | | | | | | |
|---|---|---|---|---|---|---|
| 319 | 0 | 2 | 3 | 0 | 0 | 1 |
| 332 | 0 | 1 | 1 | 0 | 0 | 0 |
| 391 | 1 | 0 | 2 | 0 | 1 | 1 |
| 233 | 0 | 0 | 1 | 0 | 0 | 0 |
| 239 | 0 | 0 | 0 | 0 | 0 | 0 |
| 139 | 0 | 0 | 0 | 0 | 0 | 0 |
| 37 | 0 | 0 | 0 | 0 | 0 | 0 |
| 32 | 0 | 0 | 1 | 0 | 0 | 0 |
| 42 | 0 | 0 | 0 | 0 | 0 | 0 |
| 3557 | 54 | | | | | |
| 546 | 1 | 1 | 2 | 0 | 0 | 1 |
| 417 | 1 | 0 | 3 | 2 | 1 | 0 |
| 222 | 0 | 4 | 0 | 3 | 0 | 0 |
| 155 | 0 | 0 | 1 | 0 | 0 | 0 |
| 158 | 0 | 0 | 0 | 0 | 0 | 0 |
| 66 | 0 | 0 | 0 | 0 | 0 | 0 |
| 105 | 3 | 0 | 0 | 0 | 0 | 0 |
| 49 | 0 | 0 | 0 | 0 | 0 | 0 |
| 41 | 1 | 0 | 0 | 0 | 0 | 0 |
| 71 | 0 | 0 | 0 | 0 | 0 | 0 |
| 37 | 0 | 0 | 0 | 0 | 0 | 0 |
| 39 | 0 | 0 | 0 | 0 | 0 | 0 |
| 1906 | 41 | | | | | |
| 540 | 14 | 2 | 4 | 3 | 1 | 1 |
| 360 | 4 | 0 | 3 | 0 | 0 | 1 |
| 417 | 2 | 1 | 8 | 5 | 7 | 3 |
| 376 | 1 | 0 | 4 | 1 | 0 | 5 |
| 419 | 1 | 1 | 1 | 0 | 0 | 1 |
| 252 | 4 | 2 | 5 | 2 | 4 | 0 |
| 228 | 2 | 0 | 1 | 1 | 0 | 0 |
| 203 | 1 | 0 | 2 | 0 | 0 | 1 |
| 106 | 0 | 0 | 0 | 0 | 1 | 0 |
| 0 | 0 | 0 | 0 | 0 | 0 | 0 |
| 4 | 0 | 0 | 0 | 0 | 0 | 0 |
| 105 | 0 | 0 | 1 | 0 | 0 | 0 |
| 3010 | 187 | | | | | |
| 612 | 2 | 6 | 4 | 3 | 1 | 1 |
| 512 | 3 | 1 | 2 | 0 | 0 | 2 |
| 392 | 2 | 1 | 2 | 1 | 1 | 1 |
| 339 | 3 | 0 | 4 | 0 | 0 | 0 |
| 333 | 2 | 1 | 5 | 2 | 0 | 2 |

| | | | | | | |
|---|---|---|---|---|---|---|
| 239 | 0 | 2 | 2 | 0 | 0 | 0 |
| 305 | 1 | 1 | 0 | 0 | 0 | 0 |
| 230 | 0 | 0 | 0 | 0 | 1 | 1 |
| 131 | 0 | 0 | 0 | 1 | 0 | 0 |
| 46 | 0 | 0 | 0 | 0 | 0 | 0 |
| 1 | 0 | 0 | 0 | 0 | 0 | 0 |
| 70 | 0 | 0 | 0 | 0 | 0 | 0 |
| 3210 | 13 | | | | | |
| 443 | 2 | 1 | 1 | 0 | 0 | 1 |
| 176 | 0 | 0 | 0 | 1 | 0 | 0 |
| 101 | 0 | 0 | 0 | 0 | 0 | 0 |
| 53 | 0 | 0 | 0 | 0 | 0 | 0 |
| 53 | 0 | 0 | 0 | 0 | 0 | 0 |
| 22 | 0 | 0 | 0 | 0 | 0 | 0 |
| 227 | 0 | 2 | 0 | 1 | 0 | 0 |
| 213 | 1 | 0 | 1 | 0 | 0 | 0 |
| 37 | 0 | 0 | 0 | 0 | 0 | 0 |
| 5 | 0 | 0 | 0 | 0 | 0 | 0 |
| 1 | 0 | 0 | 0 | 0 | 0 | 0 |
| 15 | 0 | 0 | 1 | 0 | 0 | 0 |
| 1346 | 3 | | | | | |
| 794 | 7 | 5 | 5 | 8 | 0 | 0 |
| 564 | 0 | 0 | 4 | 1 | 0 | 1 |
| 471 | 15 | 5 | 6 | 3 | 0 | 3 |
| 398 | 4 | 1 | 1 | 1 | 4 | 0 |
| 310 | 1 | 1 | 0 | 1 | 0 | 4 |
| 428 | 1 | 0 | 1 | 0 | 1 | 4 |
| 376 | 2 | 4 | 2 | 4 | 3 | 0 |
| 116 | 0 | 0 | 0 | 0 | 0 | 0 |
| 53 | 0 | 0 | 1 | 0 | 0 | 0 |
| 3 | 0 | 0 | 0 | 0 | 0 | 0 |
| 1 | 0 | 0 | 0 | 0 | 0 | 0 |
| 117 | 0 | 0 | 0 | 0 | 0 | 0 |
| 3631 | 30 | | | | | |
| 521 | 4 | 0 | 2 | 4 | 1 | 0 |
| 475 | 2 | 1 | 5 | 1 | 0 | 6 |
| 349 | 1 | 1 | 0 | 1 | 0 | 0 |
| 248 | 0 | 0 | 0 | 0 | 0 | 0 |
| 162 | 3 | 0 | 0 | 0 | 0 | 0 |
| 48 | 0 | 0 | 0 | 0 | 0 | 0 |
| 108 | 0 | 0 | 0 | 0 | 0 | 0 |

| | | | | | | |
|---|---|---|---|---|---|---|
| 43 | 0 | 0 | 0 | 0 | 0 | 0 |
| 4 | 0 | 0 | 0 | 0 | 0 | 0 |
| 28 | 0 | 0 | 0 | 0 | 0 | 0 |
| 8 | 0 | 0 | 0 | 0 | 0 | 0 |
| 32 | 0 | 0 | 0 | 0 | 0 | 0 |
| 2026 | 10 | | | | | |
| 240 | 0 | 0 | 1 | 0 | 2 | 0 |
| 247 | 2 | 1 | 0 | 0 | 0 | 0 |
| 114 | 0 | 0 | 0 | 0 | 0 | 0 |
| 111 | 0 | 0 | 0 | 0 | 0 | 0 |
| 161 | 2 | 2 | 0 | 2 | 0 | 1 |
| 158 | 0 | 0 | 0 | 0 | 0 | 0 |
| 112 | 0 | 0 | 0 | 0 | 0 | 0 |
| 304 | 2 | 0 | 0 | 0 | 3 | 2 |
| 248 | 0 | 0 | 2 | 0 | 0 | 0 |
| 116 | 0 | 0 | 3 | 2 | 1 | 0 |
| 158 | 0 | 2 | 3 | 2 | 0 | 0 |
| 34 | 0 | 0 | 0 | 0 | 0 | 0 |
| 2003 | 6 | | | | | |
| 652 | 10 | 2 | 7 | 1 | 0 | 1 |
| 618 | 5 | 3 | 11 | 1 | 0 | 3 |
| 531 | 5 | 3 | 9 | 2 | 4 | 4 |
| 411 | 7 | 0 | 8 | 3 | 0 | 0 |
| 336 | 5 | 2 | 6 | 2 | 1 | 2 |
| 321 | 12 | 0 | 0 | 1 | 0 | 1 |
| 306 | 2 | 2 | 1 | 0 | 0 | 1 |
| 253 | 7 | 2 | 0 | 0 | 3 | 0 |
| 201 | 0 | 1 | 0 | 0 | 0 | 0 |
| 35 | 0 | 0 | 0 | 0 | 0 | 0 |
| 80 | 3 | 0 | 0 | 0 | 0 | 0 |
| 276 | 2 | 0 | 0 | 0 | 0 | 0 |
| 4020 | 58 | | | | | |
| 428 | 1 | 1 | 0 | 0 | 1 | 0 |
| 422 | 0 | 3 | 2 | 3 | 0 | 1 |
| 288 | 4 | 0 | 0 | 0 | 0 | 0 |
| 229 | 0 | 0 | 0 | 0 | 0 | 0 |
| 132 | 1 | 0 | 0 | 0 | 0 | 0 |
| 81 | 0 | 0 | 0 | 0 | 0 | 0 |
| 355 | 6 | 2 | 4 | 0 | 0 | 2 |
| 305 | 0 | 0 | 1 | 0 | 0 | 1 |
| 119 | 2 | 0 | 0 | 0 | 0 | 0 |

| | | | | | | |
|---|---|---|---|---|---|---|
| 44 | 0 | 0 | 0 | 0 | 0 | 0 |
| 11 | 0 | 0 | 0 | 0 | 0 | 0 |
| 31 | 0 | 0 | 0 | 0 | 0 | 0 |
| 2445 | 14 | | | | | |
| 823 | 8 | 4 | 20 | 0 | 3 | 6 |
| 571 | 3 | 3 | 12 | 6 | 6 | 3 |
| 536 | 1 | 0 | 5 | 0 | 3 | 7 |
| 511 | 6 | 2 | 5 | 1 | 3 | 2 |
| 460 | 0 | 2 | 6 | 1 | 4 | 1 |
| 390 | 5 | 1 | 15 | 3 | 2 | 3 |
| 467 | 6 | 2 | 8 | 6 | 3 | 2 |
| 482 | 4 | 3 | 11 | 0 | 2 | 5 |
| 345 | 7 | 0 | 11 | 1 | 3 | 3 |
| 263 | 6 | 1 | 4 | 3 | 0 | 4 |
| 249 | 2 | 1 | 7 | 0 | 0 | 0 |
| 39 | 0 | 0 | 0 | 0 | 0 | 0 |
| 5136 | 48 | | | | | |
| 482 | 4 | 1 | 7 | 2 | 1 | 2 |
| 326 | 1 | 0 | 0 | 2 | 0 | 0 |
| 209 | 0 | 0 | 0 | 0 | 0 | 0 |
| 247 | 2 | 2 | 11 | 1 | 0 | 3 |
| 59 | 0 | 0 | 4 | 0 | 0 | 0 |
| 211 | 1 | 0 | 1 | 0 | 0 | 0 |
| 79 | 0 | 0 | 2 | 0 | 0 | 0 |
| 199 | 1 | 1 | 1 | 0 | 0 | 1 |
| 93 | 1 | 0 | 0 | 0 | 0 | 0 |
| 66 | 0 | 0 | 3 | 0 | 0 | 0 |
| 150 | 3 | 0 | 0 | 2 | 3 | 0 |
| 218 | 2 | 1 | 5 | 2 | 4 | 2 |
| 2339 | 15 | | | | | |
| 986 | 6 | 4 | 4 | 1 | 1 | 1 |
| 802 | 5 | 1 | 1 | 0 | 4 | 1 |
| 718 | 5 | 3 | 4 | 6 | 0 | 1 |
| 708 | 0 | 2 | 2 | 2 | 3 | 2 |
| 698 | 4 | 0 | 2 | 2 | 1 | 0 |
| 647 | 1 | 1 | 3 | 0 | 0 | 0 |
| 462 | 5 | 4 | 1 | 1 | 3 | 2 |
| 504 | 0 | 0 | 0 | 0 | 0 | 1 |
| 160 | 4 | 0 | 3 | 0 | 0 | 0 |
| 535 | 4 | 2 | 1 | 0 | 0 | 0 |
| 193 | 0 | 0 | 2 | 0 | 0 | 0 |

| | | | | | | |
|---|---|---|---|---|---|---|
| 25 | 0 | 0 | 0 | 0 | 0 | 0 |
| 6438 | 181 | | | | | |
| 845 | 3 | 3 | 4 | 1 | 0 | 1 |
| 679 | 0 | 0 | 3 | 0 | 2 | 0 |
| 395 | 3 | 0 | 1 | 0 | 0 | 2 |
| 511 | 2 | 1 | 3 | 5 | 0 | 0 |
| 235 | 0 | 0 | 1 | 0 | 0 | 0 |
| 98 | 0 | 0 | 0 | 0 | 0 | 0 |
| 132 | 0 | 0 | 0 | 0 | 0 | 0 |
| 72 | 0 | 0 | 0 | 0 | 0 | 0 |
| 0 | 0 | 0 | 0 | 0 | 0 | 0 |
| 7 | 0 | 0 | 0 | 0 | 0 | 0 |
| 44 | 0 | 0 | 0 | 0 | 0 | 0 |
| 64 | 0 | 0 | 0 | 0 | 0 | 0 |
| 3082 | 61 | | | | | |
| 655 | 2 | 0 | 3 | 0 | 0 | 0 |
| 310 | 2 | 0 | 4 | 0 | 0 | 0 |
| 294 | 0 | 2 | 0 | 0 | 0 | 1 |
| 86 | 0 | 0 | 0 | 0 | 0 | 0 |
| 47 | 0 | 0 | 0 | 0 | 0 | 0 |
| 52 | 3 | 0 | 0 | 0 | 0 | 0 |
| 48 | 0 | 0 | 0 | 0 | 0 | 0 |
| 55 | 3 | 0 | 0 | 0 | 0 | 0 |
| 46 | 0 | 0 | 0 | 0 | 0 | 0 |
| 43 | 0 | 0 | 0 | 0 | 0 | 0 |
| 46 | 0 | 0 | 0 | 0 | 0 | 0 |
| 20 | 0 | 0 | 0 | 0 | 0 | 0 |
| 1702 | 24 | | | | | |
| 695 | 7 | 3 | 4 | 1 | 0 | 2 |
| 609 | 6 | 8 | 11 | 1 | 0 | 0 |
| 573 | 13 | 3 | 6 | 1 | 0 | 1 |
| 394 | 7 | 1 | 4 | 3 | 4 | 3 |
| 427 | 5 | 0 | 0 | 1 | 1 | 0 |
| 311 | 0 | 1 | 0 | 1 | 1 | 0 |
| 212 | 0 | 0 | 0 | 1 | 0 | 0 |
| 374 | 5 | 5 | 2 | 1 | 2 | 4 |
| 135 | 0 | 0 | 0 | 0 | 0 | 0 |
| 278 | 0 | 0 | 1 | 0 | 0 | 0 |
| 132 | 0 | 0 | 0 | 0 | 0 | 0 |
| 2 | 0 | 0 | 0 | 0 | 0 | 0 |
| 4142 | 215 | | | | | |

| | | | | | | |
|---|---|---|---|---|---|---|
| 590 | 2 | 1 | 3 | 1 | 0 | 1 |
| 466 | 2 | 0 | 0 | 1 | 0 | 0 |
| 543 | 5 | 2 | 5 | 3 | 2 | 1 |
| 324 | 1 | 1 | 2 | 0 | 0 | 2 |
| 229 | 0 | 0 | 2 | 0 | 0 | 0 |
| 74 | 0 | 0 | 0 | 0 | 0 | 0 |
| 22 | 0 | 0 | 0 | 0 | 0 | 0 |
| 23 | 0 | 0 | 0 | 0 | 0 | 0 |
| 0 | 0 | 0 | 0 | 0 | 0 | 0 |
| 82 | 0 | 0 | 0 | 0 | 0 | 0 |
| 13 | 0 | 0 | 0 | 0 | 0 | 0 |
| 7 | 0 | 0 | 0 | 0 | 0 | 0 |
| 2373 | 78 | | | | | |
| 642 | 3 | 4 | 3 | 0 | 0 | 4 |
| 613 | 4 | 3 | 7 | 3 | 0 | 1 |
| 432 | 0 | 0 | 5 | 0 | 1 | 2 |
| 387 | 0 | 1 | 3 | 2 | 2 | 0 |
| 363 | 0 | 0 | 2 | 0 | 0 | 0 |
| 130 | 0 | 0 | 0 | 0 | 0 | 0 |
| 158 | 0 | 0 | 0 | 1 | 0 | 0 |
| 30 | 0 | 0 | 0 | 0 | 0 | 0 |
| 27 | 0 | 0 | 0 | 0 | 0 | 0 |
| 18 | 0 | 0 | 0 | 0 | 0 | 0 |
| 9 | 0 | 0 | 0 | 0 | 0 | 0 |
| 0 | 0 | 0 | 0 | 0 | 0 | 0 |
| 2809 | 90 | | | | | |
| 69 | 0 | 0 | 0 | 0 | 0 | 0 |
| 11 | 0 | 0 | 0 | 0 | 0 | 0 |
| 3 | 0 | 0 | 0 | 0 | 0 | 0 |
| 7 | 0 | 0 | 0 | 0 | 0 | 0 |
| 4 | 0 | 0 | 0 | 0 | 0 | 0 |
| 1 | 0 | 0 | 0 | 0 | 0 | 0 |
| 2 | 0 | 0 | 0 | 0 | 0 | 0 |
| 2 | 0 | 0 | 0 | 0 | 0 | 0 |
| 1 | 0 | 0 | 0 | 0 | 0 | 0 |
| 0 | 0 | 0 | 0 | 0 | 0 | 0 |
| 1 | 0 | 0 | 0 | 0 | 0 | 0 |
| 0 | 0 | 0 | 0 | 0 | 0 | 0 |
| 101 | 0 | | | | | |
| 201 | 0 | 0 | 0 | 0 | 0 | 0 |
| 96 | 0 | 0 | 0 | 0 | 0 | 0 |

| | | | | | | |
|---|---|---|---|---|---|---|
| 192 | 0 | 0 | 0 | 0 | 0 | 0 |
| 114 | 0 | 0 | 0 | 0 | 0 | 0 |
| 64 | 0 | 0 | 0 | 0 | 0 | 0 |
| 191 | 7 | 0 | 0 | 6 | 0 | 0 |
| 289 | 3 | 3 | 8 | 2 | 0 | 2 |
| 116 | 1 | 0 | 2 | 0 | 0 | 0 |
| 103 | 0 | 0 | 0 | 0 | 0 | 0 |
| 159 | 0 | 0 | 1 | 0 | 0 | 0 |
| 201 | 0 | 0 | 0 | 1 | 0 | 0 |
| 125 | 0 | 0 | 0 | 0 | 0 | 0 |
| 1851 | 11 | | | | | |
| 1213 | 7 | 4 | 6 | 2 | 2 | 4 |
| 855 | 9 | 2 | 3 | 2 | 8 | 3 |
| 801 | 4 | 7 | 9 | 2 | 5 | 4 |
| 624 | 9 | 7 | 3 | 3 | 3 | 1 |
| 688 | 8 | 6 | 4 | 4 | 4 | 3 |
| 637 | 5 | 3 | 2 | 2 | 1 | 2 |
| 499 | 1 | 0 | 1 | 0 | 1 | 3 |
| 552 | 1 | 0 | 0 | 0 | 0 | 0 |
| 453 | 2 | 0 | 2 | 0 | 0 | 0 |
| 137 | 0 | 0 | 0 | 0 | 0 | 1 |
| 30 | 0 | 0 | 0 | 0 | 0 | 0 |
| 5 | 0 | 0 | 0 | 0 | 0 | 0 |
| 6494 | 46 | | | | | |
| 840 | 4 | 3 | 7 | 2 | 4 | 2 |
| 783 | 4 | 0 | 1 | 2 | 0 | 2 |
| 574 | 2 | 2 | 7 | 0 | 1 | 1 |
| 430 | 1 | 1 | 6 | 1 | 0 | 0 |
| 423 | 1 | 2 | 6 | 2 | 0 | 1 |
| 158 | 0 | 0 | 1 | 0 | 0 | 0 |
| 112 | 0 | 0 | 0 | 0 | 0 | 0 |
| 99 | 0 | 0 | 1 | 0 | 0 | 1 |
| 111 | 0 | 0 | 0 | 0 | 0 | 0 |
| 0 | 0 | 0 | 0 | 0 | 0 | 0 |
| 18 | 0 | 0 | 0 | 0 | 0 | 0 |
| 0 | 0 | 0 | 0 | 0 | 0 | 0 |
| 3548 | 12 | | | | | |
| 660 | 2 | 2 | 2 | 1 | 0 | 0 |
| 378 | 1 | 0 | 0 | 0 | 0 | 0 |
| 348 | 0 | 2 | 0 | 0 | 0 | 0 |
| 170 | 1 | 0 | 0 | 0 | 0 | 0 |

| | | | | | | |
|---|---|---|---|---|---|---|
| 37 | 0 | 0 | 0 | 0 | 0 | 0 |
| 205 | 0 | 0 | 0 | 0 | 0 | 0 |
| 101 | 0 | 0 | 0 | 0 | 0 | 0 |
| 53 | 0 | 0 | 0 | 0 | 0 | 0 |
| 0 | 0 | 0 | 0 | 0 | 0 | 0 |
| 5 | 0 | 0 | 0 | 0 | 0 | 0 |
| 136 | 0 | 0 | 0 | 0 | 0 | 0 |
| 266 | 0 | 0 | 0 | 1 | 0 | 0 |
| 2359 | 4 | | | | | |
| 380 | 2 | 0 | 20 | 1 | 0 | 1 |
| 295 | 0 | 0 | 0 | 1 | 0 | 2 |
| 249 | 0 | 0 | 3 | 0 | 0 | 0 |
| 128 | 0 | 0 | 13 | 0 | 0 | 1 |
| 208 | 3 | 3 | 16 | 2 | 0 | 2 |
| 72 | 0 | 0 | 0 | 0 | 0 | 0 |
| 107 | 5 | 2 | 2 | 2 | 0 | 0 |
| 83 | 0 | 0 | 0 | 0 | 0 | 0 |
| 30 | 0 | 0 | 0 | 0 | 0 | 0 |
| 17 | 0 | 0 | 0 | 0 | 0 | 0 |
| 32 | 0 | 0 | 0 | 0 | 0 | 0 |
| 19 | 0 | 0 | 0 | 0 | 0 | 0 |
| 1620 | 171 | | | | | |
| 406 | 3 | 1 | 1 | 1 | 0 | 1 |
| 257 | 0 | 0 | 0 | 0 | 0 | 0 |
| 151 | 0 | 0 | 0 | 2 | 0 | 0 |
| 71 | 0 | 0 | 0 | 0 | 0 | 0 |
| 28 | 0 | 0 | 0 | 0 | 0 | 0 |
| 41 | 0 | 0 | 0 | 0 | 0 | 0 |
| 63 | 0 | 0 | 0 | 0 | 0 | 0 |
| 176 | 2 | 0 | 2 | 0 | 0 | 0 |
| 23 | 0 | 0 | 0 | 0 | 0 | 0 |
| 15 | 0 | 0 | 0 | 0 | 0 | 0 |
| 10 | 0 | 0 | 0 | 0 | 0 | 0 |
| 19 | 0 | 0 | 0 | 0 | 0 | 0 |
| 1260 | 36 | | | | | |
| 275 | 1 | 0 | 3 | 0 | 0 | 1 |
| 373 | 5 | 4 | 3 | 3 | 2 | 0 |
| 307 | 1 | 2 | 3 | 2 | 1 | 4 |
| 252 | 1 | 2 | 2 | 0 | 0 | 0 |
| 102 | 0 | 0 | 0 | 0 | 0 | 0 |
| 107 | 0 | 0 | 0 | 0 | 0 | 0 |

| | | | | | | |
|---|---|---|---|---|---|---|
| 32 | 0 | 0 | 0 | 0 | 0 | 0 |
| 51 | 0 | 0 | 0 | 0 | 0 | 0 |
| 38 | 0 | 0 | 0 | 0 | 0 | 0 |
| 20 | 0 | 0 | 0 | 0 | 0 | 0 |
| 23 | 0 | 0 | 0 | 0 | 0 | 0 |
| 13 | 0 | 0 | 0 | 0 | 0 | 0 |
| 1593 | 54 | | | | | |
| 199 | 0 | 0 | 11 | 0 | 0 | 20 |
| 219 | 12 | 4 | 4 | 0 | 0 | 0 |
| 297 | 1 | 0 | 0 | 2 | 0 | 0 |
| 247 | 4 | 0 | 7 | 0 | 0 | 0 |
| 199 | 3 | 0 | 2 | 0 | 0 | 2 |
| 212 | 0 | 0 | 3 | 0 | 0 | 0 |
| 258 | 4 | 2 | 3 | 0 | 0 | 1 |
| 99 | 0 | 0 | 0 | 0 | 0 | 0 |
| 7 | 0 | 0 | 0 | 0 | 0 | 0 |
| 1 | 0 | 0 | 0 | 0 | 0 | 0 |
| 34 | 0 | 0 | 0 | 0 | 0 | 0 |
| 42 | 0 | 0 | 0 | 0 | 0 | 0 |
| 1814 | 104 | | | | | |
| 735 | 6 | 0 | 4 | 0 | 0 | 0 |
| 595 | 8 | 4 | 2 | 1 | 3 | 3 |
| 519 | 2 | 0 | 1 | 0 | 2 | 0 |
| 647 | 5 | 5 | 8 | 4 | 3 | 1 |
| 315 | 3 | 1 | 2 | 0 | 0 | 0 |
| 403 | 0 | 1 | 0 | 0 | 0 | 1 |
| 250 | 2 | 0 | 0 | 3 | 0 | 0 |
| 110 | 0 | 0 | 0 | 0 | 0 | 0 |
| 120 | 0 | 0 | 0 | 0 | 0 | 0 |
| 49 | 0 | 0 | 0 | 0 | 0 | 0 |
| 39 | 0 | 0 | 0 | 0 | 0 | 0 |
| 4 | 0 | 0 | 0 | 0 | 0 | 0 |
| 3786 | 111 | | | | | |
| 382 | 1 | 1 | 0 | 0 | 0 | 0 |
| 256 | 2 | 0 | 2 | 0 | 0 | 0 |
| 300 | 2 | 0 | 0 | 0 | 1 | 0 |
| 151 | 0 | 0 | 0 | 0 | 0 | 0 |
| 95 | 0 | 0 | 0 | 0 | 0 | 0 |
| 0 | 0 | 0 | 0 | 0 | 0 | 0 |
| 17 | 0 | 0 | 0 | 0 | 0 | 0 |
| 1 | 0 | 0 | 0 | 0 | 0 | 0 |

| | | | | | | |
|---|---|---|---|---|---|---|
| 1 | 0 | 0 | 0 | 0 | 0 | 0 |
| 0 | 0 | 0 | 0 | 0 | 0 | 0 |
| 27 | 0 | 0 | 0 | 0 | 0 | 0 |
| 179 | 0 | 1 | 0 | 0 | 0 | 1 |
| 1409 | 25 | | | | | |
| 434 | 0 | 0 | 7 | 0 | 0 | 4 |
| 465 | 4 | 0 | 0 | 2 | 4 | 0 |
| 401 | 1 | 3 | 11 | 0 | 1 | 9 |
| 339 | 0 | 0 | 1 | 1 | 1 | 8 |
| 330 | 0 | 1 | 7 | 0 | 11 | 0 |
| 265 | 6 | 7 | 2 | 0 | 0 | 1 |
| 270 | 3 | 0 | 0 | 2 | 5 | 0 |
| 164 | 1 | 0 | 0 | 0 | 0 | 0 |
| 103 | 0 | 0 | 0 | 1 | 0 | 0 |
| 13 | 0 | 0 | 0 | 0 | 0 | 0 |
| 82 | 0 | 0 | 0 | 0 | 0 | 0 |
| 0 | 0 | 0 | 0 | 0 | 0 | 0 |
| 2866 | 192 | | | | | |
| 420 | 5 | 2 | 4 | 2 | 0 | 0 |
| 337 | 2 | 3 | 3 | 0 | 0 | 3 |
| 248 | 0 | 0 | 0 | 3 | 0 | 0 |
| 132 | 0 | 0 | 0 | 0 | 0 | 0 |
| 125 | 0 | 0 | 1 | 0 | 0 | 0 |
| 57 | 0 | 0 | 0 | 0 | 0 | 0 |
| 57 | 6 | 0 | 0 | 0 | 0 | 0 |
| 9 | 0 | 0 | 0 | 0 | 0 | 0 |
| 3 | 0 | 0 | 0 | 0 | 0 | 0 |
| 0 | 0 | 0 | 0 | 0 | 0 | 0 |
| 6 | 0 | 0 | 0 | 0 | 0 | 0 |
| 0 | 0 | 0 | 0 | 0 | 0 | 0 |
| 1394 | 52 | | | | | |
| 272 | 0 | 0 | 0 | 0 | 0 | 0 |
| 184 | 3 | 5 | 0 | 0 | 0 | 0 |
| 95 | 0 | 0 | 0 | 0 | 0 | 0 |
| 144 | 0 | 0 | 0 | 0 | 0 | 2 |
| 186 | 0 | 0 | 2 | 0 | 0 | 0 |
| 37 | 0 | 0 | 0 | 0 | 0 | 0 |
| 20 | 0 | 0 | 0 | 0 | 0 | 0 |
| 18 | 0 | 0 | 0 | 0 | 0 | 0 |
| 130 | 7 | 0 | 0 | 0 | 0 | 0 |
| 271 | 8 | 4 | 5 | 4 | 0 | 0 |

| | | | | | | |
|---|---|---|---|---|---|---|
| 96 | 0 | 0 | 0 | 0 | 0 | 0 |
| 120 | 0 | 0 | 0 | 0 | 0 | 0 |
| 1573 | 80 | | | | | |
| 804 | 10 | 2 | 6 | 2 | 0 | 8 |
| 688 | 3 | 0 | 4 | 0 | 0 | 0 |
| 731 | 5 | 6 | 2 | 4 | 3 | 3 |
| 619 | 8 | 3 | 3 | 2 | 1 | 3 |
| 552 | 1 | 2 | 2 | 2 | 1 | 1 |
| 469 | 1 | 2 | 2 | 0 | 0 | 0 |
| 426 | 1 | 2 | 3 | 2 | 3 | 1 |
| 186 | 0 | 0 | 0 | 0 | 0 | 0 |
| 27 | 0 | 0 | 0 | 0 | 0 | 0 |
| 54 | 0 | 0 | 0 | 0 | 0 | 0 |
| 329 | 0 | 0 | 1 | 0 | 0 | 0 |
| 167 | 0 | 0 | 1 | 0 | 1 | 0 |
| 5052 | 167 | | | | | |
| 862 | 2 | 1 | 3 | 0 | 2 | 1 |
| 666 | 3 | 5 | 3 | 2 | 1 | 1 |
| 581 | 5 | 3 | 2 | 0 | 1 | 0 |
| 476 | 4 | 0 | 2 | 1 | 1 | 1 |
| 322 | 2 | 1 | 3 | 1 | 0 | 0 |
| 459 | 3 | 0 | 2 | 1 | 1 | 4 |
| 396 | 1 | 0 | 2 | 0 | 2 | 5 |
| 176 | 1 | 0 | 0 | 1 | 0 | 0 |
| 103 | 3 | 0 | 0 | 0 | 0 | 0 |
| 14 | 0 | 0 | 0 | 0 | 0 | 0 |
| 27 | 0 | 0 | 0 | 0 | 0 | 0 |
| 351 | 4 | 0 | 1 | 0 | 0 | 0 |
| 4433 | 152 | | | | | |
| 698 | 0 | 1 | 5 | 4 | 2 | 4 |
| 492 | 3 | 12 | 8 | 2 | 1 | 2 |
| 522 | 1 | 5 | 14 | 2 | 0 | 7 |
| 451 | 4 | 2 | 12 | 6 | 2 | 1 |
| 459 | 3 | 4 | 9 | 1 | 3 | 1 |
| 371 | 5 | 3 | 4 | 1 | 2 | 3 |
| 329 | 1 | 0 | 3 | 1 | 1 | 1 |
| 428 | 8 | 3 | 1 | 2 | 0 | 0 |
| 219 | 0 | 0 | 3 | 0 | 0 | 0 |
| 432 | 3 | 2 | 21 | 0 | 2 | 0 |
| 299 | 0 | 0 | 1 | 4 | 0 | 1 |
| 51 | 0 | 0 | 10 | 0 | 0 | 1 |

| | | | | | | |
|---|---|---|---|---|---|---|
| 4751 | 305 | | | | | |
| 332 | 0 | 0 | 0 | 0 | 4 | 0 |
| 368 | 4 | 0 | 2 | 0 | 0 | 0 |
| 276 | 0 | 1 | 7 | 0 | 0 | 0 |
| 236 | 0 | 1 | 0 | 0 | 0 | 0 |
| 242 | 0 | 0 | 0 | 0 | 0 | 1 |
| 150 | 1 | 0 | 4 | 0 | 0 | 0 |
| 2 | 0 | 0 | 0 | 0 | 0 | 0 |
| 8 | 0 | 0 | 0 | 0 | 0 | 0 |
| 9 | 0 | 0 | 0 | 0 | 0 | 0 |
| 5 | 0 | 0 | 0 | 0 | 0 | 0 |
| 24 | 0 | 0 | 0 | 0 | 0 | 0 |
| 18 | 0 | 0 | 0 | 0 | 0 | 0 |
| 1670 | 57 | | | | | |
| 577 | 3 | 2 | 4 | 1 | 0 | 2 |
| 421 | 1 | 0 | 3 | 2 | 0 | 2 |
| 367 | 3 | 1 | 2 | 0 | 0 | 0 |
| 156 | 0 | 0 | 0 | 0 | 0 | 0 |
| 123 | 0 | 0 | 0 | 0 | 0 | 0 |
| 21 | 0 | 0 | 0 | 0 | 0 | 0 |
| 148 | 4 | 0 | 1 | 0 | 0 | 0 |
| 8 | 0 | 0 | 0 | 0 | 0 | 0 |
| 104 | 0 | 0 | 0 | 0 | 0 | 0 |
| 5 | 0 | 0 | 0 | 0 | 0 | 0 |
| 11 | 0 | 0 | 0 | 0 | 0 | 0 |
| 15 | 0 | 0 | 0 | 0 | 0 | 0 |
| 1956 | 63 | | | | | |
| 268 | 0 | 0 | 1 | 1 | 0 | 1 |
| 143 | 0 | 0 | 0 | 0 | 0 | 0 |
| 108 | 2 | 0 | 0 | 0 | 0 | 0 |
| 16 | 0 | 0 | 0 | 0 | 0 | 0 |
| 7 | 0 | 0 | 0 | 0 | 0 | 0 |
| 0 | 0 | 0 | 0 | 0 | 0 | 0 |
| 9 | 0 | 0 | 0 | 0 | 0 | 0 |
| 67 | 0 | 0 | 0 | 0 | 0 | 0 |
| 47 | 0 | 0 | 0 | 0 | 0 | 0 |
| 21 | 0 | 0 | 0 | 0 | 0 | 0 |
| 53 | 0 | 0 | 0 | 0 | 0 | 0 |
| 54 | 0 | 0 | 0 | 0 | 0 | 0 |
| 793 | 5 | | | | | |
| 901 | 3 | 0 | 2 | 0 | 0 | 1 |

| | | | | | | |
|---|---|---|---|---|---|---|
| 836 | 6 | 4 | 2 | 3 | 3 | 5 |
| 878 | 5 | 6 | 5 | 3 | 6 | 5 |
| 796 | 2 | 0 | 3 | 1 | 1 | 3 |
| 611 | 11 | 6 | 3 | 3 | 7 | 4 |
| 637 | 5 | 6 | 3 | 3 | 4 | 2 |
| 703 | 2 | 1 | 1 | 2 | 3 | 2 |
| 575 | 4 | 4 | 2 | 1 | 4 | 4 |
| 576 | 1 | 3 | 3 | 4 | 0 | 6 |
| 555 | 3 | 1 | 5 | 0 | 0 | 2 |
| 353 | 0 | 1 | 0 | 0 | 0 | 2 |
| 332 | 1 | 0 | 2 | 1 | 1 | 0 |
| 7753 | 309 | | | | | |
| 768 | 4 | 1 | 7 | 4 | 1 | 2 |
| 567 | 6 | 2 | 3 | 4 | 1 | 0 |
| 460 | 2 | 1 | 2 | 5 | 1 | 6 |
| 478 | 2 | 1 | 0 | 0 | 0 | 0 |
| 477 | 2 | 1 | 2 | 7 | 0 | 1 |
| 560 | 0 | 0 | 6 | 0 | 1 | 0 |
| 574 | 5 | 2 | 2 | 2 | 1 | 0 |
| 396 | 0 | 0 | 1 | 0 | 0 | 0 |
| 501 | 1 | 0 | 1 | 0 | 0 | 0 |
| 286 | 2 | 1 | 0 | 5 | 2 | 0 |
| 285 | 0 | 1 | 0 | 0 | 0 | 0 |
| 217 | 0 | 0 | 0 | 0 | 0 | 1 |
| 5569 | 195 | | | | | |
| 461 | 5 | 1 | 0 | 0 | 0 | 0 |
| 407 | 3 | 2 | 3 | 0 | 1 | 0 |
| 456 | 0 | 0 | 1 | 1 | 0 | 0 |
| 459 | 1 | 1 | 1 | 2 | 1 | 4 |
| 409 | 2 | 0 | 0 | 0 | 0 | 4 |
| 452 | 0 | 0 | 0 | 0 | 0 | 0 |
| 539 | 0 | 1 | 0 | 0 | 0 | 0 |
| 441 | 0 | 0 | 1 | 1 | 1 | 0 |
| 418 | 4 | 0 | 0 | 3 | 0 | 0 |
| 379 | 3 | 1 | 1 | 1 | 2 | 2 |
| 362 | 0 | 1 | 2 | 0 | 0 | 1 |
| 195 | 1 | 2 | 0 | 0 | 0 | 0 |
| 4978 | 103 | | | | | |
| 997 | 3 | 3 | 5 | 1 | 3 | 4 |
| 833 | 2 | 3 | 9 | 4 | 4 | 6 |
| 799 | 6 | 2 | 5 | 1 | 5 | 4 |

| | | | | | | |
|---|---|---|---|---|---|---|
| 654 | 2 | 4 | 5 | 2 | 1 | 8 |
| 458 | 3 | 3 | 3 | 1 | 1 | 2 |
| 878 | 4 | 1 | 6 | 2 | 2 | 3 |
| 553 | 2 | 6 | 7 | 2 | 0 | 12 |
| 799 | 6 | 0 | 2 | 0 | 1 | 2 |
| 696 | 3 | 3 | 6 | 3 | 2 | 3 |
| 480 | 1 | 0 | 2 | 1 | 1 | 4 |
| 561 | 2 | 1 | 2 | 3 | 0 | 3 |
| 505 | 3 | 2 | 4 | 1 | 4 | 2 |
| 8213 | 330 | | | | | |
| 889 | 7 | 1 | 10 | 2 | 0 | 2 |
| 665 | 5 | 5 | 7 | 1 | 3 | 8 |
| 766 | 6 | 0 | 2 | 0 | 1 | 3 |
| 611 | 6 | 5 | 2 | 1 | 1 | 2 |
| 652 | 2 | 2 | 2 | 1 | 0 | 1 |
| 620 | 0 | 3 | 2 | 0 | 1 | 2 |
| 634 | 4 | 2 | 5 | 1 | 2 | 8 |
| 548 | 1 | 1 | 2 | 0 | 0 | 0 |
| 535 | 0 | 0 | 2 | 0 | 0 | 0 |
| 547 | 10 | 1 | 2 | 1 | 0 | 0 |
| 543 | 4 | 1 | 3 | 2 | 2 | 1 |
| 355 | 0 | 6 | 1 | 0 | 0 | 4 |
| 7365 | 232 | | | | | |
| 742 | 5 | 1 | 4 | 3 | 0 | 3 |
| 697 | 2 | 5 | 3 | 1 | 3 | 5 |
| 530 | 0 | 4 | 0 | 0 | 0 | 0 |
| 442 | 3 | 3 | 5 | 2 | 2 | 4 |
| 346 | 2 | 0 | 1 | 1 | 2 | 1 |
| 346 | 0 | 2 | 1 | 0 | 4 | 1 |
| 383 | 2 | 2 | 0 | 1 | 1 | 1 |
| 332 | 5 | 3 | 0 | 0 | 2 | 2 |
| 247 | 1 | 0 | 3 | 0 | 0 | 0 |
| 205 | 0 | 0 | 0 | 0 | 0 | 0 |
| 351 | 2 | 3 | 1 | 1 | 5 | 3 |
| 273 | 2 | 1 | 4 | 0 | 0 | 0 |
| 4894 | 183 | | | | | |
| 835 | 6 | 3 | 6 | 2 | 1 | 1 |
| 554 | 0 | 7 | 8 | 0 | 3 | 6 |
| 551 | 2 | 1 | 5 | 0 | 1 | 4 |
| 399 | 4 | 7 | 9 | 2 | 2 | 4 |
| 556 | 1 | 0 | 3 | 0 | 1 | 3 |

| | | | | | | |
|---|---|---|---|---|---|---|
| 415 | 4 | 2 | 2 | 3 | 0 | 1 |
| 437 | 1 | 0 | 1 | 1 | 1 | 1 |
| 338 | 1 | 1 | 2 | 1 | 4 | 11 |
| 323 | 1 | 0 | 0 | 1 | 0 | 0 |
| 287 | 4 | 0 | 2 | 0 | 3 | 1 |
| 127 | 0 | 0 | 0 | 0 | 0 | 0 |
| 78 | 0 | 0 | 0 | 0 | 0 | 0 |
| 4900 | 194 | | | | | |
| 771 | 3 | 5 | 5 | 0 | 1 | 0 |
| 602 | 6 | 3 | 10 | 2 | 0 | 4 |
| 426 | 1 | 7 | 5 | 0 | 2 | 0 |
| 433 | 3 | 2 | 2 | 0 | 1 | 1 |
| 502 | 4 | 2 | 3 | 0 | 0 | 2 |
| 470 | 4 | 2 | 9 | 0 | 0 | 0 |
| 518 | 0 | 0 | 2 | 0 | 0 | 0 |
| 256 | 0 | 0 | 0 | 0 | 0 | 0 |
| 286 | 0 | 0 | 0 | 0 | 0 | 0 |
| 357 | 6 | 2 | 1 | 0 | 0 | 0 |
| 330 | 0 | 0 | 0 | 0 | 0 | 0 |
| 486 | 0 | 0 | 1 | 0 | 0 | 2 |
| 5437 | 137 | | | | | |
| 763 | 3 | 1 | 5 | 1 | 2 | 1 |
| 491 | 2 | 1 | 11 | 0 | 0 | 1 |
| 363 | 5 | 1 | 4 | 3 | 0 | 1 |
| 388 | 0 | 0 | 4 | 0 | 0 | 1 |
| 361 | 0 | 0 | 0 | 0 | 0 | 0 |
| 304 | 0 | 3 | 2 | 0 | 0 | 0 |
| 363 | 0 | 0 | 3 | 0 | 0 | 1 |
| 273 | 0 | 0 | 1 | 0 | 0 | 0 |
| 210 | 0 | 0 | 0 | 0 | 0 | 0 |
| 186 | 3 | 0 | 0 | 0 | 0 | 0 |
| 149 | 0 | 0 | 1 | 0 | 0 | 0 |
| 76 | 0 | 0 | 0 | 0 | 0 | 0 |
| 3927 | 84 | | | | | |
| 307 | 2 | 0 | 0 | 0 | 0 | 2 |
| 373 | 1 | 1 | 3 | 0 | 2 | 2 |
| 383 | 1 | 0 | 2 | 0 | 5 | 7 |
| 169 | 0 | 0 | 3 | 0 | 0 | 1 |
| 202 | 0 | 3 | 0 | 0 | 0 | 0 |
| 121 | 0 | 0 | 0 | 0 | 0 | 0 |
| 127 | 0 | 0 | 1 | 0 | 0 | 0 |

| | | | | | | |
|---|---|---|---|---|---|---|
| 47 | 0 | 0 | 0 | 0 | 0 | 0 |
| 78 | 0 | 0 | 0 | 0 | 0 | 0 |
| 162 | 0 | 0 | 0 | 0 | 2 | 0 |
| 37 | 0 | 0 | 0 | 0 | 0 | 0 |
| 0 | 0 | 0 | 0 | 0 | 0 | 0 |
| 2006 | 81 | | | | | |
| 340 | 0 | 0 | 0 | 0 | 3 | 0 |
| 443 | 6 | 1 | 1 | 1 | 0 | 0 |
| 182 | 1 | 0 | 1 | 0 | 0 | 0 |
| 129 | 0 | 0 | 0 | 0 | 1 | 0 |
| 112 | 0 | 0 | 0 | 0 | 0 | 0 |
| 110 | 0 | 0 | 0 | 0 | 0 | 0 |
| 77 | 0 | 0 | 0 | 0 | 0 | 0 |
| 30 | 0 | 0 | 0 | 0 | 0 | 0 |
| 29 | 0 | 0 | 0 | 0 | 0 | 0 |
| 91 | 0 | 0 | 0 | 0 | 0 | 0 |
| 83 | 0 | 0 | 0 | 0 | 0 | 0 |
| 61 | 0 | 0 | 0 | 0 | 0 | 0 |
| 1687 | 33 | | | | | |
| 212 | 0 | 0 | 0 | 0 | 1 | 0 |
| 274 | 2 | 1 | 0 | 0 | 0 | 0 |
| 157 | 0 | 1 | 0 | 0 | 0 | 0 |
| 102 | 0 | 0 | 0 | 0 | 0 | 0 |
| 30 | 0 | 0 | 0 | 0 | 0 | 0 |
| 57 | 0 | 0 | 0 | 0 | 0 | 0 |
| 108 | 0 | 0 | 1 | 0 | 0 | 0 |
| 52 | 0 | 0 | 0 | 0 | 0 | 0 |
| 13 | 0 | 0 | 0 | 0 | 0 | 0 |
| 11 | 0 | 0 | 0 | 0 | 0 | 0 |
| 94 | 0 | 0 | 0 | 0 | 0 | 0 |
| 70 | 0 | 0 | 0 | 0 | 0 | 0 |
| 1180 | 20 | | | | | |
| 212 | 0 | 0 | 25 | 0 | 0 | 0 |
| 213 | 0 | 0 | 9 | 0 | 0 | 3 |
| 405 | 6 | 5 | 3 | 4 | 0 | 2 |
| 358 | 5 | 0 | 7 | 5 | 2 | 2 |
| 234 | 0 | 0 | 2 | 0 | 0 | 2 |
| 206 | 8 | 1 | 0 | 1 | 0 | 0 |
| 166 | 0 | 0 | 0 | 0 | 0 | 0 |
| 9 | 0 | 0 | 0 | 0 | 0 | 0 |
| 1 | 0 | 0 | 0 | 0 | 0 | 0 |

| | | | | | | |
|---|---|---|---|---|---|---|
| 51 | 0 | 0 | 0 | 0 | 0 | 0 |
| 6 | 0 | 0 | 0 | 0 | 0 | 0 |
| 2 | 0 | 0 | 0 | 0 | 0 | 0 |
| 1863 | 167 | | | | | |
| 235 | 0 | 0 | 2 | 0 | 0 | 0 |
| 185 | 1 | 0 | 0 | 0 | 0 | 0 |
| 246 | 0 | 0 | 0 | 3 | 0 | 0 |
| 232 | 0 | 3 | 4 | 0 | 0 | 0 |
| 192 | 2 | 0 | 0 | 0 | 0 | 0 |
| 145 | 0 | 0 | 0 | 0 | 0 | 0 |
| 147 | 0 | 0 | 0 | 0 | 0 | 0 |
| 167 | 0 | 0 | 0 | 0 | 0 | 1 |
| 91 | 0 | 0 | 0 | 0 | 0 | 0 |
| 25 | 0 | 0 | 0 | 0 | 0 | 0 |
| 17 | 0 | 0 | 0 | 0 | 0 | 0 |
| 93 | 0 | 0 | 0 | 0 | 0 | 0 |
| 1775 | 42 | | | | | |
| 111 | 0 | 0 | 0 | 0 | 0 | 0 |
| 111 | 0 | 0 | 0 | 0 | 0 | 0 |
| 148 | 0 | 0 | 4 | 0 | 0 | 0 |
| 22 | 0 | 0 | 0 | 0 | 0 | 0 |
| 85 | 0 | 0 | 0 | 0 | 0 | 0 |
| 38 | 0 | 0 | 0 | 0 | 0 | 0 |
| 33 | 0 | 0 | 0 | 0 | 0 | 0 |
| 20 | 0 | 0 | 0 | 0 | 0 | 0 |
| 20 | 0 | 0 | 0 | 0 | 0 | 0 |
| 18 | 0 | 0 | 0 | 0 | 0 | 0 |
| 19 | 0 | 0 | 0 | 0 | 0 | 0 |
| 29 | 0 | 0 | 0 | 0 | 0 | 0 |
| 654 | 4 | | | | | |
| 641 | 13 | 8 | 8 | 1 | 0 | 1 |
| 518 | 9 | 9 | 4 | 0 | 1 | 0 |
| 384 | 12 | 7 | 4 | 5 | 1 | 0 |
| 418 | 3 | 3 | 5 | 2 | 3 | 1 |
| 638 | 6 | 1 | 3 | 0 | 0 | 0 |
| 505 | 3 | 3 | 4 | 4 | 0 | 0 |
| 364 | 4 | 11 | 0 | 2 | 2 | 0 |
| 223 | 2 | 4 | 1 | 0 | 0 | 0 |
| 169 | 1 | 0 | 2 | 0 | 1 | 0 |
| 63 | 3 | 0 | 0 | 0 | 0 | 0 |
| 27 | 3 | 0 | 0 | 0 | 0 | 0 |

| | | | | | | |
|---|---|---|---|---|---|---|
| 26 | 0 | 0 | 0 | 0 | 0 | 0 |
| 3976 | 203 | | | | | |
| 425 | 1 | 0 | 3 | 0 | 0 | 0 |
| 353 | 0 | 0 | 1 | 0 | 0 | 2 |
| 438 | 2 | 2 | 0 | 0 | 2 | 0 |
| 356 | 0 | 2 | 1 | 0 | 0 | 0 |
| 390 | 8 | 1 | 0 | 3 | 0 | 0 |
| 421 | 0 | 0 | 0 | 0 | 0 | 0 |
| 189 | 0 | 0 | 0 | 0 | 0 | 0 |
| 266 | 0 | 0 | 0 | 0 | 0 | 0 |
| 170 | 0 | 0 | 0 | 0 | 0 | 0 |
| 189 | 2 | 0 | 2 | 0 | 0 | 0 |
| 68 | 0 | 0 | 1 | 0 | 0 | 0 |
| 100 | 0 | 0 | 0 | 0 | 0 | 0 |
| 3365 | 42 | | | | | |
| 463 | 2 | 0 | 0 | 0 | 0 | 0 |
| 449 | 0 | 0 | 1 | 0 | 2 | 0 |
| 283 | 1 | 0 | 1 | 0 | 0 | 0 |
| 210 | 0 | 0 | 0 | 0 | 0 | 0 |
| 107 | 0 | 0 | 0 | 0 | 0 | 0 |
| 102 | 0 | 0 | 0 | 0 | 0 | 0 |
| 78 | 0 | 0 | 0 | 0 | 0 | 0 |
| 43 | 0 | 0 | 0 | 0 | 0 | 0 |
| 5 | 0 | 0 | 0 | 0 | 0 | 0 |
| 0 | 0 | 0 | 0 | 0 | 0 | 0 |
| 4 | 0 | 0 | 0 | 0 | 0 | 0 |
| 54 | 1 | 0 | 0 | 0 | 0 | 0 |
| 1798 | 12 | | | | | |
| 568 | 7 | 2 | 9 | 2 | 0 | 1 |
| 530 | 3 | 1 | 8 | 0 | 2 | 2 |
| 493 | 6 | 1 | 3 | 2 | 2 | 0 |
| 400 | 5 | 8 | 10 | 1 | 2 | 2 |
| 336 | 1 | 5 | 7 | 6 | 0 | 1 |
| 352 | 13 | 7 | 2 | 0 | 2 | 3 |
| 311 | 2 | 1 | 6 | 0 | 2 | 0 |
| 129 | 1 | 0 | 0 | 0 | 0 | 0 |
| 70 | 0 | 0 | 0 | 0 | 0 | 0 |
| 13 | 0 | 0 | 0 | 0 | 0 | 0 |
| 1 | 0 | 0 | 0 | 0 | 0 | 0 |
| 0 | 0 | 0 | 0 | 0 | 0 | 0 |
| 3203 | 213 | | | | | |

| | | | | | | |
|---|---|---|---|---|---|---|
| 609 | 4 | 2 | 6 | 0 | 0 | 1 |
| 571 | 3 | 1 | 4 | 2 | 1 | 2 |
| 481 | 4 | 3 | 12 | 0 | 1 | 0 |
| 398 | 4 | 1 | 2 | 3 | 1 | 5 |
| 346 | 3 | 2 | 10 | 1 | 0 | 1 |
| 312 | 3 | 18 | 20 | 1 | 0 | 2 |
| 429 | 5 | 5 | 3 | 7 | 5 | 0 |
| 362 | 2 | 2 | 4 | 2 | 0 | 1 |
| 421 | 5 | 2 | 15 | 0 | 0 | 4 |
| 256 | 6 | 0 | 6 | 4 | 0 | 1 |
| 172 | 0 | 0 | 14 | 0 | 0 | 0 |
| 113 | 0 | 0 | 4 | 0 | 0 | 0 |
| 4470 | 297 | | | | | |
| 546 | 7 | 5 | 8 | 3 | 2 | 2 |
| 384 | 4 | 0 | 10 | 2 | 2 | 3 |
| 374 | 1 | 1 | 4 | 1 | 1 | 5 |
| 241 | 1 | 4 | 7 | 0 | 0 | 4 |
| 285 | 5 | 3 | 4 | 1 | 0 | 2 |
| 316 | 1 | 3 | 4 | 0 | 0 | 3 |
| 407 | 1 | 4 | 4 | 4 | 6 | 7 |
| 226 | 0 | 0 | 10 | 0 | 0 | 0 |
| 168 | 0 | 0 | 2 | 0 | 0 | 1 |
| 183 | 4 | 0 | 7 | 0 | 0 | 1 |
| 166 | 0 | 0 | 0 | 0 | 0 | 0 |
| 158 | 0 | 1 | 1 | 0 | 0 | 0 |
| 3454 | 202 | | | | | |
| 768 | 20 | 7 | 6 | 3 | 5 | 6 |
| 586 | 13 | 3 | 5 | 2 | 4 | 1 |
| 595 | 6 | 1 | 3 | 4 | 2 | 6 |
| 504 | 5 | 1 | 2 | 3 | 2 | 1 |
| 478 | 12 | 3 | 4 | 8 | 1 | 3 |
| 405 | 20 | 0 | 1 | 2 | 1 | 1 |
| 418 | 4 | 5 | 2 | 4 | 0 | 0 |
| 454 | 24 | 4 | 1 | 3 | 0 | 1 |
| 427 | 6 | 1 | 1 | 5 | 0 | 1 |
| 441 | 7 | 4 | 2 | 5 | 2 | 1 |
| 398 | 18 | 3 | 2 | 1 | 2 | 2 |
| 317 | 11 | 2 | 2 | 3 | 1 | 1 |
| 5791 | 443 | | | | | |
| 647 | 8 | 7 | 6 | 3 | 1 | 2 |
| 496 | 12 | 5 | 1 | 1 | 2 | 3 |

| | | | | | | |
|---|---|---|---|---|---|---|
| 506 | 4 | 1 | 3 | 0 | 0 | 0 |
| 531 | 5 | 5 | 2 | 1 | 2 | 3 |
| 340 | 2 | 4 | 6 | 0 | 0 | 6 |
| 448 | 7 | 7 | 11 | 2 | 2 | 2 |
| 504 | 7 | 1 | 4 | 3 | 1 | 1 |
| 425 | 4 | 5 | 6 | 1 | 1 | 5 |
| 347 | 5 | 2 | 8 | 2 | 3 | 2 |
| 341 | 3 | 8 | 5 | 0 | 0 | 3 |
| 450 | 3 | 5 | 2 | 0 | 0 | 0 |
| 463 | 2 | 1 | 0 | 0 | 4 | 0 |
| 5498 | 287 | | | | | |
| 449 | 5 | 2 | 5 | 0 | 0 | 1 |
| 305 | 0 | 3 | 1 | 1 | 1 | 0 |
| 352 | 4 | 0 | 1 | 1 | 2 | 0 |
| 282 | 0 | 0 | 0 | 0 | 0 | 1 |
| 248 | 1 | 1 | 2 | 1 | 0 | 1 |
| 285 | 4 | 0 | 0 | 1 | 0 | 1 |
| 228 | 1 | 1 | 7 | 3 | 0 | 0 |
| 151 | 0 | 0 | 1 | 0 | 0 | 4 |
| 283 | 7 | 0 | 1 | 4 | 0 | 0 |
| 288 | 14 | 1 | 0 | 6 | 0 | 2 |
| 188 | 8 | 6 | 2 | 4 | 1 | 0 |
| 188 | 1 | 0 | 0 | 3 | 0 | 0 |
| 3247 | 157 | | | | | |
| 750 | 15 | 8 | 5 | 3 | 0 | 4 |
| 536 | 4 | 1 | 7 | 0 | 2 | 0 |
| 555 | 7 | 2 | 1 | 3 | 2 | 2 |
| 304 | 5 | 4 | 5 | 3 | 1 | 0 |
| 425 | 6 | 3 | 5 | 2 | 6 | 2 |
| 694 | 4 | 5 | 4 | 3 | 1 | 6 |
| 429 | 2 | 1 | 3 | 2 | 3 | 4 |
| 401 | 2 | 2 | 16 | 1 | 2 | 3 |
| 465 | 4 | 0 | 2 | 4 | 0 | 1 |
| 533 | 1 | 2 | 2 | 3 | 2 | 2 |
| 467 | 4 | 3 | 6 | 1 | 3 | 8 |
| 384 | 1 | 9 | 5 | 1 | 3 | 2 |
| 5943 | 361 | | | | | |
| 660 | 6 | 0 | 5 | 1 | 0 | 1 |
| 574 | 4 | 7 | 0 | 1 | 0 | 0 |
| 719 | 8 | 3 | 2 | 0 | 3 | 2 |
| 672 | 2 | 1 | 4 | 1 | 0 | 1 |

| | | | | | | |
|---|---|---|---|---|---|---|
| 684 | 2 | 1 | 2 | 0 | 0 | 2 |
| 488 | 5 | 3 | 5 | 3 | 2 | 7 |
| 494 | 2 | 0 | 0 | 6 | 1 | 0 |
| 745 | 4 | 2 | 1 | 4 | 2 | 1 |
| 483 | 6 | 3 | 5 | 3 | 1 | 3 |
| 540 | 0 | 1 | 3 | 1 | 1 | 2 |
| 466 | 1 | 6 | 1 | 4 | 2 | 1 |
| 467 | 1 | 0 | 2 | 2 | 5 | 2 |
| 6992 | 212 | | | | | |
| 552 | 3 | 2 | 3 | 1 | 0 | 2 |
| 436 | 0 | 0 | 0 | 0 | 1 | 1 |
| 385 | 2 | 1 | 6 | 1 | 1 | 1 |
| 417 | 0 | 0 | 0 | 0 | 0 | 0 |
| 332 | 0 | 0 | 0 | 0 | 0 | 0 |
| 457 | 0 | 0 | 0 | 0 | 0 | 0 |
| 420 | 2 | 2 | 2 | 2 | 2 | 3 |
| 542 | 2 | 1 | 1 | 0 | 2 | 1 |
| 467 | 1 | 0 | 0 | 0 | 1 | 1 |
| 671 | 0 | 0 | 0 | 0 | 0 | 0 |
| 465 | 2 | 1 | 2 | 2 | 2 | 1 |
| 506 | 0 | 1 | 3 | 0 | 0 | 2 |
| 5650 | 112 | | | | | |
| 863 | 10 | 2 | 8 | 5 | 2 | 1 |
| 590 | 17 | 5 | 7 | 12 | 0 | 3 |
| 477 | 4 | 2 | 7 | 2 | 1 | 2 |
| 430 | 0 | 1 | 1 | 1 | 0 | 0 |
| 530 | 8 | 3 | 8 | 1 | 0 | 5 |
| 361 | 7 | 0 | 3 | 1 | 0 | 0 |
| 250 | 1 | 0 | 1 | 0 | 1 | 3 |
| 316 | 3 | 0 | 0 | 0 | 0 | 0 |
| 110 | 8 | 0 | 0 | 0 | 0 | 0 |
| 24 | 0 | 0 | 0 | 0 | 0 | 0 |
| 28 | 0 | 0 | 0 | 0 | 0 | 0 |
| 144 | 3 | 0 | 1 | 0 | 0 | 0 |
| 4123 | 202 | | | | | |
| 620 | 4 | 4 | 6 | 0 | 0 | 0 |
| 484 | 1 | 0 | 1 | 2 | 0 | 0 |
| 399 | 1 | 2 | 1 | 0 | 0 | 0 |
| 374 | 1 | 4 | 1 | 0 | 0 | 0 |
| 269 | 5 | 0 | 0 | 0 | 0 | 0 |
| 157 | 1 | 2 | 2 | 0 | 0 | 0 |

| | | | | | | |
|---|---|---|---|---|---|---|
| 206 | 1 | 0 | 0 | 0 | 0 | 0 |
| 164 | 1 | 0 | 0 | 0 | 0 | 0 |
| 45 | 1 | 0 | 0 | 0 | 0 | 0 |
| 295 | 0 | 0 | 0 | 0 | 0 | 0 |
| 311 | 0 | 0 | 1 | 0 | 0 | 0 |
| 287 | 1 | 0 | 2 | 0 | 0 | 0 |
| 3611 | 79 | | | | | |
| 527 | 4 | 2 | 4 | 8 | 0 | 0 |
| 207 | 0 | 0 | 3 | 0 | 0 | 0 |
| 147 | 0 | 0 | 0 | 0 | 0 | 0 |
| 125 | 0 | 0 | 0 | 0 | 0 | 0 |
| 76 | 0 | 0 | 0 | 0 | 0 | 0 |
| 12 | 0 | 0 | 0 | 0 | 0 | 0 |
| 10 | 0 | 0 | 0 | 0 | 0 | 0 |
| 161 | 0 | 0 | 0 | 0 | 0 | 0 |
| 41 | 0 | 0 | 0 | 0 | 0 | 0 |
| 96 | 0 | 1 | 0 | 0 | 0 | 0 |
| 459 | 0 | 1 | 4 | 2 | 1 | 0 |
| 263 | 2 | 2 | 15 | 0 | 0 | 0 |
| 2124 | 65 | | | | | |
| 617 | 20 | 4 | 15 | 4 | 0 | 6 |
| 642 | 10 | 0 | 1 | 0 | 0 | 2 |
| 583 | 5 | 6 | 6 | 2 | 7 | 5 |
| 551 | 3 | 2 | 5 | 0 | 4 | 3 |
| 661 | 11 | 4 | 7 | 3 | 4 | 1 |
| 420 | 2 | 0 | 6 | 4 | 0 | 1 |
| 390 | 7 | 1 | 2 | 3 | 0 | 1 |
| 297 | 3 | 0 | 1 | 0 | 0 | 0 |
| 371 | 5 | 1 | 3 | 3 | 1 | 0 |
| 302 | 0 | 0 | 4 | 0 | 0 | 0 |
| 291 | 1 | 1 | 4 | 1 | 0 | 0 |
| 187 | 3 | 3 | 18 | 0 | 0 | 0 |
| 5312 | 317 | | | | | |
| 767 | 1 | 2 | 6 | 0 | 2 | 2 |
| 596 | 3 | 1 | 4 | 0 | 2 | 1 |
| 494 | 1 | 0 | 4 | 1 | 0 | 0 |
| 394 | 2 | 0 | 0 | 1 | 0 | 0 |
| 276 | 0 | 0 | 1 | 0 | 0 | 0 |
| 283 | 3 | 2 | 2 | 3 | 1 | 2 |
| 154 | 0 | 0 | 1 | 0 | 0 | 0 |
| 318 | 0 | 0 | 0 | 0 | 0 | 0 |

| | | | | | | |
|---|---|---|---|---|---|---|
| 39 | 0 | 0 | 0 | 0 | 0 | 0 |
| 332 | 0 | 0 | 0 | 0 | 0 | 0 |
| 566 | 0 | 0 | 0 | 0 | 0 | 1 |
| 414 | 1 | 0 | 3 | 0 | 0 | 0 |
| 4633 | 102 | | | | | |
| 370 | 0 | 0 | 3 | 0 | 0 | 0 |
| 325 | 1 | 1 | 0 | 0 | 1 | 1 |
| 313 | 0 | 0 | 1 | 0 | 1 | 0 |
| 241 | 0 | 0 | 0 | 0 | 0 | 1 |
| 108 | 0 | 0 | 0 | 0 | 0 | 0 |
| 123 | 0 | 0 | 0 | 0 | 0 | 0 |
| 240 | 0 | 0 | 0 | 1 | 0 | 0 |
| 278 | 0 | 0 | 2 | 0 | 0 | 0 |
| 412 | 8 | 1 | 2 | 0 | 8 | 1 |
| 180 | 0 | 0 | 0 | 0 | 0 | 0 |
| 88 | 0 | 0 | 0 | 0 | 0 | 0 |
| 1 | 0 | 0 | 0 | 0 | 0 | 0 |
| 2679 | 53 | | | | | |
| 202 | 0 | 8 | 17 | 2 | 0 | 0 |
| 151 | 0 | 0 | 8 | 0 | 0 | 0 |
| 228 | 20 | 0 | 3 | 4 | 0 | 0 |
| 313 | 2 | 0 | 3 | 1 | 0 | 2 |
| 252 | 3 | 0 | 6 | 0 | 0 | 1 |
| 137 | 0 | 0 | 2 | 1 | 0 | 0 |
| 171 | 0 | 0 | 1 | 1 | 0 | 0 |
| 67 | 0 | 0 | 0 | 0 | 0 | 0 |
| 159 | 0 | 0 | 0 | 0 | 0 | 0 |
| 174 | 0 | 0 | 1 | 0 | 0 | 0 |
| 99 | 2 | 0 | 4 | 0 | 0 | 0 |
| 64 | 1 | 0 | 0 | 0 | 0 | 0 |
| 2017 | 119 | | | | | |
| 405 | 6 | 7 | 3 | 1 | 0 | 0 |
| 337 | 0 | 0 | 1 | 0 | 0 | 4 |
| 206 | 1 | 0 | 0 | 0 | 0 | 0 |
| 204 | 0 | 0 | 1 | 0 | 0 | 0 |
| 272 | 4 | 0 | 1 | 1 | 0 | 0 |
| 141 | 0 | 0 | 1 | 0 | 0 | 0 |
| 74 | 0 | 0 | 0 | 0 | 0 | 0 |
| 65 | 0 | 0 | 0 | 0 | 0 | 0 |
| 30 | 0 | 0 | 0 | 0 | 0 | 0 |
| 48 | 0 | 0 | 0 | 0 | 0 | 0 |

| | | | | | | |
|---|---|---|---|---|---|---|
| 4 | 0 | 0 | 0 | 0 | 0 | 0 |
| 27 | 0 | 0 | 0 | 0 | 0 | 0 |
| 1813 | 57 | | | | | |
| 468 | 0 | 3 | 2 | 0 | 0 | 1 |
| 263 | 5 | 0 | 0 | 0 | 0 | 0 |
| 312 | 4 | 1 | 2 | 0 | 2 | 1 |
| 219 | 0 | 0 | 0 | 0 | 0 | 0 |
| 183 | 1 | 0 | 0 | 0 | 0 | 0 |
| 102 | 1 | 0 | 0 | 0 | 0 | 0 |
| 63 | 0 | 0 | 0 | 0 | 0 | 0 |
| 72 | 0 | 0 | 0 | 0 | 0 | 0 |
| 223 | 0 | 0 | 0 | 0 | 0 | 0 |
| 39 | 0 | 0 | 0 | 0 | 0 | 0 |
| 73 | 0 | 0 | 0 | 0 | 0 | 0 |
| 49 | 0 | 0 | 0 | 0 | 0 | 0 |
| 2066 | 32 | | | | | |
| 669 | 4 | 1 | 4 | 1 | 0 | 2 |
| 554 | 3 | 3 | 8 | 1 | 0 | 0 |
| 393 | 2 | 1 | 7 | 0 | 0 | 2 |
| 639 | 3 | 4 | 9 | 4 | 4 | 4 |
| 500 | 3 | 2 | 7 | 2 | 0 | 2 |
| 493 | 3 | 0 | 2 | 5 | 6 | 2 |
| 372 | 2 | 1 | 4 | 2 | 0 | 1 |
| 398 | 2 | 0 | 3 | 4 | 0 | 0 |
| 412 | 2 | 0 | 0 | 2 | 5 | 1 |
| 301 | 4 | 0 | 0 | 4 | 0 | 0 |
| 259 | 4 | 1 | 2 | 1 | 0 | 0 |
| 275 | 1 | 1 | 0 | 6 | 3 | 1 |
| 5265 | 261 | | | | | |
| 553 | 4 | 4 | 3 | 1 | 2 | 0 |
| 404 | 1 | 0 | 3 | 4 | 1 | 3 |
| 456 | 9 | 6 | 2 | 0 | 3 | 0 |
| 413 | 0 | 2 | 3 | 2 | 7 | 0 |
| 377 | 2 | 0 | 6 | 5 | 0 | 2 |
| 445 | 1 | 2 | 3 | 0 | 1 | 2 |
| 399 | 2 | 0 | 0 | 0 | 0 | 3 |
| 330 | 2 | 8 | 3 | 4 | 7 | 1 |
| 318 | 1 | 1 | 0 | 0 | 1 | 1 |
| 205 | 0 | 0 | 0 | 1 | 0 | 0 |
| 124 | 0 | 0 | 0 | 0 | 0 | 1 |
| 27 | 0 | 0 | 0 | 0 | 0 | 0 |

| | | | | | | |
|---|---|---|---|---|---|---|
| 4051 | 189 | | | | | |
| 568 | 2 | 2 | 1 | 0 | 4 | 1 |
| 470 | 2 | 0 | 2 | 2 | 0 | 0 |
| 485 | 16 | 6 | 0 | 1 | 6 | 1 |
| 377 | 9 | 2 | 5 | 1 | 1 | 0 |
| 449 | 7 | 4 | 3 | 1 | 1 | 1 |
| 300 | 8 | 4 | 2 | 0 | 1 | 0 |
| 387 | 0 | 1 | 2 | 0 | 0 | 1 |
| 382 | 3 | 2 | 12 | 0 | 2 | 0 |
| 422 | 2 | 3 | 2 | 5 | 0 | 9 |
| 234 | 0 | 1 | 4 | 1 | 1 | 3 |
| 265 | 1 | 3 | 1 | 0 | 1 | 0 |
| 190 | 0 | 0 | 1 | 0 | 0 | 0 |
| 4529 | 221 | | | | | |
| 527 | 4 | 0 | 8 | 2 | 0 | 2 |
| 535 | 6 | 5 | 4 | 2 | 0 | 0 |
| 568 | 19 | 3 | 4 | 1 | 4 | 0 |
| 466 | 4 | 7 | 15 | 0 | 0 | 0 |
| 569 | 3 | 2 | 2 | 2 | 1 | 3 |
| 441 | 1 | 2 | 2 | 0 | 2 | 9 |
| 332 | 0 | 0 | 2 | 0 | 0 | 0 |
| 421 | 7 | 6 | 10 | 1 | 3 | 3 |
| 485 | 3 | 2 | 3 | 0 | 0 | 1 |
| 303 | 1 | 2 | 5 | 1 | 0 | 0 |
| 296 | 0 | 2 | 4 | 1 | 0 | 0 |
| 293 | 0 | 0 | 0 | 0 | 0 | 0 |
| 5236 | 282 | | | | | |
| 452 | 2 | 1 | 2 | 0 | 0 | 0 |
| 442 | 5 | 1 | 6 | 3 | 0 | 0 |
| 430 | 7 | 2 | 10 | 2 | 0 | 3 |
| 417 | 3 | 2 | 0 | 1 | 0 | 0 |
| 325 | 3 | 1 | 1 | 0 | 0 | 0 |
| 340 | 1 | 2 | 5 | 1 | 0 | 3 |
| 321 | 0 | 0 | 5 | 0 | 0 | 1 |
| 220 | 1 | 1 | 0 | 0 | 0 | 0 |
| 266 | 0 | 0 | 3 | 0 | 0 | 0 |
| 213 | 3 | 1 | 3 | 0 | 0 | 0 |
| 149 | 0 | 0 | 0 | 0 | 0 | 0 |
| 162 | 0 | 0 | 0 | 0 | 0 | 0 |
| 3737 | 131 | | | | | |
| 598 | 4 | 0 | 3 | 5 | 0 | 2 |

| | | | | | | |
|---|---|---|---|---|---|---|
| 530 | 5 | 1 | 8 | 2 | 1 | 2 |
| 322 | 10 | 1 | 8 | 2 | 0 | 2 |
| 422 | 3 | 1 | 3 | 0 | 3 | 1 |
| 411 | 1 | 2 | 5 | 4 | 1 | 0 |
| 299 | 1 | 1 | 3 | 0 | 0 | 2 |
| 271 | 1 | 0 | 0 | 0 | 0 | 0 |
| 346 | 2 | 0 | 4 | 1 | 0 | 1 |
| 265 | 0 | 0 | 0 | 0 | 0 | 1 |
| 281 | 0 | 0 | 0 | 0 | 0 | 0 |
| 213 | 1 | 3 | 2 | 0 | 0 | 0 |
| 134 | 0 | 0 | 0 | 0 | 0 | 0 |
| 4092 | 142 | | | | | |
| 674 | 9 | 2 | 6 | 0 | 0 | 2 |
| 638 | 5 | 8 | 10 | 3 | 4 | 2 |
| 538 | 6 | 3 | 7 | 1 | 3 | 1 |
| 481 | 5 | 3 | 7 | 1 | 1 | 5 |
| 537 | 5 | 4 | 8 | 3 | 1 | 2 |
| 376 | 2 | 0 | 2 | 1 | 1 | 3 |
| 421 | 8 | 3 | 19 | 3 | 2 | 5 |
| 349 | 6 | 1 | 2 | 1 | 0 | 0 |
| 277 | 5 | 0 | 0 | 1 | 1 | 1 |
| 172 | 3 | 3 | 3 | 0 | 0 | 0 |
| 229 | 0 | 0 | 1 | 0 | 0 | 0 |
| 251 | 3 | 0 | 0 | 2 | 1 | 0 |
| 4943 | 279 | | | | | |
| 608 | 2 | 1 | 5 | 3 | 1 | 2 |
| 530 | 4 | 1 | 4 | 2 | 1 | 2 |
| 617 | 3 | 2 | 14 | 3 | 1 | 1 |
| 514 | 0 | 0 | 4 | 3 | 1 | 2 |
| 460 | 6 | 2 | 2 | 1 | 2 | 2 |
| 553 | 7 | 2 | 2 | 0 | 0 | 0 |
| 433 | 6 | 3 | 2 | 2 | 0 | 0 |
| 369 | 6 | 0 | 6 | 0 | 1 | 4 |
| 344 | 10 | 1 | 0 | 0 | 0 | 2 |
| 426 | 0 | 0 | 1 | 1 | 0 | 0 |
| 295 | 1 | 0 | 3 | 1 | 2 | 0 |
| 314 | 4 | 5 | 6 | 1 | 0 | 2 |
| 5463 | 261 | | | | | |
| 648 | 3 | 2 | 3 | 1 | 1 | 1 |
| 502 | 5 | 1 | 4 | 2 | 2 | 1 |
| 452 | 6 | 3 | 3 | 1 | 0 | 3 |

| | | | | | | |
|---|---|---|---|---|---|---|
| 486 | 1 | 1 | 2 | 0 | 0 | 1 |
| 346 | 4 | 3 | 3 | 0 | 1 | 0 |
| 403 | 1 | 0 | 1 | 1 | 2 | 1 |
| 420 | 0 | 1 | 1 | 0 | 0 | 0 |
| 384 | 1 | 0 | 0 | 0 | 1 | 1 |
| 333 | 5 | 3 | 5 | 3 | 0 | 1 |
| 313 | 0 | 0 | 2 | 0 | 0 | 2 |
| 293 | 1 | 5 | 5 | 0 | 1 | 4 |
| 367 | 1 | 0 | 4 | 0 | 0 | 0 |
| 4947 | 188 | | | | | |
| 213 | 0 | 1 | 27 | 0 | 0 | 0 |
| 110 | 0 | 0 | 14 | 0 | 0 | 0 |
| 42 | 0 | 0 | 10 | 0 | 0 | 0 |
| 80 | 0 | 0 | 12 | 0 | 0 | 0 |
| 69 | 0 | 0 | 8 | 0 | 0 | 0 |
| 62 | 0 | 0 | 9 | 0 | 0 | 0 |
| 49 | 0 | 0 | 6 | 0 | 0 | 0 |
| 92 | 0 | 0 | 35 | 0 | 0 | 0 |
| 122 | 0 | 0 | 29 | 0 | 0 | 0 |
| 100 | 0 | 0 | 0 | 0 | 0 | 0 |
| 147 | 8 | 0 | 1 | 0 | 0 | 0 |
| 317 | 7 | 10 | 2 | 0 | 0 | 2 |
| 1403 | 195 | | | | | |
| 459 | 5 | 1 | 7 | 1 | 0 | 1 |
| 452 | 3 | 1 | 1 | 1 | 1 | 0 |
| 206 | 2 | 0 | 2 | 0 | 0 | 5 |
| 260 | 0 | 0 | 0 | 0 | 3 | 0 |
| 388 | 0 | 0 | 1 | 0 | 0 | 0 |
| 192 | 0 | 0 | 0 | 1 | 0 | 0 |
| 76 | 0 | 0 | 0 | 0 | 0 | 0 |
| 12 | 0 | 0 | 0 | 0 | 0 | 0 |
| 0 | 0 | 0 | 0 | 0 | 0 | 0 |
| 8 | 0 | 0 | 0 | 0 | 0 | 0 |
| 0 | 0 | 0 | 0 | 0 | 0 | 0 |
| 3 | 0 | 0 | 0 | 0 | 0 | 0 |
| 2056 | 61 | | | | | |
| 357 | 3 | 0 | 2 | 4 | 0 | 3 |
| 255 | 4 | 0 | 2 | 0 | 1 | 0 |
| 251 | 4 | 0 | 0 | 0 | 0 | 0 |
| 87 | 0 | 0 | 0 | 0 | 0 | 0 |
| 179 | 4 | 0 | 2 | 0 | 0 | 0 |

| | | | | | | |
|---|---|---|---|---|---|---|
| 42 | 0 | 0 | 0 | 0 | 0 | 0 |
| 32 | 0 | 0 | 1 | 0 | 0 | 0 |
| 36 | 0 | 0 | 0 | 0 | 0 | 0 |
| 21 | 0 | 0 | 0 | 0 | 0 | 0 |
| 0 | 0 | 0 | 0 | 0 | 0 | 0 |
| 0 | 0 | 0 | 0 | 0 | 0 | 0 |
| 0 | 0 | 0 | 0 | 0 | 0 | 0 |
| 1260 | 36 | | | | | |
| 466 | 12 | 3 | 6 | 1 | 0 | 2 |
| 201 | 5 | 0 | 0 | 0 | 0 | 0 |
| 48 | 1 | 0 | 0 | 0 | 0 | 0 |
| 42 | 0 | 0 | 0 | 0 | 0 | 0 |
| 20 | 0 | 0 | 0 | 0 | 0 | 0 |
| 9 | 0 | 0 | 0 | 0 | 0 | 0 |
| 55 | 0 | 0 | 0 | 0 | 0 | 0 |
| 33 | 0 | 0 | 0 | 0 | 0 | 0 |
| 0 | 0 | 0 | 0 | 0 | 0 | 0 |
| 0 | 0 | 0 | 0 | 0 | 0 | 0 |
| 0 | 0 | 0 | 0 | 0 | 0 | 0 |
| 0 | 0 | 0 | 0 | 0 | 0 | 0 |
| 874 | 34 | | | | | |
| 330 | 4 | 0 | 2 | 0 | 0 | 0 |
| 113 | 0 | 0 | 0 | 0 | 0 | 0 |
| 178 | 3 | 0 | 1 | 0 | 0 | 1 |
| 24 | 0 | 0 | 0 | 0 | 0 | 0 |
| 71 | 0 | 0 | 0 | 0 | 0 | 0 |
| 11 | 0 | 0 | 0 | 0 | 0 | 0 |
| 11 | 0 | 0 | 0 | 0 | 0 | 0 |
| 17 | 0 | 0 | 0 | 0 | 0 | 0 |
| 7 | 0 | 0 | 0 | 0 | 0 | 0 |
| 16 | 0 | 0 | 0 | 0 | 0 | 0 |
| 9 | 0 | 0 | 0 | 0 | 0 | 0 |
| 6 | 0 | 0 | 0 | 0 | 0 | 0 |
| 793 | 19 | | | | | |
| 295 | 2 | 2 | 1 | 0 | 1 | 1 |
| 273 | 1 | 0 | 5 | 1 | 0 | 0 |
| 229 | 1 | 1 | 3 | 0 | 0 | 0 |
| 127 | 0 | 0 | 0 | 0 | 0 | 0 |
| 194 | 3 | 0 | 0 | 0 | 0 | 0 |
| 67 | 0 | 0 | 0 | 0 | 0 | 0 |
| 16 | 0 | 0 | 0 | 0 | 0 | 0 |

| | | | | | | |
|---|---|---|---|---|---|---|
| 13 | 0 | 0 | 0 | 0 | 0 | 0 |
| 0 | 0 | 0 | 0 | 0 | 0 | 0 |
| 0 | 0 | 0 | 0 | 0 | 0 | 0 |
| 0 | 0 | 0 | 0 | 0 | 0 | 0 |
| 5 | 0 | 0 | 0 | 0 | 0 | 0 |
| 1219 | 36 | | | | | |
| 690 | 9 | 6 | 20 | 1 | 1 | 1 |
| 416 | 7 | 3 | 17 | 3 | 3 | 2 |
| 387 | 2 | 3 | 14 | 1 | 1 | 0 |
| 258 | 2 | 0 | 2 | 3 | 0 | 0 |
| 372 | 6 | 3 | 1 | 0 | 0 | 1 |
| 220 | 1 | 0 | 4 | 0 | 0 | 0 |
| 227 | 2 | 0 | 2 | 0 | 0 | 1 |
| 225 | 3 | 0 | 0 | 0 | 0 | 0 |
| 191 | 4 | 0 | 2 | 0 | 0 | 0 |
| 151 | 0 | 0 | 0 | 0 | 0 | 0 |
| 64 | 0 | 0 | 0 | 0 | 0 | 0 |
| 52 | 0 | 0 | 1 | 0 | 0 | 0 |
| 3253 | 161 | | | | | |
| 567 | 15 | 0 | 5 | 0 | 0 | 1 |
| 508 | 4 | 2 | 4 | 0 | 0 | 0 |
| 334 | 2 | 0 | 0 | 0 | 0 | 0 |
| 224 | 2 | 0 | 1 | 0 | 0 | 0 |
| 128 | 0 | 0 | 0 | 0 | 0 | 0 |
| 155 | 2 | 0 | 0 | 0 | 0 | 0 |
| 111 | 0 | 0 | 0 | 0 | 0 | 0 |
| 37 | 0 | 0 | 0 | 0 | 0 | 0 |
| 20 | 0 | 0 | 0 | 0 | 0 | 0 |
| 1 | 0 | 0 | 0 | 0 | 0 | 0 |
| 155 | 1 | 0 | 2 | 0 | 0 | 0 |
| 174 | 0 | 0 | 0 | 0 | 0 | 0 |
| 2414 | 52 | | | | | |
| 582 | 13 | 4 | 5 | 2 | 0 | 4 |
| 398 | 5 | 1 | 6 | 5 | 5 | 1 |
| 436 | 6 | 1 | 1 | 3 | 1 | 1 |
| 238 | 0 | 0 | 3 | 0 | 0 | 1 |
| 242 | 20 | 2 | 0 | 1 | 0 | 0 |
| 286 | 0 | 1 | 0 | 0 | 1 | 0 |
| 189 | 0 | 0 | 0 | 0 | 1 | 0 |
| 106 | 2 | 0 | 0 | 0 | 0 | 0 |
| 14 | 0 | 0 | 0 | 0 | 0 | 0 |

| | | | | | | |
|---|---|---|---|---|---|---|
| 76 | 0 | 0 | 0 | 0 | 0 | 0 |
| 12 | 0 | 0 | 0 | 0 | 0 | 0 |
| 14 | 0 | 0 | 0 | 0 | 0 | 0 |
| 2593 | 146 | | | | | |
| 505 | 5 | 3 | 2 | 3 | 2 | 1 |
| 379 | 3 | 5 | 6 | 3 | 3 | 1 |
| 412 | 5 | 2 | 4 | 10 | 1 | 2 |
| 191 | 0 | 0 | 2 | 0 | 0 | 0 |
| 157 | 2 | 0 | 10 | 0 | 0 | 0 |
| 120 | 1 | 3 | 0 | 0 | 0 | 0 |
| 185 | 2 | 0 | 2 | 0 | 0 | 0 |
| 63 | 0 | 0 | 0 | 0 | 0 | 0 |
| 16 | 0 | 0 | 0 | 0 | 0 | 0 |
| 24 | 0 | 0 | 0 | 0 | 0 | 0 |
| 19 | 0 | 0 | 0 | 0 | 0 | 0 |
| 28 | 0 | 0 | 0 | 0 | 0 | 0 |
| 2099 | 121 | | | | | |
| 494 | 27 | 3 | 4 | 0 | 0 | 0 |
| 381 | 13 | 1 | 8 | 4 | 0 | 3 |
| 366 | 2 | 3 | 8 | 0 | 4 | 4 |
| 353 | 3 | 0 | 3 | 3 | 0 | 1 |
| 290 | 3 | 0 | 3 | 1 | 0 | 0 |
| 207 | 2 | 0 | 1 | 0 | 0 | 0 |
| 129 | 0 | 0 | 0 | 0 | 0 | 0 |
| 204 | 0 | 0 | 8 | 0 | 0 | 0 |
| 204 | 7 | 0 | 0 | 1 | 0 | 0 |
| 133 | 0 | 0 | 0 | 0 | 0 | 0 |
| 31 | 0 | 0 | 0 | 0 | 0 | 0 |
| 40 | 0 | 0 | 0 | 0 | 0 | 0 |
| 2832 | 159 | | | | | |
| 280 | 5 | 0 | 4 | 0 | 0 | 0 |
| 207 | 1 | 0 | 4 | 0 | 0 | 0 |
| 104 | 0 | 0 | 0 | 0 | 0 | 0 |
| 59 | 2 | 0 | 0 | 0 | 0 | 0 |
| 233 | 5 | 1 | 4 | 0 | 0 | 2 |
| 144 | 0 | 0 | 0 | 0 | 0 | 0 |
| 174 | 0 | 2 | 0 | 0 | 0 | 0 |
| 282 | 2 | 0 | 5 | 0 | 0 | 0 |
| 105 | 0 | 0 | 0 | 0 | 0 | 0 |
| 70 | 0 | 0 | 0 | 0 | 0 | 0 |
| 45 | 0 | 0 | 0 | 0 | 0 | 0 |

| | | | | | | |
|---|---|---|---|---|---|---|
| 27 | 0 | 0 | 0 | 0 | 0 | 0 |
| 1730 | 54 | | | | | |
| 557 | 16 | 6 | 4 | 7 | 0 | 1 |
| 585 | 8 | 9 | 12 | 0 | 1 | 2 |
| 555 | 16 | 5 | 12 | 2 | 3 | 3 |
| 580 | 14 | 7 | 4 | 3 | 3 | 2 |
| 558 | 2 | 5 | 8 | 3 | 3 | 5 |
| 562 | 4 | 4 | 4 | 4 | 1 | 1 |
| 455 | 5 | 2 | 7 | 2 | 0 | 4 |
| 515 | 2 | 7 | 5 | 1 | 3 | 3 |
| 359 | 4 | 3 | 4 | 0 | 1 | 0 |
| 397 | 3 | 5 | 11 | 0 | 0 | 0 |
| 383 | 2 | 1 | 7 | 0 | 0 | 1 |
| 348 | 13 | 3 | 8 | 0 | 1 | 1 |
| 5854 | 399 | | | | | |
| 660 | 3 | 5 | 8 | 2 | 2 | 1 |
| 507 | 6 | 15 | 6 | 2 | 1 | 2 |
| 404 | 2 | 4 | 4 | 0 | 0 | 1 |
| 370 | 5 | 7 | 3 | 2 | 0 | 2 |
| 326 | 8 | 3 | 9 | 3 | 1 | 0 |
| 307 | 7 | 4 | 5 | 0 | 0 | 0 |
| 227 | 0 | 0 | 1 | 0 | 0 | 0 |
| 211 | 0 | 1 | 0 | 0 | 0 | 0 |
| 139 | 0 | 0 | 0 | 0 | 0 | 0 |
| 157 | 0 | 0 | 0 | 0 | 0 | 0 |
| 223 | 3 | 5 | 2 | 0 | 0 | 1 |
| 47 | 0 | 0 | 0 | 0 | 4 | 0 |
| 3578 | 192 | | | | | |
| 553 | 14 | 3 | 14 | 4 | 1 | 4 |
| 382 | 7 | 3 | 19 | 0 | 1 | 1 |
| 283 | 6 | 0 | 8 | 0 | 0 | 0 |
| 301 | 6 | 1 | 17 | 2 | 0 | 1 |
| 210 | 3 | 0 | 2 | 0 | 0 | 0 |
| 216 | 9 | 0 | 8 | 0 | 0 | 0 |
| 150 | 0 | 0 | 4 | 0 | 0 | 0 |
| 139 | 3 | 0 | 5 | 0 | 0 | 0 |
| 40 | 0 | 0 | 0 | 0 | 0 | 0 |
| 39 | 0 | 0 | 0 | 0 | 0 | 0 |
| 57 | 0 | 0 | 0 | 0 | 0 | 0 |
| 93 | 0 | 0 | 0 | 0 | 0 | 0 |
| 2463 | 167 | | | | | |

| | | | | | | |
|---|---|---|---|---|---|---|
| 654 | 9 | 2 | 9 | 1 | 0 | 1 |
| 502 | 7 | 8 | 6 | 0 | 1 | 1 |
| 354 | 2 | 4 | 2 | 0 | 0 | 0 |
| 378 | 1 | 0 | 0 | 1 | 1 | 4 |
| 368 | 7 | 0 | 0 | 0 | 0 | 0 |
| 132 | 0 | 0 | 0 | 0 | 0 | 0 |
| 71 | 0 | 0 | 0 | 0 | 0 | 0 |
| 20 | 0 | 0 | 0 | 0 | 0 | 0 |
| 26 | 0 | 0 | 0 | 0 | 0 | 0 |
| 36 | 0 | 0 | 0 | 0 | 0 | 0 |
| 15 | 0 | 0 | 0 | 0 | 0 | 0 |
| 7 | 0 | 0 | 0 | 0 | 0 | 0 |
| 2563 | 96 | | | | | |
| 669 | 9 | 4 | 6 | 2 | 0 | 4 |
| 442 | 3 | 1 | 5 | 1 | 0 | 2 |
| 344 | 1 | 2 | 4 | 1 | 2 | 0 |
| 261 | 3 | 0 | 4 | 0 | 0 | 0 |
| 164 | 2 | 0 | 0 | 0 | 0 | 0 |
| 90 | 1 | 0 | 0 | 0 | 0 | 0 |
| 56 | 0 | 0 | 0 | 0 | 0 | 0 |
| 20 | 0 | 0 | 0 | 0 | 0 | 0 |
| 270 | 3 | 0 | 2 | 0 | 0 | 0 |
| 24 | 0 | 0 | 0 | 0 | 0 | 0 |
| 3 | 0 | 0 | 0 | 0 | 0 | 0 |
| 86 | 0 | 0 | 0 | 0 | 0 | 0 |
| 2429 | 103 | | | | | |
| 514 | 3 | 0 | 2 | 1 | 0 | 1 |
| 534 | 4 | 4 | 3 | 0 | 3 | 4 |
| 421 | 2 | 0 | 2 | 1 | 0 | 0 |
| 423 | 1 | 0 | 3 | 0 | 0 | 4 |
| 327 | 0 | 0 | 1 | 6 | 0 | 2 |
| 271 | 0 | 0 | 0 | 0 | 0 | 0 |
| 431 | 1 | 0 | 1 | 0 | 1 | 0 |
| 22 | 0 | 0 | 0 | 0 | 0 | 0 |
| 5 | 0 | 0 | 0 | 0 | 0 | 0 |
| 4 | 0 | 0 | 0 | 0 | 0 | 0 |
| 8 | 0 | 0 | 0 | 0 | 0 | 0 |
| 2 | 0 | 0 | 0 | 0 | 0 | 0 |
| 2962 | 83 | | | | | |
| 873 | 4 | 4 | 10 | 3 | 0 | 3 |
| 609 | 1 | 2 | 4 | 4 | 0 | 1 |

| | | | | | | |
|---|---|---|---|---|---|---|
| 458 | 3 | 0 | 3 | 0 | 8 | 0 |
| 425 | 3 | 0 | 0 | 2 | 1 | 0 |
| 448 | 0 | 0 | 4 | 0 | 0 | 0 |
| 391 | 16 | 4 | 10 | 1 | 0 | 1 |
| 391 | 1 | 0 | 0 | 0 | 0 | 0 |
| 128 | 1 | 0 | 0 | 0 | 0 | 0 |
| 60 | 0 | 0 | 0 | 0 | 0 | 0 |
| 1 | 0 | 0 | 0 | 0 | 0 | 0 |
| 3 | 0 | 0 | 0 | 0 | 0 | 0 |
| 45 | 1 | 0 | 0 | 1 | 0 | 0 |
| 3832 | 156 | | | | | |
| 772 | 6 | 1 | 4 | 4 | 0 | 0 |
| 380 | 0 | 0 | 2 | 0 | 0 | 2 |
| 366 | 2 | 0 | 2 | 0 | 0 | 0 |
| 92 | 0 | 0 | 0 | 0 | 0 | 0 |
| 183 | 0 | 0 | 0 | 0 | 0 | 0 |
| 272 | 0 | 0 | 2 | 0 | 0 | 0 |
| 236 | 2 | 1 | 0 | 0 | 0 | 0 |
| 22 | 0 | 0 | 0 | 0 | 0 | 0 |
| 7 | 0 | 0 | 0 | 0 | 0 | 0 |
| 3 | 0 | 0 | 0 | 0 | 0 | 0 |
| 14 | 0 | 0 | 0 | 0 | 0 | 0 |
| 16 | 0 | 0 | 0 | 0 | 0 | 0 |
| 2363 | 45 | | | | | |
| 514 | 0 | 1 | 6 | 2 | 0 | 0 |
| 211 | 1 | 1 | 2 | 0 | 0 | 0 |
| 51 | 0 | 0 | 0 | 0 | 0 | 0 |
| 54 | 0 | 0 | 0 | 0 | 0 | 0 |
| 27 | 0 | 0 | 0 | 0 | 0 | 0 |
| 25 | 0 | 0 | 0 | 0 | 0 | 0 |
| 12 | 0 | 0 | 0 | 0 | 0 | 0 |
| 136 | 0 | 0 | 0 | 0 | 0 | 0 |
| 14 | 0 | 0 | 0 | 0 | 0 | 0 |
| 122 | 0 | 0 | 0 | 0 | 0 | 0 |
| 138 | 3 | 0 | 0 | 0 | 0 | 0 |
| 100 | 0 | 0 | 0 | 0 | 0 | 0 |
| 1404 | 22 | | | | | |
| 624 | 8 | 0 | 5 | 2 | 0 | 0 |
| 649 | 4 | 0 | 3 | 0 | 0 | 0 |
| 604 | 3 | 4 | 3 | 0 | 0 | 0 |
| 428 | 6 | 2 | 9 | 1 | 0 | 2 |

| | | | | | | |
|---|---|---|---|---|---|---|
| 446 | 4 | 1 | 6 | 3 | 0 | 2 |
| 425 | 4 | 0 | 1 | 5 | 0 | 0 |
| 471 | 2 | 0 | 0 | 0 | 0 | 0 |
| 297 | 0 | 0 | 2 | 0 | 0 | 0 |
| 162 | 0 | 0 | 0 | 0 | 0 | 0 |
| 154 | 0 | 0 | 0 | 0 | 0 | 0 |
| 23 | 0 | 0 | 0 | 0 | 0 | 0 |
| 1 | 0 | 0 | 0 | 0 | 0 | 0 |
| 4284 | 134 | | | | | |
| 877 | 1 | 0 | 3 | 0 | 0 | 0 |
| 415 | 0 | 1 | 0 | 0 | 0 | 0 |
| 386 | 1 | 0 | 0 | 0 | 0 | 0 |
| 410 | 4 | 2 | 2 | 0 | 0 | 0 |
| 467 | 0 | 0 | 1 | 2 | 0 | 3 |
| 275 | 1 | 0 | 0 | 0 | 0 | 0 |
| 222 | 0 | 0 | 0 | 0 | 0 | 0 |
| 214 | 1 | 0 | 0 | 3 | 0 | 0 |
| 218 | 1 | 0 | 0 | 0 | 0 | 0 |
| 177 | 0 | 0 | 0 | 0 | 0 | 0 |
| 208 | 0 | 0 | 3 | 0 | 0 | 0 |
| 214 | 0 | 0 | 0 | 0 | 0 | 0 |
| 4083 | 52 | | | | | |
| 473 | 5 | 1 | 7 | 1 | 0 | 0 |
| 222 | 0 | 3 | 2 | 0 | 0 | 0 |
| 237 | 0 | 0 | 3 | 0 | 0 | 0 |
| 346 | 3 | 0 | 3 | 0 | 0 | 2 |
| 220 | 0 | 0 | 1 | 2 | 0 | 1 |
| 69 | 0 | 0 | 0 | 0 | 0 | 0 |
| 185 | 0 | 0 | 1 | 0 | 0 | 0 |
| 111 | 0 | 0 | 0 | 0 | 0 | 0 |
| 162 | 4 | 0 | 0 | 1 | 0 | 0 |
| 17 | 0 | 0 | 0 | 0 | 0 | 0 |
| 25 | 0 | 0 | 0 | 0 | 0 | 0 |
| 201 | 0 | 0 | 6 | 0 | 0 | 0 |
| 2268 | 67 | | | | | |
| 644 | 10 | 6 | 14 | 6 | 3 | 5 |
| 496 | 8 | 7 | 9 | 1 | 0 | 4 |
| 456 | 7 | 22 | 10 | 6 | 5 | 6 |
| 541 | 3 | 2 | 6 | 3 | 5 | 5 |
| 322 | 2 | 14 | 20 | 1 | 2 | 3 |
| 358 | 5 | 4 | 6 | 4 | 2 | 0 |

| | | | | | | |
|---|---|---|---|---|---|---|
| 240 | 0 | 0 | 0 | 0 | 0 | 0 |
| 349 | 1 | 3 | 2 | 0 | 0 | 1 |
| 229 | 2 | 3 | 5 | 0 | 0 | 0 |
| 224 | 0 | 0 | 0 | 0 | 1 | 1 |
| 17 | 0 | 0 | 0 | 0 | 0 | 0 |
| 62 | 0 | 0 | 1 | 0 | 0 | 0 |
| 3938 | 303 | | | | | |
| 725 | 5 | 3 | 4 | 5 | 1 | 2 |
| 639 | 2 | 8 | 6 | 2 | 2 | 2 |
| 660 | 4 | 3 | 4 | 3 | 0 | 6 |
| 743 | 2 | 3 | 2 | 3 | 0 | 0 |
| 602 | 2 | 3 | 10 | 0 | 0 | 2 |
| 687 | 3 | 2 | 8 | 1 | 1 | 1 |
| 669 | 5 | 4 | 6 | 0 | 0 | 2 |
| 412 | 8 | 4 | 10 | 1 | 1 | 0 |
| 504 | 0 | 2 | 2 | 0 | 0 | 0 |
| 373 | 0 | 0 | 3 | 0 | 0 | 0 |
| 417 | 2 | 2 | 1 | 0 | 1 | 3 |
| 520 | 2 | 5 | 13 | 2 | 1 | 6 |
| 6951 | 281 | | | | | |
| 577 | 4 | 3 | 4 | 1 | 1 | 1 |
| 309 | 1 | 0 | 0 | 2 | 0 | 0 |
| 380 | 2 | 1 | 3 | 0 | 0 | 0 |
| 328 | 4 | 2 | 3 | 0 | 0 | 0 |
| 209 | 1 | 0 | 2 | 0 | 0 | 0 |
| 30 | 0 | 0 | 0 | 0 | 0 | 0 |
| 144 | 1 | 0 | 1 | 0 | 0 | 0 |
| 121 | 0 | 0 | 0 | 0 | 0 | 0 |
| 44 | 0 | 0 | 0 | 0 | 0 | 0 |
| 105 | 0 | 0 | 1 | 0 | 0 | 0 |
| 134 | 0 | 0 | 0 | 0 | 0 | 0 |
| 26 | 0 | 0 | 0 | 0 | 0 | 0 |
| 2407 | 63 | | | | | |
| 708 | 6 | 3 | 6 | 0 | 1 | 1 |
| 466 | 8 | 4 | 9 | 3 | 1 | 2 |
| 488 | 5 | 9 | 3 | 2 | 4 | 4 |
| 339 | 9 | 5 | 6 | 0 | 2 | 2 |
| 433 | 1 | 3 | 6 | 1 | 0 | 0 |
| 399 | 5 | 1 | 4 | 0 | 2 | 1 |
| 330 | 10 | 7 | 6 | 2 | 0 | 0 |
| 350 | 0 | 0 | 1 | 0 | 0 | 0 |

| | | | | | | |
|---|---|---|---|---|---|---|
| 147 | 1 | 0 | 0 | 0 | 0 | 0 |
| 64 | 0 | 0 | 0 | 0 | 0 | 0 |
| 13 | 0 | 0 | 0 | 0 | 0 | 0 |
| 19 | 0 | 0 | 0 | 0 | 0 | 0 |
| 3756 | 235 | | | | | |
| 706 | 3 | 4 | 7 | 1 | 2 | 3 |
| 446 | 1 | 0 | 5 | 1 | 0 | 5 |
| 391 | 8 | 2 | 1 | 2 | 1 | 1 |
| 318 | 1 | 2 | 5 | 1 | 0 | 3 |
| 292 | 1 | 0 | 1 | 5 | 0 | 0 |
| 190 | 0 | 0 | 1 | 1 | 1 | 0 |
| 130 | 1 | 0 | 0 | 0 | 0 | 0 |
| 22 | 0 | 0 | 0 | 0 | 0 | 0 |
| 17 | 0 | 0 | 0 | 0 | 0 | 0 |
| 30 | 0 | 0 | 0 | 0 | 0 | 0 |
| 8 | 0 | 0 | 0 | 0 | 0 | 0 |
| 18 | 0 | 0 | 0 | 0 | 0 | 0 |
| 2568 | 125 | | | | | |
| 644 | 2 | 2 | 3 | 1 | 0 | 2 |
| 450 | 0 | 0 | 0 | 0 | 1 | 0 |
| 394 | 2 | 0 | 4 | 1 | 0 | 0 |
| 387 | 0 | 1 | 0 | 0 | 0 | 0 |
| 197 | 2 | 0 | 0 | 2 | 0 | 0 |
| 73 | 0 | 0 | 0 | 0 | 0 | 0 |
| 120 | 0 | 0 | 0 | 0 | 0 | 0 |
| 57 | 0 | 0 | 0 | 0 | 0 | 0 |
| 74 | 0 | 0 | 0 | 0 | 0 | 0 |
| 30 | 0 | 0 | 0 | 0 | 0 | 0 |
| 122 | 0 | 0 | 1 | 0 | 0 | 0 |
| 9 | 0 | 0 | 0 | 0 | 0 | 0 |
| 2557 | 57 | | | | | |
| 352 | 3 | 0 | 5 | 1 | 0 | 1 |
| 282 | 2 | 7 | 13 | 0 | 0 | 0 |
| 333 | 3 | 3 | 4 | 0 | 1 | 1 |
| 321 | 1 | 5 | 4 | 0 | 1 | 0 |
| 365 | 5 | 1 | 7 | 5 | 3 | 4 |
| 233 | 5 | 0 | 5 | 0 | 0 | 0 |
| 188 | 8 | 0 | 0 | 0 | 0 | 0 |
| 250 | 6 | 7 | 14 | 0 | 2 | 2 |
| 133 | 0 | 0 | 4 | 0 | 1 | 0 |
| 69 | 0 | 0 | 0 | 0 | 0 | 0 |

| | | | | | | |
|---|---|---|---|---|---|---|
| 100 | 0 | 0 | 0 | 0 | 0 | 0 |
| 43 | 0 | 0 | 0 | 0 | 0 | 0 |
| 2669 | 183 | | | | | |
| 348 | 1 | 2 | 4 | 0 | 0 | 0 |
| 258 | 3 | 0 | 2 | 1 | 0 | 0 |
| 166 | 0 | 0 | 3 | 0 | 0 | 0 |
| 75 | 0 | 0 | 2 | 0 | 0 | 0 |
| 165 | 1 | 0 | 0 | 0 | 0 | 0 |
| 210 | 0 | 1 | 1 | 0 | 0 | 0 |
| 191 | 1 | 0 | 0 | 0 | 0 | 3 |
| 144 | 2 | 0 | 0 | 0 | 0 | 0 |
| 159 | 0 | 0 | 4 | 0 | 0 | 0 |
| 109 | 2 | 0 | 1 | 0 | 0 | 0 |
| 119 | 0 | 0 | 0 | 0 | 0 | 0 |
| 111 | 0 | 0 | 0 | 0 | 0 | 0 |
| 2055 | 49 | | | | | |
| 529 | 11 | 2 | 10 | 2 | 0 | 1 |
| 204 | 0 | 1 | 1 | 0 | 0 | 0 |
| 314 | 4 | 0 | 2 | 0 | 0 | 1 |
| 170 | 1 | 0 | 5 | 0 | 0 | 0 |
| 281 | 1 | 0 | 2 | 1 | 0 | 0 |
| 221 | 0 | 1 | 2 | 0 | 2 | 0 |
| 239 | 0 | 1 | 0 | 0 | 0 | 0 |
| 174 | 2 | 0 | 0 | 0 | 0 | 0 |
| 79 | 0 | 0 | 0 | 0 | 0 | 0 |
| 250 | 3 | 1 | 11 | 1 | 0 | 0 |
| 170 | 2 | 0 | 3 | 0 | 0 | 1 |
| 98 | 0 | 0 | 0 | 0 | 0 | 0 |
| 2729 | 94 | | | | | |
| 506 | 15 | 3 | 6 | 1 | 0 | 2 |
| 429 | 6 | 3 | 1 | 3 | 7 | 3 |
| 345 | 1 | 0 | 1 | 5 | 0 | 0 |
| 512 | 4 | 3 | 5 | 2 | 4 | 5 |
| 446 | 0 | 0 | 13 | 1 | 2 | 0 |
| 386 | 6 | 1 | 0 | 3 | 2 | 0 |
| 366 | 1 | 0 | 2 | 1 | 0 | 0 |
| 337 | 6 | 0 | 3 | 0 | 0 | 0 |
| 243 | 16 | 5 | 2 | 1 | 2 | 1 |
| 288 | 4 | 2 | 1 | 0 | 0 | 0 |
| 271 | 0 | 0 | 1 | 1 | 0 | 1 |
| 190 | 4 | 1 | 0 | 0 | 1 | 0 |

| | | | | | | |
|---|---|---|---|---|---|---|
| 4319 | 257 | | | | | |
| 366 | 3 | 1 | 0 | 0 | 0 | 0 |
| 372 | 2 | 0 | 0 | 0 | 0 | 0 |
| 276 | 1 | 2 | 5 | 1 | 0 | 4 |
| 358 | 6 | 3 | 2 | 2 | 4 | 4 |
| 296 | 3 | 0 | 2 | 1 | 0 | 0 |
| 333 | 3 | 0 | 0 | 0 | 1 | 1 |
| 151 | 0 | 0 | 0 | 0 | 0 | 0 |
| 179 | 2 | 0 | 2 | 1 | 0 | 0 |
| 323 | 4 | 2 | 1 | 2 | 0 | 0 |
| 239 | 0 | 0 | 1 | 1 | 2 | 1 |
| 78 | 0 | 0 | 1 | 0 | 0 | 0 |
| 127 | 0 | 0 | 0 | 0 | 0 | 0 |
| 3098 | 112 | | | | | |
| 376 | 5 | 3 | 0 | 0 | 0 | 0 |
| 196 | 1 | 2 | 1 | 0 | 0 | 0 |
| 105 | 0 | 0 | 2 | 0 | 0 | 0 |
| 36 | 0 | 0 | 0 | 0 | 0 | 0 |
| 209 | 4 | 0 | 2 | 1 | 1 | 1 |
| 118 | 0 | 2 | 1 | 0 | 0 | 0 |
| 30 | 0 | 0 | 0 | 0 | 0 | 0 |
| 7 | 0 | 0 | 0 | 0 | 0 | 0 |
| 16 | 0 | 0 | 0 | 0 | 0 | 0 |
| 0 | 0 | 0 | 0 | 0 | 0 | 0 |
| 27 | 0 | 0 | 0 | 0 | 0 | 0 |
| 85 | 0 | 0 | 0 | 0 | 0 | 0 |
| 1205 | 32 | | | | | |
| 312 | 2 | 0 | 3 | 3 | 0 | 0 |
| 281 | 2 | 3 | 6 | 0 | 0 | 0 |
| 211 | 3 | 3 | 9 | 0 | 0 | 0 |
| 340 | 1 | 0 | 3 | 1 | 1 | 2 |
| 225 | 0 | 1 | 21 | 0 | 0 | 0 |
| 194 | 2 | 0 | 1 | 1 | 0 | 0 |
| 205 | 0 | 1 | 1 | 0 | 0 | 1 |
| 123 | 0 | 0 | 0 | 0 | 0 | 0 |
| 178 | 1 | 0 | 2 | 0 | 0 | 0 |
| 187 | 7 | 0 | 1 | 0 | 1 | 0 |
| 153 | 2 | 2 | 0 | 0 | 0 | 0 |
| 141 | 0 | 0 | 0 | 0 | 0 | 0 |
| 2550 | 105 | | | | | |
| 407 | 4 | 1 | 2 | 0 | 0 | 1 |

| | | | | | | |
|---|---|---|---|---|---|---|
| 287 | 1 | 0 | 11 | 0 | 0 | 0 |
| 234 | 0 | 1 | 4 | 2 | 0 | 0 |
| 165 | 0 | 0 | 1 | 0 | 0 | 0 |
| 145 | 4 | 0 | 0 | 0 | 0 | 0 |
| 235 | 0 | 1 | 2 | 0 | 0 | 1 |
| 127 | 0 | 0 | 0 | 0 | 0 | 0 |
| 47 | 0 | 0 | 0 | 0 | 0 | 0 |
| 10 | 0 | 0 | 0 | 0 | 0 | 0 |
| 86 | 0 | 0 | 1 | 0 | 0 | 0 |
| 5 | 0 | 0 | 0 | 0 | 0 | 0 |
| 20 | 0 | 0 | 0 | 0 | 0 | 0 |
| 1768 | 48 | | | | | |
| 378 | 1 | 0 | 4 | 0 | 0 | 0 |
| 136 | 0 | 0 | 0 | 0 | 0 | 0 |
| 152 | 0 | 0 | 1 | 0 | 0 | 0 |
| 53 | 0 | 0 | 0 | 0 | 0 | 0 |
| 42 | 0 | 0 | 0 | 0 | 0 | 0 |
| 185 | 8 | 1 | 0 | 0 | 0 | 0 |
| 48 | 0 | 0 | 0 | 0 | 0 | 0 |
| 95 | 3 | 0 | 0 | 0 | 0 | 0 |
| 53 | 0 | 0 | 0 | 0 | 0 | 0 |
| 47 | 0 | 0 | 0 | 0 | 0 | 0 |
| 37 | 0 | 0 | 0 | 0 | 0 | 0 |
| 179 | 0 | 0 | 13 | 0 | 0 | 0 |
| 1405 | 35 | | | | | |
| 129 | 13 | 0 | 0 | 1 | 0 | 0 |
| 60 | 2 | 0 | 0 | 0 | 0 | 0 |
| 106 | 8 | 0 | 0 | 0 | 0 | 0 |
| 84 | 0 | 0 | 0 | 0 | 0 | 0 |
| 62 | 4 | 0 | 0 | 0 | 0 | 0 |
| 158 | 0 | 21 | 7 | 0 | 0 | 6 |
| 1 | 0 | 0 | 0 | 0 | 0 | 0 |
| 10 | 0 | 0 | 0 | 0 | 0 | 0 |
| 7 | 0 | 0 | 0 | 0 | 0 | 0 |
| 31 | 0 | 0 | 0 | 0 | 0 | 0 |
| 110 | 0 | 0 | 0 | 0 | 0 | 1 |
| 37 | 0 | 0 | 0 | 0 | 0 | 1 |
| 795 | 110 | | | | | |
| 331 | 12 | 2 | 7 | 4 | 0 | 3 |
| 291 | 0 | 1 | 5 | 2 | 1 | 1 |
| 195 | 3 | 4 | 13 | 0 | 2 | 2 |

| | | | | | | |
|---|---|---|---|---|---|---|
| 147 | 0 | 1 | 2 | 0 | 0 | 0 |
| 183 | 3 | 1 | 1 | 2 | 0 | 2 |
| 102 | 0 | 0 | 0 | 0 | 0 | 1 |
| 157 | 2 | 5 | 4 | 0 | 0 | 5 |
| 78 | 0 | 0 | 0 | 0 | 0 | 1 |
| 18 | 0 | 0 | 0 | 0 | 0 | 0 |
| 29 | 0 | 0 | 0 | 0 | 0 | 0 |
| 80 | 0 | 0 | 0 | 0 | 0 | 0 |
| 160 | 1 | 0 | 1 | 2 | 0 | 1 |
| 1771 | 138 | | | | | |
| 202 | 3 | 1 | 0 | 6 | 0 | 1 |
| 267 | 0 | 0 | 4 | 2 | 1 | 0 |
| 165 | 3 | 1 | 4 | 0 | 0 | 1 |
| 108 | 3 | 0 | 3 | 0 | 0 | 0 |
| 210 | 3 | 0 | 1 | 0 | 0 | 0 |
| 98 | 0 | 0 | 1 | 0 | 0 | 0 |
| 251 | 2 | 3 | 2 | 0 | 1 | 0 |
| 66 | 0 | 6 | 4 | 0 | 0 | 0 |
| 128 | 2 | 0 | 2 | 1 | 0 | 0 |
| 98 | 0 | 0 | 0 | 0 | 0 | 0 |
| 1 | 3 | 0 | 0 | 0 | 0 | 0 |
| 52 | 11 | 0 | 0 | 0 | 0 | 0 |
| 1646 | 93 | | | | | |
| 358 | 3 | 0 | 4 | 0 | 0 | 3 |
| 232 | 2 | 1 | 0 | 2 | 0 | 1 |
| 213 | 0 | 0 | 0 | 0 | 0 | 0 |
| 176 | 3 | 0 | 2 | 1 | 0 | 1 |
| 174 | 4 | 2 | 2 | 0 | 0 | 1 |
| 201 | 0 | 0 | 1 | 0 | 0 | 3 |
| 106 | 0 | 0 | 0 | 0 | 0 | 0 |
| 35 | 0 | 0 | 0 | 0 | 0 | 0 |
| 197 | 0 | 3 | 2 | 0 | 0 | 1 |
| 103 | 0 | 0 | 1 | 0 | 0 | 0 |
| 6 | 0 | 0 | 0 | 0 | 0 | 0 |
| 7 | 0 | 0 | 0 | 0 | 0 | 0 |
| 1808 | 74 | | | | | |
| 309 | 1 | 2 | 2 | 0 | 0 | 0 |
| 297 | 3 | 3 | 5 | 1 | 3 | 1 |
| 148 | 0 | 0 | 0 | 0 | 0 | 0 |
| 254 | 3 | 1 | 1 | 1 | 1 | 3 |
| 221 | 0 | 1 | 0 | 0 | 0 | 0 |

| | | | | | | |
|---|---|---|---|---|---|---|
| 175 | 0 | 0 | 0 | 0 | 0 | 0 |
| 269 | 6 | 0 | 0 | 0 | 0 | 1 |
| 164 | 3 | 0 | 0 | 0 | 0 | 0 |
| 68 | 0 | 0 | 0 | 0 | 0 | 0 |
| 95 | 0 | 0 | 0 | 0 | 0 | 0 |
| 65 | 0 | 0 | 0 | 0 | 0 | 0 |
| 153 | 0 | 0 | 0 | 0 | 0 | 0 |
| 2218 | 57 | | | | | |
| 322 | 3 | 0 | 0 | 1 | 0 | 1 |
| 291 | 1 | 2 | 4 | 0 | 0 | 1 |
| 255 | 4 | 0 | 2 | 0 | 0 | 0 |
| 295 | 12 | 1 | 0 | 4 | 0 | 0 |
| 205 | 0 | 0 | 2 | 0 | 0 | 0 |
| 117 | 0 | 0 | 0 | 0 | 0 | 0 |
| 25 | 0 | 0 | 0 | 0 | 0 | 0 |
| 137 | 0 | 0 | 0 | 0 | 0 | 0 |
| 58 | 0 | 0 | 0 | 0 | 0 | 0 |
| 25 | 0 | 0 | 0 | 0 | 0 | 0 |
| 29 | 0 | 0 | 0 | 0 | 0 | 0 |
| 25 | 0 | 0 | 0 | 0 | 0 | 0 |
| 1784 | 45 | | | | | |
| 496 | 7 | 3 | 12 | 2 | 0 | 1 |
| 422 | 5 | 0 | 2 | 0 | 0 | 2 |
| 409 | 1 | 0 | 2 | 6 | 4 | 4 |
| 456 | 0 | 6 | 3 | 2 | 1 | 3 |
| 256 | 3 | 0 | 1 | 0 | 3 | 1 |
| 316 | 5 | 2 | 2 | 4 | 0 | 0 |
| 319 | 0 | 2 | 0 | 0 | 0 | 0 |
| 244 | 5 | 0 | 2 | 1 | 0 | 1 |
| 191 | 0 | 0 | 0 | 0 | 0 | 0 |
| 140 | 12 | 0 | 0 | 0 | 0 | 0 |
| 309 | 0 | 0 | 1 | 1 | 0 | 0 |
| 209 | 2 | 2 | 1 | 0 | 0 | 0 |
| 3767 | 155 | | | | | |
| 543 | 2 | 1 | 1 | 3 | 2 | 3 |
| 382 | 2 | 2 | 0 | 0 | 0 | 2 |
| 397 | 2 | 0 | 4 | 0 | 0 | 0 |
| 400 | 2 | 0 | 0 | 0 | 0 | 1 |
| 341 | 1 | 0 | 2 | 0 | 3 | 0 |
| 229 | 1 | 0 | 0 | 0 | 0 | 0 |
| 246 | 1 | 0 | 0 | 0 | 0 | 0 |

| | | | | | | |
|---|---|---|---|---|---|---|
| 229 | 5 | 0 | 0 | 0 | 0 | 0 |
| 108 | 0 | 0 | 0 | 1 | 0 | 0 |
| 66 | 0 | 0 | 0 | 0 | 0 | 0 |
| 71 | 0 | 0 | 0 | 0 | 0 | 0 |
| 199 | 0 | 0 | 0 | 0 | 0 | 0 |
| 3211 | 71 | | | | | |
| 510 | 0 | 0 | 2 | 6 | 0 | 0 |
| 425 | 0 | 1 | 1 | 0 | 0 | 1 |
| 220 | 1 | 0 | 0 | 0 | 2 | 0 |
| 357 | 1 | 0 | 1 | 0 | 0 | 0 |
| 280 | 0 | 0 | 0 | 0 | 0 | 0 |
| 56 | 0 | 0 | 0 | 0 | 0 | 0 |
| 230 | 0 | 0 | 0 | 0 | 0 | 0 |
| 313 | 13 | 0 | 0 | 3 | 0 | 0 |
| 265 | 9 | 2 | 2 | 2 | 0 | 0 |
| 210 | 0 | 0 | 0 | 0 | 0 | 0 |
| 75 | 0 | 0 | 0 | 0 | 0 | 0 |
| 44 | 0 | 0 | 0 | 0 | 0 | 0 |
| 2985 | 66 | | | | | |
| 839 | 7 | 3 | 7 | 2 | 0 | 6 |
| 525 | 5 | 5 | 6 | 0 | 0 | 4 |
| 460 | 3 | 2 | 5 | 1 | 0 | 4 |
| 507 | 3 | 3 | 6 | 1 | 2 | 0 |
| 329 | 0 | 0 | 3 | 0 | 0 | 1 |
| 381 | 0 | 2 | 4 | 0 | 2 | 1 |
| 299 | 1 | 1 | 3 | 0 | 0 | 0 |
| 69 | 0 | 0 | 0 | 0 | 0 | 0 |
| 480 | 0 | 0 | 6 | 0 | 0 | 0 |
| 84 | 0 | 0 | 0 | 0 | 0 | 0 |
| 455 | 7 | 1 | 2 | 1 | 0 | 0 |
| 177 | 0 | 0 | 0 | 0 | 0 | 0 |
| 4605 | 150 | | | | | |
| 663 | 7 | 2 | 1 | 0 | 0 | 0 |
| 512 | 3 | 1 | 6 | 0 | 0 | 1 |
| 612 | 5 | 2 | 0 | 2 | 0 | 1 |
| 400 | 1 | 0 | 0 | 2 | 0 | 0 |
| 394 | 0 | 0 | 1 | 0 | 0 | 2 |
| 465 | 4 | 0 | 0 | 0 | 0 | 1 |
| 627 | 2 | 0 | 0 | 0 | 0 | 0 |
| 524 | 1 | 0 | 1 | 1 | 0 | 0 |
| 533 | 0 | 0 | 0 | 0 | 0 | 0 |

| | | | | | | |
|---|---|---|---|---|---|---|
| 267 | 2 | 0 | 0 | 0 | 0 | 0 |
| 330 | 0 | 0 | 4 | 0 | 0 | 0 |
| 272 | 0 | 1 | 2 | 0 | 0 | 1 |
| 5599 | 87 | | | | | |
| 563 | 1 | 1 | 1 | 0 | 0 | 0 |
| 243 | 0 | 0 | 0 | 0 | 0 | 0 |
| 281 | 0 | 0 | 0 | 0 | 0 | 0 |
| 265 | 0 | 0 | 0 | 0 | 0 | 0 |
| 151 | 0 | 0 | 0 | 0 | 0 | 0 |
| 176 | 5 | 0 | 0 | 0 | 0 | 0 |
| 71 | 0 | 0 | 0 | 0 | 0 | 0 |
| 89 | 0 | 0 | 0 | 0 | 0 | 0 |
| 51 | 0 | 0 | 0 | 0 | 0 | 0 |
| 48 | 0 | 0 | 0 | 0 | 0 | 0 |
| 40 | 0 | 0 | 0 | 0 | 0 | 0 |
| 0 | 0 | 0 | 0 | 0 | 0 | 0 |
| 1978 | 11 | | | | | |
| 536 | 9 | 5 | 17 | 3 | 0 | 2 |
| 359 | 13 | 2 | 5 | 2 | 0 | 2 |
| 339 | 10 | 2 | 7 | 1 | 0 | 0 |
| 363 | 6 | 7 | 1 | 2 | 3 | 0 |
| 324 | 5 | 2 | 5 | 1 | 4 | 0 |
| 149 | 4 | 1 | 0 | 0 | 0 | 0 |
| 56 | 4 | 0 | 0 | 0 | 0 | 0 |
| 31 | 0 | 0 | 0 | 0 | 0 | 0 |
| 139 | 0 | 0 | 0 | 0 | 0 | 1 |
| 183 | 1 | 0 | 3 | 0 | 0 | 0 |
| 19 | 0 | 0 | 0 | 0 | 0 | 0 |
| 21 | 3 | 0 | 0 | 0 | 0 | 0 |
| 2519 | 183 | | | | | |
| 640 | 9 | 3 | 9 | 3 | 0 | 2 |
| 571 | 7 | 6 | 12 | 2 | 1 | 5 |
| 337 | 10 | 2 | 4 | 5 | 0 | 2 |
| 290 | 4 | 2 | 6 | 0 | 0 | 0 |
| 293 | 2 | 3 | 0 | 0 | 1 | 1 |
| 343 | 9 | 2 | 2 | 1 | 2 | 0 |
| 312 | 1 | 0 | 2 | 0 | 2 | 1 |
| 384 | 0 | 3 | 2 | 0 | 0 | 1 |
| 360 | 2 | 0 | 2 | 2 | 0 | 0 |
| 254 | 0 | 4 | 13 | 0 | 0 | 1 |
| 45 | 0 | 0 | 2 | 0 | 0 | 0 |

| | | | | | | |
|---|---|---|---|---|---|---|
| 118 | 0 | 0 | 1 | 2 | 0 | 0 |
| 3947 | 217 | | | | | |
| 302 | 1 | 0 | 5 | 0 | 0 | 0 |
| 28 | 0 | 0 | 0 | 0 | 0 | 0 |
| 38 | 0 | 0 | 1 | 0 | 0 | 0 |
| 139 | 0 | 0 | 1 | 1 | 0 | 1 |
| 333 | 1 | 0 | 3 | 2 | 1 | 0 |
| 144 | 4 | 3 | 1 | 2 | 0 | 0 |
| 284 | 0 | 0 | 0 | 2 | 0 | 0 |
| 36 | 0 | 0 | 0 | 0 | 0 | 0 |
| 47 | 0 | 0 | 1 | 0 | 0 | 0 |
| 20 | 0 | 0 | 0 | 0 | 0 | 0 |
| 56 | 0 | 0 | 0 | 0 | 0 | 0 |
| 115 | 0 | 0 | 0 | 0 | 0 | 0 |
| 1542 | 61 | | | | | |
| 1141 | 6 | 10 | 7 | 1 | 3 | 6 |
| 746 | 1 | 8 | 10 | 2 | 1 | 0 |
| 572 | 3 | 6 | 5 | 2 | 1 | 3 |
| 487 | 5 | 12 | 12 | 2 | 4 | 4 |
| 608 | 4 | 4 | 3 | 3 | 0 | 4 |
| 456 | 2 | 2 | 6 | 0 | 2 | 1 |
| 406 | 9 | 9 | 8 | 1 | 3 | 2 |
| 662 | 2 | 8 | 8 | 1 | 2 | 3 |
| 445 | 2 | 1 | 6 | 0 | 2 | 1 |
| 548 | 4 | 3 | 4 | 1 | 2 | 2 |
| 613 | 1 | 4 | 3 | 0 | 1 | 1 |
| 542 | 5 | 5 | 6 | 8 | 1 | 1 |
| 7226 | 360 | | | | | |
| 845 | 2 | 3 | 6 | 2 | 2 | 1 |
| 721 | 5 | 4 | 2 | 2 | 2 | 2 |
| 605 | 1 | 2 | 4 | 0 | 1 | 2 |
| 591 | 1 | 4 | 4 | 1 | 1 | 1 |
| 562 | 5 | 2 | 5 | 1 | 2 | 4 |
| 755 | 1 | 0 | 0 | 0 | 0 | 0 |
| 663 | 2 | 1 | 2 | 0 | 2 | 0 |
| 645 | 1 | 3 | 2 | 5 | 0 | 2 |
| 745 | 2 | 1 | 4 | 1 | 0 | 2 |
| 616 | 1 | 1 | 5 | 0 | 1 | 2 |
| 490 | 3 | 3 | 1 | 0 | 0 | 0 |
| 508 | 0 | 3 | 1 | 1 | 1 | 2 |
| 7746 | 197 | | | | | |

| | | | | | | |
|---|---|---|---|---|---|---|
| 1000 | 2 | 2 | 4 | 2 | 1 | 1 |
| 751 | 1 | 3 | 4 | 0 | 0 | 0 |
| 651 | 2 | 8 | 5 | 2 | 0 | 0 |
| 724 | 0 | 3 | 2 | 0 | 1 | 1 |
| 619 | 8 | 0 | 0 | 0 | 2 | 2 |
| 494 | 1 | 2 | 0 | 5 | 0 | 2 |
| 529 | 5 | 0 | 2 | 3 | 0 | 0 |
| 621 | 7 | 11 | 0 | 1 | 1 | 1 |
| 577 | 1 | 0 | 1 | 0 | 0 | 2 |
| 611 | 2 | 3 | 1 | 2 | 0 | 2 |
| 562 | 0 | 1 | 3 | 0 | 0 | 1 |
| 490 | 3 | 3 | 1 | 2 | 0 | 1 |
| 7629 | 156 | | | | | |
| 656 | 10 | 1 | 12 | 5 | 1 | 7 |
| 501 | 4 | 2 | 11 | 0 | 5 | 1 |
| 488 | 2 | 4 | 10 | 0 | 2 | 3 |
| 408 | 2 | 4 | 4 | 3 | 2 | 2 |
| 341 | 2 | 1 | 4 | 1 | 3 | 1 |
| 286 | 4 | 2 | 3 | 6 | 1 | 2 |
| 284 | 0 | 1 | 1 | 1 | 1 | 2 |
| 247 | 0 | 0 | 2 | 0 | 0 | 2 |
| 262 | 0 | 1 | 4 | 2 | 1 | 0 |
| 333 | 1 | 2 | 5 | 3 | 1 | 2 |
| 178 | 0 | 0 | 0 | 0 | 0 | 0 |
| 176 | 0 | 0 | 0 | 0 | 0 | 0 |
| 4160 | 277 | | | | | |
| 501 | 2 | 1 | 5 | 0 | 1 | 4 |
| 389 | 4 | 3 | 8 | 2 | 0 | 0 |
| 362 | 4 | 1 | 4 | 0 | 1 | 3 |
| 388 | 2 | 7 | 10 | 0 | 0 | 1 |
| 390 | 8 | 0 | 3 | 3 | 2 | 0 |
| 294 | 4 | 0 | 1 | 3 | 0 | 0 |
| 224 | 1 | 0 | 0 | 1 | 0 | 0 |
| 155 | 0 | 0 | 0 | 0 | 0 | 0 |
| 136 | 0 | 0 | 0 | 2 | 0 | 0 |
| 156 | 1 | 2 | 1 | 0 | 0 | 1 |
| 0 | 0 | 0 | 0 | 0 | 0 | 0 |
| 12 | 0 | 0 | 0 | 0 | 0 | 0 |
| 3007 | 169 | | | | | |
| 438 | 7 | 4 | 6 | 2 | 0 | 1 |
| 349 | 6 | 0 | 2 | 2 | 2 | 1 |

| | | | | | | |
|---|---|---|---|---|---|---|
| 283 | 1 | 2 | 4 | 0 | 0 | 2 |
| 157 | 0 | 0 | 0 | 0 | 0 | 0 |
| 106 | 0 | 0 | 0 | 0 | 0 | 0 |
| 81 | 0 | 0 | 0 | 0 | 0 | 0 |
| 25 | 0 | 0 | 0 | 0 | 0 | 0 |
| 4 | 0 | 0 | 0 | 0 | 0 | 0 |
| 18 | 0 | 0 | 0 | 0 | 0 | 0 |
| 17 | 0 | 0 | 0 | 0 | 0 | 0 |
| 0 | 0 | 0 | 0 | 0 | 0 | 0 |
| 0 | 0 | 0 | 0 | 0 | 0 | 0 |
| 1478 | 53 | | | | | |
| 3886 | 213 | | | | | |
| 4654 | 228 | | | | | |
| 2510 | 173 | | | | | |
| 2675 | 157 | | | | | |

| poke 7 | poke 8 | poke 9 | rearing |
|---|---|---|---|
| 1 | 5 | 7 | 1 |
| 0 | 1 | 2 | 0 |
| 5 | 3 | 10 | 0 |
| 0 | 1 | 1 | 11 |
| 1 | 0 | 4 | 0 |
| 3 | 1 | 6 | 2 |
| 0 | 0 | 0 | 1 |
| 0 | 0 | 0 | 1 |
| 0 | 0 | 0 | 0 |
| 0 | 0 | 0 | 0 |
| 0 | 0 | 0 | 0 |
| 0 | 0 | 0 | 0 |
| | | | 16 |
| 6 | 2 | 7 | 0 |
| 0 | 1 | 1 | 1 |
| 3 | 1 | 0 | 0 |
| 1 | 0 | 1 | 0 |
| 0 | 0 | 0 | 0 |
| 1 | 0 | 0 | 4 |
| 0 | 0 | 0 | 0 |
| 0 | 0 | 0 | 0 |
| 0 | 0 | 0 | 0 |
| 0 | 0 | 0 | 0 |
| 0 | 0 | 0 | 0 |
| 0 | 0 | 0 | 0 |
| | | | 5 |
| 3 | 0 | 1 | 0 |
| 0 | 0 | 1 | 0 |
| 2 | 0 | 1 | 0 |
| 0 | 0 | 0 | 0 |
| 0 | 0 | 0 | 0 |
| 0 | 0 | 0 | 0 |
| 0 | 0 | 0 | 0 |
| 0 | 0 | 0 | 0 |
| 0 | 0 | 0 | 0 |
| 0 | 0 | 0 | 0 |
| 0 | 0 | 0 | 0 |
| 0 | 0 | 1 | 9 |
| | | | 9 |
| 4 | 4 | 10 | 9 |

| | | | |
|---|---|---|---|
| 4 | 1 | 4 | 14 |
| 0 | 1 | 0 | 28 |
| 1 | 2 | 2 | 4 |
| 0 | 2 | 2 | 11 |
| 0 | 0 | 0 | 5 |
| 3 | 0 | 4 | 9 |
| 0 | 0 | 0 | 0 |
| 0 | 0 | 0 | 21 |
| 0 | 0 | 0 | 2 |
| 0 | 0 | 0 | 2 |
| 0 | 0 | 0 | 0 |
| | | | 105 |
| 4 | 0 | 2 | 14 |
| 2 | 3 | 3 | 16 |
| 1 | 2 | 0 | 12 |
| 1 | 1 | 2 | 11 |
| 1 | 0 | 0 | 3 |
| 3 | 0 | 1 | 11 |
| 0 | 0 | 5 | 5 |
| 0 | 0 | 5 | 6 |
| 0 | 0 | 1 | 7 |
| 0 | 0 | 1 | 2 |
| 0 | 0 | 0 | 3 |
| 0 | 0 | 0 | 0 |
| | | | 90 |
| 0 | 0 | 0 | 9 |
| 3 | 0 | 0 | 7 |
| 2 | 2 | 0 | 4 |
| 2 | 0 | 1 | 0 |
| 0 | 0 | 0 | 0 |
| 0 | 0 | 0 | 0 |
| 0 | 0 | 0 | 1 |
| 0 | 0 | 0 | 0 |
| 0 | 0 | 0 | 0 |
| 0 | 0 | 0 | 2 |
| 0 | 0 | 0 | 0 |
| 2 | 0 | 0 | 4 |
| | | | 27 |
| 2 | 2 | 5 | 3 |
| 0 | 0 | 0 | 7 |
| 0 | 0 | 2 | 1 |

| | | | |
|---|---|---|---|
| 0 | 0 | 0 | 0 |
| 3 | 0 | 0 | 1 |
| 0 | 0 | 0 | 0 |
| 0 | 0 | 0 | 0 |
| 0 | 0 | 0 | 0 |
| 0 | 0 | 0 | 0 |
| 2 | 0 | 0 | 0 |
| 0 | 0 | 0 | 0 |
| 0 | 0 | 0 | 0 |
| | | | 12 |
| 3 | 1 | 2 | 2 |
| 3 | 1 | 2 | 4 |
| 1 | 1 | 1 | 3 |
| 0 | 0 | 1 | 15 |
| 5 | 1 | 2 | 2 |
| 2 | 0 | 1 | 7 |
| 2 | 0 | 0 | 0 |
| 1 | 0 | 0 | 0 |
| 0 | 0 | 0 | 0 |
| 1 | 0 | 0 | 1 |
| 0 | 0 | 0 | 13 |
| 0 | 0 | 2 | 1 |
| | | | 48 |
| 3 | 9 | 11 | 1 |
| 6 | 4 | 15 | 6 |
| 8 | 4 | 5 | 24 |
| 2 | 1 | 7 | 29 |
| 4 | 3 | 8 | 13 |
| 7 | 4 | 15 | 15 |
| 3 | 4 | 6 | 12 |
| 1 | 2 | 1 | 49 |
| 1 | 0 | 16 | 30 |
| 0 | 0 | 3 | 10 |
| 0 | 1 | 1 | 9 |
| 0 | 0 | 1 | 4 |
| | | | 202 |
| 6 | 1 | 5 | 20 |
| 2 | 3 | 5 | 14 |
| 3 | 1 | 5 | 20 |
| 5 | 2 | 2 | 10 |
| 1 | 1 | 3 | 14 |

| | | | |
|---|---|---|---|
| 7 | 0 | 0 | 6 |
| 1 | 2 | 0 | 9 |
| 0 | 1 | 0 | 2 |
| 1 | 0 | 0 | 1 |
| 2 | 1 | 4 | 5 |
| 0 | 1 | 1 | 3 |
| 0 | 0 | 0 | 0 |
| | | | 104 |
| 4 | 0 | 1 | 12 |
| 11 | 2 | 4 | 4 |
| 2 | 1 | 0 | 1 |
| 0 | 1 | 1 | 8 |
| 2 | 2 | 10 | 10 |
| 0 | 0 | 5 | 3 |
| 3 | 3 | 3 | 2 |
| 1 | 2 | 2 | 0 |
| 1 | 0 | 0 | 1 |
| 0 | 0 | 0 | 4 |
| 0 | 0 | 0 | 1 |
| 0 | 0 | 0 | 0 |
| | | | 46 |
| 3 | 3 | 7 | 1 |
| 6 | 2 | 8 | 6 |
| 2 | 2 | 3 | 28 |
| 1 | 2 | 3 | 16 |
| 6 | 2 | 6 | 11 |
| 0 | 0 | 5 | 9 |
| 2 | 2 | 2 | 5 |
| 2 | 2 | 5 | 8 |
| 0 | 1 | 8 | 4 |
| 1 | 0 | 0 | 4 |
| 1 | 0 | 0 | 6 |
| 0 | 0 | 0 | 3 |
| | | | 101 |
| 3 | 2 | 6 | 13 |
| 2 | 1 | 1 | 15 |
| 4 | 3 | 5 | 22 |
| 2 | 0 | 3 | 12 |
| 4 | 1 | 1 | 8 |
| 4 | 1 | 5 | 11 |
| 0 | 2 | 0 | 31 |

| | | | |
|---|---|---|---|
| 1 | 1 | 3 | 6 |
| 1 | 0 | 2 | 11 |
| 1 | 13 | 3 | 10 |
| 1 | 0 | 0 | 15 |
| 0 | 0 | 0 | 5 |
| | | | 159 |
| 2 | 4 | 7 | 7 |
| 3 | 3 | 6 | 5 |
| 1 | 3 | 1 | 23 |
| 3 | 2 | 4 | 16 |
| 0 | 1 | 2 | 19 |
| 3 | 3 | 1 | 8 |
| 3 | 2 | 2 | 14 |
| 2 | 2 | 6 | 22 |
| 0 | 0 | 0 | 7 |
| 3 | 1 | 1 | 14 |
| 0 | 0 | 0 | 10 |
| 0 | 0 | 0 | 0 |
| | | | 145 |
| 4 | 5 | 4 | 18 |
| 3 | 1 | 1 | 7 |
| 3 | 0 | 2 | 7 |
| 2 | 1 | 0 | 10 |
| 1 | 2 | 8 | 10 |
| 1 | 1 | 4 | 5 |
| 0 | 0 | 1 | 13 |
| 0 | 0 | 4 | 11 |
| 4 | 1 | 1 | 9 |
| 0 | 1 | 1 | 4 |
| 1 | 0 | 1 | 2 |
| 0 | 2 | 0 | 0 |
| | | | 96 |
| 0 | 0 | 3 | 4 |
| 11 | 0 | 4 | 2 |
| 1 | 1 | 5 | 6 |
| 1 | 2 | 7 | 2 |
| 2 | 1 | 1 | 8 |
| 3 | 3 | 4 | 5 |
| 1 | 1 | 4 | 5 |
| 0 | 3 | 0 | 2 |
| 0 | 0 | 2 | 2 |

| | | | |
|---|---|---|---|
| 0 | 0 | 3 | 5 |
| 0 | 0 | 0 | 0 |
| 0 | 0 | 0 | 1 |
| | | | 42 |
| 3 | 1 | 2 | 5 |
| 2 | 1 | 4 | 9 |
| 3 | 0 | 2 | 7 |
| 6 | 0 | 1 | 5 |
| 0 | 2 | 1 | 6 |
| 6 | 0 | 0 | 0 |
| 1 | 0 | 0 | 0 |
| 0 | 0 | 0 | 0 |
| 0 | 0 | 0 | 3 |
| 5 | 1 | 1 | 0 |
| 3 | 0 | 0 | 2 |
| 0 | 0 | 0 | 0 |
| | | | 37 |
| 2 | 0 | 0 | 3 |
| 0 | 1 | 1 | 9 |
| 0 | 0 | 1 | 2 |
| 3 | 2 | 2 | 7 |
| 5 | 0 | 3 | 4 |
| 0 | 0 | 0 | 3 |
| 1 | 0 | 1 | 1 |
| 0 | 0 | 0 | 5 |
| 0 | 1 | 0 | 1 |
| 4 | 0 | 0 | 0 |
| 2 | 0 | 0 | 1 |
| 0 | 0 | 1 | 6 |
| | | | 42 |
| 12 | 0 | 5 | 1 |
| 0 | 0 | 0 | 0 |
| 0 | 2 | 0 | 2 |
| 0 | 0 | 2 | 5 |
| 2 | 0 | 0 | 25 |
| 2 | 1 | 4 | 11 |
| 0 | 0 | 0 | 2 |
| 0 | 0 | 0 | 0 |
| 0 | 0 | 0 | 0 |
| 0 | 0 | 0 | 0 |
| 0 | 0 | 0 | 0 |

| | | | |
|---|---|---|---|
| 0 | 0 | 0 | 0 |
| | | | 46 |
| 4 | 1 | 0 | 4 |
| 1 | 0 | 4 | 9 |
| 0 | 1 | 2 | 9 |
| 1 | 0 | 0 | 2 |
| 0 | 0 | 0 | 0 |
| 0 | 0 | 0 | 0 |
| 0 | 0 | 0 | 0 |
| 0 | 0 | 0 | 0 |
| 0 | 0 | 0 | 0 |
| 0 | 0 | 0 | 0 |
| 0 | 0 | 0 | 0 |
| 0 | 0 | 0 | 0 |
| | | | 24 |
| 1 | 0 | 2 | 3 |
| 0 | 0 | 1 | 1 |
| 3 | 0 | 0 | 0 |
| 0 | 0 | 0 | 0 |
| 0 | 0 | 0 | 0 |
| 0 | 1 | 0 | 0 |
| 0 | 0 | 0 | 0 |
| 0 | 0 | 0 | 0 |
| 0 | 0 | 0 | 0 |
| 0 | 0 | 0 | 0 |
| 3 | 2 | 0 | 0 |
| 0 | 1 | 0 | 3 |
| | | | 7 |
| 3 | 0 | 5 | 0 |
| 1 | 3 | 8 | 1 |
| 4 | 0 | 2 | 4 |
| 3 | 0 | 3 | 4 |
| 4 | 0 | 1 | 14 |
| 5 | 4 | 1 | 4 |
| 0 | 1 | 0 | 10 |
| 0 | 0 | 0 | 1 |
| 0 | 0 | 0 | 7 |
| 0 | 0 | 0 | 3 |
| 0 | 0 | 1 | 2 |
| 0 | 0 | 0 | 0 |
| | | | 50 |

| | | | |
|---|---|---|---|
| 1 | 0 | 1 | 0 |
| 0 | 0 | 1 | 3 |
| 0 | 0 | 1 | 1 |
| 0 | 0 | 5 | 3 |
| 0 | 0 | 0 | 0 |
| 0 | 0 | 0 | 0 |
| 0 | 0 | 0 | 0 |
| 0 | 0 | 0 | 0 |
| 0 | 0 | 0 | 0 |
| 0 | 0 | 0 | 0 |
| 0 | 0 | 0 | 0 |
| 0 | 0 | 0 | 0 |
| | | | 7 |
| 2 | 1 | 1 | 0 |
| 0 | 3 | 1 | 9 |
| 2 | 0 | 0 | 5 |
| 0 | 0 | 0 | 3 |
| 0 | 0 | 0 | 0 |
| 0 | 0 | 0 | 0 |
| 1 | 0 | 0 | 0 |
| 0 | 0 | 0 | 3 |
| 0 | 0 | 0 | 0 |
| 0 | 0 | 0 | 0 |
| 0 | 0 | 1 | 3 |
| 0 | 2 | 1 | 4 |
| | | | 27 |
| 0 | 0 | 14 | 0 |
| 5 | 8 | 4 | 0 |
| 9 | 8 | 9 | 0 |
| 5 | 12 | 3 | 0 |
| 7 | 5 | 0 | 0 |
| 2 | 22 | 3 | 0 |
| 6 | 4 | 1 | 0 |
| 5 | 4 | 3 | 0 |
| 0 | 0 | 0 | 0 |
| 0 | 0 | 0 | 0 |
| 3 | 2 | 0 | 5 |
| 0 | 0 | 0 | 0 |
| | | | 5 |
| 0 | 0 | 2 | 0 |
| 0 | 2 | 6 | 0 |

| | | | |
|---|---|---|---|
| 0 | 1 | 1 | 0 |
| 0 | 0 | 4 | 0 |
| 0 | 0 | 0 | 0 |
| 0 | 0 | 0 | 0 |
| 0 | 0 | 0 | 0 |
| 0 | 0 | 0 | 0 |
| 0 | 0 | 0 | 0 |
| 0 | 0 | 0 | 0 |
| 0 | 0 | 0 | 0 |
| 0 | 0 | 0 | 0 |
| | | | 0 |
| 1 | 9 | 3 | 0 |
| 4 | 20 | 3 | 0 |
| 1 | 1 | 3 | 0 |
| 0 | 0 | 0 | 0 |
| 0 | 1 | 0 | 0 |
| 6 | 8 | 4 | 0 |
| 1 | 1 | 0 | 0 |
| 0 | 0 | 0 | 0 |
| 0 | 0 | 0 | 0 |
| 0 | 0 | 0 | 0 |
| 0 | 0 | 0 | 0 |
| 0 | 0 | 0 | 0 |
| | | | 0 |
| 4 | 2 | 5 | 24 |
| 0 | 0 | 0 | 9 |
| 6 | 1 | 4 | 25 |
| 2 | 11 | 5 | 5 |
| 1 | 1 | 0 | 22 |
| 4 | 1 | 4 | 13 |
| 2 | 0 | 1 | 27 |
| 2 | 1 | 0 | 10 |
| 1 | 0 | 0 | 4 |
| 1 | 0 | 0 | 3 |
| 0 | 0 | 2 | 5 |
| 0 | 0 | 0 | 1 |
| | | | 148 |
| 2 | 1 | 3 | 2 |
| 8 | 2 | 0 | 7 |
| 5 | 0 | 2 | 4 |
| 0 | 0 | 0 | 0 |

| | | | |
|---|---|---|---|
| 0 | 0 | 2 | 0 |
| 0 | 0 | 1 | 8 |
| 2 | 2 | 1 | 5 |
| 0 | 0 | 9 | 0 |
| 0 | 0 | 0 | 0 |
| 0 | 0 | 4 | 0 |
| 0 | 0 | 0 | 0 |
| 0 | 0 | 0 | 0 |
| | | | 26 |
| 2 | 0 | 1 | 12 |
| 6 | 2 | 2 | 16 |
| 6 | 1 | 2 | 14 |
| 1 | 2 | 1 | 7 |
| 0 | 1 | 0 | 7 |
| 1 | 0 | 0 | 9 |
| 1 | 0 | 1 | 13 |
| 1 | 1 | 0 | 7 |
| 3 | 2 | 2 | 17 |
| 1 | 0 | 1 | 8 |
| 1 | 0 | 5 | 9 |
| 1 | 0 | 0 | 8 |
| | | | 127 |
| 5 | 1 | 2 | 16 |
| 0 | 0 | 0 | 14 |
| 1 | 5 | 4 | 11 |
| 1 | 0 | 0 | 14 |
| 0 | 1 | 5 | 24 |
| 0 | 0 | 0 | 2 |
| 0 | 0 | 0 | 3 |
| 0 | 0 | 0 | 2 |
| 0 | 0 | 0 | 0 |
| 0 | 0 | 0 | 0 |
| 0 | 0 | 0 | 0 |
| 0 | 0 | 0 | 0 |
| | | | 86 |
| 2 | 0 | 2 | 11 |
| 3 | 3 | 4 | 10 |
| 0 | 0 | 0 | 5 |
| 3 | 0 | 3 | 2 |
| 0 | 0 | 2 | 0 |
| 4 | 3 | 1 | 2 |

| | | | |
|---|---|---|---|
| 0 | 1 | 2 | 1 |
| 3 | 0 | 0 | 6 |
| 0 | 0 | 0 | 1 |
| 0 | 0 | 0 | 1 |
| 0 | 0 | 0 | 6 |
| 0 | 0 | 0 | 8 |
| | | | 53 |
| 2 | 0 | 1 | 10 |
| 1 | 3 | 3 | 12 |
| 8 | 0 | 0 | 6 |
| 2 | 1 | 0 | 11 |
| 0 | 2 | 2 | 18 |
| 2 | 0 | 1 | 8 |
| 1 | 0 | 1 | 13 |
| 0 | 0 | 0 | 50 |
| 2 | 0 | 3 | 48 |
| 0 | 0 | 0 | 15 |
| 0 | 0 | 1 | 9 |
| 1 | 2 | 2 | 8 |
| | | | 208 |
| 3 | 1 | 3 | 7 |
| 2 | 1 | 4 | 14 |
| 0 | 2 | 12 | 2 |
| 4 | 0 | 0 | 6 |
| 1 | 0 | 5 | 5 |
| 6 | 2 | 4 | 7 |
| 2 | 3 | 6 | 7 |
| 3 | 2 | 0 | 11 |
| 1 | 4 | 7 | 5 |
| 0 | 0 | 2 | 7 |
| 2 | 0 | 0 | 12 |
| 2 | 1 | 2 | 4 |
| | | | 87 |
| 1 | 1 | 2 | 16 |
| 0 | 2 | 1 | 8 |
| 8 | 4 | 6 | 13 |
| 2 | 0 | 0 | 13 |
| 1 | 1 | 0 | 25 |
| 1 | 2 | 3 | 5 |
| 1 | 0 | 1 | 6 |
| 1 | 0 | 0 | 7 |

| | | | |
|---|---|---|---|
| 1 | 0 | 1 | 7 |
| 0 | 0 | 0 | 1 |
| 0 | 1 | 1 | 9 |
| 0 | 0 | 0 | 4 |
| | | | 114 |
| 0 | 1 | 1 | 17 |
| 2 | 3 | 1 | 5 |
| 5 | 1 | 2 | 5 |
| 0 | 0 | 0 | 7 |
| 1 | 0 | 1 | 1 |
| 0 | 0 | 0 | 8 |
| 0 | 0 | 3 | 19 |
| 1 | 0 | 1 | 9 |
| 1 | 0 | 1 | 1 |
| 2 | 1 | 0 | 0 |
| 0 | 0 | 1 | 12 |
| 0 | 0 | 1 | 2 |
| | | | 86 |
| 7 | 10 | 6 | 24 |
| 6 | 0 | 14 | 14 |
| 0 | 1 | 3 | 15 |
| 0 | 1 | 5 | 11 |
| 1 | 0 | 7 | 14 |
| 4 | 2 | 1 | 16 |
| 0 | 0 | 3 | 13 |
| 0 | 0 | 2 | 5 |
| 0 | 0 | 2 | 6 |
| 0 | 0 | 0 | 6 |
| 3 | 0 | 0 | 6 |
| 0 | 0 | 0 | 0 |
| | | | 130 |
| 3 | 0 | 7 | 16 |
| 0 | 4 | 2 | 17 |
| 0 | 1 | 0 | 10 |
| 0 | 4 | 7 | 14 |
| 0 | 0 | 0 | 6 |
| 5 | 5 | 6 | 7 |
| 0 | 0 | 0 | 12 |
| 0 | 1 | 0 | 1 |
| 0 | 0 | 0 | 1 |
| 0 | 0 | 0 | 0 |

| | | | |
|---|---|---|---|
| 0 | 0 | 1 | 5 |
| 0 | 0 | 0 | 0 |
| | | | 89 |
| 2 | 3 | 2 | 13 |
| 2 | 2 | 1 | 7 |
| 0 | 1 | 1 | 24 |
| 0 | 0 | 0 | 5 |
| 0 | 0 | 0 | 3 |
| 0 | 0 | 0 | 0 |
| 0 | 0 | 0 | 0 |
| 0 | 0 | 0 | 0 |
| 0 | 0 | 0 | 0 |
| 0 | 0 | 0 | 0 |
| 0 | 0 | 0 | 1 |
| 0 | 0 | 0 | 0 |
| | | | 53 |
| 2 | 4 | 8 | 1 |
| 6 | 0 | 2 | 6 |
| 4 | 0 | 0 | 2 |
| 2 | 4 | 3 | 8 |
| 0 | 0 | 0 | 16 |
| 0 | 0 | 0 | 1 |
| 3 | 3 | 5 | 12 |
| 5 | 0 | 0 | 12 |
| 1 | 5 | 0 | 5 |
| 3 | 2 | 0 | 11 |
| 1 | 0 | 1 | 6 |
| 2 | 0 | 0 | 10 |
| | | | 90 |
| 3 | 2 | 0 | 15 |
| 1 | 1 | 2 | 20 |
| 3 | 2 | 2 | 17 |
| 2 | 0 | 0 | 23 |
| 0 | 3 | 3 | 16 |
| 4 | 0 | 1 | 12 |
| 0 | 0 | 0 | 51 |
| 0 | 0 | 2 | 20 |
| 0 | 0 | 1 | 28 |
| 3 | 2 | 0 | 20 |
| 3 | 2 | 0 | 16 |
| 1 | 1 | 1 | 13 |

| | | | |
|---|---|---|---|
| | | | 251 |
| 6 | 2 | 4 | 26 |
| 4 | 2 | 4 | 14 |
| 1 | 0 | 2 | 17 |
| 0 | 0 | 0 | 24 |
| 12 | 1 | 5 | 26 |
| 4 | 0 | 0 | 15 |
| 1 | 0 | 2 | 9 |
| 5 | 2 | 3 | 9 |
| 0 | 0 | 0 | 9 |
| 0 | 0 | 1 | 9 |
| 3 | 0 | 0 | 24 |
| 0 | 1 | 0 | 9 |
| | | | 191 |
| 6 | 2 | 6 | 0 |
| 5 | 0 | 3 | 1 |
| 7 | 1 | 4 | 4 |
| 3 | 1 | 3 | 2 |
| 2 | 0 | 4 | 1 |
| 0 | 0 | 0 | 1 |
| 0 | 1 | 0 | 0 |
| 0 | 0 | 0 | 0 |
| 0 | 0 | 0 | 0 |
| 0 | 0 | 0 | 0 |
| 3 | 0 | 1 | 5 |
| 0 | 0 | 0 | 0 |
| | | | 14 |
| 4 | 1 | 4 | 3 |
| 1 | 4 | 3 | 8 |
| 1 | 4 | 3 | 17 |
| 3 | 4 | 8 | 1 |
| 1 | 1 | 1 | 9 |
| 4 | 0 | 1 | 8 |
| 2 | 0 | 4 | 10 |
| 0 | 0 | 1 | 4 |
| 1 | 2 | 9 | 1 |
| 0 | 0 | 5 | 7 |
| 1 | 0 | 0 | 2 |
| 1 | 0 | 2 | 0 |
| | | | 70 |
| 1 | 1 | 3 | 1 |

| | | | |
|---|---|---|---|
| 2 | 1 | 5 | 0 |
| 2 | 0 | 0 | 7 |
| 0 | 0 | 2 | 4 |
| 0 | 0 | 3 | 0 |
| 5 | 1 | 2 | 1 |
| 0 | 0 | 2 | 0 |
| 2 | 0 | 0 | 6 |
| 0 | 0 | 0 | 0 |
| 0 | 0 | 0 | 0 |
| 0 | 2 | 0 | 0 |
| 0 | 0 | 0 | 0 |
| | | | 19 |
| 2 | 0 | 0 | 2 |
| 0 | 0 | 2 | 0 |
| 4 | 2 | 4 | 6 |
| 8 | 7 | 3 | 12 |
| 0 | 2 | 2 | 5 |
| 0 | 0 | 1 | 12 |
| 0 | 0 | 0 | 3 |
| 0 | 0 | 0 | 0 |
| 7 | 2 | 3 | 23 |
| 0 | 0 | 4 | 0 |
| 0 | 0 | 0 | 0 |
| 0 | 0 | 0 | 0 |
| | | | 63 |
| 0 | 26 | 0 | 0 |
| 0 | 18 | 0 | 0 |
| 0 | 23 | 0 | 0 |
| 0 | 10 | 0 | 0 |
| 0 | 16 | 0 | 0 |
| 0 | 16 | 0 | 0 |
| 0 | 16 | 0 | 0 |
| 0 | 16 | 0 | 0 |
| 0 | 28 | 0 | 0 |
| 0 | 13 | 0 | 0 |
| 0 | 13 | 0 | 0 |
| 0 | 11 | 0 | 0 |
| | | | 0 |
| 4 | 4 | 3 | 0 |
| 3 | 2 | 5 | 0 |
| 3 | 0 | 3 | 0 |

| | | | |
|---|---|---|---|
| 2 | 0 | 0 | 0 |
| 1 | 0 | 0 | 0 |
| 0 | 0 | 0 | 0 |
| 0 | 0 | 0 | 0 |
| 3 | 0 | 0 | 0 |
| 0 | 0 | 0 | 0 |
| 0 | 0 | 0 | 0 |
| 0 | 0 | 0 | 0 |
| 0 | 0 | 0 | 0 |
| | | | 0 |
| 6 | 1 | 2 | 2 |
| 1 | 3 | 3 | 2 |
| 4 | 3 | 0 | 8 |
| 2 | 4 | 6 | 3 |
| 3 | 2 | 2 | 5 |
| 2 | 1 | 0 | 10 |
| 8 | 4 | 12 | 6 |
| 12 | 5 | 3 | 2 |
| 0 | 0 | 2 | 10 |
| 0 | 0 | 0 | 0 |
| 0 | 0 | 0 | 0 |
| 0 | 0 | 0 | 0 |
| | | | 48 |
| 6 | 4 | 2 | 0 |
| 0 | 0 | 0 | 0 |
| 1 | 1 | 0 | 0 |
| 0 | 0 | 0 | 0 |
| 0 | 0 | 0 | 0 |
| 2 | 0 | 0 | 0 |
| 0 | 0 | 0 | 0 |
| 0 | 0 | 0 | 0 |
| 1 | 2 | 2 | 14 |
| 0 | 0 | 0 | 0 |
| 0 | 0 | 0 | 0 |
| 0 | 0 | 0 | 0 |
| | | | 14 |
| 0 | 3 | 2 | 1 |
| 10 | 3 | 5 | 2 |
| 2 | 0 | 1 | 12 |
| 0 | 2 | 0 | 1 |
| 0 | 0 | 0 | 3 |

| | | | |
|---|---|---|---|
| 0 | 0 | 0 | 0 |
| 0 | 0 | 0 | 3 |
| 3 | 1 | 1 | 0 |
| 3 | 1 | 2 | 1 |
| 0 | 0 | 0 | 0 |
| 1 | 0 | 0 | 1 |
| 0 | 0 | 0 | 0 |
| | | | 24 |
| 4 | 3 | 1 | 8 |
| 8 | 1 | 6 | 13 |
| 5 | 6 | 4 | 14 |
| 8 | 0 | 0 | 21 |
| 2 | 5 | 1 | 9 |
| 3 | 7 | 0 | 15 |
| 8 | 1 | 5 | 18 |
| 0 | 0 | 2 | 31 |
| 2 | 1 | 0 | 1 |
| 0 | 1 | 0 | 0 |
| 0 | 2 | 0 | 0 |
| 3 | 2 | 2 | 5 |
| | | | 135 |
| 7 | 7 | 5 | 0 |
| 2 | 0 | 3 | 5 |
| 6 | 1 | 2 | 13 |
| 5 | 0 | 1 | 0 |
| 2 | 0 | 0 | 0 |
| 0 | 0 | 0 | 1 |
| 0 | 0 | 0 | 1 |
| 0 | 0 | 0 | 0 |
| 0 | 0 | 0 | 0 |
| 0 | 0 | 0 | 0 |
| 0 | 0 | 0 | 0 |
| 0 | 0 | 0 | 0 |
| | | | 20 |
| 2 | 2 | 5 | 9 |
| 0 | 1 | 3 | 4 |
| 2 | 1 | 1 | 4 |
| 0 | 0 | 2 | 2 |
| 0 | 0 | 0 | 0 |
| 0 | 0 | 0 | 0 |
| 0 | 0 | 0 | 0 |

| | | | |
|---|---|---|---|
| 3 | 3 | 2 | 0 |
| 0 | 0 | 0 | 0 |
| 0 | 0 | 0 | 0 |
| 0 | 0 | 0 | 0 |
| 0 | 0 | 0 | 0 |
| | | | 19 |
| 2 | 6 | 7 | 1 |
| 0 | 0 | 6 | 0 |
| 5 | 6 | 2 | 7 |
| 1 | 0 | 5 | 1 |
| 0 | 0 | 0 | 1 |
| 0 | 0 | 0 | 0 |
| 1 | 0 | 0 | 4 |
| 0 | 0 | 0 | 0 |
| 5 | 1 | 0 | 1 |
| 2 | 4 | 5 | 17 |
| 1 | 2 | 0 | 7 |
| 1 | 0 | 0 | 4 |
| | | | 43 |
| 6 | 5 | 5 | 1 |
| 3 | 5 | 4 | 3 |
| 1 | 2 | 0 | 1 |
| 4 | 1 | 1 | 1 |
| 1 | 0 | 2 | 3 |
| 4 | 1 | 1 | 1 |
| 1 | 1 | 0 | 0 |
| 0 | 0 | 0 | 0 |
| 0 | 0 | 0 | 0 |
| 3 | 0 | 0 | 1 |
| 0 | 0 | 0 | 0 |
| 0 | 0 | 0 | 0 |
| | | | 11 |
| 0 | 1 | 1 | 0 |
| 1 | 0 | 0 | 0 |
| 0 | 0 | 4 | 0 |
| 4 | 4 | 2 | 2 |
| 1 | 0 | 0 | 0 |
| 2 | 0 | 1 | 0 |
| 0 | 0 | 5 | 1 |
| 0 | 0 | 0 | 0 |
| 0 | 0 | 0 | 0 |

| | | | |
|---|---|---|---|
| 0 | 0 | 1 | 0 |
| 0 | 0 | 0 | 1 |
| 0 | 0 | 0 | 0 |
| | | | 4 |
| 7 | 9 | 16 | 5 |
| 4 | 14 | 2 | 36 |
| 0 | 4 | 7 | 26 |
| 4 | 7 | 5 | 38 |
| 2 | 5 | 3 | 16 |
| 4 | 5 | 3 | 21 |
| 1 | 0 | 1 | 12 |
| 1 | 5 | 3 | 9 |
| 2 | 2 | 2 | 5 |
| 2 | 1 | 1 | 8 |
| 0 | 0 | 0 | 2 |
| 0 | 0 | 0 | 0 |
| | | | 178 |
| 2 | 2 | 2 | 4 |
| 6 | 6 | 6 | 17 |
| 3 | 6 | 9 | 16 |
| 2 | 5 | 2 | 19 |
| 7 | 6 | 6 | 14 |
| 0 | 1 | 6 | 24 |
| 2 | 3 | 4 | 33 |
| 4 | 2 | 4 | 14 |
| 3 | 1 | 0 | 5 |
| 3 | 8 | 0 | 39 |
| 5 | 3 | 1 | 23 |
| 3 | 5 | 2 | 6 |
| | | | 214 |
| 4 | 8 | 4 | 28 |
| 3 | 1 | 2 | 34 |
| 4 | 7 | 2 | 26 |
| 3 | 5 | 2 | 53 |
| 1 | 3 | 3 | 44 |
| 1 | 2 | 3 | 48 |
| 6 | 4 | 6 | 23 |
| 3 | 3 | 4 | 20 |
| 4 | 2 | 10 | 32 |
| 5 | 2 | 2 | 17 |
| 2 | 5 | 5 | 7 |

| | | | |
|---|---|---|---|
| 4 | 1 | 0 | 26 |
| | | | 358 |
| 5 | 1 | 2 | 0 |
| 4 | 2 | 7 | 0 |
| 9 | 5 | 5 | 0 |
| 3 | 3 | 2 | 0 |
| 7 | 5 | 7 | 0 |
| 2 | 0 | 3 | 0 |
| 5 | 2 | 6 | 0 |
| 0 | 0 | 0 | 0 |
| 0 | 0 | 0 | 0 |
| 0 | 0 | 0 | 0 |
| 0 | 0 | 2 | 0 |
| 0 | 0 | 3 | 0 |
| | | | 0 |
| 5 | 0 | 4 | 0 |
| 3 | 2 | 4 | 0 |
| 0 | 0 | 5 | 1 |
| 0 | 0 | 0 | 0 |
| 6 | 0 | 0 | 0 |
| 0 | 0 | 0 | 0 |
| 0 | 0 | 0 | 0 |
| 0 | 0 | 0 | 0 |
| 0 | 0 | 0 | 0 |
| 0 | 0 | 0 | 0 |
| 0 | 0 | 1 | 0 |
| 0 | 0 | 0 | 0 |
| | | | 1 |
| 1 | 1 | 10 | 5 |
| 0 | 0 | 0 | 0 |
| 3 | 8 | 1 | 0 |
| 0 | 0 | 1 | 1 |
| 0 | 0 | 0 | 0 |
| 0 | 0 | 0 | 0 |
| 0 | 0 | 0 | 0 |
| 4 | 0 | 1 | 0 |
| 0 | 0 | 0 | 0 |
| 0 | 0 | 0 | 0 |
| 0 | 0 | 0 | 0 |
| 0 | 0 | 0 | 0 |
| | | | 6 |

| | | | |
|---|---|---|---|
| 2 | 0 | 6 | 0 |
| 0 | 1 | 1 | 0 |
| 5 | 2 | 4 | 0 |
| 4 | 1 | 2 | 0 |
| 0 | 0 | 2 | 0 |
| 0 | 0 | 0 | 0 |
| 0 | 0 | 0 | 0 |
| 0 | 0 | 0 | 0 |
| 0 | 0 | 0 | 0 |
| 0 | 0 | 0 | 0 |
| 0 | 0 | 0 | 0 |
| 0 | 0 | 0 | 0 |
| | | | 0 |
| 2 | 2 | 1 | 0 |
| 1 | 0 | 5 | 0 |
| 0 | 0 | 2 | 0 |
| 0 | 0 | 0 | 0 |
| 0 | 0 | 0 | 0 |
| 0 | 0 | 0 | 0 |
| 0 | 0 | 0 | 0 |
| 0 | 0 | 0 | 0 |
| 1 | 0 | 0 | 0 |
| 0 | 0 | 0 | 0 |
| 0 | 0 | 0 | 0 |
| 0 | 0 | 0 | 0 |
| | | | 0 |
| 2 | 1 | 3 | 1 |
| 0 | 0 | 0 | 3 |
| 1 | 0 | 0 | 0 |
| 0 | 0 | 0 | 0 |
| 0 | 0 | 0 | 0 |
| 0 | 0 | 0 | 0 |
| 2 | 3 | 1 | 0 |
| 1 | 0 | 2 | 0 |
| 0 | 0 | 0 | 0 |
| 0 | 0 | 0 | 0 |
| 0 | 0 | 0 | 0 |
| 0 | 0 | 0 | 0 |
| | | | 4 |
| 15 | 2 | 13 | 13 |
| 2 | 3 | 4 | 23 |

| | | | |
|---|---|---|---|
| 4 | 1 | 3 | 11 |
| 8 | 3 | 2 | 9 |
| 4 | 0 | 0 | 44 |
| 5 | 2 | 16 | 13 |
| 2 | 2 | 4 | 26 |
| 1 | 3 | 3 | 11 |
| 2 | 1 | 9 | 60 |
| 0 | 1 | 2 | 30 |
| 4 | 0 | 0 | 22 |
| 0 | 0 | 1 | 15 |
| | | | 277 |
| 6 | 3 | 3 | 22 |
| 1 | 4 | 2 | 26 |
| 6 | 1 | 4 | 32 |
| 2 | 1 | 1 | 20 |
| 4 | 1 | 4 | 12 |
| 1 | 0 | 3 | 24 |
| 2 | 1 | 1 | 18 |
| 1 | 0 | 3 | 19 |
| 3 | 4 | 0 | 14 |
| 0 | 0 | 1 | 2 |
| 0 | 0 | 0 | 0 |
| 0 | 0 | 0 | 3 |
| | | | 192 |
| 1 | 0 | 4 | 24 |
| 0 | 0 | 6 | 21 |
| 1 | 1 | 3 | 15 |
| 0 | 0 | 8 | 18 |
| 0 | 0 | 0 | 9 |
| 2 | 0 | 7 | 3 |
| 0 | 0 | 1 | 8 |
| 0 | 0 | 0 | 1 |
| 0 | 0 | 0 | 6 |
| 0 | 0 | 8 | 2 |
| 0 | 0 | 1 | 0 |
| 0 | 0 | 0 | 0 |
| | | | 107 |
| 8 | 6 | 7 | 0 |
| 3 | 5 | 7 | 0 |
| 3 | 1 | 1 | 0 |
| 1 | 0 | 0 | 0 |

| | | | |
|---|---|---|---|
| 1 | 3 | 7 | 0 |
| 0 | 1 | 0 | 0 |
| 0 | 0 | 0 | 0 |
| 0 | 0 | 0 | 0 |
| 0 | 0 | 0 | 0 |
| 0 | 0 | 0 | 0 |
| 0 | 0 | 0 | 0 |
| 0 | 0 | 0 | 0 |
| | | rearing result | 0 |
| 3 | 1 | 4 | 15 |
| 4 | 2 | 1 | 31 |
| 1 | 5 | 5 | 30 |
| 6 | 1 | 7 | 37 |
| 2 | 1 | 1 | 16 |
| 0 | 0 | 2 | 11 |
| 1 | 0 | 1 | 19 |
| 3 | 2 | 5 | 3 |
| 0 | 0 | 0 | 0 |
| 0 | 0 | 0 | 12 |
| 0 | 0 | 0 | 0 |
| 0 | 0 | 0 | 1 |
| | | | 175 |
| 4 | 2 | 3 | 7 |
| 3 | 0 | 1 | 10 |
| 1 | 1 | 1 | 3 |
| 1 | 0 | 4 | 0 |
| 1 | 0 | 0 | 5 |
| 0 | 0 | 0 | 0 |
| 0 | 0 | 0 | 0 |
| 0 | 0 | 2 | 1 |
| 0 | 0 | 0 | 0 |
| 0 | 0 | 0 | 0 |
| 0 | 0 | 0 | 0 |
| 0 | 0 | 0 | 0 |
| | | | 26 |
| 3 | 4 | 7 | 0 |
| 7 | 1 | 2 | 0 |
| 9 | 3 | 4 | 0 |
| 8 | 3 | 6 | 0 |
| 1 | 1 | 0 | 3 |
| 2 | 0 | 1 | 0 |

| | | | |
|---|---|---|---|
| 0 | 0 | 0 | 0 |
| 0 | 0 | 0 | 0 |
| 0 | 0 | 0 | 0 |
| 0 | 0 | 0 | 0 |
| 0 | 0 | 0 | 0 |
| 0 | 0 | 0 | 0 |
| | | rearing result | 3 |
| 7 | 3 | 7 | 4 |
| 5 | 2 | 1 | 6 |
| 2 | 2 | 1 | 5 |
| 1 | 5 | 1 | 18 |
| 2 | 0 | 1 | 13 |
| 2 | 3 | 3 | 9 |
| 0 | 6 | 2 | 19 |
| 2 | 0 | 1 | 8 |
| 0 | 0 | 0 | 8 |
| 0 | 0 | 0 | 3 |
| 2 | 2 | 5 | 1 |
| 0 | 0 | 0 | 2 |
| | | | 96 |
| 3 | 10 | 2 | 5 |
| 3 | 5 | 4 | 13 |
| 4 | 5 | 6 | 8 |
| 3 | 4 | 4 | 8 |
| 1 | 5 | 0 | 6 |
| 1 | 1 | 2 | 5 |
| 3 | 3 | 0 | 8 |
| 0 | 3 | 0 | 13 |
| 4 | 0 | 0 | 0 |
| 1 | 6 | 1 | 1 |
| 0 | 0 | 0 | 3 |
| 0 | 0 | 0 | 1 |
| | | | 71 |
| 0 | 0 | 0 | 0 |
| 0 | 0 | 0 | 0 |
| 0 | 0 | 0 | 0 |
| 0 | 0 | 0 | 0 |
| 0 | 0 | 0 | 0 |
| 5 | 0 | 0 | 0 |
| 0 | 0 | 0 | 0 |
| 0 | 0 | 5 | 0 |

| | | | |
|---|---|---|---|
| 0 | 0 | 0 | 0 |
| 0 | 0 | 0 | 0 |
| 0 | 0 | 0 | 0 |
| 0 | 0 | 0 | 0 |
| | rearing result | | 0 |
| 3 | 6 | 6 | 0 |
| 2 | 3 | 2 | 0 |
| 3 | 1 | 3 | 0 |
| 1 | 0 | 0 | 0 |
| 3 | 5 | 5 | 0 |
| 0 | 0 | 1 | 2 |
| 0 | 0 | 0 | 0 |
| 0 | 0 | 0 | 0 |
| 0 | 0 | 2 | 0 |
| 0 | 0 | 0 | 0 |
| 0 | 0 | 0 | 0 |
| 0 | 0 | 0 | 0 |
| | | | 2 |
| 1 | 1 | 3 | 0 |
| 3 | 0 | 2 | 0 |
| 4 | 0 | 0 | 0 |
| 1 | 0 | 4 | 0 |
| 0 | 1 | 0 | 0 |
| 0 | 0 | 0 | 0 |
| 0 | 0 | 0 | 0 |
| 0 | 0 | 0 | 0 |
| 0 | 0 | 0 | 0 |
| 0 | 0 | 0 | 0 |
| 0 | 0 | 0 | 0 |
| 0 | 0 | 1 | 0 |
| | | | 0 |
| 7 | 7 | 11 | 0 |
| 1 | 1 | 6 | 0 |
| 4 | 0 | 2 | 0 |
| 6 | 2 | 3 | 0 |
| 0 | 1 | 0 | 0 |
| 1 | 3 | 0 | 0 |
| 1 | 0 | 0 | 0 |
| 1 | 1 | 7 | 0 |
| 0 | 0 | 1 | 0 |
| 0 | 0 | 0 | 0 |

| | | | |
|---|---|---|---|
| 0 | 0 | 0 | 0 |
| 0 | 0 | 0 | 0 |
| | | | 0 |
| 0 | 3 | 0 | 0 |
| 2 | 0 | 6 | 0 |
| 3 | 1 | 1 | 0 |
| 0 | 0 | 1 | 0 |
| 5 | 1 | 5 | 0 |
| 0 | 0 | 3 | 0 |
| 2 | 4 | 3 | 0 |
| 0 | 1 | 2 | 0 |
| 4 | 0 | 2 | 0 |
| 0 | 0 | 0 | 0 |
| 0 | 0 | 0 | 0 |
| 0 | 0 | 0 | 0 |
| | | | 0 |
| 2 | 2 | 1 | 0 |
| 2 | 0 | 1 | 0 |
| 3 | 4 | 1 | 0 |
| 0 | 0 | 0 | 0 |
| 3 | 3 | 1 | 0 |
| 4 | 0 | 1 | 0 |
| 0 | 0 | 0 | 0 |
| 0 | 0 | 0 | 0 |
| 0 | 0 | 0 | 0 |
| 0 | 0 | 0 | 0 |
| 0 | 0 | 0 | 0 |
| 0 | 7 | 0 | 0 |
| | | | 0 |
| 6 | 2 | 6 | 1 |
| 3 | 6 | 3 | 0 |
| 1 | 0 | 8 | 0 |
| 8 | 3 | 2 | 0 |
| 0 | 0 | 1 | 1 |
| 0 | 2 | 0 | 0 |
| 0 | 0 | 0 | 0 |
| 0 | 0 | 0 | 0 |
| 0 | 0 | 0 | 0 |
| 0 | 0 | 0 | 0 |
| 0 | 0 | 0 | 0 |
| 0 | 0 | 0 | 0 |

| | | | |
|---|---|---|---|
| | | | 2 |
| 6 | 4 | 7 | 0 |
| 6 | 0 | 2 | 0 |
| 1 | 0 | 3 | 0 |
| 2 | 0 | 0 | 0 |
| 0 | 0 | 0 | 0 |
| 0 | 0 | 0 | 0 |
| 0 | 0 | 0 | 0 |
| 0 | 0 | 1 | 0 |
| 3 | 0 | 0 | 0 |
| 0 | 0 | 0 | 0 |
| 0 | 0 | 0 | 0 |
| 0 | 0 | 0 | 0 |
| | | | 0 |
| 3 | 0 | 2 | 0 |
| 2 | 2 | 0 | 6 |
| 1 | 0 | 2 | 0 |
| 2 | 0 | 0 | 0 |
| 1 | 0 | 0 | 0 |
| 0 | 0 | 1 | 0 |
| 1 | 1 | 0 | 0 |
| 0 | 0 | 0 | 0 |
| 1 | 0 | 0 | 0 |
| 0 | 0 | 0 | 0 |
| 0 | 0 | 0 | 0 |
| 0 | 0 | 0 | 0 |
| | | | 6 |
| 7 | 7 | 10 | 2 |
| 5 | 1 | 6 | 0 |
| 2 | 0 | 4 | 16 |
| 5 | 6 | 12 | 2 |
| 5 | 2 | 1 | 11 |
| 0 | 0 | 4 | 9 |
| 0 | 0 | 0 | 1 |
| 1 | 0 | 3 | 12 |
| 2 | 0 | 0 | 3 |
| 0 | 3 | 0 | 5 |
| 0 | 0 | 0 | 1 |
| 0 | 0 | 0 | 3 |
| | | | 65 |
| 6 | 0 | 4 | 0 |

| | | | |
|---|---|---|---|
| 1 | 1 | 1 | 0 |
| 0 | 0 | 1 | 0 |
| 0 | 0 | 0 | 1 |
| 0 | 0 | 0 | 0 |
| 0 | 0 | 0 | 0 |
| 0 | 0 | 0 | 0 |
| 0 | 0 | 0 | 0 |
| 0 | 0 | 0 | 0 |
| 0 | 0 | 0 | 0 |
| 0 | 0 | 0 | 0 |
| 0 | 0 | 0 | 0 |
| | | | 1 |
| 2 | 0 | 3 | 2 |
| 4 | 0 | 1 | 3 |
| 1 | 1 | 4 | 0 |
| 0 | 0 | 0 | 0 |
| 0 | 0 | 0 | 0 |
| 0 | 0 | 1 | 0 |
| 0 | 0 | 0 | 0 |
| 0 | 0 | 0 | 0 |
| 0 | 0 | 0 | 0 |
| 0 | 0 | 0 | 0 |
| 0 | 0 | 0 | 0 |
| 0 | 0 | 0 | 0 |
| | | | 5 |
| 6 | 2 | 5 | 2 |
| 4 | 7 | 6 | 9 |
| 2 | 1 | 1 | 7 |
| 2 | 3 | 7 | 5 |
| 2 | 1 | 3 | 6 |
| 2 | 0 | 3 | 8 |
| 0 | 0 | 8 | 2 |
| 1 | 1 | 3 | 4 |
| 0 | 0 | 1 | 1 |
| 0 | 0 | 1 | 0 |
| 0 | 0 | 0 | 0 |
| 0 | 1 | 0 | 3 |
| | | | 47 |
| 1 | 3 | 1 | 2 |
| 0 | 2 | 3 | 3 |
| 1 | 0 | 1 | 4 |

| | | | |
|---|---|---|---|
| 0 | 0 | 0 | 0 |
| 0 | 0 | 0 | 0 |
| 0 | 0 | 1 | 2 |
| 0 | 1 | 0 | 0 |
| 0 | 0 | 0 | 0 |
| 0 | 0 | 0 | 0 |
| 0 | 0 | 0 | 0 |
| 0 | 0 | 0 | 0 |
| 0 | 0 | 0 | 0 |
| | | | 11 |
| 1 | 12 | 1 | 1 |
| 0 | 2 | 0 | 1 |
| 0 | 0 | 1 | 0 |
| 0 | 0 | 0 | 0 |
| 0 | 0 | 0 | 0 |
| 0 | 0 | 0 | 0 |
| 0 | 0 | 0 | 0 |
| 0 | 0 | 0 | 0 |
| 0 | 0 | 0 | 0 |
| 0 | 0 | 0 | 0 |
| 0 | 0 | 0 | 0 |
| 0 | 0 | 0 | 0 |
| | | | 2 |
| 3 | 2 | 9 | 0 |
| 1 | 0 | 3 | 0 |
| 0 | 8 | 9 | 0 |
| 2 | 5 | 3 | 0 |
| 3 | 0 | 2 | 1 |
| 9 | 1 | 5 | 0 |
| 4 | 0 | 3 | 0 |
| 0 | 6 | 3 | 0 |
| 0 | 0 | 5 | 0 |
| 0 | 0 | 0 | 0 |
| 0 | 0 | 0 | 0 |
| 0 | 0 | 0 | 0 |
| | | | 1 |
| 7 | 4 | 2 | 1 |
| 0 | 1 | 1 | 5 |
| 0 | 0 | 4 | 2 |
| 0 | 2 | 0 | 2 |
| 3 | 0 | 1 | 3 |

| | | | |
|---|---|---|---|
| 5 | 0 | 1 | 1 |
| 0 | 0 | 1 | 2 |
| 1 | 0 | 0 | 4 |
| 0 | 0 | 0 | 0 |
| 0 | 0 | 0 | 0 |
| 0 | 0 | 0 | 0 |
| 1 | 0 | 0 | 0 |
| | | | 20 |
| 3 | 0 | 4 | 2 |
| 0 | 0 | 0 | 0 |
| 0 | 0 | 0 | 0 |
| 1 | 0 | 0 | 0 |
| 0 | 0 | 0 | 0 |
| 0 | 0 | 0 | 0 |
| 2 | 0 | 0 | 0 |
| 0 | 0 | 0 | 1 |
| 0 | 0 | 0 | 0 |
| 0 | 0 | 0 | 0 |
| 0 | 0 | 0 | 0 |
| 0 | 0 | 0 | 0 |
| | | | 3 |
| 6 | 8 | 4 | 10 |
| 2 | 0 | 1 | 10 |
| 6 | 0 | 3 | 3 |
| 1 | 0 | 0 | 10 |
| 2 | 5 | 9 | 3 |
| 2 | 0 | 3 | 27 |
| 0 | 2 | 1 | 5 |
| 0 | 0 | 0 | 1 |
| 0 | 0 | 2 | 1 |
| 0 | 0 | 0 | 0 |
| 0 | 0 | 0 | 0 |
| 0 | 0 | 0 | 0 |
| | | | 70 |
| 4 | 0 | 3 | 8 |
| 2 | 1 | 0 | 3 |
| 2 | 0 | 1 | 0 |
| 1 | 2 | 0 | 1 |
| 0 | 0 | 0 | 0 |
| 1 | 0 | 0 | 0 |
| 0 | 0 | 1 | 0 |

| | | | |
|---|---|---|---|
| 0 | 0 | 0 | 0 |
| 0 | 0 | 0 | 0 |
| 0 | 0 | 0 | 0 |
| 0 | 0 | 0 | 0 |
| 0 | 0 | 0 | 0 |
| | | | 12 |
| 0 | 0 | 5 | 7 |
| 1 | 0 | 0 | 1 |
| 0 | 0 | 0 | 1 |
| 1 | 0 | 0 | 4 |
| 0 | 2 | 0 | 0 |
| 0 | 1 | 0 | 0 |
| 0 | 0 | 0 | 0 |
| 1 | 2 | 1 | 4 |
| 0 | 0 | 0 | 0 |
| 0 | 0 | 0 | 1 |
| 0 | 0 | 0 | 0 |
| 0 | 0 | 0 | 0 |
| | | | 18 |
| 5 | 13 | 11 | 13 |
| 1 | 3 | 2 | 22 |
| 2 | 6 | 4 | 10 |
| 1 | 0 | 4 | 11 |
| 6 | 2 | 0 | 8 |
| 2 | 0 | 2 | 7 |
| 1 | 2 | 3 | 6 |
| 0 | 3 | 0 | 6 |
| 0 | 0 | 0 | 7 |
| 0 | 0 | 0 | 0 |
| 0 | 0 | 0 | 1 |
| 0 | 2 | 0 | 9 |
| | | | 100 |
| 2 | 2 | 1 | 24 |
| 2 | 2 | 0 | 19 |
| 1 | 1 | 0 | 3 |
| 0 | 0 | 0 | 2 |
| 0 | 0 | 0 | 0 |
| 0 | 0 | 0 | 0 |
| 1 | 1 | 1 | 16 |
| 0 | 0 | 2 | 8 |
| 0 | 0 | 0 | 0 |

| | | | |
|---|---|---|---|
| 0 | 0 | 0 | 0 |
| 0 | 0 | 0 | 0 |
| 0 | 0 | 0 | 0 |
| | | | 72 |
| 8 | 5 | 6 | 2 |
| 0 | 1 | 8 | 4 |
| 2 | 5 | 8 | 9 |
| 2 | 0 | 1 | 12 |
| 1 | 4 | 6 | 13 |
| 3 | 4 | 5 | 11 |
| 3 | 7 | 2 | 11 |
| 2 | 6 | 7 | 13 |
| 1 | 3 | 4 | 11 |
| 5 | 2 | 2 | 4 |
| 0 | 0 | 0 | 9 |
| 0 | 0 | 0 | 1 |
| | | | 100 |
| 2 | 6 | 4 | 4 |
| 0 | 1 | 4 | 0 |
| 0 | 0 | 0 | 5 |
| 3 | 4 | 2 | 5 |
| 0 | 0 | 0 | 0 |
| 0 | 2 | 0 | 1 |
| 0 | 0 | 0 | 0 |
| 1 | 0 | 0 | 0 |
| 0 | 0 | 0 | 3 |
| 0 | 2 | 1 | 0 |
| 4 | 0 | 0 | 0 |
| 0 | 8 | 2 | 2 |
| | | | 20 |
| 4 | 1 | 5 | 28 |
| 3 | 3 | 4 | 37 |
| 6 | 1 | 4 | 28 |
| 3 | 2 | 4 | 25 |
| 6 | 0 | 2 | 36 |
| 2 | 0 | 4 | 36 |
| 5 | 0 | 4 | 22 |
| 5 | 0 | 2 | 8 |
| 0 | 0 | 0 | 5 |
| 2 | 0 | 3 | 20 |
| 0 | 0 | 0 | 2 |

| | | | |
|---|---|---|---|
| 0 | 0 | 0 | 0 |
| | | | 247 |
| 8 | 1 | 3 | 19 |
| 6 | 0 | 2 | 15 |
| 0 | 0 | 0 | 6 |
| 2 | 2 | 2 | 0 |
| 0 | 0 | 0 | 1 |
| 0 | 0 | 0 | 0 |
| 0 | 0 | 0 | 0 |
| 0 | 0 | 0 | 0 |
| 0 | 0 | 0 | 0 |
| 0 | 0 | 0 | 0 |
| 0 | 0 | 0 | 0 |
| 0 | 0 | 0 | 0 |
| | | | 41 |
| 1 | 3 | 0 | 5 |
| 0 | 0 | 0 | 0 |
| 0 | 0 | 0 | 2 |
| 0 | 0 | 0 | 0 |
| 0 | 0 | 0 | 0 |
| 0 | 0 | 0 | 0 |
| 0 | 0 | 0 | 0 |
| 0 | 0 | 0 | 0 |
| 0 | 0 | 0 | 0 |
| 0 | 0 | 0 | 0 |
| 0 | 0 | 0 | 0 |
| 0 | 0 | 0 | 0 |
| | | | 7 |
| 8 | 11 | 6 | 0 |
| 4 | 2 | 2 | 0 |
| 3 | 6 | 3 | 0 |
| 1 | 9 | 1 | 0 |
| 2 | 8 | 4 | 0 |
| 5 | 3 | 0 | 0 |
| 0 | 0 | 0 | 3 |
| 4 | 7 | 3 | 3 |
| 0 | 0 | 0 | 0 |
| 1 | 2 | 0 | 1 |
| 0 | 0 | 0 | 0 |
| 0 | 0 | 0 | 0 |
| | | | 7 |

| | | | |
|---|---|---|---|
| 1 | 3 | 4 | 0 |
| 3 | 0 | 3 | 2 |
| 4 | 3 | 4 | 4 |
| 1 | 0 | 7 | 0 |
| 5 | 0 | 2 | 0 |
| 0 | 0 | 0 | 0 |
| 0 | 0 | 0 | 0 |
| 0 | 0 | 0 | 0 |
| 0 | 0 | 0 | 0 |
| 1 | 0 | 0 | 0 |
| 0 | 0 | 0 | 0 |
| 0 | 0 | 0 | 0 |
| | | | 6 |
| 3 | 2 | 3 | 1 |
| 3 | 2 | 3 | 8 |
| 0 | 0 | 5 | 5 |
| 6 | 4 | 1 | 9 |
| 1 | 0 | 3 | 0 |
| 1 | 0 | 1 | 0 |
| 1 | 0 | 0 | 0 |
| 0 | 0 | 0 | 0 |
| 0 | 0 | 0 | 0 |
| 0 | 0 | 0 | 0 |
| 0 | 0 | 0 | 0 |
| 0 | 0 | 0 | 0 |
| | | | 23 |
| 0 | 47 | 0 | 0 |
| 0 | 47 | 0 | 0 |
| 0 | 40 | 0 | 0 |
| 0 | 52 | 0 | 0 |
| 0 | 71 | 0 | 0 |
| 0 | 23 | 0 | 0 |
| 0 | 55 | 0 | 0 |
| 0 | 45 | 0 | 0 |
| 0 | 83 | 0 | 0 |
| 0 | 37 | 0 | 0 |
| 0 | 21 | 0 | 0 |
| 0 | 36 | 0 | 0 |
| | | | 0 |
| 0 | 0 | 19 | 0 |
| 0 | 0 | 4 | 0 |

| | | | |
|---|---|---|---|
| 19 | 4 | 7 | 0 |
| 0 | 0 | 9 | 0 |
| 0 | 0 | 10 | 0 |
| 2 | 0 | 0 | 0 |
| 5 | 0 | 0 | 0 |
| 0 | 0 | 0 | 0 |
| 0 | 0 | 8 | 0 |
| 0 | 0 | 0 | 8 |
| 0 | 0 | 16 | 0 |
| 0 | 0 | 0 | 1 |
| | | | 9 |
| 5 | 1 | 3 | 44 |
| 7 | 5 | 5 | 43 |
| 4 | 4 | 7 | 29 |
| 6 | 2 | 3 | 23 |
| 2 | 1 | 3 | 27 |
| 3 | 6 | 4 | 22 |
| 0 | 0 | 5 | 40 |
| 1 | 0 | 2 | 47 |
| 0 | 1 | 6 | 10 |
| 0 | 0 | 3 | 2 |
| 0 | 0 | 0 | 0 |
| 0 | 0 | 0 | 0 |
| | | | 287 |
| 6 | 2 | 3 | 47 |
| 4 | 0 | 1 | 41 |
| 1 | 1 | 1 | 16 |
| 0 | 0 | 1 | 19 |
| 3 | 1 | 1 | 9 |
| 0 | 0 | 0 | 2 |
| 1 | 0 | 0 | 1 |
| 0 | 0 | 1 | 1 |
| 0 | 0 | 0 | 4 |
| 0 | 0 | 0 | 0 |
| 0 | 0 | 0 | 0 |
| 0 | 0 | 0 | 0 |
| | | | 140 |
| 2 | 8 | 1 | 12 |
| 1 | 3 | 3 | 6 |
| 0 | 0 | 0 | 12 |
| 0 | 0 | 0 | 4 |

| | | | |
|---|---|---|---|
| 0 | 0 | 0 | 0 |
| 0 | 0 | 3 | 8 |
| 0 | 0 | 0 | 1 |
| 0 | 0 | 0 | 1 |
| 0 | 0 | 0 | 0 |
| 0 | 0 | 0 | 0 |
| 0 | 0 | 4 | 5 |
| 0 | 1 | 3 | 12 |
| | | | 61 |
| 3 | 9 | 15 | 1 |
| 10 | 5 | 7 | 0 |
| 6 | 1 | 3 | 0 |
| 2 | 1 | 4 | 0 |
| 4 | 1 | 4 | 0 |
| 1 | 0 | 0 | 0 |
| 6 | 0 | 3 | 0 |
| 2 | 0 | 0 | 0 |
| 0 | 0 | 0 | 0 |
| 0 | 0 | 0 | 0 |
| 3 | 0 | 0 | 0 |
| 0 | 0 | 0 | 0 |
| | | | 1 |
| 4 | 1 | 2 | 3 |
| 3 | 1 | 1 | 0 |
| 3 | 0 | 0 | 0 |
| 1 | 0 | 0 | 0 |
| 0 | 0 | 0 | 0 |
| 0 | 0 | 0 | 0 |
| 1 | 0 | 1 | 0 |
| 2 | 1 | 2 | 0 |
| 0 | 0 | 0 | 0 |
| 0 | 0 | 0 | 0 |
| 0 | 0 | 0 | 0 |
| 0 | 0 | 0 | 0 |
| | | | 3 |
| 5 | 0 | 6 | 1 |
| 1 | 0 | 0 | 0 |
| 1 | 0 | 1 | 0 |
| 0 | 0 | 0 | 1 |
| 0 | 0 | 0 | 0 |
| 0 | 0 | 0 | 0 |

| | | | |
|---|---|---|---|
| 0 | 0 | 0 | 0 |
| 0 | 0 | 0 | 0 |
| 0 | 0 | 0 | 0 |
| 0 | 0 | 0 | 0 |
| 0 | 0 | 0 | 0 |
| 0 | 0 | 0 | 0 |
| | | | 2 |
| 0 | 0 | 0 | 0 |
| 0 | 0 | 0 | 0 |
| 3 | 2 | 2 | 0 |
| 0 | 0 | 3 | 0 |
| 0 | 0 | 5 | 0 |
| 0 | 0 | 1 | 0 |
| 0 | 0 | 3 | 3 |
| 0 | 0 | 0 | 0 |
| 0 | 0 | 0 | 0 |
| 0 | 0 | 0 | 0 |
| 0 | 0 | 0 | 0 |
| 0 | 0 | 0 | 0 |
| | | | 3 |
| 3 | 0 | 3 | 1 |
| 1 | 4 | 6 | 0 |
| 1 | 2 | 2 | 0 |
| 1 | 4 | 4 | 7 |
| 0 | 0 | 1 | 0 |
| 3 | 0 | 0 | 2 |
| 1 | 0 | 0 | 0 |
| 0 | 0 | 0 | 0 |
| 0 | 0 | 0 | 0 |
| 0 | 0 | 0 | 0 |
| 0 | 0 | 0 | 0 |
| 0 | 0 | 0 | 0 |
| | | | 10 |
| 0 | 1 | 2 | 1 |
| 8 | 0 | 0 | 0 |
| 0 | 0 | 1 | 0 |
| 0 | 0 | 0 | 1 |
| 0 | 0 | 0 | 0 |
| 0 | 0 | 0 | 0 |
| 0 | 0 | 0 | 0 |
| 0 | 0 | 0 | 0 |

| | | | |
|---|---|---|---|
| 0 | 0 | 0 | 0 |
| 0 | 0 | 0 | 0 |
| 0 | 0 | 0 | 0 |
| 2 | 0 | 0 | 0 |
| | | | 2 |
| 20 | 3 | 10 | 0 |
| 0 | 12 | 7 | 0 |
| 3 | 2 | 2 | 7 |
| 1 | 6 | 3 | 16 |
| 1 | 0 | 1 | 9 |
| 6 | 2 | 0 | 3 |
| 1 | 2 | 5 | 8 |
| 0 | 0 | 0 | 3 |
| 1 | 0 | 0 | 0 |
| 0 | 0 | 0 | 0 |
| 0 | 0 | 0 | 0 |
| 0 | 0 | 0 | 0 |
| | | | 46 |
| 0 | 0 | 3 | 2 |
| 1 | 9 | 4 | 0 |
| 1 | 0 | 0 | 6 |
| 0 | 0 | 0 | 2 |
| 0 | 0 | 0 | 2 |
| 0 | 0 | 0 | 0 |
| 0 | 0 | 0 | 0 |
| 0 | 0 | 0 | 0 |
| 0 | 0 | 0 | 0 |
| 0 | 0 | 0 | 0 |
| 0 | 0 | 0 | 0 |
| 0 | 0 | 0 | 0 |
| | | | 12 |
| 3 | 0 | 0 | 0 |
| 0 | 0 | 0 | 0 |
| 0 | 0 | 0 | 0 |
| 0 | 0 | 3 | 1 |
| 6 | 9 | 0 | 0 |
| 0 | 0 | 0 | 0 |
| 0 | 0 | 0 | 0 |
| 0 | 0 | 0 | 0 |
| 0 | 0 | 5 | 0 |
| 9 | 3 | 2 | 0 |

| | | | |
|---|---|---|---|
| 0 | 0 | 0 | 0 |
| 0 | 0 | 0 | 0 |
| | | | 1 |
| 6 | 1 | 6 | 9 |
| 0 | 0 | 3 | 12 |
| 4 | 5 | 4 | 17 |
| 3 | 2 | 3 | 19 |
| 4 | 1 | 7 | 11 |
| 1 | 0 | 3 | 11 |
| 2 | 0 | 2 | 8 |
| 0 | 0 | 2 | 5 |
| 0 | 0 | 0 | 1 |
| 0 | 0 | 1 | 0 |
| 0 | 0 | 0 | 7 |
| 0 | 0 | 0 | 3 |
| | | | 103 |
| 5 | 7 | 5 | 9 |
| 2 | 3 | 3 | 10 |
| 5 | 5 | 4 | 11 |
| 1 | 4 | 2 | 3 |
| 3 | 0 | 3 | 4 |
| 1 | 6 | 1 | 3 |
| 2 | 3 | 1 | 5 |
| 0 | 0 | 0 | 2 |
| 0 | 0 | 0 | 1 |
| 0 | 0 | 0 | 0 |
| 0 | 0 | 0 | 0 |
| 1 | 1 | 2 | 5 |
| | | | 53 |
| 5 | 4 | 4 | 10 |
| 1 | 2 | 6 | 14 |
| 2 | 12 | 4 | 4 |
| 5 | 2 | 2 | 6 |
| 10 | 3 | 1 | 5 |
| 0 | 2 | 2 | 5 |
| 2 | 8 | 5 | 3 |
| 1 | 0 | 0 | 9 |
| 2 | 1 | 0 | 1 |
| 2 | 4 | 2 | 16 |
| 2 | 1 | 0 | 3 |
| 0 | 0 | 0 | 0 |

| | | | |
|---|---|---|---|
| | | | 76 |
| 3 | 4 | 11 | 0 |
| 0 | 0 | 1 | 0 |
| 3 | 0 | 7 | 0 |
| 0 | 0 | 0 | 0 |
| 0 | 0 | 0 | 0 |
| 0 | 0 | 3 | 0 |
| 0 | 0 | 0 | 0 |
| 0 | 0 | 0 | 0 |
| 0 | 0 | 0 | 0 |
| 0 | 0 | 0 | 0 |
| 0 | 0 | 0 | 0 |
| 0 | 0 | 0 | 0 |
| | | | 0 |
| 5 | 0 | 8 | 0 |
| 6 | 1 | 5 | 0 |
| 2 | 0 | 2 | 0 |
| 1 | 0 | 0 | 0 |
| 0 | 0 | 1 | 0 |
| 0 | 0 | 0 | 0 |
| 1 | 0 | 0 | 0 |
| 0 | 0 | 0 | 0 |
| 0 | 0 | 0 | 0 |
| 0 | 0 | 0 | 0 |
| 0 | 0 | 0 | 0 |
| 0 | 0 | 0 | 0 |
| | | | 0 |
| 0 | 0 | 0 | 0 |
| 0 | 0 | 0 | 0 |
| 0 | 0 | 0 | 0 |
| 0 | 0 | 0 | 0 |
| 0 | 0 | 0 | 0 |
| 0 | 0 | 0 | 0 |
| 0 | 0 | 0 | 0 |
| 0 | 0 | 0 | 0 |
| 0 | 0 | 0 | 0 |
| 0 | 0 | 0 | 0 |
| 0 | 0 | 0 | 0 |
| 0 | 0 | 0 | 0 |
| | | | 0 |
| 5 | 0 | 4 | 70 |

| | | | |
|---|---|---|---|
| 4 | 3 | 7 | 21 |
| 4 | 5 | 5 | 52 |
| 3 | 0 | 1 | 64 |
| 5 | 0 | 5 | 14 |
| 6 | 3 | 4 | 14 |
| 2 | 3 | 10 | 32 |
| 3 | 3 | 3 | 6 |
| 2 | 2 | 4 | 37 |
| 3 | 5 | 4 | 5 |
| 0 | 0 | 4 | 14 |
| 4 | 0 | 1 | 6 |
| | | | 335 |
| 4 | 2 | 4 | 37 |
| 5 | 1 | 3 | 49 |
| 8 | 1 | 4 | 14 |
| 3 | 4 | 2 | 24 |
| 8 | 6 | 2 | 12 |
| 3 | 2 | 3 | 73 |
| 7 | 2 | 0 | 24 |
| 1 | 2 | 2 | 37 |
| 0 | 0 | 4 | 45 |
| 3 | 0 | 1 | 13 |
| 4 | 0 | 1 | 13 |
| 1 | 0 | 0 | 12 |
| | | | 353 |
| 4 | 1 | 2 | 19 |
| 0 | 0 | 3 | 14 |
| 0 | 1 | 0 | 26 |
| 3 | 2 | 2 | 10 |
| 2 | 2 | 2 | 25 |
| 0 | 1 | 0 | 40 |
| 1 | 0 | 0 | 57 |
| 0 | 0 | 0 | 32 |
| 2 | 1 | 3 | 27 |
| 1 | 4 | 1 | 11 |
| 1 | 1 | 0 | 12 |
| 0 | 2 | 0 | 2 |
| | | | 275 |
| 1 | 4 | 6 | 26 |
| 6 | 1 | 5 | 16 |
| 5 | 3 | 8 | 25 |

| | | | |
|---|---|---|---|
| 4 | 3 | 7 | 13 |
| 1 | 0 | 0 | 5 |
| 4 | 1 | 4 | 40 |
| 1 | 3 | 1 | 24 |
| 3 | 0 | 5 | 63 |
| 7 | 1 | 5 | 35 |
| 3 | 2 | 2 | 23 |
| 1 | 0 | 2 | 51 |
| 4 | 2 | 6 | 33 |
| | | | 354 |
| 3 | 2 | 3 | 43 |
| 3 | 2 | 0 | 16 |
| 6 | 0 | 4 | 53 |
| 5 | 2 | 2 | 27 |
| 3 | 2 | 1 | 16 |
| 4 | 0 | 5 | 33 |
| 4 | 1 | 1 | 24 |
| 2 | 0 | 1 | 35 |
| 1 | 1 | 1 | 39 |
| 2 | 0 | 0 | 37 |
| 4 | 1 | 2 | 17 |
| 2 | 0 | 0 | 12 |
| | | | 352 |
| 8 | 1 | 4 | 32 |
| 1 | 3 | 4 | 31 |
| 3 | 0 | 3 | 24 |
| 2 | 0 | 1 | 25 |
| 2 | 1 | 0 | 14 |
| 2 | 0 | 1 | 13 |
| 1 | 2 | 2 | 13 |
| 0 | 0 | 0 | 11 |
| 2 | 3 | 1 | 21 |
| 1 | 3 | 0 | 13 |
| 2 | 6 | 4 | 13 |
| 2 | 0 | 0 | 9 |
| | | | 219 |
| 3 | 3 | 3 | 1 |
| 5 | 1 | 4 | 2 |
| 1 | 4 | 5 | 8 |
| 2 | 1 | 1 | 6 |
| 1 | 1 | 3 | 8 |

| | | | |
|---|---|---|---|
| 1 | 1 | 0 | 7 |
| 0 | 0 | 0 | 11 |
| 9 | 1 | 2 | 0 |
| 0 | 0 | 0 | 1 |
| 1 | 0 | 0 | 7 |
| 0 | 0 | 0 | 0 |
| 0 | 0 | 0 | 0 |
| | | | 51 |
| 2 | 5 | 1 | 2 |
| 3 | 2 | 2 | 0 |
| 0 | 1 | 0 | 0 |
| 0 | 1 | 0 | 0 |
| 0 | 4 | 3 | 6 |
| 0 | 0 | 0 | 4 |
| 0 | 3 | 0 | 2 |
| 0 | 0 | 0 | 0 |
| 0 | 0 | 0 | 2 |
| 1 | 0 | 2 | 0 |
| 0 | 0 | 1 | 0 |
| 0 | 3 | 0 | 4 |
| | | | 20 |
| 0 | 2 | 2 | 2 |
| 2 | 0 | 3 | 5 |
| 0 | 0 | 0 | 3 |
| 0 | 4 | 3 | 1 |
| 2 | 0 | 1 | 4 |
| 0 | 1 | 3 | 0 |
| 0 | 0 | 0 | 0 |
| 0 | 0 | 0 | 0 |
| 0 | 0 | 0 | 0 |
| 0 | 0 | 0 | 0 |
| 0 | 0 | 0 | 0 |
| 0 | 0 | 0 | 0 |
| | | | 15 |
| 13 | 0 | 8 | 0 |
| 0 | 1 | 3 | 0 |
| 3 | 0 | 6 | 0 |
| 4 | 0 | 0 | 0 |
| 0 | 0 | 2 | 0 |
| 0 | 0 | 0 | 0 |
| 0 | 0 | 0 | 0 |

| | | | |
|---|---|---|---|
| 0 | 0 | 0 | 0 |
| 0 | 1 | 1 | 0 |
| 0 | 1 | 0 | 0 |
| 0 | 0 | 0 | 0 |
| 0 | 0 | 0 | 0 |
| | | | 0 |
| 0 | 0 | 2 | 0 |
| 0 | 1 | 0 | 0 |
| 0 | 0 | 8 | 0 |
| 0 | 1 | 0 | 0 |
| 0 | 0 | 0 | 0 |
| 0 | 0 | 2 | 0 |
| 0 | 0 | 2 | 0 |
| 0 | 0 | 0 | 0 |
| 0 | 0 | 0 | 0 |
| 1 | 0 | 0 | 0 |
| 0 | 0 | 1 | 0 |
| 0 | 0 | 0 | 0 |
| | | | 0 |
| 0 | 1 | 0 | 0 |
| 0 | 0 | 0 | 0 |
| 4 | 0 | 0 | 0 |
| 0 | 0 | 5 | 0 |
| 0 | 0 | 0 | 0 |
| 0 | 0 | 0 | 0 |
| 0 | 0 | 0 | 0 |
| 0 | 0 | 4 | 0 |
| 0 | 0 | 0 | 0 |
| 0 | 0 | 0 | 0 |
| 0 | 0 | 0 | 0 |
| 0 | 0 | 0 | 0 |
| | | | 0 |
| 0 | 0 | 0 | 0 |
| 5 | 1 | 4 | 0 |
| 7 | 2 | 5 | 0 |
| 5 | 0 | 5 | 0 |
| 3 | 0 | 0 | 0 |
| 11 | 0 | 0 | 0 |
| 3 | 0 | 3 | 0 |
| 2 | 0 | 0 | 0 |
| 4 | 0 | 0 | 0 |

| | | | |
|---|---|---|---|
| 7 | 0 | 0 | 0 |
| 8 | 0 | 0 | 0 |
| 0 | 0 | 0 | 0 |
| | | | 0 |
| 2 | 0 | 0 | 0 |
| 0 | 0 | 0 | 0 |
| 3 | 0 | 1 | 0 |
| 6 | 3 | 7 | 0 |
| 0 | 0 | 0 | 0 |
| 0 | 0 | 0 | 0 |
| 0 | 0 | 3 | 0 |
| 0 | 0 | 0 | 0 |
| 0 | 0 | 0 | 0 |
| 0 | 0 | 0 | 0 |
| 0 | 0 | 0 | 0 |
| 0 | 0 | 1 | 0 |
| | | | 0 |
| 0 | 0 | 0 | 0 |
| 0 | 0 | 0 | 2 |
| 0 | 0 | 0 | 0 |
| 0 | 0 | 0 | 0 |
| 0 | 0 | 0 | 0 |
| 0 | 0 | 0 | 0 |
| 0 | 0 | 0 | 0 |
| 0 | 0 | 0 | 0 |
| 0 | 0 | 0 | 0 |
| 0 | 0 | 0 | 0 |
| 0 | 0 | 0 | 0 |
| 0 | 0 | 0 | 0 |
| | | | 2 |
| 8 | 4 | 8 | 4 |
| 0 | 0 | 4 | 3 |
| 0 | 0 | 3 | 5 |
| 4 | 5 | 0 | 6 |
| 0 | 1 | 3 | 17 |
| 1 | 0 | 0 | 5 |
| 1 | 0 | 1 | 2 |
| 0 | 0 | 0 | 2 |
| 0 | 0 | 0 | 2 |
| 0 | 0 | 0 | 1 |
| 0 | 0 | 0 | 5 |

| | | | |
|---|---|---|---|
| 0 | 0 | 0 | 2 |
| | | | 54 |
| 1 | 0 | 0 | 7 |
| 0 | 0 | 2 | 7 |
| 0 | 3 | 0 | 8 |
| 0 | 0 | 0 | 5 |
| 0 | 0 | 0 | 8 |
| 0 | 3 | 0 | 15 |
| 0 | 0 | 0 | 0 |
| 0 | 0 | 0 | 1 |
| 0 | 0 | 0 | 1 |
| 0 | 0 | 0 | 4 |
| 0 | 0 | 0 | 2 |
| 0 | 0 | 0 | 1 |
| | | | 59 |
| 1 | 0 | 0 | 5 |
| 1 | 1 | 1 | 2 |
| 0 | 0 | 0 | 5 |
| 0 | 0 | 0 | 1 |
| 0 | 0 | 0 | 0 |
| 0 | 0 | 0 | 0 |
| 0 | 0 | 0 | 0 |
| 0 | 0 | 0 | 0 |
| 0 | 0 | 0 | 0 |
| 0 | 0 | 0 | 0 |
| 0 | 0 | 0 | 0 |
| 0 | 0 | 0 | 0 |
| | | | 13 |
| 3 | 5 | 12 | 0 |
| 1 | 7 | 4 | 9 |
| 4 | 5 | 4 | 14 |
| 1 | 2 | 1 | 10 |
| 2 | 2 | 2 | 2 |
| 13 | 3 | 2 | 12 |
| 0 | 0 | 1 | 15 |
| 0 | 0 | 0 | 0 |
| 1 | 0 | 0 | 0 |
| 0 | 0 | 0 | 0 |
| 0 | 0 | 0 | 0 |
| 0 | 0 | 0 | 0 |
| | | | 62 |

| | | | |
|---|---|---|---|
| 0 | 1 | 3 | 3 |
| 2 | 3 | 2 | 17 |
| 2 | 1 | 2 | 6 |
| 1 | 2 | 2 | 6 |
| 0 | 0 | 1 | 10 |
| 1 | 4 | 2 | 1 |
| 0 | 12 | 6 | 4 |
| 0 | 9 | 4 | 11 |
| 1 | 3 | 6 | 5 |
| 2 | 3 | 2 | 2 |
| 0 | 0 | 0 | 4 |
| 0 | 0 | 0 | 0 |
| | | | 69 |
| 2 | 0 | 1 | 4 |
| 0 | 4 | 2 | 8 |
| 0 | 3 | 10 | 7 |
| 0 | 0 | 0 | 7 |
| 1 | 3 | 3 | 2 |
| 0 | 4 | 1 | 1 |
| 6 | 2 | 0 | 4 |
| 0 | 0 | 0 | 7 |
| 0 | 0 | 0 | 0 |
| 0 | 0 | 1 | 0 |
| 1 | 1 | 1 | 0 |
| 0 | 0 | 0 | 0 |
| | | | 40 |
| 7 | 8 | 10 | 6 |
| 9 | 1 | 1 | 11 |
| 7 | 3 | 16 | 25 |
| 8 | 1 | 10 | 12 |
| 3 | 0 | 3 | 10 |
| 2 | 1 | 3 | 21 |
| 5 | 2 | 8 | 30 |
| 6 | 1 | 3 | 28 |
| 4 | 0 | 2 | 36 |
| 3 | 1 | 1 | 22 |
| 7 | 2 | 2 | 9 |
| 2 | 2 | 1 | 11 |
| | | | 221 |
| 2 | 1 | 3 | 12 |
| 3 | 1 | 2 | 14 |

| | | | |
|---|---|---|---|
| 2 | 0 | 1 | 34 |
| 1 | 1 | 7 | 22 |
| 1 | 1 | 3 | 7 |
| 5 | 1 | 3 | 21 |
| 0 | 0 | 1 | 14 |
| 3 | 1 | 3 | 10 |
| 1 | 1 | 1 | 4 |
| 1 | 1 | 2 | 10 |
| 4 | 0 | 0 | 32 |
| 1 | 0 | 6 | 20 |
| | | | 200 |
| 2 | 2 | 3 | 10 |
| 4 | 1 | 0 | 22 |
| 3 | 1 | 1 | 14 |
| 0 | 0 | 2 | 9 |
| 2 | 1 | 2 | 6 |
| 3 | 1 | 1 | 7 |
| 0 | 1 | 0 | 6 |
| 0 | 0 | 1 | 2 |
| 3 | 0 | 0 | 17 |
| 1 | 2 | 1 | 2 |
| 0 | 0 | 0 | 5 |
| 2 | 0 | 0 | 14 |
| | | | 114 |
| 12 | 2 | 12 | 3 |
| 4 | 1 | 5 | 5 |
| 5 | 3 | 3 | 12 |
| 0 | 0 | 2 | 3 |
| 5 | 6 | 3 | 1 |
| 5 | 3 | 3 | 14 |
| 2 | 1 | 6 | 19 |
| 1 | 2 | 2 | 5 |
| 2 | 2 | 0 | 35 |
| 5 | 3 | 5 | 16 |
| 6 | 1 | 2 | 11 |
| 0 | 2 | 4 | 8 |
| | | | 132 |
| 3 | 0 | 0 | 11 |
| 0 | 2 | 0 | 22 |
| 5 | 3 | 3 | 29 |
| 5 | 2 | 3 | 33 |

| | | | |
|---|---|---|---|
| 0 | 1 | 1 | 41 |
| 1 | 1 | 4 | 25 |
| 1 | 1 | 1 | 49 |
| 4 | 2 | 1 | 13 |
| 1 | 1 | 0 | 19 |
| 1 | 0 | 2 | 35 |
| 0 | 0 | 0 | 29 |
| 0 | 0 | 0 | 15 |
| | | | 321 |
| 1 | 2 | 1 | 12 |
| 4 | 1 | 1 | 32 |
| 1 | 0 | 0 | 23 |
| 0 | 0 | 0 | 22 |
| 0 | 0 | 2 | 15 |
| 1 | 1 | 0 | 23 |
| 1 | 7 | 2 | 32 |
| 3 | 2 | 0 | 22 |
| 0 | 0 | 6 | 25 |
| 0 | 1 | 2 | 64 |
| 3 | 0 | 0 | 26 |
| 1 | 3 | 2 | 23 |
| | | | 319 |
| 5 | 3 | 2 | 6 |
| 2 | 1 | 5 | 6 |
| 3 | 0 | 7 | 8 |
| 0 | 0 | 3 | 1 |
| 6 | 3 | 8 | 6 |
| 1 | 0 | 1 | 3 |
| 0 | 2 | 0 | 3 |
| 0 | 0 | 0 | 1 |
| 0 | 0 | 0 | 0 |
| 0 | 0 | 0 | 0 |
| 0 | 0 | 0 | 0 |
| 0 | 0 | 0 | 0 |
| | | | 34 |
| 4 | 0 | 5 | 9 |
| 1 | 0 | 7 | 4 |
| 3 | 0 | 0 | 3 |
| 1 | 1 | 0 | 3 |
| 0 | 0 | 0 | 4 |
| 0 | 0 | 0 | 0 |

| | | | |
|---|---|---|---|
| 0 | 0 | 0 | 5 |
| 0 | 0 | 2 | 1 |
| 0 | 0 | 0 | 0 |
| 5 | 0 | 5 | 1 |
| 0 | 0 | 0 | 4 |
| 0 | 0 | 0 | 5 |
| | | | 39 |
| 3 | 1 | 3 | 3 |
| 0 | 0 | 0 | 8 |
| 0 | 0 | 0 | 0 |
| 0 | 0 | 0 | 0 |
| 0 | 0 | 0 | 0 |
| 0 | 0 | 0 | 0 |
| 0 | 0 | 0 | 0 |
| 0 | 0 | 0 | 0 |
| 0 | 0 | 0 | 1 |
| 3 | 0 | 0 | 3 |
| 0 | 0 | 0 | 8 |
| 0 | 0 | 6 | 4 |
| | | | 27 |
| 5 | 12 | 10 | 4 |
| 3 | 6 | 1 | 8 |
| 2 | 3 | 7 | 2 |
| 1 | 4 | 2 | 0 |
| 8 | 8 | 5 | 16 |
| 0 | 3 | 1 | 8 |
| 1 | 0 | 1 | 4 |
| 0 | 0 | 1 | 0 |
| 5 | 4 | 1 | 4 |
| 1 | 0 | 2 | 4 |
| 0 | 0 | 1 | 2 |
| 0 | 0 | 0 | 0 |
| | | | 52 |
| 2 | 6 | 4 | 16 |
| 5 | 6 | 5 | 3 |
| 0 | 1 | 0 | 11 |
| 0 | 0 | 3 | 5 |
| 0 | 0 | 1 | 2 |
| 3 | 4 | 4 | 0 |
| 0 | 0 | 0 | 0 |
| 2 | 0 | 0 | 0 |

| | | | |
|---|---|---|---|
| 0 | 0 | 0 | 0 |
| 1 | 0 | 0 | 2 |
| 0 | 0 | 1 | 5 |
| 0 | 1 | 0 | 1 |
| | | | 45 |
| 1 | 4 | 0 | 2 |
| 0 | 0 | 3 | 5 |
| 3 | 1 | 0 | 1 |
| 0 | 0 | 0 | 3 |
| 0 | 0 | 0 | 0 |
| 0 | 0 | 0 | 0 |
| 0 | 0 | 1 | 1 |
| 1 | 0 | 0 | 8 |
| 3 | 0 | 2 | 5 |
| 0 | 1 | 0 | 5 |
| 0 | 0 | 0 | 1 |
| 0 | 0 | 0 | 0 |
| | | | 31 |
| 0 | 0 | 0 | 0 |
| 0 | 0 | 0 | 0 |
| 0 | 0 | 0 | 0 |
| 2 | 0 | 12 | 0 |
| 3 | 0 | 4 | 0 |
| 0 | 0 | 0 | 0 |
| 0 | 4 | 1 | 0 |
| 0 | 0 | 0 | 0 |
| 0 | 0 | 0 | 0 |
| 0 | 0 | 0 | 0 |
| 0 | 0 | 0 | 0 |
| 0 | 0 | 0 | 0 |
| | | | 0 |
| 6 | 0 | 3 | 0 |
| 3 | 0 | 0 | 0 |
| 10 | 1 | 1 | 0 |
| 0 | 0 | 2 | 0 |
| 0 | 0 | 0 | 0 |
| 0 | 0 | 0 | 0 |
| 0 | 0 | 0 | 0 |
| 0 | 0 | 0 | 0 |
| 0 | 0 | 0 | 0 |
| 0 | 0 | 0 | 0 |

| | | | |
|---|---|---|---|
| 0 | 0 | 0 | 0 |
| 0 | 0 | 0 | 0 |
| | | | 0 |
| 1 | 0 | 1 | 1 |
| 0 | 0 | 1 | 0 |
| 1 | 0 | 0 | 0 |
| 1 | 0 | 1 | 0 |
| 0 | 0 | 1 | 0 |
| 0 | 0 | 0 | 0 |
| 0 | 0 | 0 | 0 |
| 0 | 0 | 0 | 0 |
| 0 | 0 | 2 | 1 |
| 0 | 0 | 0 | 0 |
| 0 | 0 | 0 | 0 |
| 0 | 0 | 0 | 0 |
| | | | 2 |
| 9 | 0 | 5 | 5 |
| 3 | 0 | 5 | 17 |
| 5 | 6 | 7 | 5 |
| 6 | 3 | 5 | 19 |
| 4 | 1 | 3 | 11 |
| 3 | 4 | 2 | 15 |
| 5 | 0 | 5 | 23 |
| 4 | 0 | 0 | 19 |
| 4 | 2 | 4 | 13 |
| 1 | 0 | 1 | 7 |
| 1 | 0 | 1 | 5 |
| 1 | 2 | 1 | 2 |
| | | | 141 |
| 3 | 0 | 3 | 20 |
| 6 | 2 | 2 | 11 |
| 1 | 0 | 1 | 14 |
| 2 | 1 | 1 | 17 |
| 12 | 9 | 1 | 8 |
| 2 | 0 | 4 | 18 |
| 0 | 0 | 0 | 11 |
| 4 | 0 | 9 | 3 |
| 1 | 5 | 0 | 15 |
| 0 | 0 | 0 | 4 |
| 0 | 0 | 0 | 1 |
| 0 | 0 | 0 | 0 |

| | | | |
|---|---|---|---|
| | | | 122 |
| 2 | 0 | 4 | 26 |
| 1 | 5 | 4 | 21 |
| 1 | 4 | 2 | 13 |
| 1 | 1 | 2 | 16 |
| 0 | 5 | 0 | 16 |
| 9 | 3 | 0 | 11 |
| 0 | 0 | 4 | 31 |
| 0 | 0 | 2 | 8 |
| 2 | 3 | 5 | 4 |
| 1 | 2 | 1 | 1 |
| 0 | 0 | 0 | 3 |
| 0 | 0 | 0 | 0 |
| | | | 150 |
| 7 | 0 | 1 | 6 |
| 5 | 2 | 5 | 18 |
| 6 | 6 | 6 | 21 |
| 1 | 2 | 7 | 11 |
| 5 | 4 | 4 | 16 |
| 3 | 6 | 10 | 13 |
| 0 | 0 | 0 | 15 |
| 1 | 0 | 2 | 6 |
| 5 | 1 | 2 | 28 |
| 2 | 0 | 5 | 5 |
| 1 | 0 | 6 | 14 |
| 0 | 0 | 1 | 12 |
| | | | 165 |
| 10 | 0 | 4 | 18 |
| 2 | 0 | 0 | 10 |
| 5 | 4 | 0 | 10 |
| 1 | 0 | 2 | 12 |
| 3 | 0 | 1 | 4 |
| 4 | 0 | 3 | 4 |
| 1 | 0 | 3 | 7 |
| 1 | 0 | 0 | 4 |
| 0 | 1 | 1 | 7 |
| 0 | 0 | 0 | 0 |
| 0 | 0 | 0 | 0 |
| 0 | 0 | 0 | 2 |
| | | | 78 |
| 4 | 1 | 2 | 20 |

| | | | |
|---|---|---|---|
| 2 | 2 | 0 | 16 |
| 1 | 0 | 0 | 8 |
| 2 | 0 | 6 | 15 |
| 1 | 3 | 2 | 8 |
| 0 | 0 | 1 | 11 |
| 0 | 0 | 0 | 20 |
| 1 | 2 | 4 | 6 |
| 0 | 0 | 0 | 2 |
| 0 | 0 | 5 | 1 |
| 0 | 0 | 0 | 1 |
| 0 | 0 | 0 | 0 |
| | | | 108 |
| 4 | 0 | 4 | 6 |
| 2 | 4 | 1 | 13 |
| 5 | 0 | 7 | 12 |
| 2 | 1 | 0 | 5 |
| 3 | 1 | 5 | 8 |
| 4 | 8 | 4 | 8 |
| 2 | 4 | 4 | 8 |
| 2 | 2 | 1 | 6 |
| 3 | 0 | 1 | 3 |
| 0 | 0 | 0 | 1 |
| 1 | 0 | 0 | 2 |
| 2 | 0 | 2 | 1 |
| | | | 73 |
| 5 | 6 | 2 | 23 |
| 3 | 2 | 1 | 22 |
| 4 | 3 | 5 | 14 |
| 3 | 1 | 3 | 33 |
| 6 | 2 | 3 | 22 |
| 1 | 1 | 2 | 36 |
| 2 | 3 | 2 | 31 |
| 1 | 2 | 5 | 12 |
| 6 | 0 | 2 | 21 |
| 2 | 3 | 2 | 34 |
| 5 | 4 | 2 | 15 |
| 2 | 6 | 1 | 7 |
| | | | 270 |
| 2 | 2 | 3 | 27 |
| 1 | 1 | 4 | 22 |
| 2 | 0 | 1 | 40 |

| | | | |
|---|---|---|---|
| 0 | 2 | 7 | 28 |
| 3 | 2 | 9 | 30 |
| 1 | 2 | 1 | 61 |
| 2 | 0 | 3 | 84 |
| 3 | 4 | 2 | 47 |
| 0 | 1 | 2 | 16 |
| 2 | 3 | 1 | 12 |
| 0 | 2 | 6 | 8 |
| 1 | 1 | 1 | 1 |
| | | | 376 |
| 0 | 0 | 0 | 0 |
| 0 | 0 | 0 | 0 |
| 0 | 0 | 0 | 0 |
| 0 | 0 | 0 | 0 |
| 0 | 0 | 0 | 0 |
| 0 | 0 | 0 | 0 |
| 0 | 0 | 0 | 0 |
| 0 | 0 | 0 | 0 |
| 0 | 0 | 0 | 0 |
| 0 | 0 | 0 | 0 |
| 0 | 0 | 0 | 2 |
| 2 | 6 | 6 | 0 |
| | | | 2 |
| 3 | 0 | 5 | 6 |
| 4 | 1 | 0 | 17 |
| 0 | 0 | 3 | 4 |
| 0 | 3 | 2 | 1 |
| 1 | 0 | 3 | 18 |
| 0 | 0 | 0 | 1 |
| 0 | 0 | 0 | 0 |
| 0 | 0 | 0 | 0 |
| 0 | 0 | 0 | 0 |
| 0 | 0 | 0 | 0 |
| 0 | 0 | 0 | 0 |
| 0 | 0 | 0 | 0 |
| | | | 47 |
| 3 | 1 | 0 | 3 |
| 0 | 0 | 0 | 3 |
| 0 | 0 | 2 | 1 |
| 0 | 0 | 0 | 0 |
| 0 | 0 | 0 | 3 |

| | | | |
|---|---|---|---|
| 0 | 0 | 0 | 0 |
| 0 | 0 | 0 | 0 |
| 0 | 0 | 0 | 0 |
| 0 | 0 | 0 | 0 |
| 0 | 0 | 0 | 0 |
| 0 | 0 | 0 | 0 |
| 0 | 0 | 0 | 0 |
| | | | 10 |
| 3 | 1 | 0 | 0 |
| 0 | 0 | 0 | 0 |
| 0 | 0 | 0 | 0 |
| 0 | 0 | 0 | 0 |
| 0 | 0 | 0 | 0 |
| 0 | 0 | 0 | 0 |
| 0 | 0 | 0 | 0 |
| 0 | 0 | 0 | 0 |
| 0 | 0 | 0 | 0 |
| 0 | 0 | 0 | 0 |
| 0 | 0 | 0 | 0 |
| 0 | 0 | 0 | 0 |
| | | | 0 |
| 0 | 0 | 1 | 0 |
| 0 | 0 | 0 | 0 |
| 3 | 3 | 1 | 0 |
| 0 | 0 | 0 | 0 |
| 0 | 0 | 0 | 0 |
| 0 | 0 | 0 | 0 |
| 0 | 0 | 0 | 0 |
| 0 | 0 | 0 | 0 |
| 0 | 0 | 0 | 0 |
| 0 | 0 | 0 | 0 |
| 0 | 0 | 0 | 0 |
| 0 | 0 | 0 | 0 |
| | | | 0 |
| 1 | 0 | 5 | 0 |
| 0 | 2 | 3 | 0 |
| 0 | 0 | 1 | 0 |
| 0 | 0 | 0 | 0 |
| 2 | 0 | 0 | 0 |
| 0 | 0 | 0 | 0 |
| 0 | 0 | 0 | 0 |

| | | | |
|---|---|---|---|
| 0 | 0 | 0 | 0 |
| 0 | 0 | 0 | 0 |
| 0 | 0 | 0 | 0 |
| 0 | 0 | 0 | 0 |
| 0 | 0 | 0 | 0 |
| | | | 0 |
| 5 | 3 | 6 | 1 |
| 0 | 1 | 0 | 2 |
| 4 | 0 | 3 | 3 |
| 2 | 0 | 1 | 1 |
| 0 | 0 | 4 | 8 |
| 0 | 0 | 0 | 3 |
| 0 | 0 | 0 | 3 |
| 0 | 0 | 0 | 3 |
| 0 | 0 | 0 | 0 |
| 0 | 0 | 0 | 0 |
| 0 | 0 | 0 | 0 |
| 0 | 0 | 0 | 1 |
| | | | 25 |
| 3 | 0 | 1 | 9 |
| 3 | 2 | 0 | 15 |
| 2 | 0 | 0 | 11 |
| 0 | 0 | 0 | 3 |
| 0 | 0 | 0 | 4 |
| 0 | 0 | 0 | 2 |
| 0 | 0 | 0 | 1 |
| 0 | 0 | 0 | 0 |
| 0 | 0 | 0 | 0 |
| 0 | 0 | 0 | 0 |
| 0 | 0 | 0 | 1 |
| 0 | 0 | 0 | 3 |
| | | | 49 |
| 4 | 4 | 5 | 0 |
| 2 | 7 | 10 | 0 |
| 4 | 2 | 2 | 5 |
| 0 | 0 | 0 | 3 |
| 2 | 0 | 2 | 0 |
| 2 | 0 | 4 | 4 |
| 0 | 0 | 0 | 6 |
| 0 | 0 | 0 | 0 |
| 0 | 0 | 0 | 0 |

| | | | |
|---|---|---|---|
| 0 | 0 | 0 | 1 |
| 0 | 0 | 0 | 0 |
| 0 | 0 | 0 | 0 |
| | | | 19 |
| 4 | 1 | 3 | 0 |
| 3 | 4 | 4 | 0 |
| 5 | 2 | 11 | 0 |
| 0 | 0 | 1 | 2 |
| 0 | 0 | 0 | 0 |
| 0 | 0 | 0 | 3 |
| 0 | 0 | 0 | 0 |
| 0 | 0 | 0 | 2 |
| 0 | 0 | 0 | 0 |
| 0 | 0 | 0 | 0 |
| 0 | 0 | 0 | 0 |
| 0 | 0 | 0 | 0 |
| | | | 7 |
| 3 | 0 | 4 | 0 |
| 3 | 3 | 2 | 0 |
| 3 | 3 | 1 | 0 |
| 3 | 2 | 4 | 0 |
| 1 | 1 | 1 | 0 |
| 1 | 0 | 0 | 0 |
| 0 | 0 | 0 | 0 |
| 0 | 0 | 1 | 0 |
| 2 | 1 | 0 | 0 |
| 0 | 0 | 0 | 0 |
| 0 | 0 | 0 | 0 |
| 0 | 0 | 0 | 0 |
| | | | 0 |
| 1 | 0 | 0 | 1 |
| 1 | 2 | 0 | 0 |
| 0 | 0 | 0 | 0 |
| 0 | 0 | 0 | 0 |
| 0 | 2 | 1 | 0 |
| 0 | 0 | 5 | 0 |
| 4 | 0 | 0 | 0 |
| 0 | 0 | 1 | 0 |
| 0 | 0 | 0 | 0 |
| 0 | 0 | 0 | 0 |
| 0 | 0 | 0 | 0 |

| | | | |
|---|---|---|---|
| 0 | 0 | 0 | 0 |
| | | | 1 |
| 0 | 0 | 2 | 1 |
| 1 | 1 | 6 | 2 |
| 6 | 6 | 6 | 12 |
| 5 | 6 | 5 | 12 |
| 3 | 1 | 6 | 7 |
| 4 | 4 | 3 | 16 |
| 3 | 4 | 11 | 7 |
| 4 | 2 | 6 | 4 |
| 0 | 0 | 3 | 14 |
| 2 | 0 | 3 | 5 |
| 0 | 0 | 1 | 5 |
| 1 | 1 | 0 | 1 |
| | | | 86 |
| 1 | 4 | 1 | 13 |
| 3 | 2 | 7 | 22 |
| 1 | 0 | 0 | 9 |
| 3 | 1 | 6 | 10 |
| 1 | 2 | 5 | 3 |
| 2 | 1 | 4 | 4 |
| 0 | 0 | 0 | 13 |
| 1 | 0 | 1 | 2 |
| 0 | 0 | 0 | 2 |
| 1 | 1 | 0 | 0 |
| 1 | 1 | 2 | 1 |
| 0 | 0 | 0 | 0 |
| | | | 79 |
| 1 | 0 | 3 | 0 |
| 0 | 1 | 2 | 0 |
| 1 | 2 | 1 | 0 |
| 2 | 0 | 1 | 0 |
| 3 | 0 | 1 | 0 |
| 2 | 0 | 1 | 1 |
| 0 | 0 | 0 | 0 |
| 0 | 0 | 0 | 0 |
| 0 | 0 | 0 | 0 |
| 0 | 0 | 0 | 0 |
| 0 | 0 | 0 | 0 |
| 0 | 0 | 0 | 0 |
| | | | 1 |

| | | | |
|---|---|---|---|
| 7 | 0 | 3 | 0 |
| 3 | 3 | 3 | 0 |
| 2 | 0 | 0 | 1 |
| 4 | 1 | 3 | 1 |
| 0 | 0 | 0 | 0 |
| 0 | 0 | 0 | 0 |
| 0 | 0 | 0 | 0 |
| 0 | 0 | 0 | 0 |
| 0 | 0 | 0 | 0 |
| 0 | 0 | 0 | 0 |
| 0 | 0 | 0 | 0 |
| 0 | 0 | 0 | 0 |
| | | | 2 |
| 3 | 0 | 5 | 1 |
| 2 | 4 | 4 | 2 |
| 5 | 7 | 5 | 1 |
| 0 | 0 | 2 | 0 |
| 1 | 0 | 0 | 1 |
| 0 | 0 | 0 | 0 |
| 0 | 0 | 0 | 1 |
| 0 | 0 | 0 | 0 |
| 1 | 0 | 2 | 3 |
| 0 | 0 | 0 | 0 |
| 0 | 0 | 0 | 0 |
| 0 | 0 | 0 | 0 |
| | | | 9 |
| 1 | 1 | 5 | 3 |
| 1 | 1 | 5 | 9 |
| 3 | 2 | 2 | 8 |
| 1 | 0 | 3 | 7 |
| 1 | 0 | 0 | 0 |
| 1 | 1 | 0 | 5 |
| 1 | 0 | 4 | 3 |
| 0 | 0 | 0 | 0 |
| 0 | 0 | 0 | 0 |
| 0 | 0 | 0 | 0 |
| 0 | 0 | 0 | 0 |
| 0 | 0 | 0 | 0 |
| | | | 35 |
| 12 | 4 | 6 | 6 |
| 5 | 2 | 8 | 1 |

| | | | |
|---|---|---|---|
| 5 | 0 | 6 | 7 |
| 0 | 0 | 1 | 13 |
| 0 | 0 | 0 | 9 |
| 8 | 0 | 2 | 4 |
| 1 | 0 | 0 | 11 |
| 0 | 0 | 0 | 1 |
| 0 | 0 | 0 | 0 |
| 0 | 0 | 0 | 0 |
| 0 | 0 | 0 | 0 |
| 0 | 0 | 0 | 0 |
| | | | 52 |
| 1 | 0 | 1 | 11 |
| 1 | 0 | 3 | 3 |
| 2 | 0 | 0 | 3 |
| 0 | 0 | 0 | 0 |
| 0 | 3 | 6 | 1 |
| 0 | 0 | 0 | 6 |
| 0 | 0 | 0 | 2 |
| 0 | 0 | 0 | 0 |
| 0 | 0 | 0 | 0 |
| 0 | 0 | 0 | 0 |
| 0 | 0 | 0 | 0 |
| 0 | 0 | 0 | 0 |
| | | | 26 |
| 3 | 3 | 0 | 10 |
| 0 | 0 | 0 | 2 |
| 0 | 0 | 0 | 0 |
| 0 | 0 | 0 | 0 |
| 0 | 0 | 0 | 0 |
| 0 | 0 | 0 | 0 |
| 0 | 0 | 0 | 0 |
| 0 | 0 | 0 | 0 |
| 0 | 0 | 0 | 0 |
| 0 | 0 | 0 | 0 |
| 0 | 0 | 0 | 1 |
| 0 | 0 | 0 | 0 |
| | | | 13 |
| 10 | 1 | 1 | 1 |
| 1 | 0 | 4 | 1 |
| 5 | 3 | 2 | 0 |
| 1 | 3 | 2 | 0 |

| | | | |
|---|---|---|---|
| 0 | 2 | 4 | 3 |
| 4 | 1 | 3 | 6 |
| 1 | 0 | 1 | 2 |
| 1 | 0 | 0 | 2 |
| 0 | 0 | 0 | 4 |
| 2 | 0 | 0 | 1 |
| 0 | 0 | 0 | 0 |
| 0 | 0 | 0 | 0 |
| | | | 20 |
| 4 | 1 | 1 | 3 |
| 2 | 0 | 0 | 0 |
| 1 | 0 | 0 | 4 |
| 2 | 0 | 2 | 13 |
| 2 | 3 | 3 | 9 |
| 0 | 0 | 0 | 2 |
| 0 | 0 | 0 | 2 |
| 0 | 0 | 0 | 6 |
| 0 | 0 | 0 | 3 |
| 0 | 0 | 2 | 2 |
| 0 | 0 | 0 | 4 |
| 0 | 0 | 0 | 6 |
| | | | 54 |
| 1 | 0 | 0 | 1 |
| 0 | 0 | 1 | 7 |
| 0 | 0 | 5 | 2 |
| 7 | 1 | 2 | 1 |
| 0 | 0 | 0 | 2 |
| 0 | 0 | 0 | 0 |
| 1 | 0 | 1 | 1 |
| 0 | 0 | 0 | 0 |
| 1 | 0 | 0 | 2 |
| 0 | 0 | 0 | 0 |
| 0 | 0 | 0 | 0 |
| 1 | 0 | 0 | 6 |
| | | | 22 |
| 8 | 4 | 7 | 0 |
| 3 | 1 | 8 | 0 |
| 2 | 2 | 5 | 1 |
| 1 | 2 | 8 | 7 |
| 0 | 5 | 1 | 2 |
| 0 | 0 | 3 | 2 |

| | | | |
|---|---|---|---|
| 2 | 0 | 0 | 6 |
| 0 | 0 | 3 | 7 |
| 0 | 0 | 0 | 4 |
| 0 | 0 | 2 | 0 |
| 0 | 0 | 0 | 0 |
| 0 | 0 | 0 | 0 |
| | | | 29 |
| 6 | 1 | 5 | 19 |
| 1 | 4 | 9 | 33 |
| 4 | 0 | 9 | 18 |
| 2 | 2 | 4 | 35 |
| 2 | 0 | 9 | 19 |
| 2 | 2 | 6 | 22 |
| 1 | 2 | 5 | 16 |
| 0 | 0 | 1 | 0 |
| 1 | 0 | 1 | 9 |
| 0 | 0 | 1 | 1 |
| 0 | 0 | 0 | 4 |
| 4 | 2 | 4 | 3 |
| | | | 179 |
| 6 | 2 | 3 | 6 |
| 0 | 0 | 3 | 2 |
| 0 | 0 | 1 | 3 |
| 1 | 3 | 3 | 1 |
| 0 | 0 | 1 | 1 |
| 0 | 0 | 0 | 0 |
| 0 | 0 | 2 | 0 |
| 0 | 0 | 0 | 1 |
| 0 | 0 | 0 | 0 |
| 0 | 0 | 0 | 0 |
| 0 | 0 | 0 | 0 |
| 0 | 0 | 0 | 0 |
| | | | 14 |
| 4 | 6 | 1 | 1 |
| 9 | 3 | 7 | 0 |
| 1 | 2 | 7 | 0 |
| 18 | 9 | 2 | 4 |
| 4 | 1 | 3 | 2 |
| 5 | 2 | 1 | 4 |
| 0 | 0 | 0 | 0 |
| 0 | 2 | 2 | 0 |

| | | | |
|---|---|---|---|
| 0 | 0 | 0 | 1 |
| 0 | 0 | 0 | 2 |
| 0 | 0 | 0 | 0 |
| 0 | 0 | 0 | 0 |
| | | | 14 |
| 5 | 6 | 3 | 0 |
| 3 | 7 | 4 | 0 |
| 5 | 5 | 4 | 0 |
| 3 | 1 | 6 | 0 |
| 1 | 1 | 0 | 5 |
| 0 | 1 | 0 | 0 |
| 0 | 0 | 0 | 0 |
| 0 | 0 | 0 | 0 |
| 0 | 0 | 0 | 0 |
| 0 | 0 | 0 | 0 |
| 0 | 0 | 0 | 0 |
| 0 | 0 | 0 | 0 |
| | | | 5 |
| 4 | 4 | 4 | 1 |
| 6 | 2 | 2 | 8 |
| 2 | 3 | 0 | 8 |
| 1 | 0 | 1 | 5 |
| 1 | 0 | 0 | 3 |
| 0 | 0 | 0 | 0 |
| 0 | 3 | 0 | 0 |
| 0 | 0 | 0 | 0 |
| 0 | 0 | 0 | 0 |
| 0 | 0 | 0 | 0 |
| 0 | 0 | 0 | 0 |
| 0 | 0 | 0 | 0 |
| | | | 25 |
| 13 | 0 | 2 | 0 |
| 0 | 0 | 0 | 0 |
| 0 | 4 | 15 | 0 |
| 0 | 0 | 0 | 0 |
| 4 | 3 | 3 | 1 |
| 0 | 0 | 0 | 3 |
| 3 | 0 | 0 | 1 |
| 0 | 1 | 1 | 0 |
| 0 | 0 | 0 | 0 |
| 0 | 0 | 0 | 0 |

| | | | |
|---|---|---|---|
| 0 | 0 | 0 | 0 |
| 0 | 0 | 0 | 0 |
| | | | 5 |
| 0 | 0 | 0 | 0 |
| 0 | 2 | 2 | 0 |
| 0 | 0 | 1 | 0 |
| 0 | 0 | 0 | 0 |
| 2 | 0 | 1 | 0 |
| 0 | 0 | 1 | 0 |
| 4 | 0 | 2 | 1 |
| 0 | 0 | 0 | 0 |
| 0 | 0 | 0 | 2 |
| 0 | 0 | 0 | 0 |
| 0 | 0 | 0 | 0 |
| 0 | 0 | 0 | 0 |
| | | | 3 |
| 2 | 0 | 3 | 4 |
| 0 | 1 | 1 | 0 |
| 0 | 0 | 0 | 6 |
| 2 | 0 | 1 | 3 |
| 3 | 0 | 0 | 0 |
| 0 | 1 | 2 | 1 |
| 0 | 0 | 0 | 5 |
| 0 | 0 | 0 | 0 |
| 0 | 0 | 0 | 0 |
| 0 | 0 | 3 | 1 |
| 0 | 0 | 0 | 0 |
| 0 | 0 | 0 | 0 |
| | | | 20 |
| 8 | 0 | 4 | 0 |
| 2 | 4 | 5 | 0 |
| 5 | 0 | 0 | 11 |
| 6 | 7 | 4 | 22 |
| 2 | 2 | 6 | 23 |
| 1 | 4 | 10 | 19 |
| 3 | 1 | 2 | 25 |
| 3 | 1 | 1 | 16 |
| 0 | 0 | 3 | 6 |
| 1 | 0 | 0 | 19 |
| 2 | 0 | 0 | 16 |
| 0 | 6 | 0 | 7 |

| | | | |
|---|---|---|---|
| | | | 164 |
| 5 | 7 | 1 | 0 |
| 1 | 0 | 4 | 4 |
| 0 | 1 | 2 | 1 |
| 0 | 1 | 4 | 7 |
| 0 | 0 | 0 | 7 |
| 3 | 0 | 2 | 0 |
| 0 | 0 | 0 | 1 |
| 1 | 2 | 1 | 3 |
| 1 | 0 | 2 | 5 |
| 1 | 0 | 2 | 1 |
| 0 | 0 | 0 | 0 |
| 0 | 0 | 0 | 0 |
| | | | 29 |
| 2 | 0 | 1 | 0 |
| 0 | 0 | 0 | 0 |
| 0 | 0 | 0 | 4 |
| 0 | 0 | 0 | 0 |
| 1 | 0 | 1 | 3 |
| 1 | 0 | 0 | 0 |
| 0 | 0 | 0 | 0 |
| 0 | 0 | 0 | 0 |
| 0 | 0 | 0 | 0 |
| 0 | 0 | 0 | 0 |
| 0 | 0 | 0 | 0 |
| 0 | 0 | 0 | 0 |
| | | | 7 |
| 2 | 0 | 0 | 0 |
| 0 | 0 | 0 | 0 |
| 0 | 0 | 1 | 0 |
| 3 | 1 | 4 | 0 |
| 0 | 0 | 0 | 0 |
| 1 | 0 | 0 | 0 |
| 1 | 0 | 0 | 0 |
| 0 | 0 | 0 | 0 |
| 1 | 0 | 0 | 0 |
| 2 | 0 | 2 | 0 |
| 0 | 0 | 0 | 0 |
| 0 | 0 | 0 | 0 |
| | | | 0 |
| 3 | 0 | 1 | 0 |

| | | | |
|---|---|---|---|
| 0 | 1 | 2 | 0 |
| 1 | 0 | 0 | 0 |
| 0 | 0 | 0 | 2 |
| 1 | 0 | 0 | 0 |
| 2 | 0 | 0 | 0 |
| 0 | 0 | 0 | 0 |
| 0 | 0 | 0 | 0 |
| 0 | 0 | 0 | 0 |
| 0 | 0 | 0 | 0 |
| 0 | 0 | 0 | 0 |
| 0 | 0 | 0 | 0 |
| | | | 2 |
| 1 | 0 | 1 | 1 |
| 0 | 0 | 0 | 0 |
| 0 | 0 | 0 | 0 |
| 0 | 0 | 0 | 0 |
| 0 | 0 | 0 | 0 |
| 1 | 0 | 1 | 0 |
| 0 | 0 | 0 | 1 |
| 0 | 0 | 0 | 0 |
| 0 | 0 | 0 | 0 |
| 0 | 0 | 0 | 0 |
| 0 | 0 | 0 | 0 |
| 0 | 0 | 0 | 0 |
| | | | 2 |
| 0 | 0 | 0 | 0 |
| 0 | 0 | 0 | 0 |
| 0 | 0 | 0 | 0 |
| 0 | 0 | 0 | 0 |
| 0 | 0 | 0 | 0 |
| 6 | 4 | 3 | 0 |
| 6 | 0 | 0 | 0 |
| 8 | 0 | 0 | 0 |
| 3 | 0 | 0 | 0 |
| 7 | 0 | 0 | 0 |
| 0 | 0 | 9 | 0 |
| 0 | 0 | 0 | 0 |
| | | | 0 |
| 5 | 2 | 2 | 0 |
| 0 | 0 | 1 | 0 |
| 0 | 1 | 4 | 0 |

| | | | |
|---|---|---|---|
| 4 | 1 | 2 | 0 |
| 0 | 1 | 0 | 0 |
| 0 | 0 | 3 | 0 |
| 2 | 2 | 4 | 0 |
| 0 | 0 | 4 | 0 |
| 0 | 0 | 2 | 0 |
| 0 | 0 | 0 | 0 |
| 0 | 0 | 1 | 0 |
| 0 | 0 | 0 | 0 |
| | | | 0 |
| 6 | 0 | 1 | 0 |
| 1 | 1 | 2 | 0 |
| 0 | 0 | 1 | 0 |
| 1 | 0 | 0 | 0 |
| 1 | 0 | 1 | 0 |
| 0 | 0 | 0 | 1 |
| 1 | 0 | 0 | 0 |
| 0 | 0 | 0 | 0 |
| 2 | 0 | 0 | 0 |
| 0 | 0 | 0 | 0 |
| 0 | 0 | 0 | 0 |
| 0 | 0 | 0 | 0 |
| | | | 1 |
| 1 | 0 | 7 | 0 |
| 0 | 0 | 0 | 0 |
| 1 | 1 | 0 | 0 |
| 3 | 2 | 1 | 0 |
| 1 | 0 | 2 | 0 |
| 1 | 1 | 6 | 0 |
| 0 | 0 | 0 | 0 |
| 0 | 0 | 0 | 0 |
| 0 | 2 | 2 | 0 |
| 0 | 0 | 0 | 0 |
| 0 | 0 | 0 | 0 |
| 0 | 0 | 0 | 0 |
| | | | 0 |
| 2 | 0 | 3 | 0 |
| 0 | 0 | 0 | 3 |
| 0 | 0 | 4 | 2 |
| 0 | 1 | 2 | 2 |
| 1 | 0 | 0 | 5 |

| | | | |
|---|---|---|---|
| 0 | 0 | 0 | 0 |
| 1 | 0 | 0 | 0 |
| 0 | 0 | 0 | 0 |
| 0 | 0 | 0 | 0 |
| 1 | 0 | 0 | 0 |
| 0 | 0 | 0 | 0 |
| 0 | 0 | 0 | 0 |
| | | | 12 |
| 1 | 0 | 0 | 1 |
| 1 | 1 | 0 | 2 |
| 2 | 0 | 0 | 1 |
| 1 | 0 | 1 | 1 |
| 0 | 0 | 0 | 0 |
| 0 | 0 | 0 | 0 |
| 0 | 0 | 0 | 0 |
| 0 | 0 | 0 | 0 |
| 0 | 0 | 0 | 0 |
| 0 | 0 | 0 | 0 |
| 0 | 0 | 0 | 0 |
| 0 | 0 | 0 | 0 |
| | | | 5 |
| 1 | 1 | 2 | 0 |
| 1 | 2 | 6 | 1 |
| 3 | 2 | 3 | 5 |
| 1 | 4 | 1 | 10 |
| 0 | 0 | 1 | 8 |
| 3 | 2 | 3 | 0 |
| 0 | 0 | 0 | 8 |
| 0 | 0 | 1 | 2 |
| 1 | 0 | 0 | 2 |
| 0 | 0 | 0 | 0 |
| 0 | 0 | 0 | 2 |
| 0 | 0 | 0 | 1 |
| | | | 39 |
| 1 | 2 | 2 | 14 |
| 0 | 0 | 6 | 13 |
| 4 | 1 | 2 | 4 |
| 0 | 1 | 1 | 8 |
| 3 | 1 | 3 | 5 |
| 0 | 0 | 0 | 4 |
| 0 | 0 | 0 | 0 |

| | | | |
|---|---|---|---|
| 3 | 0 | 0 | 1 |
| 0 | 0 | 0 | 0 |
| 0 | 0 | 0 | 0 |
| 0 | 0 | 0 | 0 |
| 0 | 0 | 0 | 3 |
| | | | 52 |
| 6 | 0 | 1 | 11 |
| 0 | 0 | 2 | 9 |
| 0 | 0 | 0 | 2 |
| 0 | 0 | 0 | 5 |
| 1 | 0 | 0 | 3 |
| 0 | 0 | 0 | 0 |
| 0 | 0 | 0 | 6 |
| 1 | 0 | 0 | 7 |
| 1 | 3 | 4 | 2 |
| 0 | 0 | 0 | 0 |
| 0 | 0 | 0 | 0 |
| 0 | 0 | 0 | 0 |
| | | | 45 |
| 12 | 8 | 5 | 20 |
| 0 | 0 | 2 | 11 |
| 1 | 0 | 0 | 20 |
| 1 | 0 | 0 | 15 |
| 0 | 0 | 2 | 7 |
| 0 | 0 | 0 | 11 |
| 0 | 0 | 1 | 7 |
| 0 | 0 | 0 | 4 |
| 0 | 1 | 0 | 16 |
| 0 | 0 | 0 | 0 |
| 7 | 0 | 0 | 13 |
| 0 | 0 | 0 | 10 |
| | | | 134 |
| 1 | 0 | 0 | 23 |
| 4 | 0 | 4 | 16 |
| 10 | 2 | 3 | 12 |
| 2 | 1 | 0 | 4 |
| 0 | 0 | 0 | 6 |
| 0 | 0 | 0 | 20 |
| 2 | 0 | 1 | 38 |
| 0 | 0 | 0 | 31 |
| 0 | 0 | 0 | 34 |

| | | | |
|---|---|---|---|
| 0 | 0 | 0 | 6 |
| 0 | 0 | 0 | 9 |
| 0 | 0 | 0 | 9 |
| | | | 208 |
| 1 | 2 | 0 | 2 |
| 0 | 0 | 0 | 3 |
| 0 | 0 | 0 | 0 |
| 0 | 0 | 0 | 3 |
| 0 | 0 | 0 | 0 |
| 0 | 0 | 0 | 0 |
| 0 | 0 | 0 | 0 |
| 0 | 0 | 0 | 0 |
| 0 | 0 | 0 | 0 |
| 0 | 0 | 0 | 0 |
| 0 | 0 | 0 | 0 |
| 0 | 0 | 0 | 0 |
| | | | 8 |
| 4 | 0 | 1 | 1 |
| 1 | 2 | 7 | 0 |
| 2 | 0 | 3 | 3 |
| 3 | 7 | 3 | 0 |
| 8 | 1 | 1 | 2 |
| 0 | 0 | 1 | 2 |
| 0 | 0 | 0 | 0 |
| 0 | 0 | 0 | 0 |
| 4 | 1 | 1 | 0 |
| 0 | 0 | 0 | 0 |
| 0 | 0 | 0 | 0 |
| 0 | 0 | 0 | 0 |
| | | | 8 |
| 4 | 2 | 4 | 1 |
| 4 | 5 | 4 | 0 |
| 5 | 0 | 4 | 1 |
| 1 | 0 | 0 | 0 |
| 5 | 1 | 4 | 4 |
| 5 | 2 | 0 | 0 |
| 0 | 0 | 0 | 5 |
| 3 | 0 | 2 | 19 |
| 1 | 1 | 1 | 2 |
| 0 | 0 | 0 | 0 |
| 0 | 0 | 0 | 0 |

| | | | |
|---|---|---|---|
| 1 | 0 | 0 | 0 |
| | | | 32 |
| 1 | 0 | 2 | 1 |
| 0 | 0 | 0 | 0 |
| 0 | 0 | 0 | 0 |
| 2 | 2 | 2 | 0 |
| 5 | 3 | 2 | 0 |
| 1 | 1 | 1 | 0 |
| 7 | 0 | 0 | 0 |
| 0 | 0 | 0 | 0 |
| 0 | 0 | 0 | 0 |
| 0 | 0 | 0 | 0 |
| 0 | 0 | 0 | 0 |
| 2 | 0 | 0 | 0 |
| | | | 1 |
| 11 | 5 | 7 | 20 |
| 1 | 3 | 3 | 24 |
| 0 | 1 | 3 | 12 |
| 4 | 3 | 1 | 9 |
| 2 | 1 | 4 | 29 |
| 4 | 0 | 2 | 15 |
| 3 | 1 | 2 | 13 |
| 1 | 1 | 1 | 27 |
| 1 | 1 | 4 | 38 |
| 1 | 8 | 4 | 12 |
| 1 | 2 | 1 | 22 |
| 3 | 1 | 4 | 11 |
| | | | 232 |
| 6 | 1 | 7 | 25 |
| 3 | 3 | 0 | 29 |
| 1 | 0 | 3 | 20 |
| 3 | 1 | 7 | 21 |
| 1 | 0 | 1 | 20 |
| 0 | 0 | 0 | 59 |
| 4 | 1 | 2 | 21 |
| 1 | 0 | 0 | 24 |
| 3 | 1 | 2 | 23 |
| 0 | 1 | 3 | 31 |
| 0 | 1 | 2 | 7 |
| 5 | 0 | 4 | 18 |
| | | | 298 |

| | | | |
|---|---|---|---|
| 1 | 1 | 1 | 25 |
| 0 | 0 | 2 | 31 |
| 2 | 1 | 1 | 26 |
| 1 | 0 | 0 | 24 |
| 3 | 2 | 3 | 11 |
| 0 | 0 | 1 | 7 |
| 0 | 1 | 1 | 18 |
| 0 | 0 | 0 | 29 |
| 0 | 0 | 0 | 25 |
| 3 | 1 | 0 | 18 |
| 0 | 0 | 2 | 34 |
| 1 | 2 | 0 | 26 |
| | | | 274 |
| 6 | 4 | 3 | 0 |
| 2 | 2 | 3 | 10 |
| 2 | 4 | 2 | 8 |
| 3 | 6 | 9 | 11 |
| 3 | 11 | 4 | 6 |
| 3 | 1 | 1 | 2 |
| 6 | 4 | 2 | 5 |
| 4 | 1 | 5 | 4 |
| 4 | 1 | 0 | 3 |
| 4 | 5 | 4 | 4 |
| 0 | 1 | 2 | 12 |
| 5 | 1 | 0 | 2 |
| | | | 67 |
| 6 | 5 | 3 | 10 |
| 4 | 5 | 6 | 3 |
| 3 | 3 | 1 | 6 |
| 9 | 4 | 2 | 13 |
| 3 | 3 | 2 | 12 |
| 1 | 3 | 3 | 3 |
| 3 | 0 | 0 | 3 |
| 0 | 0 | 0 | 2 |
| 1 | 0 | 0 | 1 |
| 0 | 2 | 1 | 0 |
| 0 | 0 | 0 | 0 |
| 0 | 0 | 0 | 0 |
| | | | 53 |
| 0 | 1 | 1 | 4 |
| 5 | 0 | 1 | 3 |

| | | | |
|---|---|---|---|
| 0 | 0 | 0 | 10 |
| 2 | 0 | 1 | 5 |
| 0 | 0 | 0 | 2 |
| 0 | 0 | 0 | 0 |
| 0 | 0 | 0 | 0 |
| 0 | 0 | 0 | 0 |
| 0 | 0 | 0 | 0 |
| 0 | 0 | 0 | 0 |
| 0 | 0 | 0 | 0 |
| 0 | 0 | 0 | 0 |
| | | | 24 |
| | | | 58 |
| | | | 102 |
| | | | 10 |
| | | | 23 |
